# High-Throughput Functional Annotation of Natural Products by Integrated Activity Profiling

**DOI:** 10.1101/748129

**Authors:** Suzie K. Hight, Trevor N. Clark, Kenji L. Kurita, Elizabeth A. McMillan, Walter Bray, Anam F. Shaikh, F. P. Jake Haeckl, Fausto Carnevale-Neto, Scott La, Akshar Lohith, Rachel M. Vaden, Jeon Lee, Shuguang Wei, R. Scott Lokey, Michael A. White, Roger G. Linington, John B. MacMillan

## Abstract

Determining mechanism of action (MOA) is one of the biggest challenges in natural products discovery. Here, we report a comprehensive platform that uses Similarity Network Fusion (SNF) to improve MOA predictions by integrating data from the cytological profiling high-content imaging platform and the gene expression platform FUSION, and pairs these data with untargeted metabolomics analysis for *de novo* bioactive compound discovery. The predictive value of the integrative approach was assessed using a library of target-annotated small molecules as benchmarks. Using Kolmogorov–Smirnov (KS) tests to compare in-class to out-of-class similarity, we found that SNF retains the ability to identify significant in-class similarity across a diverse set of target classes, and could also find target classes that were not detectable in either platform alone. This confirmed that integration of expression-based and image-based phenotypes can accurately report on MOA. Furthermore, we integrated untargeted metabolomics of complex natural product fractions with the SNF network to map biological signatures to specific metabolites. Three examples are presented where SNF coupled with metabolomics was used to directly functionally characterize natural products and accelerate identification of bioactive metabolites, including the discovery of the novel azoxy-containing biaryl compounds parkamycins A and B. Our results support SNF integration of multiple phenotypic screening approaches along with untargeted metabolomics as a powerful approach for advancing natural products drug discovery.

**Significance statement:** New data-driven methods to aid in the discovery and biological characterization of natural products are necessary to advance the field. Assigning the mechanism of action (MOA) to novel bioactive compounds is an essential step in drug discovery and a major challenge in chemical biology. Despite technological advances in isolation, synthesis and screening strategies that make many bioactive substances readily available, in most cases their biological targets remain unknown. Additionally, a major bottleneck in natural products discovery efforts is de-replication of the large number of known compounds that predominate in crude extracts and fraction libraries. Advances in metabolomics has provided a better understanding of the constituents present in these libraries, but is not sufficient in itself to drive the discovery of novel biologically active metabolites. Here we describe an unbiased, data-driven strategy which integrates phenotypic screening with metabolomics into a single platform that provides rapid identification and functional annotation of natural products. This approach can be applied to any cohort of uncharacterized chemicals and represents a strategy that could significantly accelerate the process of drug discovery.

## Introduction & Background

Assigning the mechanism of action (MOA) to botanicals, natural products and synthetic chemicals is an essential step in drug discovery and remains a major challenge in chemical biology. Despite the technological advances in isolation, synthesis and screening strategies that make many bioactive substances available, in most cases their biological targets remain unknown (1, 2). This challenge is exacerbated when taking a systems-level approach to gain mechanistic information about entire collections of molecules, such as natural product libraries from microorganisms, marine invertebrates or plants. Due to the difficulties associated with isolation of natural products, the majority of academic and industrial natural product collections are complex fractions or crude extracts. Thus, in addition to the biological complexity of assigning mechanism of action, there is the added complication that typical one-compound-one-well assay formats are not applicable (3).

There has been a concerted effort in academia as well as biopharma to return to phenotypic screening approaches in drug discovery efforts (4, 5). This paradigm shift has come with the development of novel, information-rich phenotypic approaches that have the ability to provide an unprecedented level of mechanistic understanding. These methods take advantage of gene expression profiling (6–8), high content imaging (9–11), yeast chemical genetics (12), proteomics (13, 14), and others (5, 15). While these platforms are valuable individually, each one is subject to the limitations of the reporter system used and thus can still fail to detect real biological associations. Here we test the hypothesis that using computational tools to integrate screening results from orthogonal screening platforms will allow for simultaneous leverage of divergent phenotypic coverage to inform MOA predictions. In this study, we integrate gene expression-based (Functional Signature Ontology; FUSION) and high content imaging-based (Cytological Profiling; CP) screening platforms using Similarity Network Fusion (SNF), and use this fused network to annotate high-resolution mass spectrometric profiling of a library of complex natural product fractions. The result is a novel framework for the functional annotation of natural products that demonstrates the power of leveraging multiple data types.

Our interest in the functional characterization of microbe-derived natural products and botanicals led our groups to independently develop phenotypic screening strategies to evaluate natural product fraction libraries from marine bacteria. One platform, termed Functional Signature Ontology (FUSION), utilizes perturbation-induced gene expression signatures coupled with pattern-matching tools to produce verifiable guilt-by-association MOA hypotheses (8). The initial platform used the colon cancer cell line HCT116 and six reporter genes to generate expression signatures of “known” perturbagens such as siRNAs or miRNAs, for comparison to “unknown” perturbagens such as natural product fractions and pure natural products. This method has been used to characterize a series of microbially-derived molecules with unique mechanisms of action (8, 16–19). In this study, we have adapted the FUSION approach to a non-small cell lung cancer context using the cell line NCI-H23 and a new set of 14 reporter genes that form the basis for pattern-matching between known and unknown perturbagens. A limitation of this approach, as well as other gene expression-based approaches such as the Connectivity Map (LINCS Consortium) (7), is that the sensitivity and specificity of the signature a bioactive molecule can produce is dependent on the biological context of the assay. For example, interrogation of signaling pathways which are not intact in the chosen biological model may result in either a null or off-target pharmacological effect. Consideration of these factors is important when utilizing these types of data to inform MOA hypotheses.

A biologically orthogonal platform, cytological profiling (CP), utilizes high-content image analysis of perturbation-treated cells stained with a panel of fluorescent probes to extract sets of cytological features that are then used with pattern-matching tools to predict MOA (20). The CP platform utilizes unsynchronized HeLa cells, which, after treatment with perturbagen are fixed and stained with probes for S-phase progression, total DNA, phospho-histone H3, tubulin and actin (3). A total of 251 unique cytological features are then extracted for each perturbagen from automated fluorescence microscopy images. Clustering compounds by their CP fingerprints has revealed both well-established associations among compounds with the same target or MOA, as well as novel or unexpected associations and unique phenotypes of natural products (3, 11). As with FUSION, while CP and related high-content imaging platforms have demonstrated success for evaluating mechanism of action, they are subject to limited resolution of bioactive compounds with broad cellular effects, or limited sensitivity to bioactive compounds with that engage morphologically silent mechanisms (11).

Both of these technologies have been productively utilized by our respective groups to evaluate the mechanism of action of complex natural product mixtures and pure natural products (3, 8, 11, 16–19, 21). However the inherent limitations of these platforms can be especially problematic when exploring large, uncharacterized libraries whose active metabolites may span a wide and divergent range of biological activities. Tools which can be applied to such systematic explorations would greatly accelerate natural product characterization and drug discovery. We hypothesized that a bioinformatic approach to integrate the two platforms could expand the biological space covered while retaining the information from both platforms. A challenge with integration of diverse data types is how to handle disparate numbers of features and data scales in individual datasets. To solve this problem, we adopted similarity network fusion (SNF) (22), which overcomes this challenge by constructing similarity networks individually for each available data type and then fusing these into a single network based on shared similarity across both datasets. SNF has been used efficiently in a range of applications, including but not limited to integration of image-based profiling data to capture cell-to-cell heterogeneity (23), MOA inference by combining drug sensitivity with structural data (24), and integration of multi-omic molecular profiles for identification of subtypes within cancer, COVID-19 and other diseases (22, 25–33).

Unlike synthetic screening libraries, which are arrayed in a one-compound-one-well format, natural products screening libraries are typically prepared as complex mixtures. To relate phenotypes to specific components in these mixtures, we required both a detailed description of the chemical constitution of each fraction, and informatics tools to define the associations between constituents and phenotypes. Accurate characterization of chemical composition for natural products libraries remains an unsolved challenge in the field. This objective is particularly difficult because natural products can be present at a large range of concentrations, with variable ionization efficiencies and a wide range of polarities and molecular weights.

Current mass spectrometry-based methods often yield peak lists with very high false discovery rates, making it difficult to identify biologically relevant features from these large results files (34). To address this, we developed bespoke acquisition and data processing methods designed to describe the chemical constitution of the natural products fraction library while removing interference signals caused by instrument noise and systemic contaminants from the sample processing workflow. These methods included appropriate replicates, blanks, and sample preparation workflow, employing an ion-mobility spectroscopy-enabled ESI-qTOF. The inclusion of ion mobility spectroscopy improved separation of co-eluting components and improved the binning of analytes between samples by providing a third axis of separation (collision cross-sectional area) over traditional methods (35). Using a modified version of our Compound Activity Mapping platform (21), we then defined activity scores for all analytes based on SNF clustering results and developed a custom data visualization platform to directly relate analytes to specific biological phenotypes.

Here we describe the integration of Cytological Profiling and FUSION screening data to create a platform for functional annotation that can simultaneously leverage the information from both datatypes to guide *de novo* mode of action prediction. By combining this with next-generation metabolomics analysis of natural products libraries, we have created a unique and powerful framework for natural product biological characterization (Figure 1A). The value of this platform is illustrated by three examples that demonstrate how integration of these discrete data types can help drive discovery of novel chemistry and characterization of biological activity.

**Figure 1.**
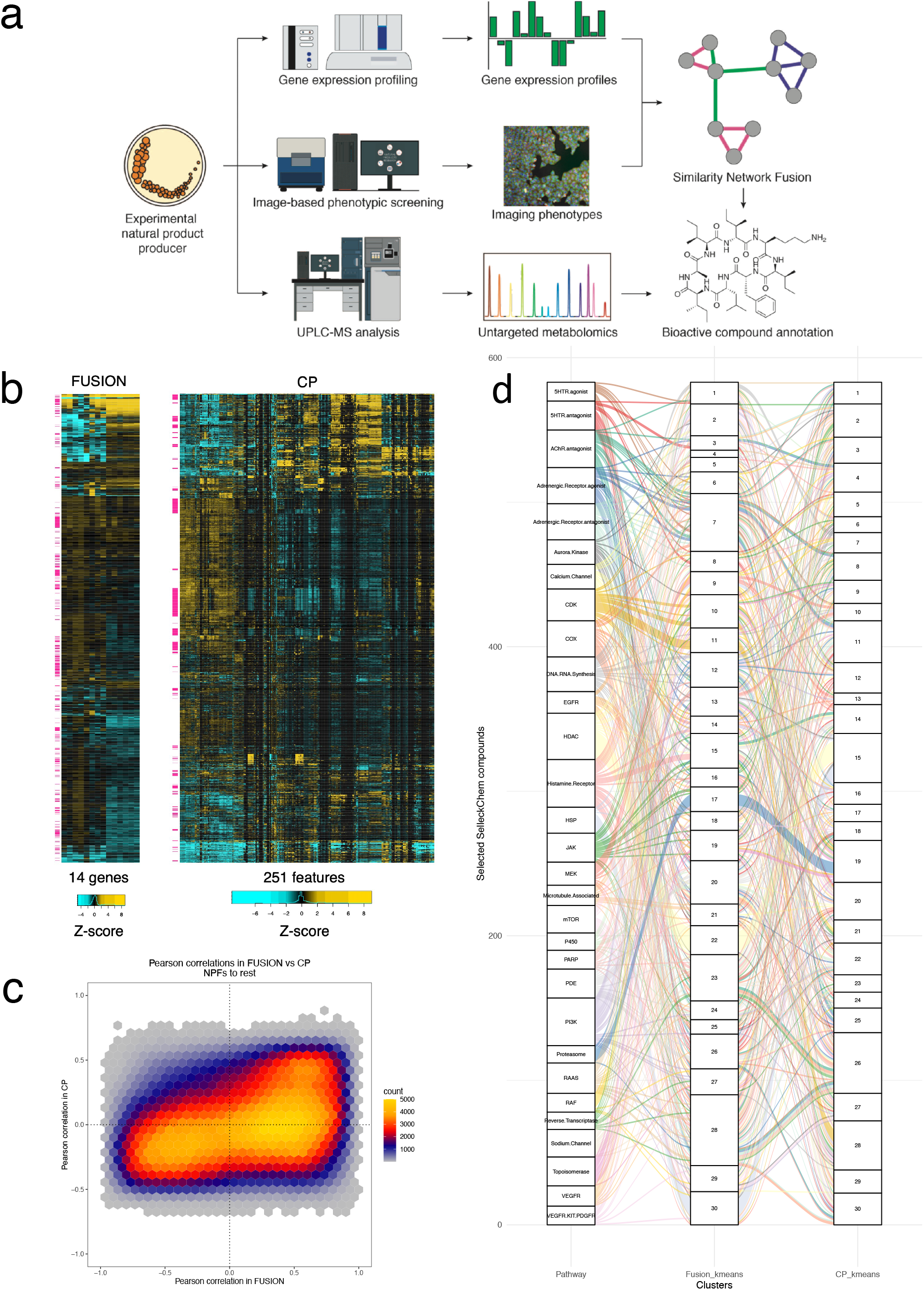
Overview of screening platforms and initial data collection. A) Experimental outline. Natural product fractions are isolated from marine bacteria, screened through two biological screening platforms (FUSION and CP), and subjected to high-resolution mass spectrometry-based metabolomics profiling. FUSION and CP data are integrated using Similarity Network Fusion (SNF) which then provides biological annotation on individual metabolites identified. B) Two-way hierarchical clustering of Z-scores from FUSION and CP using Euclidean distance and complete linkage. NPFs are indicated with pink flags. C) Heat-scatter hexplot comparing Pearson correlations between NPFs and all other perturbagens in FUSION vs CP. D) Alluvial diagram comparing k-means clustering of chemicals in the top 30 largest target classes in FUSION and CP. Each target class is represented by a different color.

## Results

### Integration of multiple platforms retains in-class target classification

As a test-of-concept, we profiled a small collection of 628 randomly selected microbial natural product fractions in the FUSION, CP, and metabolomics profiling platforms (Figure 1A, Supplementary Note 1). For reference benchmarks, we also collected FUSION and CP profiles from a library of 2027 known synthetic small molecules that were selected for known bioactivity and have been annotated for mechanism of action and/or molecular target (Supplementary Figure 1, Supplementary Table 1, Supplementary Note 1). Briefly, all perturbagens were screened in triplicate on both platforms. FUSION gene expression signatures were normalized to non-treated wells, and CP fingerprints were normalized to DMSO-treated wells (Supplementary Note 3). A Z-score transformation was then applied to both normalized datasets. To evaluate differential sensitivity of these orthogonal platforms to detection of chemical activity, we classified as “quiet” any perturbagen where all Z-scored probe values were less than |0.5|. Approximately 33% of perturbagens were quiet in FUSION (n=889), and ∼25% of perturbagens were quiet in CP (n=686). Notably, natural product fractions comprised 25% of quiet perturbagens in FUSION (n=226) but only 0.7% of quiet perturbagens in CP (n=5). In total, intersection of these lists revealed that 12% of all perturbagens were quiet in both datasets (n=329; 3 natural product fractions and 326 Selleck chemicals), suggesting that integrating the two datasets will provide active signatures for a larger percentage of the total compounds.

Next we compared the dispersion of knowns and unknowns in each dataset using two-way hierarchical clustering (Figure 1B). We observed that natural product fractions were interspersed throughout the clustering of each dataset, with more dispersion in FUSION than in CP. This confirmed that the natural product fractions produced sufficiently divergent signatures to allow for similarity analyses with benchmark chemicals that target a broad biological space. Interestingly, comparison of the pair-wise Pearson correlations between natural product fractions and all other perturbagens in FUSION vs. CP revealed that while some correlations trend in the same direction, the overall concordance between the two datasets on a perturbagen-by-perturbagen basis is relatively low (Figure 1C). In fact, there are interactions that are negatively correlated in FUSION but positively correlated in CP. We also observe that the majority of Pearson correlations in CP fall within a relatively narrow range (Pearson r values between −0.5 and 0.5), while correlations in FUSION spread across the full range (Figure 1C, Supplementary Figure 2). This suggests that while the two platforms can report on the same biological space, CP may provide less resolution between MOAs than FUSION.

In order to assess concordance between the two datasets at the level of molecular target, we selected FUSION and CP signatures from the top 30 largest target classes in the Selleck library and applied k-means clustering (k=30) to this subset. Using the hypergeometric test with Bonferonni correction for multiple comparisons to score for significant enrichment of target classes within each cluster revealed that FUSION and CP have similar levels of sensitivity in terms of total number of target classes detected (19 and 22 target classes with p<0.00167, respectively), however there is notable divergence between the two datasets in terms of which target classes are detected (Supplementary Figure 3; Supplementary Figure 4). Comparison of cluster membership between the two datasets revealed that some target classes were robustly clustered together in both platforms (i.e., HSP, proteasome, HDAC), while others are clustered more closely in one dataset than another (i.e., Aurora Kinase inhibitors cluster more closely together in CP than in FUSION, and mTOR inhibitors cluster more closely together in FUSION than CP) (Figure 1D; Supplementary Table 2; Supplementary Figure 5). Moreover, comparison on a cluster-by-cluster basis of each dataset reveals that while both FUSION and CP are capable of clustering together target classes of similar MOA, the types of MOAs they pair are different in many cases (Supplementary Figure 6). Target class pairings that were observed in FUSION but not CP include mTOR and PI3K inhibitors in cluster 2, RAF and MEK inhibitors in cluster 9, topoisomerase and CDK inhibitors in cluster 10, and pan-RTK and EGFR inhibitors in cluster 25 (Supplementary Figure 4). By contrast, several other target class pairings were observed in CP but not in FUSION, including microtubule and JAK inhibitors in cluster 2, PARP and DNA/RNA synthesis inhibitors in cluster 7, HSP and HDAC inhibitors in cluster 15, and DNA/RNA synthesis and CDK inhibitors in cluster 28. Functional evidence supporting each of these pairings can be found in the literature. Importantly, many target classes were not effectively clustered together by k=30 clustering, but the identity of these classes were also divergent between the two datasets (Supplementary Figure 3). This lack of concordance could reflect differences between gene expression-based versus image-based readouts, low target expression in the cell lines used, mis-annotation of targets, polypharmacology within target annotated classes, and/or that k=30 clustering did not offer adequate resolution to discriminate between some target classes. Taken together, these analyses suggest that at this level of resolution, each dataset is able to cover the same biological space with comparable depth, but reports on the same space very differently.

Generation of a fused similarity network across CP and FUSION signatures would allow for the orthogonal information contained in the molecular and morphological phenotypic readouts to be leveraged simultaneously in the annotation of uncharacterized compounds. However, integration of orthogonal datasets is a computational challenge due to inherent differences in experimental collection, measured features, noise, and overall scale between methods (36). In order to test the idea that combining the information from FUSION and CP would lead to an improved platform for MOA assignment, we used a data integration approach called Similarity Network Fusion (SNF) (22). This method addresses challenges associated with differences in scale and feature measurement by first constructing within-sample similarity networks for each data type. A single similarity matrix is then generated by iteratively propagating similarity information simultaneously across all individual networks to generate a single, fused similarity matrix where perturbagens with evidence of similarity across multiple datasets result in higher similarity measures (Supplementary Figure 7, and Supplementary Note 4). To optimize for our high-content bioassay data, we adapted SNF by varying the value of *k* nearest neighbors and taking an agglomerate value of similarity across all *k* to generate a final matrix of similarity weights (see Supplementary Note 4). This matrix was then used to calculate a new Euclidean distance or Pearson correlation matrix, and subjected to hierarchical affinity propagation clustering (37, 38) to group perturbagens based on each metric (Supplementary Note 4).

In order to assess the performance of the individual and fused datasets in assigning MOA, we again used our collection of commercial compounds (Selleck) and their target annotations as benchmark references. Among the 195 pre-annotated target classes within this collection, 89 classes contained five or more chemicals. A two-sample, one-sided Kolmogorov–Smirnov (KS) test was applied to each of these 89 target classes to determine if the pairwise similarities between chemicals, as determined by FUSION or CP, with the same target annotation (“in-class”) were significantly closer or more correlated than the pairwise similarities between these chemicals and those from other target classes (“out-of-class”) (Supplementary Note 4). We compared Euclidean distance and Pearson correlation as similarity metrics. Perturbagens with high Pearson correlation will have signatures whose overall trend is in the same direction, but whose magnitudes may be very different. This can be useful when considering perturbagens which may have similar biological effects but different levels of potency, but will also have the effect of dispersing noise throughout the dataset. Meanwhile perturbagens with small Euclidean distances will have signatures which are closely related in both direction and magnitude. Thus, this metric can be particularly useful to make fine distinctions between different mechanisms of action, but may group noisy signatures together.

Comparison of the KS-test p-values for each target class across all datasets revealed that that SNF using Euclidean distance identified 25 of the 29 target classes identified in FUSION (86%) and 24 of the 32 target classes identified in CP (75%). We also observed that SNF-Euclidean identified significant self-association between members of 19 additional target classes (Figure 2A, 2D; CDF plots for each target class are included in Supplementary Figure 8). By contrast, SNF-Pearson identified 36 of the 38 target classes identified in FUSION (95%) and 43 of the 57 target classes identified in CP (75%), and 2 additional target classes that were not identified in either dataset (Figure 2B; Supplementary Figure 8; Supplementary Note 4). Notably, there also was a high degree of overlap in target classes identified by SNF-Euclidean and SNF-Pearson (Figure 2C). Taken together, these analyses demonstrated that valuable associations can be found in each dataset using either similarity metric, and that SNF retains at least 75% of the information found in individual datasets. Thus SNF is a platform in which the biological associations in both datasets can be leveraged together to provide functional annotation of compounds with unknown MOA.

**Figure 2.**
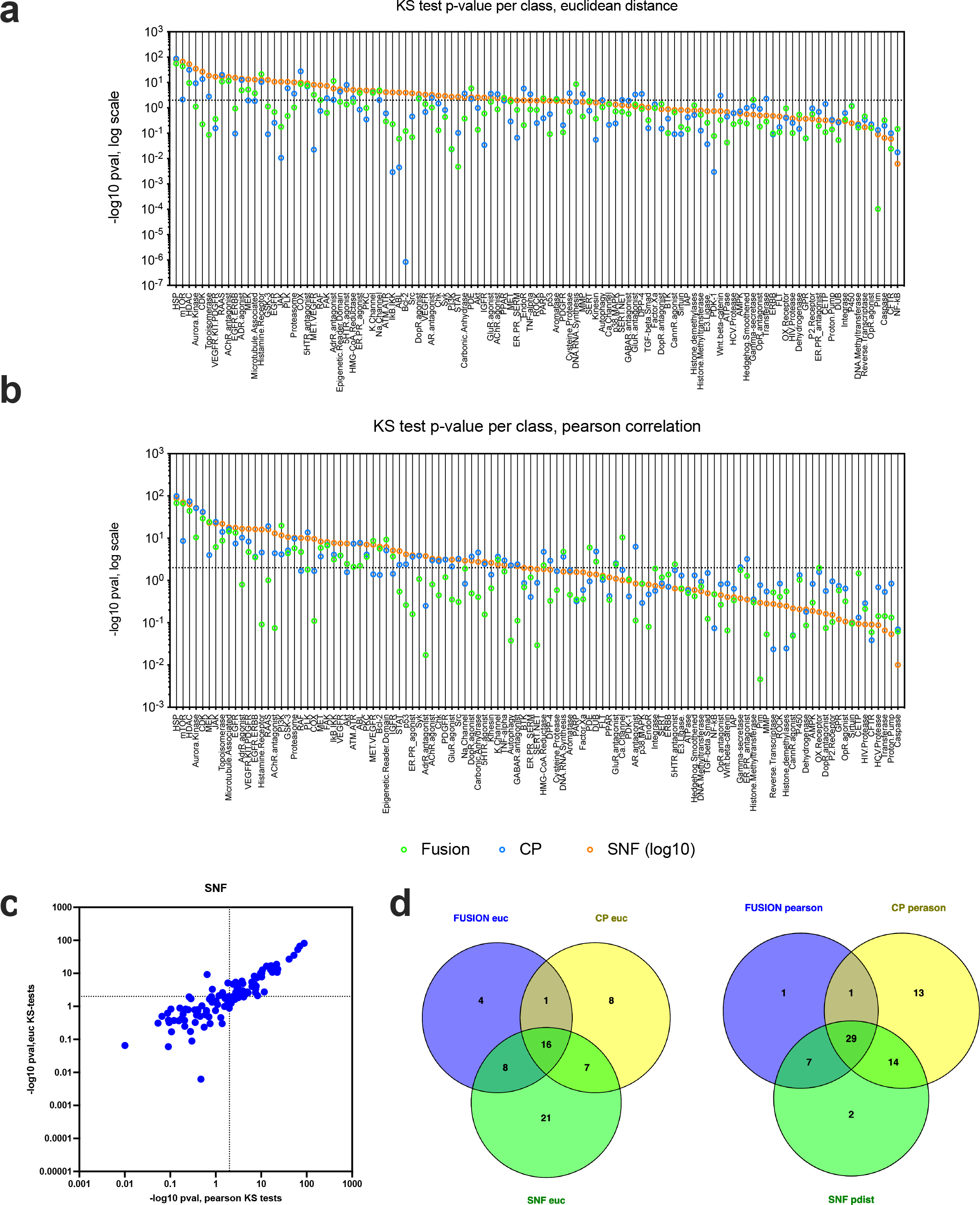
Comparison of KS-test p-values for in-class vs out-of-class target annotation in FUSION, CP, and SNF. Dot plots of −log_10_ KS-test p-values for each target annotation class in each dataset, using A) Euclidean distance or B) Pearson correlation as the similarity metric. Significance threshold is represented by the horizontal line (p=0.01). C) KS-test p-values for every target class with at least 5 members in SNF-Pearson vs. SNF-Euclidean. D) Venn diagrams illustrating the overlap in target classes scoring as significantly self-associated be KS-test (p<0.01) using Euclidean distance or Pearson correlation.

### SNF integration drives clustering of natural product fractions

We next used SNF values to construct a relational network among the reference compounds and natural product fractions using hierarchical affinity propagation clustering (APC) as described previously (38) (Figure 3). This clustering method was chosen as it is a deterministic method that defines, in a data-driven fashion, both the number and membership of clusters emerging from a given similarity matrix (37). Coloring edges based on the contribution from each individual dataset (see Supplementary Note 4) revealed that more than half of associations are supported by both datasets (∼54%), while ∼14% and ∼32% of associations are supported primarily by FUSION and CP, respectively (see Methods; Figure 3A). When these edge annotations are quantified on a per cluster basis, we observed that while most clusters are supported by both datasets, some clusters are driven by one dataset (i.e., Clusters 2 and 91 are driven by CP, while Clusters 120 and 125 are driven by FUSION; Figure 3B). Notably, most of the perturbagens that were flagged as “dead” in either platform clustered together in the SNF-Euclidean network, and this list included compounds for which cytotoxicity would be expected at the doses used in these assays (i.e., topoisomerase inhibitors; Supplementary Figure 9). In general agreement with our KS-test results, many clusters were significantly enriched for chemicals with the same target annotation, as assessed by a hypergeometric test (Figure 3C; Supplementary Table 3). We also observed that some clusters were significantly enriched for multiple classes, which may reflect similar mechanisms of action and/or convergence of downstream signaling effects (e.g., Cluster 123 is significantly enriched for PI3K, mTOR and EGFR target classes). It is also possible that overlap of multiple target classes in the same cluster may reflect a limitation of the gene set, cytological features, or the cell lines selected for profiling in both platforms, in that these reporters may not be sufficient to distinguish between those mechanistic classes. A comparison between the SNF-Euclidean and SNF-Pearson APC maps revealed that some target classes which can either be classified as closely related to other classes or be divided into subclasses have more separation in the SNF-Euclidean APC map compared to SNF-Pearson. For example, several chemicals annotated as “epigenetic reader domain” inhibitors cluster together with HDAC inhibitors in the SNF-Pearson APC map (Cluster 64), but separately from HDAC inhibitors in the SNF-Euclidean APC map (see Cluster 76). The SNF-Euclidean map is also able to cluster pan-CDK inhibitors separately from other more specific CDK inhibitors (Cluster 51). We also observe that there are more clusters in the SNF-Pearson APC map that contain multiple members of different classes than in the SNF-Euclidean map (Figure 3C; Supplementary Figure 10). The Pearson networks do clearly contain valuable associations (Figure 2B), which are likely to be informative across different biological contexts compared to the Euclidean distance networks. Our comparison of the networks suggests that SNF-Euclidean may have superior power to distinguish between related target classes than the SNF-Pearson network, and thus we chose to use this similarity metric in downstream analyses.

**Figure 3.**
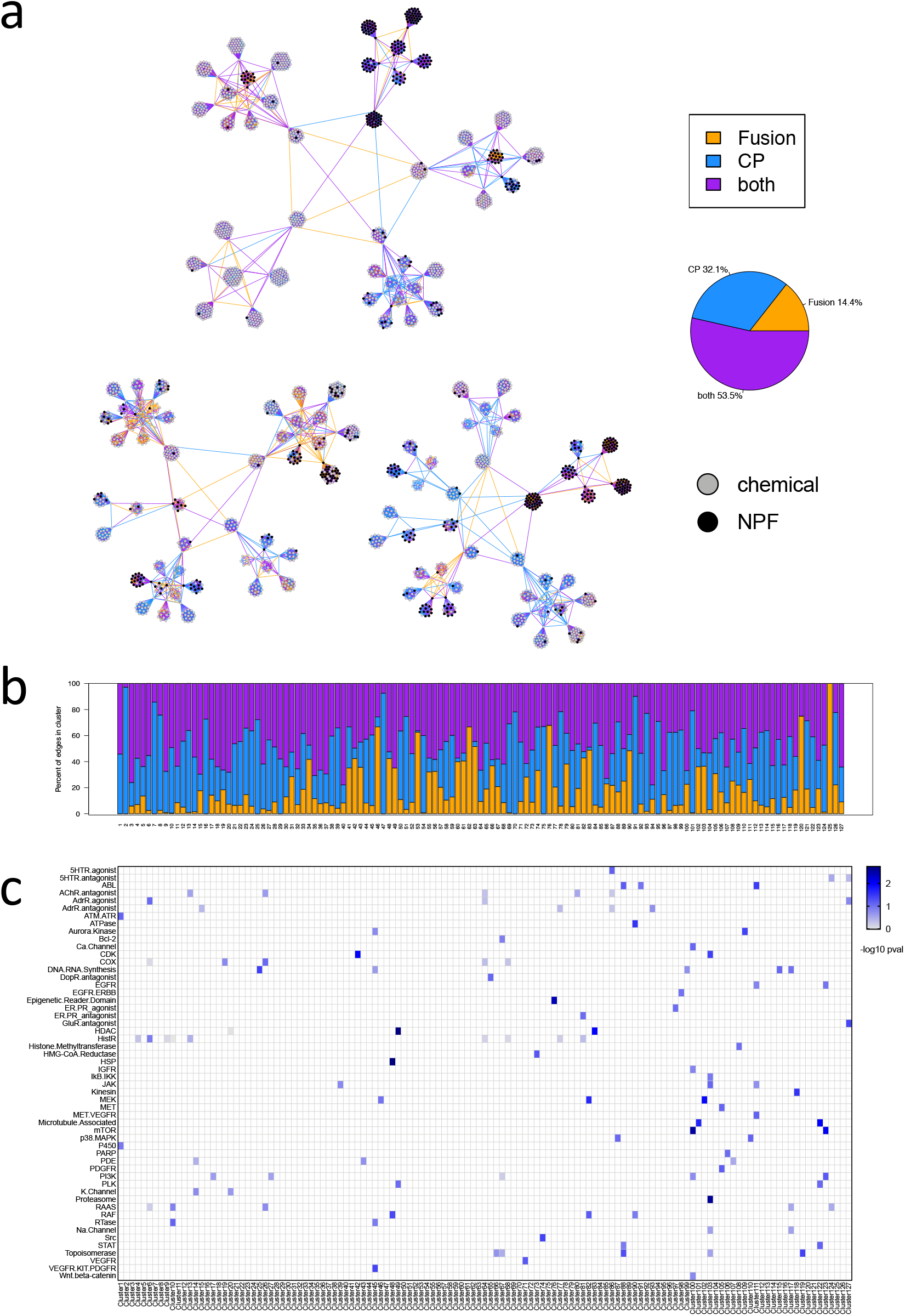
Affinity propagation clustering map of the SNF network preserves in-class target associations. A) Hierarchical affinity propagation clustering map of the SNF network using Euclidean distance as the similarity metric. Edges are colored based on contribution from individual datasets: Orange = supported by FUSION, blue = supported by CP, and purple = supported by both datasets. Perturbagen type is indicated by node color: black = NPF, gray = pure chemical. B) Barplot showing the percent of total edges in each APC cluster that are supported by FUSION, CP, or both datasets. Clusters are labeled with cluster number and the Selleck target class(es) which were significant by hypergeometric test in that cluster. C) Heatmap showing minus log10 p-values calculated by hypergeometric test for each target annotation class, per APC cluster. Target classes without significant enrichment in any cluster are omitted (Bonferroni-corrected alpha = 0.0016).

### Untargeted metabolomics relates chemical constitution to functional signatures via the SNF-similarity score

The chemical complexity of natural product fractions increases the difficulty in relating phenotypes to specific molecules or sets of molecules for a given sample. However, in most cases biological activities are driven by a single compound or a small subset of compounds in each extract (39). By determining the distribution of secondary metabolites across the full sample set, it is possible to test the hypothesis that extracts with similar phenotypes contain the same or similar bioactive species. In order to create a clear picture of chemical constitution across the sample set, we performed untargeted metabolomics on the full set of natural products extracts using a UPLC-IMS-qTOF instrument operating in data-independent acquisition mode (DIA) (Supplementary Note 2). Inclusion of ion mobility spectrometry affords an additional axis of separation over standard LCMS systems that improves separation of complex mixtures and provides an additional physicochemical measure (collisional cross-sectional area) for matching analytes between samples. Use of DIA increases the percentage of analytes that are subjected to MS^2^ fragmentation compared to traditional data-dependent acquisition (DDA). These fragmentation patterns are useful for comparing analytes between samples, and for comparing to external reference libraries for compound identification (40, 41).

Samples were analyzed as three independent technical replicates, and consensus feature lists generated for each sample using a suite of in-house data processing scripts. Mass spectrometric features were required to appear in at least two of three replicates to be included in the consensus feature list. These sample-by-sample feature lists were then ‘basketed’ to produce a single list of unique mass spectrometric features across the full sample set. This feature list included information about mass spectrometric properties (e.g. retention time, mass to charge ratio, collision cross-sectional area) as well as sample distribution (Supplementary Note 2).

For this initial study, a small set of 75 randomly selected strains of marine-derived Actinobacteria and Firmicutes from the MacMillan and Linington culture collections were grown in large-scale liquid culture, extracted using our standard extraction protocol, and prefractionated over C18 to afford 628 natural product fractions. Mass spectrometric analysis of these fractions identified a total of 8108 mass spectrometric features, of which 3498 appear only once in the sample set (43%). To examine the relationship between individual features and the SNF network, we employed a variation of our previously developed Compound Activity Mapping method to score predicted mass spectrometry feature activities (21). For each unique feature in the metabolomic dataset, we identified the subset of natural product fractions containing the feature and calculated the average of the SNF similarity scores within this set (see Supplementary Figure 11 and Supplementary Note 4). This score provides a numerical evaluation of how closely the presence of a specific mass feature is correlated with the presence of a specific biological phenotype in the APC network. In cases where a given feature is responsible for an observed activity, it is expected that the phenotypes of the associated set should be similar, and that the average SNF similarity score (SNF score, with a score range from 0 to 1) should be correspondingly high. By contrast, compounds that do not impart a biological response should not correlate with a specific biological signature, and the SNF score should be correspondingly weak. Using a 95^th^ percentile cutoff of all Euclidean distance-based SNF scores revealed 229 features with high correlation to biological activity (Supplementary Figure 12).

SNF scoring is feature-independent, meaning that a high score for one mass spectrometric feature has no impact on the scores of other features in the sample. This is important because the mass spectrometry data are not deconvoluted by either adduct (e.g. [M+H]^+^ vs [M+Na]^+^) or in-source fragments (e.g. [M-H_2_O+H]^+^). It is therefore common to identify a suite of mass spectrometry features with the same retention time that all possess strong SNF scores. These features can be used in concert to determine the correct accurate mass for the active component (which aids in dereplication) and to reconstitute mass spectrometry fragments (which can help with metabolite identification).

Calculating SNF scores for every mass spectrometric feature provides a metric to quickly identify bioactive compounds and prioritize them for subsequent isolation. A valuable visualization for these data is the Compound Activity Map (Figure 4A). In this network, extracts are represented by large nodes, while individual mass spectrometric features are represented by small nodes, color-coded by SNF score. Only mass spectrometric features with SNF scores above a set threshold are included, with edges added between extracts and the features they contain. The network is therefore arranged based on shared bioactive chemical features. Using this visualization, it is possible to identify sets of extracts with the same mechanistic prediction based on SNF annotations. Selection of clusters with similar SNF scores can be used to prioritize target molecules with shared biological properties. For example, fractions SW218953, SW218954 and SW218955 (Figure 4B) share a suite of related mass spectrometric features, including molecular ions, adducts and in source fragments, that possess similar extracted ion chromatograms (Figure 4C). These features also possess strong SNF scores (dark green nodes in Figure 4B) and identical activity predictions as shown by assignment to the same SNF APC cluster, suggestive of a single bioactive compound family in these samples. Conversely, in situations where clusters contain several classes of bioactive compounds, SNF scoring can be used to subdivide these clusters by chemical family. For example, samples SW218928, SW218929, SW218930 and SW218931 (Supplementary Figure 13) divide into two groups based on differences in mass spectrometric features, suggesting the presence of two separate compound classes within these related extracts.

**Figure 4:**
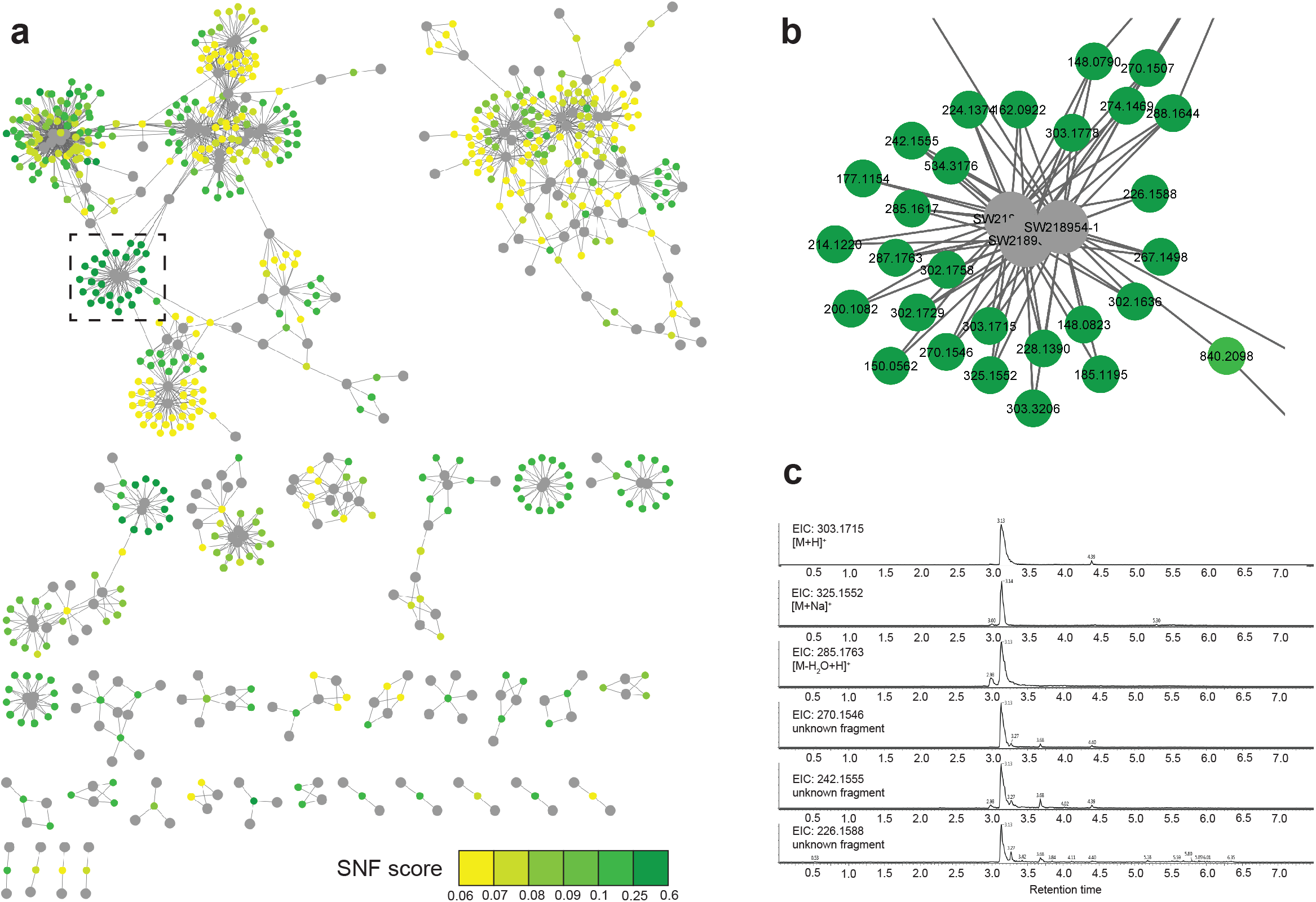
Compound Activity map for combined SNF profiles and untargeted metabolomics features. Large nodes represent extracts. Small nodes represent mass spectrometry features. Edges represent presence of mass spectrometric features in connected extracts. Only mass spectrometric features with predicted SNF scores >0.06 are included. A) Full Map. small nodes color coded by SNF score. B) Expansion of a representative region of the APC map with large nodes coloured by SNF score. C) EIC of mass spectrometric features present in adjacent fractions with similar SNF scores.

Compound Activity Maps thus provide a powerful strategy to prioritize candidate compounds for isolation. However, this visualization is centered on biological attributes, and provides less information about chemical properties. As a complement to Compound Activity Mapping we used the open source Bokeh server library to create a data visualization tool that enables direct examination and filtering of the untargeted metabolomics data with a range of data display options (Supplementary Figure 11). This platform can display metabolomics data as plots of retention time vs *m/z* ratio, filtering based on SNF score, presence in extract list, or both. This viewpoint on the data is valuable for selecting lead compounds that not only score well based on bioactivity predictions, but also display robust chemical signatures for subsequent isolation and structure elucidation.

In order to evaluate the efficiency of this new platform for *de novo* bioactivity prediction from complex mixtures, we tested two different query approaches; 1) querying the SNF network for natural products clustering predominately near other reference compounds, (biology-first discovery), and 2) filtering for metabolites with highly correlated biological activity as assessed by SNF score (chemistry-first discovery). The first case highlights compounds that possess mechanisms in line with known bioactives while the second approach identified sets of compounds with mechanisms that are not covered by the training set of known bioactives. This second approach attempts to address one of the largest challenges in natural products, that is, identification of new natural products with novel mechanisms of action. These strategies were selected to test the platform under different conditions, from simple situations where the annotations were unanimous, to complex situations with multiple reference compound types and multiple natural product fractions. In each approach, we highlight the contribution of SNF and metabolomics towards identification of both the natural product driving the signature and its biological mechanism of action.

### Identification of trichostatin A from an HDAC-inhibitor enriched cluster validates the integrated SNF platform

We first sought to validate the SNF network by querying the dataset for natural product fractions that clustered mainly with reference compounds from a single target class. In the SNF-Euclidean APC, there were 6 clusters that were highly enriched for chemicals belonging to the same target class (p<1E-10): Cluster 48 (HSP), Cluster 49 (HDAC), Cluster 76 (Epigenetic Reader Domain), Cluster 100 (mTOR), and Cluster 103 (Proteasome) (Figure 3B; Supplementary Table 4). Of these, Clusters 49, 100 and 103 contained natural product fractions (3 in Cluster 49, 1 in Cluster 100, and 1 in Cluster 103), thus identifying readily testable MOA hypotheses for the bioactive natural products present in each case. A KS-test confirmed that association between chemicals in the HDAC inhibitor target class is preserved in the full dataset, and that these associations were still significantly improved in SNF compared to FUSION or CP (p=1.8e-61; Figure 5A).

**Figure 5.**
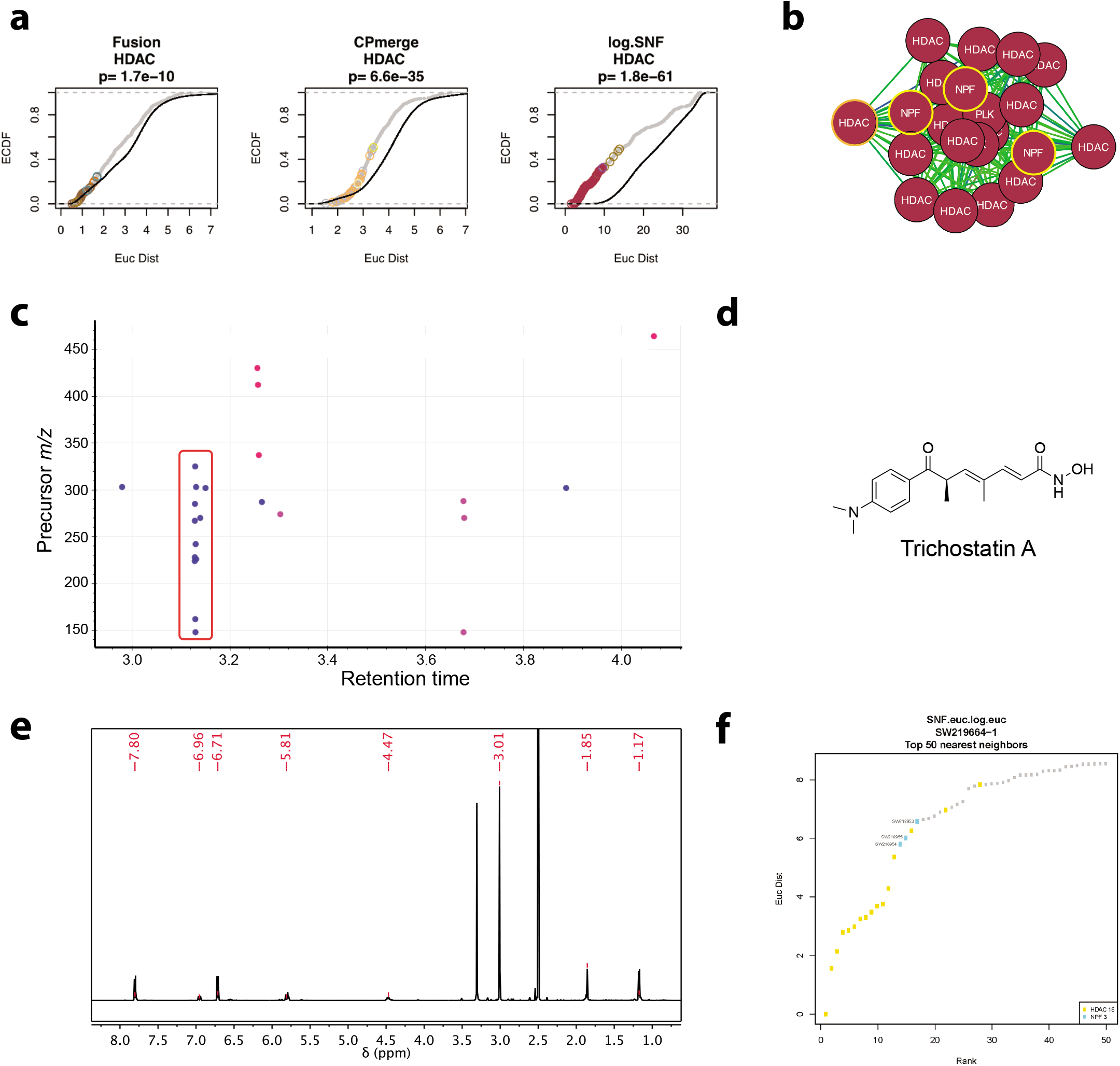
SNF correctly assigns MOA to the major metabolite in a series of natural product fractions. A) CDF plots comparing pairwise Euclidean distances between chemicals annotated as HDAC inhibitors in the full dataset, versus out-of-class associations. Gray, in-class; Black, out-of-class. Colored circles correspond to cluster membership in the associated APC map. The p-value was calculated by KS-test. B) Cluster 49 from the SNF-Euclidean map, drawn using a spring-embedded layout such that edge length is proportional to Euclidean distance. C) Retention plot showing common mass spec features present in the NPFs highlighted panel B. D) Chemical structure of trichostatin A. E) NMR spectra confirming trichostatin A. F) The top 50 nearest neighbors to trichostatin A in the SNF-Euclidean network, with the three natural product fractions found in Cluster 49 labeled.

We observed that the HDAC inhibitor cluster contained 3 sequentially isolated natural product fractions (SW218953, SW218954, and SW218955) (Figure 5). The presence of multiple NPFs from the same series suggests the presence of common metabolite profiles. Filtering the metabolomics data for features that are only present in these three natural product fractions revealed a vertical “stripe” of mass spectrometry features at 3.13 minutes with a parent mass feature of 303.1712*m/z* (Figure 5C; red box). This pattern of signals is indicative of both a parent mass and associated in-source fragments from the LCMS analysis (extracted ion chromatograms for the NP fractions are included in Supplementary Figure 14). Subsequent chromatographic optimization, purification and NMR analysis from SW218953 (Figure 5D) identified this product as the known bacterial metabolite trichostatin A (Figure 5E). Trichostatin A has been extensively studied for its activity as an HDAC inhibitor (42, 43). Notably, Cluster 49 also contained pure trichostatin A from the Selleck library (Supplementary Table 5). An analysis of the top 50 nearest neighbors to trichostatin A in the SNF-Euclidean dataset also confirms that these three natural product fractions are tightly associated with HDAC inhibitors (Figure 5F). Collectively, the natural product fractions containing trichostatin A also had a very high SNF score (0.537; 99.9^th^ percentile). Therefore, our integrated platform successfully assigned known MOA to natural product fractions in an agnostic manner, confirming the power of this annotation strategy.

### Metabolomics-guided SNF cluster selection identifies desferrioxamine class of natural products

Next we explored whether the integrated FUSION-CP dataset could be used to identify novel bioactive metabolites in our natural product libraries. Filtering the metabolomics data for SNF scores near the 80^th^ percentile (SNF score ∼0.053) identified two related molecules with parent mass-to-charge ratios of 587.3395 and 601.3567 and the same molecular fragments, present in SW218716, SW218717 and SW218718, SW218850, SW218755 and a few other natural product fractions from multiple different bacteria (Supplementary Figure 15. Chromatographic optimization, purification and structure elucidation identified these metabolites as desferrioxamines D2 and E (DFOD2 and DFOE) (Supplementary Note 5, Supplementary Figure 16). Querying the SNF-Euclidean network for the top 50 nearest neighbors to SW218850 revealed that it is closely associated with other DFOE-containing natural product fractions, as well as with several known iron chelators: two formulations of Ciclopirox (Selleck catalog numbers S2528 and S3019), and Deferasirox (Selleck S1712) (Supplementary Figure 16).

These data confirm that untargeted metabolomics with SNF score annotation can be used to quickly identify the active metabolite in fractions that share a phenotype. This is also an example of a commonly encountered scenario in natural products research, where several natural product fractions show a distinctive phenotype that is driven by the presence of a commonly encountered compound class. In this case, we were able to rapidly determine that the natural product fractions in this grouping are highly enriched by the DFO family of iron chelators.

### SNF-score driven identification of surugamide A as modulator of CDK activity

Using a top 98^th^ percentile cutoff of SNF scores also identified a single natural product (parent mass-to-charge ratio of 912.6266) that was present across multiple extracts from multiple bacterial species (SW218824, SW218835, SW218858, SW218859, and others; Figure 6A, Supplementary Figure 17). Purification and structure elucidation identified this metabolite as the cyclic octapeptide surugamide A (Figure 6B, Supplementary Note 5, Supplementary Figure 17) (44). The surugamides are a recently discovered class of cyclic peptides that appear to be widely distributed in *Streptomyces* sp. Initial biological activity reports for surugamide A show weak activity as protease inhibitors with an IC_50_ of 21 µM in an enzymatic assay for inhibition of bovine cathepsin B (44). Querying the SNF-Euclidean APC network showed several surugamide-containing fractions clustering near each other and in close proximity to CDK inhibitors (Supplementary Figure 17C). The retinoblastoma tumor suppressor (Rb) function is tightly controlled by CDK complex proteins and the phosphorylation state of Rb is indicative of cell cycle progression (45). Based on this APC clustering, we evaluated surugamide A (10 µM) for its ability to inhibit Rb phosphorylation, compared to the CDK inhibitor dinaciclib (100 nM). Western blot analysis of surugamide A treatment clearly shows strong suppression of Rb phosphorylation on Ser 807/811 relative to untreated cells (Figure 6C).

**Figure 6.**
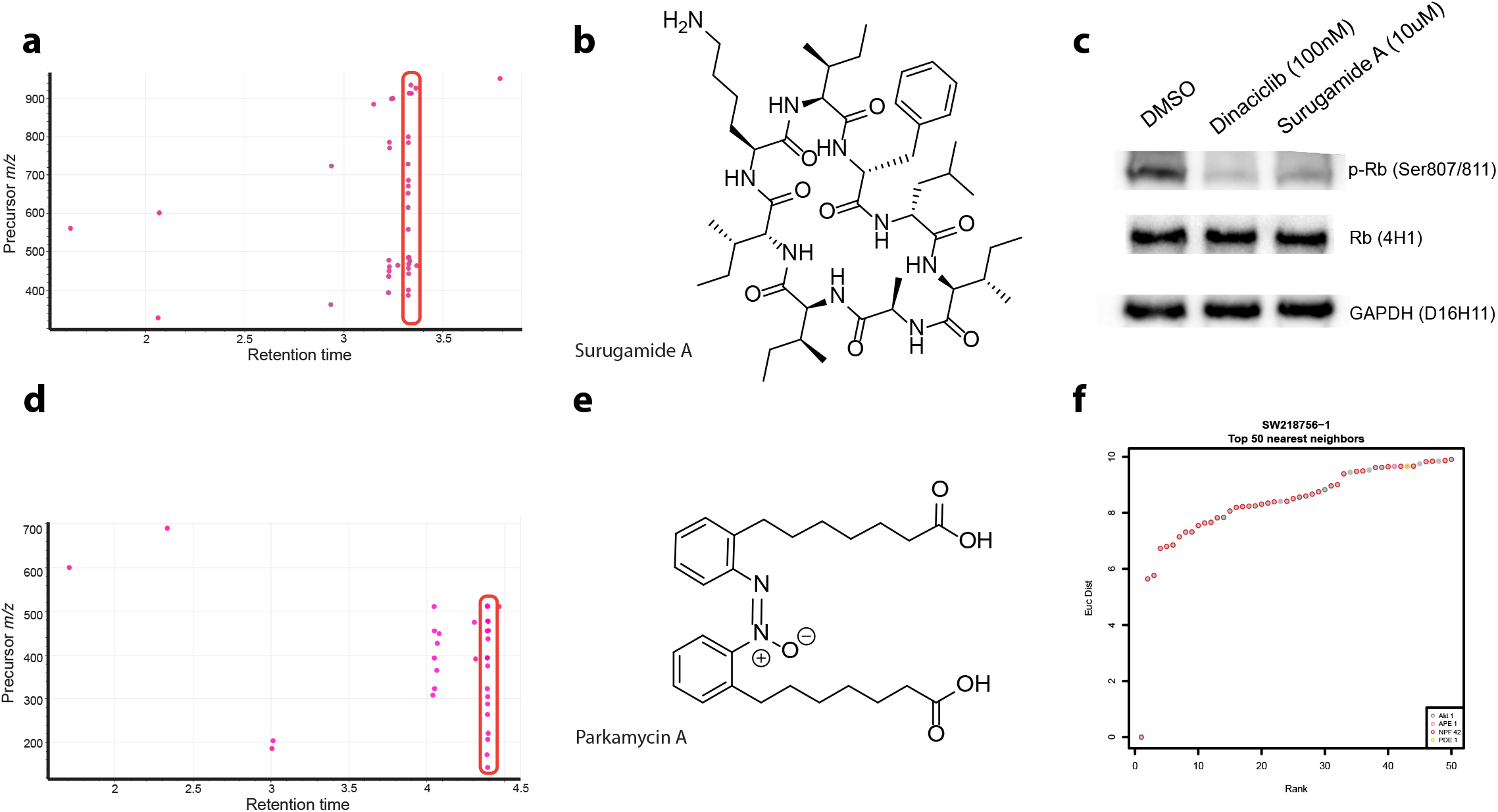
Filtering for highly correlated SNF scores can identify single metabolites with biological activity. A) Retention plot showing common mass spectrometry features associated with SNF scores above 0.5 for an APC region. B) Chemical structure of surugamide A. C) Immunoblot of changes in phospho-Rb protein expression in H23 cells after treatment with Dinaciclib for 24 hours. D) Retention plot showing common mass spectrometry features associated with SNF scores above 0.5 for a second APC region. E) Chemical structure of parkamycin A. F) The top 50 nearest neighbors to SW218756-1 in the SNF-Euclidean network. Points are colored by either target annotation or perturbagen type.

The identification of surugamide A in our dataset confirms that the use of the SNF score with untargeted metabolomics can also be used to quickly identify single bioactive metabolites in fractions that share a phenotype. In the case of surugamide A, clustering with a subset of CDK inhibitors provided a testable and validated biological hypothesis. Taken together, these data demonstrate that the bioinformatic integration of FUSION, CP, and metabolomics datasets can effectively drive the rapid discovery and characterization of bioactive natural products.

### SNF-guided discovery of Parkamycins A and B

Finally, several clusters in the APC map contained exclusively NP fractions, suggesting the presence of bioactive metabolites with mechanisms not represented in the training set. One of these clusters contained two isobaric species with strong SNF scores (*m/z* = 455.2604 and 455.2627, rt = 4.06 and 4.40 minutes; Figure 6D, Supplementary Figure 18), suggestive of a bioactive compound family. Refermentation and isolation yielded two molecules with matching UV spectra, one of which (parkamycin B) was highly unstable, rapidly converting to the more stable isomer (parkamycin A) on standing (Figure 6E). Structure determination of the more stable isomer using a full suite of NMR and spectroscopic techniques (Supplementary Note 6, NMR spectra as Supplementary Figures 19-28) identified parkamycin A as a novel natural product containing the highly unusual biphenylazoxy core pharmacophore. While azoxy motifs have some precedent in medicinal chemistry there are very few examples of this motif in nature (46). These results demonstrate the value of ‘chemistry-first’ prioritization methods for discovering novel natural product scaffolds with biological activities not represented in the reference compound training set.

## Discussion

The natural products literature contains thousands of examples of novel compounds with biological activities reported from simple end-point assays, such as cytotoxicity or antimicrobial growth inhibition assays. While this provides a handle for further investigation, the lack of detailed mechanistic information means that the majority of these molecules are never followed up on for biological characterization. This is due to the aforementioned challenges associated with characterizing the mode of action of pharmacological agents. Previous biological screening platforms developed by our laboratories (cytological profiling and FUSION) have been successful at characterizing new natural products with detailed mechanistic assignments (3, 8, 11, 16–19). While powerful, both platforms encountered scenarios where no prediction for a natural product fraction was possible due to weak signatures. Differences in both resolution and sensitivity between platforms can limit their utility, either because a given mechanism is not reported on by the assay system, or because the resolving power of the platform is insufficient to differentiate between mechanistic classes. In order to maximize the amount of information used to predict MOA, we applied an adapted version of Similarity Network Fusion (SNF) to integrate data from both cytological profiling and FUSION.

The SNF network retained the associations of many pure chemicals to their annotated target class that were observed in either FUSION or CP, delivering the capacity to leverage both datasets simultaneously for untargeted mode of action prediction. We validated the utility of this network in assigning MOA by demonstrating that natural product fractions containing trichostatin A were clustered with pure trichostatin A and other HDAC inhibitors, and similarly that fractions containing desferrioxamine derivatives clustered with other iron chelators. We then further developed a robust pipeline to assign the mechanistic annotation that the SNF network provides to specific natural product structures. Using untargeted metabolomic profiling of the full natural product fraction library and creating a scoring method (SNF score) to relate these mass spectrometry features to defined phenotypes, it is possible to directly predict the contributions of all mass spectrometry features to the biological landscape of the sample set. Development of the Bokeh server visualization suite (Supplemental Figure 11) also provides a facile platform for data filtering and visualization that enables the rapid exploration of these data using a range of different viewpoints. Using this approach, we were able to link surugamide with novel biological activity against CDKs. Thus, SNF scores provide a rich perspective on chemical and functional interpretation from the natural products library. For example, in situations where two different compound classes cause the same biological phenotype (i.e. one cluster in the APC network), SNF scores can correctly identify these two compounds as high priority candidates, even though neither compound is present in all members of the biological cluster, provided that each molecule is predominantly found within that cluster. Similarly, in situations where extracts contain many mass spectrometric features, most features will be quickly deprioritized because their distributions throughout the sample set do not correlate to specific biological phenotypes. Finally, SNF scores can be used to prioritize fractions that have biologically active natural products even in the absence of benchmark compounds, as demonstrated by the identification of parkamycins from clusters primarily enriched in other NPFs. The discovery of a novel natural product with no clear associations to the diverse biological space represented in our chemical training set confirms that our integrated approach can quickly identify high priority bioactive compounds even without biological annotation. This mechanism for compound prioritization is therefore a robust and powerful strategy for directly targeting biologically relevant compounds from large, complex natural product libraries, and can greatly accelerate the discovery of both novel chemistry and biology. An important aspect of new technologies is the ability to identify minor components in complex mixtures, which the integrated biological/metabolomics signatures are able to do.

Notwithstanding the value of this approach, there are several situations which remain difficult to resolve. Currently, the SNF score is not weighted by relative intensity of each MS feature. This is because determining relative concentrations of unknown analytes in complex samples remains an unsolved challenge in mass spectrometry-based metabolomics. In situations where a bioactive metabolite is present both above and below its EC_50_, the SNF score will be reduced, as there will be no measurable phenotype in extracts where the concentration is low. Secondly, the system cannot differentiate between active and inactive metabolites if they are always co-expressed. Review of our metabolomics dataset suggests that this circumstance is rare, however in these cases both metabolites would be scored as active candidates, requiring downstream deconvolution. Finally, in situations where bioactive compounds are frequently encountered with other unrelated bioactives, the resulting phenotypes could bear little relationship to one another. Review of the dataset suggests that this situation is also unusual, however in these cases SNF scores will also deteriorate because of the reduced similarity scores between samples with different phenotypic signatures.

The use of SNF to merge orthogonal data is limited by the breadth of space covered in the input datasets, both in terms of the reference training set and the number of features used for readout. The proliferation of information rich screening technologies, such as L1000 and cell painting, provide enhanced opportunities for chemical/ biological associations in natural product and other libraries (6, 8). While we chose to use FUSION and CP in this study, the approaches and methodology outlined here would be amenable to these platforms as well. The field of natural products will be greatly enhanced by the adoption of more complex screening platforms, as one of the major limitations in the field is the lack of mechanistic understanding of the large majority of isolated molecules.

Natural products research brings with it a number of challenges, such as the chemical complexity of extracts, re-isolation of known compounds and characterization of biological activity. These challenges limit the pace of natural product research and leave knowledge gaps around the value of a given natural product structure or class. Recent initiatives to develop resources to better understand the genomics of natural product biosynthetic gene clusters (47) and the development of the Global Natural Products Social (GNPS) molecular networking platform (41) have fundamentally changed how natural product research is conducted, but the field as a whole is far behind in leveraging ‘Big Data’ to address outstanding challenges. The approach we have detailed here provides an unbiased, data-driven platform that can be used to integrate biological assay and metabolomics results to provide a comprehensive viewpoint on chemical/biological relationships in the natural product sphere.

## Materials and Methods

Materials and methods are included in Supplemental Information.

## Acknowledgements

This research was supported by NIH grants U41 AT008718 to R.G.L, M.A.W and J.B.M (NCCIH and ODS) and R01 CA149833 to J.B.M (NCI). E.A.M was supported by a NIH training grant 5T32GM8203-27. R.M.V was supported by CPRIT training grant RP140110 and NIH training grant 5T32 CA124334-09. A.F.S was supported by NSF GRF#2017247469

## Lead Contact and Materials Availability

Further information and requests for resources and reagents should be directed to and will be fulfilled by the Lead Contacts, John B. MacMillan and Roger G. Linington.

