## Supplemental Information for "High-Throughput Functional Annotation of Natural Products by Integrated Activity Profiling"

**This PDF file includes:**

Supplementary text

Figures S1 to S28 and legends

Tables S1B to S5

SI References

**Other supplementary materials for this manuscript include the following:**

Table S1A

### SUPPORTING INFORMATION

#### Supplementary Note 1: Chemical Libraries

##### Selleck Library

The reference library is a set of 2027 molecules spanning 196 compound classes, with 789 compounds not belonging to any annotated class. The library was purchased from Selleck as a premade library, combining compounds from the Bioactive, Kinase Inhibitor, and FDA-approved drug libraries. The compounds were formatted into two sets of 384 well plates at 10 mM and 2 mM.

##### Natural product libraries

Two natural product libraries were utilized in this study. The MacMillan lab collection used in this study was comprised of ~500 fractions derived from 25 marine-derived bacterial strains. The library of microbial natural product fractions was derived from marine-derived Actinomycetes (20), Firmicutes (5). These bacteria were cultivated from marine sediment samples collected in Tonga, the Gulf of Mexico (Texas, Louisiana), estuaries in South Carolina, and the Bahamas. A variety of techniques were utilized to isolate strains, including the use of nutrient-limited isolation media, such as those composed of only humic or fulvic acid, the use of small-molecule signaling compounds (N-acylhomoserine lactones, siderophores) that mimic the natural environment of the bacteria of interest, and isolation of spores using density gradient ultracentrifugation. Selection of bacterial isolates was carried out based on morphological appearance and followed up by phylogenetic characterization using 16S rRNA analysis using the Universal 16S rRNA primers FC27 and RC 1492 for the majority of the phylogenetic analysis. 16S rRNA sequences were compared to sequences in available databases using the Basic Local Alignment Search Tool.

To generate the fraction library used in this study, bacterial strains were fermented in 5 × 2.8 L Fernbach flasks each containing 1 L of a seawater based medium (10 g starch, 4 g yeast extract, 2 g peptone, 1 g $CaCO_3$, 40 mg $Fe_2(SO_4)_3 \cdot 4H_2O$, 100 mg KBr) and shaken at 200 rpm for seven days at 27 ºC. After seven days of cultivation, XAD7-HP resin (20 g/L) was added to adsorb the organic products, and the culture and resin were shaken at 200 rpm for 2 h. The resin was filtered through cheesecloth, washed with deionized water, and eluted with acetone to give a crude extract, with an average yield of 2.0 g of crude extract/strain. Further fractionation of the bacterial crude extracts (~500 mg) was accomplished using an Isco medium pressure automatic purification system (equipped with UV and ELSD detectors) using reversed phase C18 chromatography (gradient from 90:10 $H_2O:CH_3CN$ to 0:100 $H_2O:CH_3CN$ over 25 minutes, RediSep Rf Gold High Performance column with 600 mg capacity). Fermentation of each bacterial strain gives rise to a total of either 10 or 20 natural product fractions/strain. All natural product fractions in the library are standardized to a concentration of 10 mg/mL in DMSO. All fractions have been analyzed by low resolution LC/MS using an Agilent Model 6130 single quadrupole instrument.

Due to the nature of the fractionation approach, there is by design, the potential for sequential fractions to contain the same compounds. For example, if the peak for staurosporine is split between fraction 6 and 7, one could expect to have a similar biological signature between the sequential fractions.

The Linington natural product fraction library contains >5000 microbial fractions. The library is comprised of extracts of marine sediment-derived bacterial strains isolated by the Linington laboratory over the past 10 years. The library contains a cross section of Gram-positive genera, enriched in Actinobacterial strains. These sediments were collected by hand using SCUBA from over 70 discrete dive sites on the west coast of the United States and Canada. Collection sites include locations from the Channel Islands (near Los Angeles, CA) to the San Juan Islands in northern Washington state and sites on Vancouver Island, and are part of one of the largest systematic sampling campaigns for marine microbial chemistry performed in this area. Samples were collected with 15-mL centrifuge tubes and isolates were obtained from one of 8 selection media each supplemented with 50 mg of nalidixic acid and cycloheximide. Selective media listed below which end in "F" contain 1 L of MilliQ water and media ending in "S" contain 750 mL of 0.2 μm-filtered seawater, and 250 mL of MilliQ water to a total volume of 1 L. These media are as follows: actinomycete isolation agar (AIF and AIS, Difco™), SNF and SNS[1], NTF and NTS, and HVF and HVS[2]. Plates were incubated at room temperature until the appearance of desired colony morphologies consistent with actinobacteria. Colonies were picked which displayed characteristic actinobacterial morphologies and were subcultured onto either MB (37.4 g DifcoTM Marine Broth, 18 g agar, 1 L Milli-Q water) or A1 (18 g agar, 20 g starch, 10 g glucose, 5 g yeast extract, 5 g NZ-amine, 1 g $CaCO_3$, 50 mg nalidixic acid, 50 mg cycloheximide, 31.2 g Instant Ocean, 1 L Milli-Q water) agar plates.

Isolates were inoculated from MB (18 g agar, 37.4 g marine broth (Difco™), 1 L MilliQ water) or A1 (18 g agar, 20 g soluble starch, 10 g glucose, 5 g yeast extract, 5 g NZ-amine, 1 g $CaCO_3$, 1 L MilliQ water) agar plates into 7 mL of modified SYP (mSYP) liquid media (10 g starch, 4 g peptone, 2 g yeast extract, 1 L Milli-Q water, 31.2 g Instant Ocean® sea salt) in 25 x 150 mm glass culture tubes for 2-3 days at room temperature with shaking at 200 rpm. These small-scale cultures were shaken at 25°C and 200 rpm for a minimum of three days before moving to 60-mL medium-scales. Cultures were stepped up to medium-scale by inoculating 3 mL of the small-scale culture into 60 mL of freshly prepared mSYP in wide-mouthed 250-mL Erlenmeyer flasks with small springs. Medium-scale cultures were shaken at 25°C and 200 rpm for 3-7 days. Large scale cultures were prepared by inoculating 40 mL of medium-scale culture into 1 L of freshly prepared mSYP in 2.8-L Fernbach flasks with a large spring and 20 g of pre-washed Amberlite XAD-16 adsorbent resin (DCM, MeOH, and water; Sigma). Large-scale cultures were shaken at 25°C and 200 rpm for 7-10 days. At the end of the fermentation period, cells and resin were separated from the culture medium by vacuum filtration using a Whatman® glass microfiber filter and washed with deionized water. Resin and cells from each culture flask were extracted with 250 mL of 1:1 DCM/MeOH. The organic extract was separated from the cells and resin by vacuum filtration and concentrated *in vacuo*.

Crude organic extracts from marine-derived actinobacteria were subjected to solid phase extraction using a Supelco-Discovery C18 cartridge (5 g) and eluted using a MeOH/$H_2O$ step gradient (40 mL; 10% MeOH, 20% MeOH, 40% MeOH, 60% MeOH, 80% MeOH, 100% MeOH, 100% EtOAc) to afford seven fractions. The 10% MeOH fraction was discarded and the remaining six (fractions A – F) concentrated to dryness in vacuo. Dry pre-fractions were resolubilized in 1 mL of dimethyl sulfoxide (DMSO) and transferred to deep-well 96-well plates for long-term storage at -70°C.

For biological analysis, DMSO stocks of library fractions were thawed, sonicated, and reformatted to 384-well format for high-throughput screening.

For mass spectrometry analysis, DMSO stocks of library fractions were thawed, sonicated, and diluted 1:20,000 (5:200 DMSO, 10:240 50% MeOH/$H_2O$, 10:200 50% MeOH/$H_2O$) with mixing in 96-well format. Plates were reformatted into 384-well format, sealed, centrifuged at 1000 rpm for 30s, and analyzed.

#### Supplementary Note 2: Metabolomics

*Data-Independent (DIA) UPLC-MS/MS Data Acquisition*

All measurements were performed with an Acquity UPLC H-Class (Waters) using an HSS T3 C18, 100 mm × 2.1 mm, 1.7 µm column (Waters). Separation of 5 µL sample was achieved by a gradient of (A) $H_2O$ + 0.01% FA to (B) MeCN + 0.01% FA at a flow rate of 500 µL/min and 45°C for 7.5 min (5% MeCN, 0-0.3 min; 5-90% MeCN, 0.3-4.7 min; 90-98% MeCN, 4.7-5.5 min; 98% MeCN, 5.5-5.8 min; 5% MeCN, 5.81-7.5 min). The LC flow was directly infused into a Synapt G2-Si operated in positive ion mode. Analysis was conducted using the $HDMS^E$ mode which was set to alternate between collision energies of 0eV and 30eV every 0.3 sec. The instrument was operated in electrospray mode with 20 µg/mL leucine enkephalin lockspray infusion enabled every 10 seconds. Mass spectra were acquired from 50-1500 *m/z* at a 2Hz scan rate in continuum mode without lockmass correction.

*Data Processing*

All samples were analyzed in triplicate for downstream processing. All samples were measured as 384-well plate batches. Measurements were performed over the course of 7 months. Measurements for each 384-well plate were performed within two weeks of each other, with all replicates of a single 384-well plate batch measured before moving on to the next set of samples. The instrument was mass calibrated and collisional-cross section (CCS) calibrated using MajorMix (Waters) between every 384-well set of samples run. The average *m/z* error was never greater than 0.9 ppm for the calibrant signals prior to acquisition.

##### Metabolomics processing pipeline

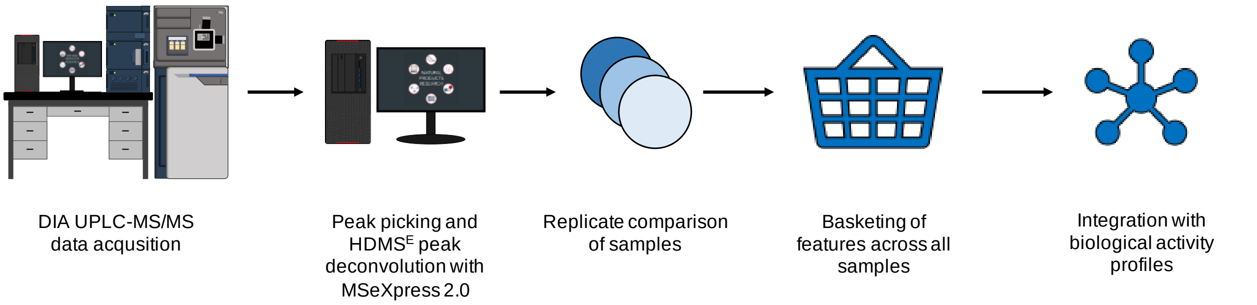

All raw data files were processed using a customized workflow developed in collaboration with Waters. The initial peak detection and $HDMS^E$ deconvolution software package MSeXpress 2.0 was employed to generate peak lists of precursor ions with their associated product ions for each sample. To ensure the validity of signals across replicates, another custom Python script (replicate comparison) removed any *m/z* feature that was not present in at least two of three analytical replicates within 0.03 Da and 0.05 min. Finally, all replicate-compared samples were basketed to give the distribution of all features across the sample set. Features within 0.03 Da and 0.05 min baskets were collapsed to single features, annotated by the samples in which they appear. Features that appeared in solvent blanks were removed prior to bioactivity integration in the Jupyter notebook.

#### Supplementary Note 3: Bioassays

##### Cell culture conditions

NCI-H23 cells were cultured in RPMI supplemented with 5% FBS (Gibco). HeLa cells were cultured in DMEM supplemented with 10% FBS (Difco, MT35015CV). All cells were cultured at 37°C under 5% $CO_2$.

##### Cytological Profiling

##### General

Briefly, HeLa cells were cultured and seeded into 384-well at 2,500 cells/well. After a 24-hour incubation, cells were pinned with test fractions (or pure compounds) using a Janus MDT robot (PerkinElmer). Two stain sets were used; Stain set 1: Hoechst, EdU-rhodamine, and anti-Phosphohistone H3, Stain Set 2: Hoechst, FITC-alpha tubulin, and rhodamine-phalloidin. For stain set 1, cells were incubated with 20 µM EdU for 1 h prior to fixing in 4 % formaldehyde solution in PBS for 20 min. Cells were then washed with PBS and permeabilized with 0.5% Triton-X in PBS for 10 min before blocking with 2% BSA-PBS solution for at least 1 h. Following this, cells were incubated with primary antibodies overnight at 4ºC. The following day, excess primary antibody was washed off with PBS and secondary antibodies and Hoechst solution were incubated for 1 h. Plates were washed with PBS and preserved with 0.1% sodium azide in PBS solution prior to imaging. For stain set 2, cells were fixed with a 4% formaldehyde solution in PBS for 20 min. Cells were then washed with PBS and permeabilized with 0.5% Triton-X in PBS for 10 min before blocking with 2% BSA-PBS solution for at least 1 h. Following this, cells were incubated with primary antibodies overnight at 4ºC. After blocking the cells were washed and incubated with FITC conjugated anti-alpha tubulin antibody overnight at 4ºC. The following day the cells are washed and then incubated with rhodamine-phalloidin and Hoechst stain for 20 minutes. The cells were then given one final wash before imaging.

Selleck chemicals were screened at both 50 µM and 10 µM in CP. In the downstream analysis, signatures that were inactive in the 10 uM dose were replaced with the signature obtained from 50 µM treatment. Otherwise the signature produced at 10 uM was used for downstream anaylsis. Macmillan natural product fractions were screened at 10 µg/mL, and Linington natural product fractions were screened at 1000 x dilution from stock. NPFs that were cytotoxic in initial screening were run in 8 pt. 3-fold dilutions starting from 50 µg/ml to obtain active but not dead CP signatures. CP signatures of the first NPF dilutions that was within three SDs of the mean control cell count replaced the initial 50 µg/ml signatures and contributed CP data for SNF merging.

##### Imaging and Analysis

Two images per well were captured with an ImageXpress Micro XLS automated epiflourescent microscope (Molecular Devices, Sunnyvale). Images were then processed as previously described[3]. Briefly, initial image processing was performed using MetaXpress image analysis software, using built-in morphometry metrics, the multiwavelength cell scoring, transfluor, and micronuclei modules. Custom written scripts were used to compare the treated measurements with the measurements of the DMSO control wells, and then convert each feature to a “histogram difference” (HD) score. This produced a 408-feature vector fingerprint which was then reduced to 251 features using additional feature reduction steps. Uninformative features with zero standard deviation across all perturbations were removed. In addition, redundancy was further reduced using the find Correlation function in the R-package caret (version 6.0-79; R-version 3.3.3). Briefly, all feature pairs with Pearson correlation coefficients greater than 0.95 were flagged and the member of each pair with the highest mean correlation to all other features was removed, resulting in the final 251-feature CP fingerprint. Compound treatment wells were labeled as ‘dead’ if the cell count for the treatment well was < 10% of the Median cell Count in the treatment plate. This resulted in 31 compounds and NPFs to be labeled as 'dead'. CP scores were calculated as the square-root of the sum of the squares of the CP features.

$$CPscore = \sqrt{\sum_{i=feature\ 1}^{n\ features} {x_i}^2}$$

#### FUSION

##### Assay and data processing

All perturbagens were screened in triplicate in the human lung cancer cell line NCI-H23 in 384-well microtiter plate format. Cells were seeded at a density of 5000 cells/well in either 50 or 60 µL of RPMI containing 5% FBS and 1000 U/mL Penicillin-Streptomycin (ThermoFisher). The next day, cells were treated with natural product fraction extract (10 µg/mL from the Macmillan library, and 1000x dilution from the Linington library) or Selleck chemicals (10 µM) using an Echo 555 Liquid Handler (Labcyte; Sunnvale, CA). After 24 hours of incubation, QuantiGene Plex 2.0 Assay Lysis Mixture (ThermoFisher) with 10 µL/mL proteinase K was added to each well for a final media:Lysis Mixture ratio of 2:1. Cells were lysed at 54°C for 30 min, and then stored at -80°C.

The FUSION assay concept was described previously[4]. We extended this concept to a lung cancer context by selecting a new set of genes that can report on the physiological state of lung cancer cell lines in particular (manuscript in preparation). Expression of 14 dynamic reporter genes (*DUSP6, FAM3C, GCNT3, GRHL2, HSD17B7, KIAA0922, LCN2, LTBR, RRM2, SIRPA, TLE2, TMEM30B, WSB2, YAP1*) and 2 static reporter genes (*EEF1A1, SIRT6*) were detected using a 16-plex QuantiGene Plex 2.0 Assay (ThermoFisher). Using a panel of four different sets of 16-plex magnetic and fluorescent beads allowed 4 experiments to be multiplexed into each well, for a final plex of 4 experiments x 16 genes = 64 measurements. An aliquot of cell lysate (~44-47% of the original volume) was then transferred to V-bottom 384-well plates and incubated with bead and hybridization probe mixes at 54°C in a MaxQ 4450 benchtop orbital shaker at 300 rpm for a minimum of 18 hours. After hybridization, samples were 4-plexed by bead set into flat-bottom 384-well plates using a Biomek robotic liquid handler and 384-well plate magnet. Plates were washed in Quantigene Assay Wash Buffer using an ELx405 CW plate washer with a 384-well plate magnet. The signal was then amplified per manufacturer's protocol using pre-amplifier, amplifier, biotin label, at a ratio of 7.5 µL/mL Label Probe Diluent, and streptavidin-phycoerythrin reagents at a ratio of 7.5 µL/mL SAPE Diluent. Plates were then washed in SAPE Wash Buffer and read on a Flexmap 3D (Luminex).

FUSION gene expression data is processed first by deconvolution into bead sets, and then normalizing the raw median fluorescence intensity (MFI) value for each gene to the median MFI of that gene on a plate-by-plate basis. To control for cell number, each gene is normalized to the geometric mean of the static controls (*EEF1A1* and *SIRT6)* on a well-by-well basis. Expression values which are > 2x the standard deviation of all values for that gene are then filtered out to eliminate spurious outliers, and only perturbagens with at least 2 replicate values for each gene are processed further. Gene signatures are then normalized to the median of "no treatment" control wells on a plate-by-plate basis, collapsed to the median of each gene, and $\log_2$ transformed to obtain the final FUSION signature.

**Compound activity assessment**

**Immunoblot analysis**

H23 cells were plated in 6-well format at a density of 150,000 cells/well and allowed to incubate overnight before drug treatment. Upon 24 hour drug treatment, cells were lysed in 1% Triton X-100 buffer mixed with 1X protease and phosphatase inhibitor cocktails (Thermo Scientific). 20 µg of each lysate was loaded and electrophoresed on a 10% SDS-PAGE gel (Bio-Rad) and transferred to a PVDF membrane using the Trans-blot turbo transfer system (Bio-Rad). After blocking with Starting Block (PBS) Blocking Buffer (Thermo Scientific), membranes were probed overnight with primary antibodies diluted at 1:1,000 at 4°C according to manufacturer recommendations. Antibodies against GAPDH (#5174), Rb (# 9309), p-Rb (# 8516) were purchased from Cell Signaling Technology. After washing and incubation with the secondary antibody (HRP-conjugated anti-mouse and anti-rabbit IgG antibodies, Jackson Immunoresearch), protein signals were visualized with ECL Select detection solution (Amersham). GAPDH was used as whole cell lysate loading control.

#### Supplementary Note 4: Statistical analysis

##### Similarity Network Fusion (SNF) Methodology

SNF is a novel similarity metric designed to aggregate information across multiple datasets and assign a similarity score to perturbations based on evidence from multiple datasets. SNF was performed as previously described[5], with some modifications (Supplementary Figure 1). Briefly, similarity within a dataset was first calculated. For perturbations i and j, similarity within a single dataset is calculated as :

$$\mathbf{W}(i,j) = \exp\left( -\frac{\rho^2(x_i, x_j)}{\mu \varepsilon_{i,j}} \right)$$

where $w_{i,j}$ gives the scaled similarity and $\rho(x, x_j)$ gives a similarity score (i.e., Euclidean distances) between perturbations i and j in one dataset. For Euclidean distances we used the dist2 function in the SNFtool package. For Pearson correlations, the distance matrix function from the R package ClassDiscovery was used to calculate a Pearson distance similarity matrix. W is calculated for each input dataset to be fused with SNF. The $\mu$ value is a hyperparameter that is empirically set and $\varepsilon$ is used to eliminate the scaling problem. $\varepsilon$ is given as:

$$\varepsilon_{i,j} = \frac{\text{mean}(\rho(x_i, N_i)) + \text{mean}(\rho(x_j, N_j)) + \rho(x_i, x_j)}{3}$$

To compute similarity across multiple sources of data with different scales of measurement a normalized weight matrix, P, was defined where $P = D^{-1}W$ where is D is the diagonal matrix whose entries D(i,i) = $\sum_j W(i,j)$. From here, K-nearest neighbors was used to measure self-similarities as follows:

$$\mathbf{S}(i,j) = \begin{cases} \frac{\mathbf{W}(i,j)}{\sum_{k \in N_i} \mathbf{W}(i,k)}, & j \in N_i \\ 0 & \text{otherwise} \end{cases}$$

In this step, the k-nearest neighbors to a perturbation will be assigned similarity scores and the others will be set to zero. We found this step to be critical in introducing errors in to the similarity measures for our purposes. Setting the similarity value to zero for similarity measures outside the k-nearest neighbors (KNN) effectively masked perturbation similarity in otherwise significantly similar associations. Thus, we chose to vary k from k=2 to k=n/2, where n is the total number of perturbations in each dataset, and use an agglomerate value of similarity across all k. Networks were then fused by prorogation information from each dataset into the other according to the following equation:

$$\mathbf{P}_{t+1}^{(1)} = \mathbf{S}^{(1)} \times \mathbf{P}_t^{(2)} \times (\mathbf{S}^{(1)})^T$$

$$\mathbf{P}_{t+1}^{(2)} = \mathbf{S}^{(2)} \times \mathbf{P}_t^{(1)} \times (\mathbf{S}^{(2)})^T$$

P and S are defined above where P carries information about the full similarity network and S carries information about similarity for KNN. The superscripts (1) and (2) indicate similarity in data types 1 or 2 (here FUSION or CP). After each iteration, the P matrices were re-normalized as described above. This process is iteratively updated for t-steps after which the final fused matrix is calculated as:

$$\mathbf{P}^{(c)} = \frac{\mathbf{P}_t^{(1)} + \mathbf{P}_t^{(2)}}{2} .$$

As stated above, we perform SNF individually for values of k varying from k=2 to k=n/2, resulting in n/2-1 matrices total. To compute an aggregate matrix, we first normalize within each matrix by dividing by the maximum non-diagonal value and then take the average value between all matrices to result in a final, fused aggregate similarity matrix. This matrix was then $\log_{10}$-transformed and clustered by affinity propagation clustering using either Euclidean distance or Pearson correlation as the similarity metric.

#### Clustering and statistics

Perturbagens were flagged as 'dead' in the CP dataset if the cell count for the treatment well was < 10% of the median cell count in the treatment plate. Perturbagens were flagged as 'dead' in the FUSION dataset if the geometric mean of the plate-median normalized MFI values for *EEF1A1* and *SIRT6* was ≤ 8.

The similarity between perturbagen profiles was measured using either Euclidean distance or Pearson correlation. To assess the ability of each dataset to identify significant associations within Selleck target classes, we used a two sample, one-sided Kolmogorov–Smirnov test (KS test) to determine if in-class associations were significantly smaller (Euclidean distance) or larger (Pearson correlation) than out-of-class associations. Only target classes with at least 5 members were considered for this analysis.

Hierarchical affinity propagation clustering was performed as previously described using Euclidean distance or Pearson correlation as the similarity metric[6]. FUSION data was clustered using the final 14 gene signatures, CP data was clustered using the final 251 features, and SNF data was clustered using the $W_{fused}$ matrix of weights after averaging over all k values. Networks were visualized in Cytoscape with edge lengths drawn proportional to the similarity metric using the Allegro Layout[5].

SNF network edges were colored according to dataset contribution as described below. For each pair-wise association, the $log_{10}$ ratio of the affinity values in each dataset (i.e., $Aff_{FUS}:Aff_{CP}$) were calculated. Ratios ≤ -0.5 were flagged as being supported by FUSION, ratios ≥ 0.5 were flagged as being supported by CP, and all other associations were flagged as being supported by both.

All data processing and statistical analyses for FUSION and SNF were carried out using the statistical platform R (http://www.R-project.org).

**Integration of Metabolomics to SNF, CP, and FUSION data**

Integration of the basketed metabolomics data was performed similarly to previous studies[7]. This approach treats every observed MS feature or basket as an individual chemical entity and asks the question, on average, what biological phenotype is expected when cells in the high content assays are treated with compound? It is acknowledged that combinations of multiple bioactive species as well as huge differences in titre are to be expected; however, the simplicity and naivety of the question serves as a reasonable starting point for further investigation.

In practice, each basket is treated as an object and assigned five numeric descriptors or attributes specific to the biological data acquisition: CP Cluster Score, CP Activity Score, FUSION Cluster Score, FUSION Activity Score, and SNF Cluster score. The Activity Score describes the strength or magnitude of the phenotype predicted when a basket is detected in an extract. The Activity Score values were calculated by first computing a numeric average of each of the observed attributes for each independent assay. The magnitude, square root of the sum of the squares, of this average predicted phenotype is the Activity Score. The Cluster Score describes the similarity of the phenotypes observed in the biological assays amongst all natural product extracts in which the basket was detected. The Cluster Score is computed using the NXN similarity matrices from each assay the combined SNF similarity matrix and is simply the average of the nondiagonal values of the sub NXN matrix consisting of all the natural product fractions containing that basket. These descriptors are then exported as a table that can be used for discovery and visualized using tools such as the custom Bokeh server.

#### Supplementary Note 5: Compound isolation

##### General

Solvents used for HPLC chromatography were Optima grade and were used without further purification. NMR spectra were acquired on a Bruker 600 MHz Avance II spectrometer equipped with a 5 mm QCI cryoprobe and referenced to residual solvent proton and carbon signals ($\delta_H$ 2.50, $\delta_C$ 39.5 for DMSO-*d6*, or $\delta_H$ 3.31, $\delta_C$ 49.0 for MeOD-*d4*). High-resolution mass spectrometry data were acquired using an electrospray ionization (ESI) accurate-mass time-of-flight (TOF) liquid chromatograph−mass spectrometer (Acquity UPLC H-Class tandem Synapt G2-Si).

##### Fermentation, and extraction

Isolate RL12-067-NTF-D (resulting extract SW218953), isolate RL12-121-HVF-C (resulting extract SW218858), isolate RL12-005-AIF-C (resulting extracts SW218716, and SW218717), and isolate RL12-115-HVF-E (resulting extracts SW218754, SW218755, SW218756) were inoculated from A1 agar plates into 10 mL of modified SYP (mSYP) liquid media (10 g starch, 4 g peptone, 2 g yeast extract, 1 L Milli-Q water, 31.2 g Instant Ocean) in 25 × 150 mm glass culture tubes for 2 days at room temperature with shaking at 200 rpm before moving to 60-mL medium-scale cultures. Cultures were stepped up to medium-scale by inoculating 3 mL of the small-scale culture into 60 mL of freshly prepared mSYP in wide-mouthed 250-mL Erlenmeyer flasks with small springs. Medium-scale cultures were shaken at 25°C and 200 rpm for 3 days. Large-scale cultures were prepared by inoculating 40 mL of medium scale culture into 1 L of freshly prepared mSYP in 2.8-L Fernbach flasks with a large spring and 20 g of pre-washed Amberlite HP-7 adsorbent resin (DCM, MeOH, and water; Sigma). Large scale cultures were shaken at 25°C and 200 rpm for 7days. At the end of the fermentation period, cells and resin were separated from the culture medium by vacuum filtration using a Whatman® glass microfiber filter and washed with deionized water. Resin and cells from each culture flask were extracted with 250 mL of 1:1 DCM/MeOH. The organic extract was separated from the cells and resin by vacuum filtration and concentrated *in vacuo*.

##### Fractionation

Crude organic extracts were subjected to solid phase extraction using a TeledyneIsco CombiFlash C18 cartridge (5 g) and eluted using a MeOH/$H_2O$ step gradient at 20 mL/min (40 mL; 10% MeOH, 20% MeOH, 40% MeOH, 60% MeOH, 80% MeOH, 100% MeOH, 100% EtOAc) to afford seven fractions. The 10% MeOH fraction was discarded and the remaining six (fractions A – F) concentrated to dryness *in vacuo*.

An aliquot (2.6 mg) of SW218953 (60 % MeOH) was found to be pure, providing structure **1** (2.6 mg, trichostatin A). An aliquot (12 mg) of SW218716 and SW218717 (40 and 60 % MeOH respectively) were combined before being subjected to RP-HPLC on a Phenomenex Synergi Fusion-RP C18 (10 x 250 mm, 10 µm) column. A gradient of acetonitrile and $H_2O$, both with 0.02% formic acid was used to purify compounds **2** (1 mg, Desferrioxamine D2) and **3** (2.2 mg Desferrioxamine E) (8 ml/min; 5% MeCN, 0-2 min; 5-

30 % MeCN, 2-20 min; 30-100% MeCN, 20-20.1 min; 100% MeCN 20.1-24 min; initial conditions). All of fraction (153 mg) SW218858 (80% MeOH) was subjected to RP-HPLC (20 ml/min, 62:38, $H_2O$: MeCN with 0.02 % formic acid) using an Atlantis T3 OBD (19 x 250 mm, 5 µm) column to give compound **4** (4.6 mg, surugamide A). All of fraction (96 mg) SW218756 was subjected to RP-HPLC (1.2 ml/min, 50:50, $H_2O$:MeCN with 0.02% formic acid) using a Phenomenex Kinetix C18 (4.6 x 300 mm, 2.6 µm) column to give compound **5** (7.9 mg, parkamycin A).

**Yields**

Approximate yields per liter of fermentation was as follows:

Compound **1**(trichostatin A) = 100 mg/L

Compound **2** (desferrioxamine D2) = 50 mg/L

Compound **3** (desferrioxamine E) = 110 mg/L

Compound **4** (surugamide A) = 4.6 mg/L

Compound **5** (parkamycin A) = 4.0 mg/L

#### Supplementary Note 6: Structure elucidation of parkamycin A

The molecular formula for Parkamycin A was determined to be $C_{36}H_{34}N_2O_5$ from HRESIMS data for the protonated and sodiated molecular ions (455.2536 and 477.2345 respectively) revealing eleven degrees of unsaturation. The $^1$H and $^{13}$C NMR revealed that the structure contained two carbonyl groups ($\delta_C$ 174.6 and 174.6) and 12 additional sp$^2$ hybridized centers ($\delta_C$ 121.1, 123.3, 126.2, 127.0, 128.5, 130.0, 130.2, 130.9, 134.7, 137.8, 142.0, and 148.9). $^1$H and $^{13}$C NMR also revealed 12 sp$^3$ hybridized methylene carbons ($\delta_C$ 24.6, 24.6, 28.4, 28.4, 28.6, 28.7, 30.2, 30.5, 30.6, 31.2, 34.0, and 34.0). Overall the spectra displayed a high degree of repetition, suggestive of a pseudo-dimeric structure. COSY and HMBC correlations combined with long-range $^4J_{HH}$ meta-couplings confirmed the presence of two ortho-substituted aromatic rings, accounting for 8 of the degrees of unsaturation and 12 of the 14 sp$^2$ hybridized carbons. COSY and HMBC correlations identified two pendant aliphatic chains, each containing six methylene units and terminating in a carbonyl group.

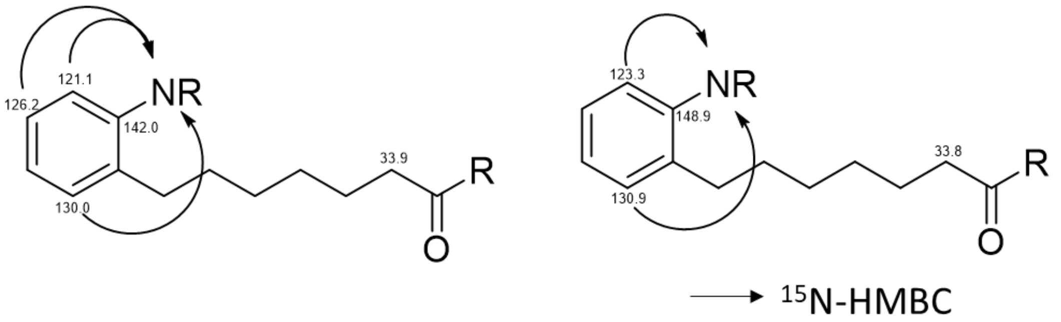

To place the remaining atoms, a $^{15}$N-HMBC spectrum was employed to determine the locations of the nitrogen atoms adjacent to each aromatic ring. This left five atoms ($H_2O_3$) remaining, as well as one double bond equivalent. This could be satisfied by either a cyclic compound with the carbonyls connected head to tail with the aryl nitrogen groups, or with an azoxy moeity joining the two aryl subunits and two carboxylic acid termini. To confirm the presense of the carboxylic acid termini, parkamycin A (0.1 mg) was treated with TMS-diazomethane (100 eq. dissovled in hexanes) in anhydrous methanol (0.5 mL) at room temperature for 15 minutes. LCMS analysis revealed both both partial (one conversion of COOH to COOMe (+14 Da)) and complete methylation products (two conversions of COOH to COOMe (+28 Da)) confirming the presence of the two acid termini.

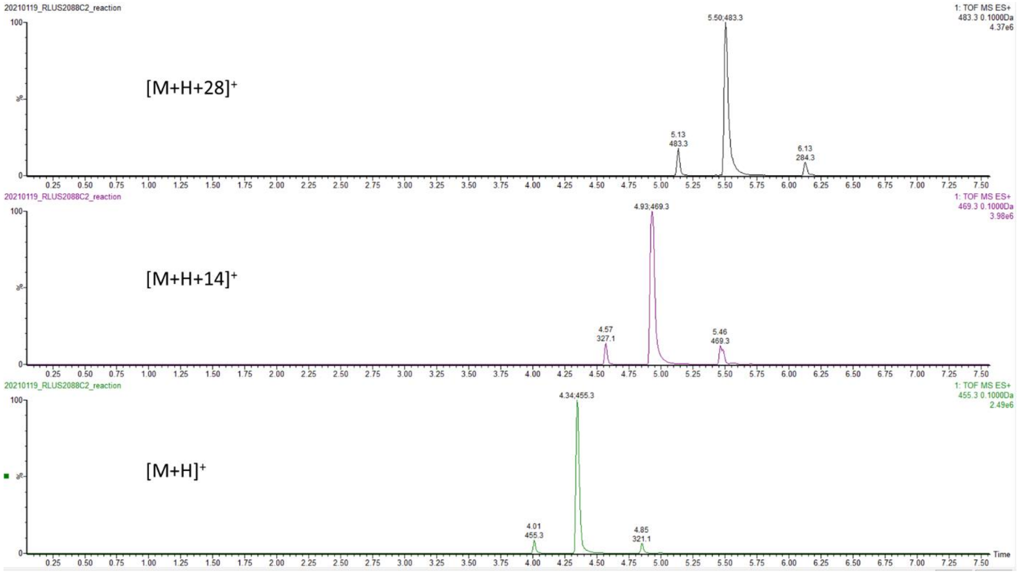

The remaining oxygen atom and double bond equivalent could only be satisfied by the presence of an azoxy moiety between the two aryl rings. This hypothesis was supported by the rapid interconversion between the cis and trans forms upon standing at room temperature. To determine which form was the more stable (parkamycin A) we employed IR spectroscopy, which showed a strong N=N stretch at 1465 $cm^{-1}$, consistent with the *cis* conformer (1468 $cm^{-1}$) rather than *trans* conformation (1415 $cm^{-1}$).[8]

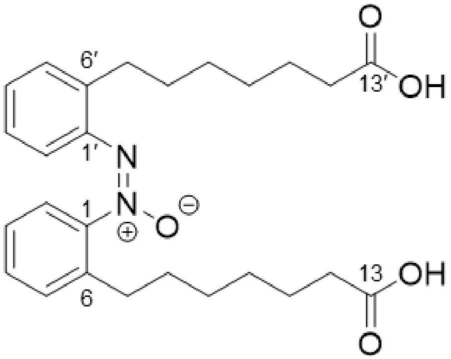

Parkamycin (**5**): UV (MeOH) $\lambda_{max}$ (log ε) 235 (2.45), 275 (2.31) nm; IR $\nu_{max}$ 2933, 1710, 1465 $cm^{-1}$. $^{1}$H NMR (DMSO-*d6*, 600 MHz) δ 7.84 (1H, d, *J* =7.1 Hz,H-2′), 7.62 (1H, d, *J* = 8.4 Hz,H-2), 7.51 (1H, t, *J* =7.5 Hz,H-4), 7.46 (1H, d, *J* =7.5 Hz,H-5), 7.42 (1H, t, *J* =7.6 Hz,H-3), 7.37 (1H, d, *J* =6.7 Hz,H-5′), 7.32 (2H, m, H-3′, H-4′), 2.73 (2H, t, *J* =7.7 Hz,H-7), 2.61 (2H, t, *J* =7.7 Hz,H-7′), 2.11 (4H, m, H-12, H-12′), 1.58 (2H, p, *J* =7.2, H-8), 1.50 (2H, *f* = 7.0 p, H-8′), 1.43 (4H, m, H-11, H-11′), 1.24 (8H, m, H-9, H-8, H-9′, H-8′) $^{13}$C NMR (DMSO-*d6,* 150 MHz) δ 174.6 (C, C-13), 174.6 (C, C-13′), 148.9 (CH, C-1), 142.0 (CH, C-1′), 137.8 (CH, C-6′), 134.7 (CH, C-6), 130.9 (CH, C-5), 130.2 (CH, C-4), 130.0 (CH, C-5′), 128.5 (CH, C-4′), 127.0 (CH, C-3), 126.2 (CH, C-3′), 123.3 (CH, C-2), 121.1 (CH, C-2′), 34.0 (CH2, C-12), 34.0 (CH2, C-12′), 31.2 (CH2, C-7′), 30.6 (CH2, C-7), 30.5 (CH2, C-8′), 30.2 (CH2, C-8), 28.7 (CH2, C-8′), 28.6 (CH2, C-8), 28.4 (CH2, C-9), 28.4 (CH2, C-9′), 24.6 (CH2, C-11), 24.6 (CH2, C-11′), 15N NMR (DMSO-d6, 600

MHz) δ 333.2, 346.8, HRESIMS *m/z* 455.2560 (calcd. for $C_{26}H_{35}N_2O_5$, 455.2540).
SMILES: OC(CCCCCCC1=CC=CC=C1/[N+]([O-
])=N\C2=CC=CC=C2CCCCCCC(O)=O)=O

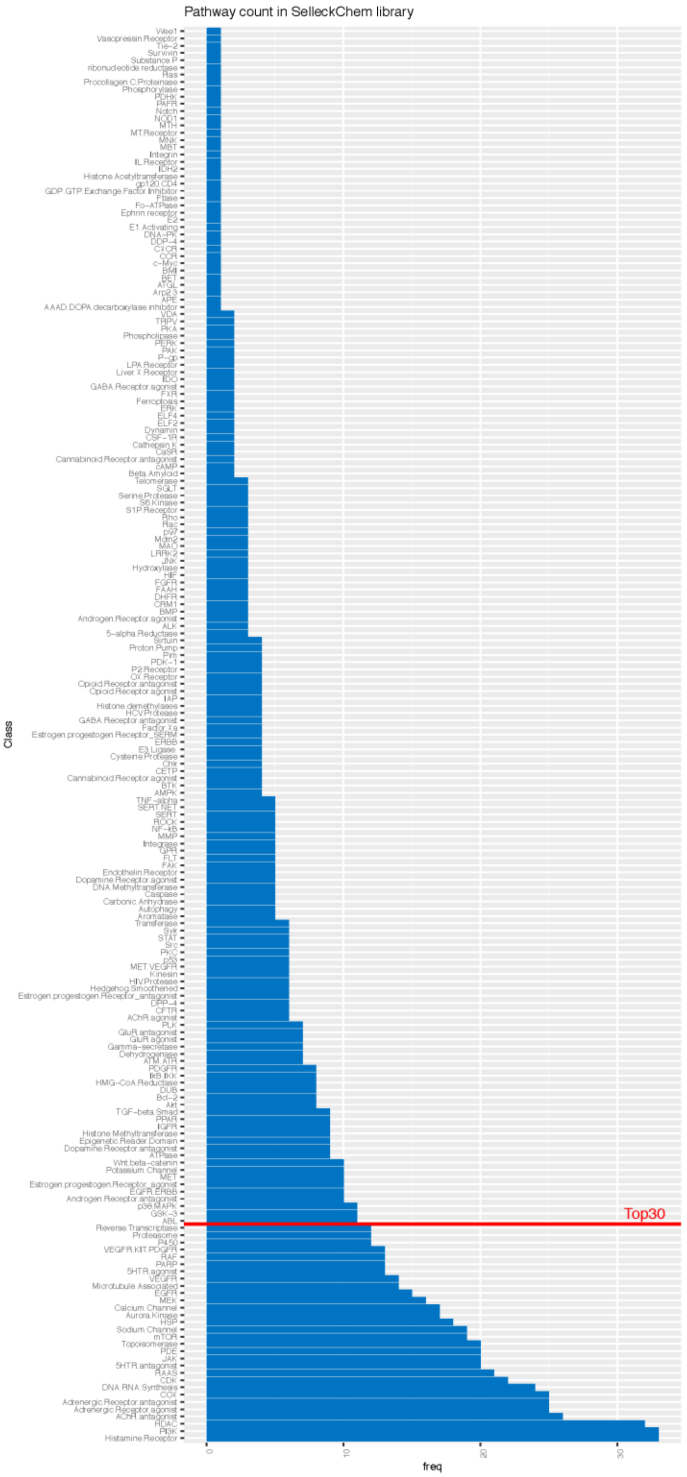

**Supplementary Figure 1: Selleck library target class membership.**
Annotated targets in the Selleck library are listed in order of the number of chemicals that are assigned to that class. The top 30 largest target classes were selected for comparison by k-means clustering in Figure 1.

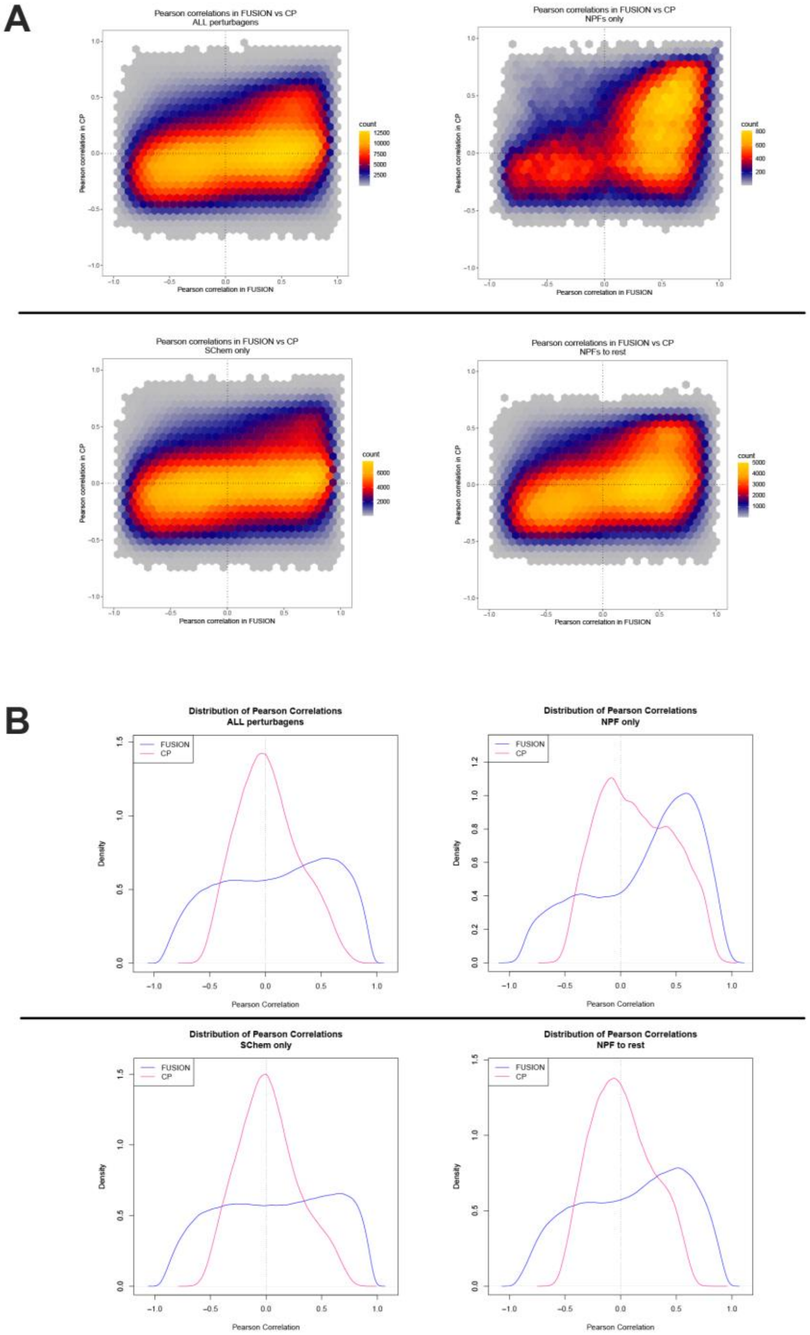

**Supplementary Figure 2: Pearson correlation distributions in FUSION and CP.**
A) Hexplots and B) density plots illustrating the trends in pearson correlations between indicated perturbagen types.

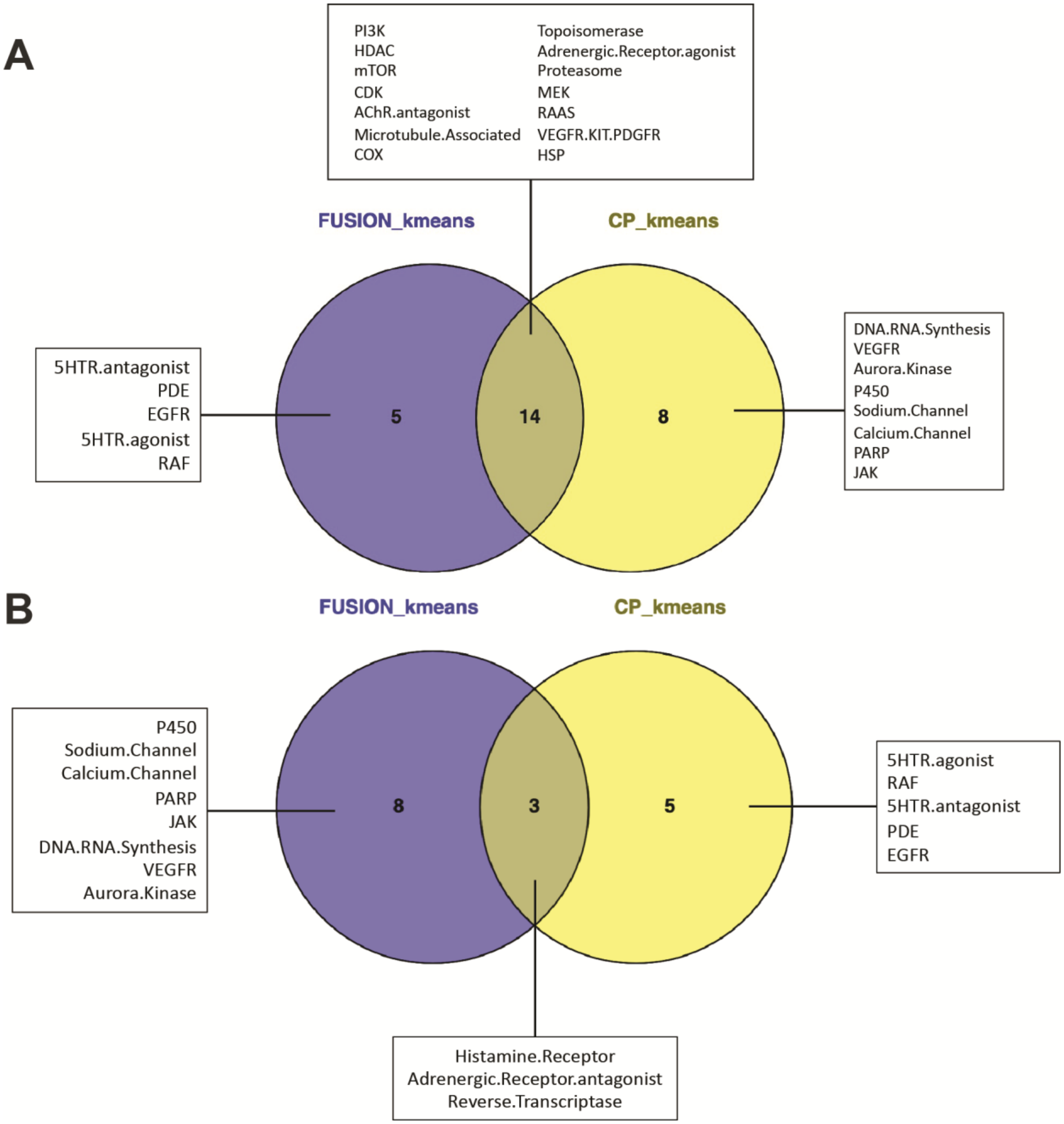

**Supplementary Figure 3: Venn diagrams illustrating overlap among target classes identified as significantly enriched in a k-means cluster.**

A) Overlap among target classes that were identified as significantly enriched in at least one kmeans (k=30) cluster by hypergeometric test with Bonferonni correction. B) Overlap among target classes that were not identified as significantly enriched by the same method.

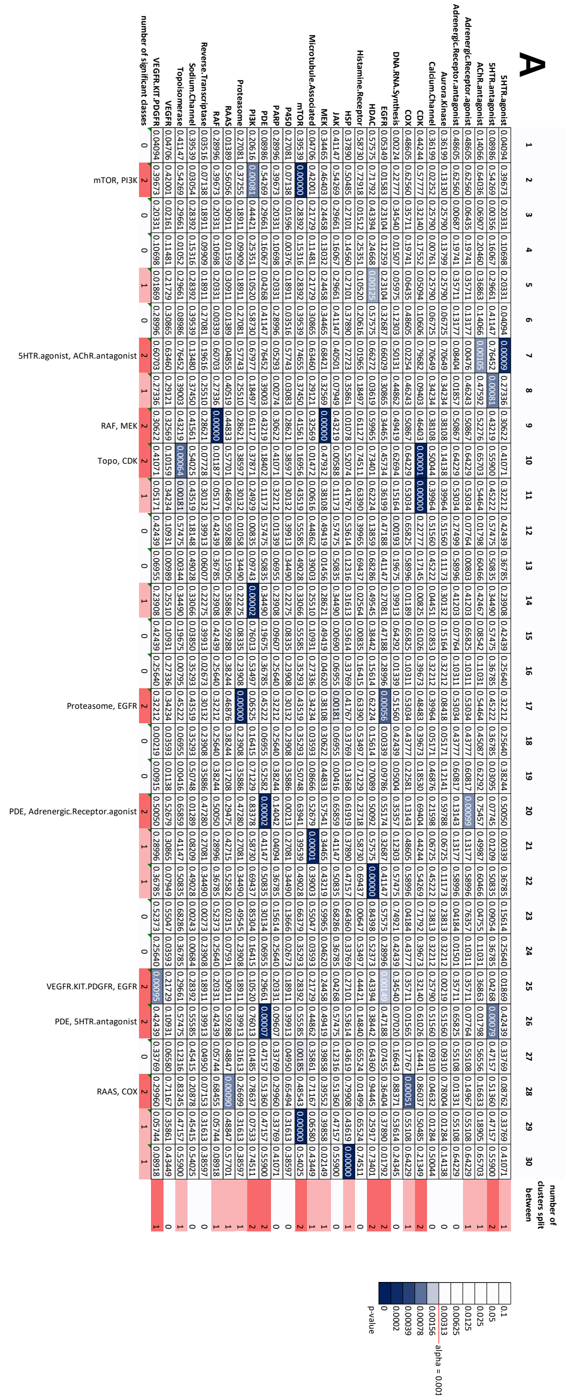

**Supplementary Figure 4: Annotated heatmaps showing which target classes are significantly enriched**

as assessed hypergeometric test (Bonferonni-corrected alpha = 0.0016667) in kmeans clusters (k=30) in A) FUSION and B) CP datasets. Lines labeled "number of significant classes" contain a count of the number of target classes found to be significantly enriched in that cluster ($p<0.00167$). Columns labeled "number of clusters split between" contain a count of the number of clusters that a target class was significantly enriched in.

**B**

| | 1 | 2 | 3 | 4 | 5 | 6 | 7 | 8 | 9 | 10 | 11 | 12 | 13 | 14 | 15 | 16 | 17 | 18 | 19 | 20 | 21 | 22 | 23 | 24 | 25 | 26 | 27 | 28 | 29 | 30 | number of clusters split between |
|---|---|---|---|---|---|---|---|---|---|---|---|---|---|---|---|---|---|---|---|---|---|---|---|---|---|---|---|---|---|---|---|
| **5HTR.agonist** | 0.28996 | 0.41071 | 0.00584 | 0.36785 | 0.32212 | 0.22138 | 0.27336 | 0.06339 | 0.30622 | 0.02673 | 0.48850 | 0.07591 | 0.16599 | 0.06955 | 0.54596 | 0.28996 | 0.23908 | 0.03119 | 0.48850 | 0.45087 | 0.30622 | 0.01046 | 0.23908 | 0.22138 | 0.05171 | 0.01036 | 0.35293 | 0.54596 | 0.30622 | 0.39673 | 0 |
| **5HTR.antagonist** | 0.08986 | 0.55900 | 0.47157 | 0.50835 | 0.11173 | 0.32116 | 0.39003 | 0.49028 | 0.43219 | 0.34490 | 0.07126 | 0.03095 | 0.02768 | 0.14685 | 0.70549 | 0.01209 | 0.34490 | 0.06955 | 0.64581 | 0.00928 | 0.43219 | 0.17145 | 0.34490 | 0.05110 | 0.45222 | 0.43006 | 0.13488 | 0.70549 | 0.43219 | 0.03505 | 0 |
| **AChR.antagonist** | 0.02510 | 0.65703 | 0.18905 | 0.05417 | 0.54464 | 0.39728 | 0.47592 | 0.04744 | 0.15647 | 0.42467 | 0.37544 | 0.62292 | 0.30735 | 0.05417 | 0.79768 | 0.49987 | 0.09592 | 0.01675 | 0.74248 | 0.32492 | 0.52276 | 0.25657 | 0.42467 | 0.08214 | 0.00051 | 0.10885 | 0.58555 | 0.79768 | 0.52276 | 0.06896 | 1 |
| **Adrenergic.Receptor.agonist** | 0.00262 | 0.64229 | 0.03713 | 0.20959 | 0.53034 | 0.38516 | 0.46243 | 0.04285 | 0.50867 | 0.41203 | 0.00564 | 0.60817 | 0.29728 | 0.58996 | 0.78455 | 0.13177 | 0.41203 | 0.43777 | 0.72837 | 0.30788 | 0.50867 | 0.24214 | 0.08956 | 0.38516 | 0.00003 | 0.26578 | 0.57094 | 0.78455 | 0.50867 | 0.06252 | 1 |
| **Adrenergic.Receptor.antagonist** | 0.13177 | 0.25854 | 0.17767 | 0.58996 | 0.16206 | 0.38516 | 0.46243 | 0.19353 | 0.50867 | 0.08956 | 0.12167 | 0.22581 | 0.29728 | 0.58996 | 0.78455 | 0.13177 | 0.41203 | 0.10311 | 0.72837 | 0.68816 | 0.50867 | 0.24214 | 0.08956 | 0.00917 | 0.53034 | 0.00614 | 0.04285 | 0.78455 | 0.14674 | 0.06252 | 0 |
| **Aurora.Kinase** | 0.36199 | 0.02543 | 0.41767 | 0.45222 | 0.39964 | 0.00000 | 0.34234 | 0.43519 | 0.38108 | 0.30132 | 0.58515 | 0.46876 | 0.21195 | 0.45222 | 0.01338 | 0.36199 | 0.30132 | 0.32212 | 0.58515 | 0.54464 | 0.00073 | 0.48483 | 0.30132 | 0.27992 | 0.39964 | 0.72460 | 0.10229 | 0.01338 | 0.38108 | 0.48483 | 2 |
| **Calcium.Channel** | 0.36199 | 0.50044 | 0.09310 | 0.01729 | 0.01090 | 0.27992 | 0.34234 | 0.43519 | 0.38108 | 0.30132 | 0.58515 | 0.01981 | 0.21195 | 0.11173 | 0.64521 | 0.00756 | 0.30132 | 0.32212 | 0.58515 | 0.17262 | 0.38108 | 0.48483 | 0.04451 | 0.27992 | 0.39964 | 0.35005 | 0.10229 | 0.64521 | 0.00073 | 0.48483 | 1 |
| **CDK** | 0.44244 | 0.59432 | 0.50485 | 0.54269 | 0.02252 | 0.06086 | 0.42001 | 0.15781 | 0.46403 | 0.37255 | 0.68139 | 0.56056 | 0.26628 | 0.54269 | 0.74001 | 0.44244 | 0.37255 | 0.39673 | 0.00311 | 0.25657 | 0.01901 | 0.57777 | 0.37255 | 0.34745 | 0.48483 | 0.48062 | 0.52413 | 0.00000 | 0.46403 | 0.57777 | 1 |
| **COX** | 0.48605 | 0.25854 | 0.03713 | 0.20959 | 0.53034 | 0.38516 | 0.46243 | 0.57094 | 0.50867 | 0.08956 | 0.35679 | 0.60817 | 0.04221 | 0.00803 | 0.78455 | 0.13177 | 0.41203 | 0.43777 | 0.72837 | 0.02076 | 0.50867 | 0.24214 | 0.41203 | 0.38516 | 0.53034 | 0.00016 | 0.57094 | 0.78455 | 0.50867 | 0.01143 | 1 |
| **DNA.RNA.Synthesis** | 0.47188 | 0.62694 | 0.53614 | 0.57475 | 0.51560 | 0.37282 | 0.00000 | 0.18148 | 0.49419 | 0.08335 | 0.11069 | 0.59288 | 0.28708 | 0.04408 | 0.77060 | 0.47188 | 0.39913 | 0.42439 | 0.11069 | 0.29079 | 0.49419 | 0.22777 | 0.39913 | 0.07122 | 0.15164 | 0.83995 | 0.55585 | 0.00158 | 0.49419 | 0.22777 | 2 |
| **EGFR** | 0.32687 | 0.01792 | 0.07455 | 0.08986 | 0.06725 | 0.02974 | 0.30865 | 0.39539 | 0.00633 | 0.03516 | 0.53926 | 0.42715 | 0.01584 | 0.41147 | 0.21586 | 0.32687 | 0.00265 | 0.04094 | 0.53926 | 0.49987 | 0.34465 | 0.44244 | 0.27081 | 0.25117 | 0.36199 | 0.67882 | 0.08209 | 0.59855 | 0.34465 | 0.44244 | 0 |
| **HDAC** | 0.57575 | 0.73401 | 0.64360 | 0.68286 | 0.06041 | 0.46555 | 0.55047 | 0.66379 | 0.00856 | 0.49545 | 0.81343 | 0.70089 | 0.36519 | 0.68286 | 0.00000 | 0.57575 | 0.49545 | 0.52373 | 0.81343 | 0.16538 | 0.59965 | 0.34323 | 0.49545 | 0.46555 | 0.23813 | 0.91467 | 0.66379 | 0.03202 | 0.05165 | 0.11355 | 1 |
| **Histamine.Receptor** | 0.04718 | 0.74511 | 0.07533 | 0.31450 | 0.06525 | 0.47622 | 0.56189 | 0.67538 | 0.61127 | 0.50644 | 0.07386 | 0.02571 | 0.07025 | 0.31450 | 0.87014 | 0.20616 | 0.50644 | 0.16415 | 0.82325 | 0.44059 | 0.61127 | 0.12189 | 0.50644 | 0.12413 | 0.01203 | 0.00629 | 0.29279 | 0.87014 | 0.22764 | 0.35751 | 0 |
| **HSP** | 0.37890 | 0.52074 | 0.43619 | 0.47157 | 0.41767 | 0.29391 | 0.35861 | 0.45415 | 0.39858 | 0.31613 | 0.60640 | 0.48847 | 0.22309 | 0.47157 | 0.00000 | 0.07455 | 0.31613 | 0.33769 | 0.05455 | 0.56556 | 0.39858 | 0.50485 | 0.31613 | 0.29391 | 0.41767 | 0.74504 | 0.45415 | 0.66653 | 0.39858 | 0.50485 | 1 |
| **JAK** | 0.08986 | 0.00064 | 0.47157 | 0.50835 | 0.11173 | 0.00479 | 0.39003 | 0.00282 | 0.43219 | 0.34490 | 0.64581 | 0.52582 | 0.24496 | 0.50835 | 0.70549 | 0.41147 | 0.34490 | 0.36785 | 0.07126 | 0.60466 | 0.01456 | 0.54269 | 0.06007 | 0.32116 | 0.45222 | 0.78156 | 0.02357 | 0.10453 | 0.43219 | 0.54269 | 1 |
| **MEK** | 0.34465 | 0.12774 | 0.39858 | 0.43219 | 0.07556 | 0.26567 | 0.32569 | 0.41561 | 0.36311 | 0.00001 | 0.56279 | 0.10946 | 0.00088 | 0.10062 | 0.62258 | 0.34465 | 0.28621 | 0.30622 | 0.56279 | 0.15647 | 0.36311 | 0.46403 | 0.00017 | 0.26567 | 0.38108 | 0.70257 | 0.41561 | 0.62258 | 0.00765 | 0.46403 | 3 |
| **Microtubule.Associated** | 0.30865 | 0.00000 | 0.35861 | 0.39003 | 0.00616 | 0.23641 | 0.29121 | 0.07253 | 0.32569 | 0.25510 | 0.51452 | 0.40519 | 0.17770 | 0.39003 | 0.57304 | 0.30865 | 0.25510 | 0.27336 | 0.51452 | 0.47592 | 0.32569 | 0.42001 | 0.25510 | 0.00163 | 0.34234 | 0.65322 | 0.00853 | 0.57304 | 0.32569 | 0.09403 | 2 |
| **mTOR** | 0.39539 | 0.16956 | 0.11288 | 0.49028 | 0.43519 | 0.30766 | 0.37450 | 0.47251 | 0.09202 | 0.33066 | 0.62661 | 0.02693 | 0.23409 | 0.49028 | 0.68659 | 0.00088 | 0.00034 | 0.35293 | 0.62661 | 0.04744 | 0.41561 | 0.52413 | 0.00538 | 0.30766 | 0.43519 | 0.76399 | 0.02046 | 0.68659 | 0.01259 | 0.52413 | 2 |
| **P450** | 0.27081 | 0.38597 | 0.31613 | 0.06007 | 0.30132 | 0.20609 | 0.25510 | 0.33066 | 0.28621 | 0.02288 | 0.00186 | 0.35886 | 0.15414 | 0.34490 | 0.51721 | 0.27081 | 0.22275 | 0.23908 | 0.46113 | 0.42467 | 0.03973 | 0.00062 | 0.22275 | 0.20609 | 0.30132 | 0.59591 | 0.33066 | 0.51721 | 0.00322 | 0.07138 | 1 |
| **PARP** | 0.28996 | 0.08918 | 0.33769 | 0.36785 | 0.32212 | 0.22138 | 0.00013 | 0.35293 | 0.04620 | 0.02673 | 0.48850 | 0.07591 | 0.16599 | 0.06955 | 0.54596 | 0.28996 | 0.23908 | 0.25640 | 0.48850 | 0.01675 | 0.30622 | 0.39673 | 0.23908 | 0.22138 | 0.05171 | 0.62564 | 0.35293 | 0.03470 | 0.30622 | 0.39673 | 1 |
| **PDE** | 0.41147 | 0.18402 | 0.12316 | 0.50835 | 0.45222 | 0.32116 | 0.39003 | 0.02357 | 0.10062 | 0.34490 | 0.07126 | 0.52582 | 0.24496 | 0.00344 | 0.70549 | 0.08986 | 0.06007 | 0.06955 | 0.64581 | 0.22255 | 0.43219 | 0.00497 | 0.06007 | 0.05110 | 0.45222 | 0.78156 | 0.49028 | 0.32762 | 0.43219 | 0.54269 | 0 |
| **PI3K** | 0.20616 | 0.74511 | 0.07533 | 0.02168 | 0.24929 | 0.12413 | 0.18497 | 0.67538 | 0.22764 | 0.14383 | 0.82325 | 0.10938 | 0.07025 | 0.69437 | 0.87014 | 0.20616 | 0.14383 | 0.03206 | 0.49922 | 0.05247 | 0.22764 | 0.35751 | 0.50644 | 0.47622 | 0.63390 | 0.70653 | 0.00033 | 0.58830 | 0.61127 | 0.12189 | 1 |
| **Proteasome** | 0.27081 | 0.38597 | 0.31613 | 0.34490 | 0.30132 | 0.20609 | 0.25510 | 0.33066 | 0.03973 | 0.22275 | 0.46113 | 0.06564 | 0.15414 | 0.34490 | 0.51721 | 0.27081 | 0.22275 | 0.23908 | 0.00000 | 0.42467 | 0.28621 | 0.37255 | 0.22275 | 0.20609 | 0.04451 | 0.21165 | 0.33066 | 0.51721 | 0.28621 | 0.37255 | 1 |
| **RAAS** | 0.01389 | 0.57701 | 0.13368 | 0.15905 | 0.46876 | 0.33443 | 0.40519 | 0.50748 | 0.44833 | 0.35886 | 0.00249 | 0.03527 | 0.25569 | 0.52582 | 0.72327 | 0.42715 | 0.35886 | 0.07591 | 0.66405 | 0.62292 | 0.44833 | 0.00598 | 0.35886 | 0.33443 | 0.46876 | 0.00033 | 0.50748 | 0.34977 | 0.44833 | 0.56056 | 1 |
| **RAF** | 0.28996 | 0.41071 | 0.05744 | 0.00795 | 0.32212 | 0.22138 | 0.00274 | 0.35293 | 0.04620 | 0.23908 | 0.48850 | 0.07591 | 0.16599 | 0.36785 | 0.54596 | 0.04094 | 0.23908 | 0.03119 | 0.48850 | 0.45087 | 0.00411 | 0.39673 | 0.23908 | 0.02256 | 0.32212 | 0.62564 | 0.35293 | 0.54596 | 0.04620 | 0.39673 | 0 |
| **Reverse.Transcriptase** | 0.27081 | 0.07728 | 0.31613 | 0.34490 | 0.30132 | 0.20609 | 0.00214 | 0.05469 | 0.03973 | 0.22275 | 0.46113 | 0.06564 | 0.01016 | 0.06007 | 0.51721 | 0.27081 | 0.22275 | 0.23908 | 0.11577 | 0.42467 | 0.28621 | 0.37255 | 0.22275 | 0.20609 | 0.30132 | 0.00757 | 0.33066 | 0.51721 | 0.28621 | 0.37255 | 0 |
| **Sodium.Channel** | 0.08209 | 0.54025 | 0.45415 | 0.13488 | 0.43519 | 0.30766 | 0.37450 | 0.47251 | 0.09202 | 0.33066 | 0.00153 | 0.14624 | 0.23409 | 0.00282 | 0.68659 | 0.39539 | 0.33066 | 0.35293 | 0.62661 | 0.20571 | 0.01259 | 0.15781 | 0.05469 | 0.04647 | 0.43519 | 0.76399 | 0.47251 | 0.68659 | 0.41561 | 0.03054 | 1 |
| **Topoisomerase** | 0.41147 | 0.55900 | 0.47157 | 0.14685 | 0.45222 | 0.32116 | 0.39003 | 0.49028 | 0.01456 | 0.34490 | 0.26182 | 0.52582 | 0.24496 | 0.50835 | 0.70549 | 0.41147 | 0.34490 | 0.36785 | 0.00000 | 0.22255 | 0.43219 | 0.54269 | 0.34490 | 0.32116 | 0.45222 | 0.16828 | 0.13488 | 0.02402 | 0.43219 | 0.17145 | 1 |
| **VEGFR** | 0.04706 | 0.10159 | 0.35861 | 0.00989 | 0.34234 | 0.23641 | 0.29121 | 0.00066 | 0.32569 | 0.25510 | 0.51452 | 0.40519 | 0.17770 | 0.39003 | 0.57304 | 0.30865 | 0.00009 | 0.27336 | 0.15019 | 0.12525 | 0.32569 | 0.42001 | 0.03083 | 0.23641 | 0.34234 | 0.65322 | 0.37450 | 0.57304 | 0.05304 | 0.42001 | 2 |
| **VEGFR.KIT.PDGFR** | 0.28996 | 0.08918 | 0.33769 | 0.06955 | 0.05171 | 0.02256 | 0.27336 | 0.06339 | 0.30622 | 0.23908 | 0.48850 | 0.38244 | 0.16599 | 0.36785 | 0.54596 | 0.28996 | 0.02673 | 0.03119 | 0.48850 | 0.45087 | 0.00023 | 0.39673 | 0.23908 | 0.22138 | 0.32212 | 0.62564 | 0.35293 | 0.03470 | 0.04620 | 0.39673 | 1 |
| **number of significant classes** | 0 | 2 | 0 | 0 | 0 | 1 | 2 | 1 | 0 | 1 | 1 | 0 | 1 | 0 | 2 | 1 | 2 | 0 | 2 | 0 | 2 | 1 | 1 | 1 | 2 | 2 | 1 | 2 | 1 | 0 | |
| | | Microtubule, JAK | | | | | PARP, DNA.RNA.Synthesis | | | | | | | | HSP, HDAC | | VEGFR, mTOR | | Topo, proteasome | | VEGFR.KIT.PDGFR, Aurk | | | | AChR.antagonist, Adrenergic.Receptor.antagonist | RAAS, COX | | DNA.RNA.Synthesis, CDK | | | |

0.1
0.05
0.025
0.0125
0.00625
0.00313
0.00156 — alpha = 0.001
0.00078
0.00039
0.0002
0
p-value

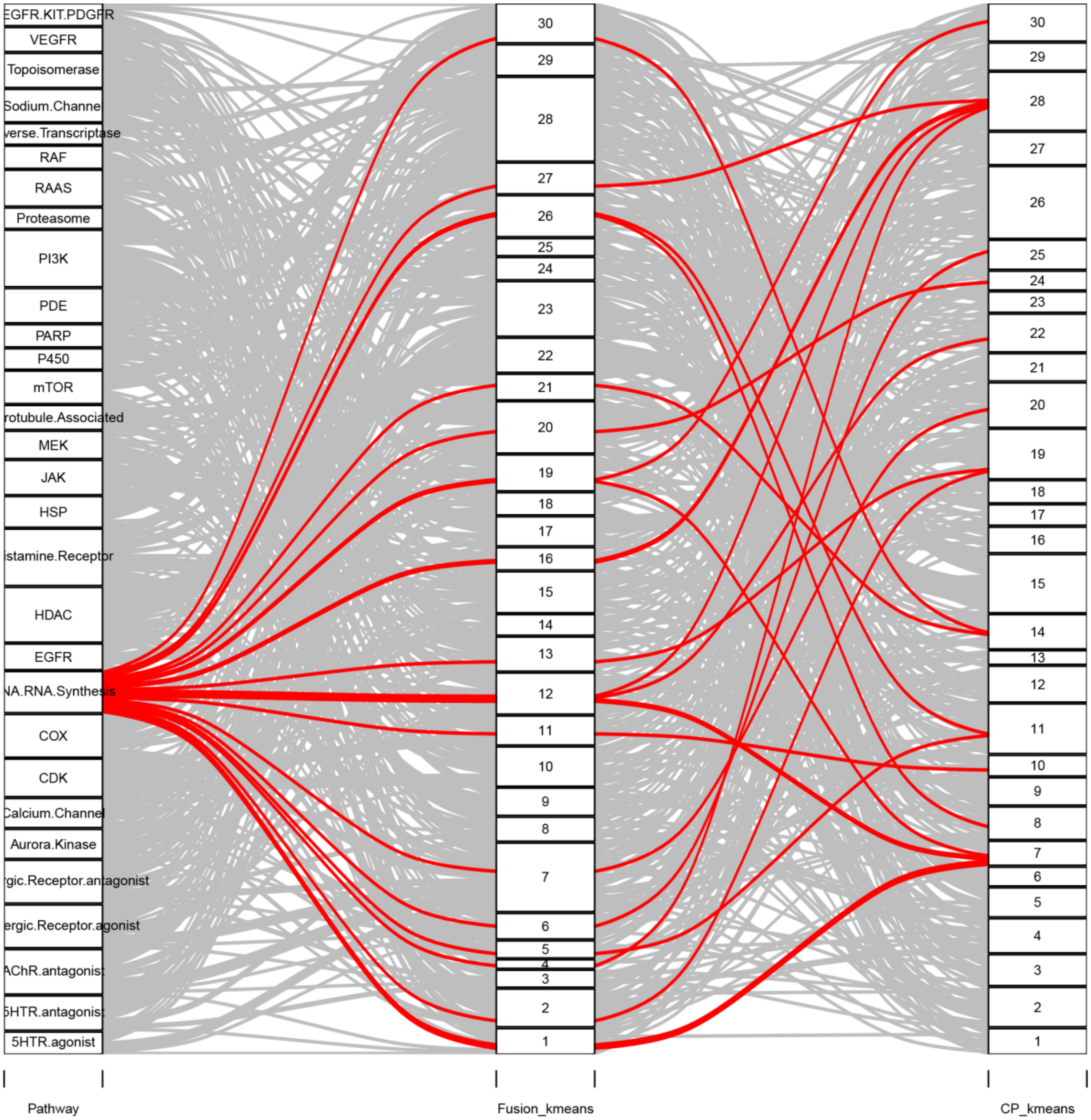

**Supplementary Figure 5. Alluvial diagrams for each target class using k-means clustering of the FUSION and CP datasets.**

Comparison of k-means clustering (k=30) of FUSION and CP signatures for the top 30 target classes in the Selleck chemical library. Each line represents a compound, and each panel displays a separate target class with chemicals belonging to that class highlighted in red.

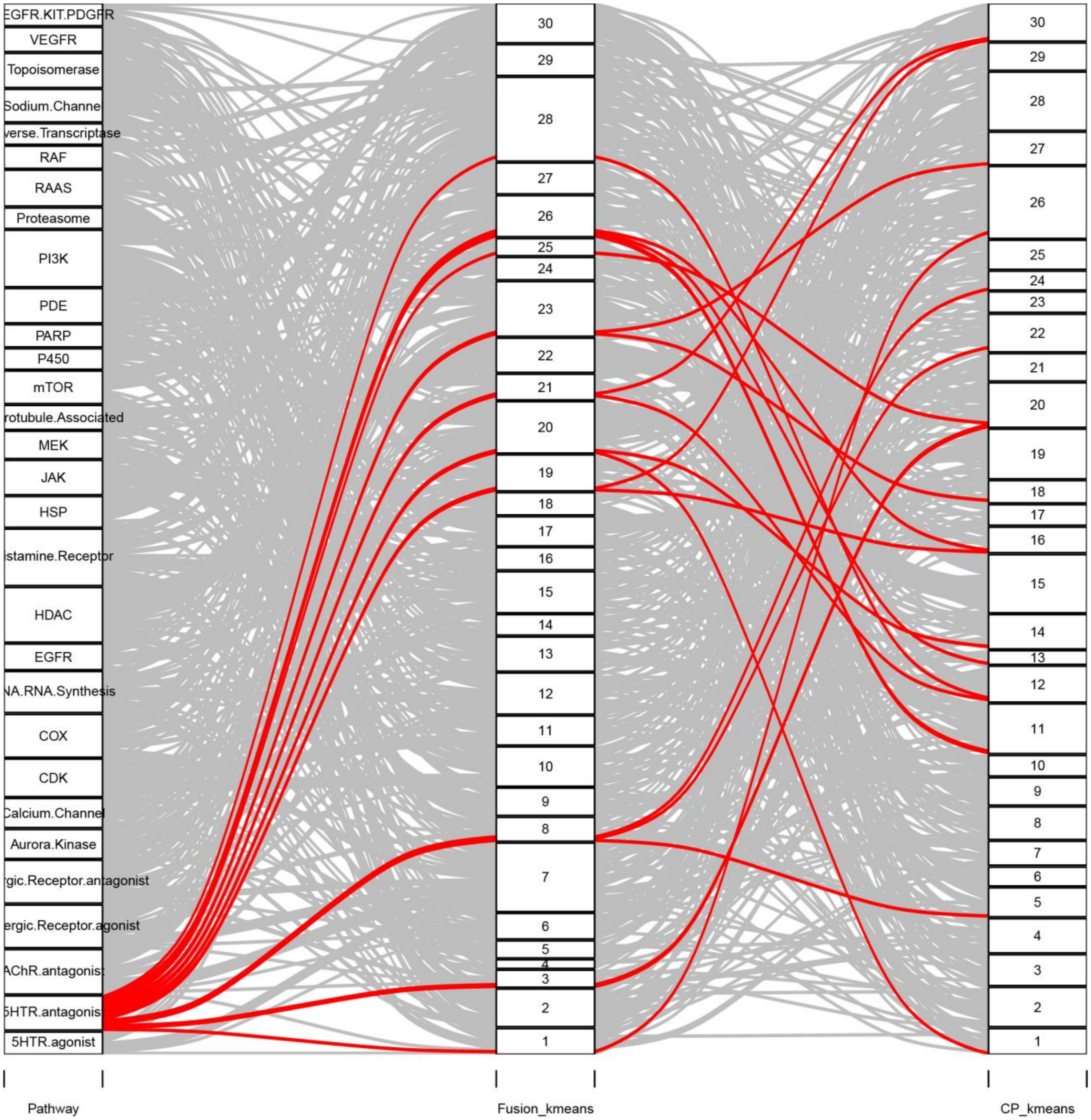
EGFR.KIT.PDGFR
VEGFR
Topoisomerase
Sodium.Channel
verse.Transcriptase
RAF
RAAS
Proteasome
PI3K
PDE
PARP
P450
mTOR
rotubule.Associated
MEK
JAK
HSP
istamine.Receptor
HDAC
EGFR
NA.RNA.Synthesis
COX
CDK
Calcium.Channel
Aurora.Kinase
rgic.Receptor.antagonist
ergic.Receptor.agonist
AChR.antagonist
5HTR.antagonist
5HTR.agonist
Pathway
Fusion_kmeans
CP_kmeans

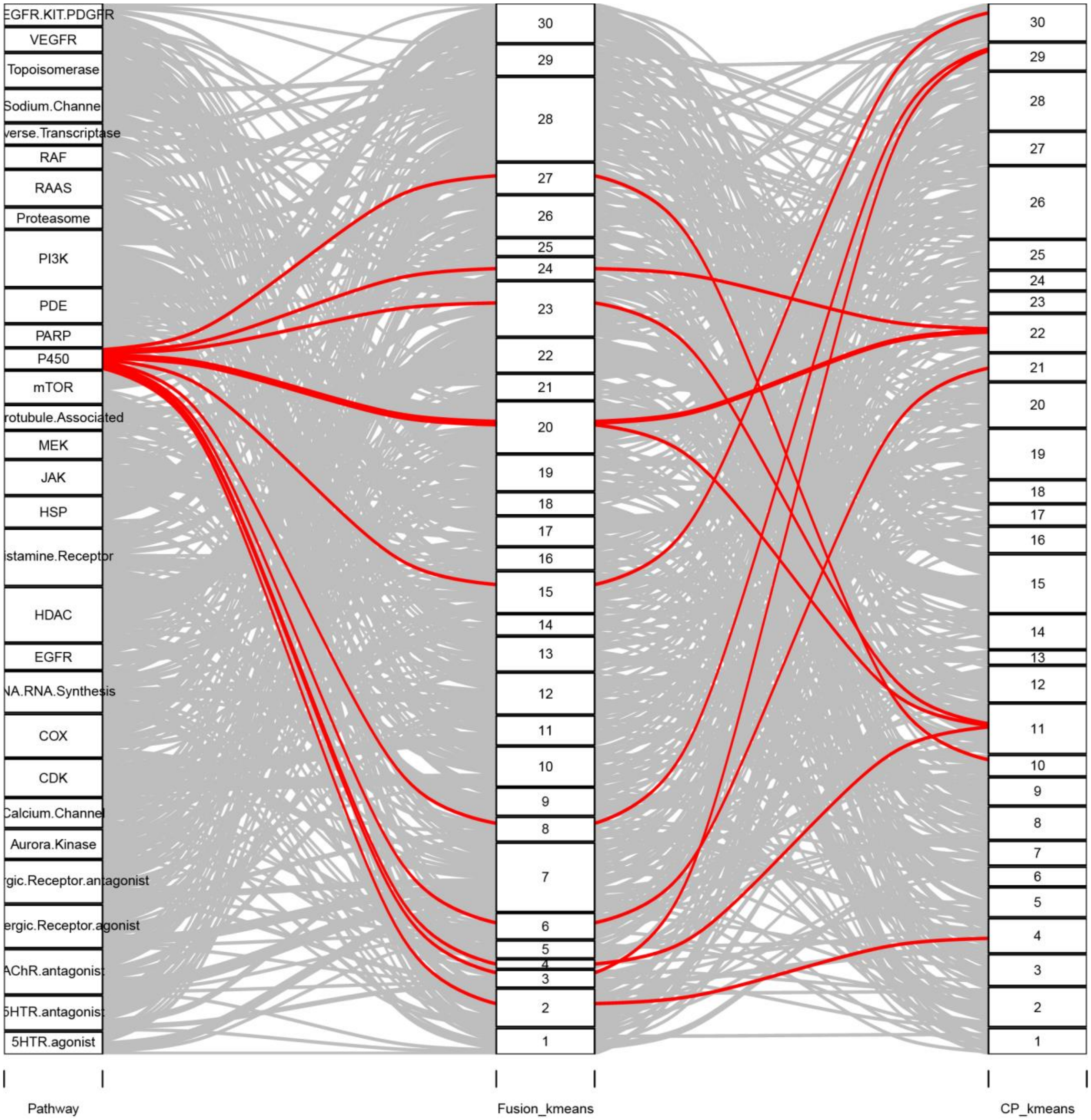
EGFR.KIT.PDGFR
VEGFR
Topoisomerase
Sodium.Channel
verse.Transcriptase
RAF
RAAS
Proteasome
PI3K
PDE
PARP
P450
mTOR
rotubule.Associated
MEK
JAK
HSP
istamine.Receptor
HDAC
EGFR
NA.RNA.Synthesis
COX
CDK
Calcium.Channel
Aurora.Kinase
rgic.Receptor.antagonist
ergic.Receptor.agonist
AChR.antagonist
5HTR.antagonist
5HTR.agonist
Pathway
Fusion_kmeans
CP_kmeans

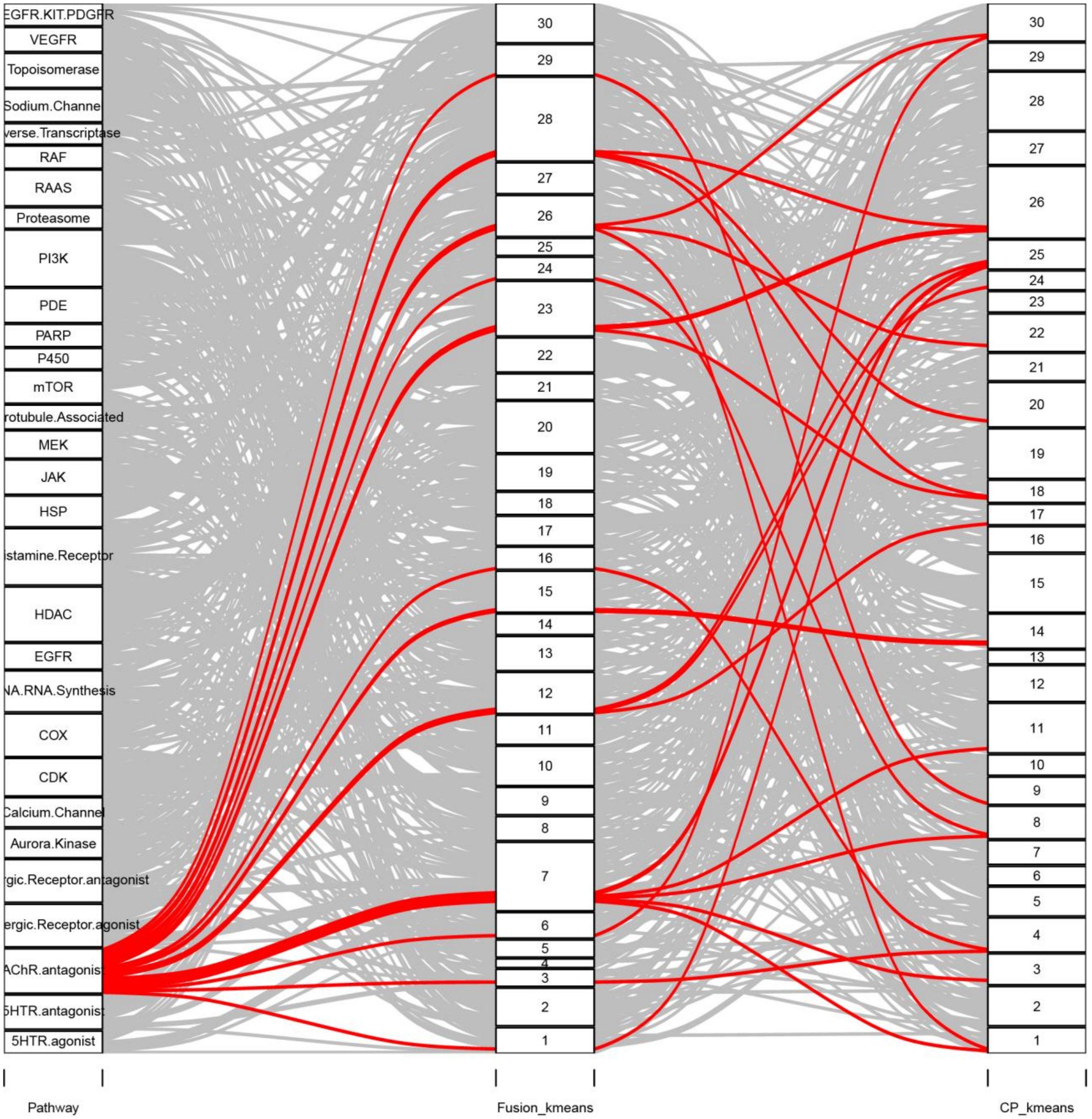
EGFR.KIT.PDGFR
VEGFR
Topoisomerase
Sodium.Channel
verse.Transcriptase
RAF
RAAS
Proteasome
PI3K
PDE
PARP
P450
mTOR
rotubule.Associated
MEK
JAK
HSP
istamine.Receptor
HDAC
EGFR
NA.RNA.Synthesis
COX
CDK
Calcium.Channel
Aurora.Kinase
rgic.Receptor.antagonist
ergic.Receptor.agonist
AChR.antagonist
5HTR.antagonist
5HTR.agonist
Pathway
Fusion_kmeans
CP_kmeans

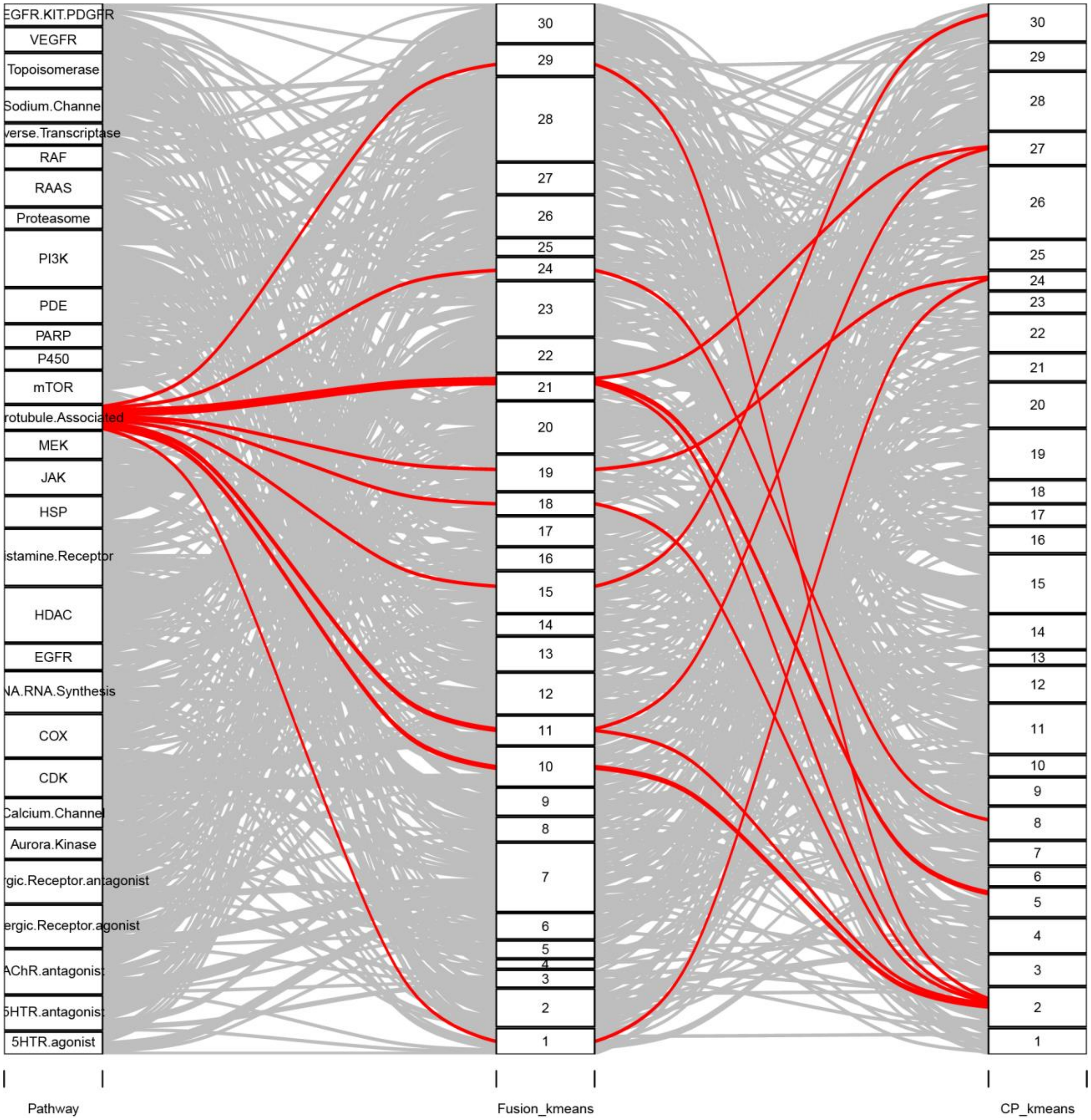
EGFR.KIT.PDGFR
VEGFR
Topoisomerase
Sodium.Channel
verse.Transcriptase
RAF
RAAS
Proteasome
PI3K
PDE
PARP
P450
mTOR
rotubule.Associated
MEK
JAK
HSP
istamine.Receptor
HDAC
EGFR
NA.RNA.Synthesis
COX
CDK
Calcium.Channel
Aurora.Kinase
rgic.Receptor.antagonist
ergic.Receptor.agonist
AChR.antagonist
5HTR.antagonist
5HTR.agonist
Pathway
Fusion_kmeans
CP_kmeans

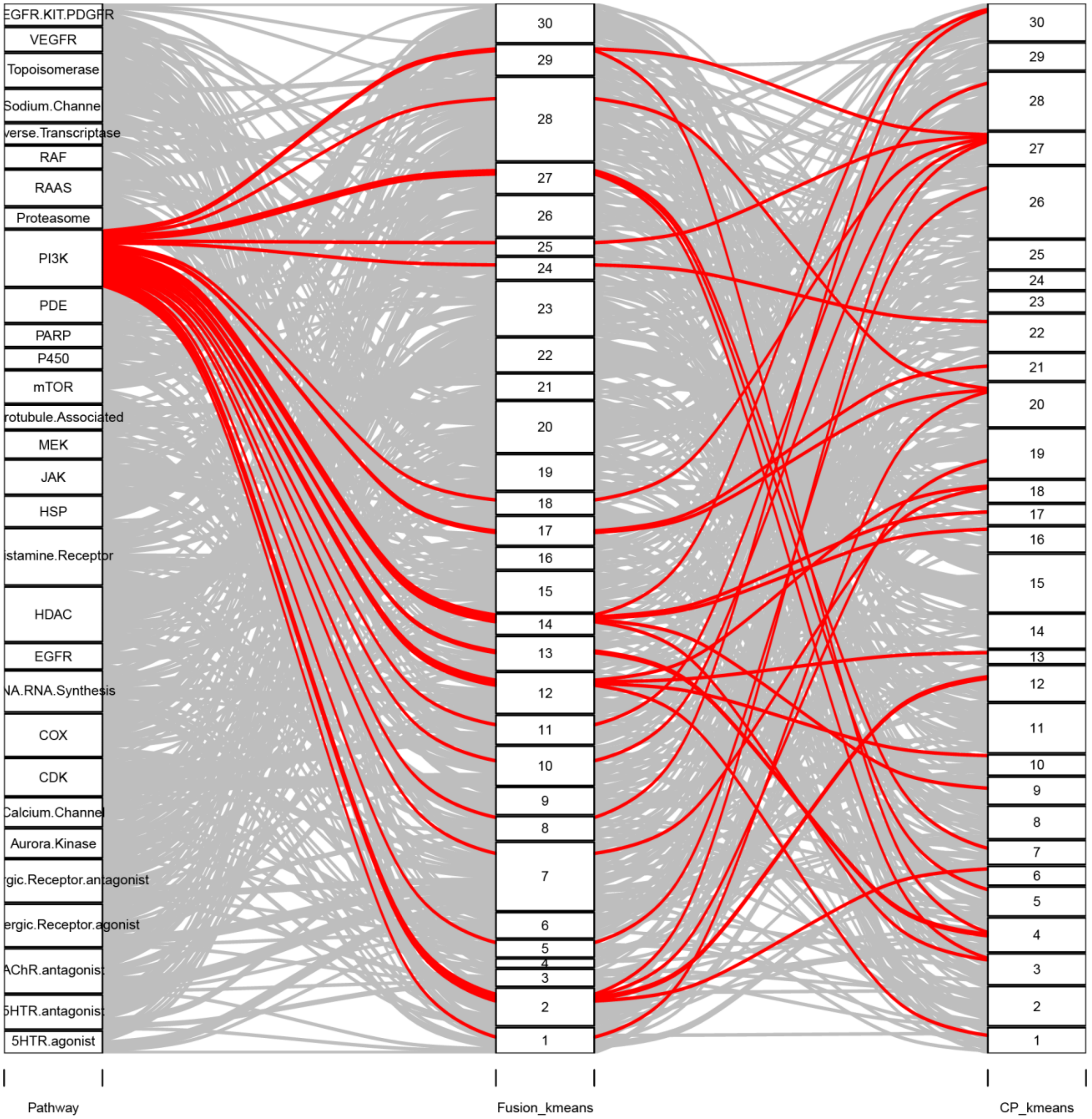
EGFR.KIT.PDGFR
VEGFR
Topoisomerase
Sodium.Channel
verse.Transcriptase
RAF
RAAS
Proteasome
PI3K
PDE
PARP
P450
mTOR
rotubule.Associated
MEK
JAK
HSP
istamine.Receptor
HDAC
EGFR
NA.RNA.Synthesis
COX
CDK
Calcium.Channel
Aurora.Kinase
rgic.Receptor.antagonist
ergic.Receptor.agonist
AChR.antagonist
5HTR.antagonist
5HTR.agonist
30
29
28
27
26
25
24
23
22
21
20
19
18
17
16
15
14
13
12
11
10
9
8
7
6
5
4
3
2
1
Pathway
Fusion_kmeans
CP_kmeans

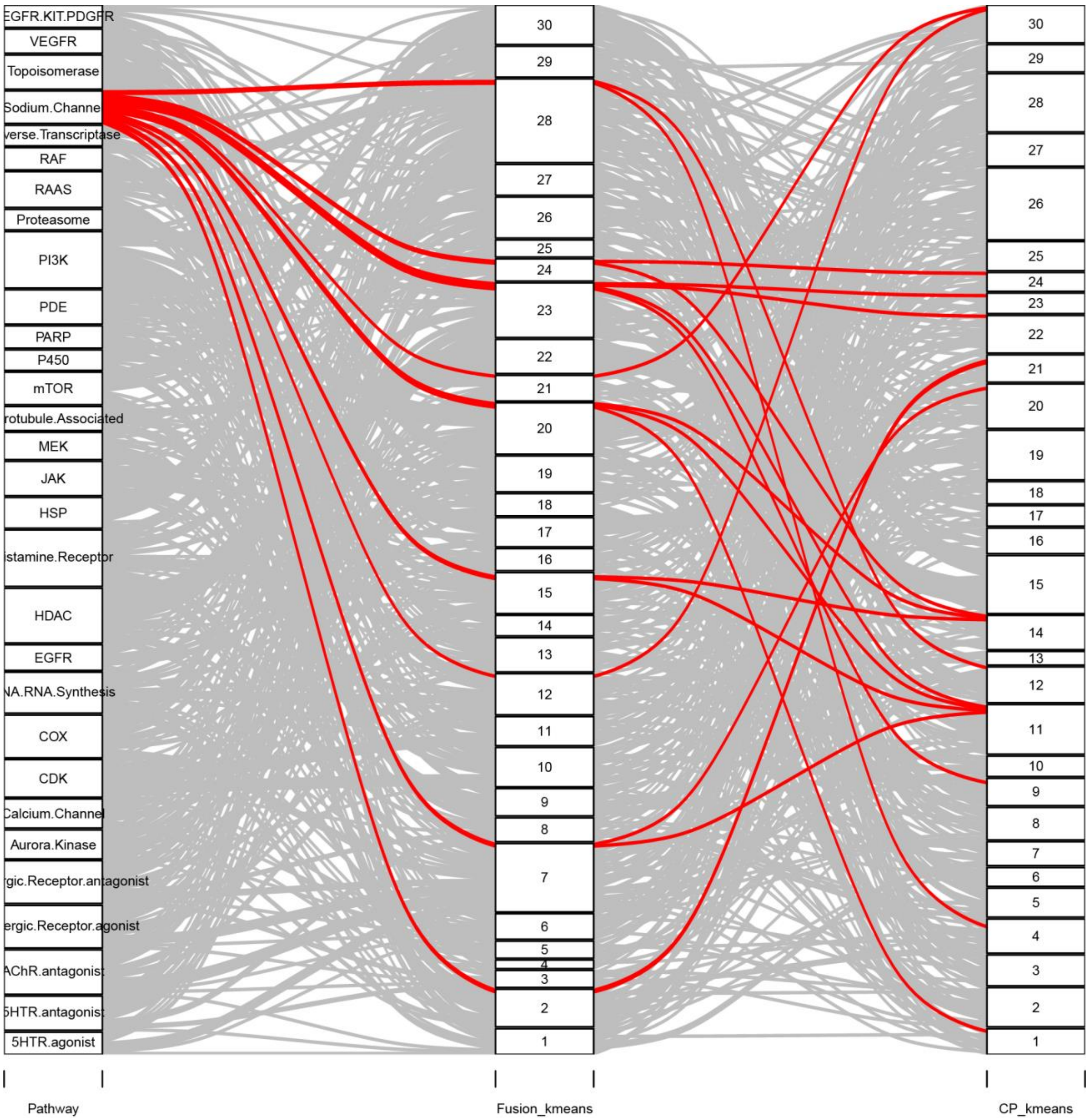

EGFR.KIT.PDGFR
VEGFR
Topoisomerase
Sodium.Channel
verse.Transcriptase
RAF
RAAS
Proteasome
PI3K
PDE
PARP
P450
mTOR
rotubule.Associated
MEK
JAK
HSP
istamine.Receptor
HDAC
EGFR
NA.RNA.Synthesis
COX
CDK
Calcium.Channel
Aurora.Kinase
rgic.Receptor.antagonist
ergic.Receptor.agonist
AChR.antagonist
5HTR.antagonist
5HTR.agonist
30
29
28
27
26
25
24
23
22
21
20
19
18
17
16
15
14
13
12
11
10
9
8
7
6
5
4
3
2
1
Pathway
Fusion_kmeans
CP_kmeans

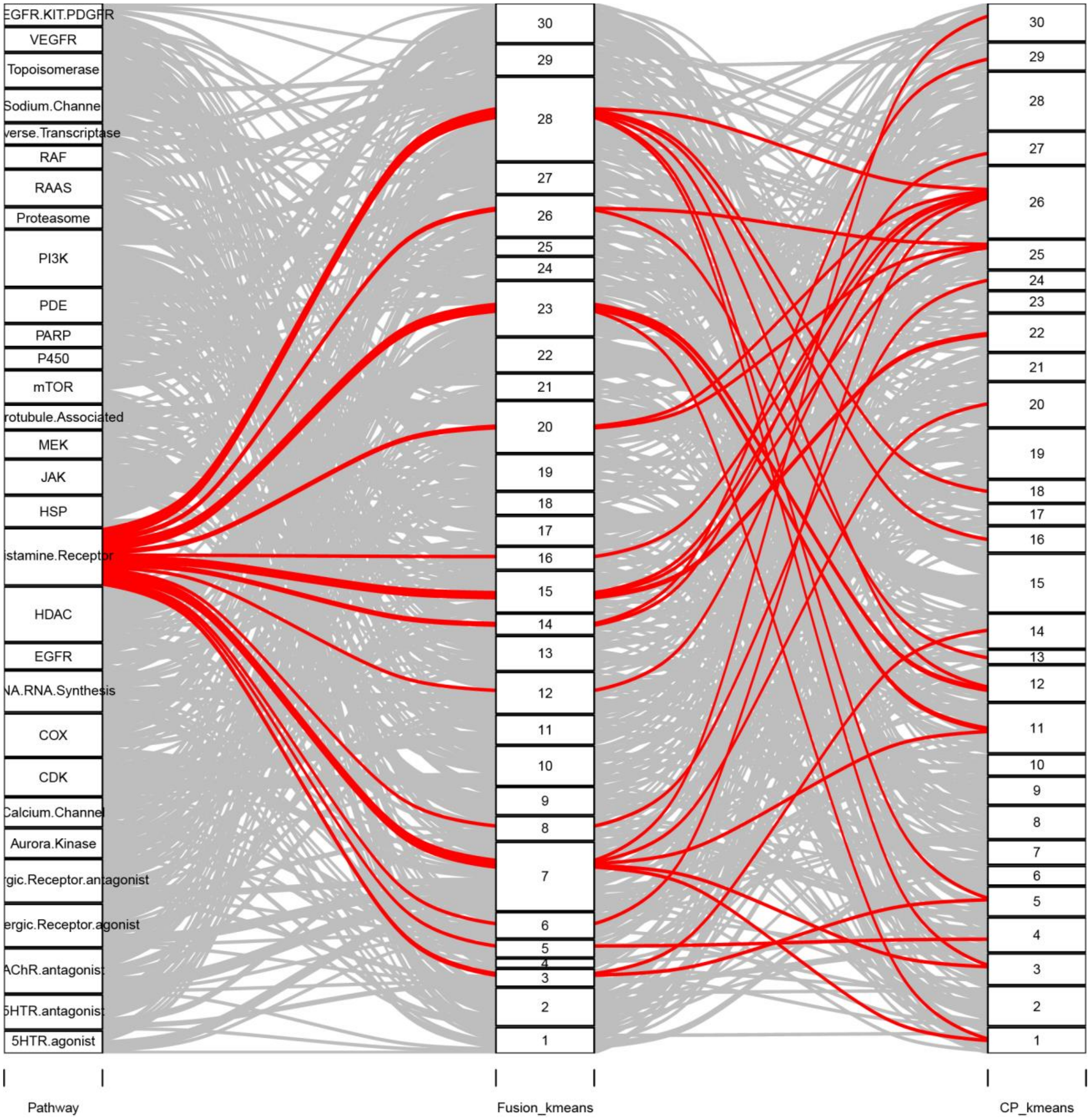
EGFR.KIT.PDGFR
VEGFR
Topoisomerase
Sodium.Channel
verse.Transcriptase
RAF
RAAS
Proteasome
PI3K
PDE
PARP
P450
mTOR
rotubule.Associated
MEK
JAK
HSP
istamine.Receptor
HDAC
EGFR
NA.RNA.Synthesis
COX
CDK
Calcium.Channel
Aurora.Kinase
rgic.Receptor.antagonist
ergic.Receptor.agonist
AChR.antagonist
5HTR.antagonist
5HTR.agonist
Pathway
Fusion_kmeans
CP_kmeans

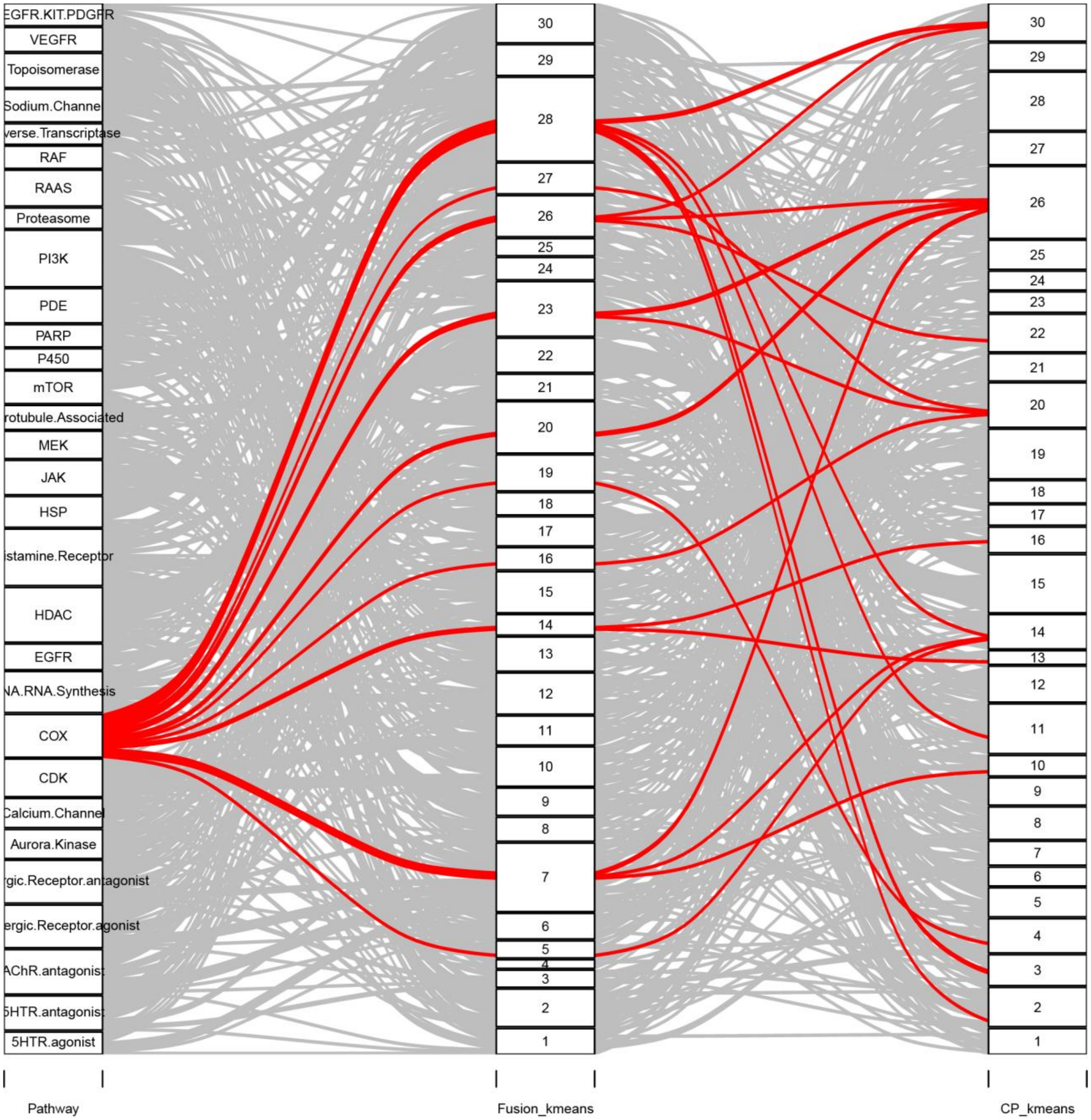
EGFR.KIT.PDGFR
VEGFR
Topoisomerase
Sodium.Channel
verse.Transcriptase
RAF
RAAS
Proteasome
PI3K
PDE
PARP
P450
mTOR
rotubule.Associated
MEK
JAK
HSP
istamine.Receptor
HDAC
EGFR
NA.RNA.Synthesis
COX
CDK
Calcium.Channel
Aurora.Kinase
rgic.Receptor.antagonist
ergic.Receptor.agonist
AChR.antagonist
5HTR.antagonist
5HTR.agonist
Pathway
Fusion_kmeans
CP_kmeans

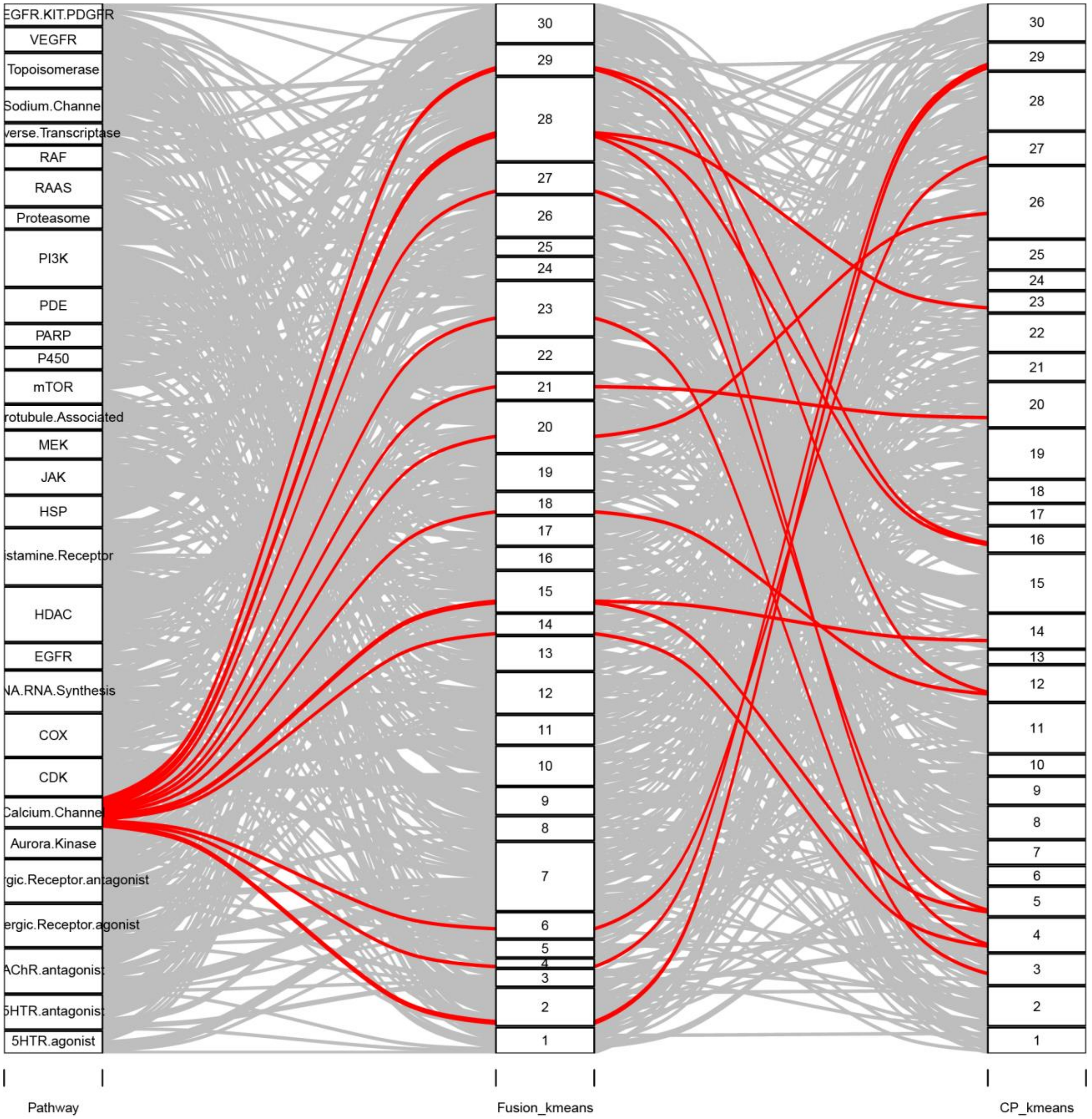
EGFR.KIT.PDGFR
VEGFR
Topoisomerase
Sodium.Channe
verse.Transcriptase
RAF
RAAS
Proteasome
PI3K
PDE
PARP
P450
mTOR
rotubule.Associated
MEK
JAK
HSP
istamine.Receptor
HDAC
EGFR
NA.RNA.Synthesis
COX
CDK
Calcium.Channe
Aurora.Kinase
rgic.Receptor.antagonist
ergic.Receptor.agonist
AChR.antagonist
5HTR.antagonist
5HTR.agonist
Pathway
Fusion_kmeans
CP_kmeans

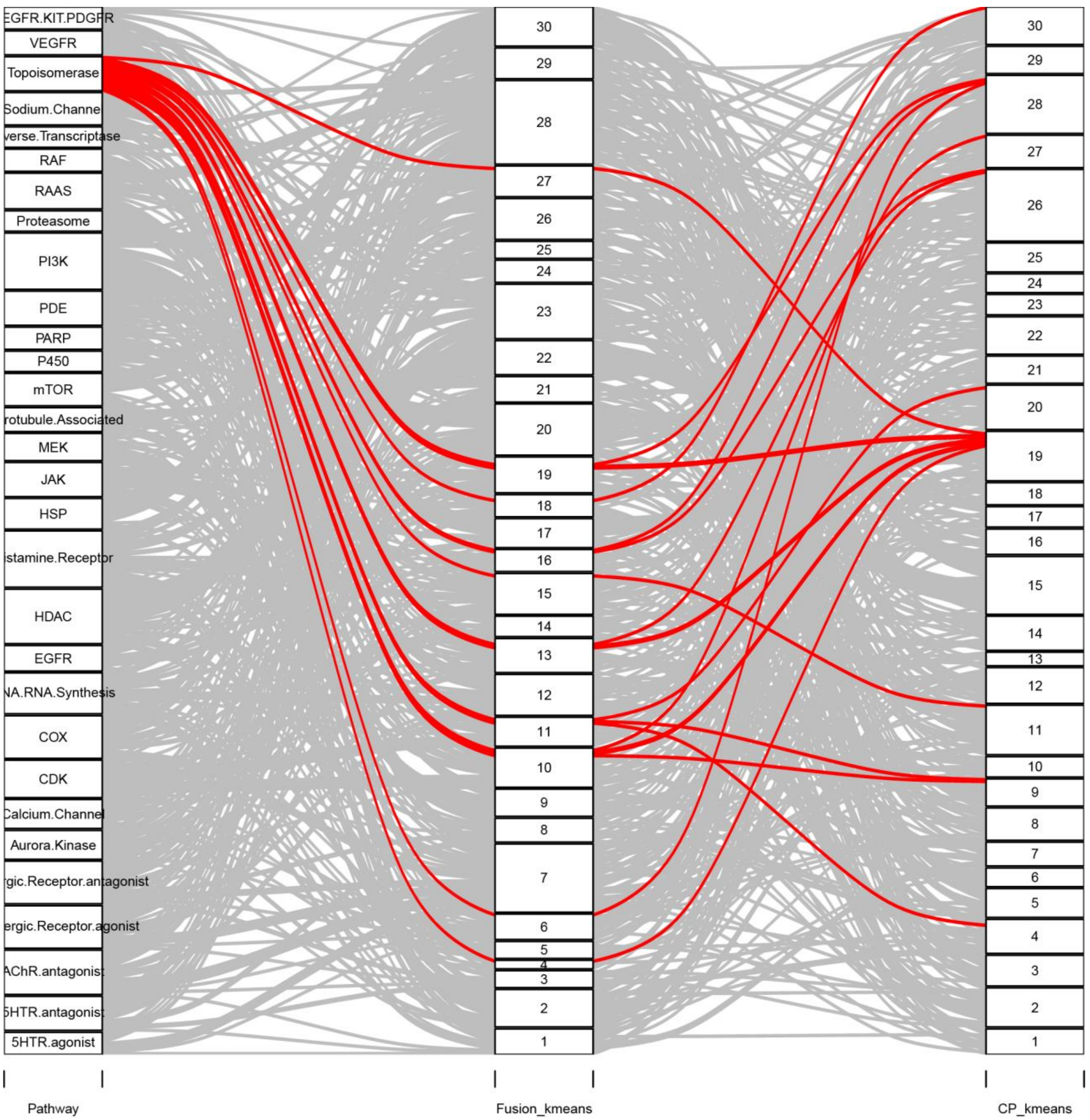
EGFR.KIT.PDGFR
VEGFR
Topoisomerase
Sodium.Channel
verse.Transcriptase
RAF
RAAS
Proteasome
PI3K
PDE
PARP
P450
mTOR
rotubule.Associated
MEK
JAK
HSP
istamine.Receptor
HDAC
EGFR
NA.RNA.Synthesis
COX
CDK
Calcium.Channel
Aurora.Kinase
rgic.Receptor.antagonist
ergic.Receptor.agonist
AChR.antagonist
5HTR.antagonist
5HTR.agonist
Pathway
Fusion_kmeans
CP_kmeans

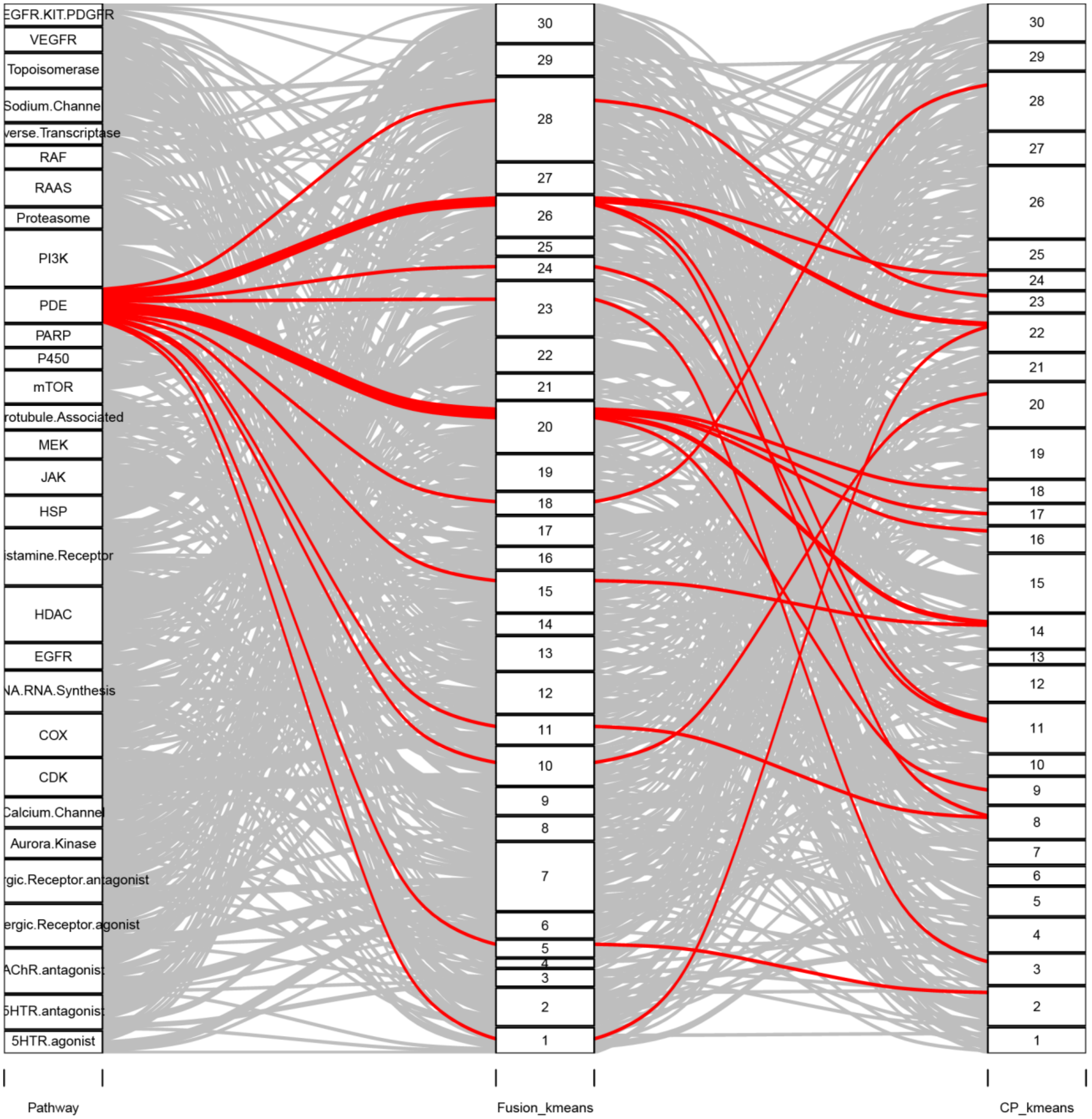
EGFR.KIT.PDGFR
VEGFR
Topoisomerase
Sodium.Channel
verse.Transcriptase
RAF
RAAS
Proteasome
PI3K
PDE
PARP
P450
mTOR
rotubule.Associated
MEK
JAK
HSP
istamine.Receptor
HDAC
EGFR
NA.RNA.Synthesis
COX
CDK
Calcium.Channel
Aurora.Kinase
gic.Receptor.antagonist
ergic.Receptor.agonist
AChR.antagonist
5HTR.antagonist
5HTR.agonist
Pathway
Fusion_kmeans
CP_kmeans

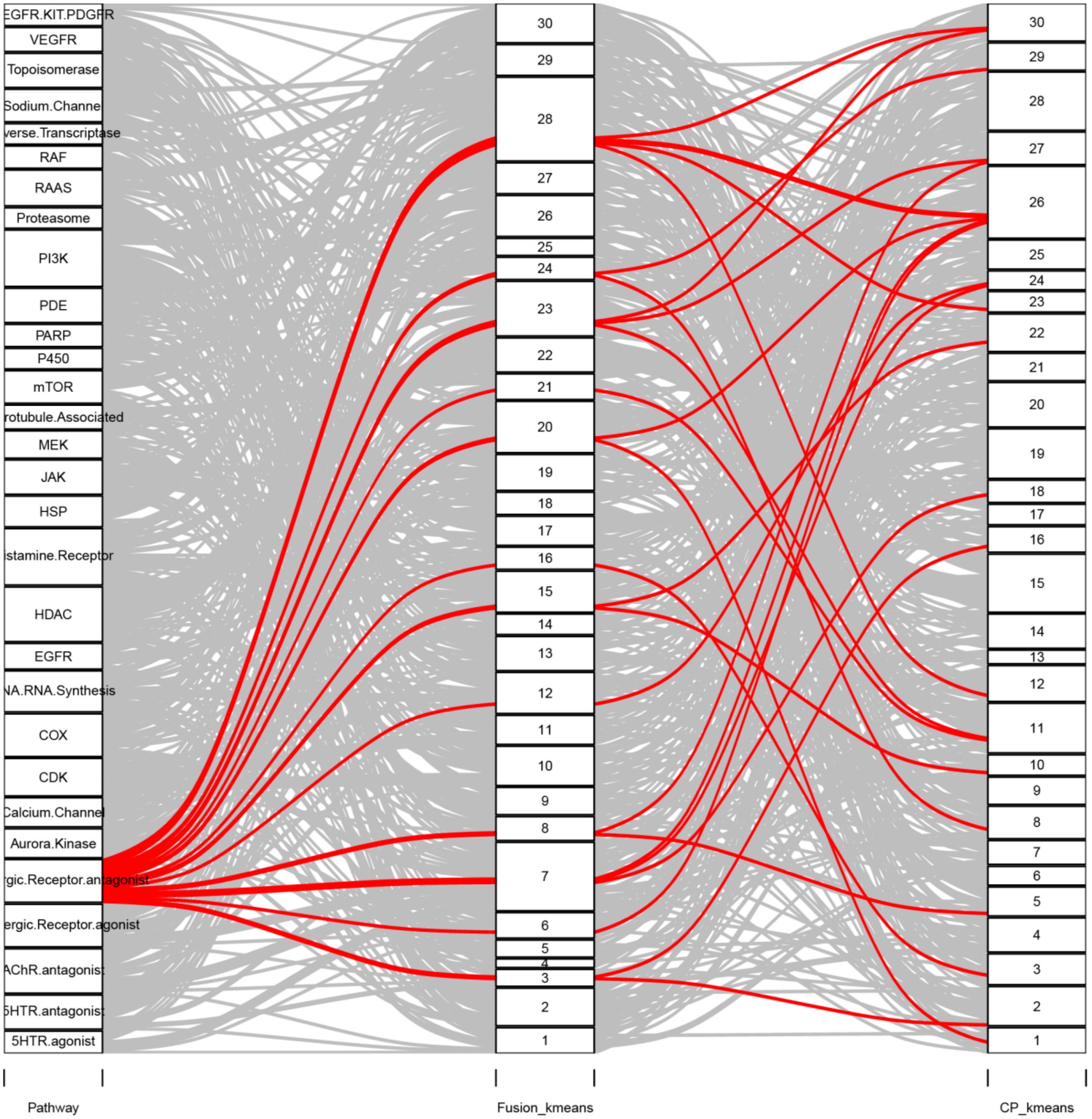
EGFR.KIT.PDGFR
VEGFR
Topoisomerase
Sodium.Channel
verse.Transcriptase
RAF
RAAS
Proteasome
PI3K
PDE
PARP
P450
mTOR
rotubule.Associated
MEK
JAK
HSP
istamine.Receptor
HDAC
EGFR
NA.RNA.Synthesis
COX
CDK
Calcium.Channel
Aurora.Kinase
gic.Receptor.antagonist
ergic.Receptor.agonist
AChR.antagonist
5HTR.antagonist
5HTR.agonist
Pathway
Fusion_kmeans
CP_kmeans

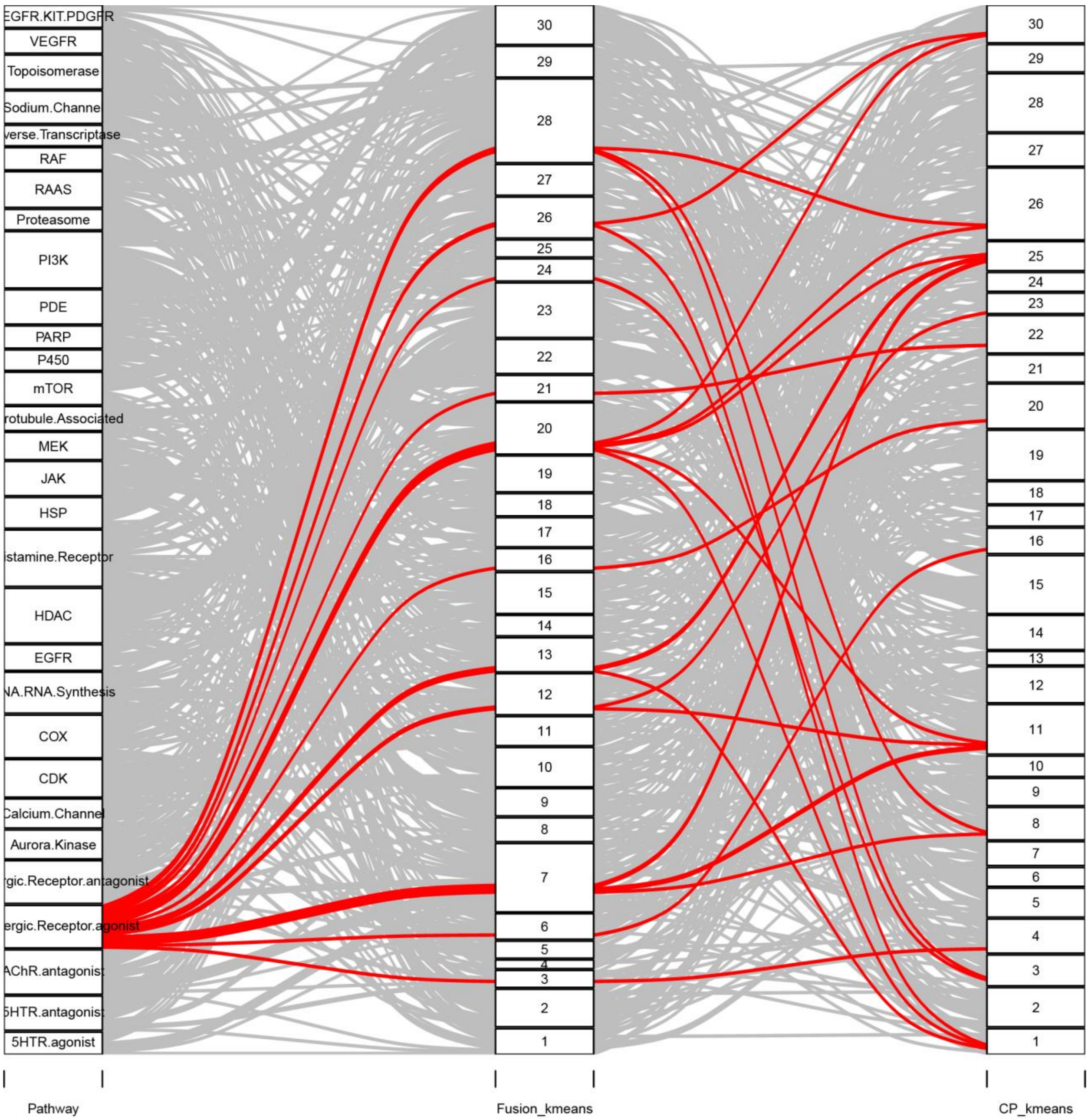
EGFR.KIT.PDGFR
VEGFR
Topoisomerase
Sodium.Channel
verse.Transcriptase
RAF
RAAS
Proteasome
PI3K
PDE
PARP
P450
mTOR
rotubule.Associated
MEK
JAK
HSP
istamine.Receptor
HDAC
EGFR
NA.RNA.Synthesis
COX
CDK
Calcium.Channel
Aurora.Kinase
rgic.Receptor.antagonist
ergic.Receptor.agonist
AChR.antagonist
5HTR.antagonist
5HTR.agonist
30
29
28
27
26
25
24
23
22
21
20
19
18
17
16
15
14
13
12
11
10
9
8
7
6
5
4
3
2
1
Pathway
Fusion_kmeans
CP_kmeans

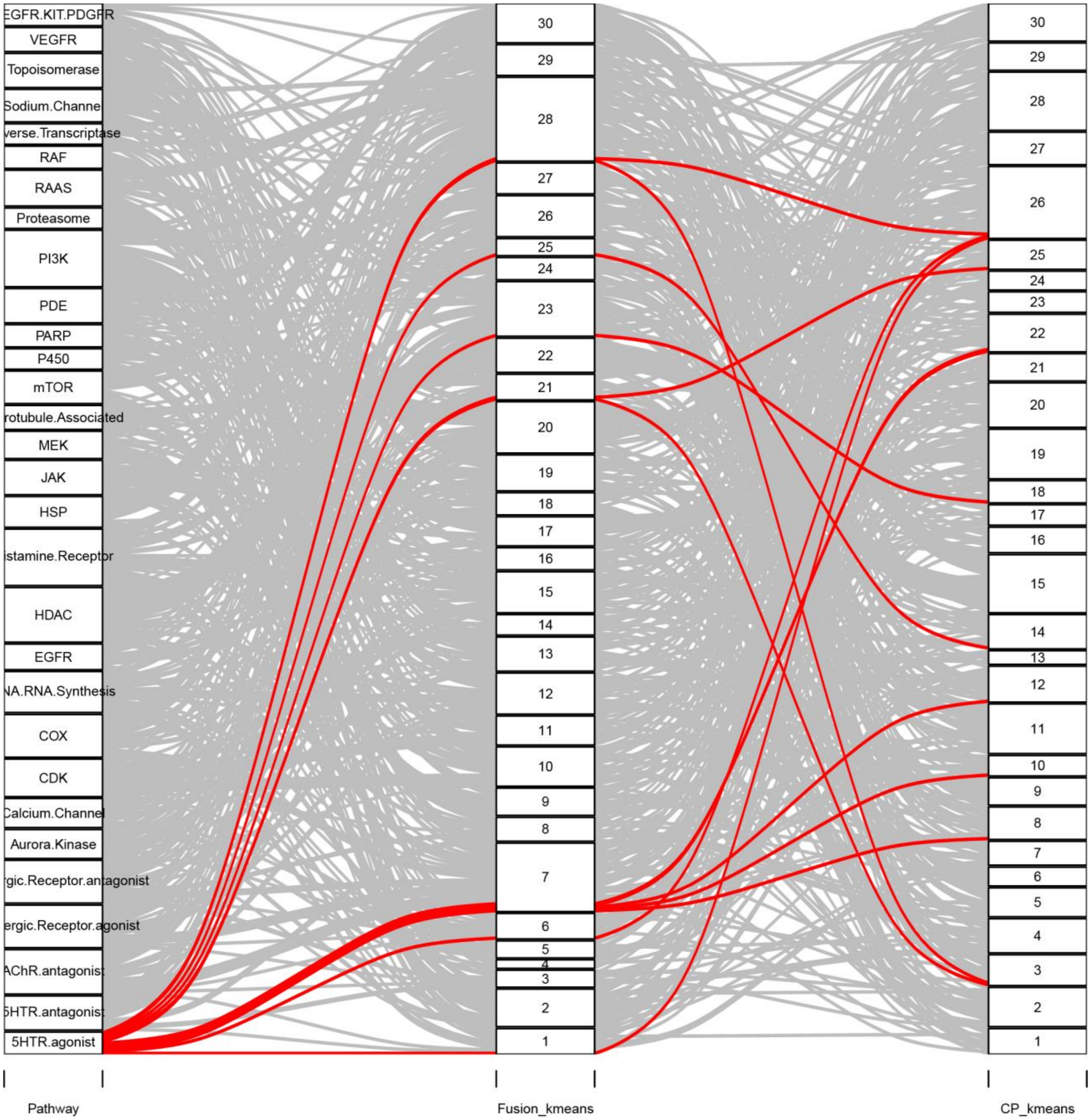

EGFR.KIT.PDGFR
VEGFR
Topoisomerase
Sodium.Channe
verse.Transcriptase
RAF
RAAS
Proteasome
PI3K
PDE
PARP
P450
mTOR
rotubule.Associated
MEK
JAK
HSP
istamine.Receptor
HDAC
EGFR
NA.RNA.Synthesis
COX
CDK
Calcium.Channel
Aurora.Kinase
rgic.Receptor.antagonist
ergic.Receptor.agonist
AChR.antagonist
5HTR.antagonist
5HTR.agonist
Pathway
Fusion_kmeans
CP_kmeans

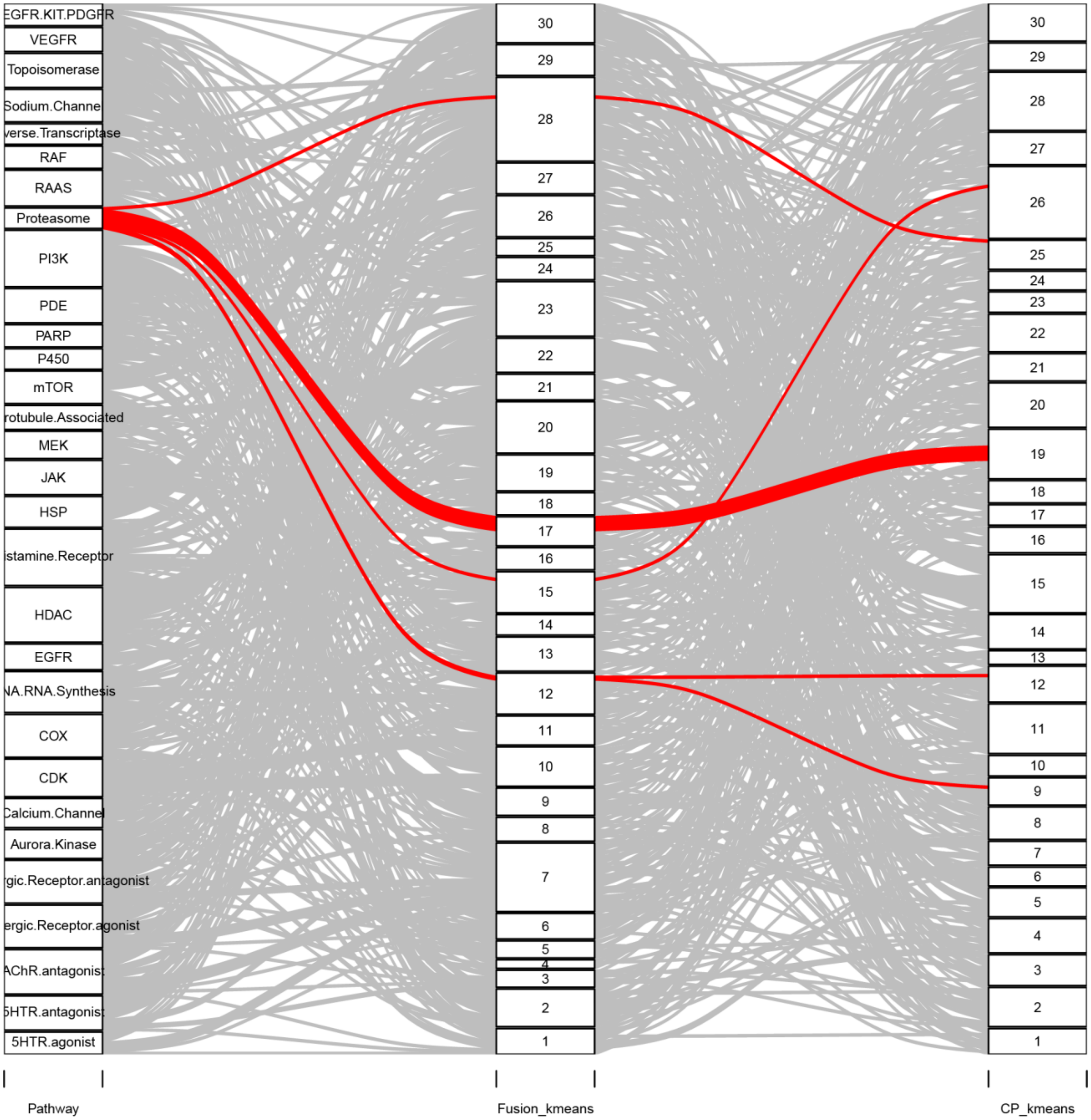

EGFR.KIT.PDGFR
VEGFR
Topoisomerase
Sodium.Channel
verse.Transcriptase
RAF
RAAS
Proteasome
PI3K
PDE
PARP
P450
mTOR
rotubule.Associated
MEK
JAK
HSP
istamine.Receptor
HDAC
EGFR
NA.RNA.Synthesis
COX
CDK
Calcium.Channel
Aurora.Kinase
gic.Receptor.antagonist
ergic.Receptor.agonist
AChR.antagonist
5HTR.antagonist
5HTR.agonist
Pathway
Fusion_kmeans
CP_kmeans

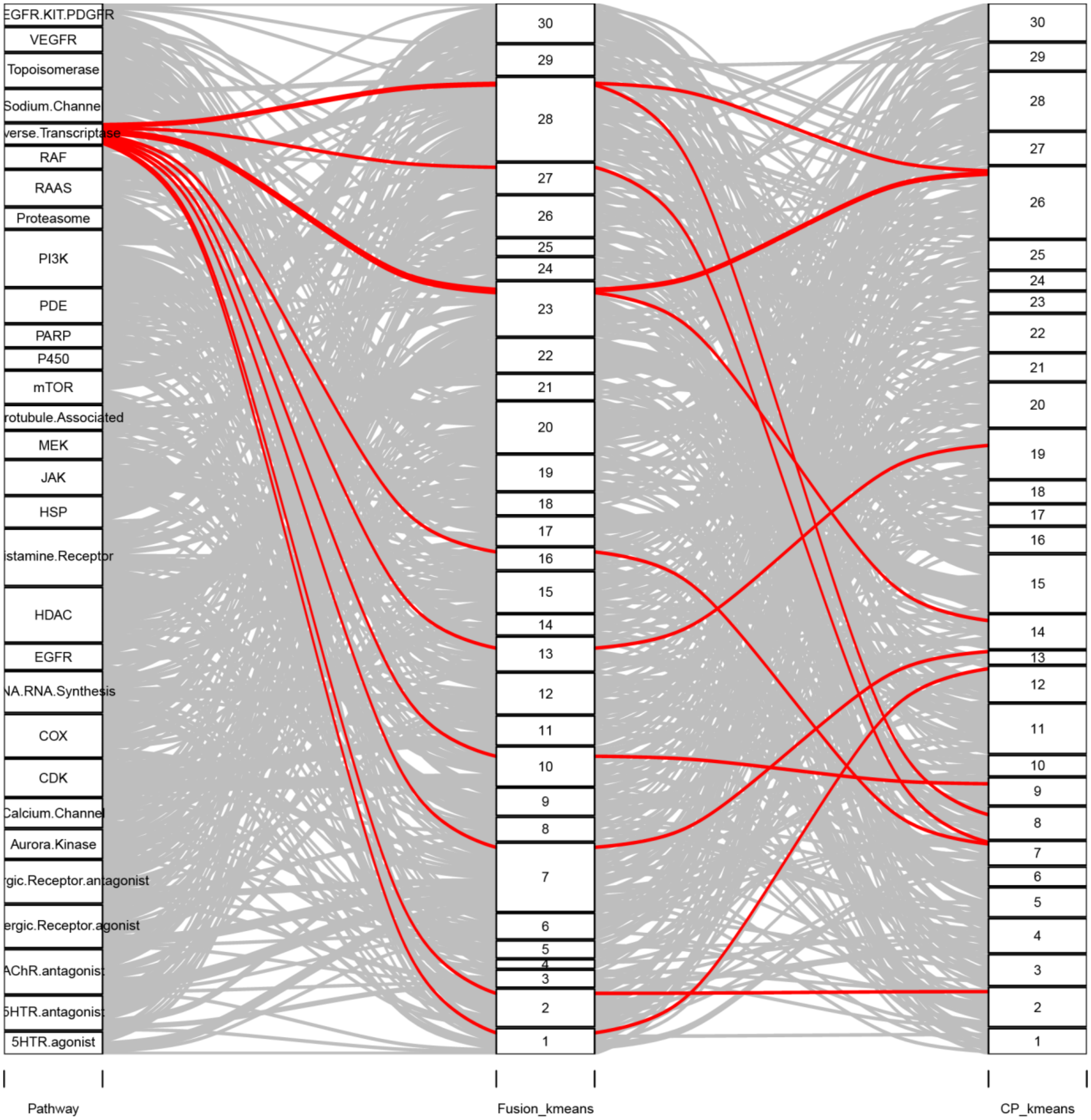

EGFR.KIT.PDGFR
VEGFR
Topoisomerase
Sodium.Channel
verse.Transcriptase
RAF
RAAS
Proteasome
PI3K
PDE
PARP
P450
mTOR
rotubule.Associated
MEK
JAK
HSP
istamine.Receptor
HDAC
EGFR
NA.RNA.Synthesis
COX
CDK
Calcium.Channel
Aurora.Kinase
gic.Receptor.antagonist
ergic.Receptor.agonist
AChR.antagonist
5HTR.antagonist
5HTR.agonist
30
29
28
27
26
25
24
23
22
21
20
19
18
17
16
15
14
13
12
11
10
9
8
7
6
5
4
3
2
1
Pathway
Fusion_kmeans
CP_kmeans

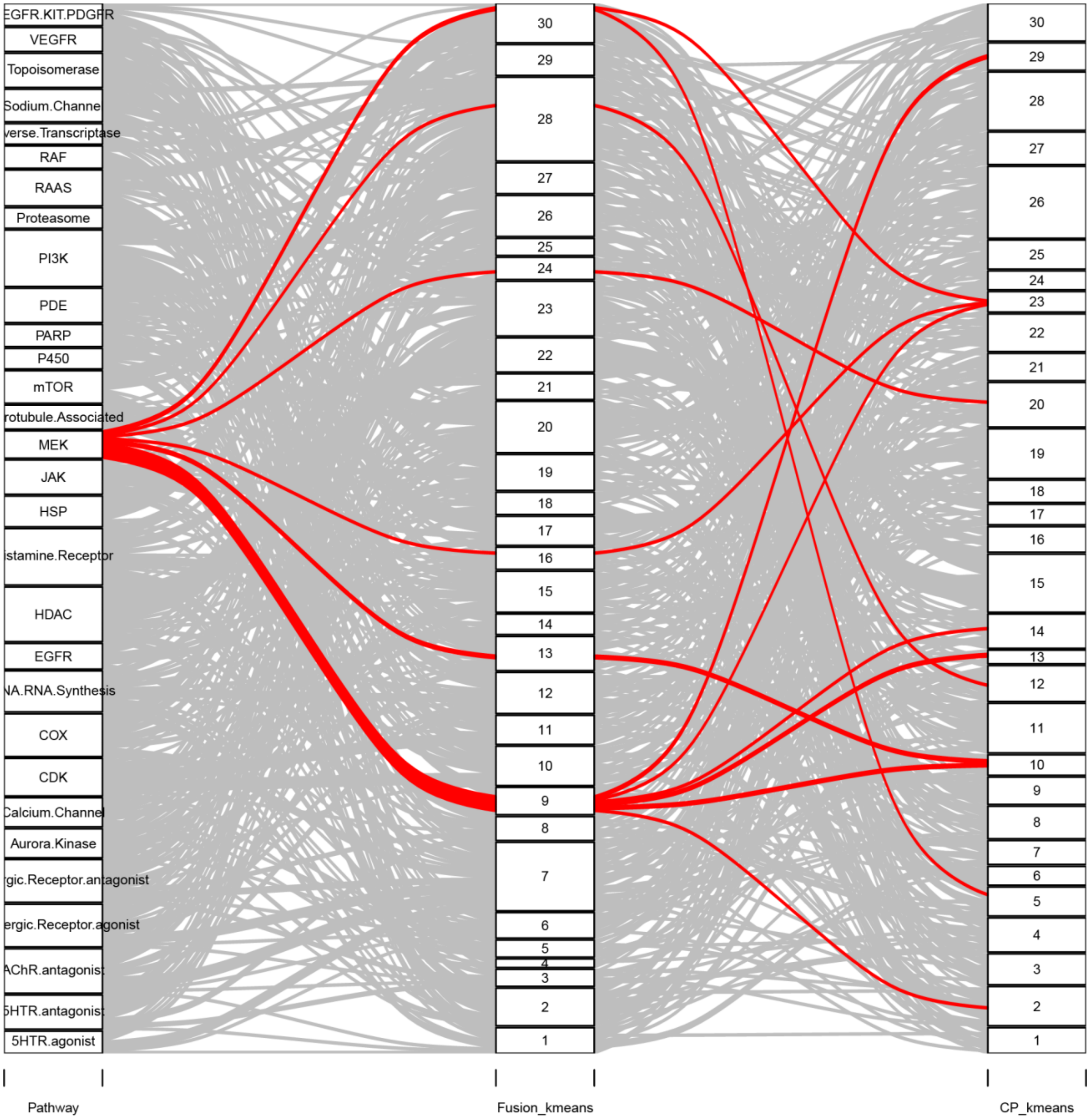

EGFR.KIT.PDGFR
VEGFR
Topoisomerase
Sodium.Channel
verse.Transcriptase
RAF
RAAS
Proteasome
PI3K
PDE
PARP
P450
mTOR
rotubule.Associated
MEK
JAK
HSP
istamine.Receptor
HDAC
EGFR
NA.RNA.Synthesis
COX
CDK
Calcium.Channel
Aurora.Kinase
gic.Receptor.antagonist
ergic.Receptor.agonist
AChR.antagonist
5HTR.antagonist
5HTR.agonist
Pathway
Fusion_kmeans
CP_kmeans

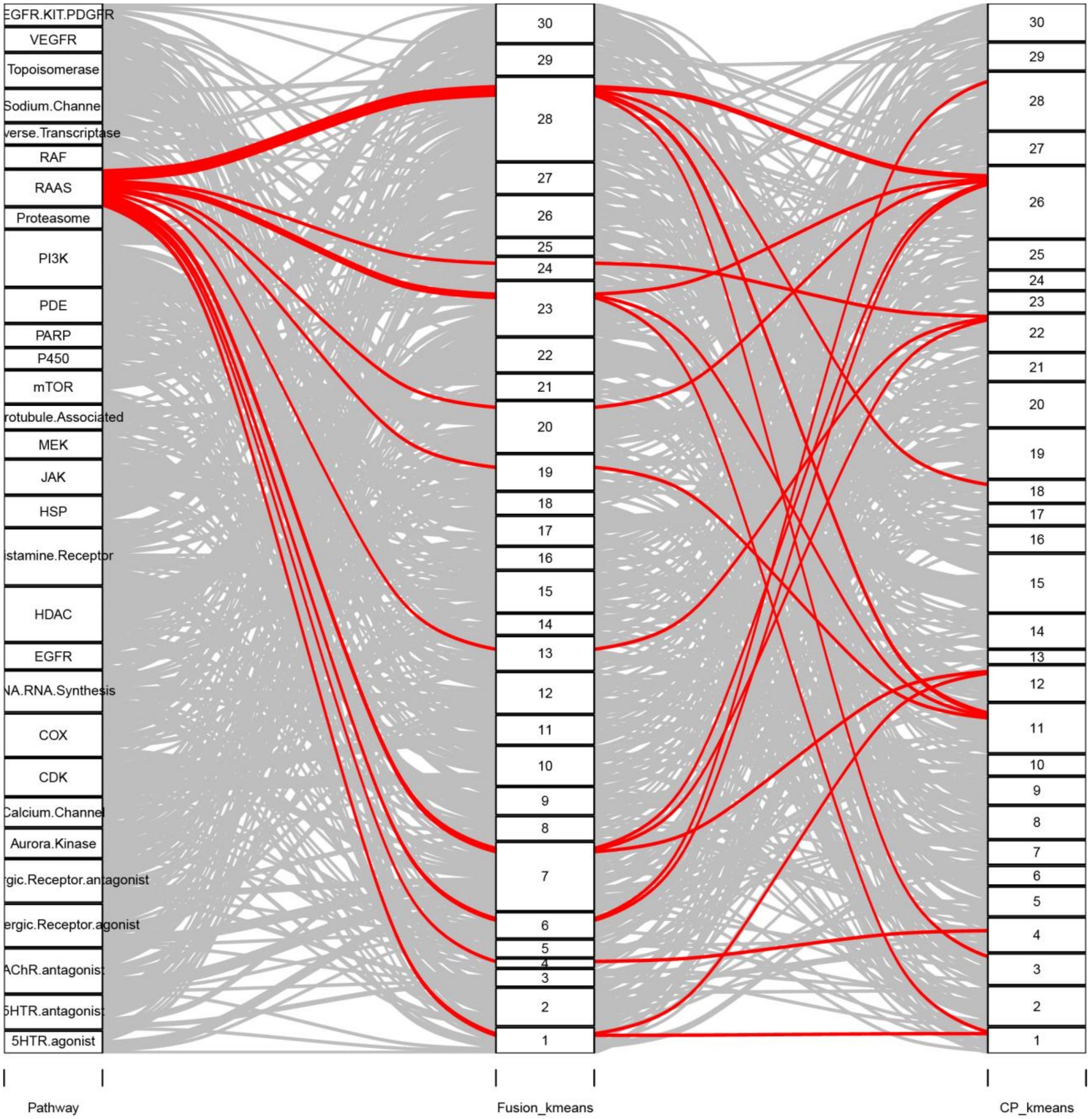
EGFR.KIT.PDGFR
VEGFR
Topoisomerase
Sodium.Channel
verse.Transcriptase
RAF
RAAS
Proteasome
PI3K
PDE
PARP
P450
mTOR
rotubule.Associated
MEK
JAK
HSP
istamine.Receptor
HDAC
EGFR
NA.RNA.Synthesis
COX
CDK
Calcium.Channel
Aurora.Kinase
rgic.Receptor.antagonist
ergic.Receptor.agonist
AChR.antagonist
5HTR.antagonist
5HTR.agonist
Pathway
Fusion_kmeans
CP_kmeans

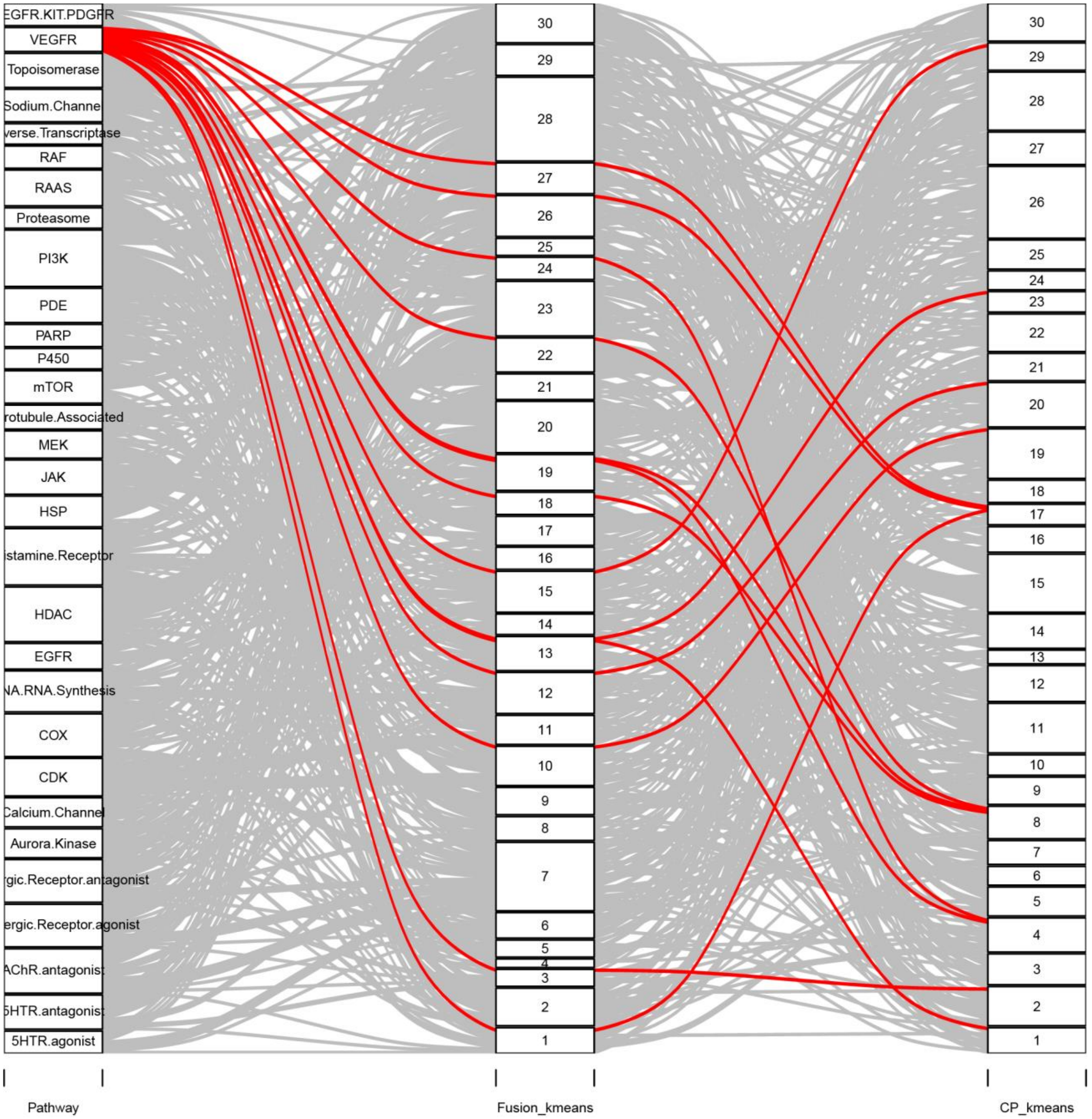
EGFR.KIT.PDGFR
VEGFR
Topoisomerase
Sodium.Channel
verse.Transcriptase
RAF
RAAS
Proteasome
PI3K
PDE
PARP
P450
mTOR
rotubule.Associated
MEK
JAK
HSP
istamine.Receptor
HDAC
EGFR
NA.RNA.Synthesis
COX
CDK
Calcium.Channel
Aurora.Kinase
rgic.Receptor.antagonist
ergic.Receptor.agonist
AChR.antagonist
5HTR.antagonist
5HTR.agonist
Pathway
Fusion_kmeans
CP_kmeans

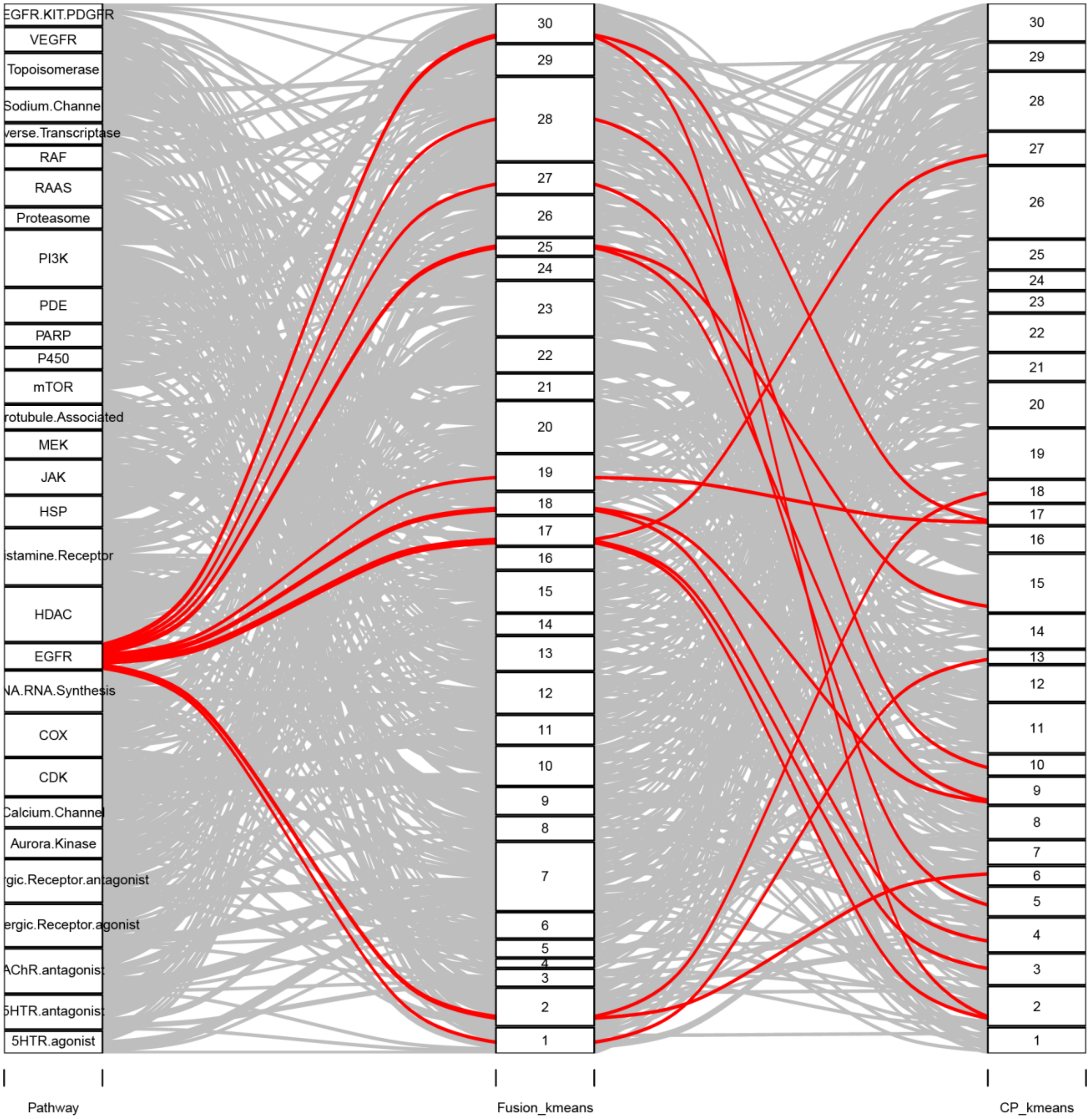
EGFR.KIT.PDGFR
VEGFR
Topoisomerase
Sodium.Channel
verse.Transcriptase
RAF
RAAS
Proteasome
PI3K
PDE
PARP
P450
mTOR
rotubule.Associated
MEK
JAK
HSP
istamine.Receptor
HDAC
EGFR
NA.RNA.Synthesis
COX
CDK
Calcium.Channel
Aurora.Kinase
rgic.Receptor.antagonist
ergic.Receptor.agonist
AChR.antagonist
5HTR.antagonist
5HTR.agonist
Pathway
Fusion_kmeans
CP_kmeans

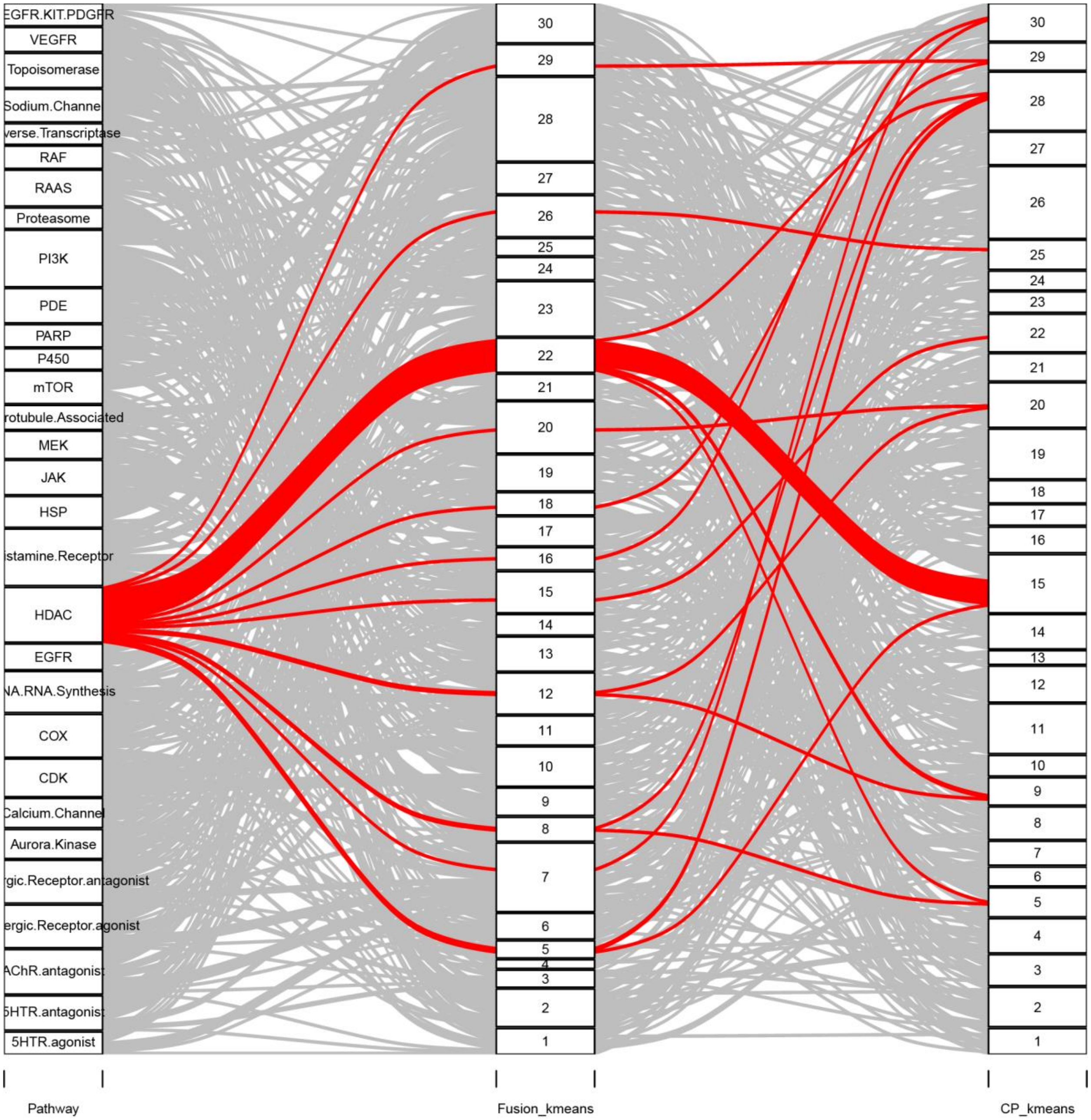
EGFR.KIT.PDGFR
VEGFR
Topoisomerase
Sodium.Channel
verse.Transcriptase
RAF
RAAS
Proteasome
PI3K
PDE
PARP
P450
mTOR
rotubule.Associated
MEK
JAK
HSP
istamine.Receptor
HDAC
EGFR
NA.RNA.Synthesis
COX
CDK
Calcium.Channel
Aurora.Kinase
rgic.Receptor.antagonist
ergic.Receptor.agonist
AChR.antagonist
5HTR.antagonist
5HTR.agonist
Pathway
Fusion_kmeans
CP_kmeans

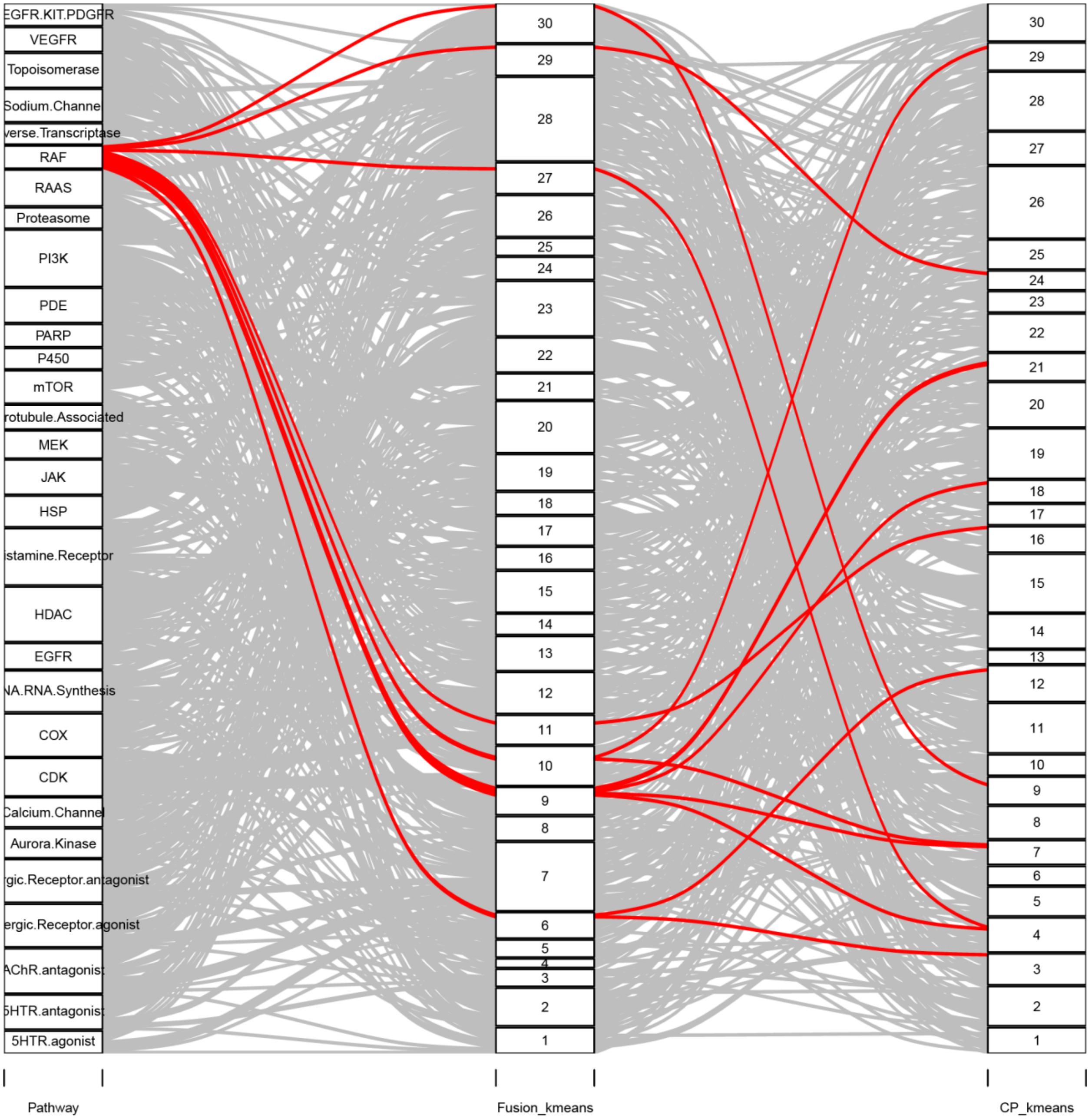

EGFR.KIT.PDGFR
VEGFR
Topoisomerase
Sodium.Channel
verse.Transcriptase
RAF
RAAS
Proteasome
PI3K
PDE
PARP
P450
mTOR
rotubule.Associated
MEK
JAK
HSP
istamine.Receptor
HDAC
EGFR
NA.RNA.Synthesis
COX
CDK
Calcium.Channel
Aurora.Kinase
gic.Receptor.antagonist
ergic.Receptor.agonist
AChR.antagonist
5HTR.antagonist
5HTR.agonist
30 29 28 27 26 25 24 23 22 21 20 19 18 17 16 15 14 13 12 11 10 9 8 7 6 5 4 3 2 1
Pathway
Fusion_kmeans
CP_kmeans

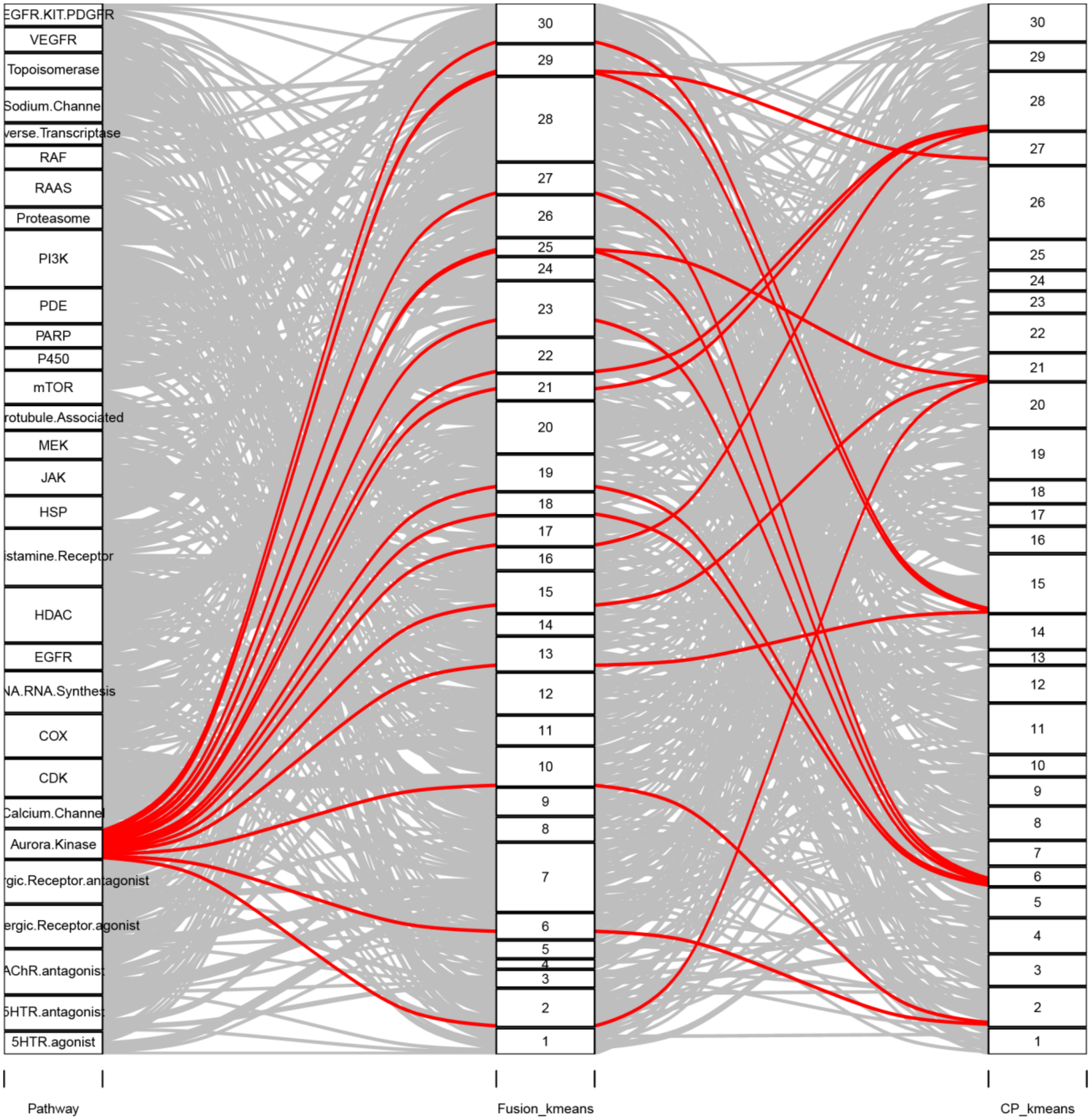

EGFR.KIT.PDGFR
VEGFR
Topoisomerase
Sodium.Channel
verse.Transcriptase
RAF
RAAS
Proteasome
PI3K
PDE
PARP
P450
mTOR
rotubule.Associated
MEK
JAK
HSP
istamine.Receptor
HDAC
EGFR
NA.RNA.Synthesis
COX
CDK
Calcium.Channel
Aurora.Kinase
gic.Receptor.antagonist
ergic.Receptor.agonist
AChR.antagonist
5HTR.antagonist
5HTR.agonist
Pathway
Fusion_kmeans
CP_kmeans

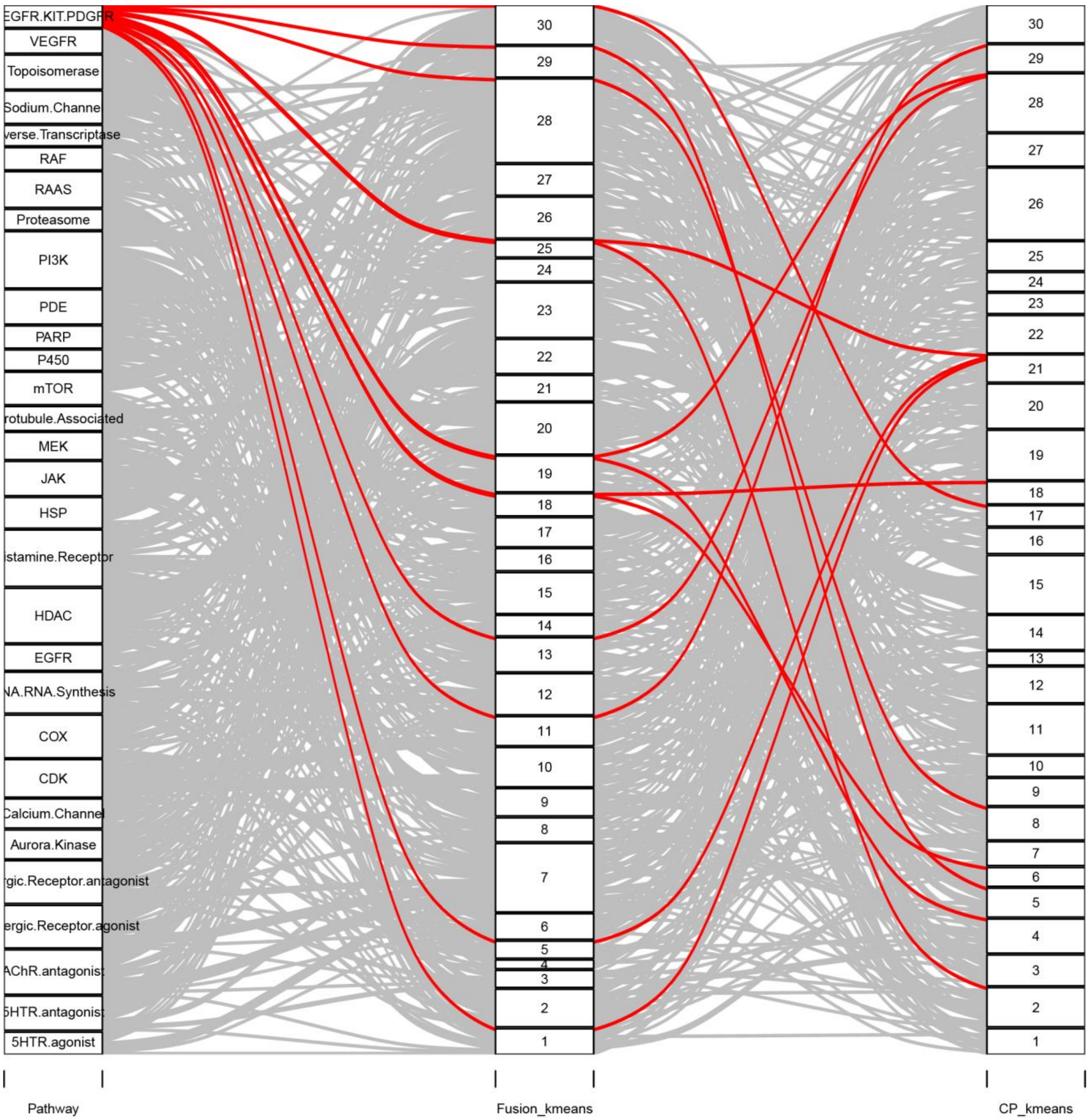
EGFR.KIT.PDGFR
VEGFR
Topoisomerase
Sodium.Channel
verse.Transcriptase
RAF
RAAS
Proteasome
PI3K
PDE
PARP
P450
mTOR
rotubule.Associated
MEK
JAK
HSP
istamine.Receptor
HDAC
EGFR
NA.RNA.Synthesis
COX
CDK
Calcium.Channel
Aurora.Kinase
rgic.Receptor.antagonist
ergic.Receptor.agonist
AChR.antagonist
5HTR.antagonist
5HTR.agonist
Pathway
Fusion_kmeans
CP_kmeans

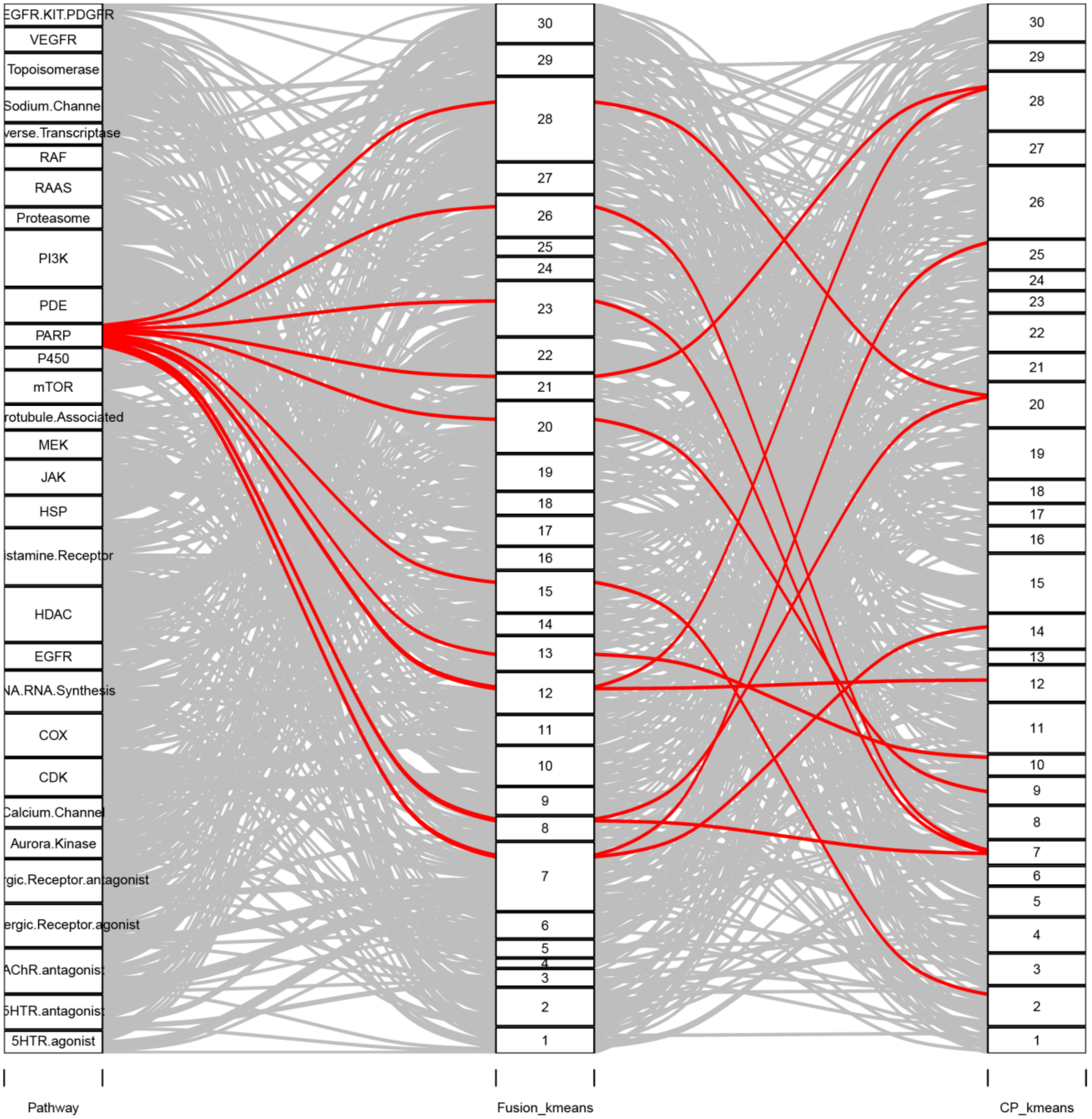

EGFR.KIT.PDGFR
VEGFR
Topoisomerase
Sodium.Channel
verse.Transcriptase
RAF
RAAS
Proteasome
PI3K
PDE
PARP
P450
mTOR
rotubule.Associated
MEK
JAK
HSP
istamine.Receptor
HDAC
EGFR
NA.RNA.Synthesis
COX
CDK
Calcium.Channel
Aurora.Kinase
gic.Receptor.antagonist
ergic.Receptor.agonist
AChR.antagonist
5HTR.antagonist
5HTR.agonist
Pathway
Fusion_kmeans
CP_kmeans

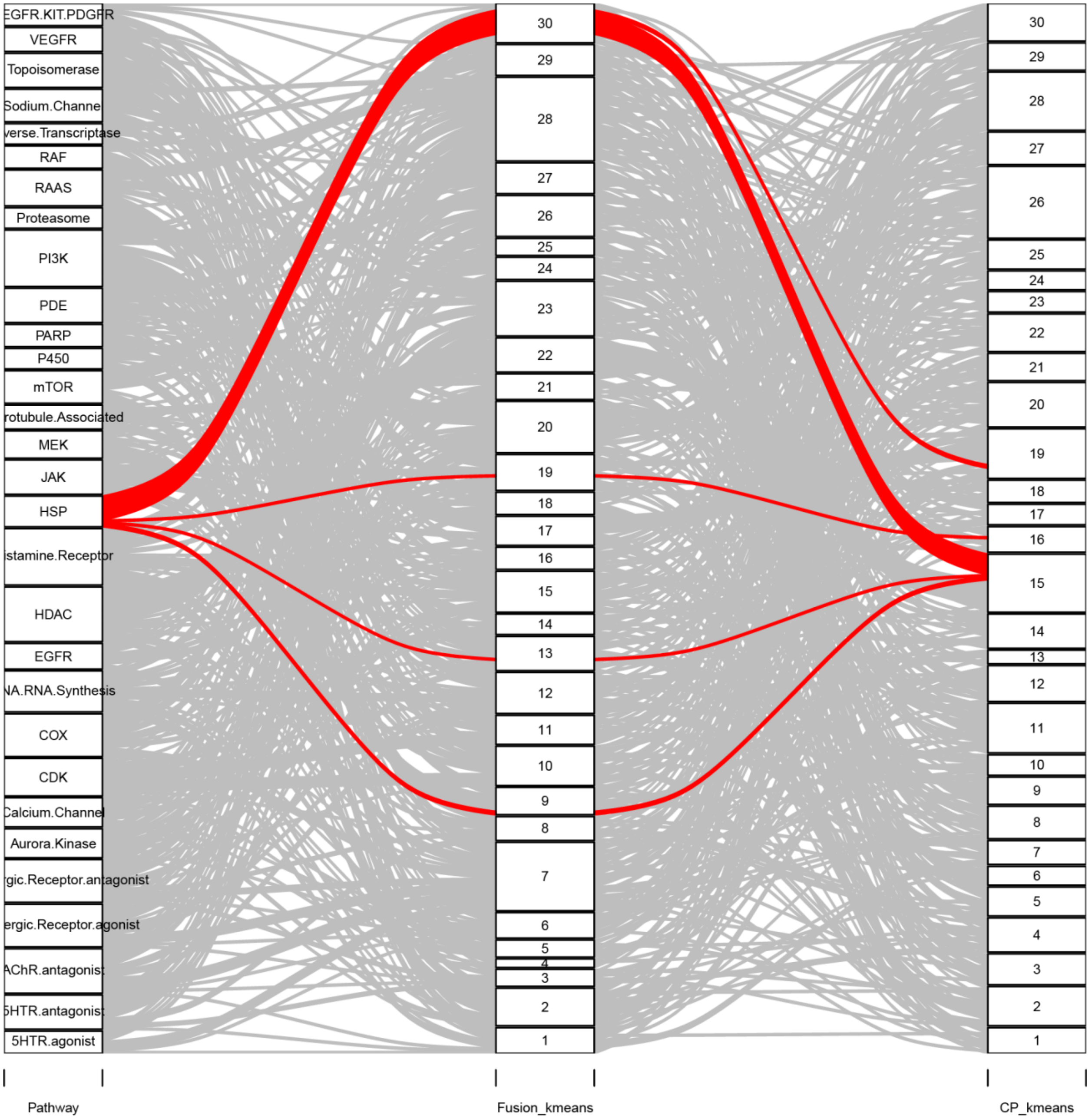
EGFR.KIT.PDGFR
VEGFR
Topoisomerase
Sodium.Channel
verse.Transcriptase
RAF
RAAS
Proteasome
PI3K
PDE
PARP
P450
mTOR
rotubule.Associated
MEK
JAK
HSP
istamine.Receptor
HDAC
EGFR
NA.RNA.Synthesis
COX
CDK
Calcium.Channel
Aurora.Kinase
rgic.Receptor.antagonist
ergic.Receptor.agonist
AChR.antagonist
5HTR.antagonist
5HTR.agonist
30
29
28
27
26
25
24
23
22
21
20
19
18
17
16
15
14
13
12
11
10
9
8
7
6
5
4
3
2
1
Pathway
Fusion_kmeans
CP_kmeans

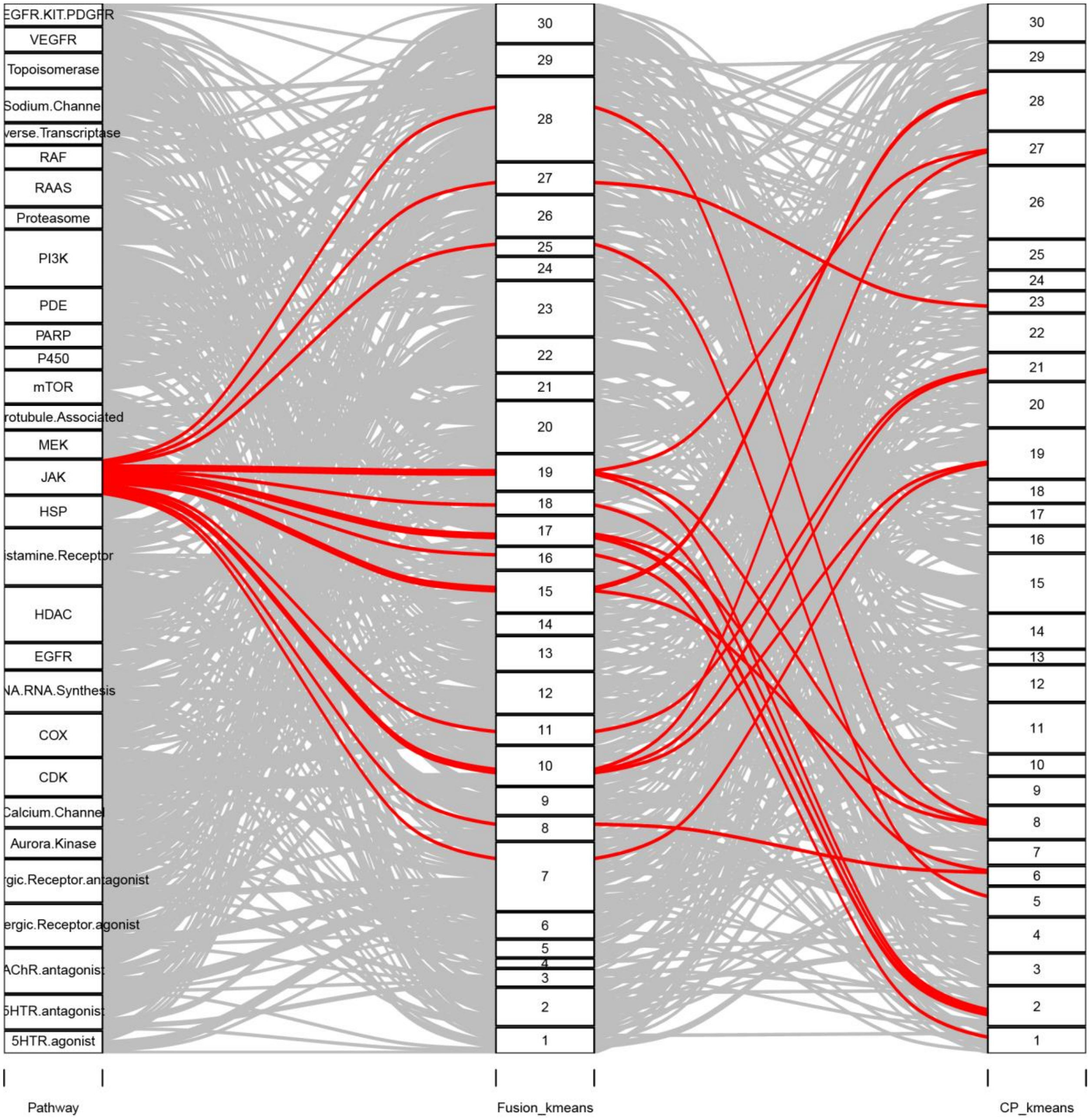

EGFR.KIT.PDGFR
VEGFR
Topoisomerase
Sodium.Channel
verse.Transcriptase
RAF
RAAS
Proteasome
PI3K
PDE
PARP
P450
mTOR
rotubule.Associated
MEK
JAK
HSP
istamine.Receptor
HDAC
EGFR
NA.RNA.Synthesis
COX
CDK
Calcium.Channel
Aurora.Kinase
rgic.Receptor.antagonist
ergic.Receptor.agonist
AChR.antagonist
5HTR.antagonist
5HTR.agonist
Pathway
Fusion_kmeans
CP_kmeans

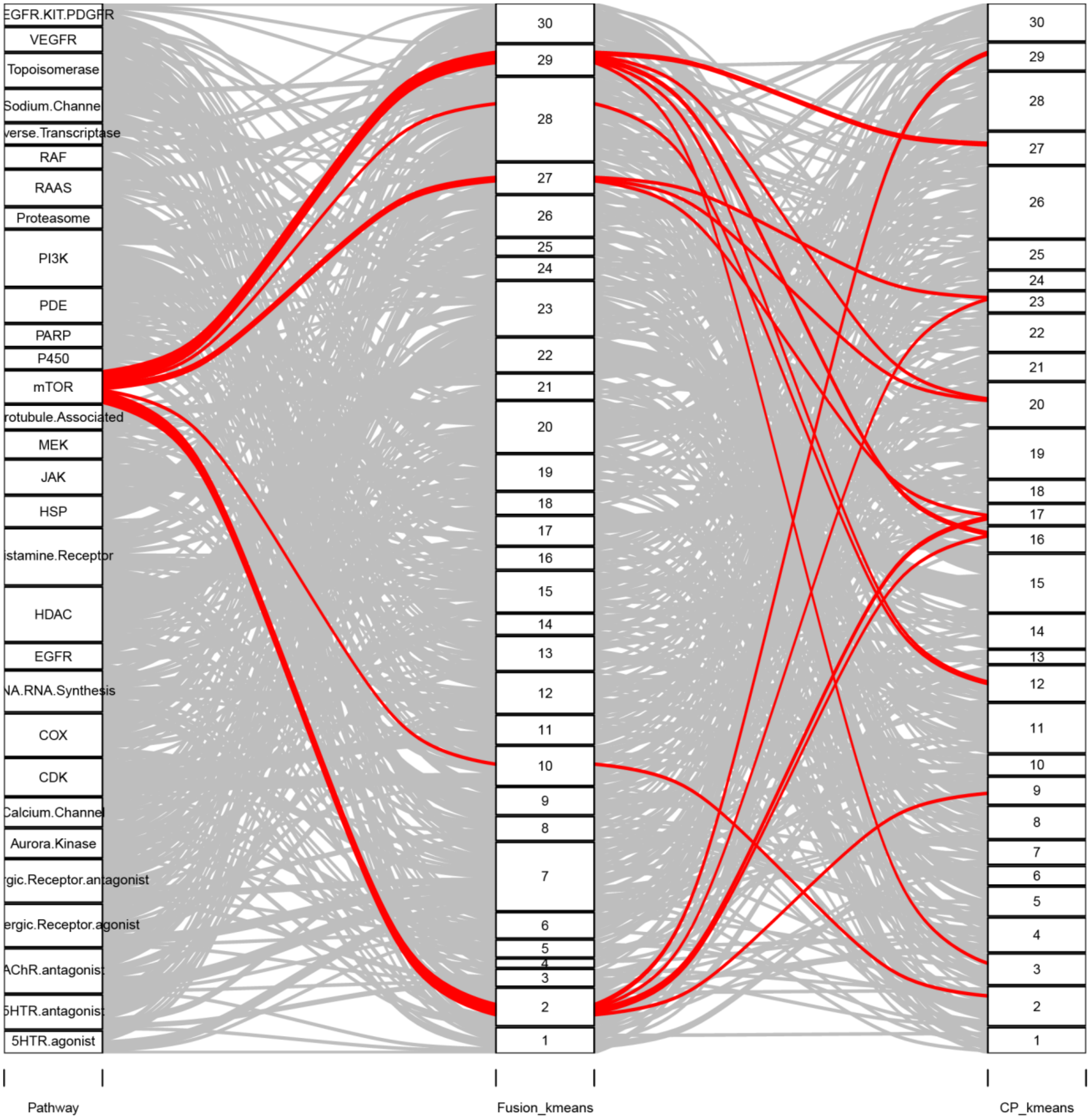

EGFR.KIT.PDGFR
VEGFR
Topoisomerase
Sodium.Channel
verse.Transcriptase
RAF
RAAS
Proteasome
PI3K
PDE
PARP
P450
mTOR
rotubule.Associated
MEK
JAK
HSP
istamine.Receptor
HDAC
EGFR
NA.RNA.Synthesis
COX
CDK
Calcium.Channel
Aurora.Kinase
gic.Receptor.antagonist
ergic.Receptor.agonist
AChR.antagonist
5HTR.antagonist
5HTR.agonist
Pathway
Fusion_kmeans
CP_kmeans

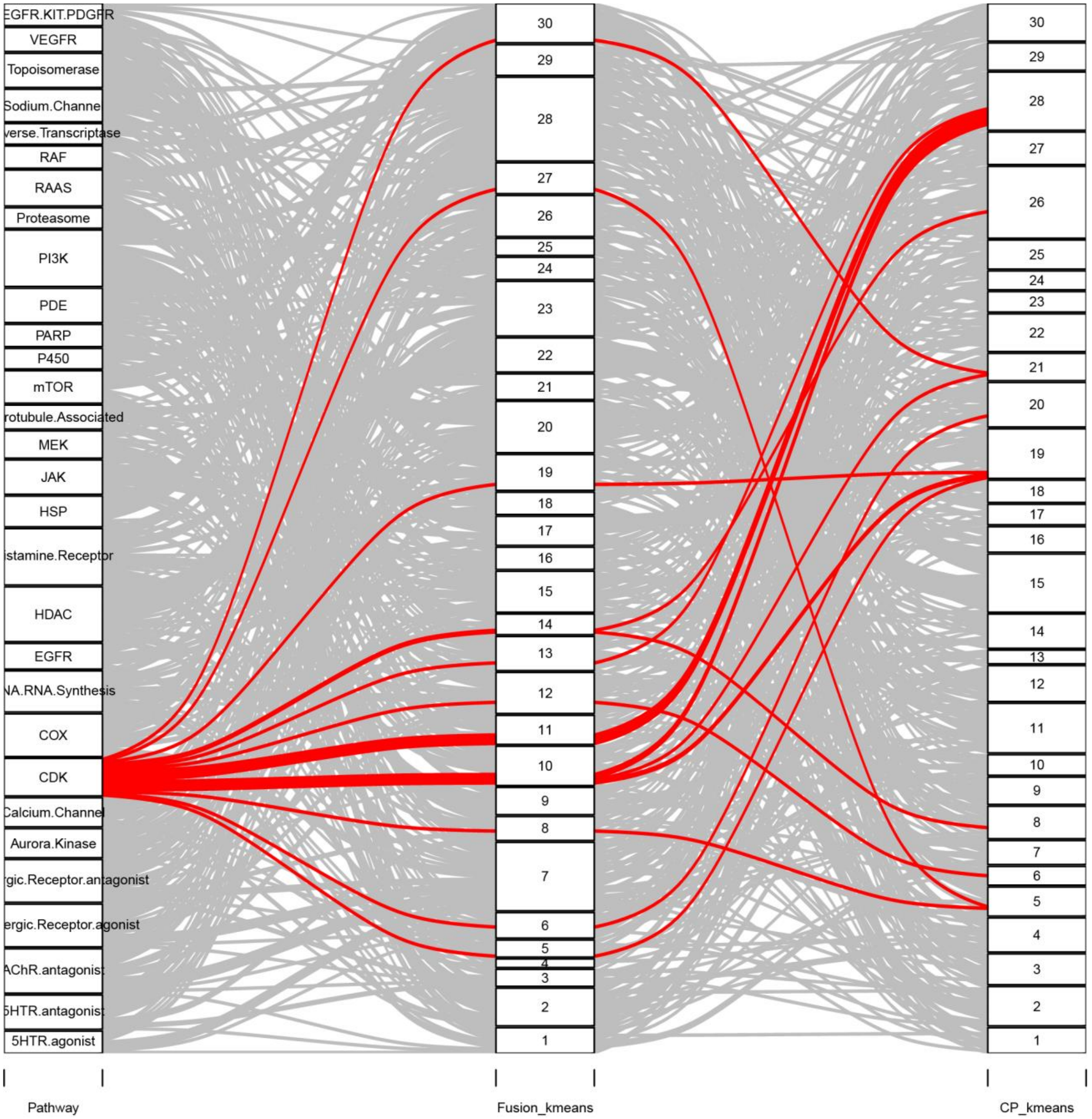
EGFR.KIT.PDGFR
VEGFR
Topoisomerase
Sodium.Channel
verse.Transcriptase
RAF
RAAS
Proteasome
PI3K
PDE
PARP
P450
mTOR
rotubule.Associated
MEK
JAK
HSP
istamine.Receptor
HDAC
EGFR
NA.RNA.Synthesis
COX
CDK
Calcium.Channel
Aurora.Kinase
rgic.Receptor.antagonist
ergic.Receptor.agonist
AChR.antagonist
5HTR.antagonist
5HTR.agonist
Pathway
Fusion_kmeans
CP_kmeans

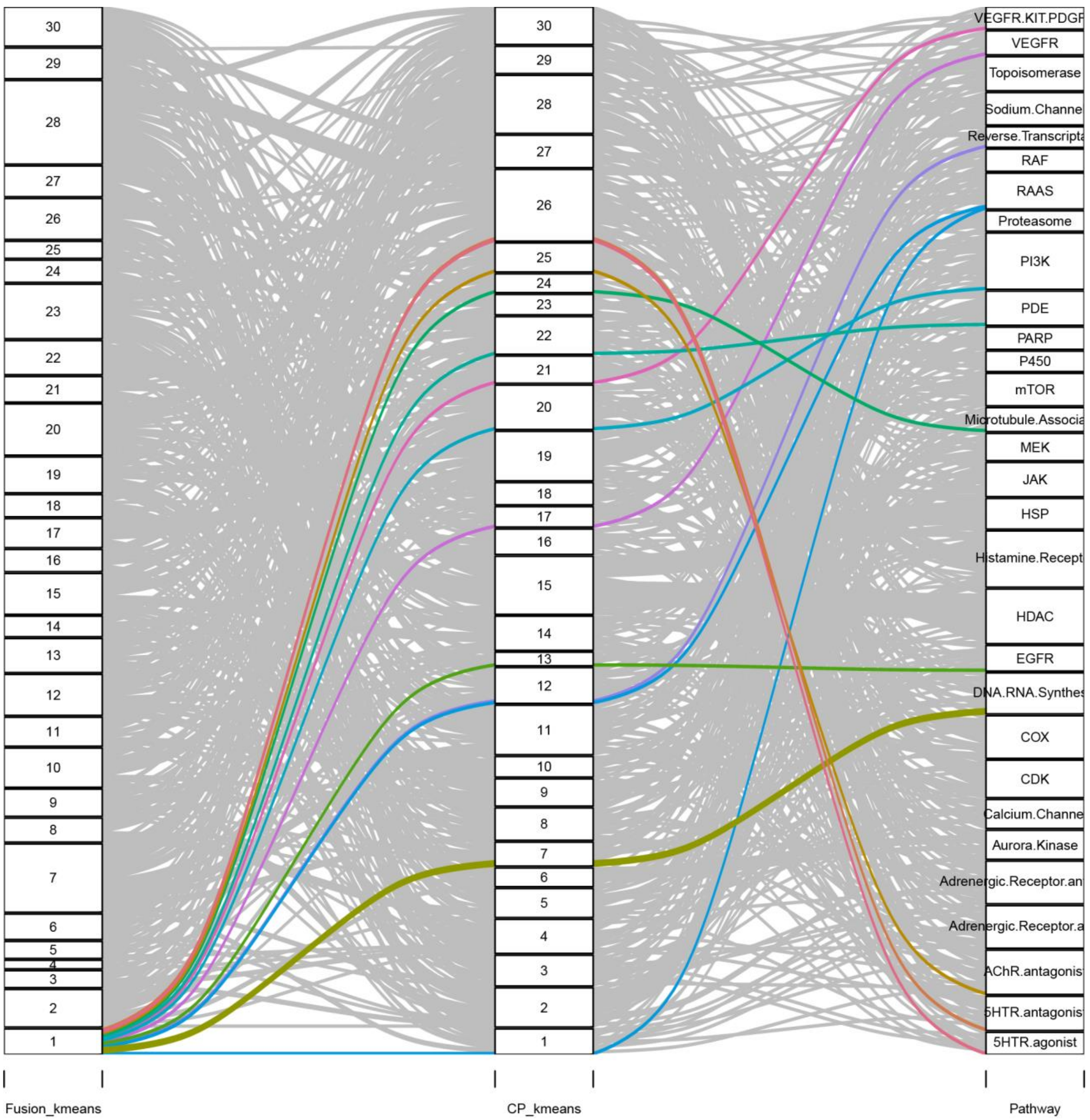

**Supplementary Figure 6. Alluvial diagrams for each cluster in FUSION and each cluster in CP.** Alluvial diagrams for each k-means cluster in the FUSION (A) and CP (B) datasets. Comparison of target class associations using k-means clustering (k=30) of FUSION and CP signatures for the top 30 target classes in the Selleck chemical library. Each line represents a compound, and each panel highlights a different cluster. Chemicals belonging to different target classes are indicated by different colors.

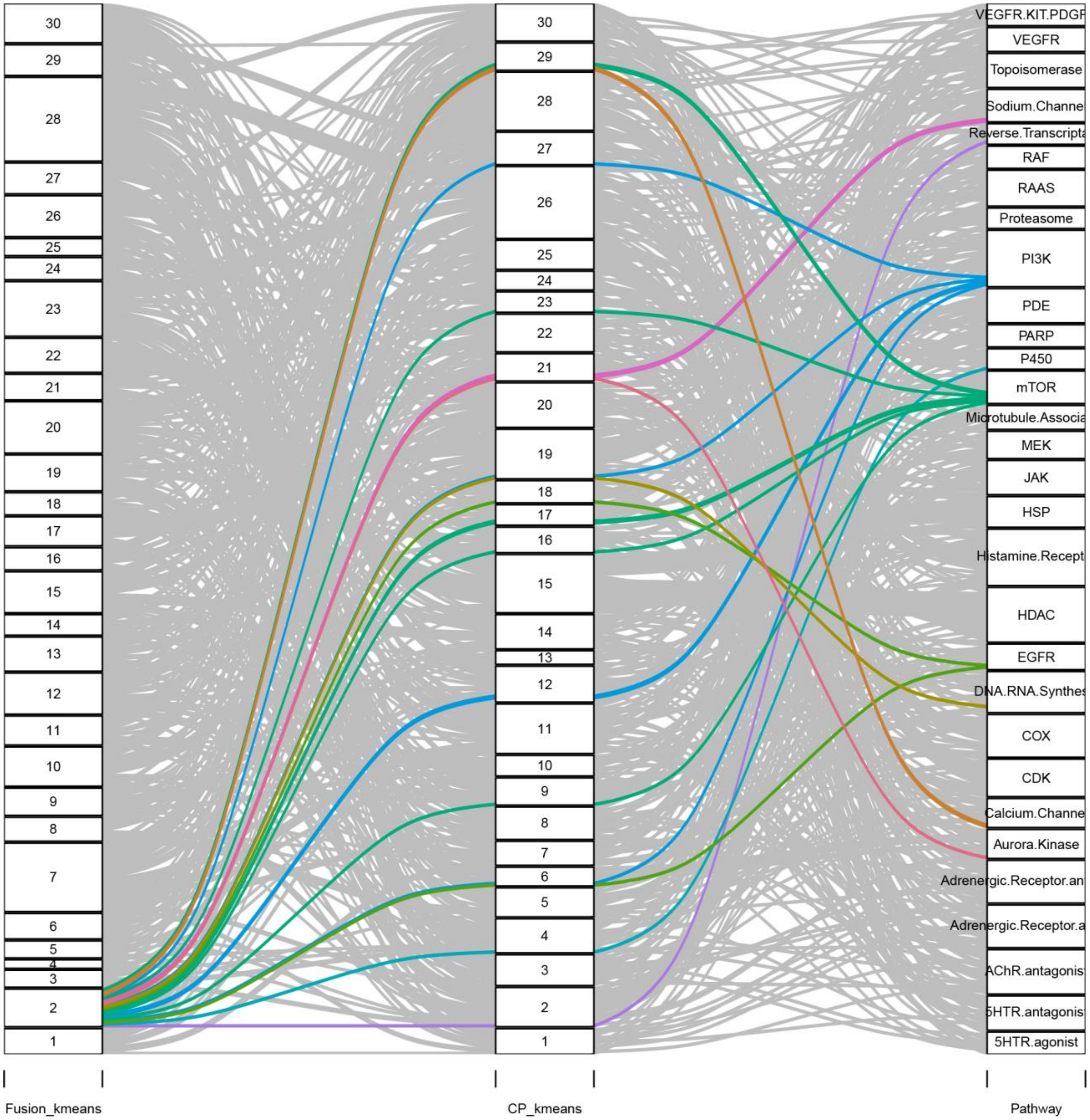

Fusion_kmeans
CP_kmeans
Pathway
30
29
28
27
26
25
24
23
22
21
20
19
18
17
16
15
14
13
12
11
10
9
8
7
6
5
4
3
2
1
VEGFR.KIT.PDGF
VEGFR
Topoisomerase
Sodium.Channe
Reverse.Transcripta
RAF
RAAS
Proteasome
PI3K
PDE
PARP
P450
mTOR
Microtubule.Associa
MEK
JAK
HSP
Histamine.Recept
HDAC
EGFR
DNA.RNA.Synthes
COX
CDK
Calcium.Channe
Aurora.Kinase
Adrenergic.Receptor.an
Adrenergic.Receptor.a
AChR.antagonis
5HTR.antagonis
5HTR.agonist

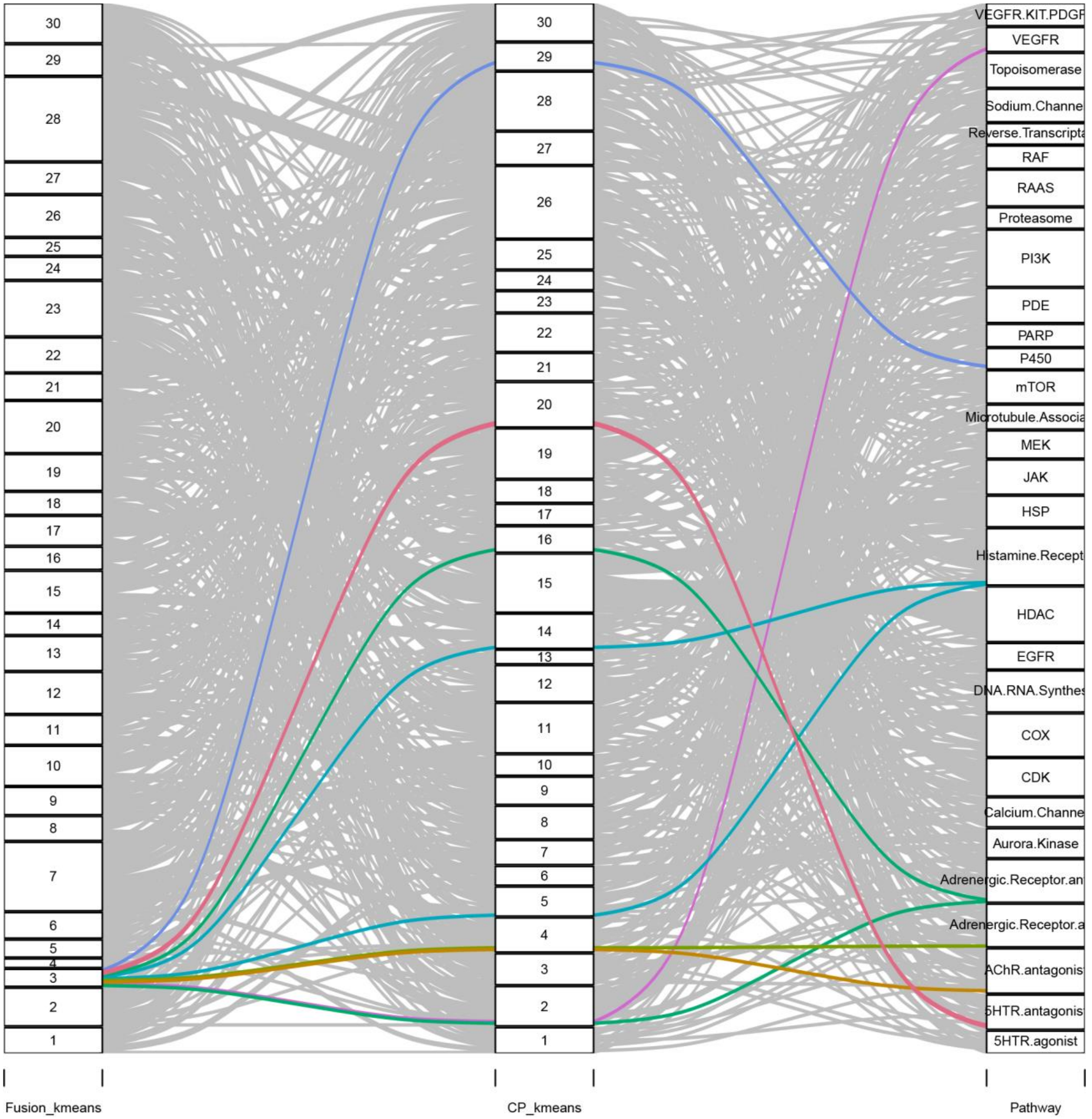
30
29
28
27
26
25
24
23
22
21
20
19
18
17
16
15
14
13
12
11
10
9
8
7
6
5
4
3
2
1
Fusion_kmeans
CP_kmeans
Pathway
VEGFR.KIT.PDGF
VEGFR
Topoisomerase
Sodium.Channe
Reverse.Transcripta
RAF
RAAS
Proteasome
PI3K
PDE
PARP
P450
mTOR
Microtubule.Associa
MEK
JAK
HSP
Histamine.Recept
HDAC
EGFR
DNA.RNA.Synthes
COX
CDK
Calcium.Channe
Aurora.Kinase
Adrenergic.Receptor.an
Adrenergic.Receptor.a
AChR.antagonis
5HTR.antagonis
5HTR.agonist

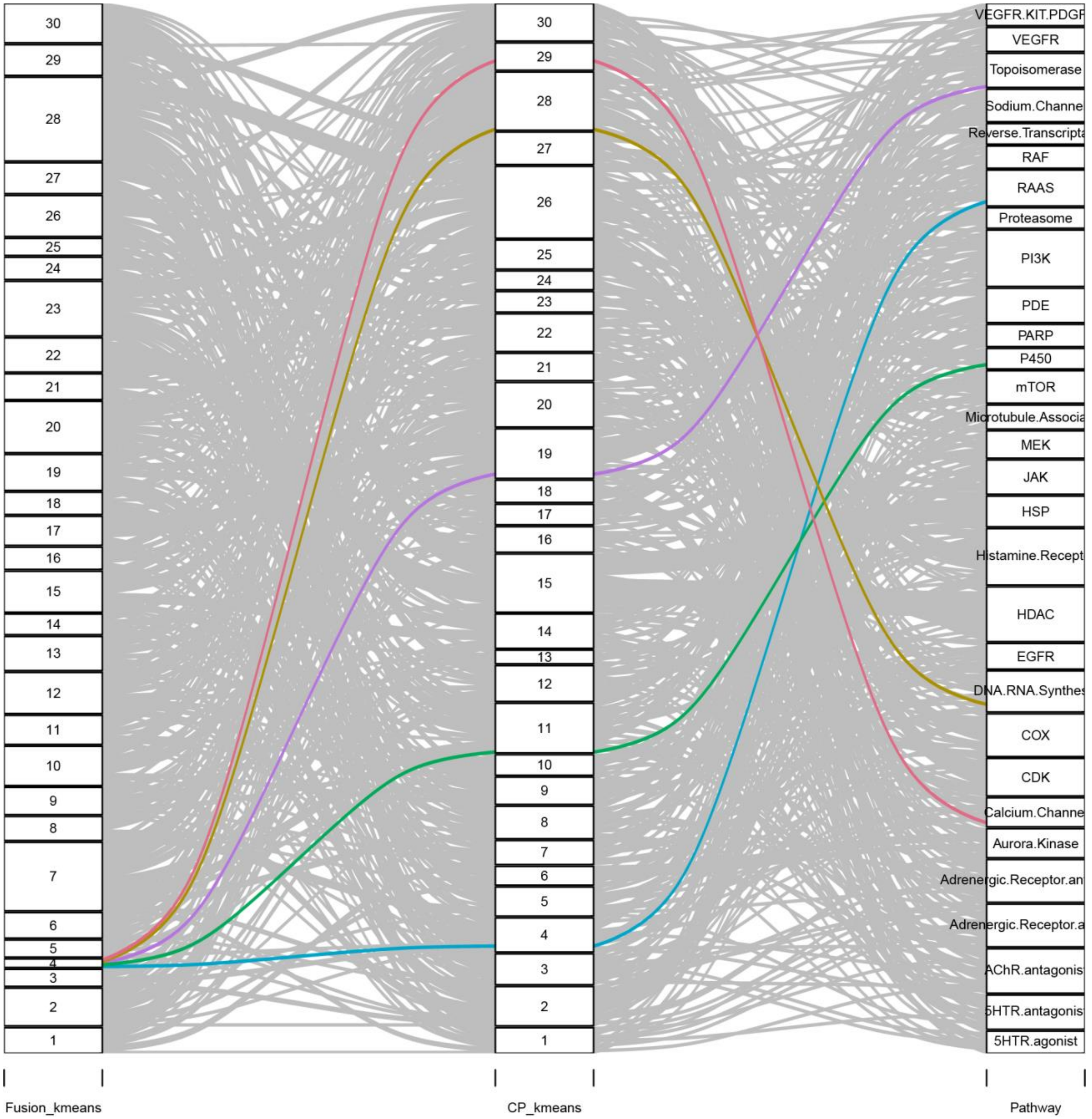

30
29
28
27
26
25
24
23
22
21
20
19
18
17
16
15
14
13
12
11
10
9
8
7
6
5
4
3
2
1
VEGFR.KIT.PDGF
VEGFR
Topoisomerase
Sodium.Channe
Reverse.Transcripta
RAF
RAAS
Proteasome
PI3K
PDE
PARP
P450
mTOR
Microtubule.Associa
MEK
JAK
HSP
Histamine.Recept
HDAC
EGFR
DNA.RNA.Synthes
COX
CDK
Calcium.Channe
Aurora.Kinase
Adrenergic.Receptor.an
Adrenergic.Receptor.a
AChR.antagonis
5HTR.antagonis
5HTR.agonist
Fusion_kmeans
CP_kmeans
Pathway

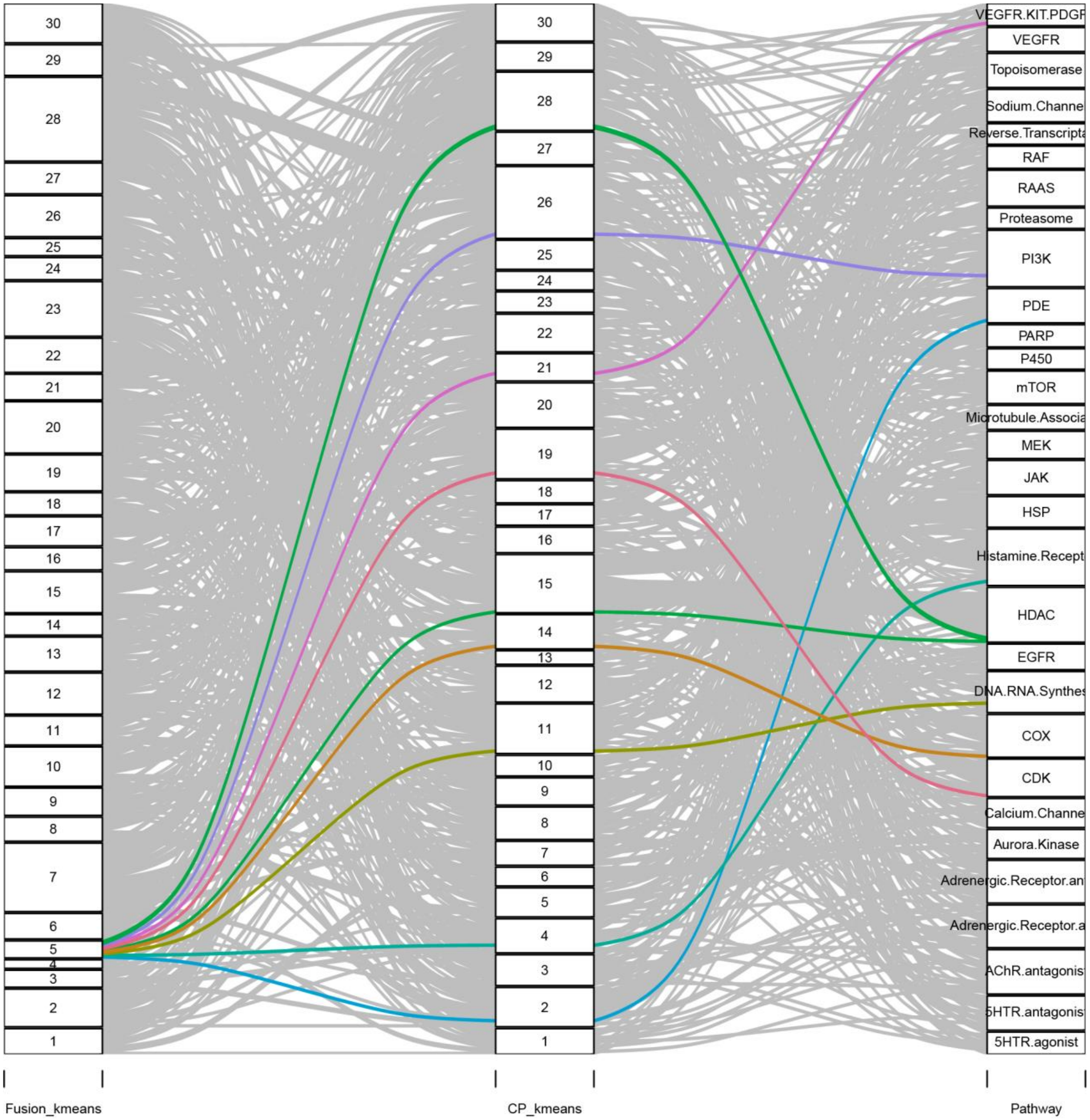
Fusion_kmeans
CP_kmeans
Pathway
30
29
28
27
26
25
24
23
22
21
20
19
18
17
16
15
14
13
12
11
10
9
8
7
6
5
4
3
2
1
VEGFR.KIT.PDGF
VEGFR
Topoisomerase
Sodium.Channe
Reverse.Transcript
RAF
RAAS
Proteasome
PI3K
PDE
PARP
P450
mTOR
Microtubule.Associa
MEK
JAK
HSP
Histamine.Recept
HDAC
EGFR
DNA.RNA.Synthes
COX
CDK
Calcium.Channe
Aurora.Kinase
Adrenergic.Receptor.an
Adrenergic.Receptor.a
AChR.antagonis
5HTR.antagonis
5HTR.agonist

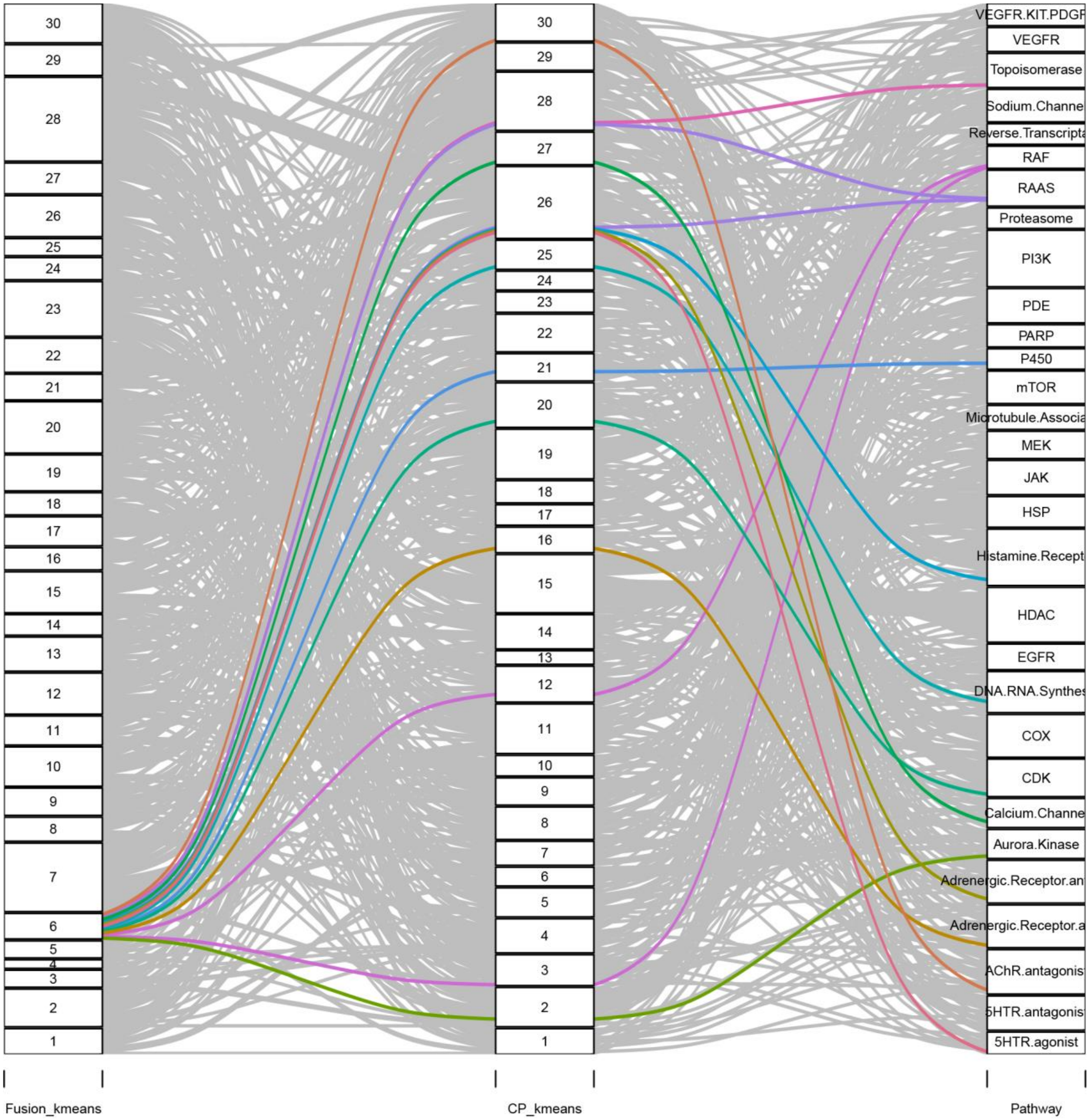
30
29
28
27
26
25
24
23
22
21
20
19
18
17
16
15
14
13
12
11
10
9
8
7
6
5
4
3
2
1
VEGFR.KIT.PDGF
VEGFR
Topoisomerase
Sodium.Channe
Reverse.Transcripta
RAF
RAAS
Proteasome
PI3K
PDE
PARP
P450
mTOR
Microtubule.Associa
MEK
JAK
HSP
Histamine.Recept
HDAC
EGFR
DNA.RNA.Synthes
COX
CDK
Calcium.Channe
Aurora.Kinase
Adrenergic.Receptor.an
Adrenergic.Receptor.a
AChR.antagonis
5HTR.antagonis
5HTR.agonist
Fusion_kmeans
CP_kmeans
Pathway

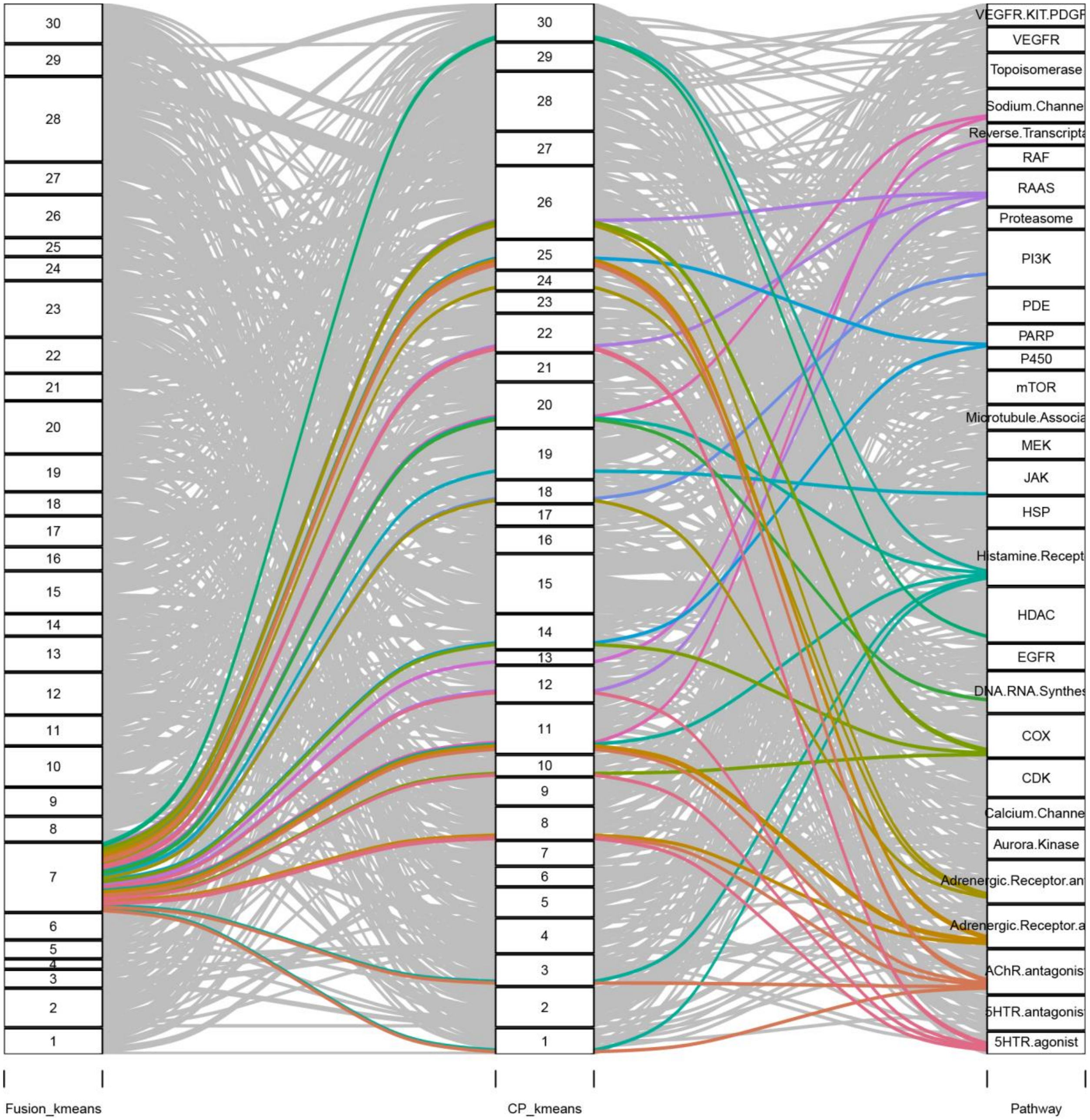
30
29
28
27
26
25
24
23
22
21
20
19
18
17
16
15
14
13
12
11
10
9
8
7
6
5
4
3
2
1
VEGFR.KIT.PDGF
VEGFR
Topoisomerase
Sodium.Channe
Reverse.Transcripta
RAF
RAAS
Proteasome
PI3K
PDE
PARP
P450
mTOR
Microtubule.Associa
MEK
JAK
HSP
Histamine.Recept
HDAC
EGFR
DNA.RNA.Synthes
COX
CDK
Calcium.Channe
Aurora.Kinase
Adrenergic.Receptor.an
Adrenergic.Receptor.a
AChR.antagonis
5HTR.antagonis
5HTR.agonist
Fusion_kmeans
CP_kmeans
Pathway

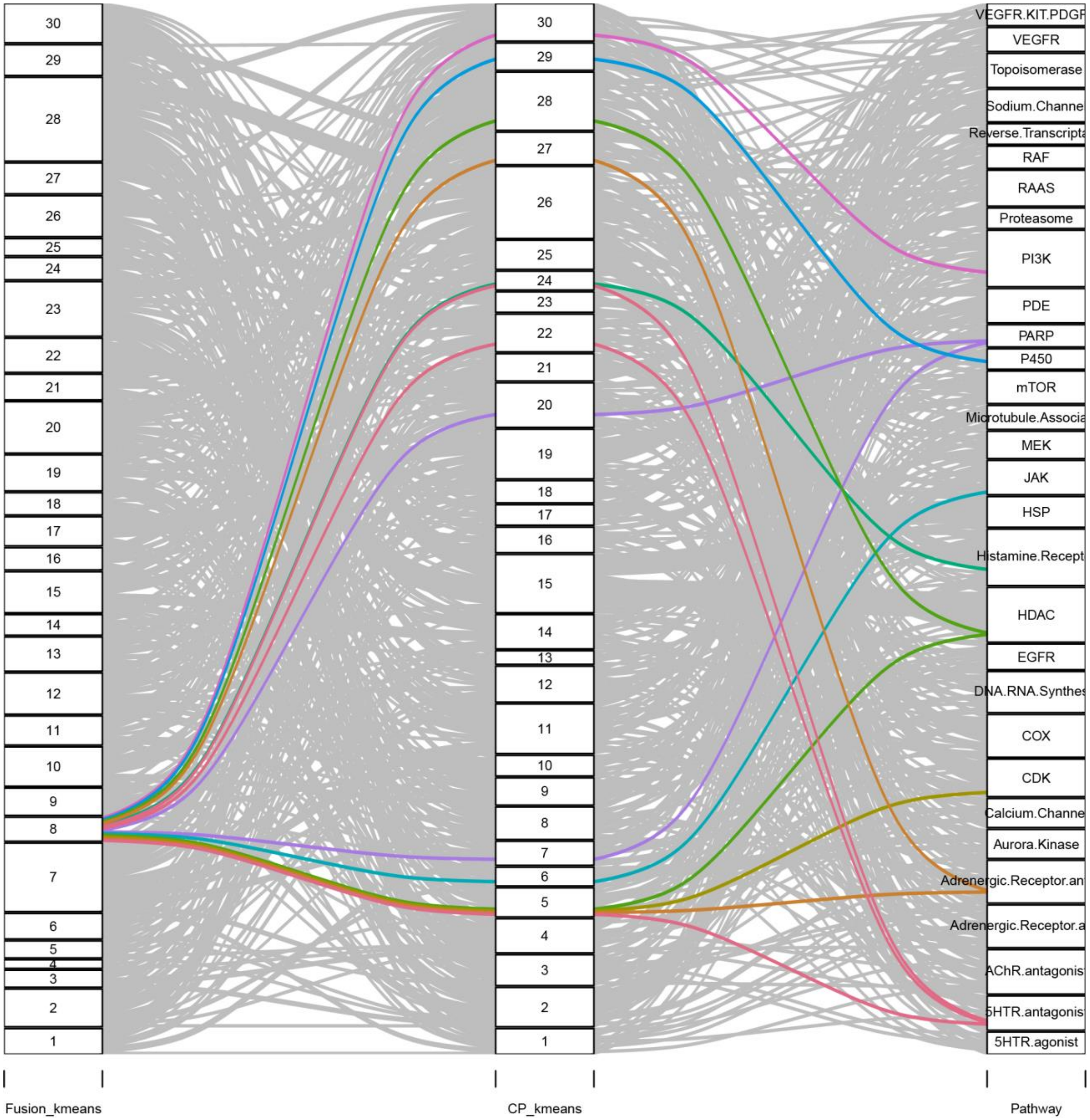
Fusion_kmeans
CP_kmeans
Pathway
30
29
28
27
26
25
24
23
22
21
20
19
18
17
16
15
14
13
12
11
10
9
8
7
6
5
4
3
2
1
VEGFR.KIT.PDGF
VEGFR
Topoisomerase
Sodium.Channe
Reverse.Transcripta
RAF
RAAS
Proteasome
PI3K
PDE
PARP
P450
mTOR
Microtubule.Associa
MEK
JAK
HSP
Histamine.Recept
HDAC
EGFR
DNA.RNA.Synthes
COX
CDK
Calcium.Channe
Aurora.Kinase
Adrenergic.Receptor.an
Adrenergic.Receptor.a
AChR.antagonis
5HTR.antagonis
5HTR.agonist

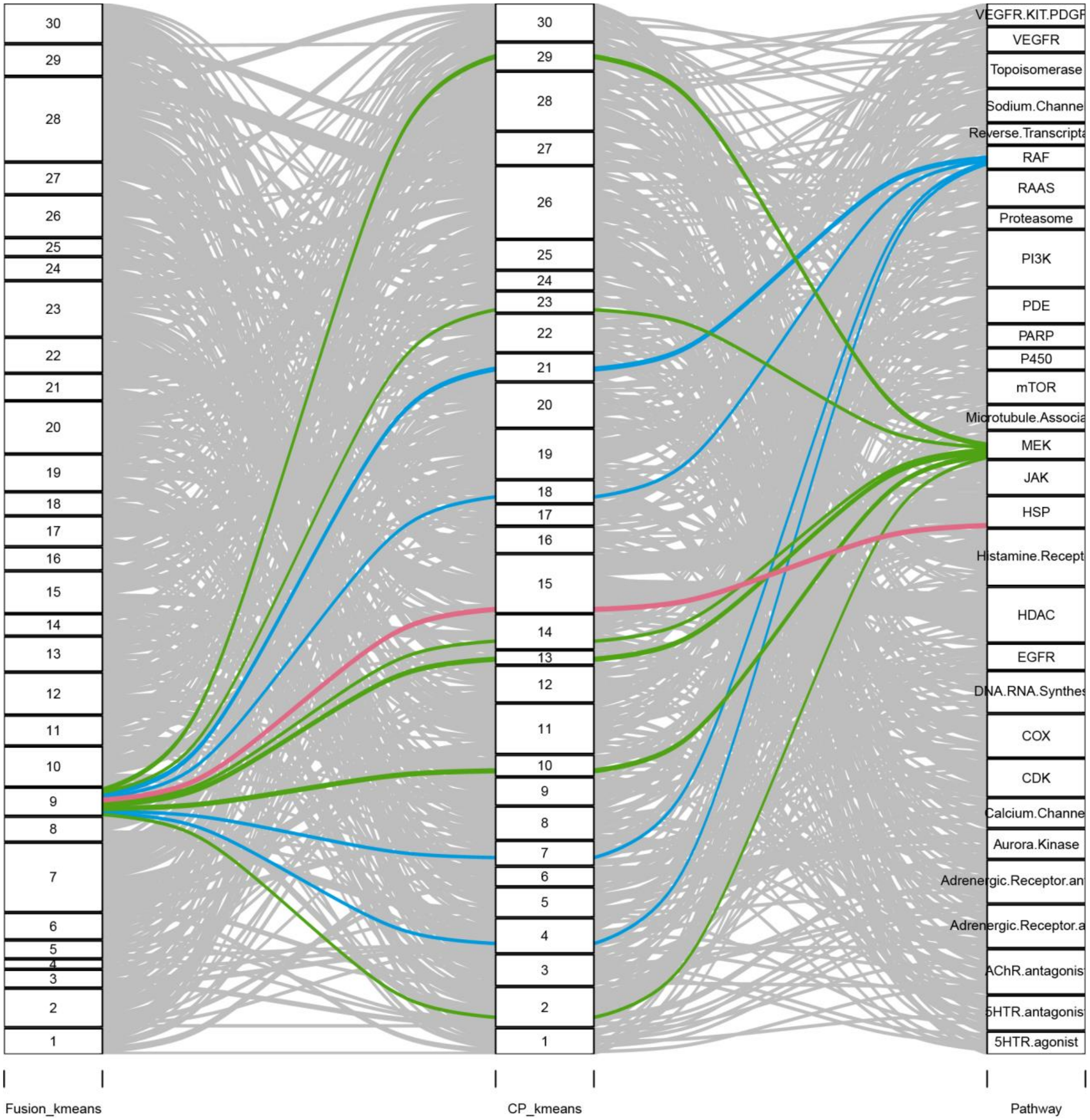

Fusion_kmeans
CP_kmeans
Pathway
VEGFR.KIT.PDGF
VEGFR
Topoisomerase
Sodium.Channe
Reverse.Transcripta
RAF
RAAS
Proteasome
PI3K
PDE
PARP
P450
mTOR
Microtubule.Associa
MEK
JAK
HSP
Histamine.Recept
HDAC
EGFR
DNA.RNA.Synthes
COX
CDK
Calcium.Channe
Aurora.Kinase
Adrenergic.Receptor.an
Adrenergic.Receptor.a
AChR.antagonis
5HTR.antagonis
5HTR.agonist

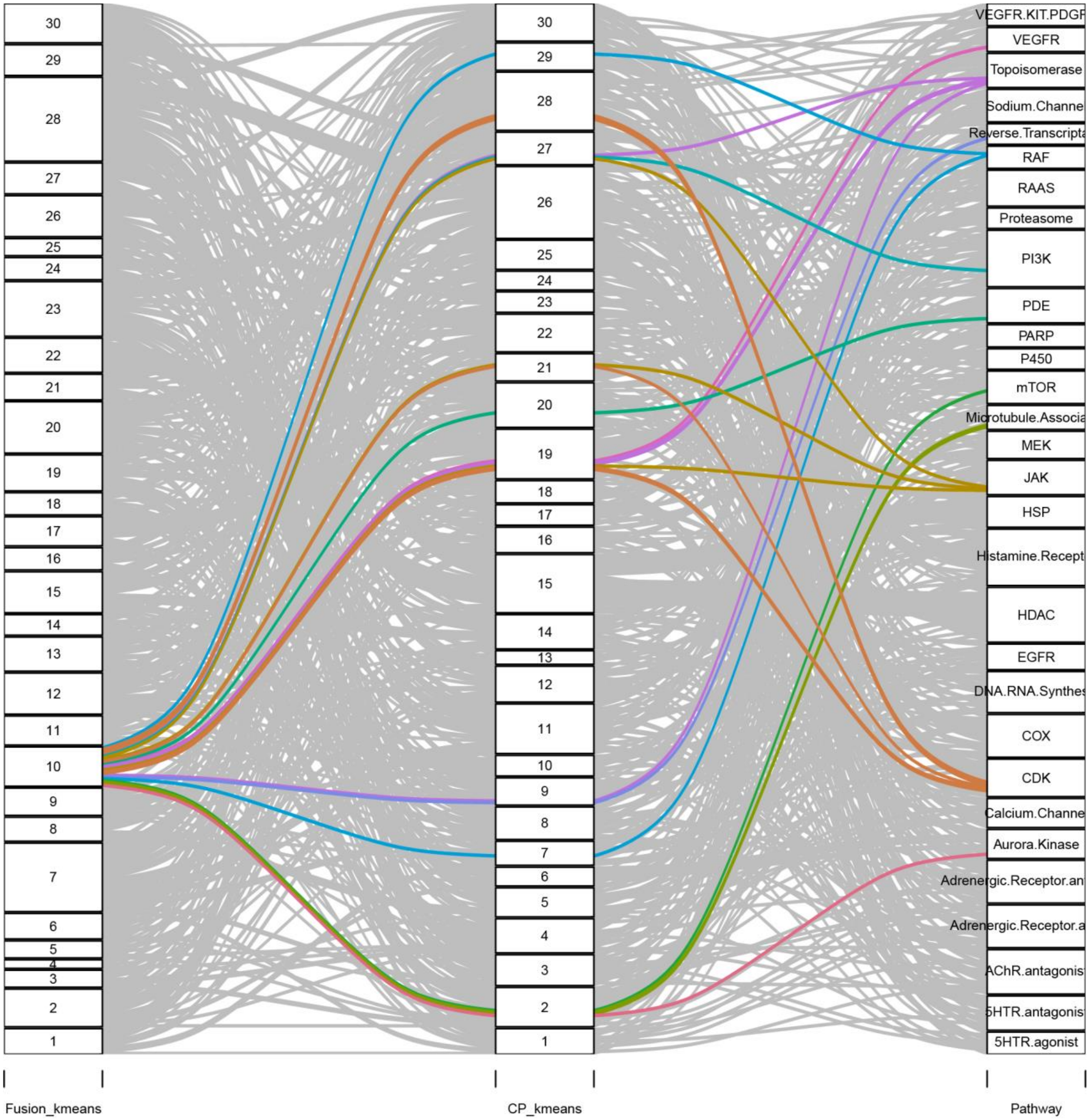

Fusion_kmeans
CP_kmeans
Pathway
30
29
28
27
26
25
24
23
22
21
20
19
18
17
16
15
14
13
12
11
10
9
8
7
6
5
4
3
2
1
VEGFR.KIT.PDGF
VEGFR
Topoisomerase
Sodium.Channe
Reverse.Transcripta
RAF
RAAS
Proteasome
PI3K
PDE
PARP
P450
mTOR
Microtubule.Associa
MEK
JAK
HSP
Histamine.Recept
HDAC
EGFR
DNA.RNA.Synthes
COX
CDK
Calcium.Channe
Aurora.Kinase
Adrenergic.Receptor.an
Adrenergic.Receptor.a
AChR.antagonis
5HTR.antagonis
5HTR.agonist

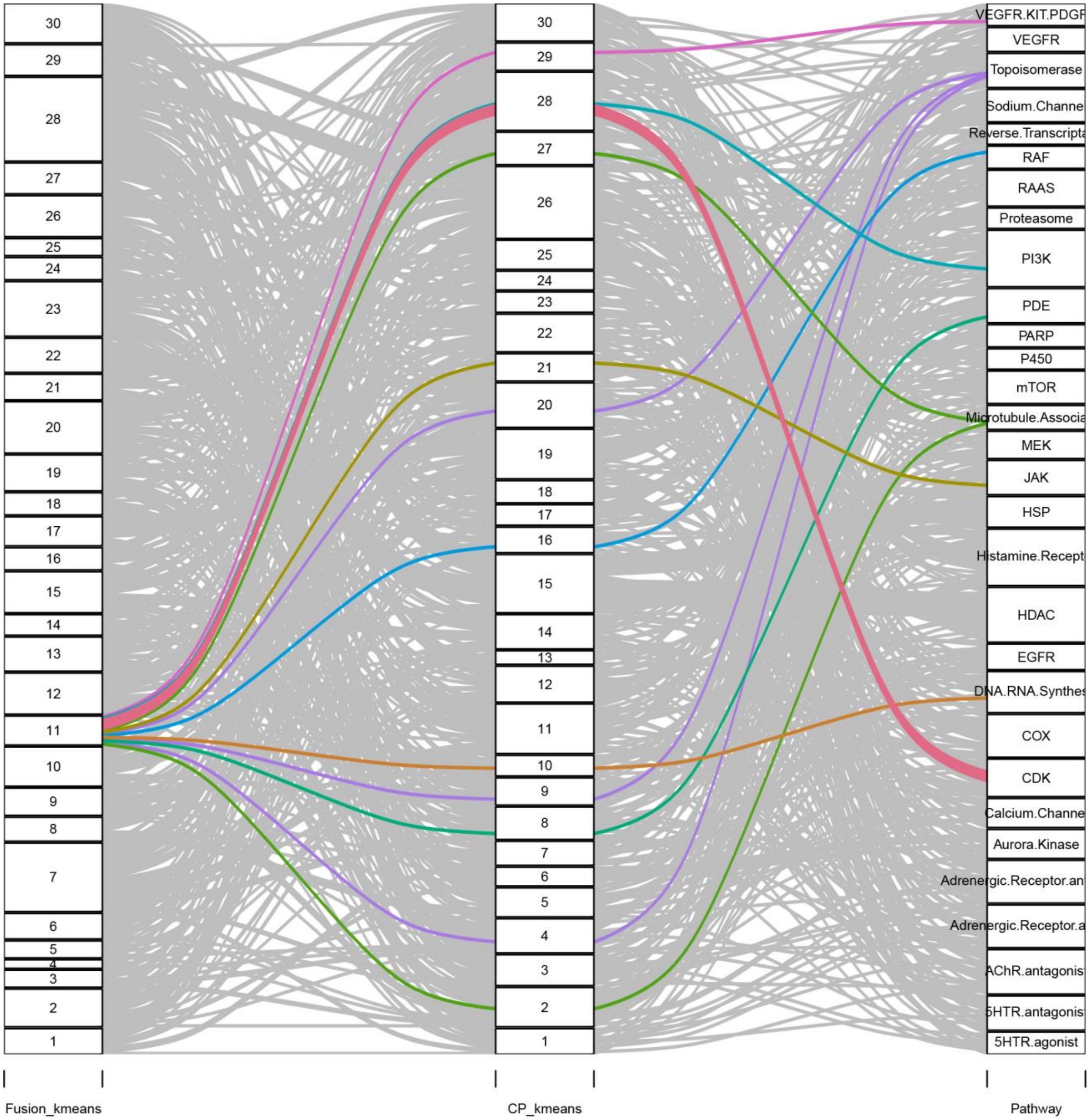
Fusion_kmeans
CP_kmeans
Pathway
VEGFR.KIT.PDGF
VEGFR
Topoisomerase
Sodium.Channe
Reverse.Transcripta
RAF
RAAS
Proteasome
PI3K
PDE
PARP
P450
mTOR
Microtubule.Associa
MEK
JAK
HSP
Histamine.Recept
HDAC
EGFR
DNA.RNA.Synthes
COX
CDK
Calcium.Channe
Aurora.Kinase
Adrenergic.Receptor.an
Adrenergic.Receptor.a
AChR.antagonis
5HTR.antagonis
5HTR.agonist

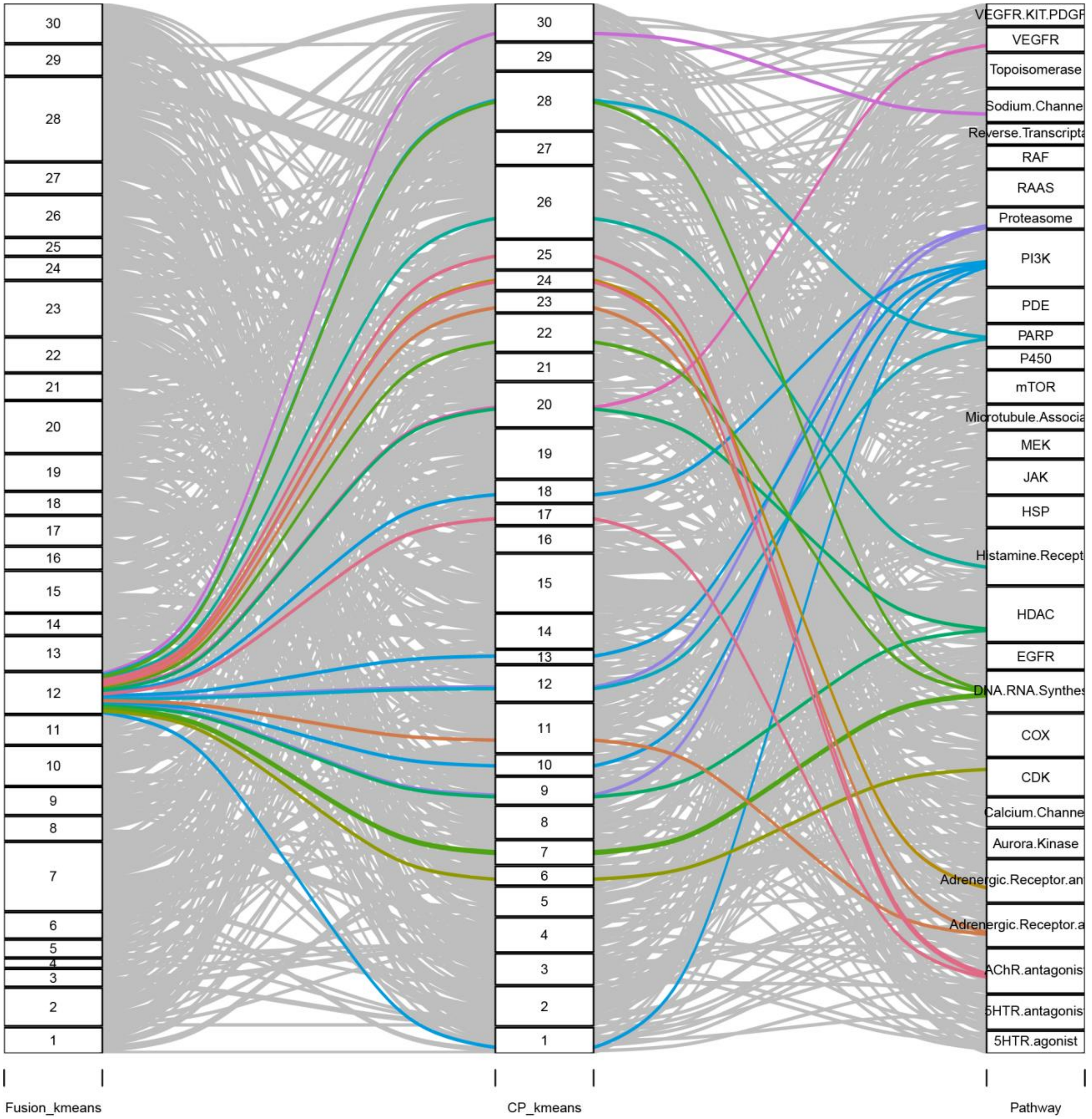
Fusion_kmeans
CP_kmeans
Pathway
30
29
28
27
26
25
24
23
22
21
20
19
18
17
16
15
14
13
12
11
10
9
8
7
6
5
4
3
2
1
VEGFR.KIT.PDGF
VEGFR
Topoisomerase
Sodium.Channe
Reverse.Transcripta
RAF
RAAS
Proteasome
PI3K
PDE
PARP
P450
mTOR
Microtubule.Associa
MEK
JAK
HSP
Histamine.Recept
HDAC
EGFR
DNA.RNA.Synthes
COX
CDK
Calcium.Channe
Aurora.Kinase
Adrenergic.Receptor.an
Adrenergic.Receptor.a
AChR.antagonis
5HTR.antagonis
5HTR.agonist

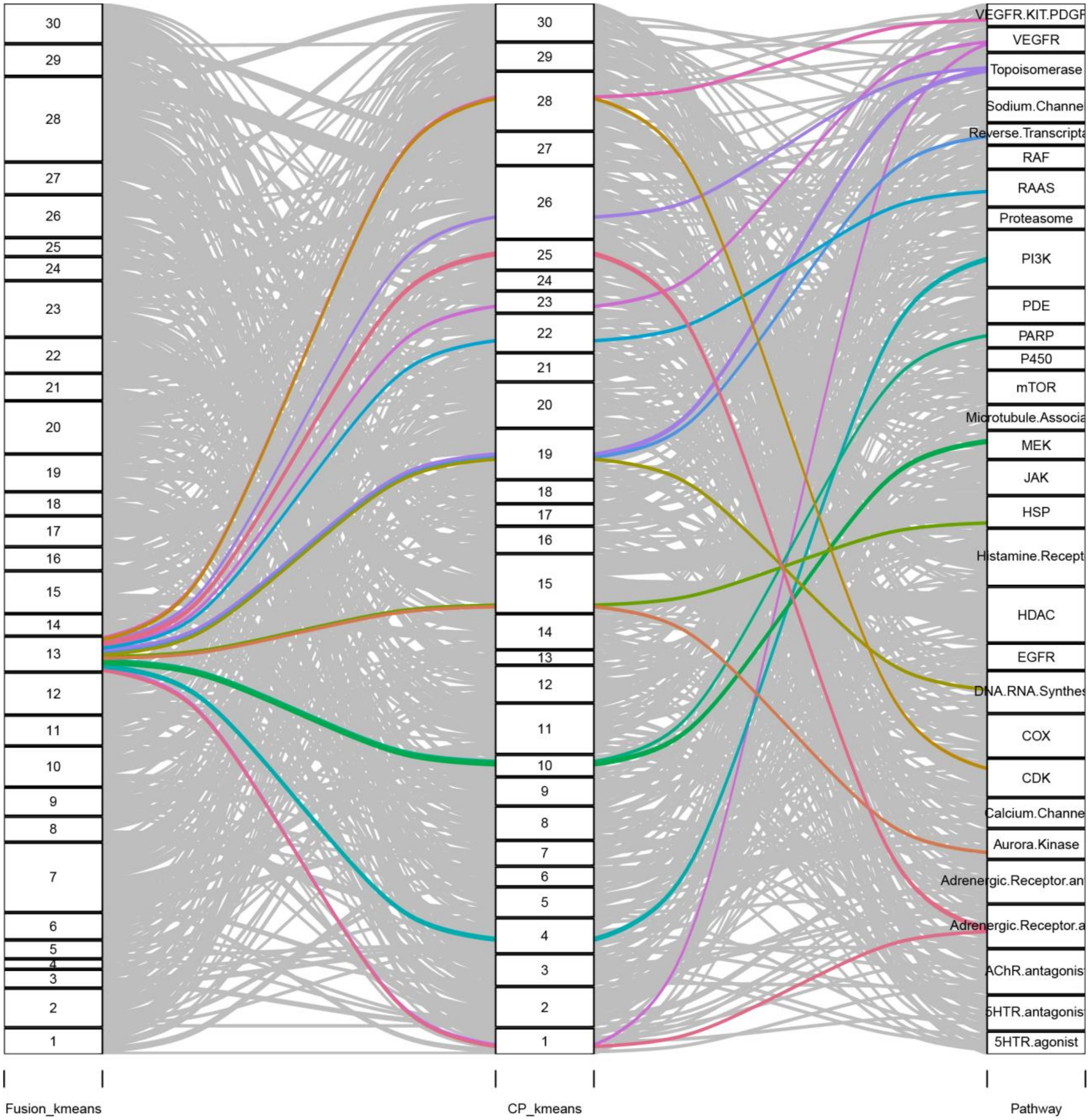
Fusion_kmeans
CP_kmeans
Pathway
VEGFR.KIT.PDGF
VEGFR
Topoisomerase
Sodium.Channe
Reverse.Transcripta
RAF
RAAS
Proteasome
PI3K
PDE
PARP
P450
mTOR
Microtubule.Associa
MEK
JAK
HSP
Histamine.Recept
HDAC
EGFR
DNA.RNA.Synthes
COX
CDK
Calcium.Channe
Aurora.Kinase
Adrenergic.Receptor.an
Adrenergic.Receptor.a
AChR.antagonis
5HTR.antagonis
5HTR.agonist

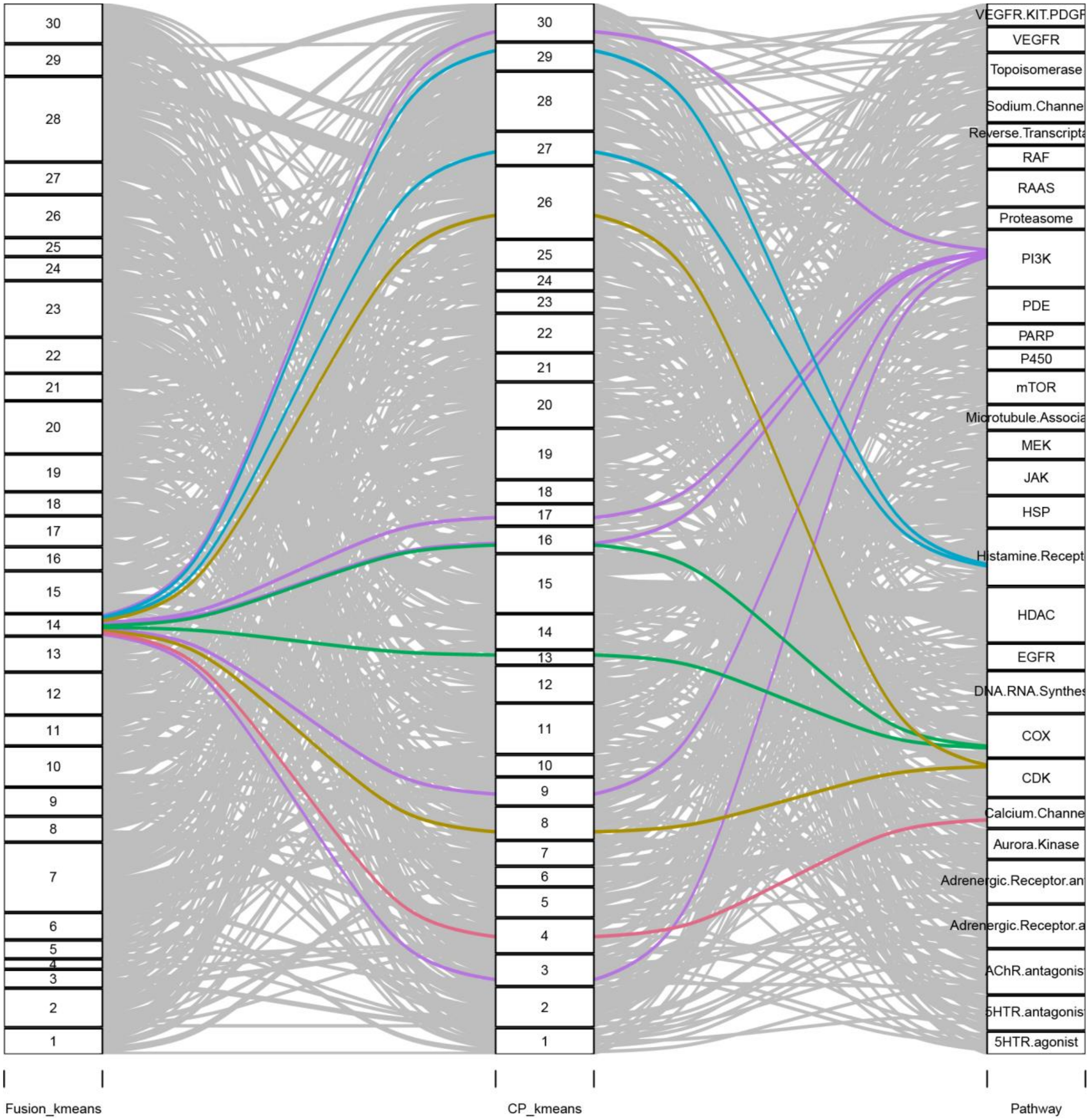
Fusion_kmeans
CP_kmeans
Pathway
VEGFR.KIT.PDGF
VEGFR
Topoisomerase
Sodium.Channe
Reverse.Transcripta
RAF
RAAS
Proteasome
PI3K
PDE
PARP
P450
mTOR
Microtubule.Associa
MEK
JAK
HSP
Histamine.Recept
HDAC
EGFR
DNA.RNA.Synthes
COX
CDK
Calcium.Channe
Aurora.Kinase
Adrenergic.Receptor.an
Adrenergic.Receptor.a
AChR.antagonis
5HTR.antagonis
5HTR.agonist

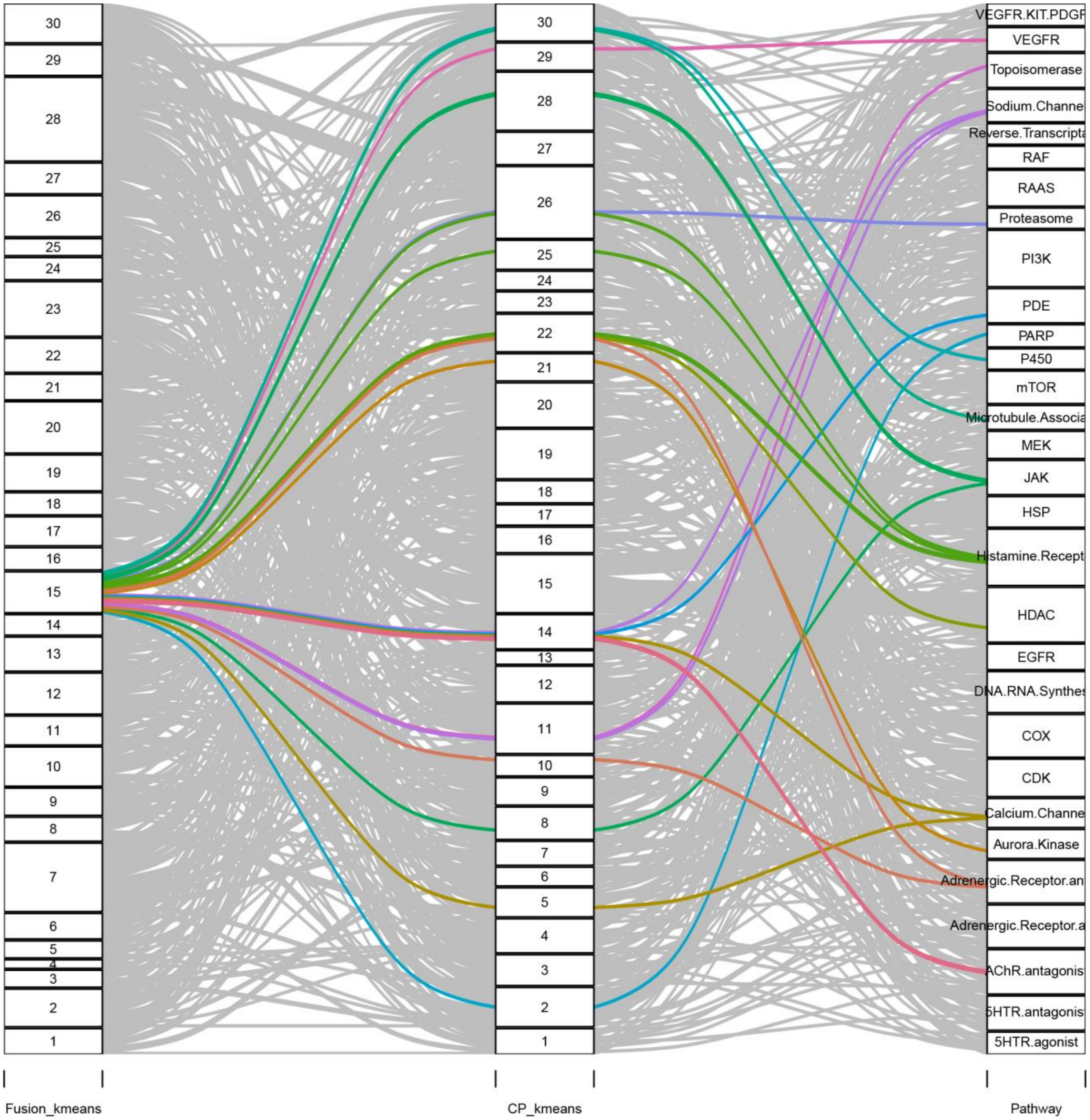
30
29
28
27
26
25
24
23
22
21
20
19
18
17
16
15
14
13
12
11
10
9
8
7
6
5
4
3
2
1
VEGFR.KIT.PDGF
VEGFR
Topoisomerase
Sodium.Channe
Reverse.Transcripta
RAF
RAAS
Proteasome
PI3K
PDE
PARP
P450
mTOR
Microtubule.Associa
MEK
JAK
HSP
Histamine.Recept
HDAC
EGFR
DNA.RNA.Synthes
COX
CDK
Calcium.Channe
Aurora.Kinase
Adrenergic.Receptor.an
Adrenergic.Receptor.a
AChR.antagonis
5HTR.antagonis
5HTR.agonist
Fusion_kmeans
CP_kmeans
Pathway

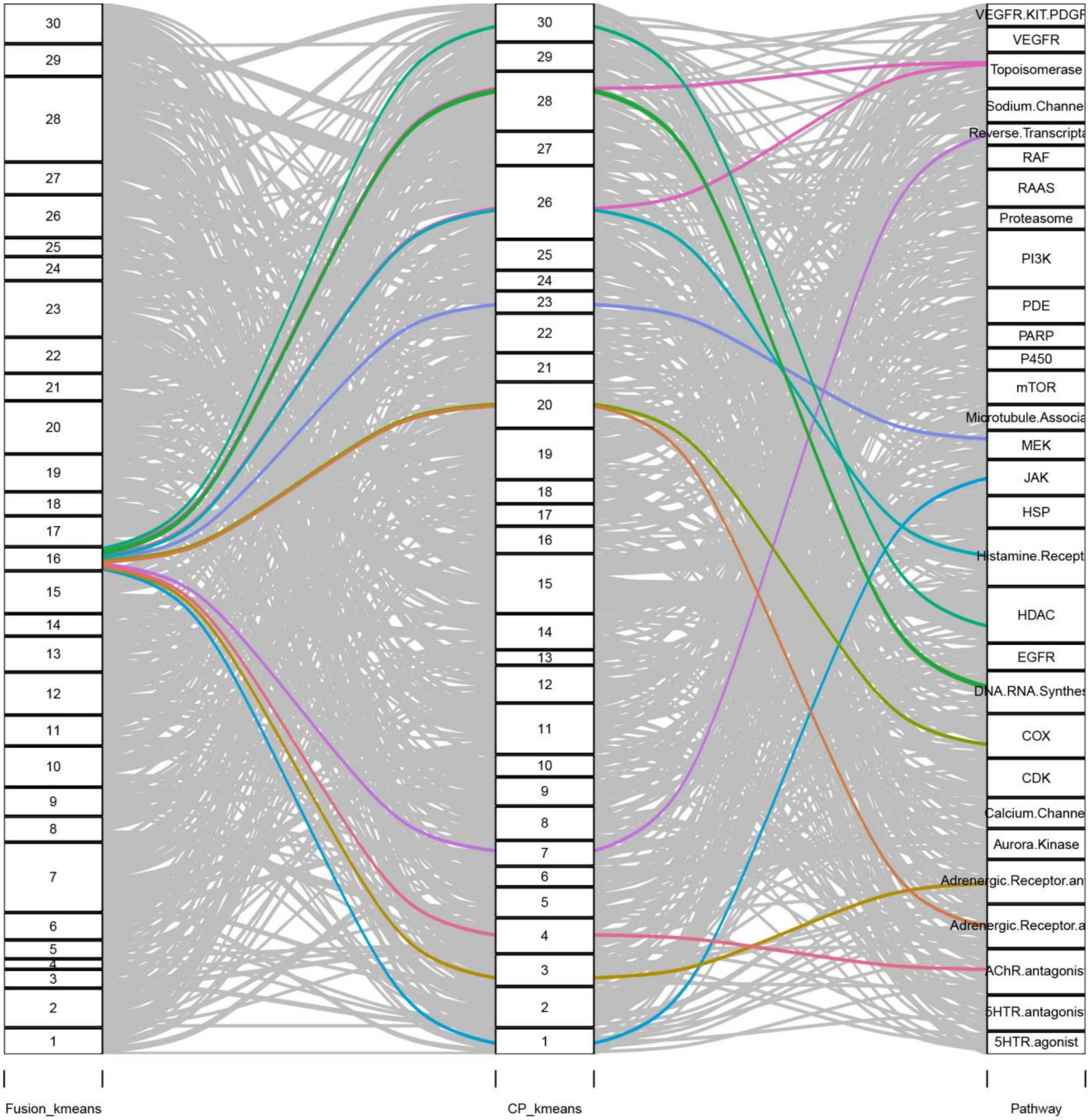

Fusion_kmeans
CP_kmeans
Pathway
VEGFR.KIT.PDGF
VEGFR
Topoisomerase
Sodium.Channe
Reverse.Transcripta
RAF
RAAS
Proteasome
PI3K
PDE
PARP
P450
mTOR
Microtubule.Associa
MEK
JAK
HSP
Histamine.Recept
HDAC
EGFR
DNA.RNA.Synthes
COX
CDK
Calcium.Channe
Aurora.Kinase
Adrenergic.Receptor.an
Adrenergic.Receptor.a
AChR.antagonis
5HTR.antagonis
5HTR.agonist

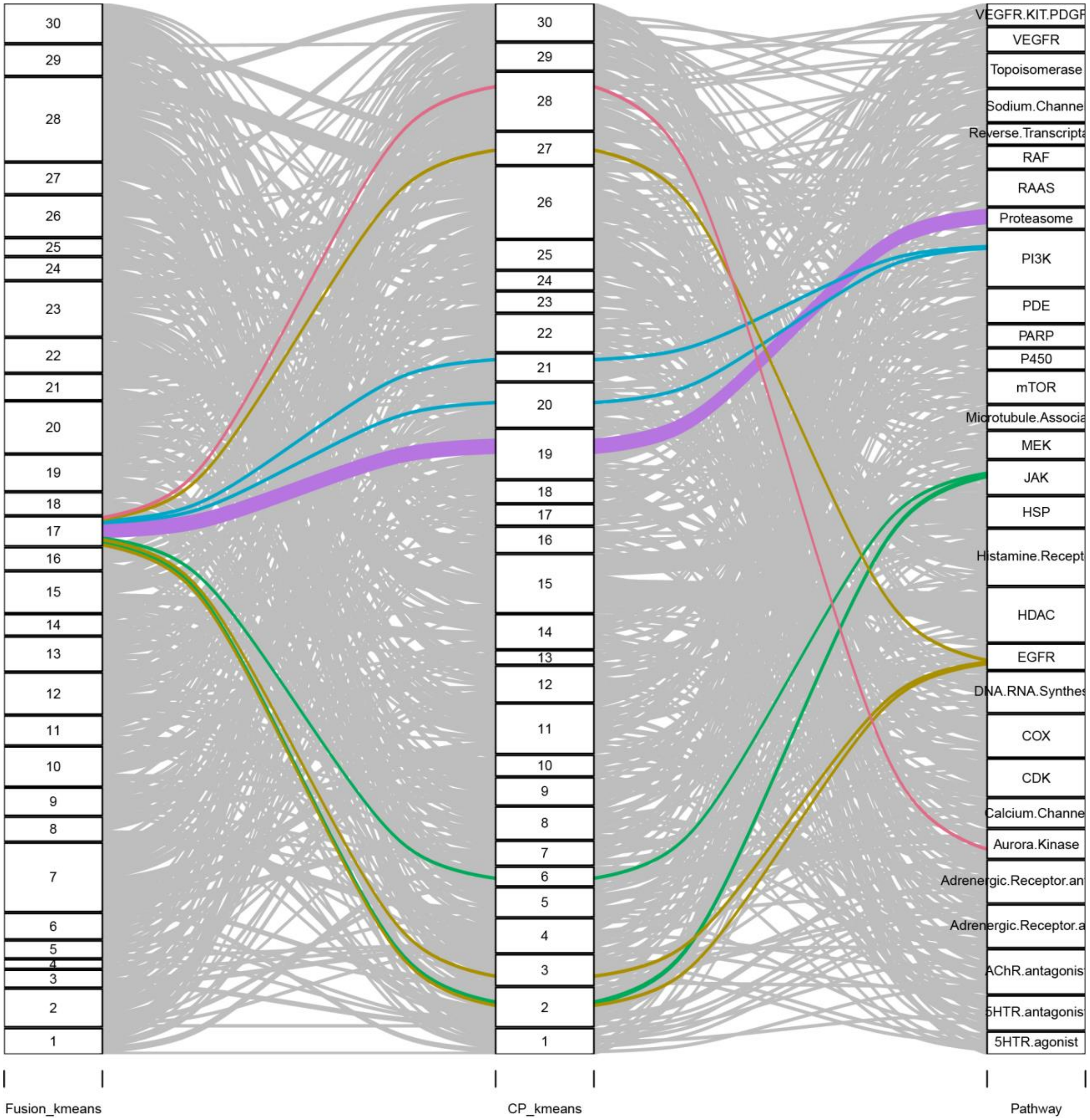
Fusion_kmeans
CP_kmeans
Pathway
VEGFR.KIT.PDGF
VEGFR
Topoisomerase
Sodium.Channe
Reverse.Transcripta
RAF
RAAS
Proteasome
PI3K
PDE
PARP
P450
mTOR
Microtubule.Associa
MEK
JAK
HSP
Histamine.Recept
HDAC
EGFR
DNA.RNA.Synthes
COX
CDK
Calcium.Channe
Aurora.Kinase
Adrenergic.Receptor.an
Adrenergic.Receptor.a
AChR.antagonis
5HTR.antagonis
5HTR.agonist

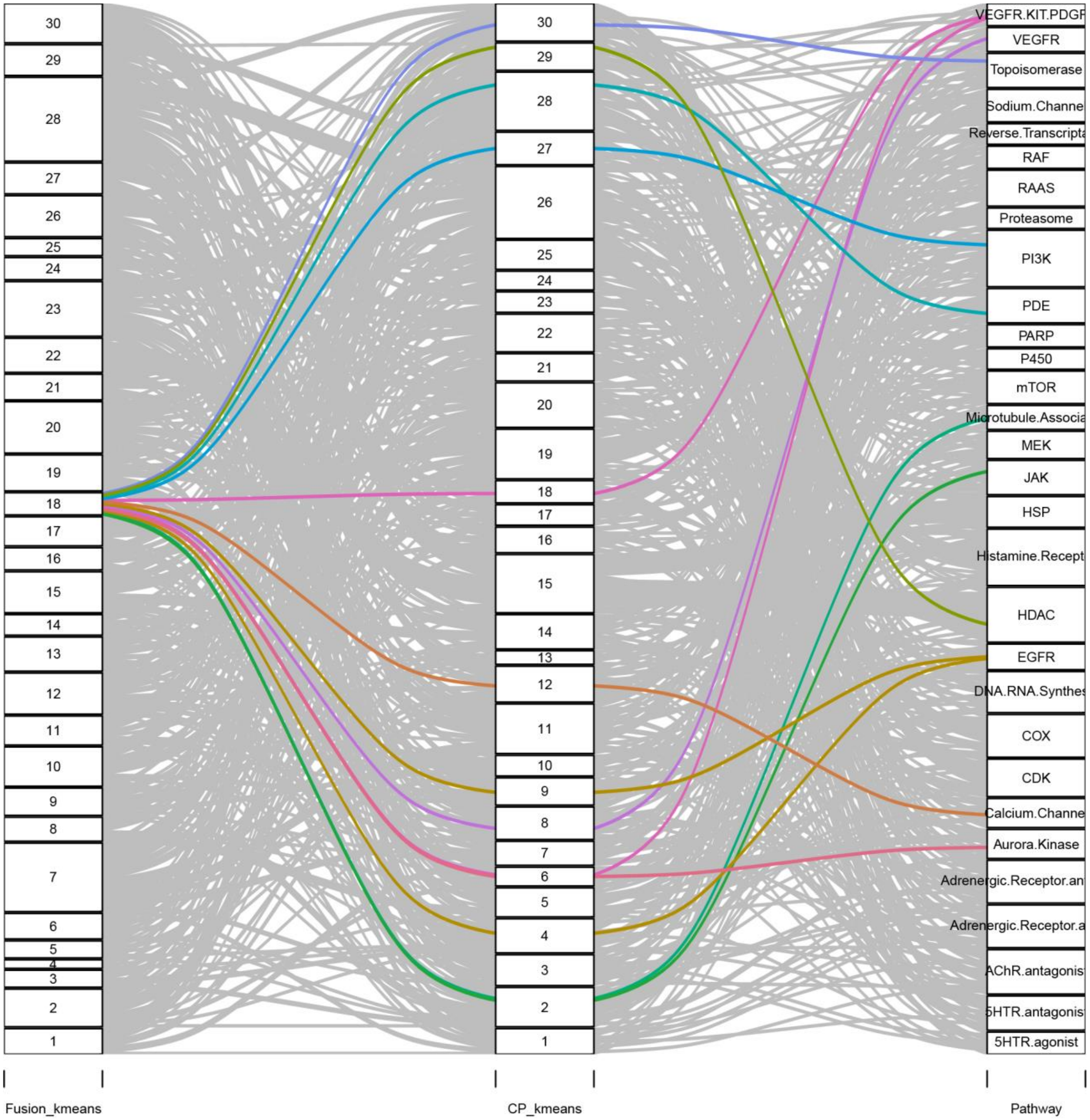
30
29
28
27
26
25
24
23
22
21
20
19
18
17
16
15
14
13
12
11
10
9
8
7
6
5
4
3
2
1
VEGFR.KIT.PDGF
VEGFR
Topoisomerase
Sodium.Channe
Reverse.Transcripta
RAF
RAAS
Proteasome
PI3K
PDE
PARP
P450
mTOR
Microtubule.Associa
MEK
JAK
HSP
Histamine.Recept
HDAC
EGFR
DNA.RNA.Synthes
COX
CDK
Calcium.Channe
Aurora.Kinase
Adrenergic.Receptor.an
Adrenergic.Receptor.a
AChR.antagonis
5HTR.antagonis
5HTR.agonist
Fusion_kmeans
CP_kmeans
Pathway

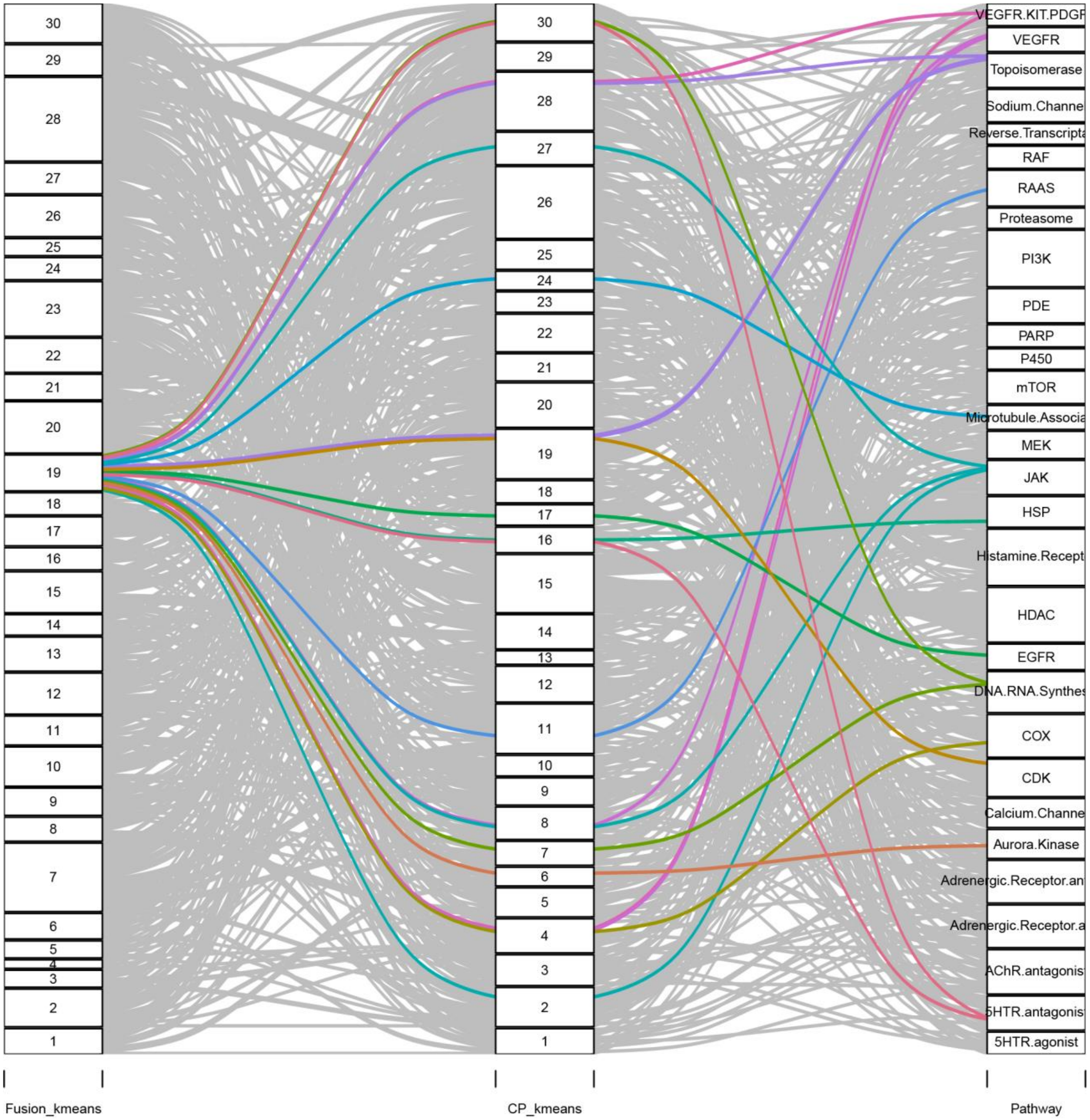
Fusion_kmeans
CP_kmeans
Pathway
VEGFR.KIT.PDGF
VEGFR
Topoisomerase
Sodium.Channe
Reverse.Transcripta
RAF
RAAS
Proteasome
PI3K
PDE
PARP
P450
mTOR
Microtubule.Associa
MEK
JAK
HSP
Histamine.Recept
HDAC
EGFR
DNA.RNA.Synthes
COX
CDK
Calcium.Channe
Aurora.Kinase
Adrenergic.Receptor.an
Adrenergic.Receptor.a
AChR.antagonis
5HTR.antagonis
5HTR.agonist

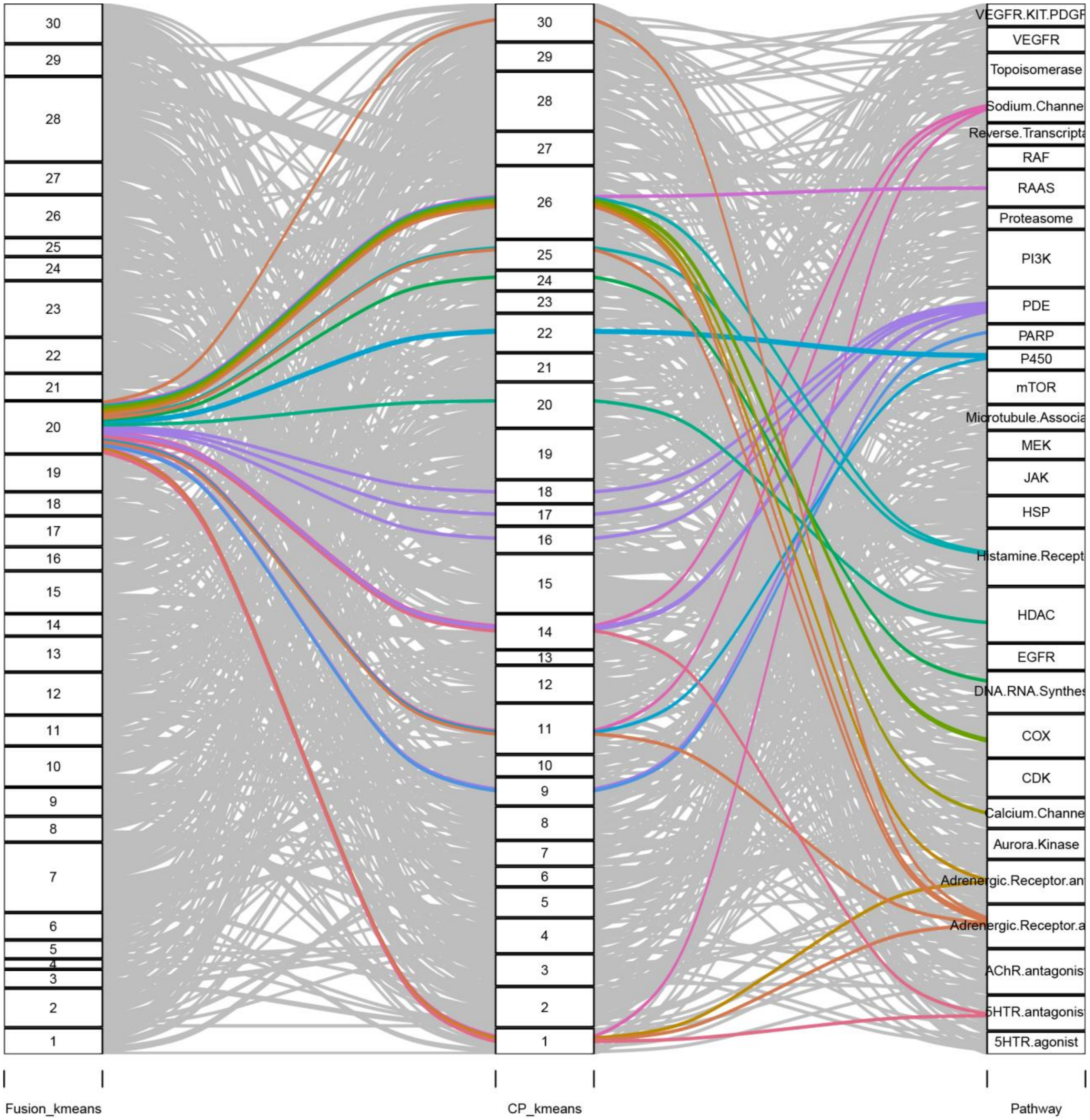
30
29
28
27
26
25
24
23
22
21
20
19
18
17
16
15
14
13
12
11
10
9
8
7
6
5
4
3
2
1
VEGFR.KIT.PDGF
VEGFR
Topoisomerase
Sodium.Channe
Reverse.Transcripta
RAF
RAAS
Proteasome
PI3K
PDE
PARP
P450
mTOR
Microtubule.Associa
MEK
JAK
HSP
Histamine.Recept
HDAC
EGFR
DNA.RNA.Synthes
COX
CDK
Calcium.Channe
Aurora.Kinase
Adrenergic.Receptor.an
Adrenergic.Receptor.a
AChR.antagonis
5HTR.antagonis
5HTR.agonist
Fusion_kmeans
CP_kmeans
Pathway

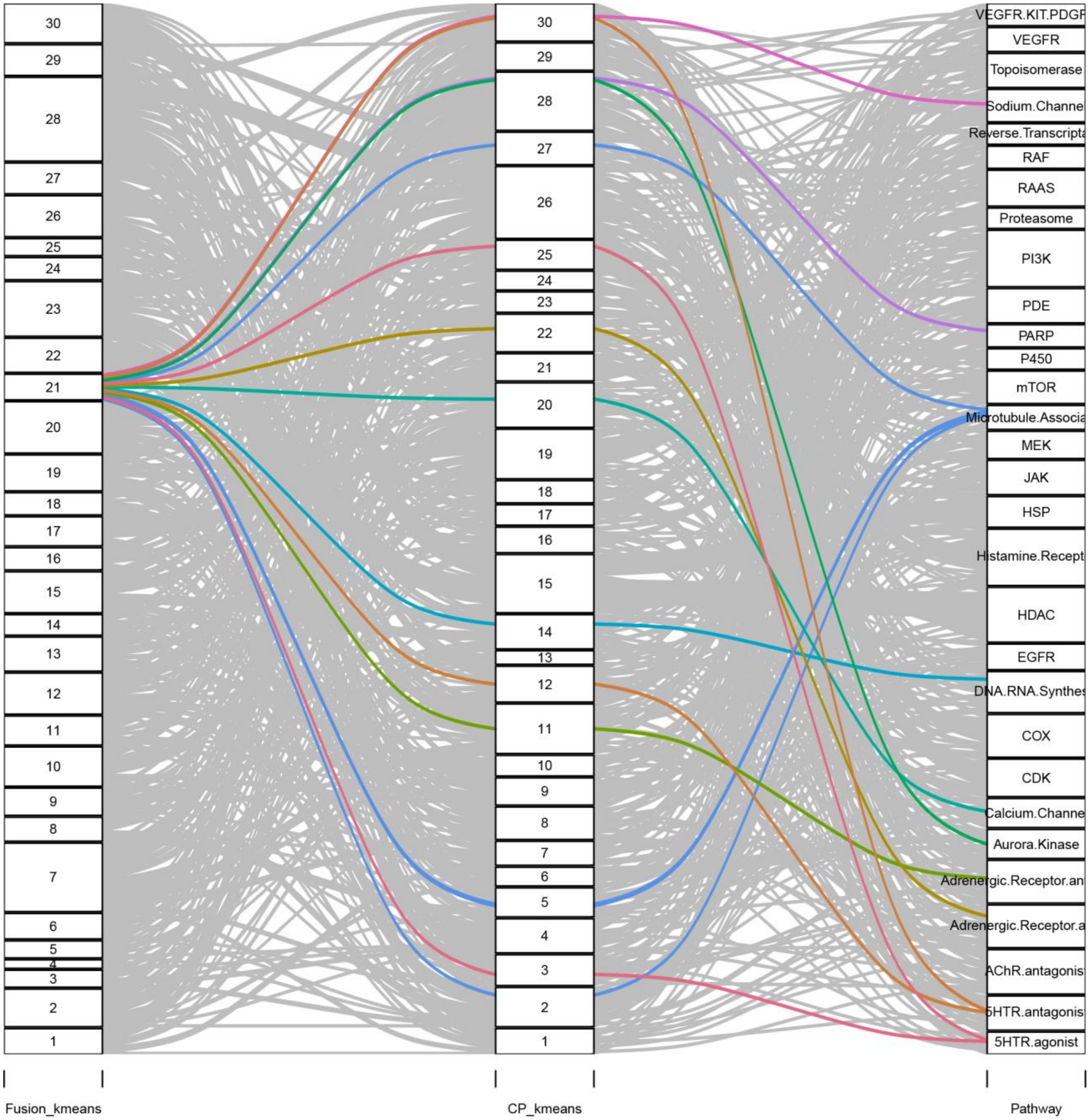

30
29
28
27
26
25
24
23
22
21
20
19
18
17
16
15
14
13
12
11
10
9
8
7
6
5
4
3
2
1
Fusion_kmeans
CP_kmeans
VEGFR.KIT.PDGF
VEGFR
Topoisomerase
Sodium.Channe
Reverse.Transcripta
RAF
RAAS
Proteasome
PI3K
PDE
PARP
P450
mTOR
Microtubule.Associa
MEK
JAK
HSP
Histamine.Recept
HDAC
EGFR
DNA.RNA.Synthes
COX
CDK
Calcium.Channe
Aurora.Kinase
Adrenergic.Receptor.an
Adrenergic.Receptor.a
AChR.antagonis
5HTR.antagonis
5HTR.agonist
Pathway

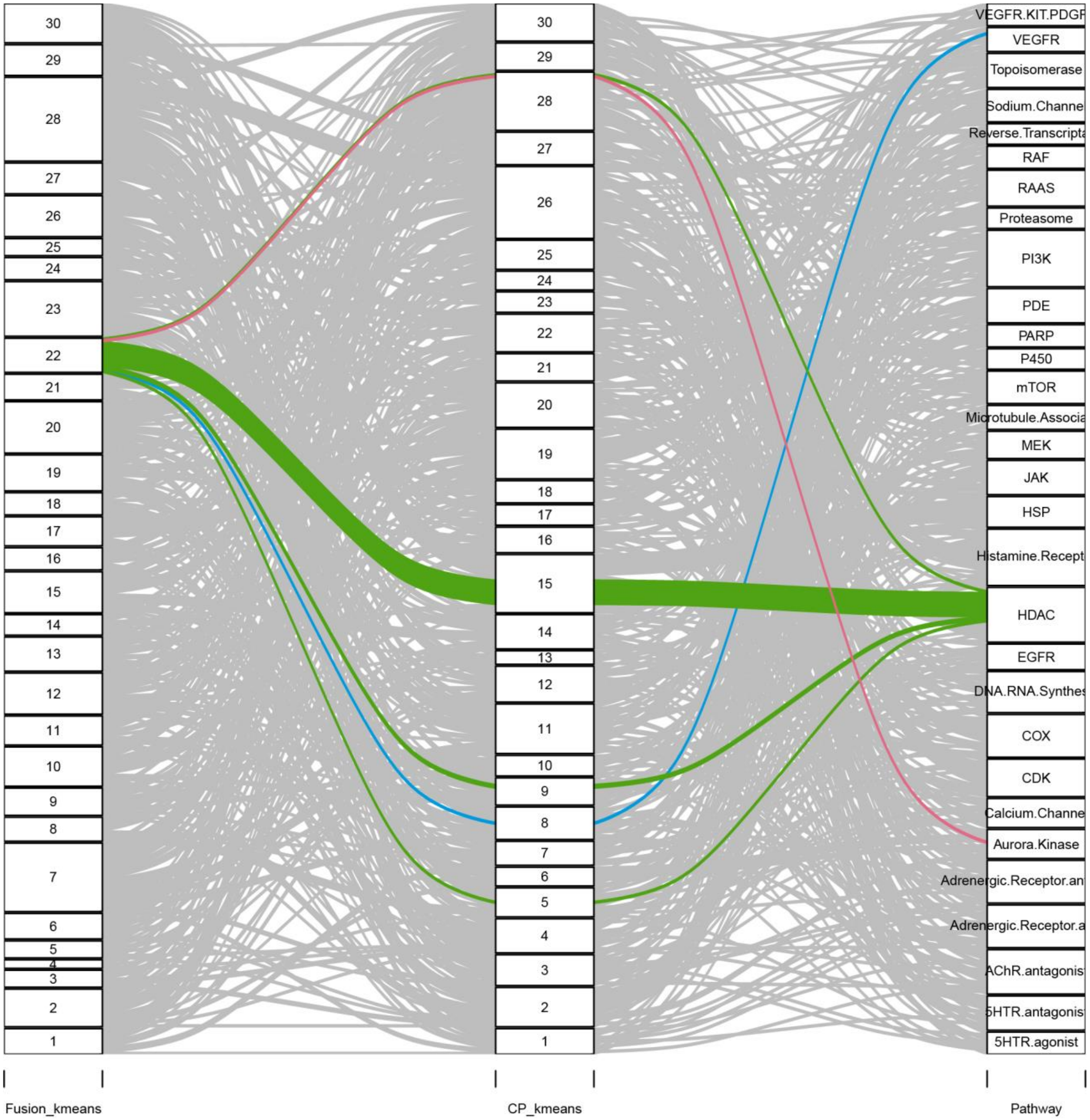
Fusion_kmeans
CP_kmeans
Pathway
VEGFR.KIT.PDGF
VEGFR
Topoisomerase
Sodium.Channe
Reverse.Transcripta
RAF
RAAS
Proteasome
PI3K
PDE
PARP
P450
mTOR
Microtubule.Associa
MEK
JAK
HSP
Histamine.Recept
HDAC
EGFR
DNA.RNA.Synthes
COX
CDK
Calcium.Channe
Aurora.Kinase
Adrenergic.Receptor.an
Adrenergic.Receptor.a
AChR.antagonis
5HTR.antagonis
5HTR.agonist

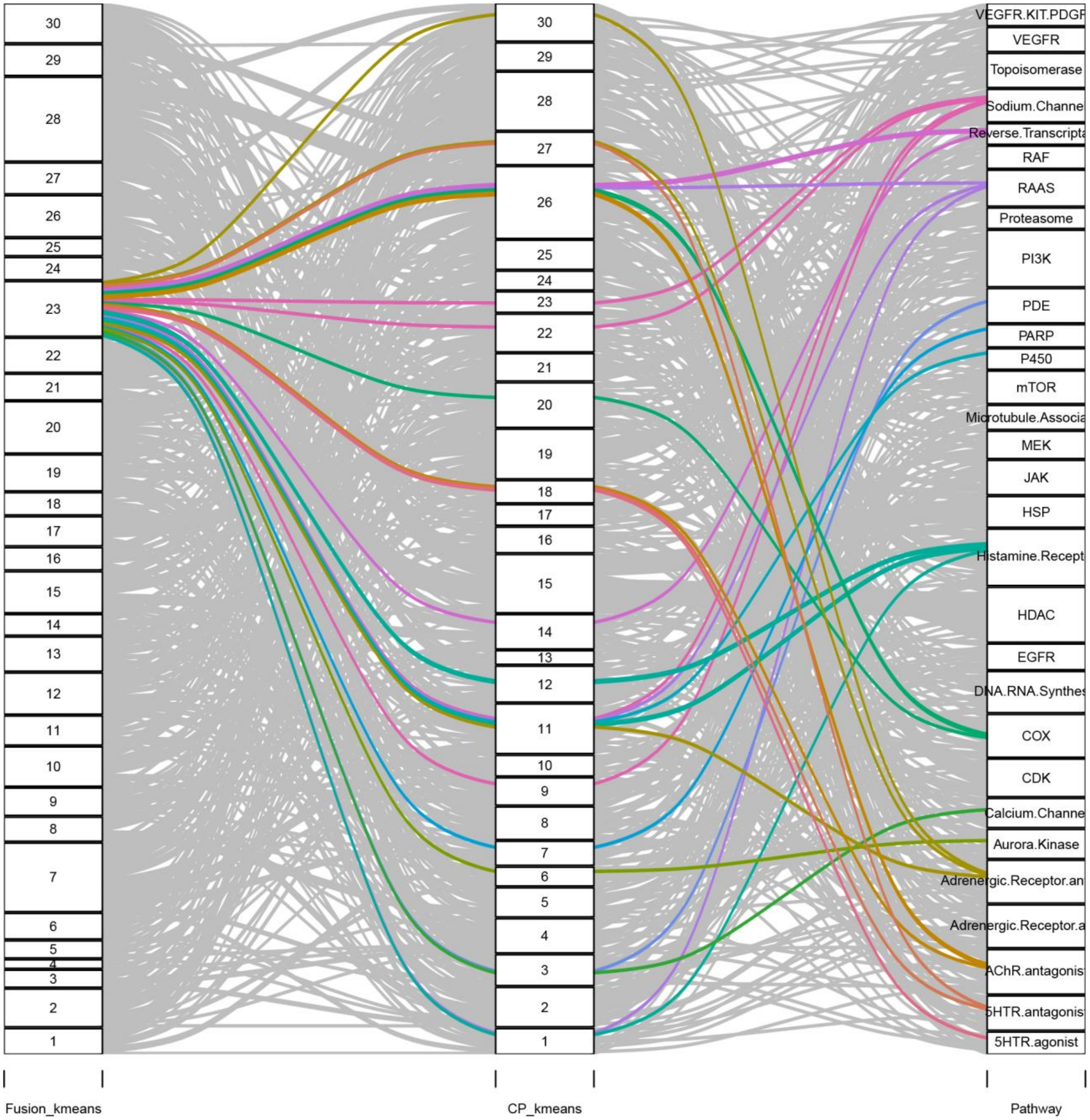
Fusion_kmeans
CP_kmeans
Pathway
VEGFR.KIT.PDGF
VEGFR
Topoisomerase
Sodium.Channe
Reverse.Transcripta
RAF
RAAS
Proteasome
PI3K
PDE
PARP
P450
mTOR
Microtubule.Associa
MEK
JAK
HSP
Histamine.Recept
HDAC
EGFR
DNA.RNA.Synthes
COX
CDK
Calcium.Channe
Aurora.Kinase
Adrenergic.Receptor.an
Adrenergic.Receptor.a
AChR.antagonist
5HTR.antagonist
5HTR.agonist

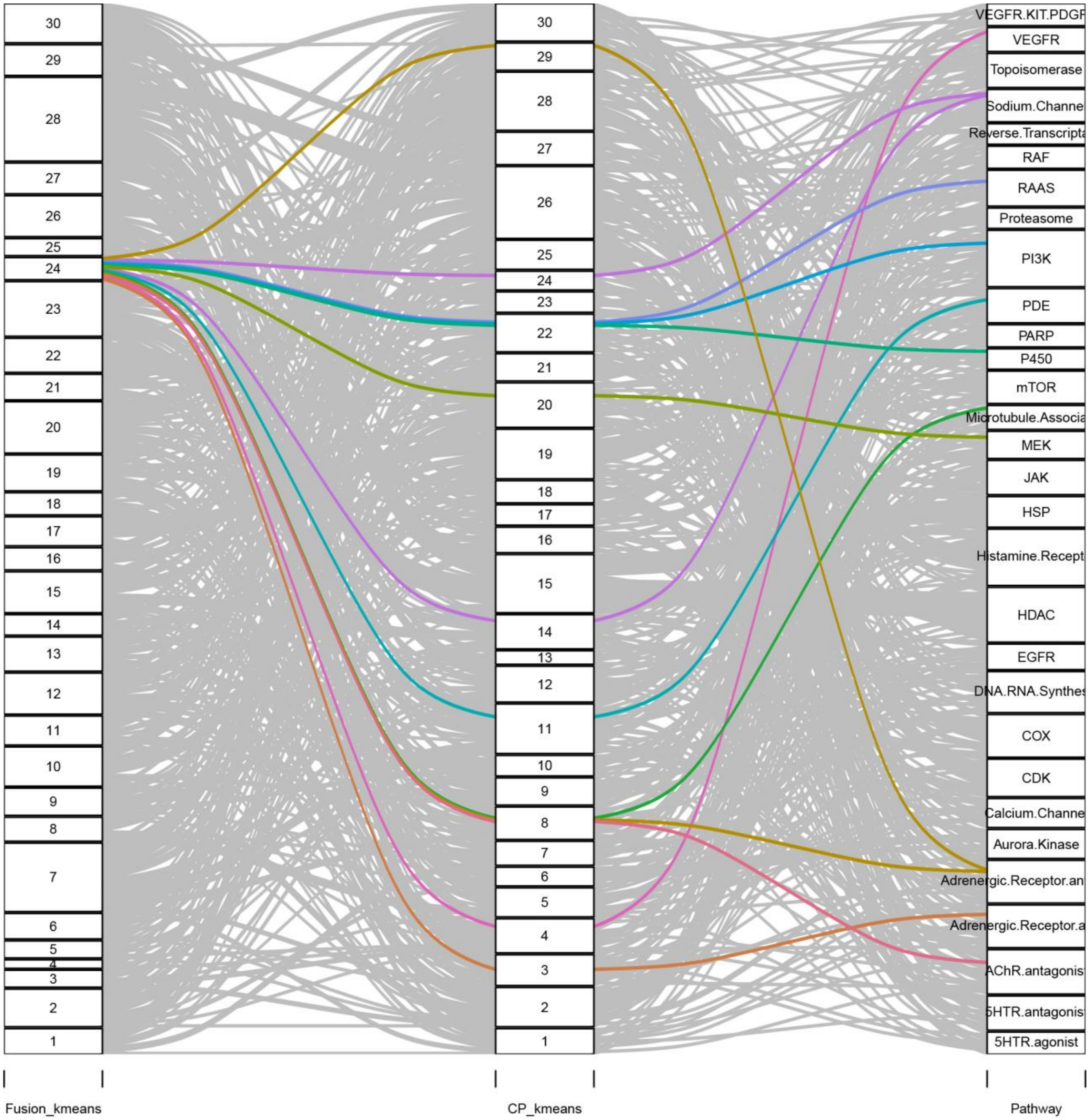
Fusion_kmeans
CP_kmeans
Pathway
VEGFR.KIT.PDGF
VEGFR
Topoisomerase
Sodium.Channe
Reverse.Transcripta
RAF
RAAS
Proteasome
PI3K
PDE
PARP
P450
mTOR
Microtubule.Associa
MEK
JAK
HSP
Histamine.Recept
HDAC
EGFR
DNA.RNA.Synthes
COX
CDK
Calcium.Channe
Aurora.Kinase
Adrenergic.Receptor.an
Adrenergic.Receptor.a
AChR.antagonis
5HTR.antagonis
5HTR.agonist

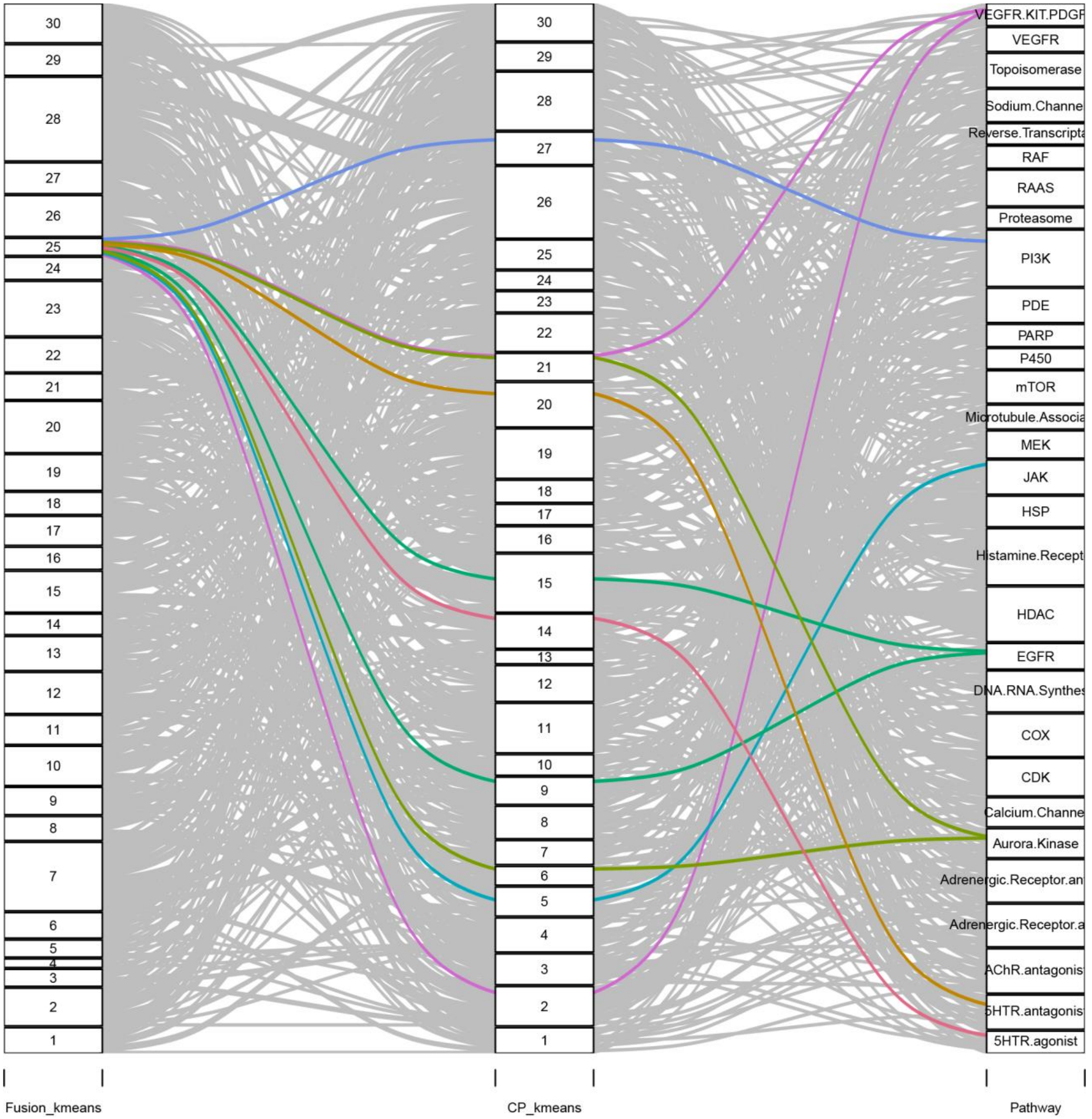
30
29
28
27
26
25
24
23
22
21
20
19
18
17
16
15
14
13
12
11
10
9
8
7
6
5
4
3
2
1
VEGFR.KIT.PDGF
VEGFR
Topoisomerase
Sodium.Channe
Reverse.Transcripta
RAF
RAAS
Proteasome
PI3K
PDE
PARP
P450
mTOR
Microtubule.Associa
MEK
JAK
HSP
Histamine.Recept
HDAC
EGFR
DNA.RNA.Synthes
COX
CDK
Calcium.Channe
Aurora.Kinase
Adrenergic.Receptor.an
Adrenergic.Receptor.a
AChR.antagonis
5HTR.antagonis
5HTR.agonist
Fusion_kmeans
CP_kmeans
Pathway

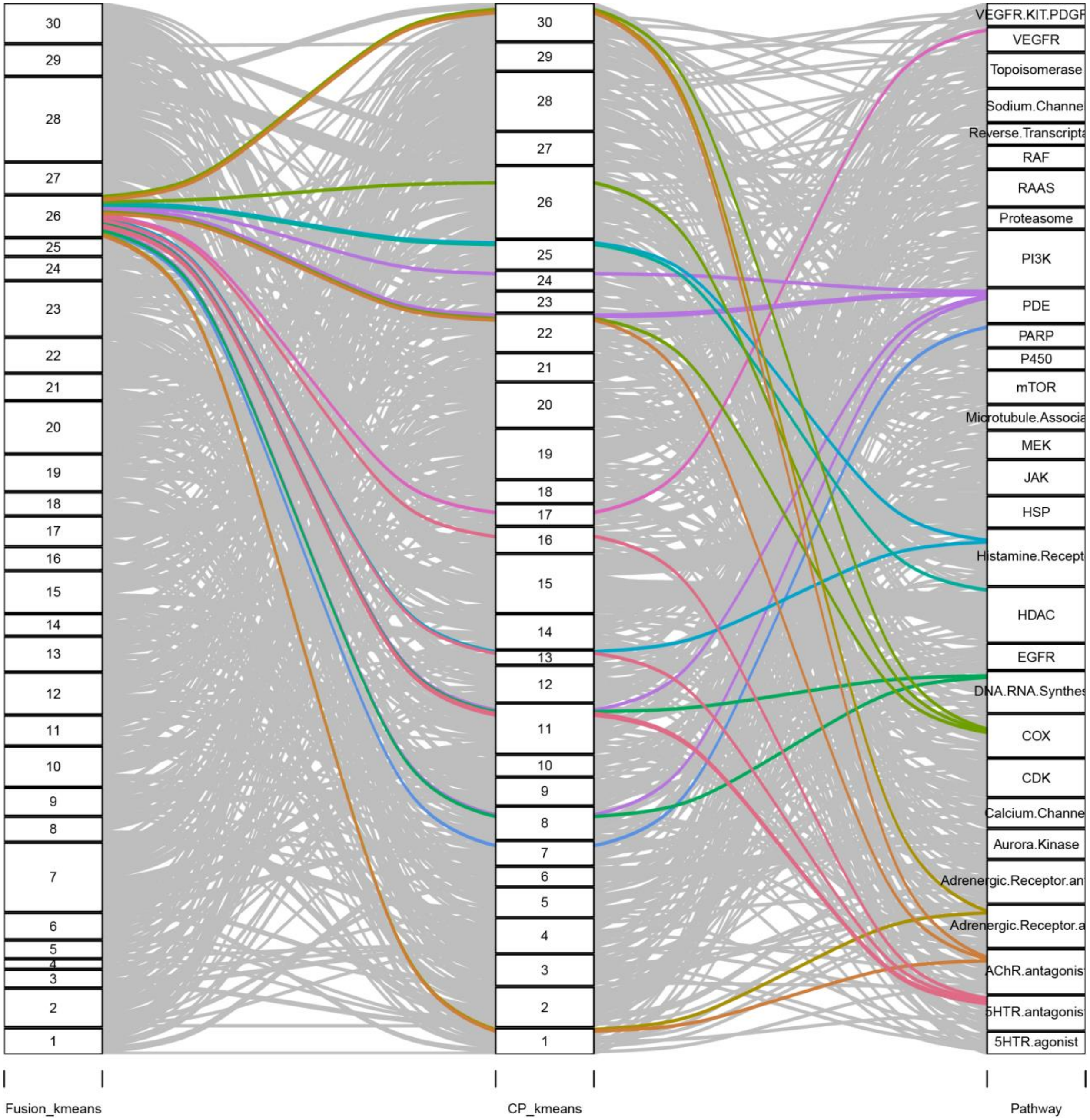

30
29
28
27
26
25
24
23
22
21
20
19
18
17
16
15
14
13
12
11
10
9
8
7
6
5
4
3
2
1
VEGFR.KIT.PDGF
VEGFR
Topoisomerase
Sodium.Channe
Reverse.Transcripta
RAF
RAAS
Proteasome
PI3K
PDE
PARP
P450
mTOR
Microtubule.Associa
MEK
JAK
HSP
Histamine.Recept
HDAC
EGFR
DNA.RNA.Synthes
COX
CDK
Calcium.Channe
Aurora.Kinase
Adrenergic.Receptor.an
Adrenergic.Receptor.a
AChR.antagonis
5HTR.antagonis
5HTR.agonist
Fusion_kmeans
CP_kmeans
Pathway

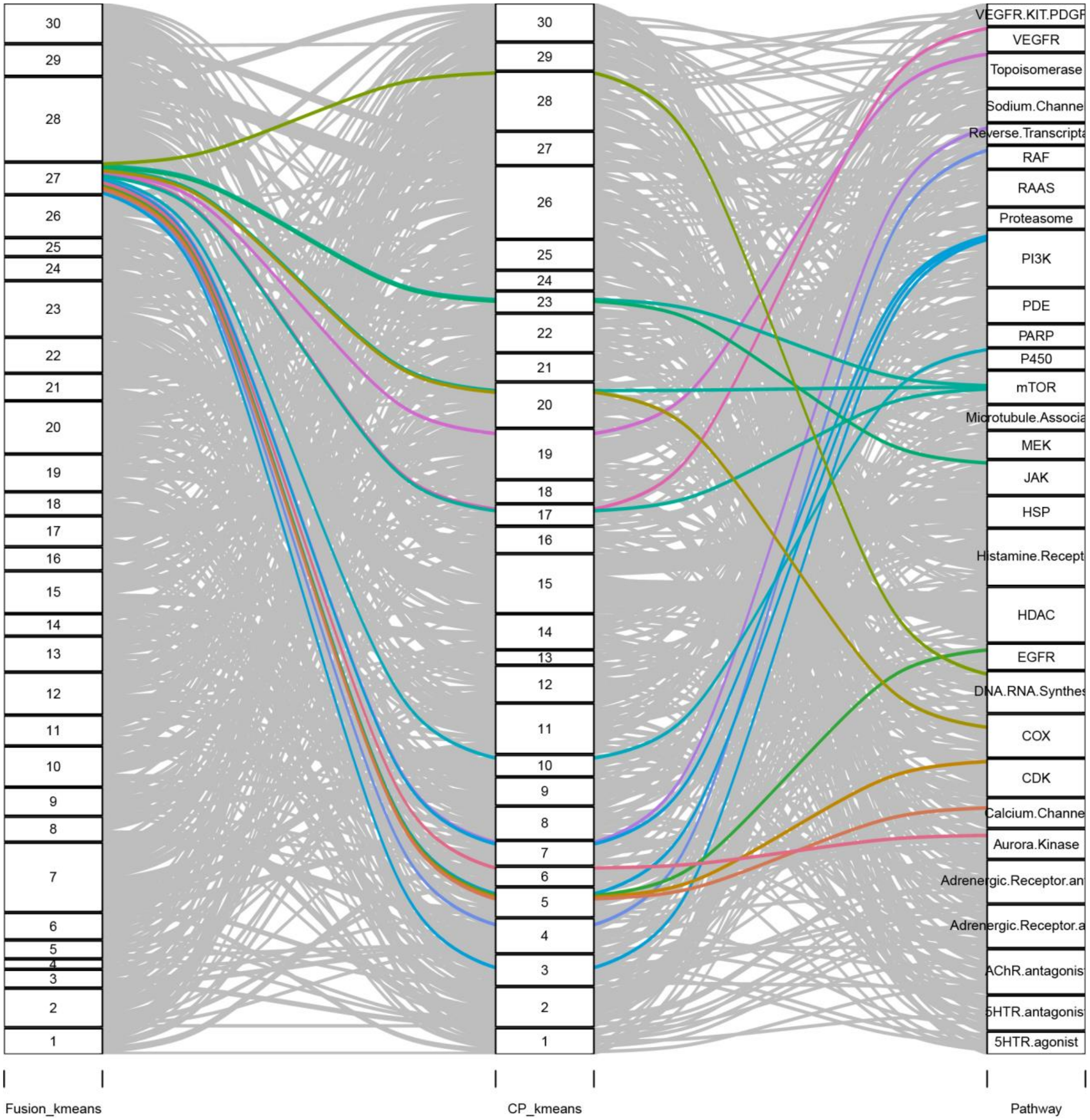

Fusion_kmeans
CP_kmeans
Pathway
30
29
28
27
26
25
24
23
22
21
20
19
18
17
16
15
14
13
12
11
10
9
8
7
6
5
4
3
2
1
VEGFR.KIT.PDGF
VEGFR
Topoisomerase
Sodium.Channe
Reverse.Transcripta
RAF
RAAS
Proteasome
PI3K
PDE
PARP
P450
mTOR
Microtubule.Associa
MEK
JAK
HSP
Histamine.Recept
HDAC
EGFR
DNA.RNA.Synthes
COX
CDK
Calcium.Channe
Aurora.Kinase
Adrenergic.Receptor.an
Adrenergic.Receptor.a
AChR.antagonis
5HTR.antagonis
5HTR.agonist

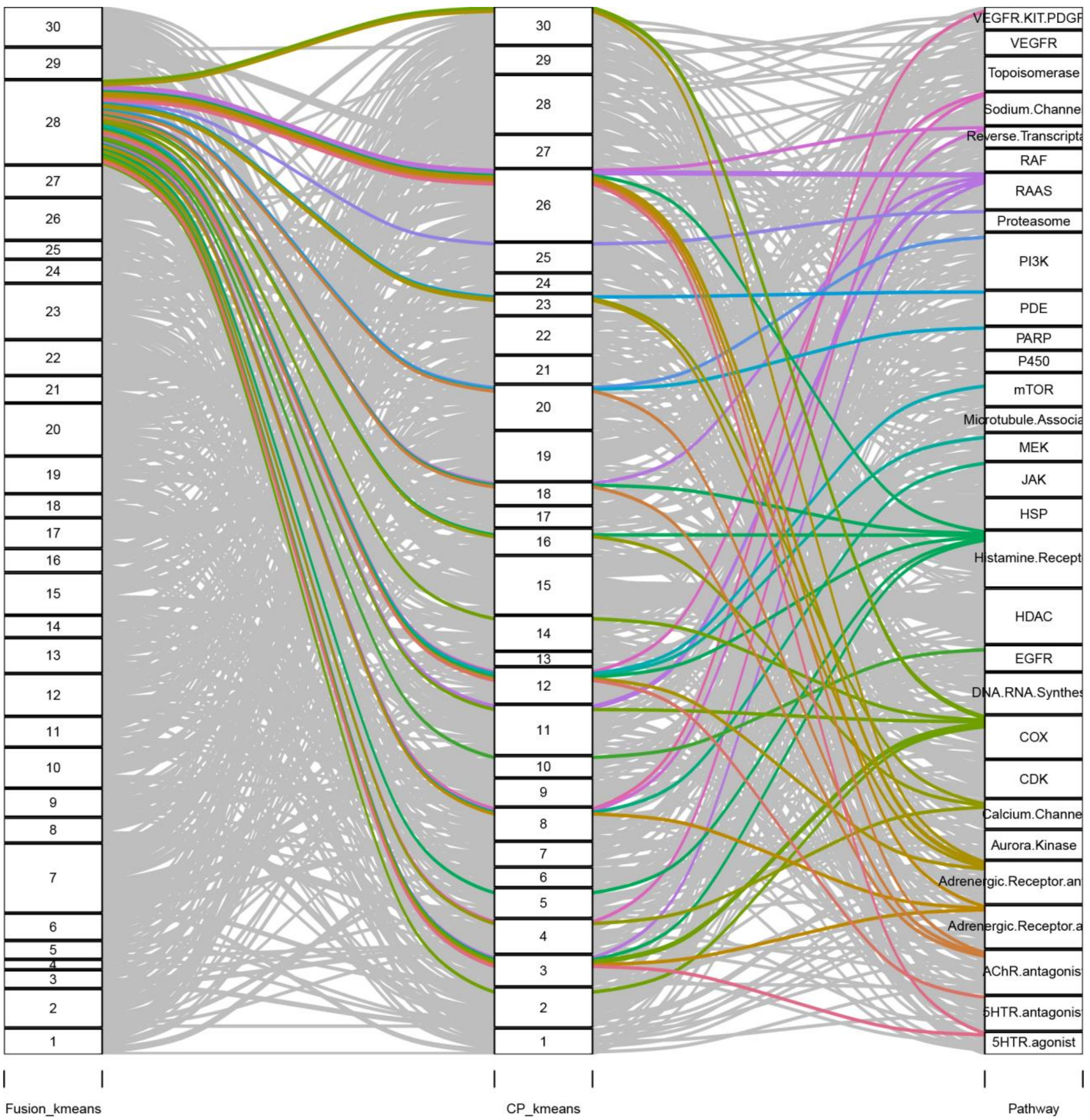
Fusion_kmeans
CP_kmeans
Pathway
VEGFR.KIT.PDGF
VEGFR
Topoisomerase
Sodium.Channe
Reverse.Transcripta
RAF
RAAS
Proteasome
PI3K
PDE
PARP
P450
mTOR
Microtubule.Associa
MEK
JAK
HSP
Histamine.Recept
HDAC
EGFR
DNA.RNA.Synthes
COX
CDK
Calcium.Channe
Aurora.Kinase
Adrenergic.Receptor.an
Adrenergic.Receptor.a
AChR.antagonis
5HTR.antagonis
5HTR.agonist

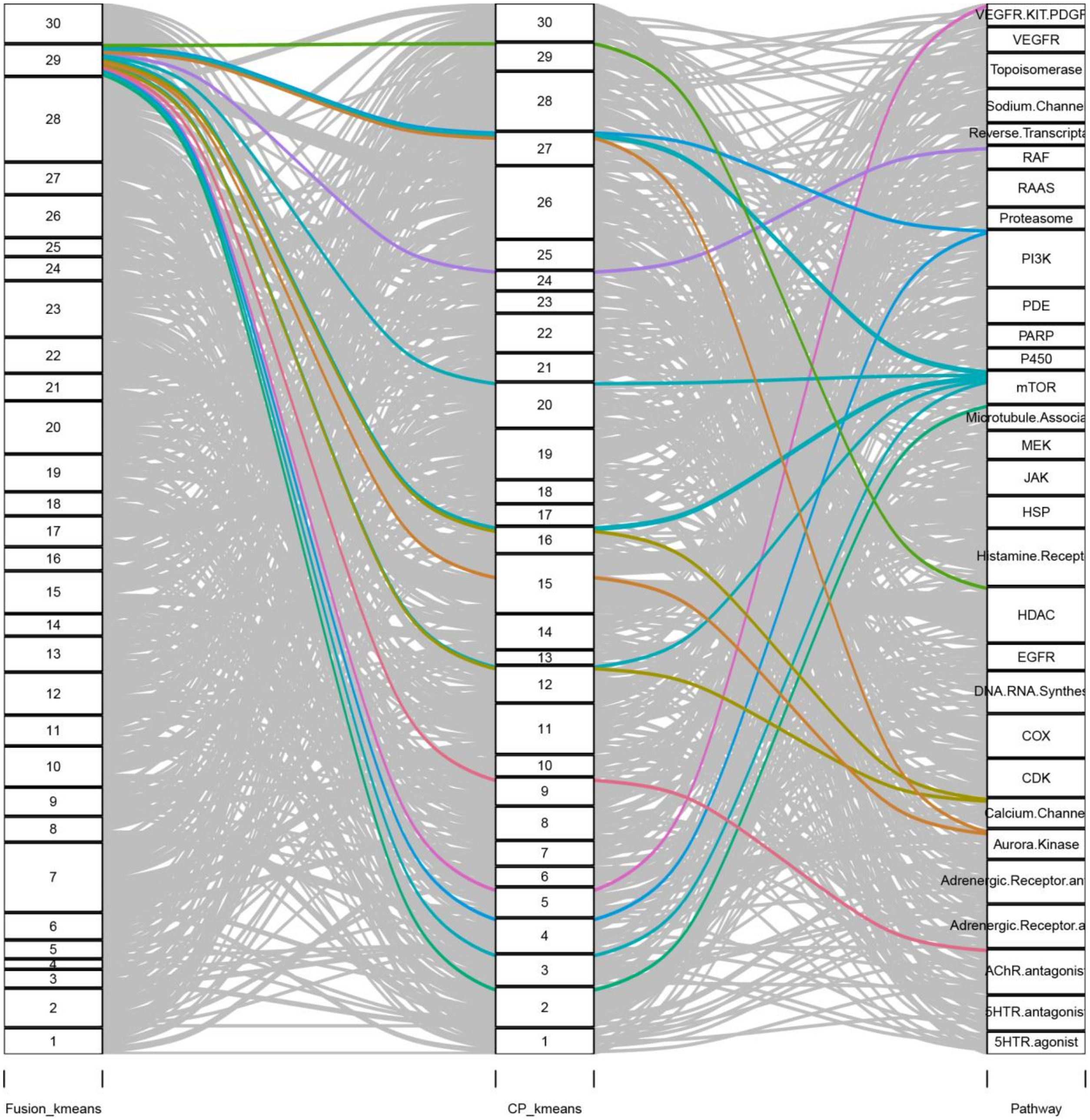
Fusion_kmeans
CP_kmeans
Pathway
VEGFR.KIT.PDGF
VEGFR
Topoisomerase
Sodium.Channe
Reverse.Transcripta
RAF
RAAS
Proteasome
PI3K
PDE
PARP
P450
mTOR
Microtubule.Associa
MEK
JAK
HSP
Histamine.Recept
HDAC
EGFR
DNA.RNA.Synthes
COX
CDK
Calcium.Channe
Aurora.Kinase
Adrenergic.Receptor.an
Adrenergic.Receptor.a
AChR.antagonis
5HTR.antagonis
5HTR.agonist

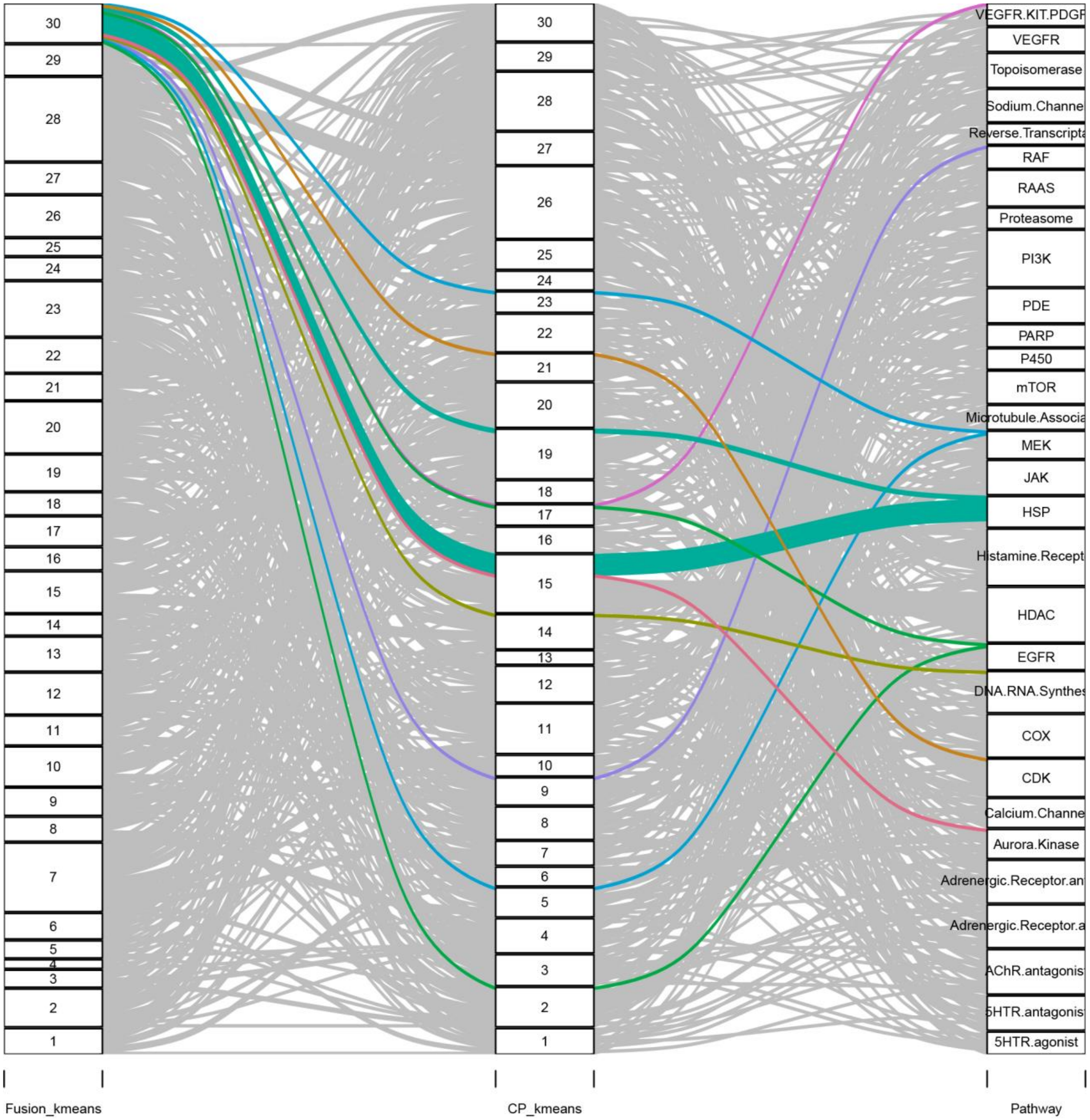
30
29
28
27
26
25
24
23
22
21
20
19
18
17
16
15
14
13
12
11
10
9
8
7
6
5
4
3
2
1
VEGFR.KIT.PDGF
VEGFR
Topoisomerase
Sodium.Channe
Reverse.Transcripta
RAF
RAAS
Proteasome
PI3K
PDE
PARP
P450
mTOR
Microtubule.Associa
MEK
JAK
HSP
Histamine.Recept
HDAC
EGFR
DNA.RNA.Synthes
COX
CDK
Calcium.Channe
Aurora.Kinase
Adrenergic.Receptor.an
Adrenergic.Receptor.a
AChR.antagonis
5HTR.antagonis
5HTR.agonist
Fusion_kmeans
CP_kmeans
Pathway

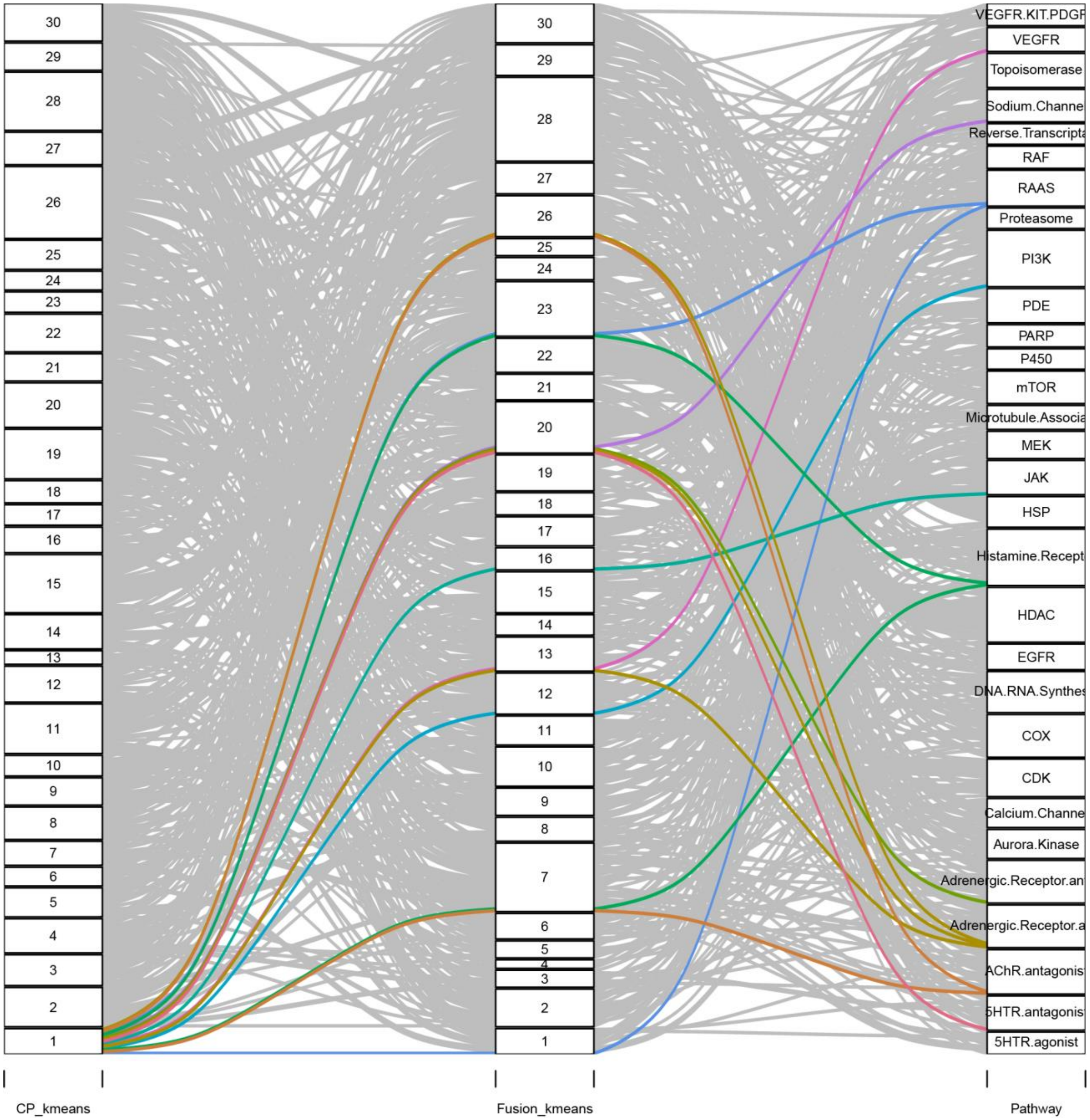
30
29
28
27
26
25
24
23
22
21
20
19
18
17
16
15
14
13
12
11
10
9
8
7
6
5
4
3
2
1
VEGFR.KIT.PDGF
VEGFR
Topoisomerase
Sodium.Channe
Reverse.Transcripta
RAF
RAAS
Proteasome
PI3K
PDE
PARP
P450
mTOR
Microtubule.Associa
MEK
JAK
HSP
Histamine.Recept
HDAC
EGFR
DNA.RNA.Synthes
COX
CDK
Calcium.Channe
Aurora.Kinase
Adrenergic.Receptor.an
Adrenergic.Receptor.a
AChR.antagonis
5HTR.antagonis
5HTR.agonist
CP_kmeans
Fusion_kmeans
Pathway

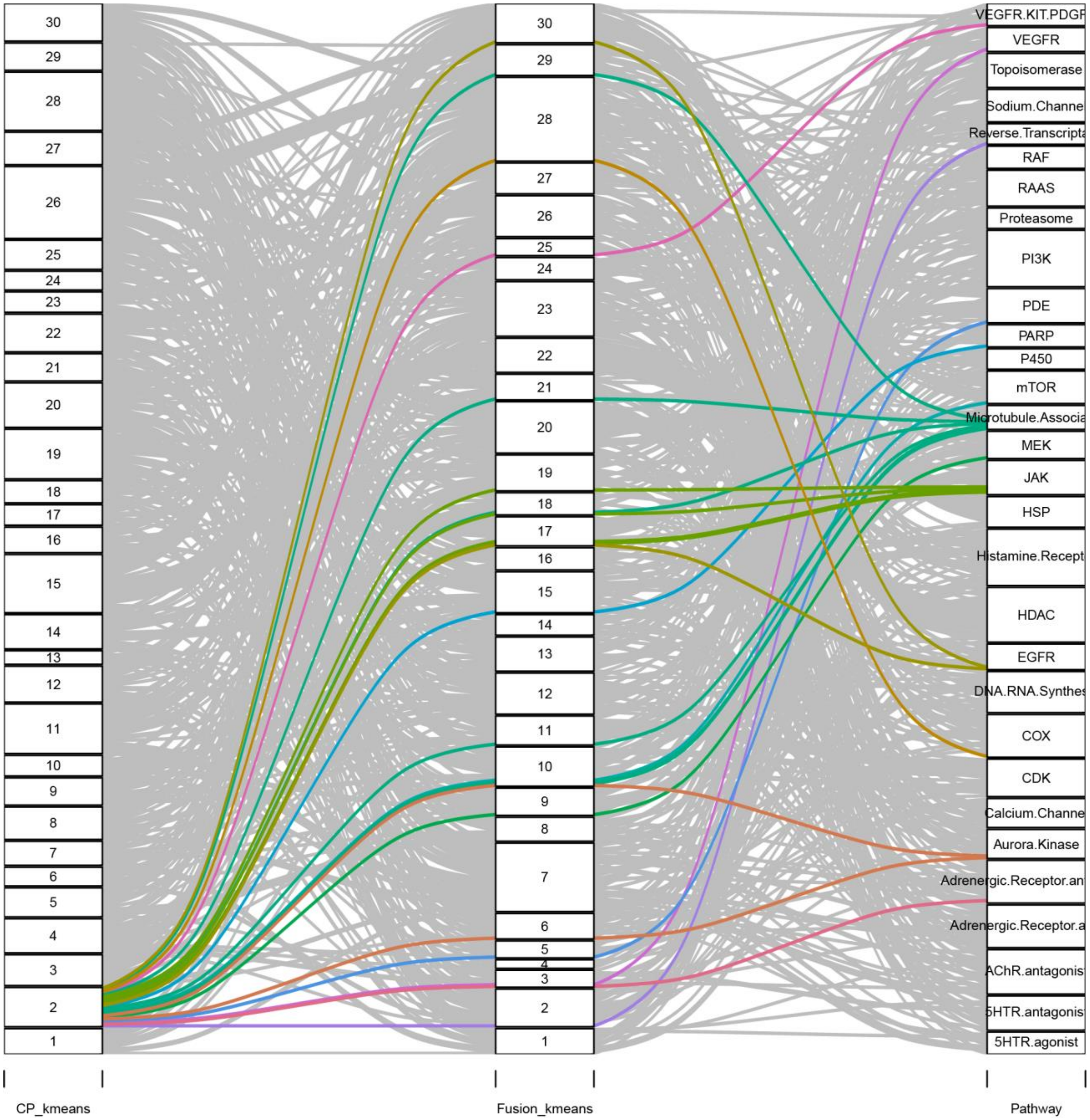
CP_kmeans
Fusion_kmeans
Pathway
30
29
28
27
26
25
24
23
22
21
20
19
18
17
16
15
14
13
12
11
10
9
8
7
6
5
4
3
2
1
VEGFR.KIT.PDGF
VEGFR
Topoisomerase
Sodium.Channe
Reverse.Transcripta
RAF
RAAS
Proteasome
PI3K
PDE
PARP
P450
mTOR
Microtubule.Associa
MEK
JAK
HSP
Histamine.Recept
HDAC
EGFR
DNA.RNA.Synthes
COX
CDK
Calcium.Channe
Aurora.Kinase
Adrenergic.Receptor.an
Adrenergic.Receptor.a
AChR.antagonis
5HTR.antagonis
5HTR.agonist

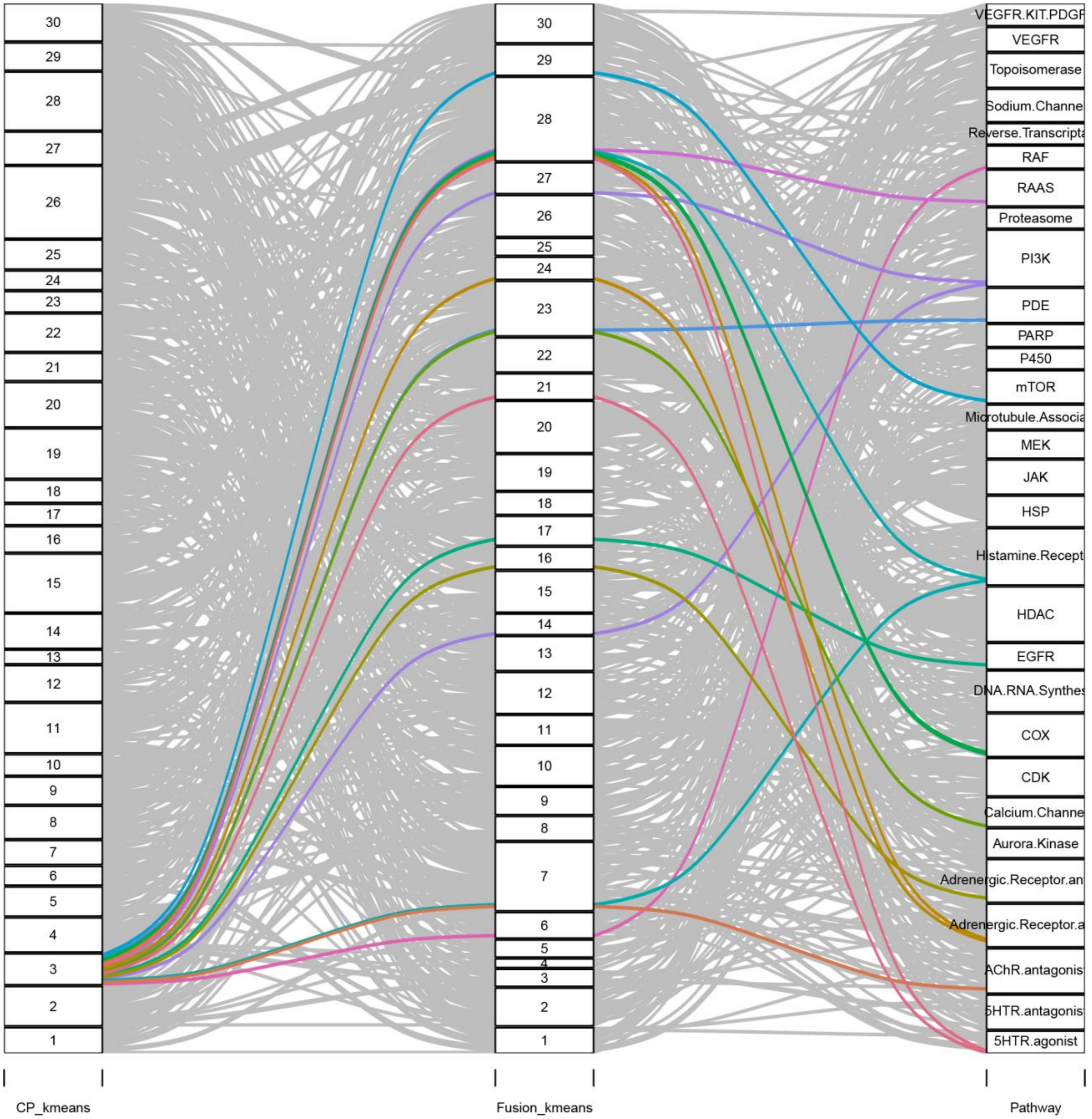

30
29
28
27
26
25
24
23
22
21
20
19
18
17
16
15
14
13
12
11
10
9
8
7
6
5
4
3
2
1
VEGFR.KIT.PDGF
VEGFR
Topoisomerase
Sodium.Channe
Reverse.Transcripta
RAF
RAAS
Proteasome
PI3K
PDE
PARP
P450
mTOR
Microtubule.Associa
MEK
JAK
HSP
Histamine.Recept
HDAC
EGFR
DNA.RNA.Synthes
COX
CDK
Calcium.Channe
Aurora.Kinase
Adrenergic.Receptor.an
Adrenergic.Receptor.a
AChR.antagonis
5HTR.antagonis
5HTR.agonist
CP_kmeans
Fusion_kmeans
Pathway

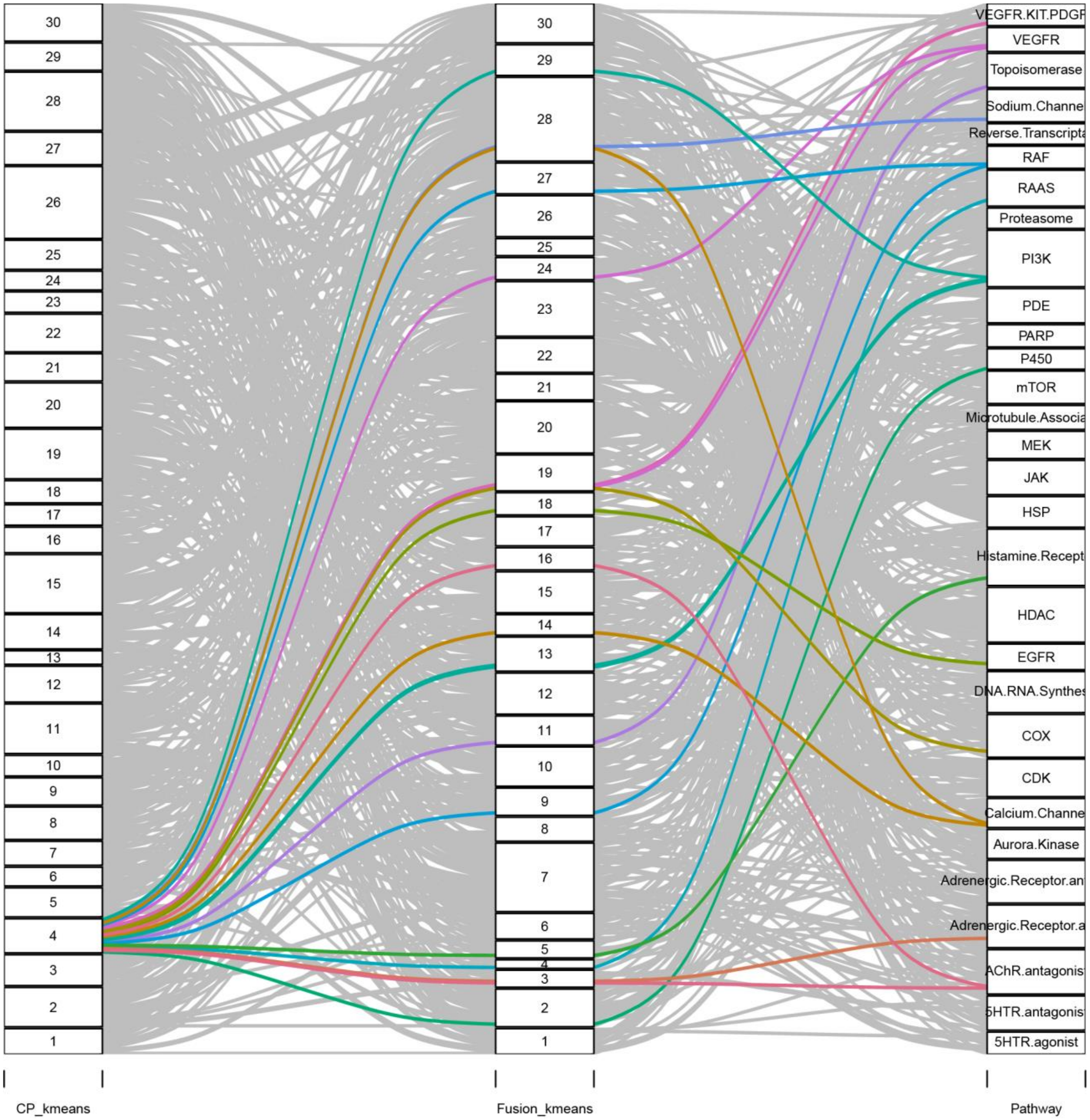
CP_kmeans
Fusion_kmeans
Pathway
VEGFR.KIT.PDGF
VEGFR
Topoisomerase
Sodium.Channe
Reverse.Transcripta
RAF
RAAS
Proteasome
PI3K
PDE
PARP
P450
mTOR
Microtubule.Associa
MEK
JAK
HSP
Histamine.Recept
HDAC
EGFR
DNA.RNA.Synthes
COX
CDK
Calcium.Channe
Aurora.Kinase
Adrenergic.Receptor.an
Adrenergic.Receptor.a
AChR.antagonis
5HTR.antagonis
5HTR.agonist

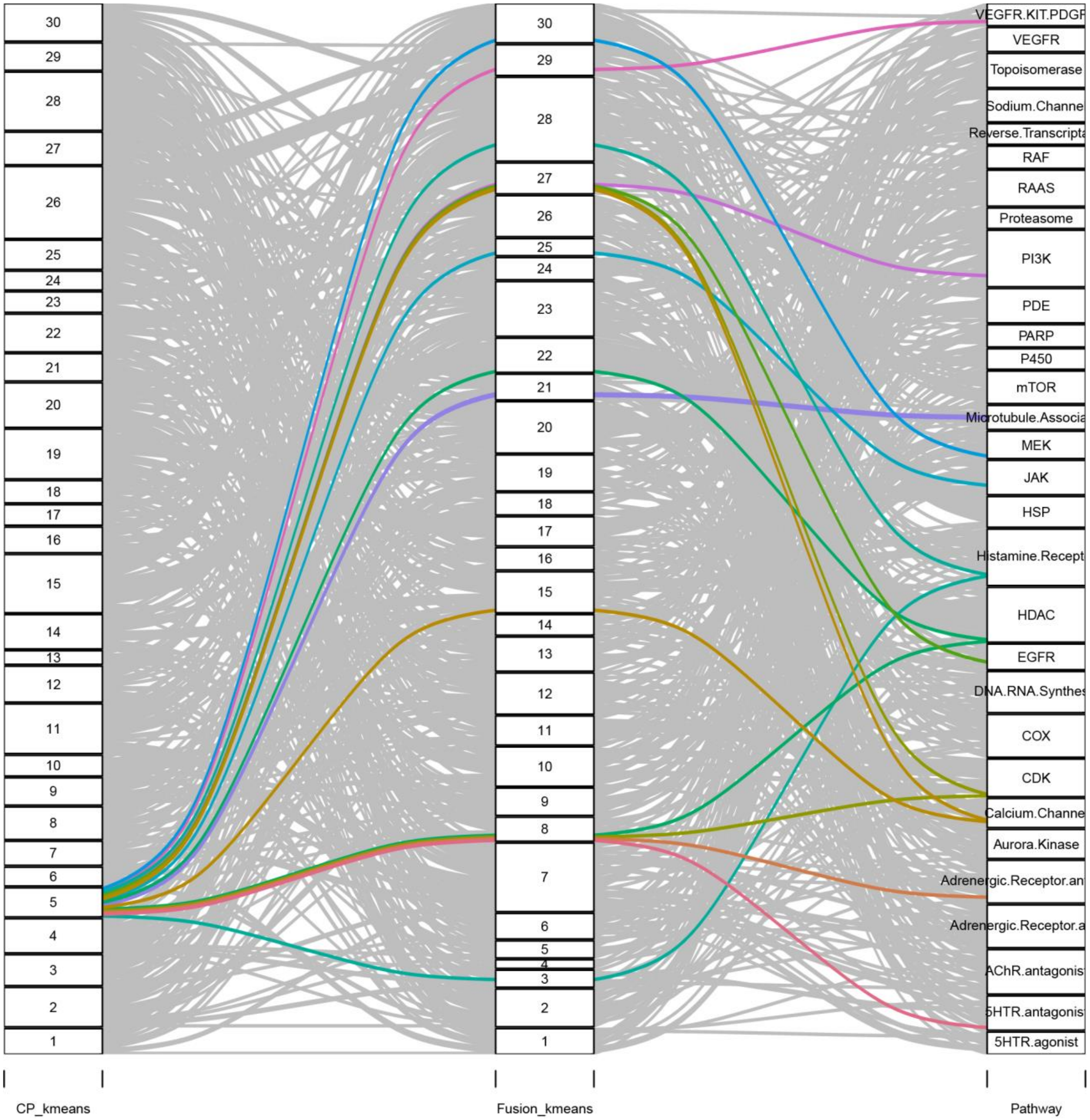
30
29
28
27
26
25
24
23
22
21
20
19
18
17
16
15
14
13
12
11
10
9
8
7
6
5
4
3
2
1
VEGFR.KIT.PDGF
VEGFR
Topoisomerase
Sodium.Channe
Reverse.Transcripta
RAF
RAAS
Proteasome
PI3K
PDE
PARP
P450
mTOR
Microtubule.Associa
MEK
JAK
HSP
Histamine.Recept
HDAC
EGFR
DNA.RNA.Synthes
COX
CDK
Calcium.Channe
Aurora.Kinase
Adrenergic.Receptor.an
Adrenergic.Receptor.a
AChR.antagonis
5HTR.antagonis
5HTR.agonist
CP_kmeans
Fusion_kmeans
Pathway

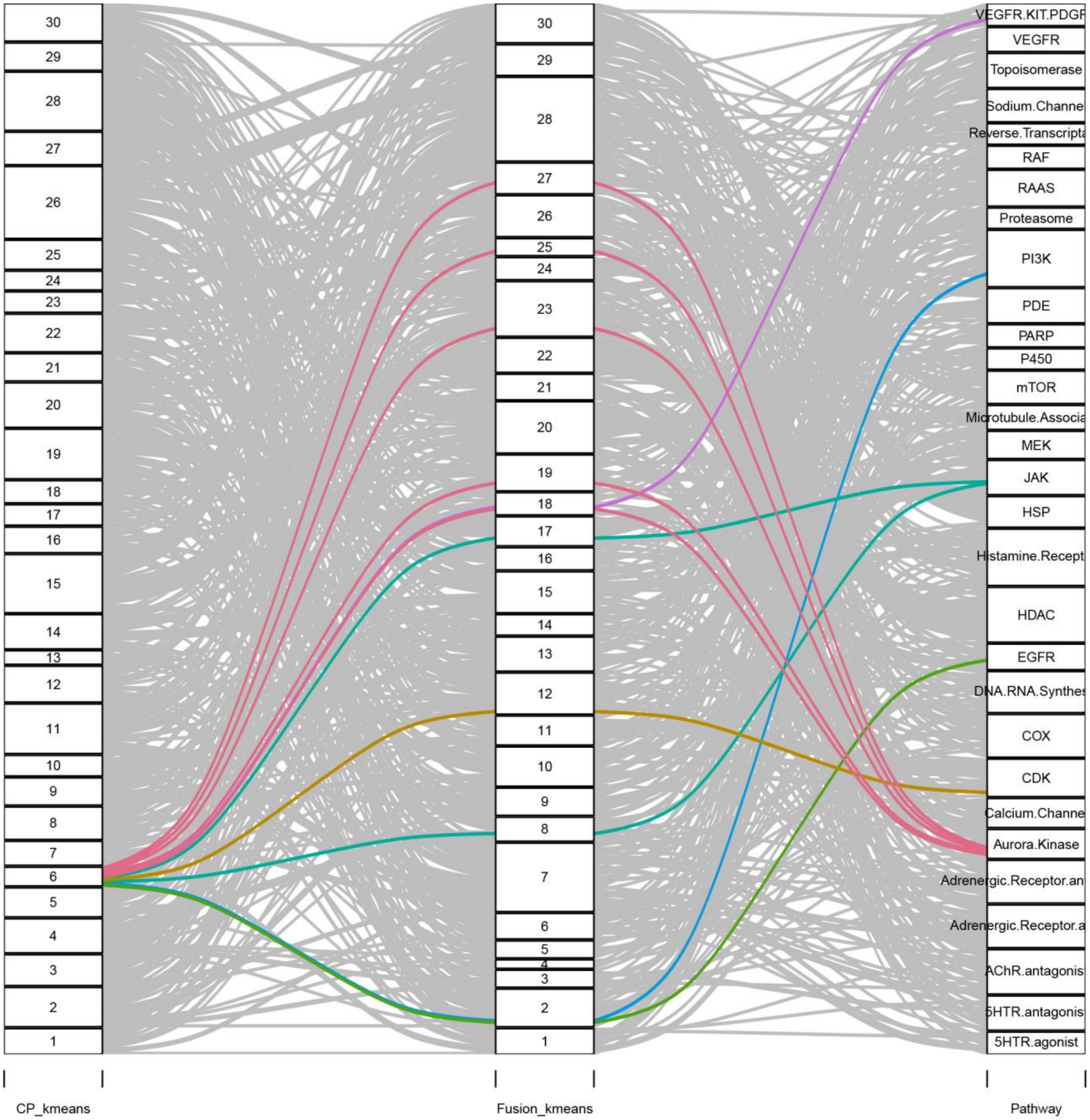

30
29
28
27
26
25
24
23
22
21
20
19
18
17
16
15
14
13
12
11
10
9
8
7
6
5
4
3
2
1
VEGFR.KIT.PDGF
VEGFR
Topoisomerase
Sodium.Channe
Reverse.Transcripta
RAF
RAAS
Proteasome
PI3K
PDE
PARP
P450
mTOR
Microtubule.Associa
MEK
JAK
HSP
Histamine.Recept
HDAC
EGFR
DNA.RNA.Synthes
COX
CDK
Calcium.Channe
Aurora.Kinase
Adrenergic.Receptor.an
Adrenergic.Receptor.a
AChR.antagonis
5HTR.antagonis
5HTR.agonist
CP_kmeans
Fusion_kmeans
Pathway

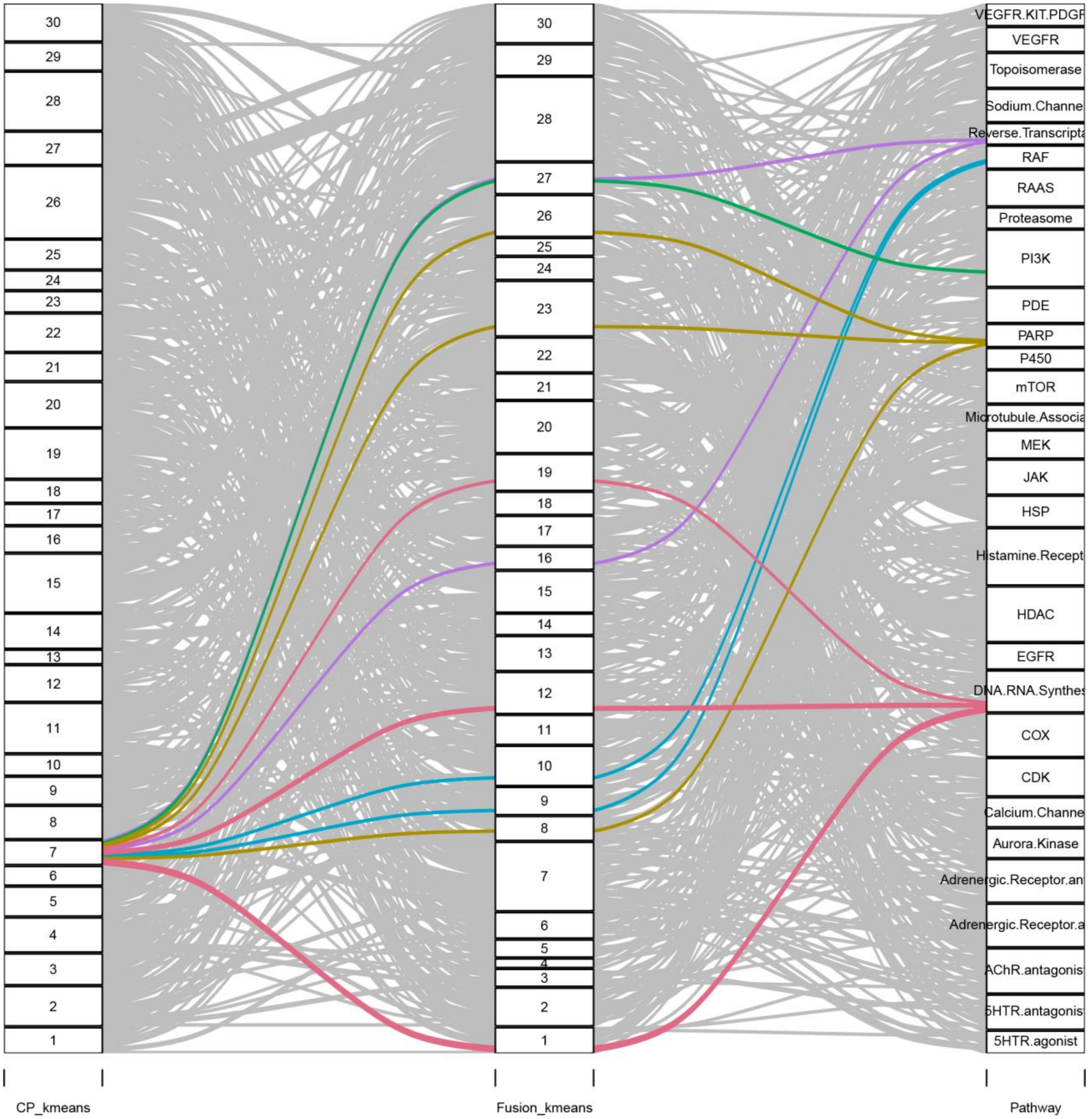
CP_kmeans
Fusion_kmeans
Pathway
VEGFR.KIT.PDGF
VEGFR
Topoisomerase
RAF
RAAS
Proteasome
PI3K
PDE
PARP
P450
mTOR
MEK
JAK
HSP
HDAC
EGFR
COX
CDK
Aurora.Kinase
5HTR.agonist

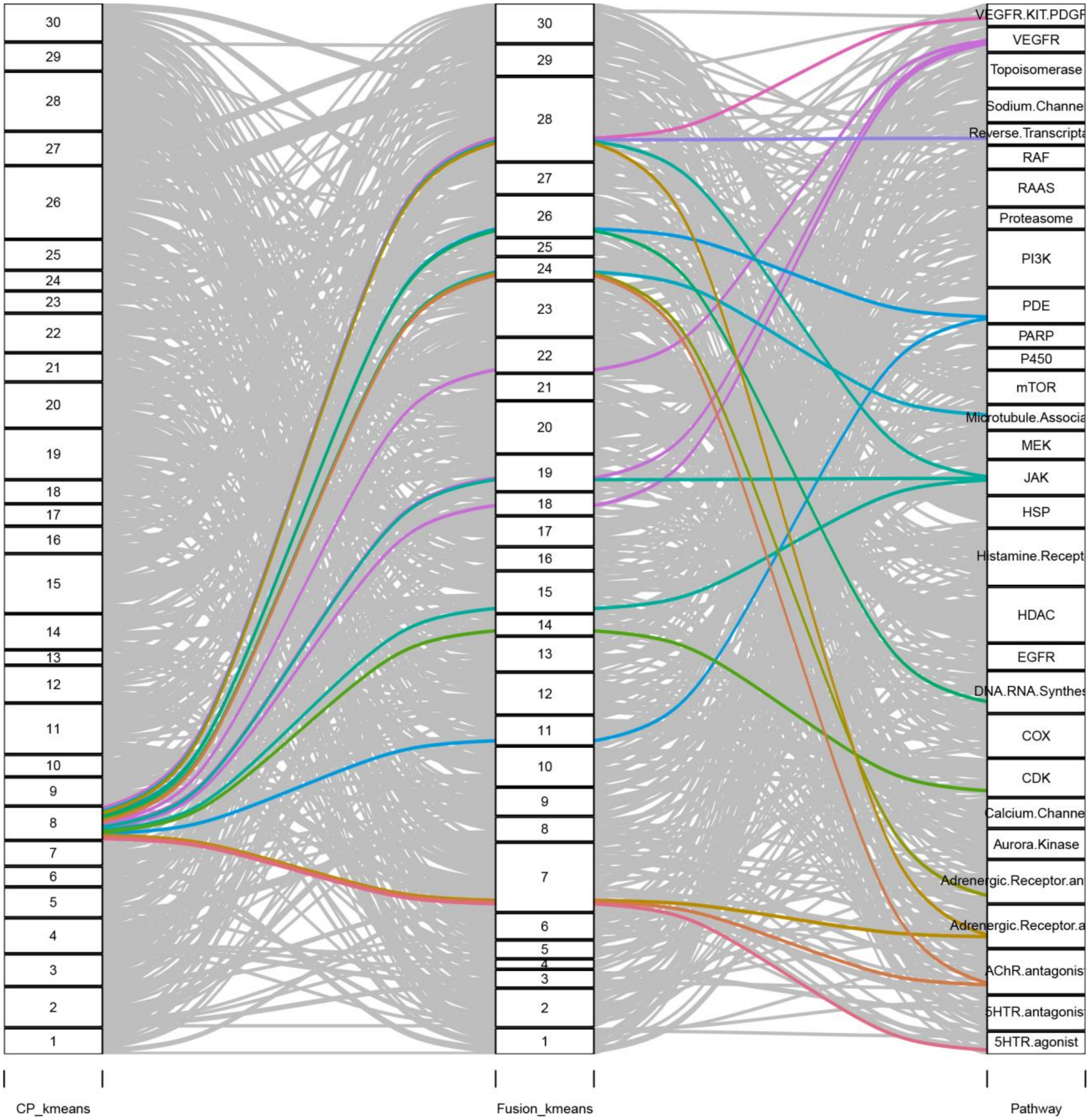
30
29
28
27
26
25
24
23
22
21
20
19
18
17
16
15
14
13
12
11
10
9
8
7
6
5
4
3
2
1
VEGFR.KIT.PDGF
VEGFR
Topoisomerase
Sodium.Channe
Reverse.Transcripta
RAF
RAAS
Proteasome
PI3K
PDE
PARP
P450
mTOR
Microtubule.Associa
MEK
JAK
HSP
Histamine.Recept
HDAC
EGFR
DNA.RNA.Synthes
COX
CDK
Calcium.Channe
Aurora.Kinase
Adrenergic.Receptor.an
Adrenergic.Receptor.a
AChR.antagonis
5HTR.antagonis
5HTR.agonist
CP_kmeans
Fusion_kmeans
Pathway

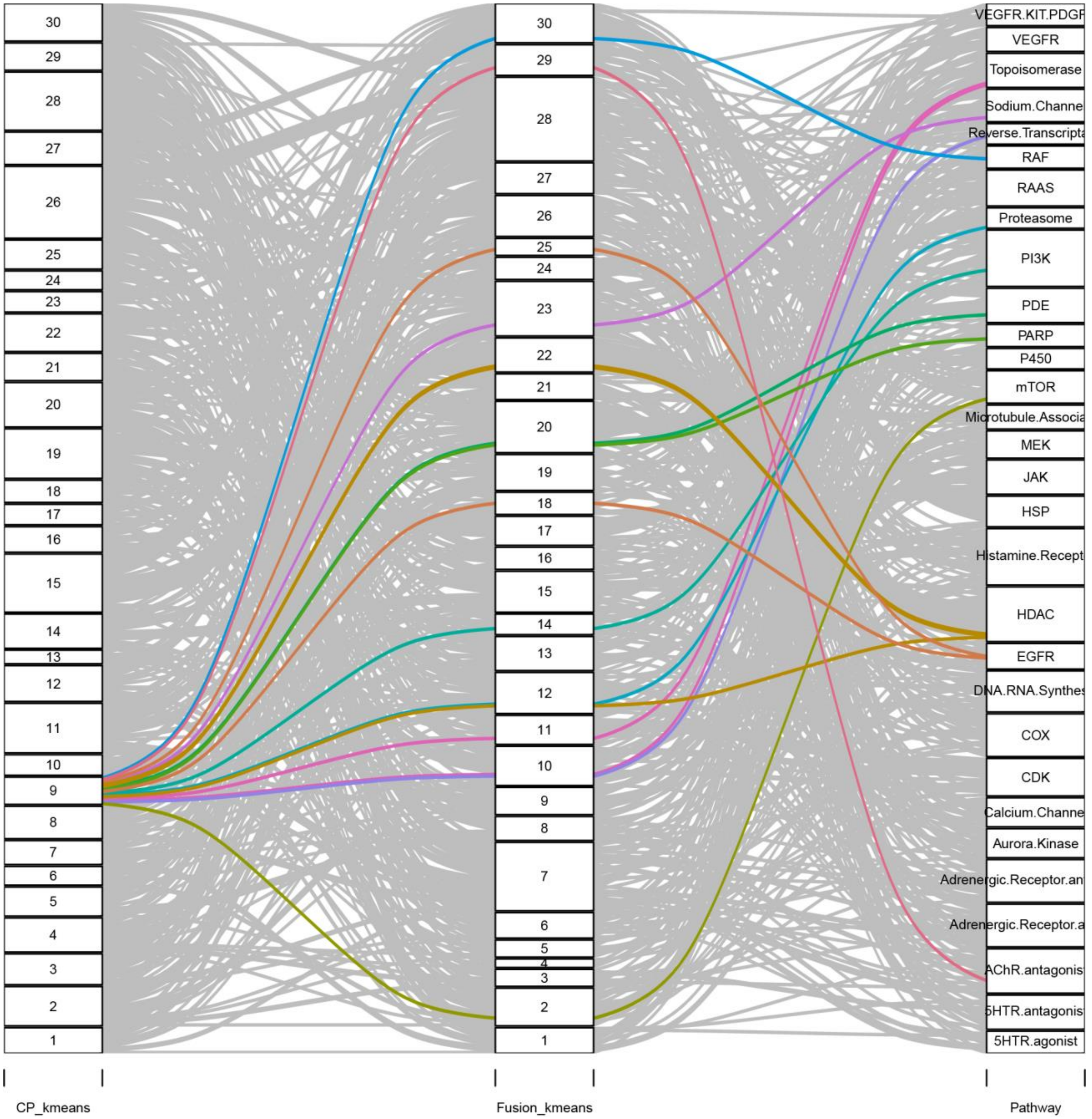
CP_kmeans
Fusion_kmeans
Pathway
30
29
28
27
26
25
24
23
22
21
20
19
18
17
16
15
14
13
12
11
10
9
8
7
6
5
4
3
2
1
VEGFR.KIT.PDGF
VEGFR
Topoisomerase
Sodium.Channe
Reverse.Transcripta
RAF
RAAS
Proteasome
PI3K
PDE
PARP
P450
mTOR
Microtubule.Associa
MEK
JAK
HSP
Histamine.Recept
HDAC
EGFR
DNA.RNA.Synthes
COX
CDK
Calcium.Channe
Aurora.Kinase
Adrenergic.Receptor.an
Adrenergic.Receptor.a
AChR.antagonis
5HTR.antagonis
5HTR.agonist

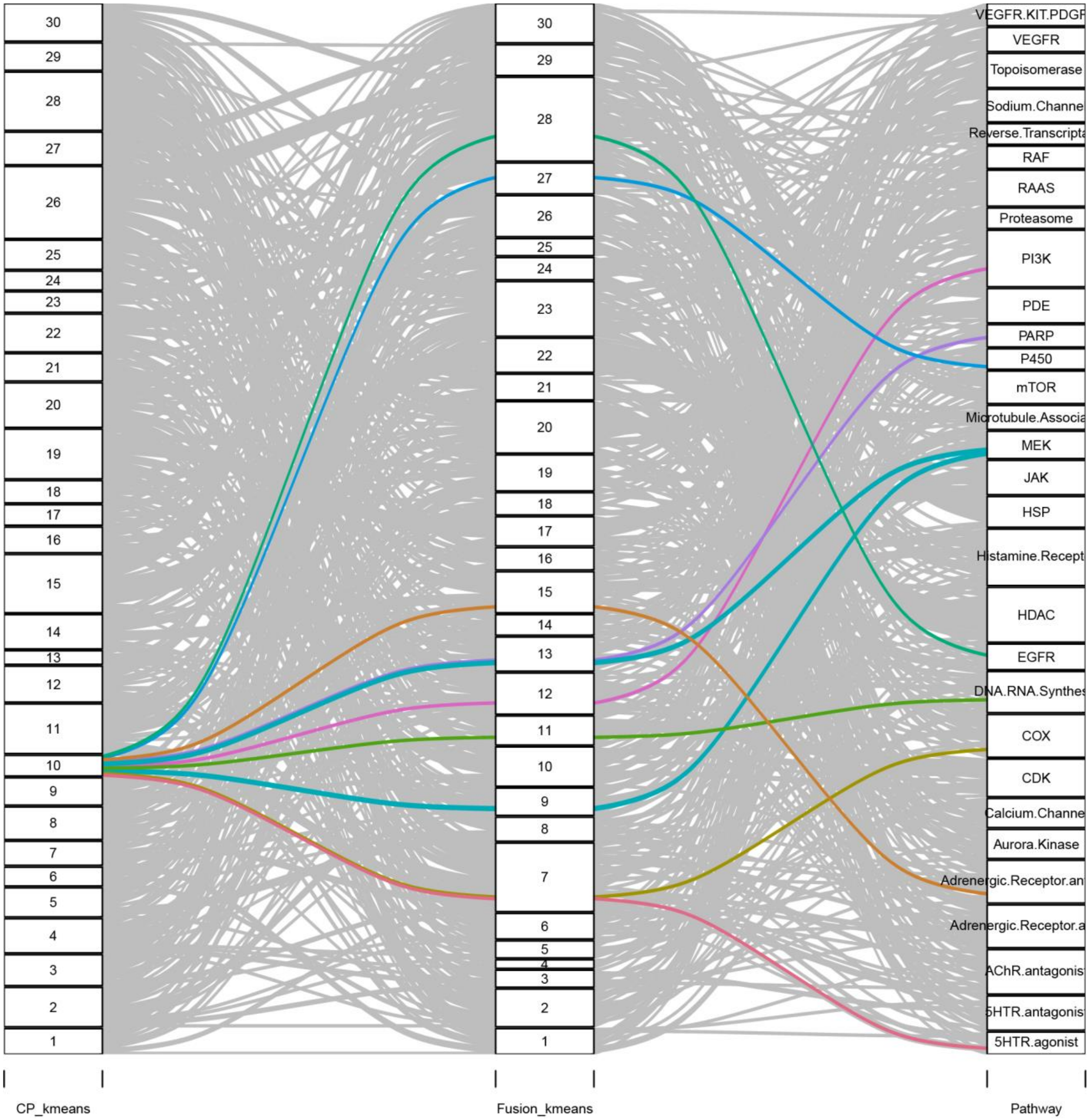
CP_kmeans
Fusion_kmeans
Pathway
VEGFR.KIT.PDGF
VEGFR
Topoisomerase
Sodium.Channe
Reverse.Transcripta
RAF
RAAS
Proteasome
PI3K
PDE
PARP
P450
mTOR
Microtubule.Associa
MEK
JAK
HSP
Histamine.Recept
HDAC
EGFR
DNA.RNA.Synthes
COX
CDK
Calcium.Channe
Aurora.Kinase
Adrenergic.Receptor.an
Adrenergic.Receptor.a
AChR.antagonis
5HTR.antagonis
5HTR.agonist

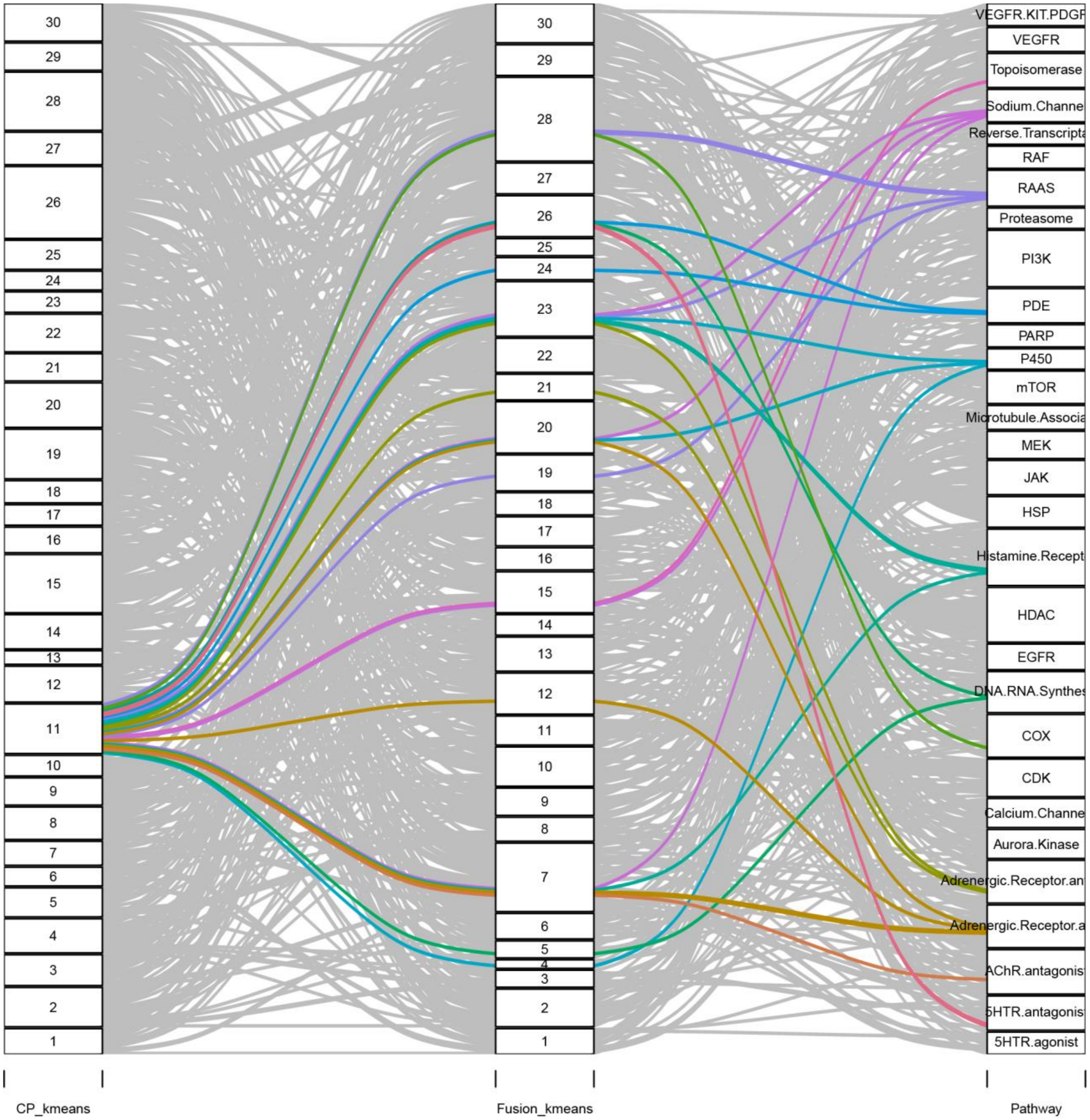

CP_kmeans
Fusion_kmeans
Pathway
VEGFR.KIT.PDGF
VEGFR
Topoisomerase
Sodium.Channe
Reverse.Transcripta
RAF
RAAS
Proteasome
PI3K
PDE
PARP
P450
mTOR
Microtubule.Associa
MEK
JAK
HSP
Histamine.Recept
HDAC
EGFR
DNA.RNA.Synthes
COX
CDK
Calcium.Channe
Aurora.Kinase
Adrenergic.Receptor.an
Adrenergic.Receptor.a
AChR.antagonis
5HTR.antagonis
5HTR.agonist

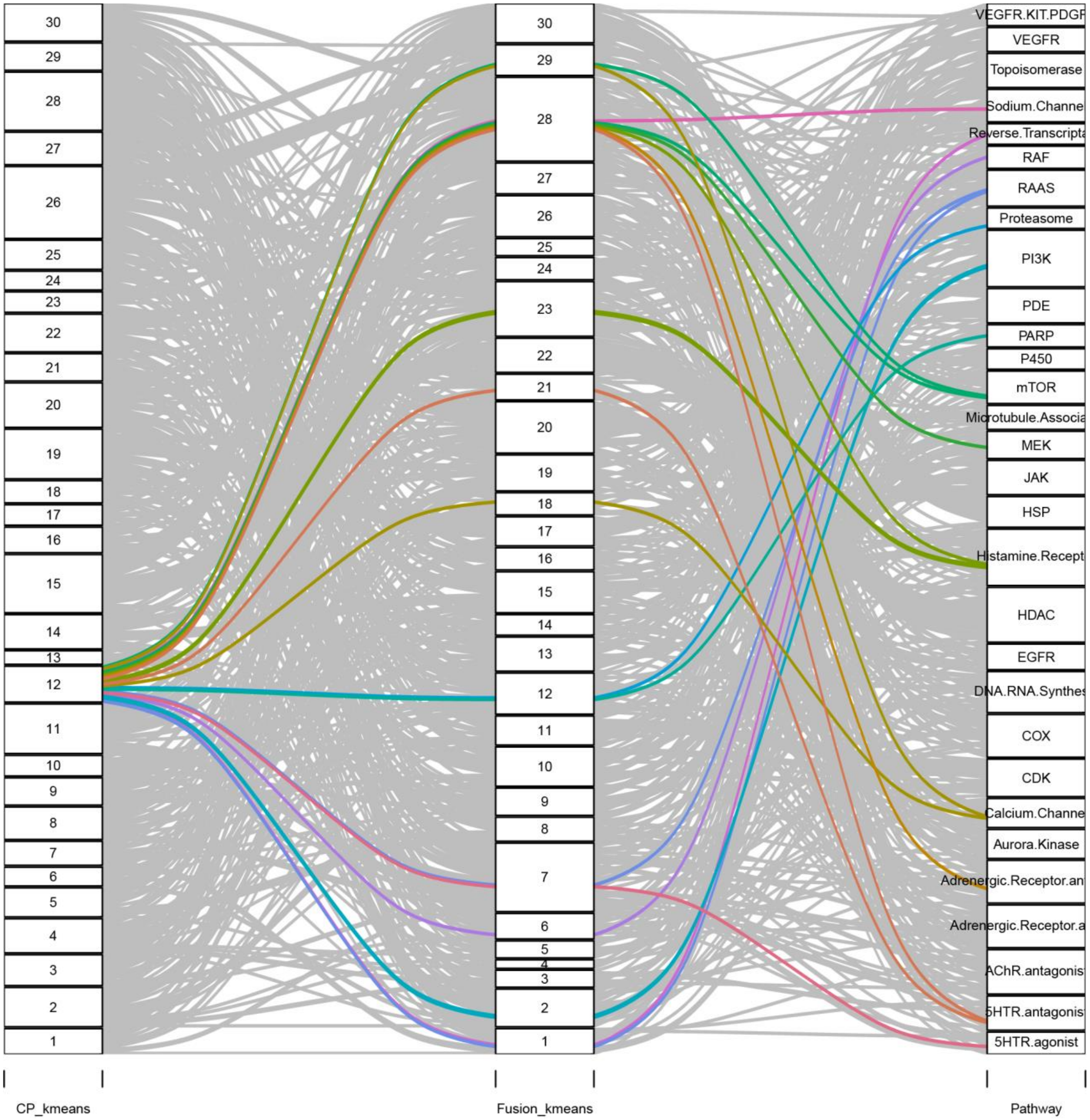
30
29
28
27
26
25
24
23
22
21
20
19
18
17
16
15
14
13
12
11
10
9
8
7
6
5
4
3
2
1
VEGFR.KIT.PDGF
VEGFR
Topoisomerase
Sodium.Channe
Reverse.Transcripta
RAF
RAAS
Proteasome
PI3K
PDE
PARP
P450
mTOR
Microtubule.Associa
MEK
JAK
HSP
Histamine.Recept
HDAC
EGFR
DNA.RNA.Synthes
COX
CDK
Calcium.Channe
Aurora.Kinase
Adrenergic.Receptor.an
Adrenergic.Receptor.a
AChR.antagonis
5HTR.antagonis
5HTR.agonist
CP_kmeans
Fusion_kmeans
Pathway

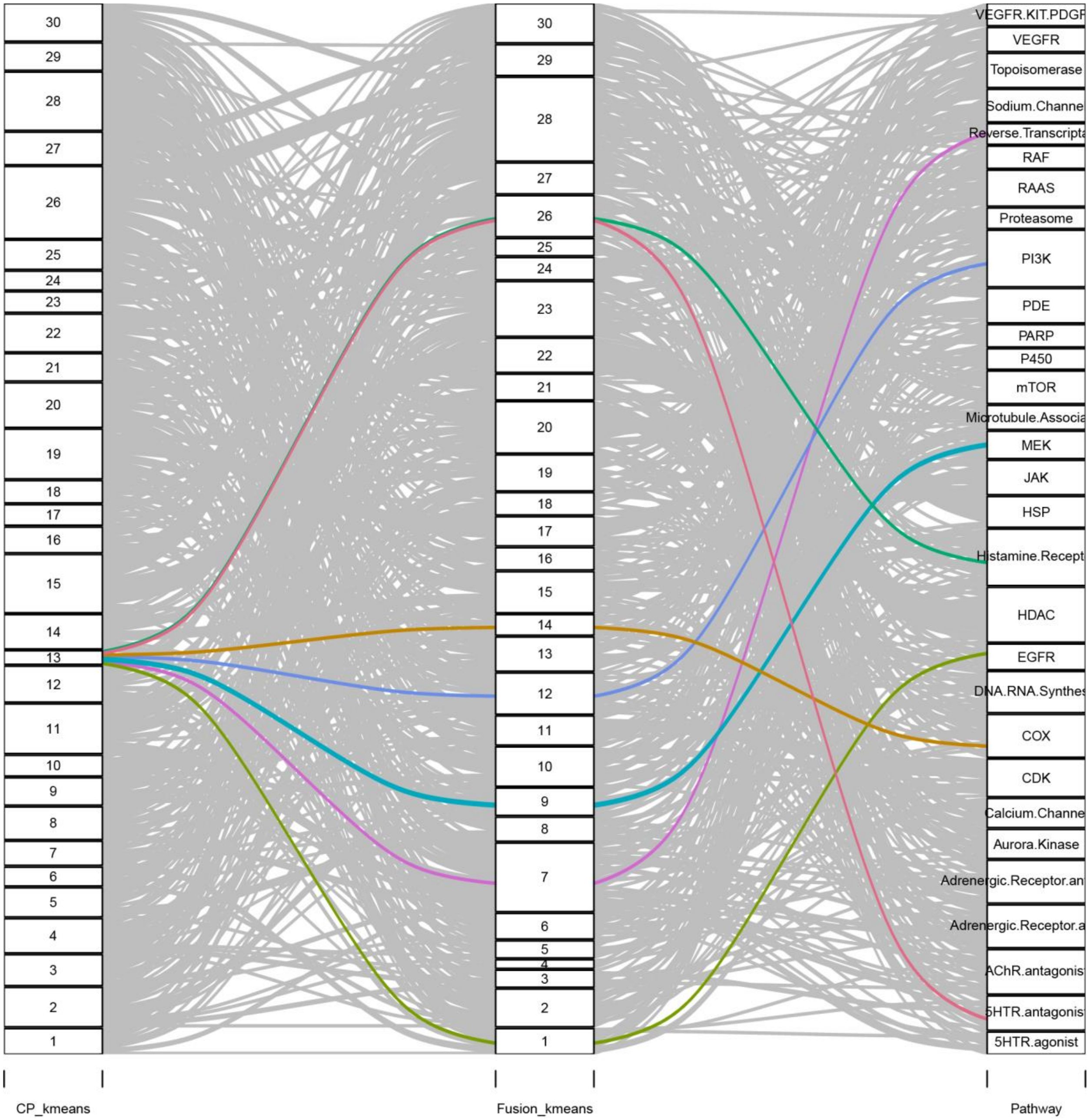
30
29
28
27
26
25
24
23
22
21
20
19
18
17
16
15
14
13
12
11
10
9
8
7
6
5
4
3
2
1
CP_kmeans
Fusion_kmeans
Pathway
VEGFR.KIT.PDGF
VEGFR
Topoisomerase
Sodium.Channe
Reverse.Transcripta
RAF
RAAS
Proteasome
PI3K
PDE
PARP
P450
mTOR
Microtubule.Associa
MEK
JAK
HSP
Histamine.Recept
HDAC
EGFR
DNA.RNA.Synthes
COX
CDK
Calcium.Channe
Aurora.Kinase
Adrenergic.Receptor.an
Adrenergic.Receptor.a
AChR.antagonis
5HTR.antagonis
5HTR.agonist

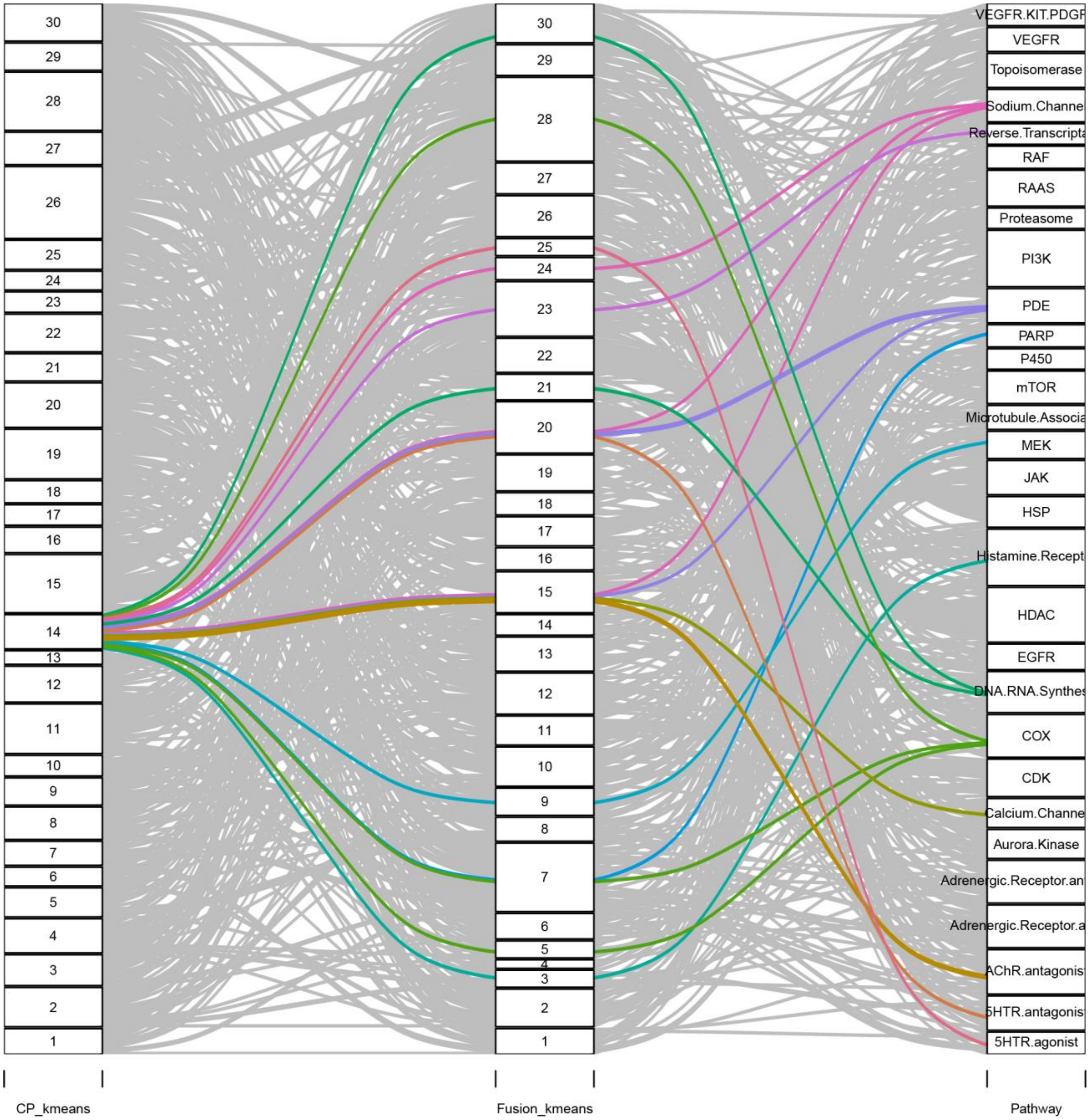
30
29
28
27
26
25
24
23
22
21
20
19
18
17
16
15
14
13
12
11
10
9
8
7
6
5
4
3
2
1
VEGFR.KIT.PDGF
VEGFR
Topoisomerase
Sodium.Channe
Reverse.Transcripta
RAF
RAAS
Proteasome
PI3K
PDE
PARP
P450
mTOR
Microtubule.Associa
MEK
JAK
HSP
Histamine.Recept
HDAC
EGFR
DNA.RNA.Synthes
COX
CDK
Calcium.Channe
Aurora.Kinase
Adrenergic.Receptor.an
Adrenergic.Receptor.a
AChR.antagonis
5HTR.antagonis
5HTR.agonist
CP_kmeans
Fusion_kmeans
Pathway

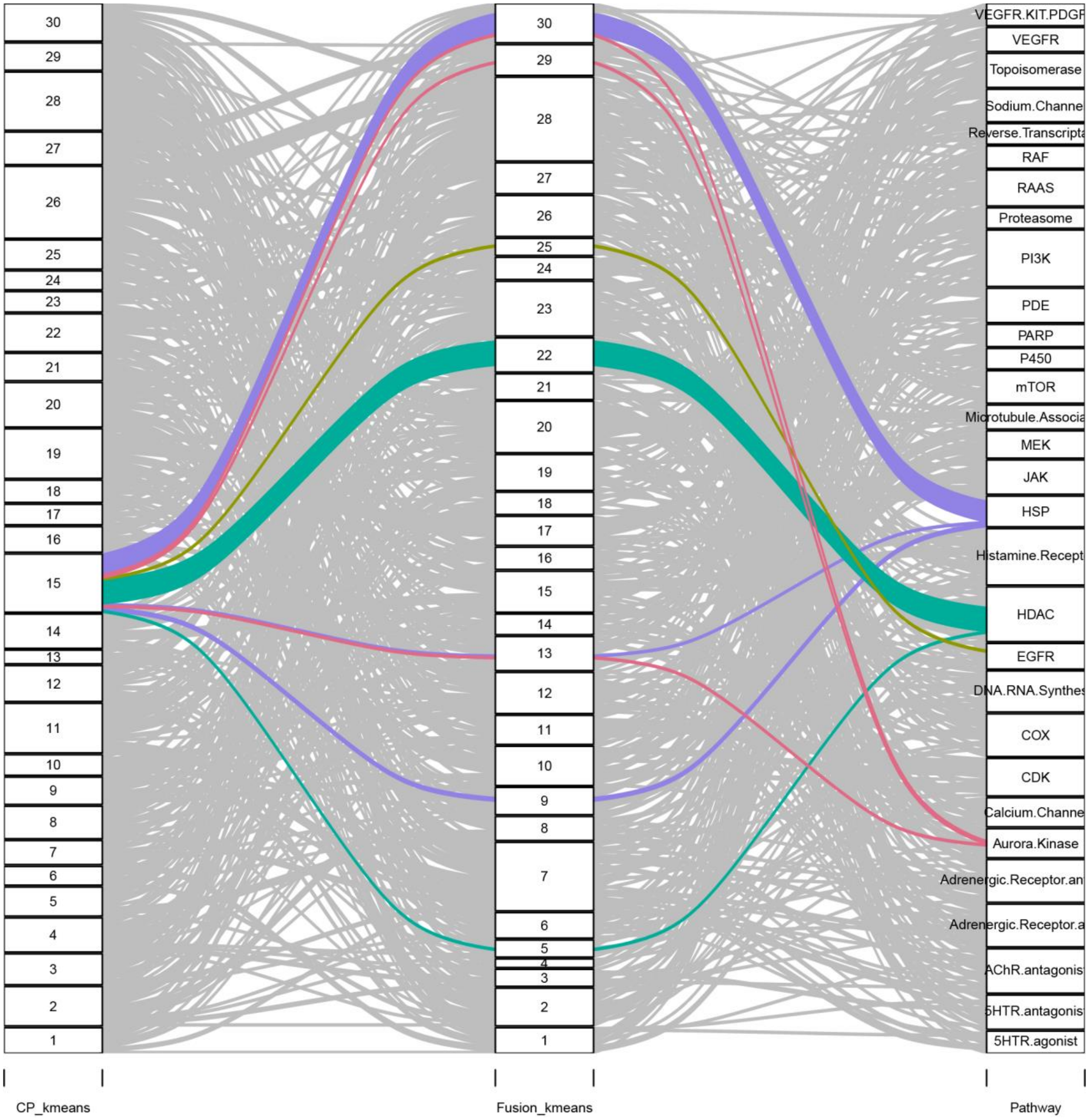
CP_kmeans
Fusion_kmeans
Pathway
VEGFR.KIT.PDGF
VEGFR
Topoisomerase
Sodium.Channe
Reverse.Transcripta
RAF
RAAS
Proteasome
PI3K
PDE
PARP
P450
mTOR
Microtubule.Associa
MEK
JAK
HSP
Histamine.Recept
HDAC
EGFR
DNA.RNA.Synthes
COX
CDK
Calcium.Channe
Aurora.Kinase
Adrenergic.Receptor.an
Adrenergic.Receptor.a
AChR.antagonis
5HTR.antagonis
5HTR.agonist

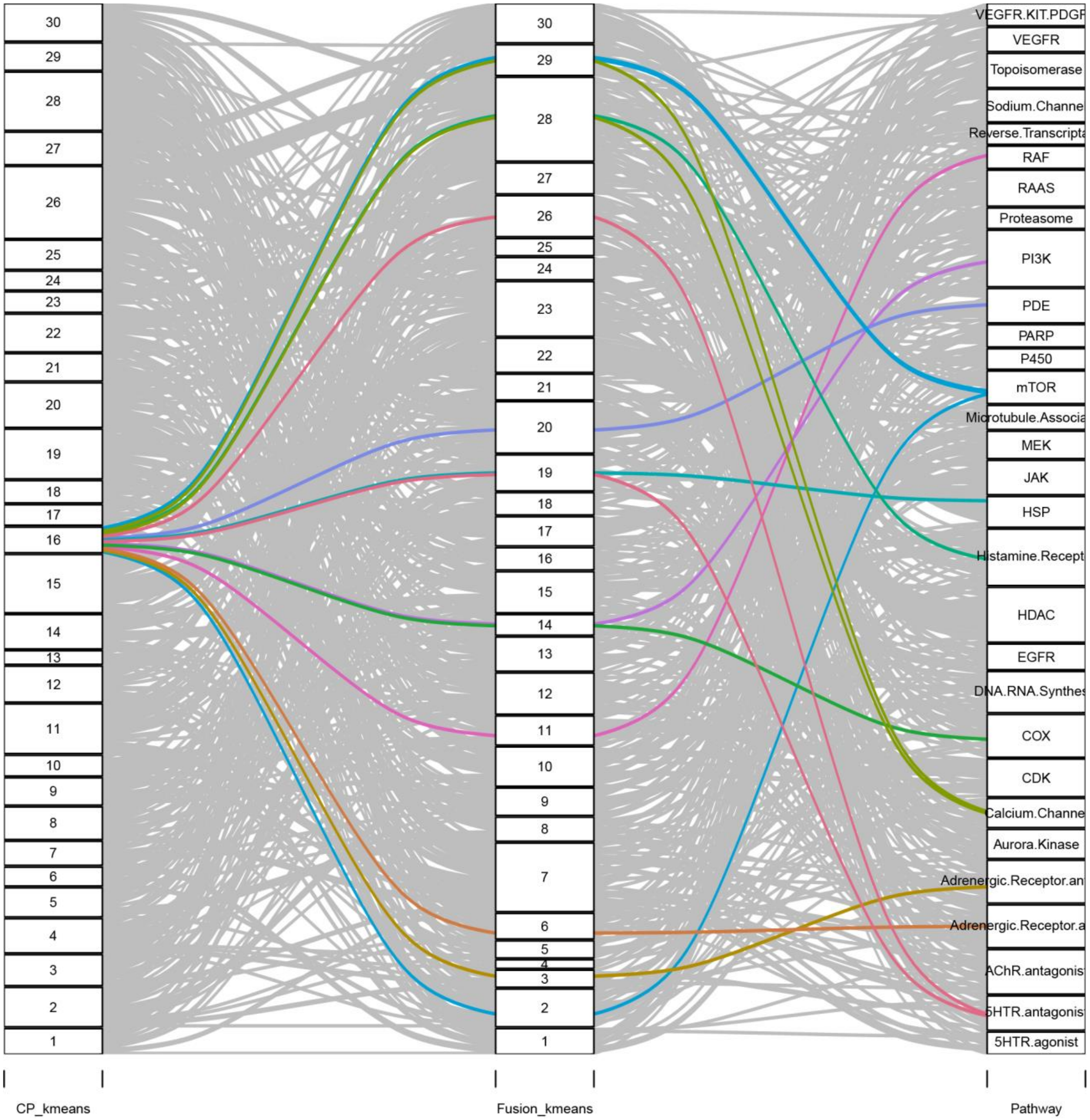

CP_kmeans
Fusion_kmeans
Pathway
30
29
28
27
26
25
24
23
22
21
20
19
18
17
16
15
14
13
12
11
10
9
8
7
6
5
4
3
2
1
VEGFR.KIT.PDGF
VEGFR
Topoisomerase
Sodium.Channe
Reverse.Transcripta
RAF
RAAS
Proteasome
PI3K
PDE
PARP
P450
mTOR
Microtubule.Associa
MEK
JAK
HSP
Histamine.Recept
HDAC
EGFR
DNA.RNA.Synthes
COX
CDK
Calcium.Channe
Aurora.Kinase
Adrenergic.Receptor.an
Adrenergic.Receptor.a
AChR.antagonis
5HTR.antagonis
5HTR.agonist

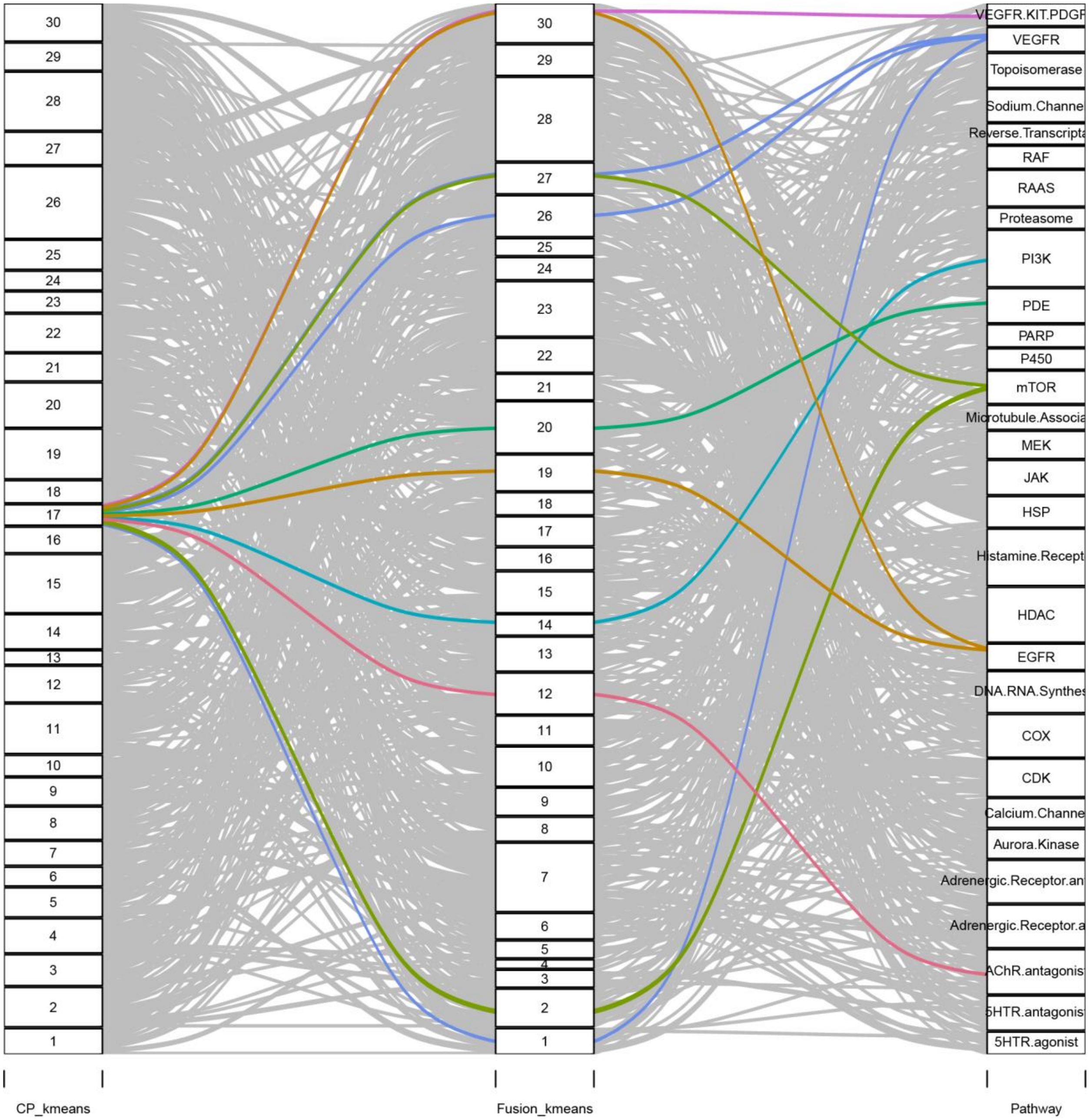

CP_kmeans
Fusion_kmeans
Pathway
VEGFR.KIT.PDGF
VEGFR
Topoisomerase
Sodium.Channe
Reverse.Transcripta
RAF
RAAS
Proteasome
PI3K
PDE
PARP
P450
mTOR
Microtubule.Associa
MEK
JAK
HSP
Histamine.Recept
HDAC
EGFR
DNA.RNA.Synthes
COX
CDK
Calcium.Channe
Aurora.Kinase
Adrenergic.Receptor.an
Adrenergic.Receptor.a
AChR.antagonis
5HTR.antagonis
5HTR.agonist

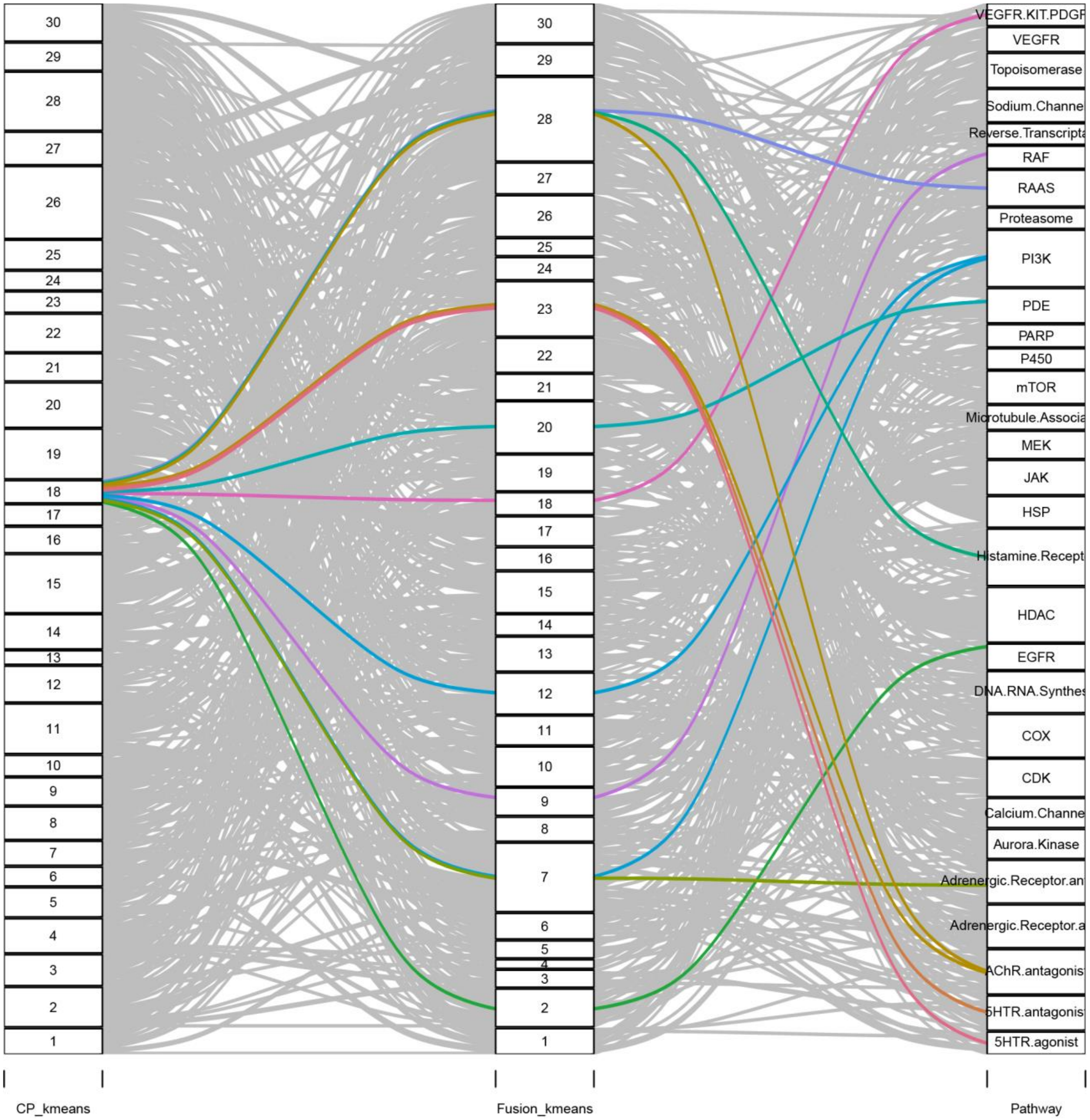
30
29
28
27
26
25
24
23
22
21
20
19
18
17
16
15
14
13
12
11
10
9
8
7
6
5
4
3
2
1
VEGFR.KIT.PDGF
VEGFR
Topoisomerase
Sodium.Channe
Reverse.Transcripta
RAF
RAAS
Proteasome
PI3K
PDE
PARP
P450
mTOR
Microtubule.Associa
MEK
JAK
HSP
Histamine.Recept
HDAC
EGFR
DNA.RNA.Synthes
COX
CDK
Calcium.Channe
Aurora.Kinase
Adrenergic.Receptor.an
Adrenergic.Receptor.a
AChR.antagonis
5HTR.antagonis
5HTR.agonist
CP_kmeans
Fusion_kmeans
Pathway

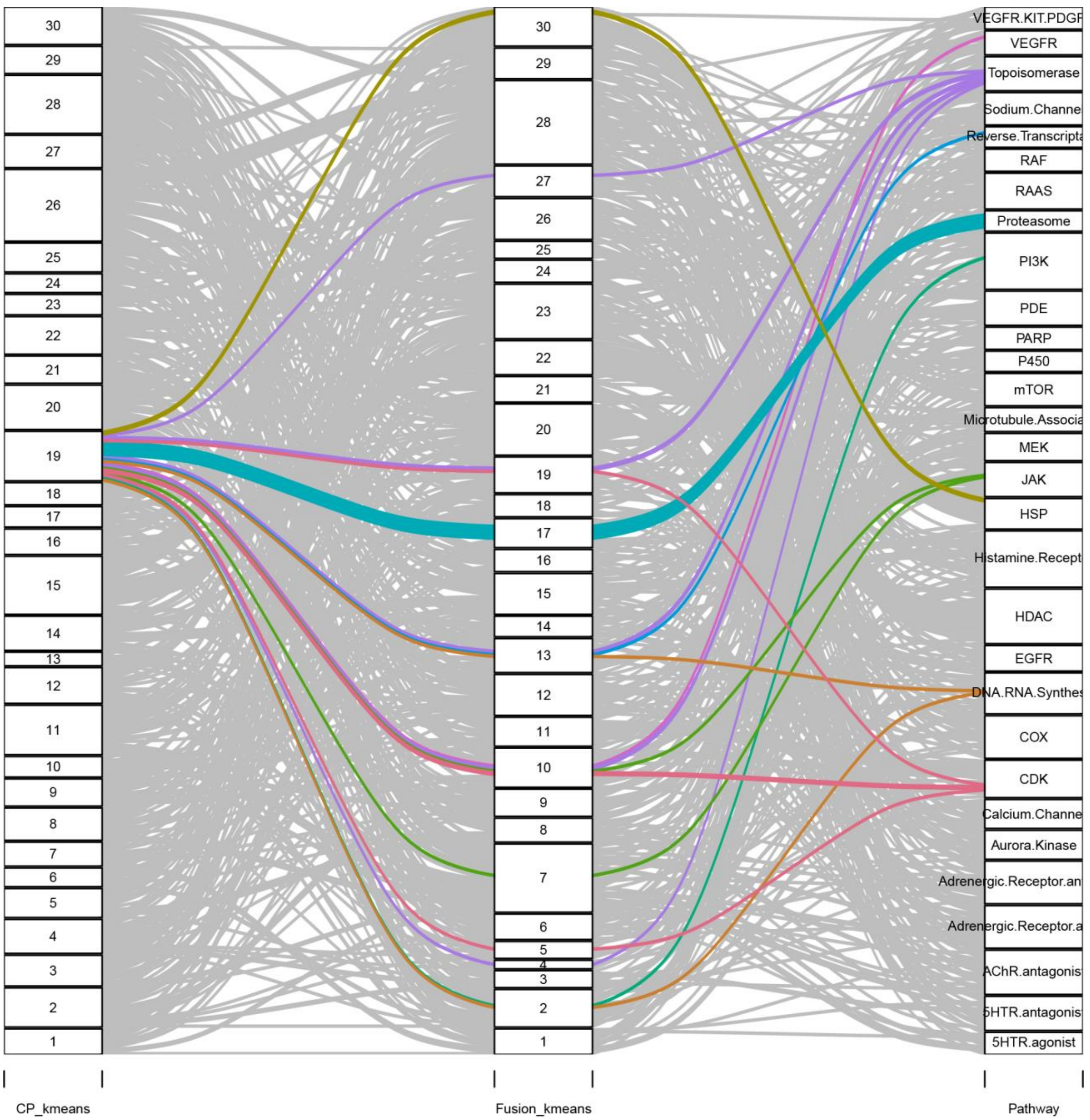
30
29
28
27
26
25
24
23
22
21
20
19
18
17
16
15
14
13
12
11
10
9
8
7
6
5
4
3
2
1
VEGFR.KIT.PDGF
VEGFR
Topoisomerase
Sodium.Channe
Reverse.Transcripta
RAF
RAAS
Proteasome
PI3K
PDE
PARP
P450
mTOR
Microtubule.Associa
MEK
JAK
HSP
Histamine.Recept
HDAC
EGFR
DNA.RNA.Synthes
COX
CDK
Calcium.Channe
Aurora.Kinase
Adrenergic.Receptor.an
Adrenergic.Receptor.a
AChR.antagonis
5HTR.antagonis
5HTR.agonist
CP_kmeans
Fusion_kmeans
Pathway

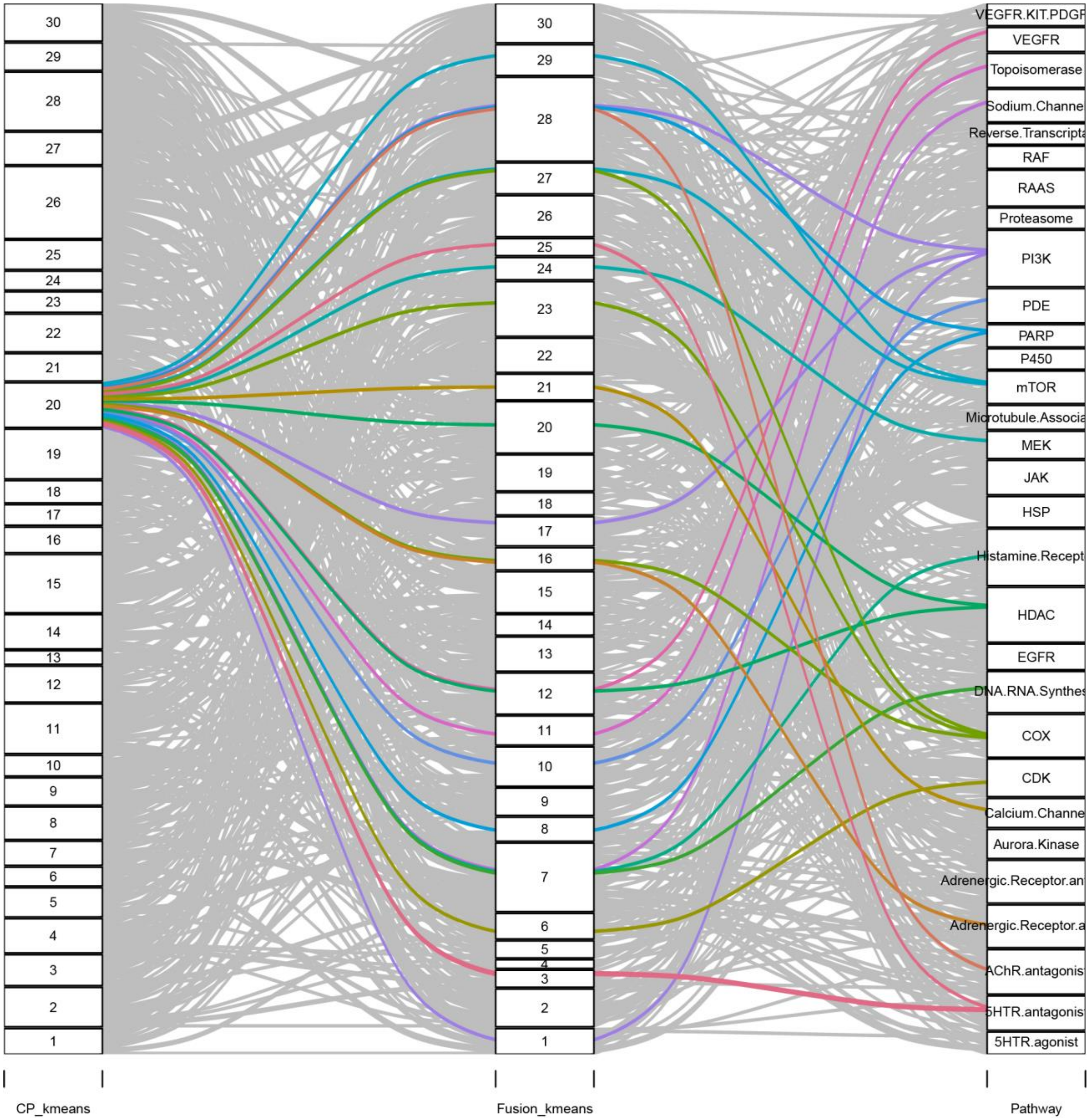
CP_kmeans
Fusion_kmeans
Pathway
30
29
28
27
26
25
24
23
22
21
20
19
18
17
16
15
14
13
12
11
10
9
8
7
6
5
4
3
2
1
VEGFR.KIT.PDGF
VEGFR
Topoisomerase
Sodium.Channe
Reverse.Transcripta
RAF
RAAS
Proteasome
PI3K
PDE
PARP
P450
mTOR
Microtubule.Associa
MEK
JAK
HSP
Histamine.Recept
HDAC
EGFR
DNA.RNA.Synthes
COX
CDK
Calcium.Channe
Aurora.Kinase
Adrenergic.Receptor.an
Adrenergic.Receptor.a
AChR.antagonis
5HTR.antagonis
5HTR.agonist

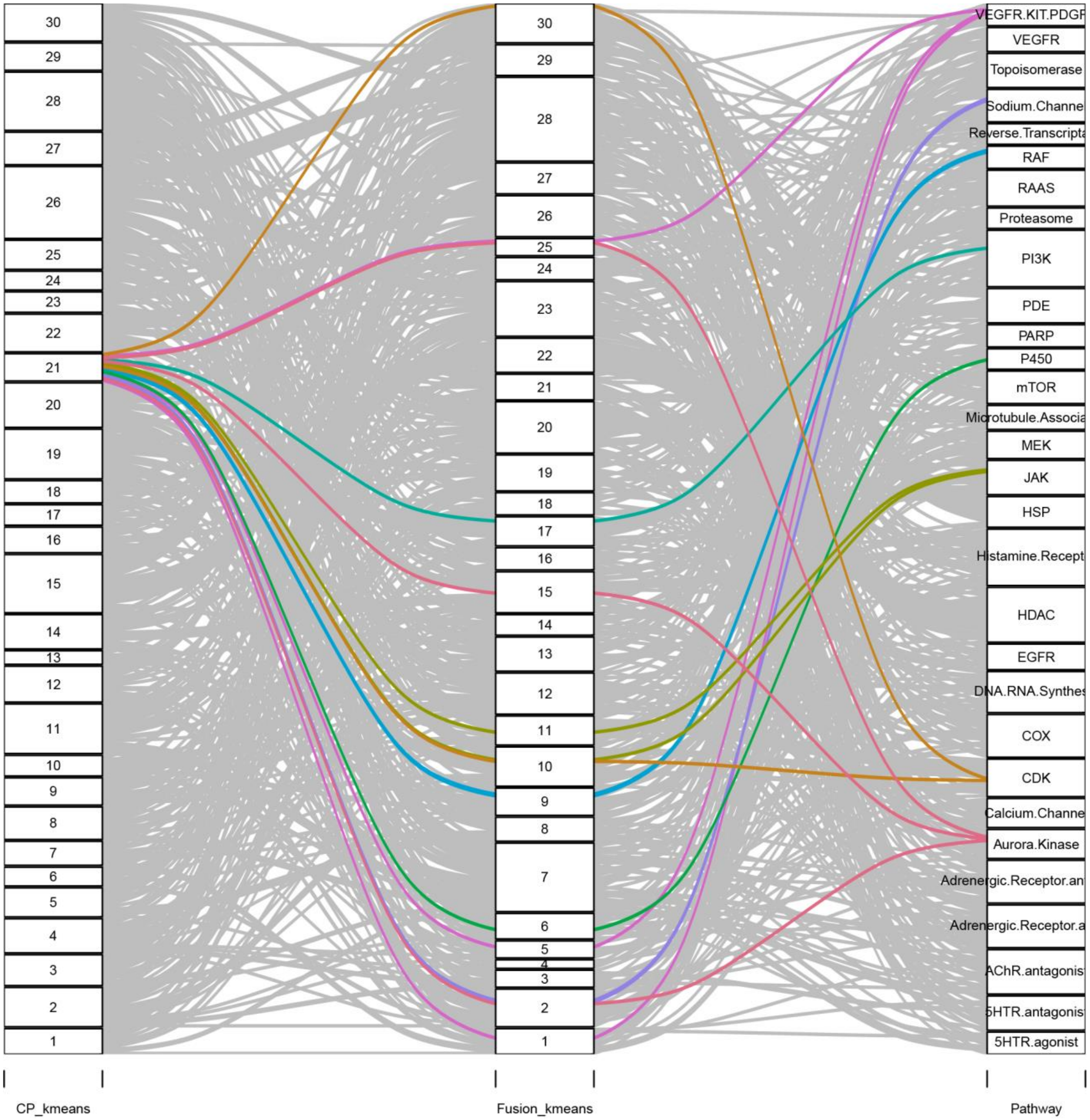

30
29
28
27
26
25
24
23
22
21
20
19
18
17
16
15
14
13
12
11
10
9
8
7
6
5
4
3
2
1
CP_kmeans
Fusion_kmeans
Pathway
VEGFR.KIT.PDGF
VEGFR
Topoisomerase
Sodium.Channe
Reverse.Transcripta
RAF
RAAS
Proteasome
PI3K
PDE
PARP
P450
mTOR
Microtubule.Associa
MEK
JAK
HSP
Histamine.Recept
HDAC
EGFR
DNA.RNA.Synthes
COX
CDK
Calcium.Channe
Aurora.Kinase
Adrenergic.Receptor.an
Adrenergic.Receptor.a
AChR.antagonis
5HTR.antagonis
5HTR.agonist

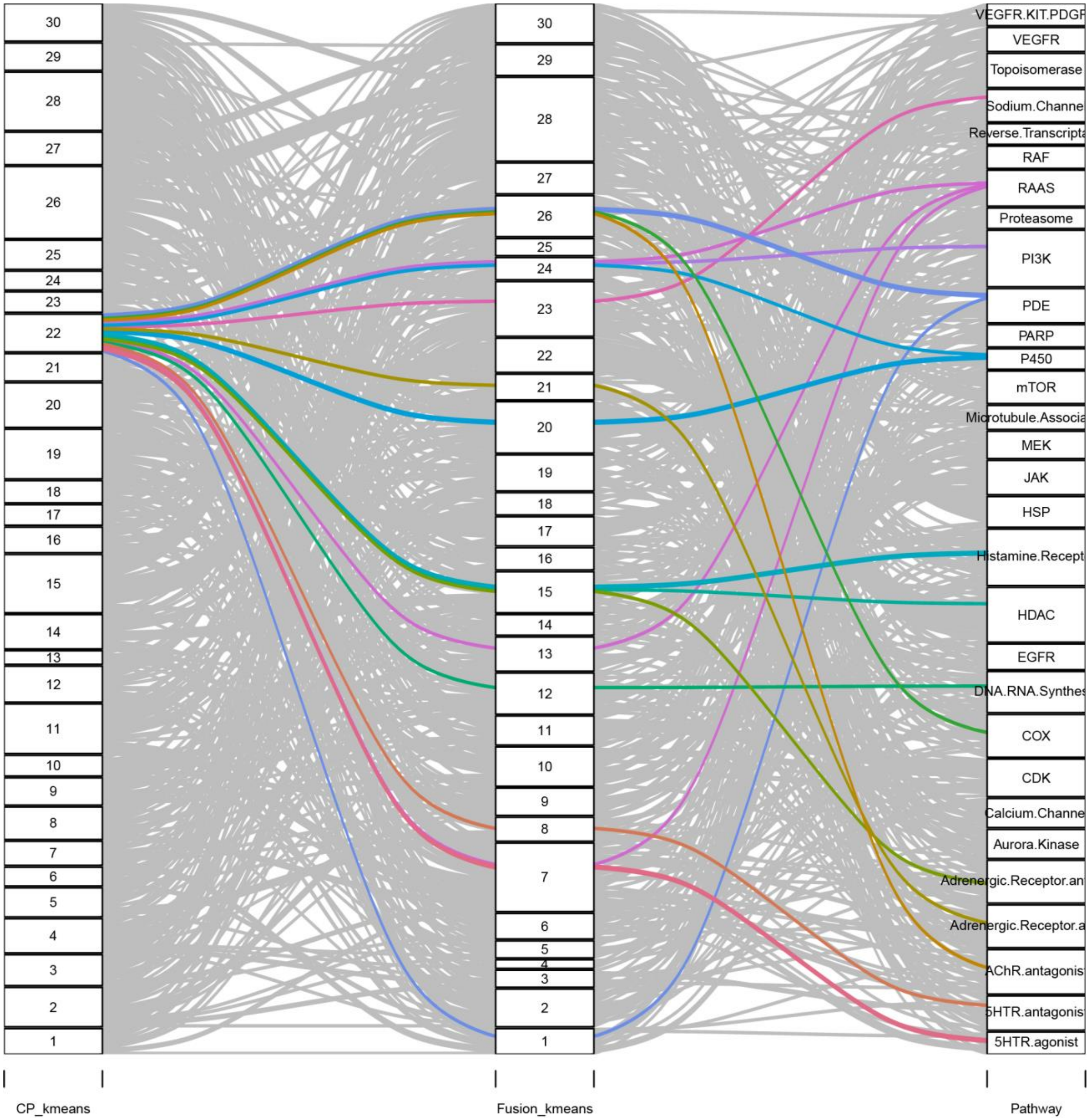

30
29
28
27
26
25
24
23
22
21
20
19
18
17
16
15
14
13
12
11
10
9
8
7
6
5
4
3
2
1
VEGFR.KIT.PDGF
VEGFR
Topoisomerase
Sodium.Channe
Reverse.Transcripta
RAF
RAAS
Proteasome
PI3K
PDE
PARP
P450
mTOR
Microtubule.Associa
MEK
JAK
HSP
Histamine.Recept
HDAC
EGFR
DNA.RNA.Synthes
COX
CDK
Calcium.Channe
Aurora.Kinase
Adrenergic.Receptor.an
Adrenergic.Receptor.a
AChR.antagonis
5HTR.antagonis
5HTR.agonist
CP_kmeans
Fusion_kmeans
Pathway

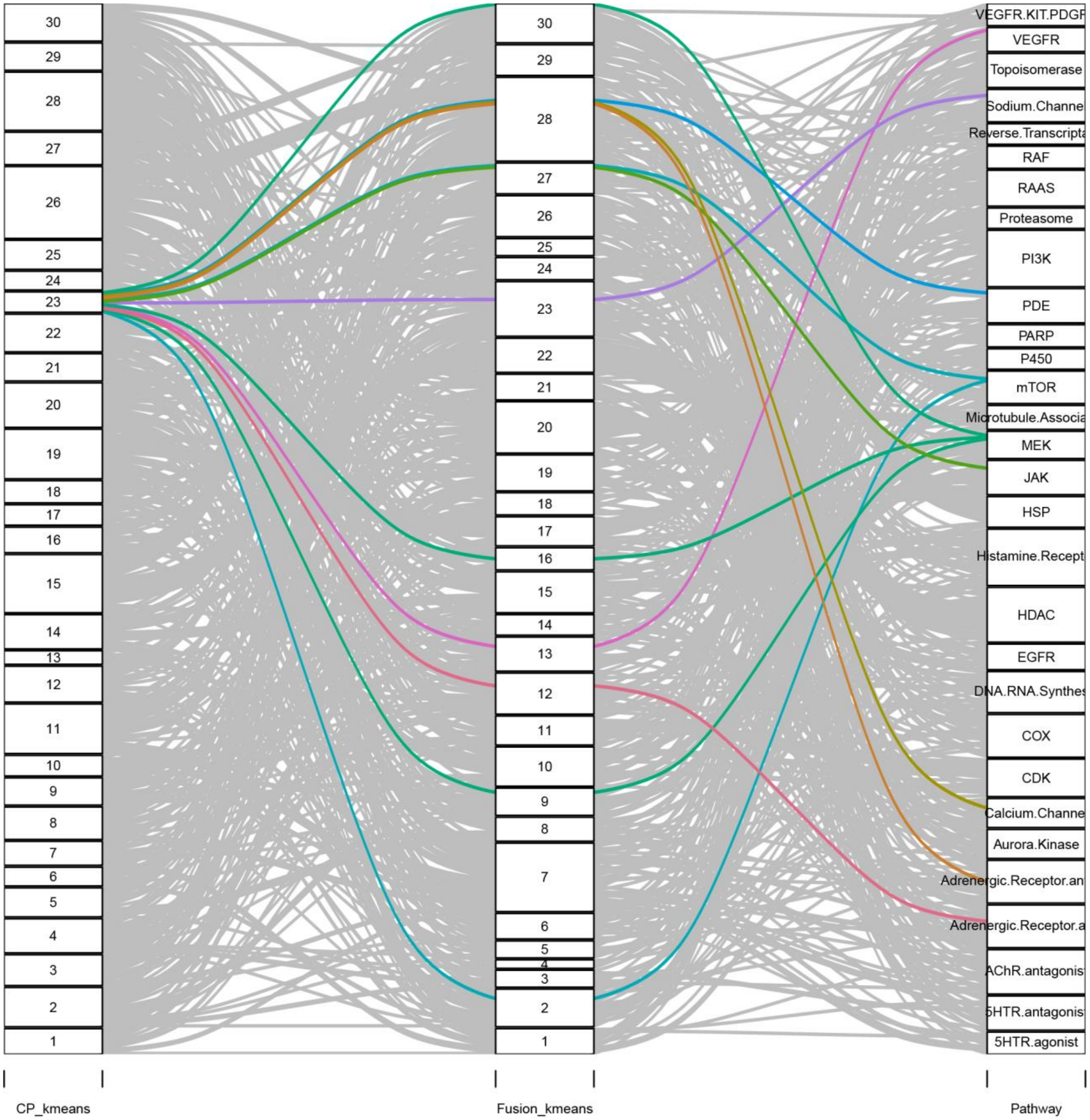
30
29
28
27
26
25
24
23
22
21
20
19
18
17
16
15
14
13
12
11
10
9
8
7
6
5
4
3
2
1
VEGFR.KIT.PDGF
VEGFR
Topoisomerase
Sodium.Channe
Reverse.Transcripta
RAF
RAAS
Proteasome
PI3K
PDE
PARP
P450
mTOR
Microtubule.Associa
MEK
JAK
HSP
Histamine.Recept
HDAC
EGFR
DNA.RNA.Synthes
COX
CDK
Calcium.Channe
Aurora.Kinase
Adrenergic.Receptor.an
Adrenergic.Receptor.a
AChR.antagonis
5HTR.antagonis
5HTR.agonist
CP_kmeans
Fusion_kmeans
Pathway

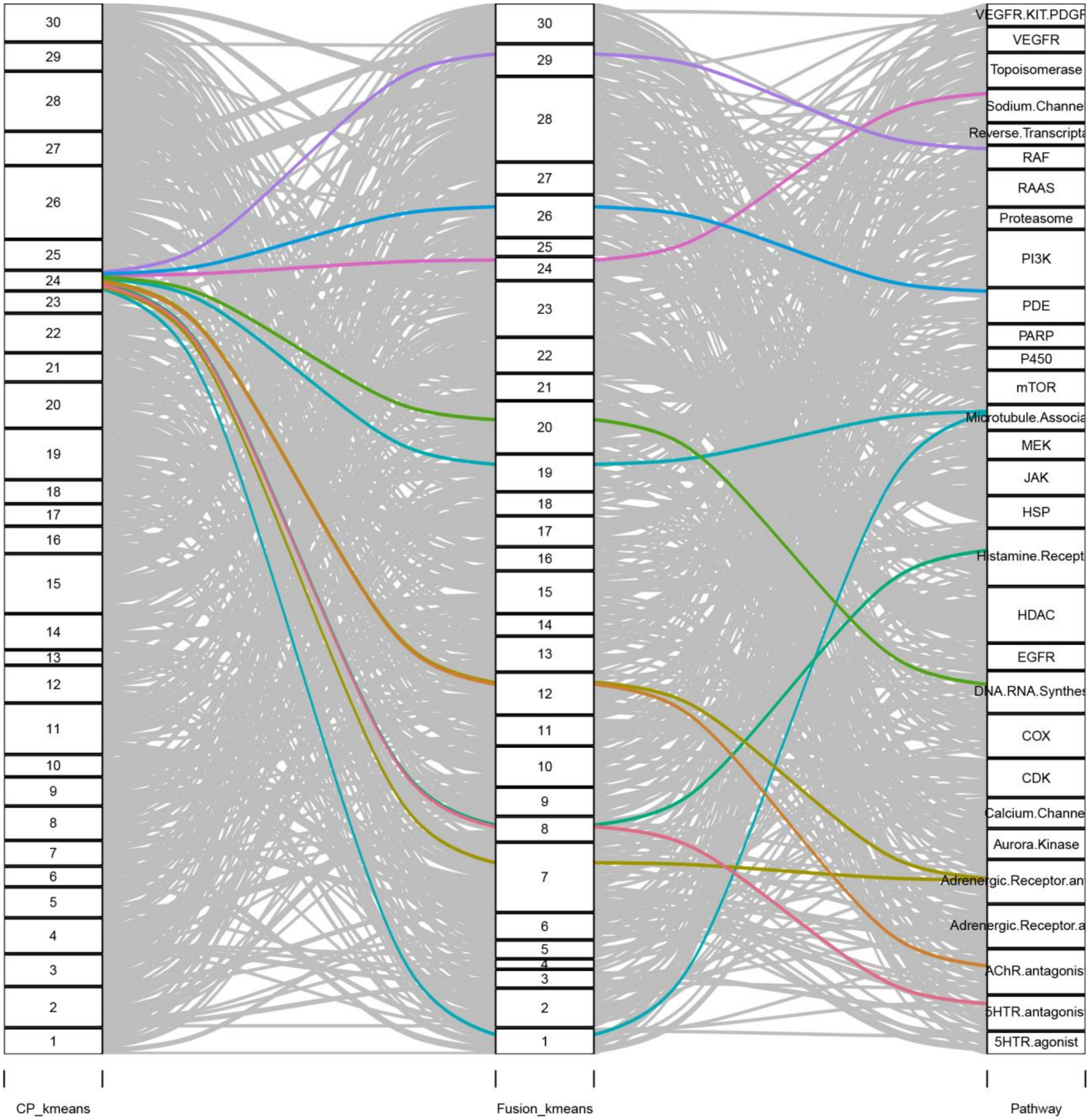
30
29
28
27
26
25
24
23
22
21
20
19
18
17
16
15
14
13
12
11
10
9
8
7
6
5
4
3
2
1
VEGFR.KIT.PDGF
VEGFR
Topoisomerase
Sodium.Channe
Reverse.Transcripta
RAF
RAAS
Proteasome
PI3K
PDE
PARP
P450
mTOR
Microtubule.Associa
MEK
JAK
HSP
Histamine.Recept
HDAC
EGFR
DNA.RNA.Synthes
COX
CDK
Calcium.Channe
Aurora.Kinase
Adrenergic.Receptor.an
Adrenergic.Receptor.a
AChR.antagonis
5HTR.antagonis
5HTR.agonist
CP_kmeans
Fusion_kmeans
Pathway

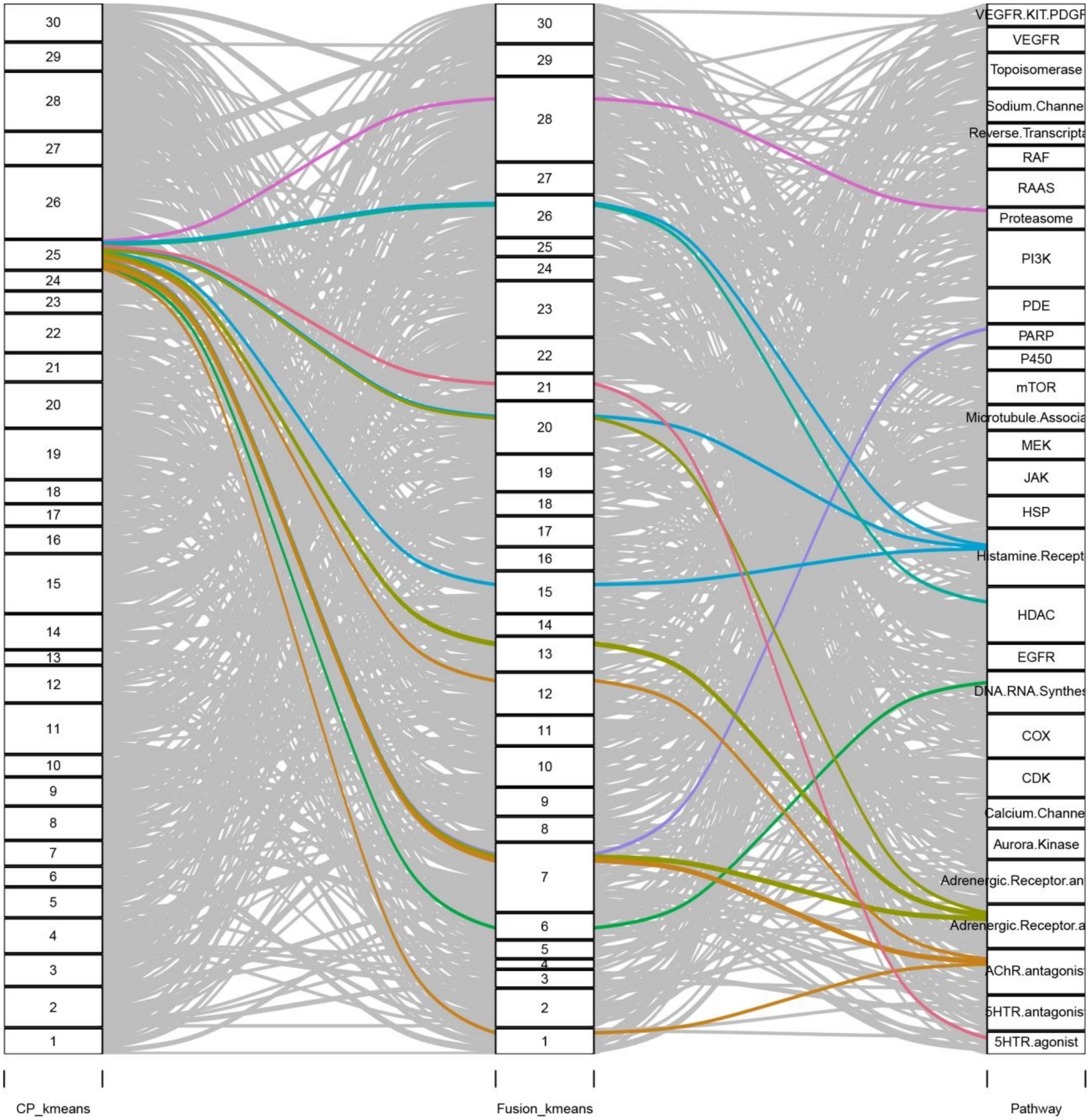
CP_kmeans
Fusion_kmeans
Pathway
VEGFR.KIT.PDGF
VEGFR
Topoisomerase
Sodium.Channe
Reverse.Transcripta
RAF
RAAS
Proteasome
PI3K
PDE
PARP
P450
mTOR
Microtubule.Associa
MEK
JAK
HSP
Histamine.Recept
HDAC
EGFR
DNA.RNA.Synthes
COX
CDK
Calcium.Channe
Aurora.Kinase
Adrenergic.Receptor.an
Adrenergic.Receptor.a
AChR.antagonis
5HTR.antagonis
5HTR.agonist

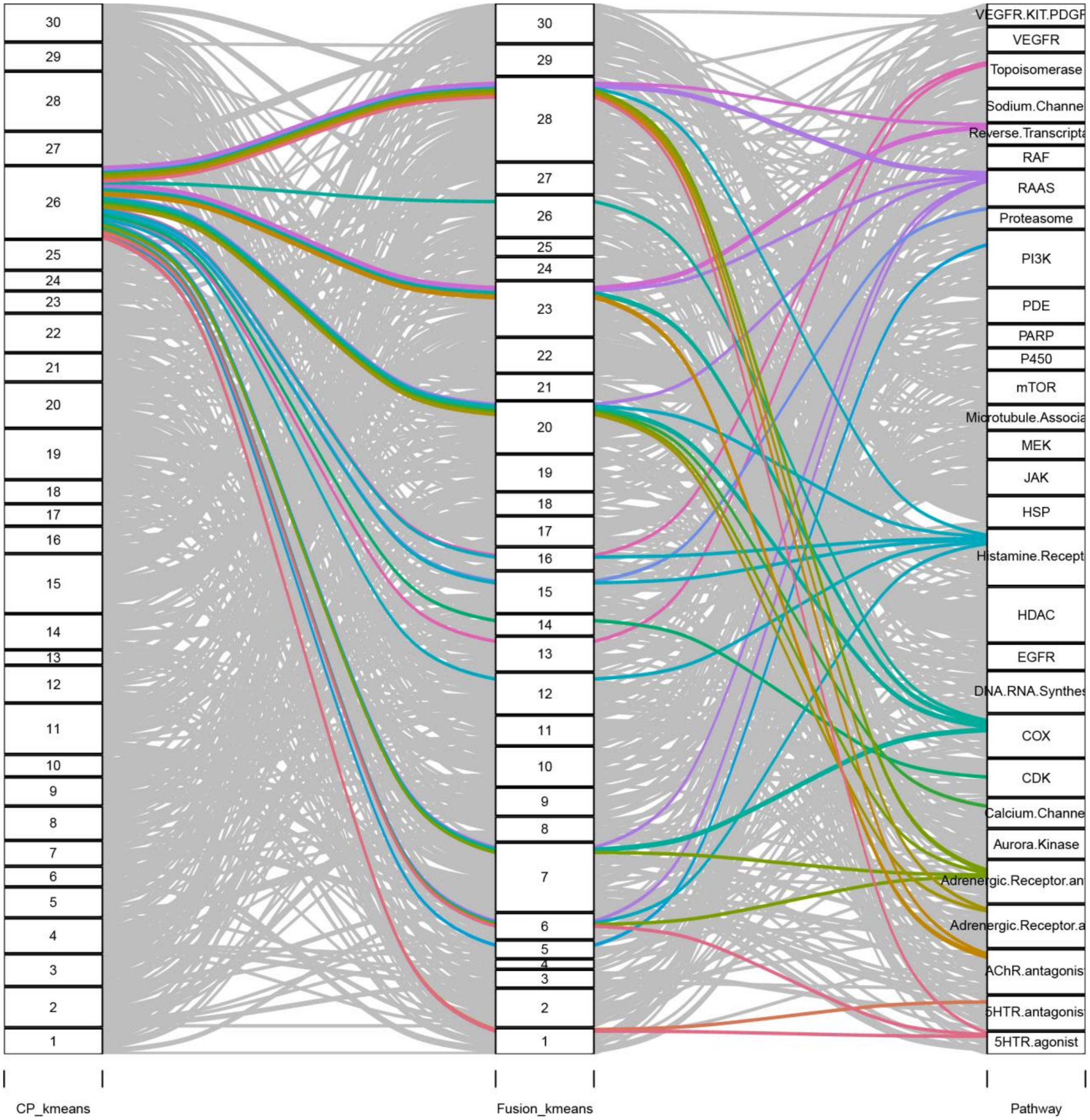

CP_kmeans
Fusion_kmeans
Pathway
VEGFR.KIT.PDGF
VEGFR
Topoisomerase
Sodium.Channe
Reverse.Transcripta
RAF
RAAS
Proteasome
PI3K
PDE
PARP
P450
mTOR
Microtubule.Associa
MEK
JAK
HSP
Histamine.Recept
HDAC
EGFR
DNA.RNA.Synthes
COX
CDK
Calcium.Channe
Aurora.Kinase
Adrenergic.Receptor.an
Adrenergic.Receptor.a
AChR.antagonis
5HTR.antagonis
5HTR.agonist

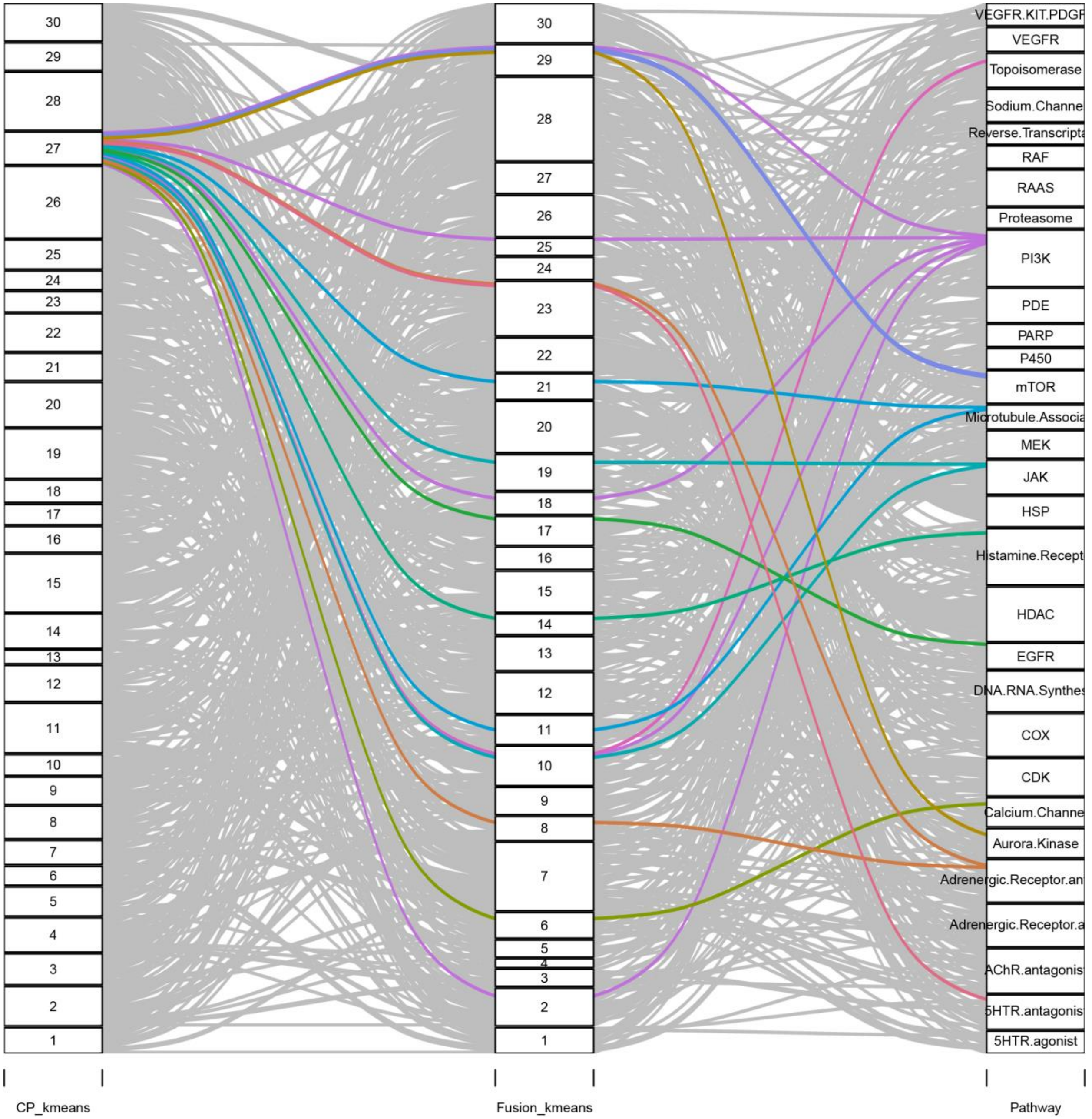

30
29
28
27
26
25
24
23
22
21
20
19
18
17
16
15
14
13
12
11
10
9
8
7
6
5
4
3
2
1
VEGFR.KIT.PDGF
VEGFR
Topoisomerase
Sodium.Channe
Reverse.Transcripta
RAF
RAAS
Proteasome
PI3K
PDE
PARP
P450
mTOR
Microtubule.Associa
MEK
JAK
HSP
Histamine.Recept
HDAC
EGFR
DNA.RNA.Synthes
COX
CDK
Calcium.Channe
Aurora.Kinase
Adrenergic.Receptor.an
Adrenergic.Receptor.a
AChR.antagonis
5HTR.antagonis
5HTR.agonist
CP_kmeans
Fusion_kmeans
Pathway

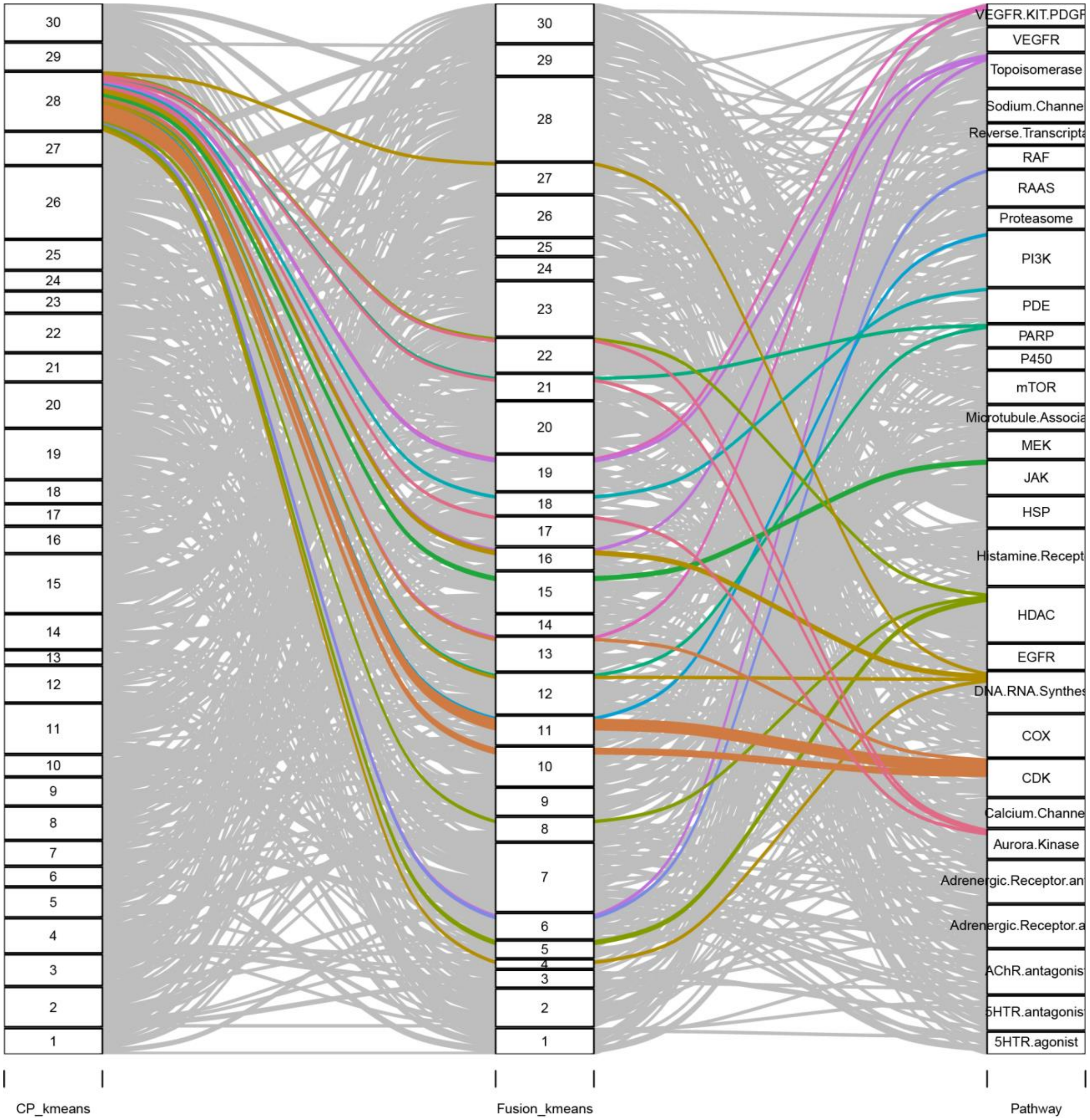
30
29
28
27
26
25
24
23
22
21
20
19
18
17
16
15
14
13
12
11
10
9
8
7
6
5
4
3
2
1
VEGFR.KIT.PDGF
VEGFR
Topoisomerase
Sodium.Channe
Reverse.Transcripta
RAF
RAAS
Proteasome
PI3K
PDE
PARP
P450
mTOR
Microtubule.Associa
MEK
JAK
HSP
Histamine.Recept
HDAC
EGFR
DNA.RNA.Synthes
COX
CDK
Calcium.Channe
Aurora.Kinase
Adrenergic.Receptor.an
Adrenergic.Receptor.a
AChR.antagonis
5HTR.antagonis
5HTR.agonist
CP_kmeans
Fusion_kmeans
Pathway

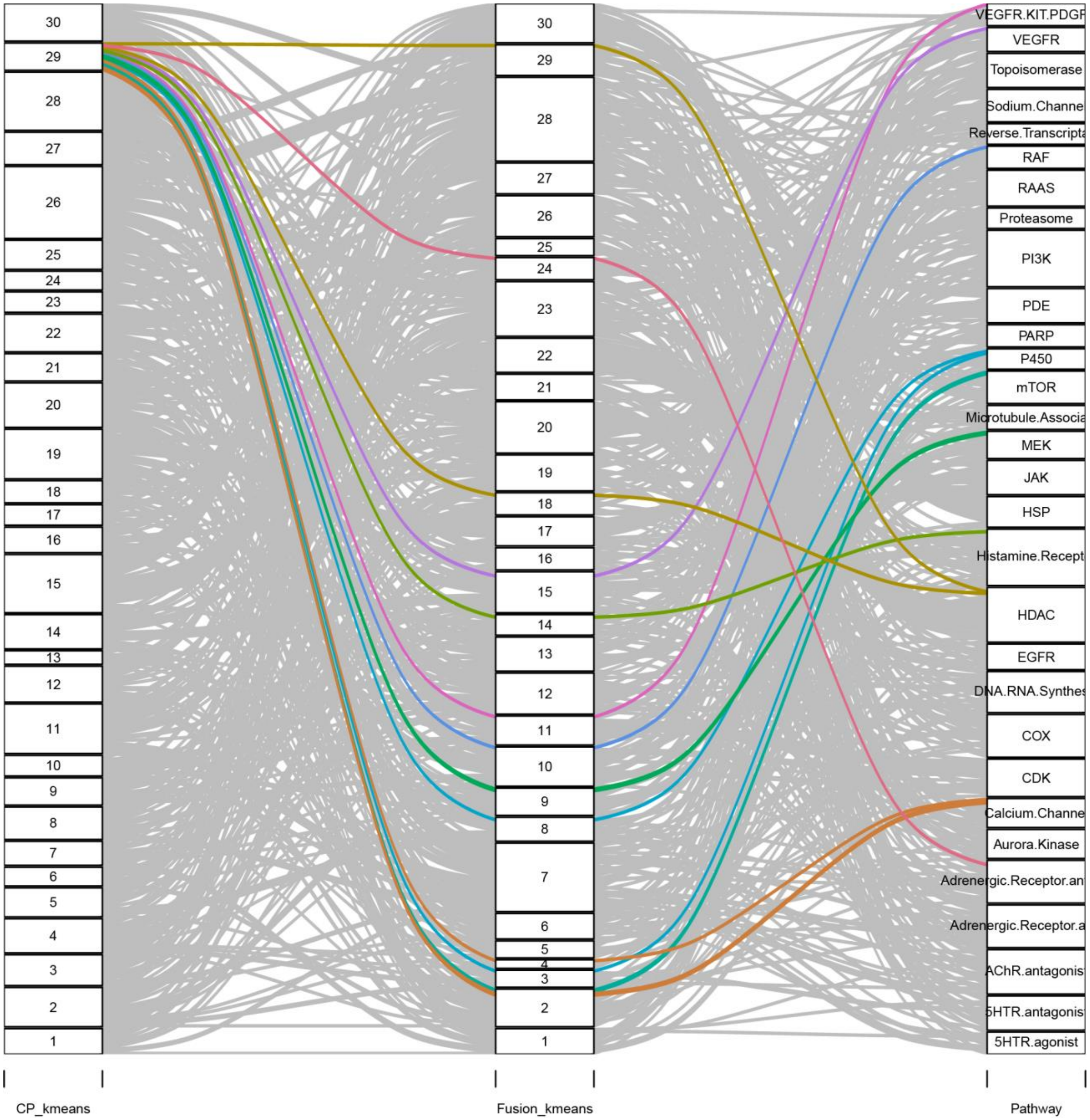
CP_kmeans
Fusion_kmeans
Pathway
30
29
28
27
26
25
24
23
22
21
20
19
18
17
16
15
14
13
12
11
10
9
8
7
6
5
4
3
2
1
VEGFR.KIT.PDGF
VEGFR
Topoisomerase
Sodium.Channe
Reverse.Transcripta
RAF
RAAS
Proteasome
PI3K
PDE
PARP
P450
mTOR
Microtubule.Associa
MEK
JAK
HSP
Histamine.Recept
HDAC
EGFR
DNA.RNA.Synthes
COX
CDK
Calcium.Channe
Aurora.Kinase
Adrenergic.Receptor.an
Adrenergic.Receptor.a
AChR.antagonis
5HTR.antagonis
5HTR.agonist

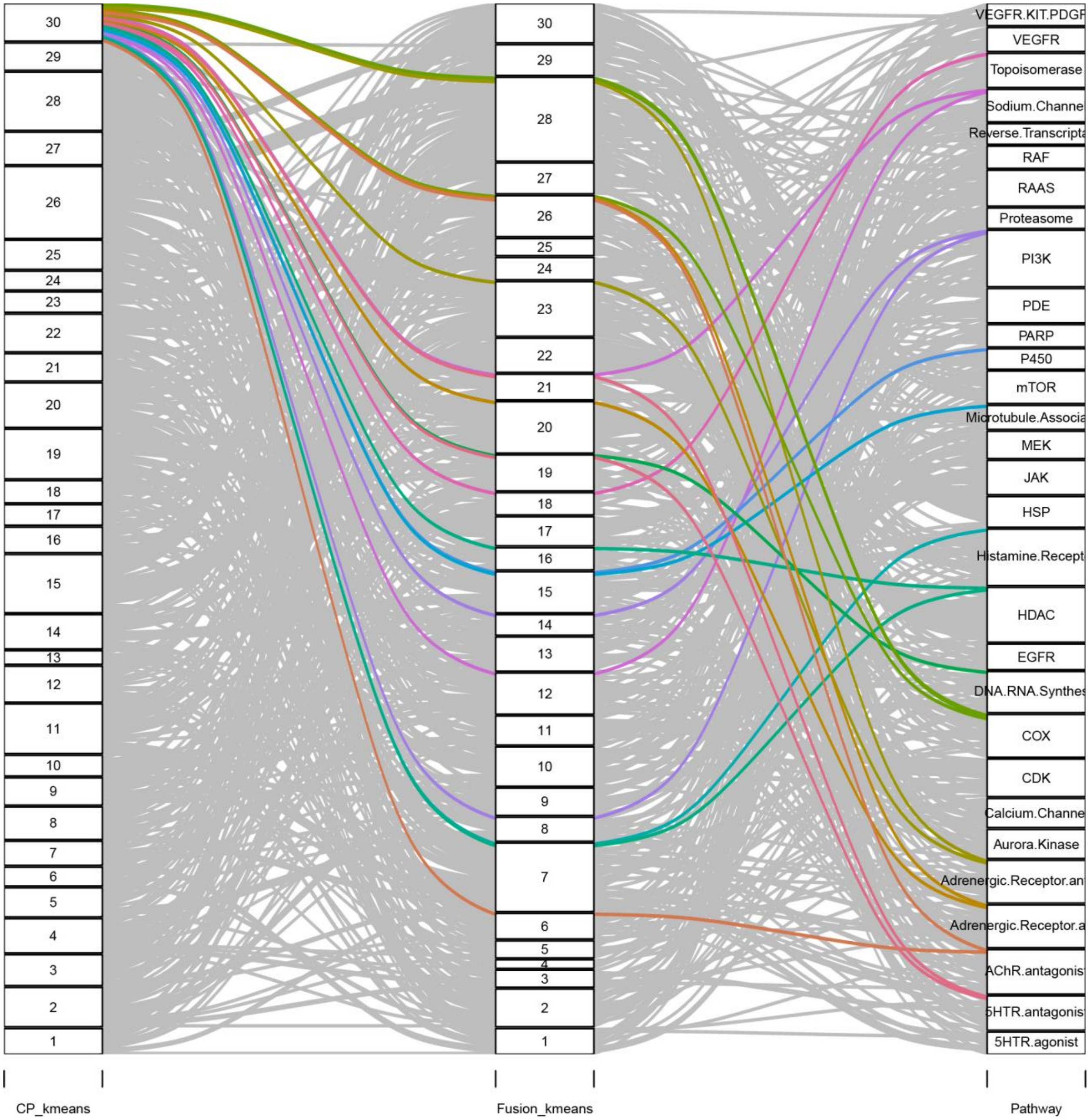
CP_kmeans
Fusion_kmeans
Pathway
30
29
28
27
26
25
24
23
22
21
20
19
18
17
16
15
14
13
12
11
10
9
8
7
6
5
4
3
2
1
VEGFR.KIT.PDGF
VEGFR
Topoisomerase
Sodium.Channe
Reverse.Transcripta
RAF
RAAS
Proteasome
PI3K
PDE
PARP
P450
mTOR
Microtubule.Associa
MEK
JAK
HSP
Histamine.Recept
HDAC
EGFR
DNA.RNA.Synthes
COX
CDK
Calcium.Channe
Aurora.Kinase
Adrenergic.Receptor.an
Adrenergic.Receptor.a
AChR.antagonis
5HTR.antagonis
5HTR.agonist

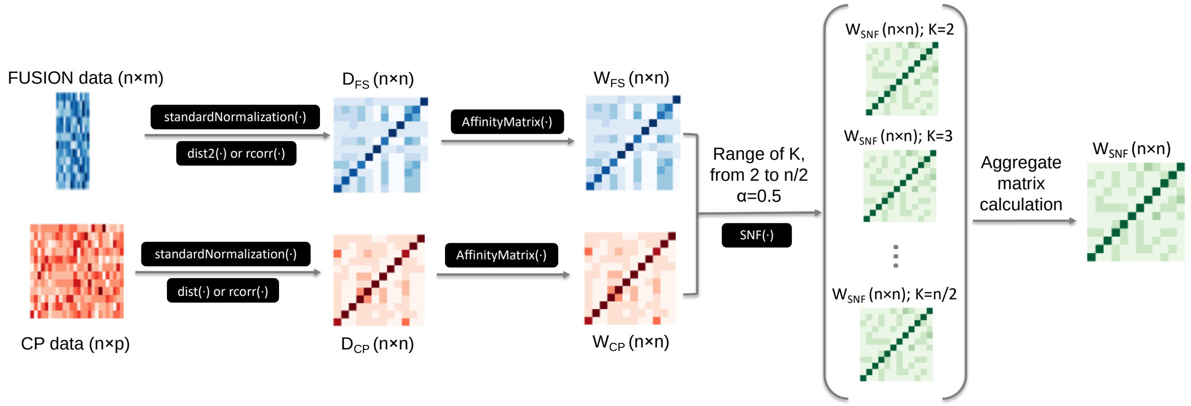

**Supplementary Figure 7: Adapted similarity network fusion workflow.**
All functions are from the SNFtool R package unless otherwise indicated. Normalized FUSION expression signatures and CP fingerprints are subjected to Z-score transformation by the standardNormalization function, followed by similarity matrix calculation using either the dist2 function for Euclidean Distance, or distanceMatrix function (ClassDiscovery package) for pearson distance. Affinity matrices were then calculated by the AffinityMatrix function. Similarity network fusion (K=number of neighbors, α=hyperparameter) is applied to $W_{FS}$ and $W_{CP}$ across a range of K values from 2 to n/2 to generate a set of $W_{SNF\text{-}K}$ matrices. An aggregate matrix is then calculated as described in Supplemental Note 4.

**a**

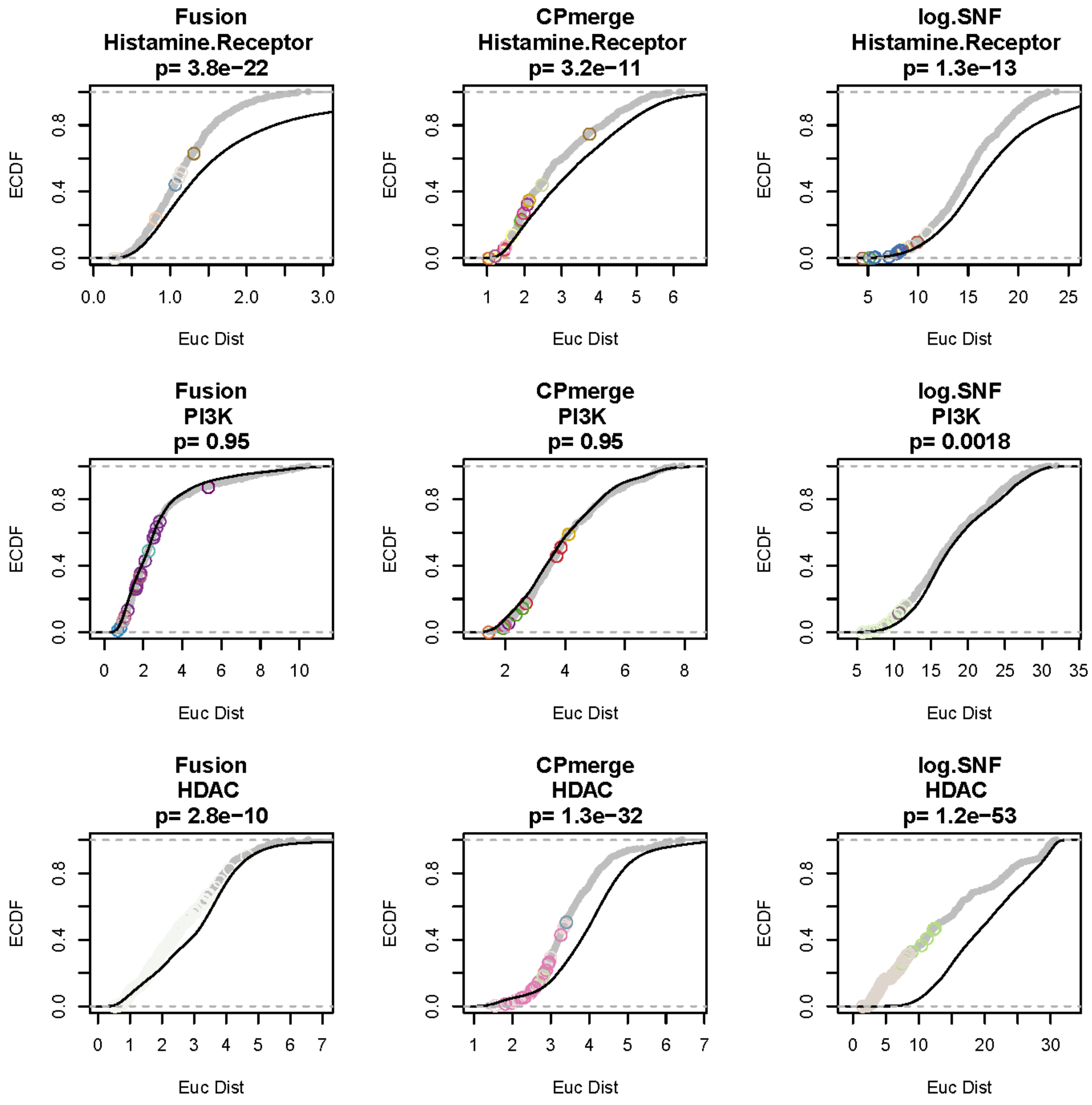

**Supplementary Figure 8: CDF plots comparing pairwise associations between in-class and out-of-class associations.** In-class-associations are colored in gray, with colored circles corresponding to cluster membership in the associated APC maps. Out-of-class associations are colored in black. KS-test p-values are shown above each plot. (**a**) CDF plots comparing pairwise Euclidean distances and (**b**) pairwise Pearson correlations.

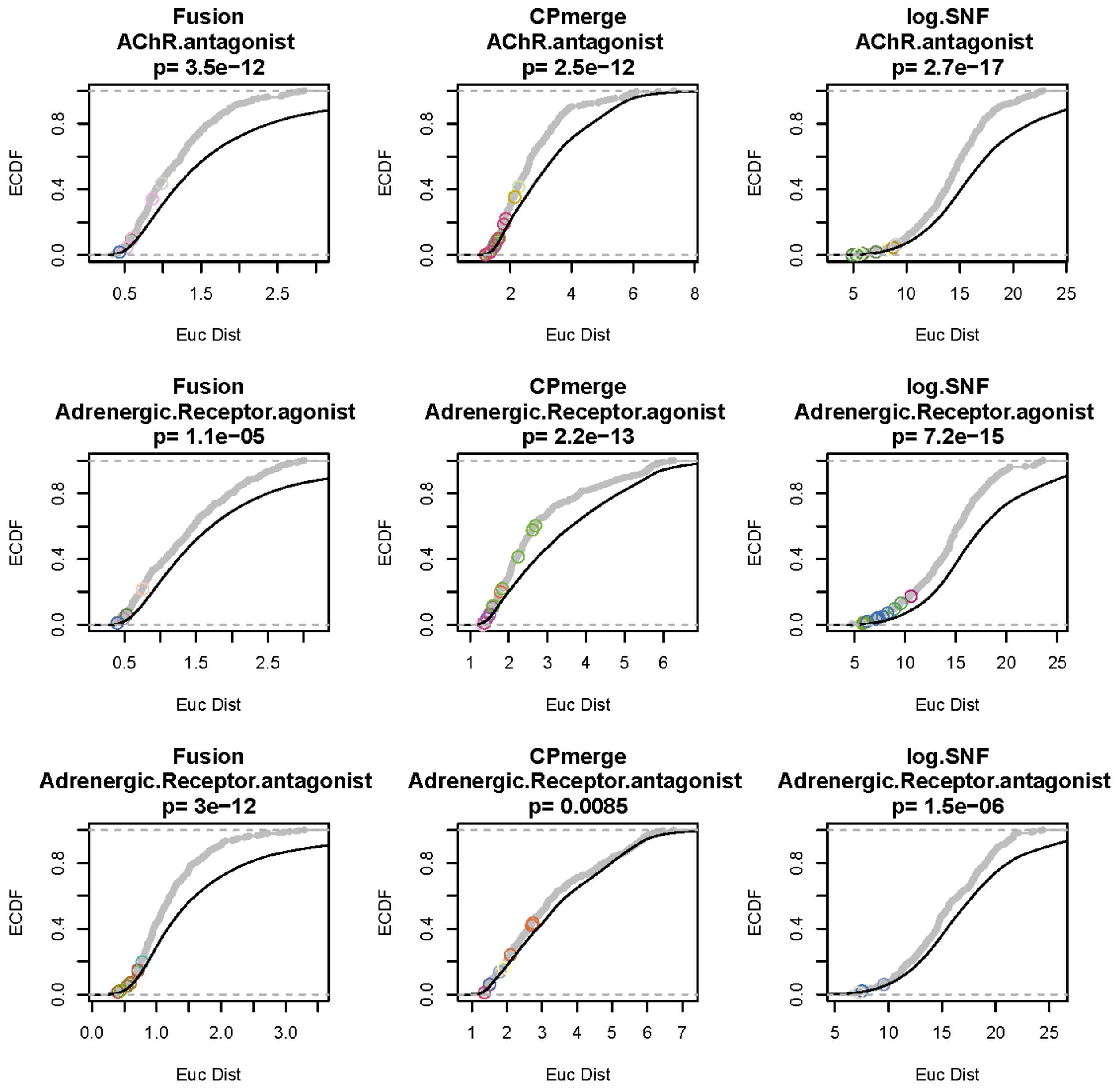
Fusion
AChR.antagonist
p= 3.5e−12
CPmerge
AChR.antagonist
p= 2.5e−12
log.SNF
AChR.antagonist
p= 2.7e−17
Fusion
Adrenergic.Receptor.agonist
p= 1.1e−05
CPmerge
Adrenergic.Receptor.agonist
p= 2.2e−13
log.SNF
Adrenergic.Receptor.agonist
p= 7.2e−15
Fusion
Adrenergic.Receptor.antagonist
p= 3e−12
CPmerge
Adrenergic.Receptor.antagonist
p= 0.0085
log.SNF
Adrenergic.Receptor.antagonist
p= 1.5e−06
ECDF
Euc Dist

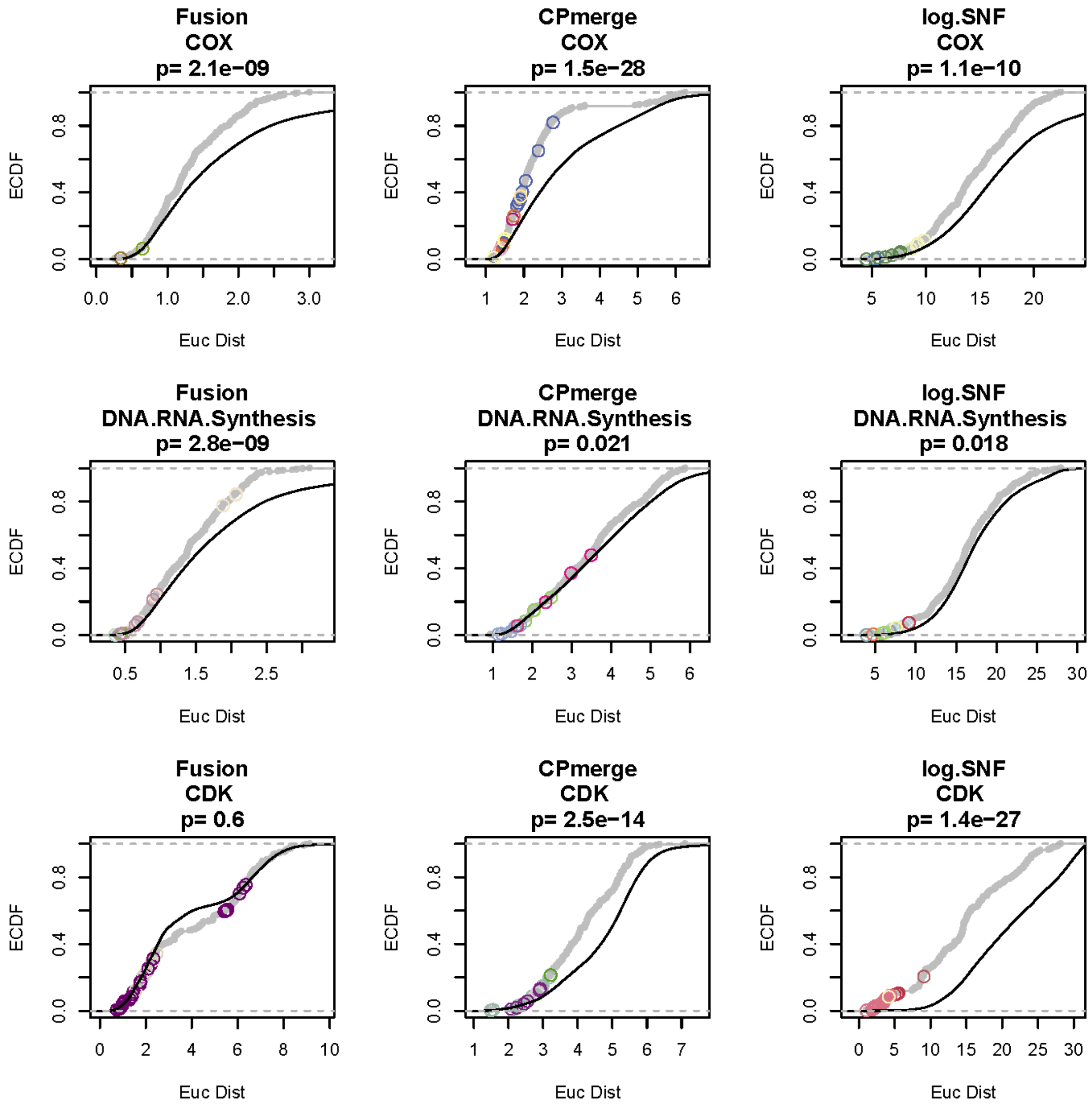
Fusion
COX
p= 2.1e−09
CPmerge
COX
p= 1.5e−28
log.SNF
COX
p= 1.1e−10
Fusion
DNA.RNA.Synthesis
p= 2.8e−09
CPmerge
DNA.RNA.Synthesis
p= 0.021
log.SNF
DNA.RNA.Synthesis
p= 0.018
Fusion
CDK
p= 0.6
CPmerge
CDK
p= 2.5e−14
log.SNF
CDK
p= 1.4e−27
ECDF
Euc Dist

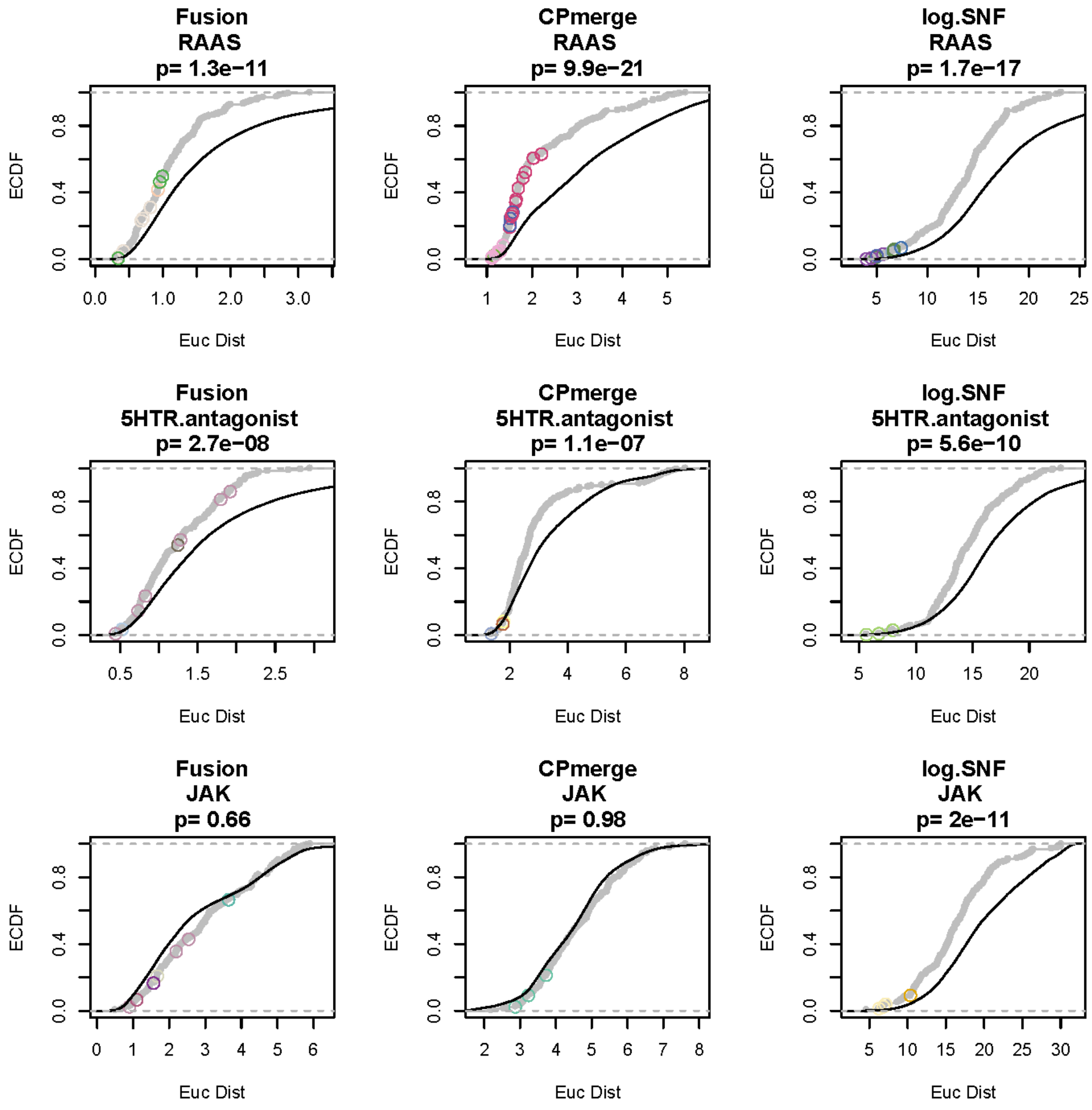
Fusion
RAAS
p= 1.3e−11
CPmerge
RAAS
p= 9.9e−21
log.SNF
RAAS
p= 1.7e−17
Fusion
5HTR.antagonist
p= 2.7e−08
CPmerge
5HTR.antagonist
p= 1.1e−07
log.SNF
5HTR.antagonist
p= 5.6e−10
Fusion
JAK
p= 0.66
CPmerge
JAK
p= 0.98
log.SNF
JAK
p= 2e−11
ECDF
Euc Dist

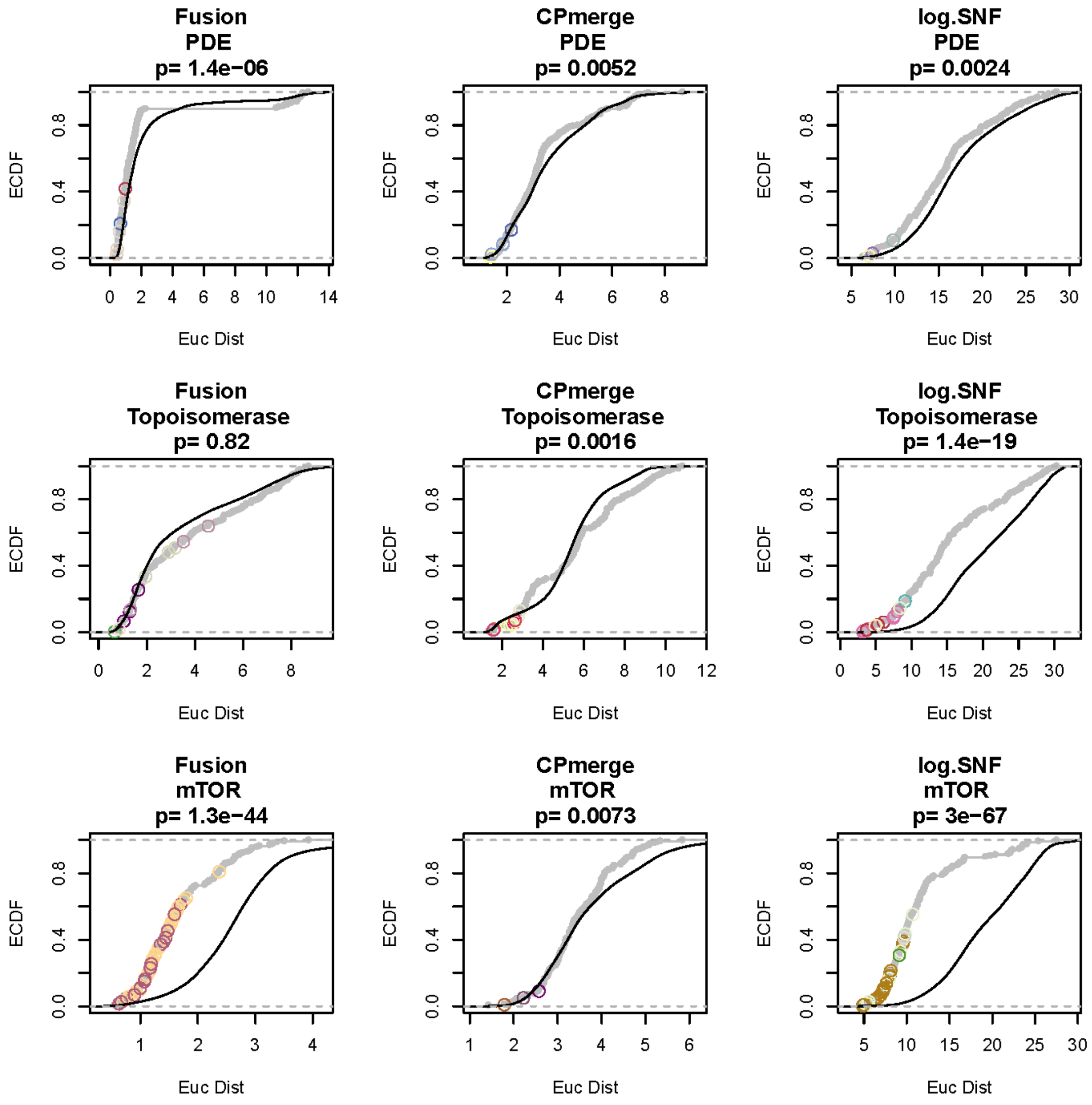
Fusion
PDE
p= 1.4e−06
CPmerge
PDE
p= 0.0052
log.SNF
PDE
p= 0.0024
Fusion
Topoisomerase
p= 0.82
CPmerge
Topoisomerase
p= 0.0016
log.SNF
Topoisomerase
p= 1.4e−19
Fusion
mTOR
p= 1.3e−44
CPmerge
mTOR
p= 0.0073
log.SNF
mTOR
p= 3e−67
ECDF
Euc Dist

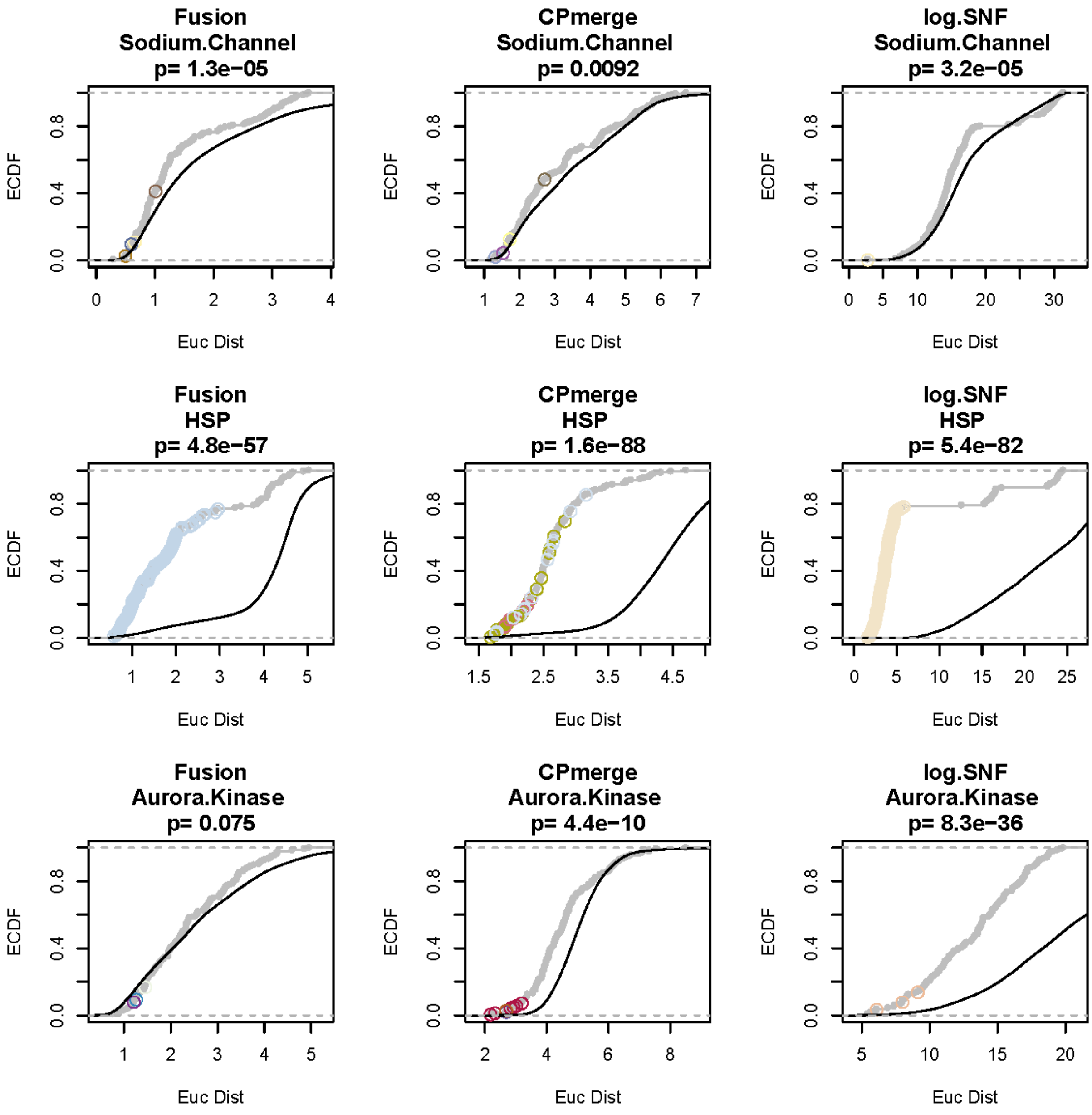
Fusion
Sodium.Channel
p= 1.3e−05
CPmerge
Sodium.Channel
p= 0.0092
log.SNF
Sodium.Channel
p= 3.2e−05
Fusion
HSP
p= 4.8e−57
CPmerge
HSP
p= 1.6e−88
log.SNF
HSP
p= 5.4e−82
Fusion
Aurora.Kinase
p= 0.075
CPmerge
Aurora.Kinase
p= 4.4e−10
log.SNF
Aurora.Kinase
p= 8.3e−36
ECDF
Euc Dist

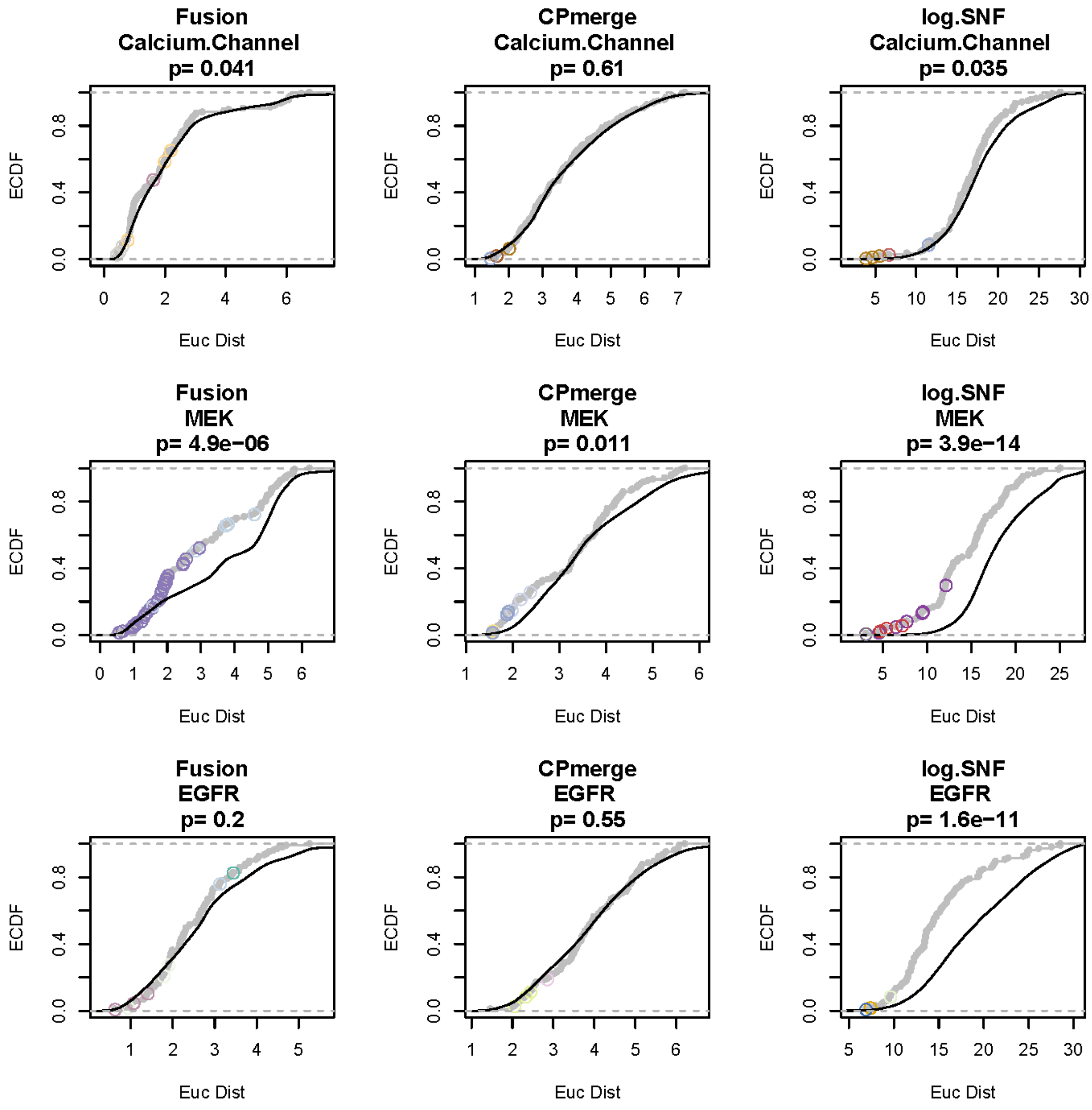
Fusion
Calcium.Channel
p= 0.041
CPmerge
Calcium.Channel
p= 0.61
log.SNF
Calcium.Channel
p= 0.035
Fusion
MEK
p= 4.9e−06
CPmerge
MEK
p= 0.011
log.SNF
MEK
p= 3.9e−14
Fusion
EGFR
p= 0.2
CPmerge
EGFR
p= 0.55
log.SNF
EGFR
p= 1.6e−11
ECDF
Euc Dist

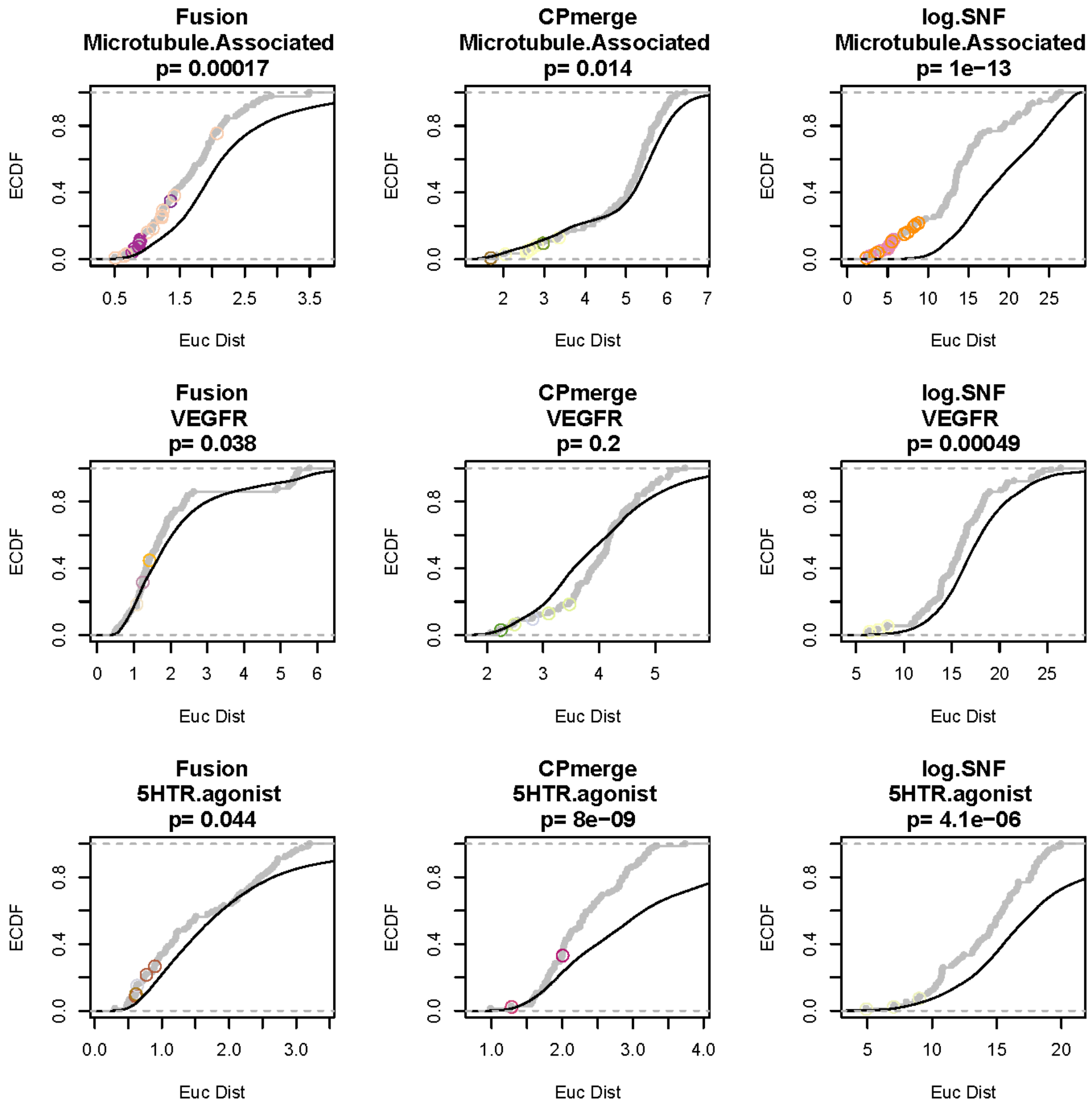

Fusion
Microtubule.Associated
p= 0.00017
CPmerge
Microtubule.Associated
p= 0.014
log.SNF
Microtubule.Associated
p= 1e−13
Fusion
VEGFR
p= 0.038
CPmerge
VEGFR
p= 0.2
log.SNF
VEGFR
p= 0.00049
Fusion
5HTR.agonist
p= 0.044
CPmerge
5HTR.agonist
p= 8e−09
log.SNF
5HTR.agonist
p= 4.1e−06
ECDF
Euc Dist

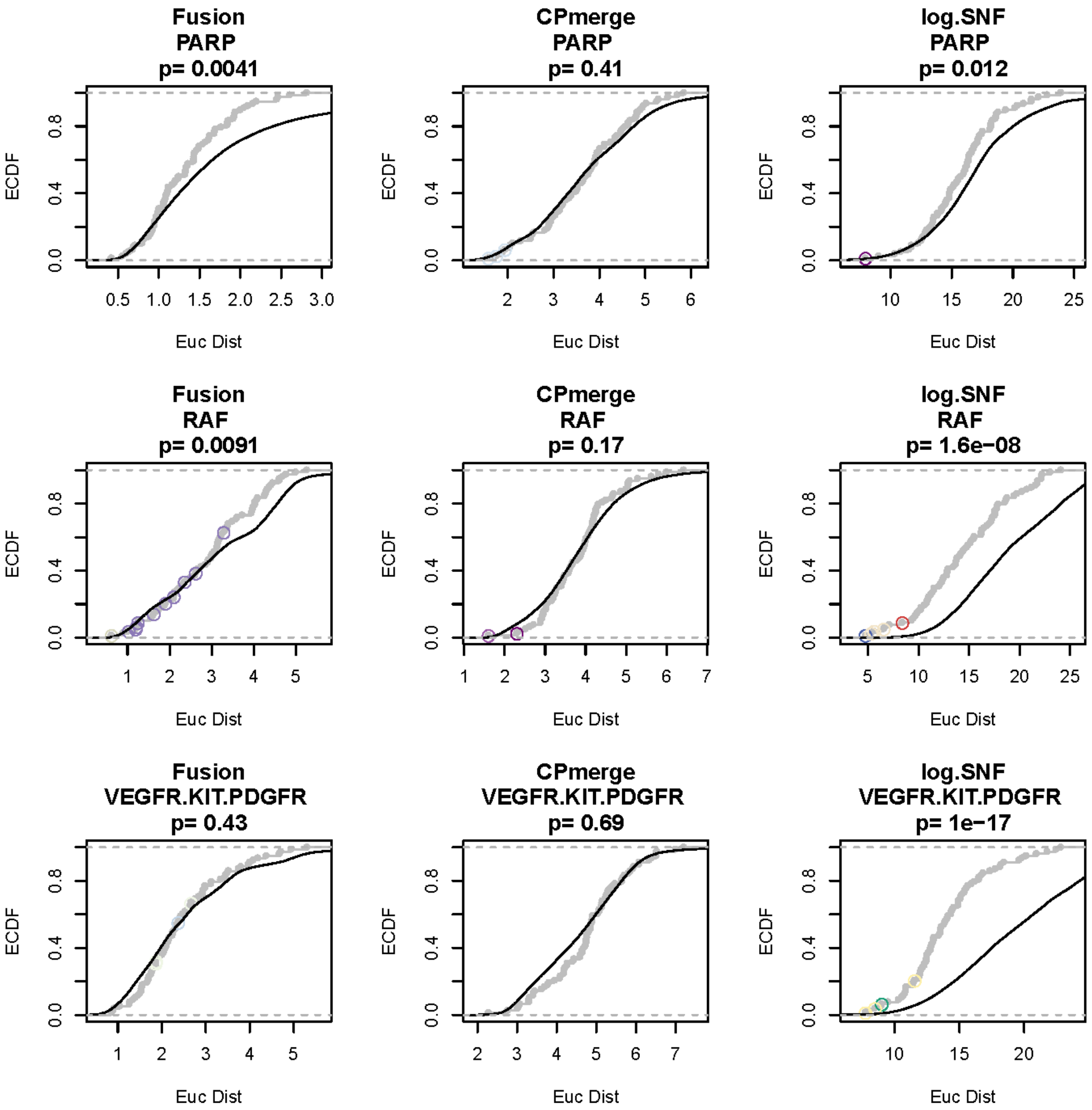
Fusion
PARP
p= 0.0041
CPmerge
PARP
p= 0.41
log.SNF
PARP
p= 0.012
Fusion
RAF
p= 0.0091
CPmerge
RAF
p= 0.17
log.SNF
RAF
p= 1.6e−08
Fusion
VEGFR.KIT.PDGFR
p= 0.43
CPmerge
VEGFR.KIT.PDGFR
p= 0.69
log.SNF
VEGFR.KIT.PDGFR
p= 1e−17
ECDF
Euc Dist

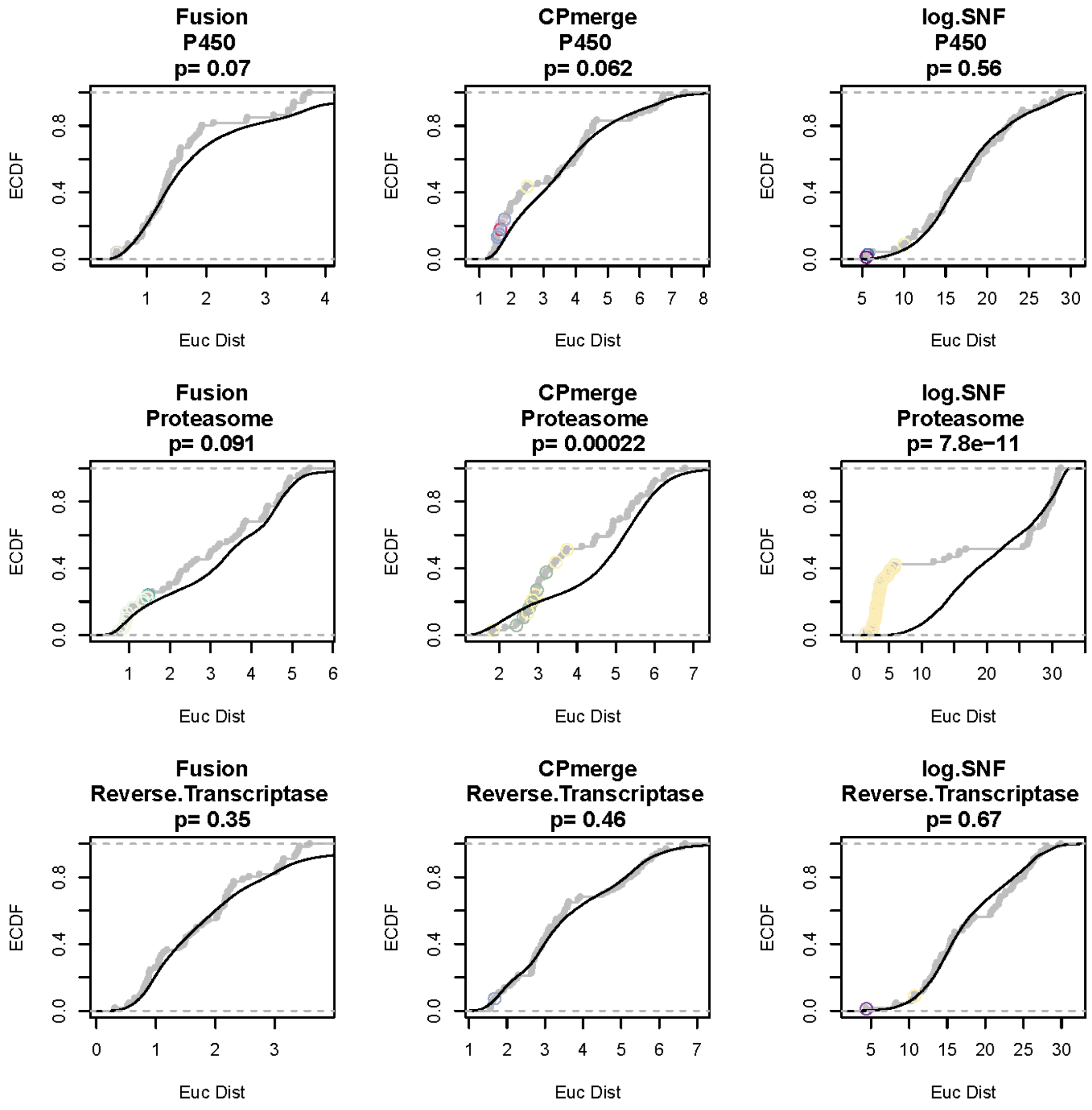
Fusion
P450
p= 0.07
CPmerge
P450
p= 0.062
log.SNF
P450
p= 0.56
Fusion
Proteasome
p= 0.091
CPmerge
Proteasome
p= 0.00022
log.SNF
Proteasome
p= 7.8e−11
Fusion
Reverse.Transcriptase
p= 0.35
CPmerge
Reverse.Transcriptase
p= 0.46
log.SNF
Reverse.Transcriptase
p= 0.67
ECDF
Euc Dist

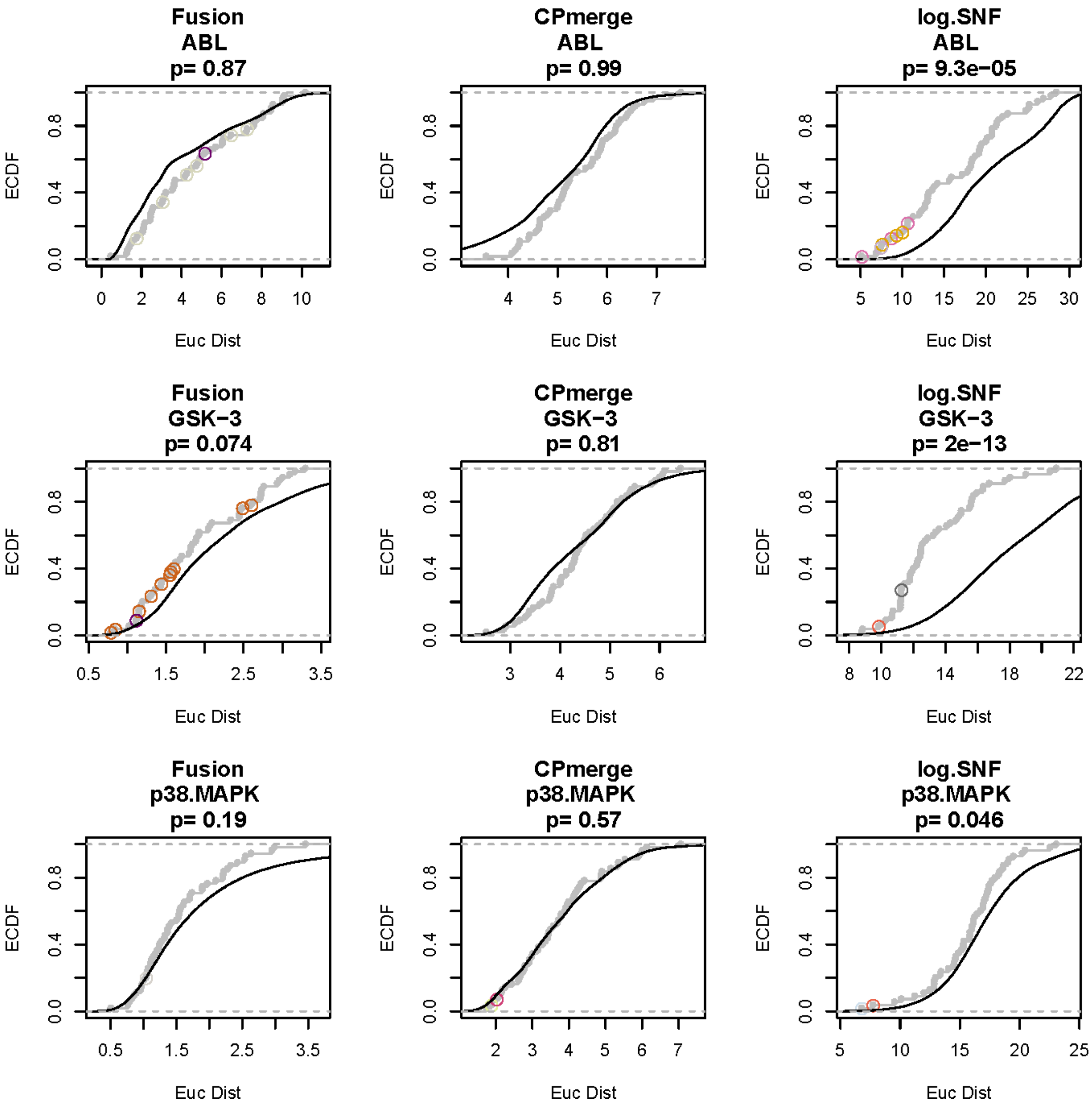
Fusion
ABL
p= 0.87
CPmerge
ABL
p= 0.99
log.SNF
ABL
p= 9.3e−05
Fusion
GSK−3
p= 0.074
CPmerge
GSK−3
p= 0.81
log.SNF
GSK−3
p= 2e−13
Fusion
p38.MAPK
p= 0.19
CPmerge
p38.MAPK
p= 0.57
log.SNF
p38.MAPK
p= 0.046
ECDF
Euc Dist

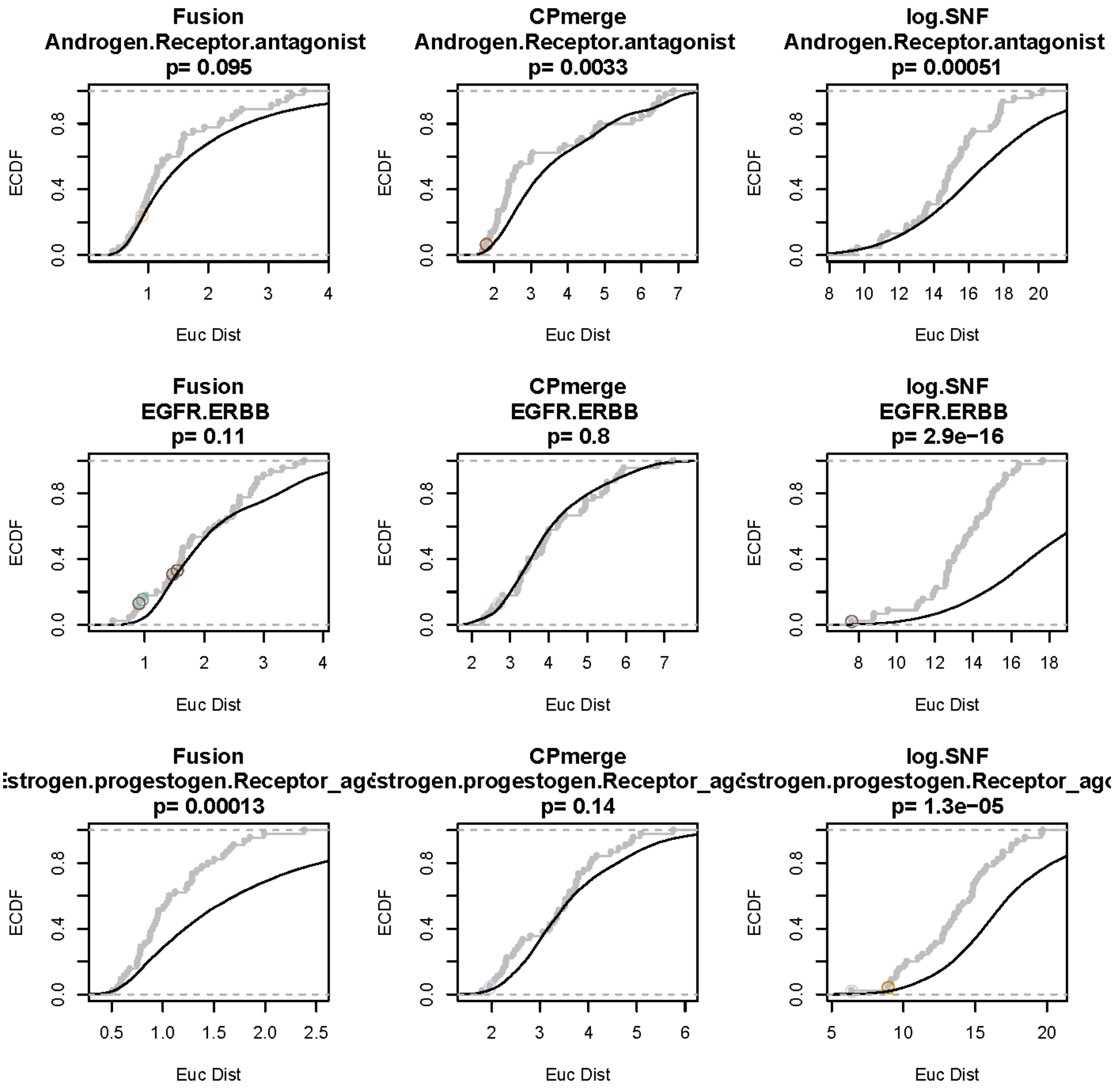

Fusion
Androgen.Receptor.antagonist
p= 0.095
CPmerge
Androgen.Receptor.antagonist
p= 0.0033
log.SNF
Androgen.Receptor.antagonist
p= 0.00051
Fusion
EGFR.ERBB
p= 0.11
CPmerge
EGFR.ERBB
p= 0.8
log.SNF
EGFR.ERBB
p= 2.9e−16
Fusion
strogen.progestogen.Receptor_ag
p= 0.00013
CPmerge
strogen.progestogen.Receptor_ag
p= 0.14
log.SNF
strogen.progestogen.Receptor_ag
p= 1.3e−05
ECDF
Euc Dist

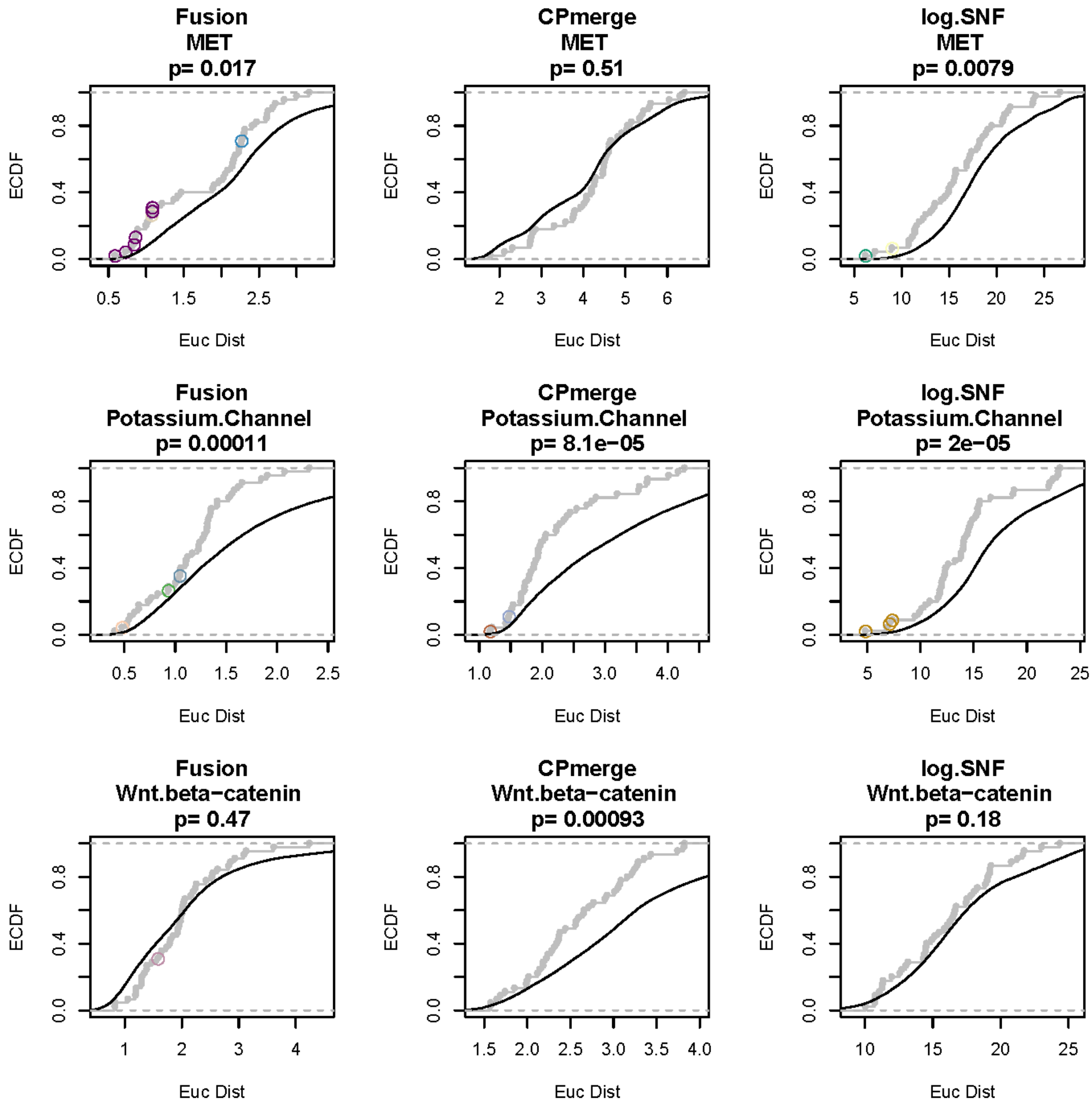
Fusion
MET
p= 0.017
CPmerge
MET
p= 0.51
log.SNF
MET
p= 0.0079
Fusion
Potassium.Channel
p= 0.00011
CPmerge
Potassium.Channel
p= 8.1e−05
log.SNF
Potassium.Channel
p= 2e−05
Fusion
Wnt.beta−catenin
p= 0.47
CPmerge
Wnt.beta−catenin
p= 0.00093
log.SNF
Wnt.beta−catenin
p= 0.18
ECDF
Euc Dist

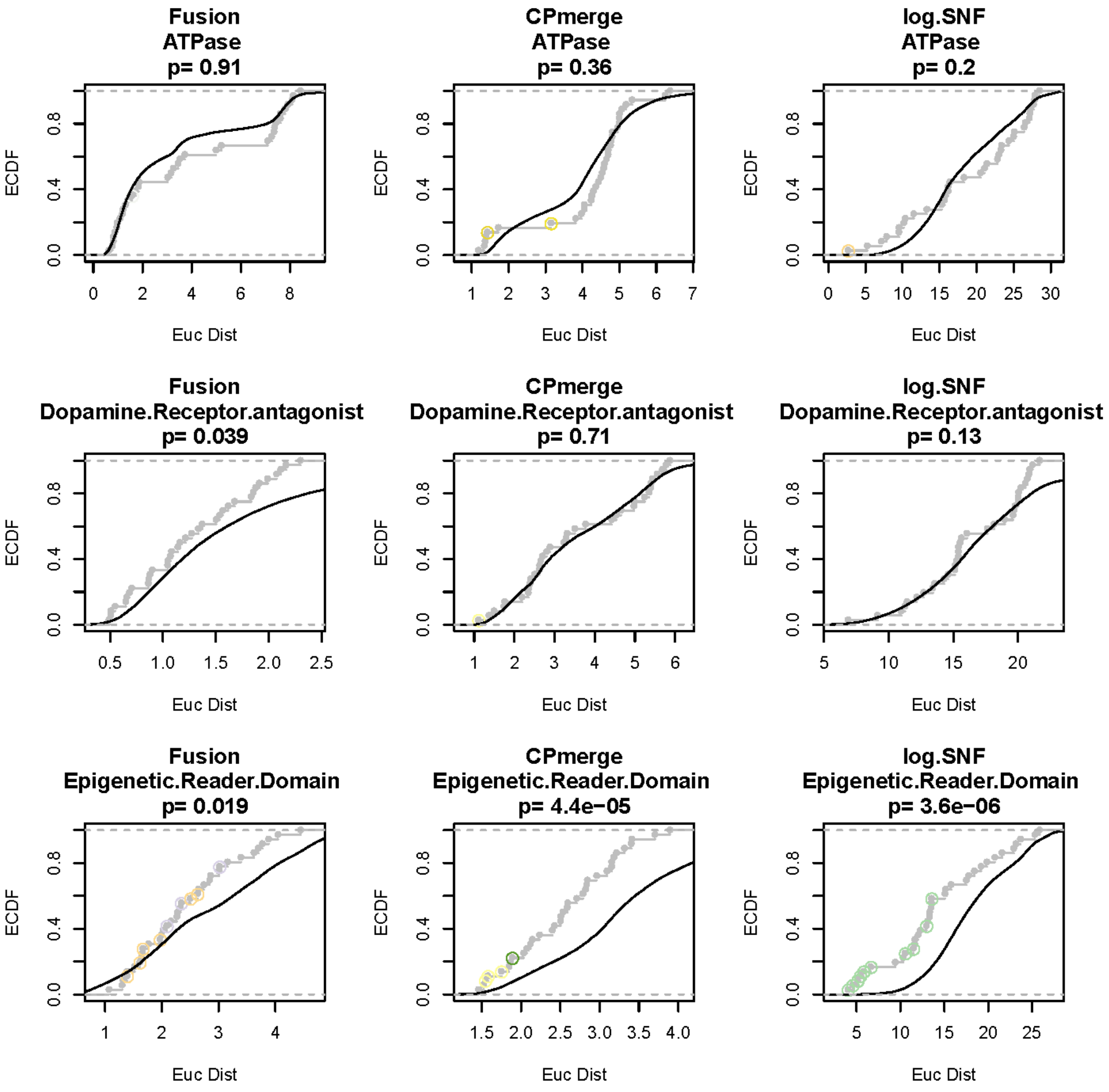
Fusion
ATPase
p= 0.91
CPmerge
ATPase
p= 0.36
log.SNF
ATPase
p= 0.2
Fusion
Dopamine.Receptor.antagonist
p= 0.039
CPmerge
Dopamine.Receptor.antagonist
p= 0.71
log.SNF
Dopamine.Receptor.antagonist
p= 0.13
Fusion
Epigenetic.Reader.Domain
p= 0.019
CPmerge
Epigenetic.Reader.Domain
p= 4.4e−05
log.SNF
Epigenetic.Reader.Domain
p= 3.6e−06
ECDF
Euc Dist

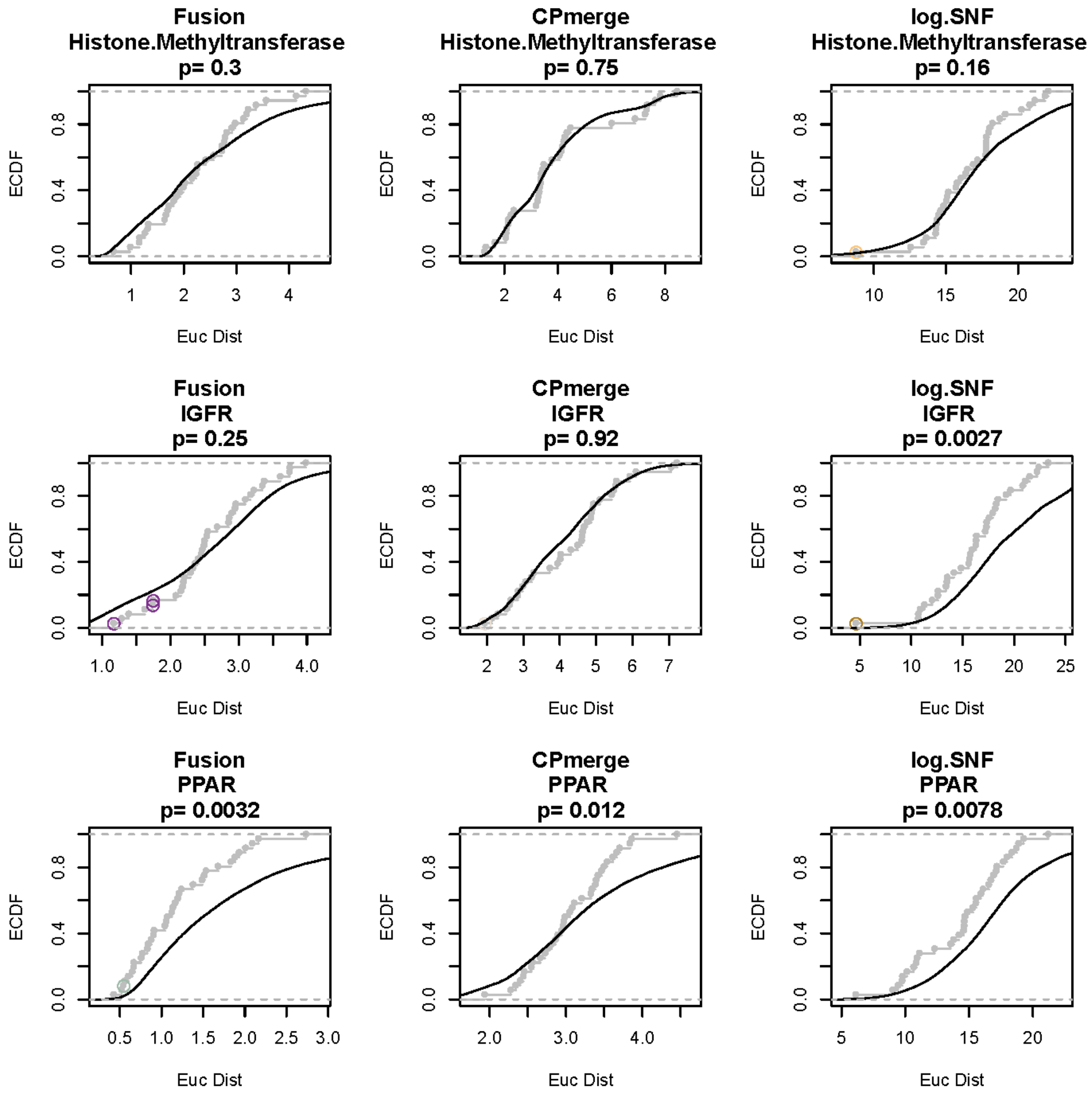
Fusion
Histone.Methyltransferase
p= 0.3
CPmerge
Histone.Methyltransferase
p= 0.75
log.SNF
Histone.Methyltransferase
p= 0.16
Fusion
IGFR
p= 0.25
CPmerge
IGFR
p= 0.92
log.SNF
IGFR
p= 0.0027
Fusion
PPAR
p= 0.0032
CPmerge
PPAR
p= 0.012
log.SNF
PPAR
p= 0.0078
ECDF
Euc Dist

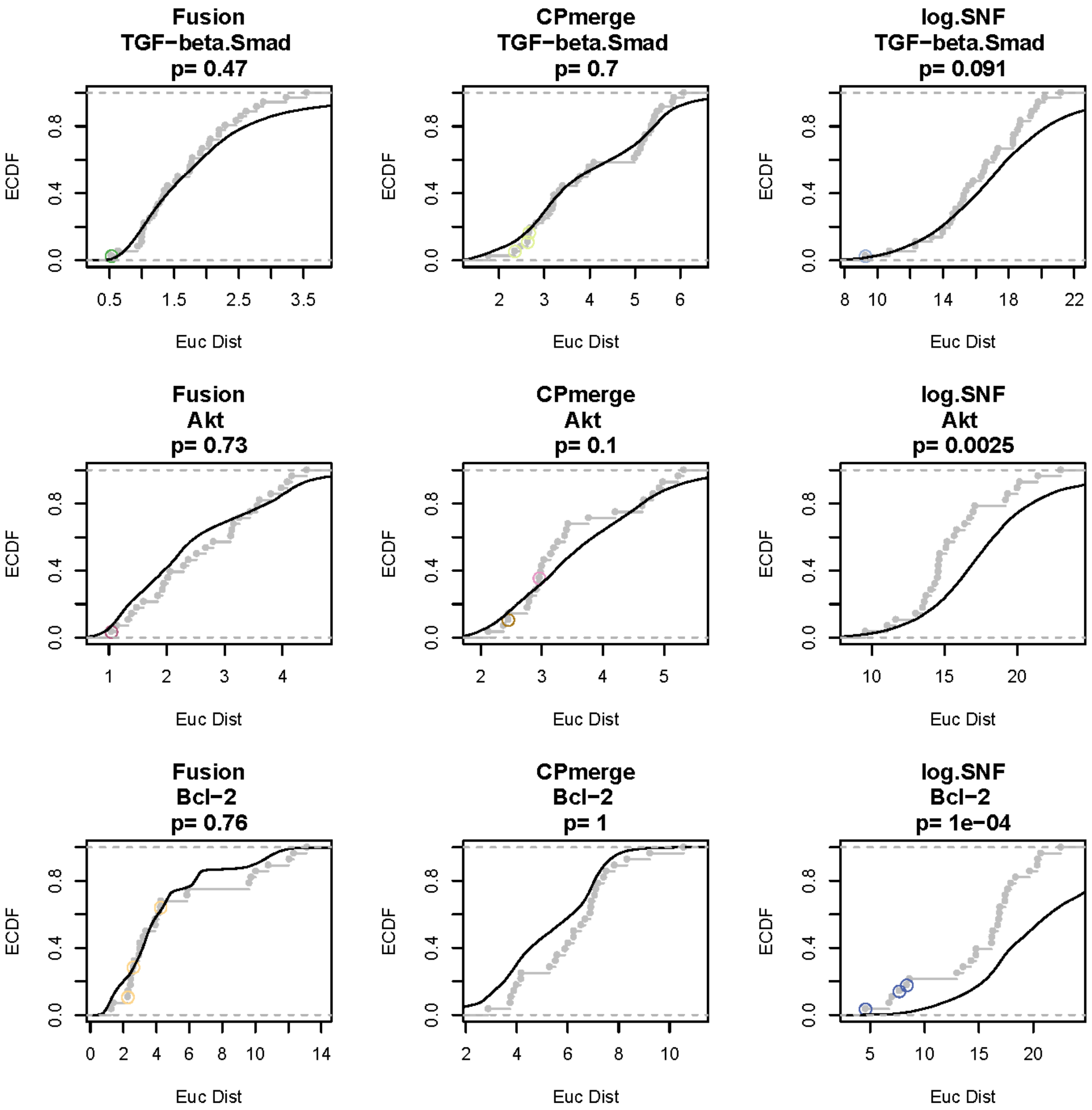
Fusion
TGF−beta.Smad
p= 0.47
CPmerge
TGF−beta.Smad
p= 0.7
log.SNF
TGF−beta.Smad
p= 0.091
Fusion
Akt
p= 0.73
CPmerge
Akt
p= 0.1
log.SNF
Akt
p= 0.0025
Fusion
Bcl−2
p= 0.76
CPmerge
Bcl−2
p= 1
log.SNF
Bcl−2
p= 1e−04
ECDF
Euc Dist

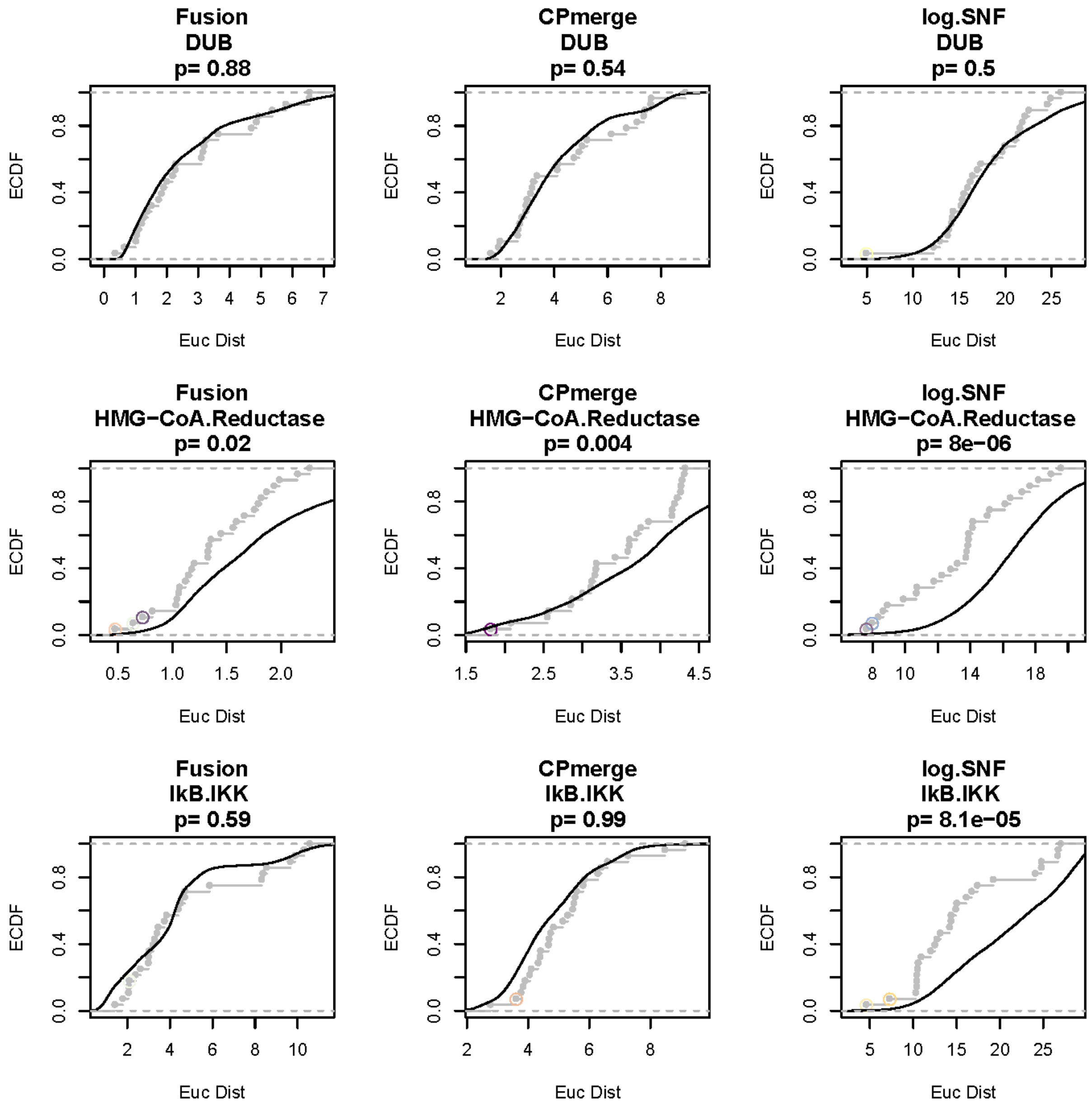
Fusion
DUB
p= 0.88
CPmerge
DUB
p= 0.54
log.SNF
DUB
p= 0.5
Fusion
HMG−CoA.Reductase
p= 0.02
CPmerge
HMG−CoA.Reductase
p= 0.004
log.SNF
HMG−CoA.Reductase
p= 8e−06
Fusion
IkB.IKK
p= 0.59
CPmerge
IkB.IKK
p= 0.99
log.SNF
IkB.IKK
p= 8.1e−05
ECDF
Euc Dist

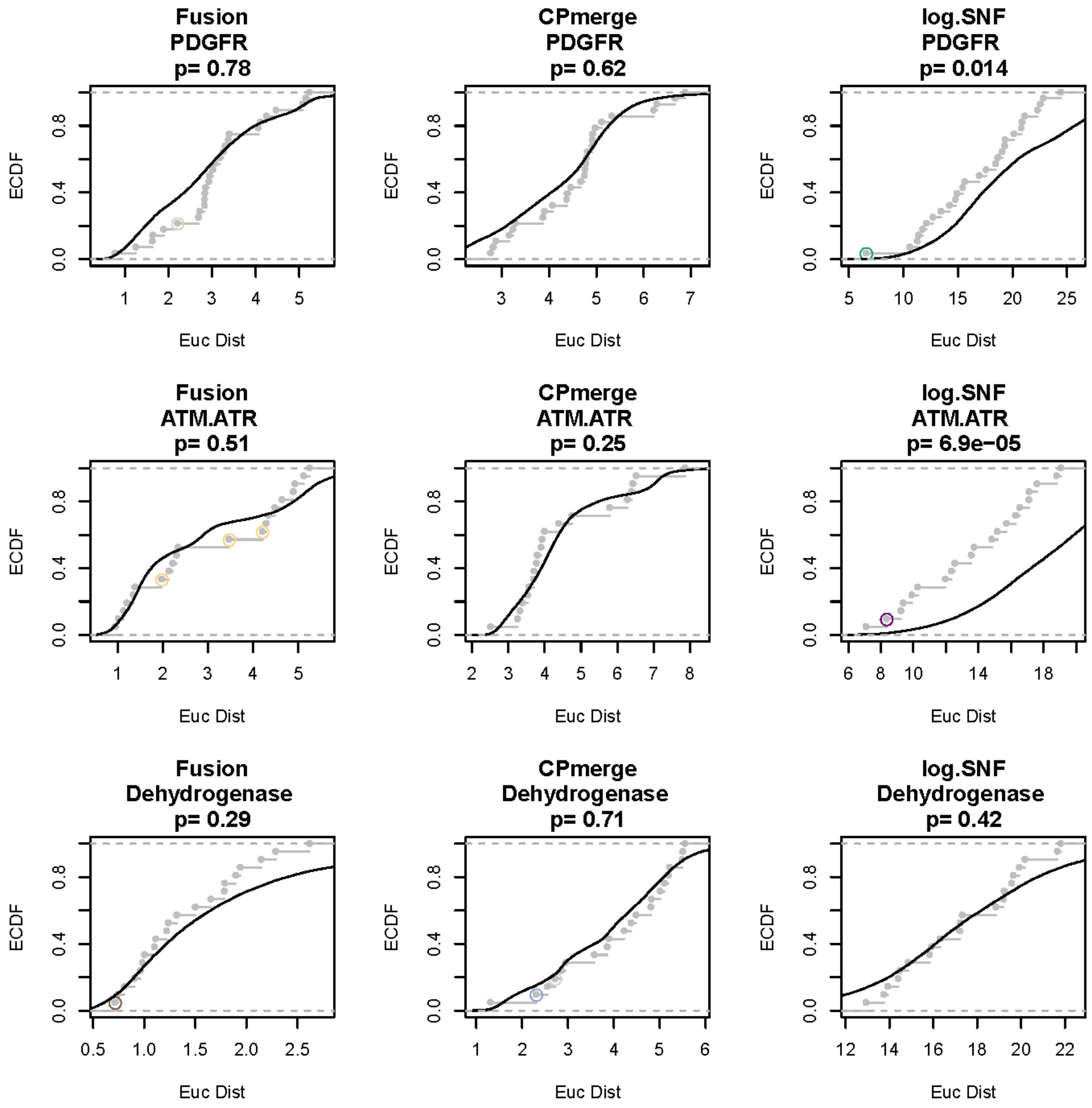
Fusion
PDGFR
p= 0.78
CPmerge
PDGFR
p= 0.62
log.SNF
PDGFR
p= 0.014
Fusion
ATM.ATR
p= 0.51
CPmerge
ATM.ATR
p= 0.25
log.SNF
ATM.ATR
p= 6.9e−05
Fusion
Dehydrogenase
p= 0.29
CPmerge
Dehydrogenase
p= 0.71
log.SNF
Dehydrogenase
p= 0.42
ECDF
Euc Dist

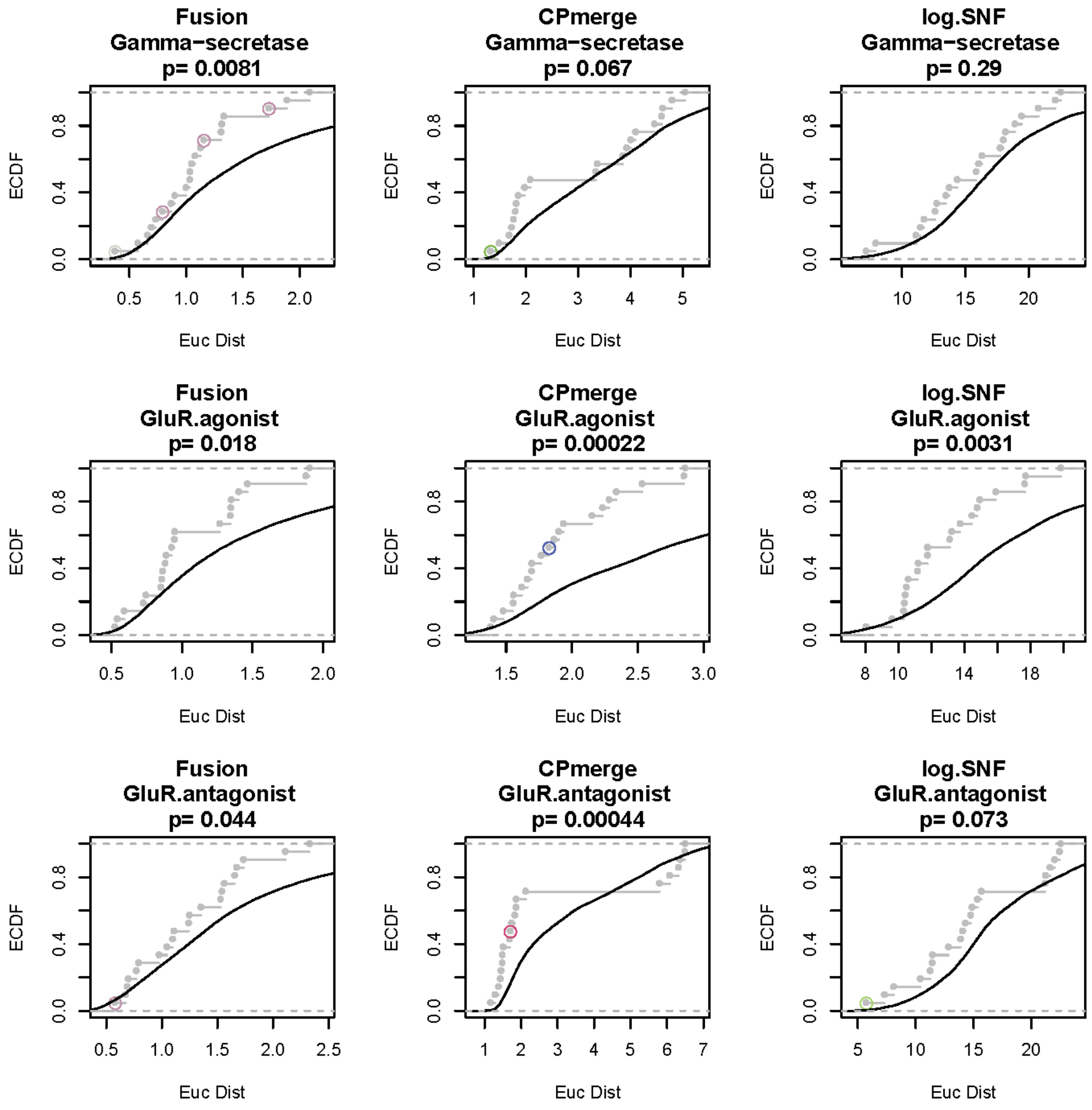
Fusion
Gamma−secretase
p= 0.0081
CPmerge
Gamma−secretase
p= 0.067
log.SNF
Gamma−secretase
p= 0.29
Fusion
GluR.agonist
p= 0.018
CPmerge
GluR.agonist
p= 0.00022
log.SNF
GluR.agonist
p= 0.0031
Fusion
GluR.antagonist
p= 0.044
CPmerge
GluR.antagonist
p= 0.00044
log.SNF
GluR.antagonist
p= 0.073
ECDF
Euc Dist

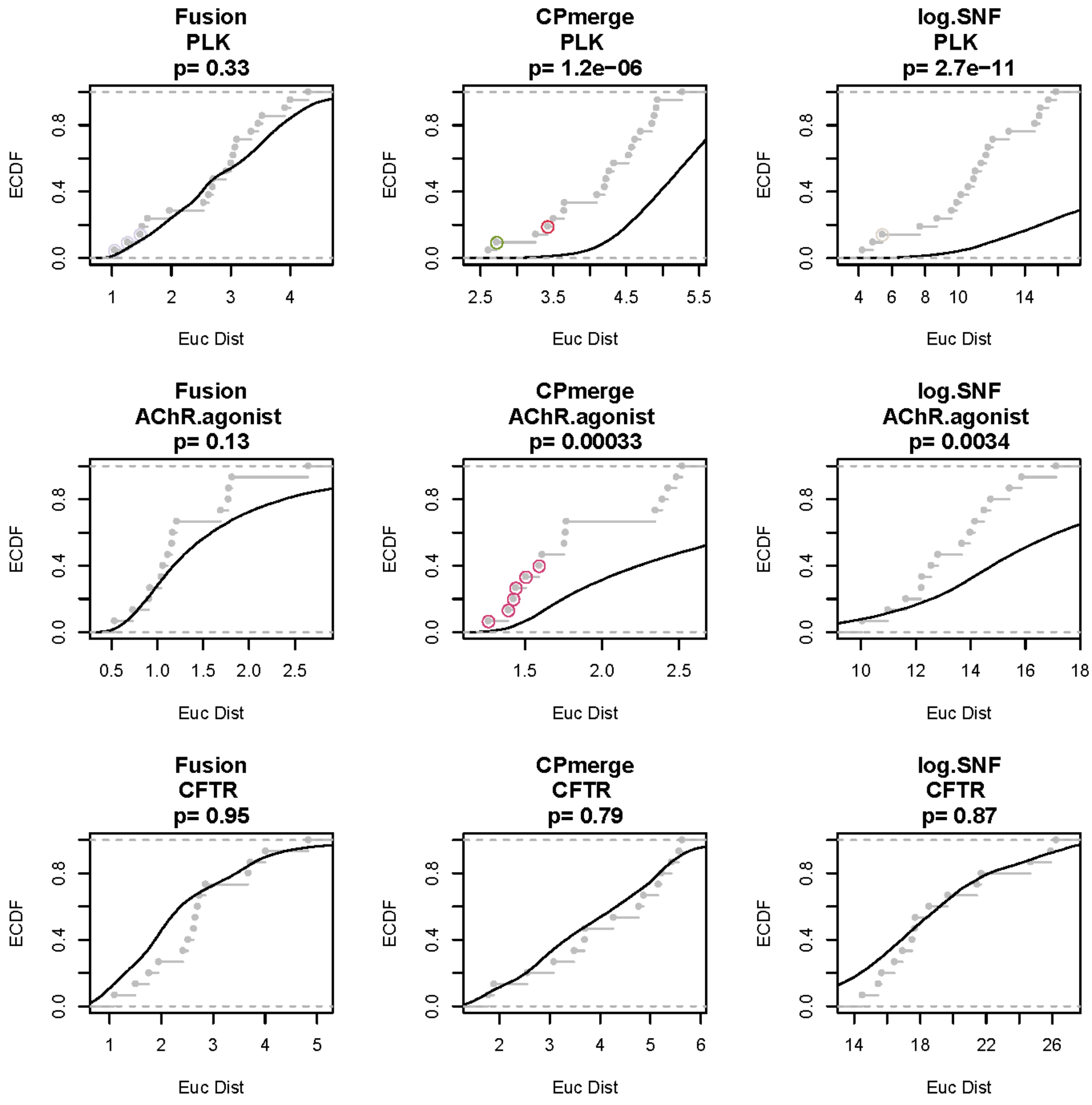
Fusion
PLK
p= 0.33
CPmerge
PLK
p= 1.2e−06
log.SNF
PLK
p= 2.7e−11
Fusion
AChR.agonist
p= 0.13
CPmerge
AChR.agonist
p= 0.00033
log.SNF
AChR.agonist
p= 0.0034
Fusion
CFTR
p= 0.95
CPmerge
CFTR
p= 0.79
log.SNF
CFTR
p= 0.87
ECDF
Euc Dist

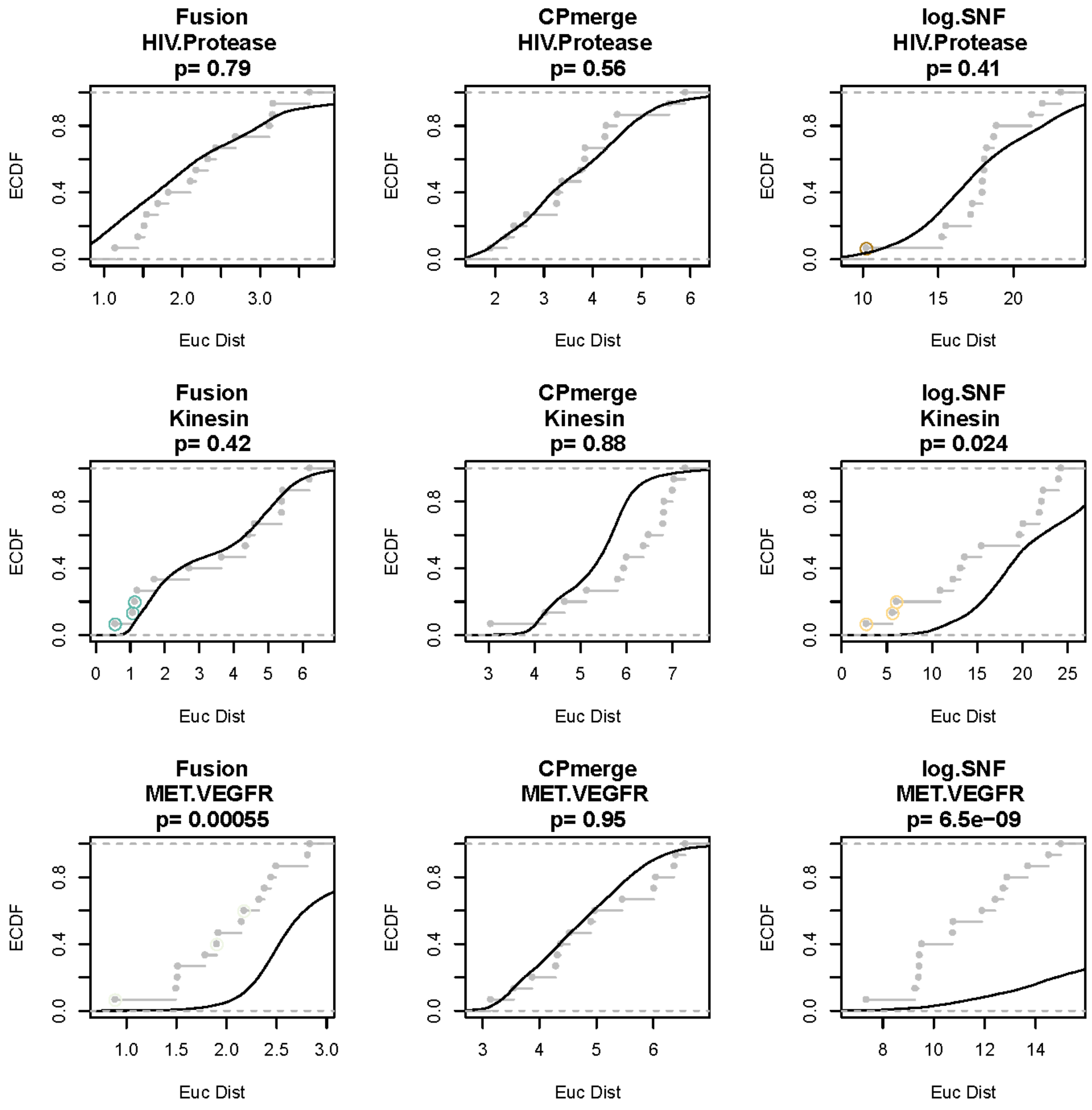
Fusion
HIV.Protease
p= 0.79
CPmerge
HIV.Protease
p= 0.56
log.SNF
HIV.Protease
p= 0.41
Fusion
Kinesin
p= 0.42
CPmerge
Kinesin
p= 0.88
log.SNF
Kinesin
p= 0.024
Fusion
MET.VEGFR
p= 0.00055
CPmerge
MET.VEGFR
p= 0.95
log.SNF
MET.VEGFR
p= 6.5e−09
ECDF
Euc Dist

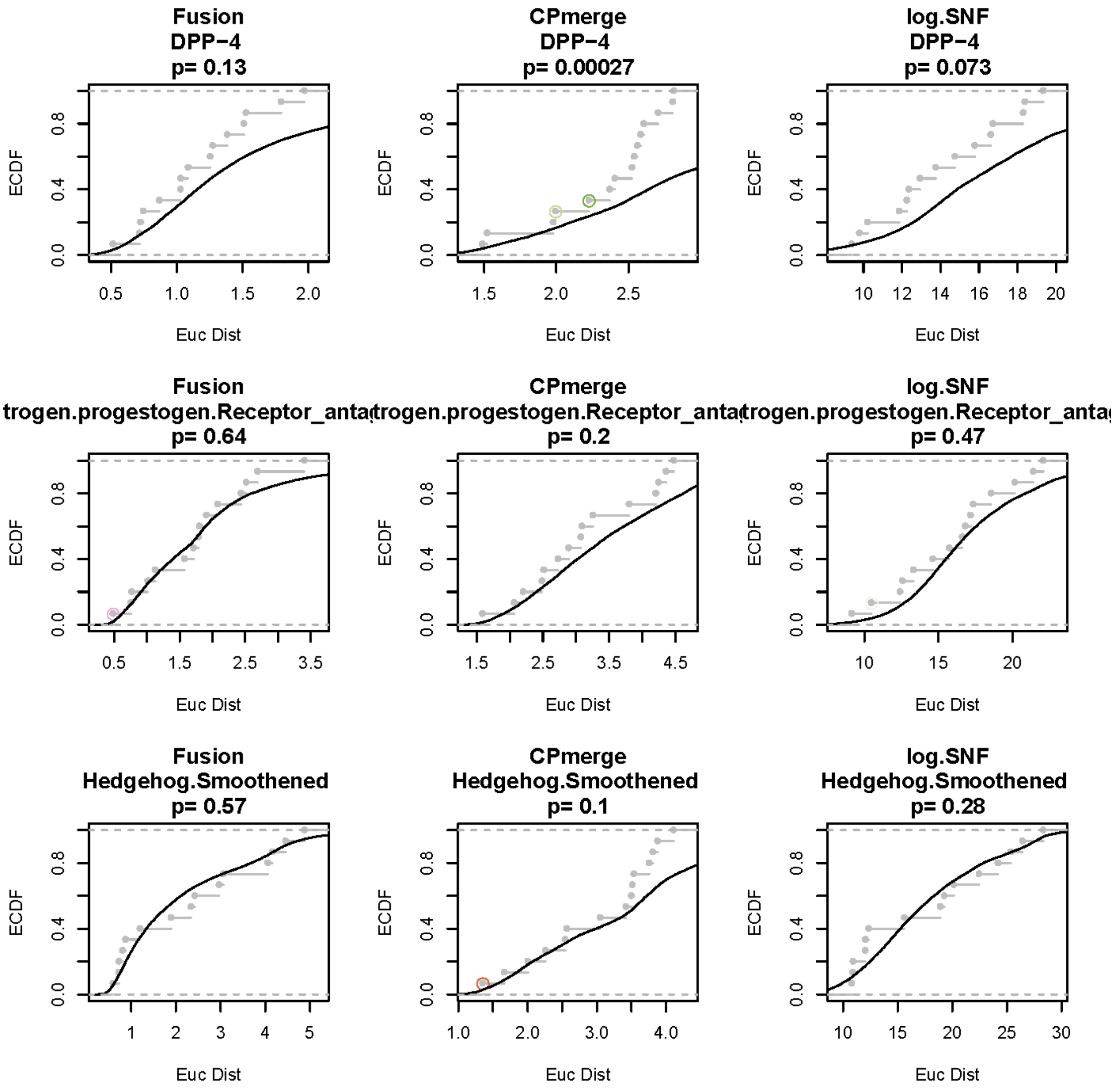
Fusion
DPP−4
p= 0.13
CPmerge
DPP−4
p= 0.00027
log.SNF
DPP−4
p= 0.073
Fusion
trogen.progestogen.Receptor_anta
p= 0.64
CPmerge
trogen.progestogen.Receptor_anta
p= 0.2
log.SNF
trogen.progestogen.Receptor_anta
p= 0.47
Fusion
Hedgehog.Smoothened
p= 0.57
CPmerge
Hedgehog.Smoothened
p= 0.1
log.SNF
Hedgehog.Smoothened
p= 0.28
ECDF
Euc Dist

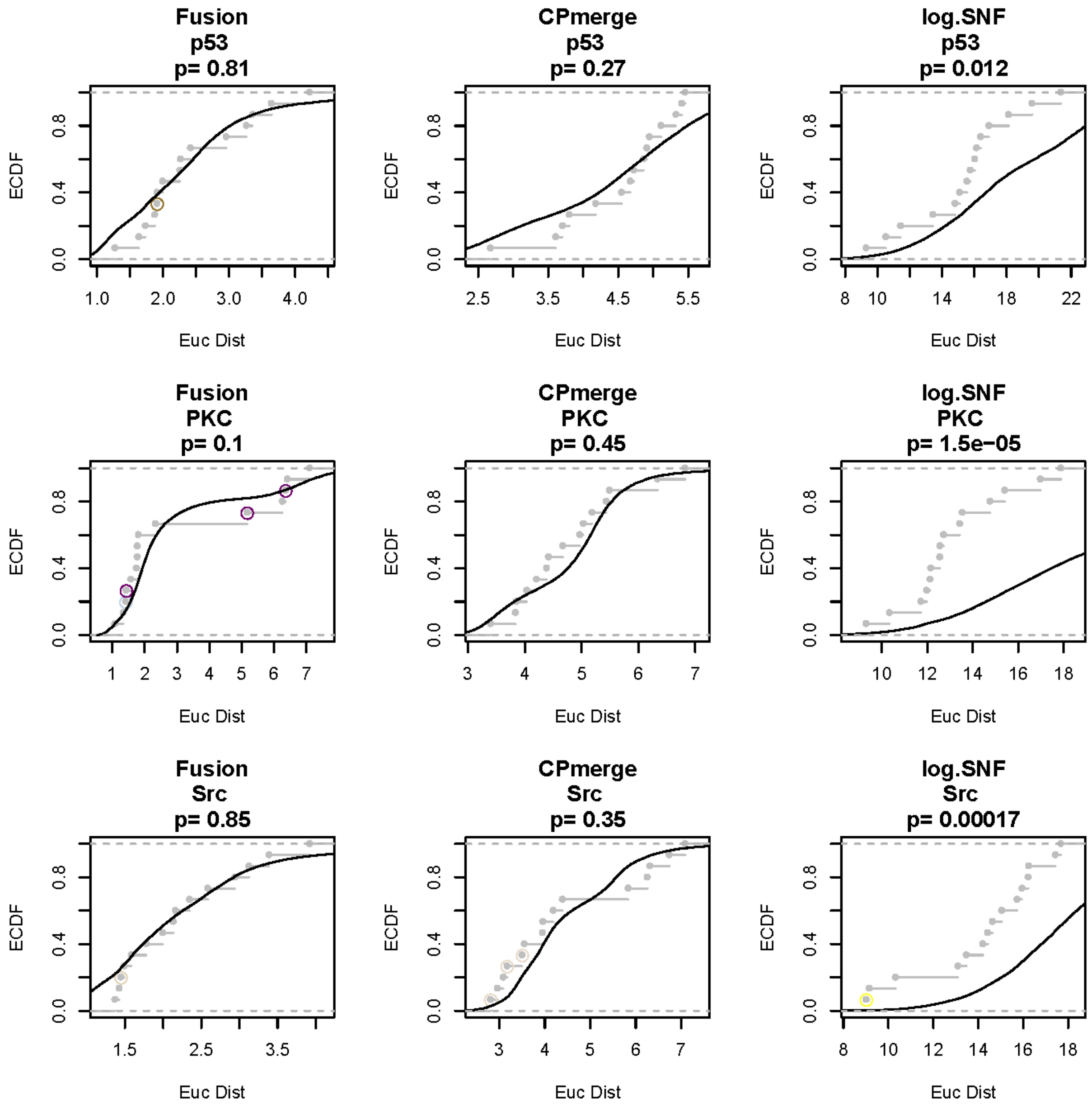
Fusion
p53
p= 0.81
CPmerge
p53
p= 0.27
log.SNF
p53
p= 0.012
Fusion
PKC
p= 0.1
CPmerge
PKC
p= 0.45
log.SNF
PKC
p= 1.5e−05
Fusion
Src
p= 0.85
CPmerge
Src
p= 0.35
log.SNF
Src
p= 0.00017
ECDF
Euc Dist

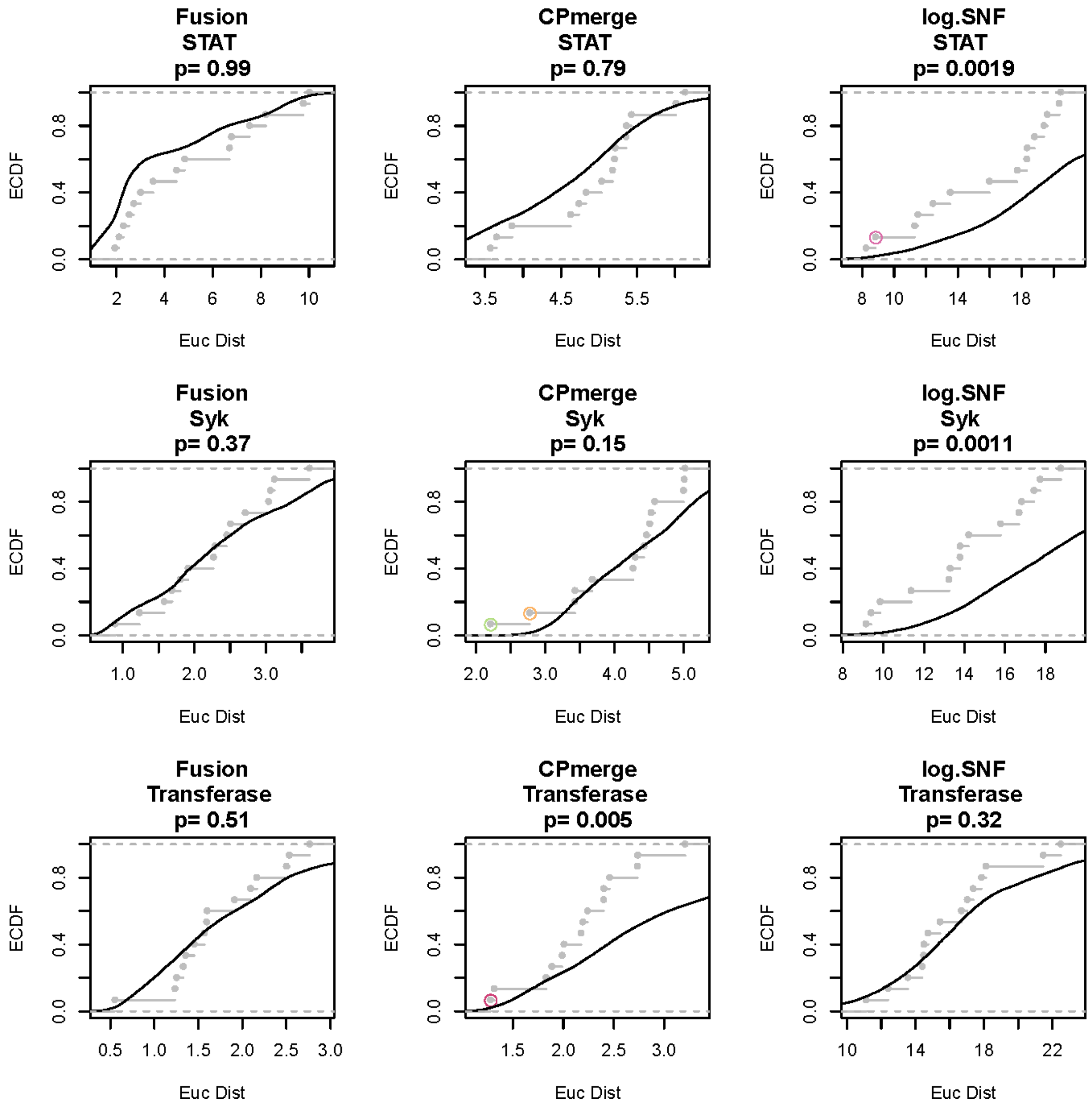
Fusion
STAT
p= 0.99
CPmerge
STAT
p= 0.79
log.SNF
STAT
p= 0.0019
Fusion
Syk
p= 0.37
CPmerge
Syk
p= 0.15
log.SNF
Syk
p= 0.0011
Fusion
Transferase
p= 0.51
CPmerge
Transferase
p= 0.005
log.SNF
Transferase
p= 0.32
ECDF
Euc Dist

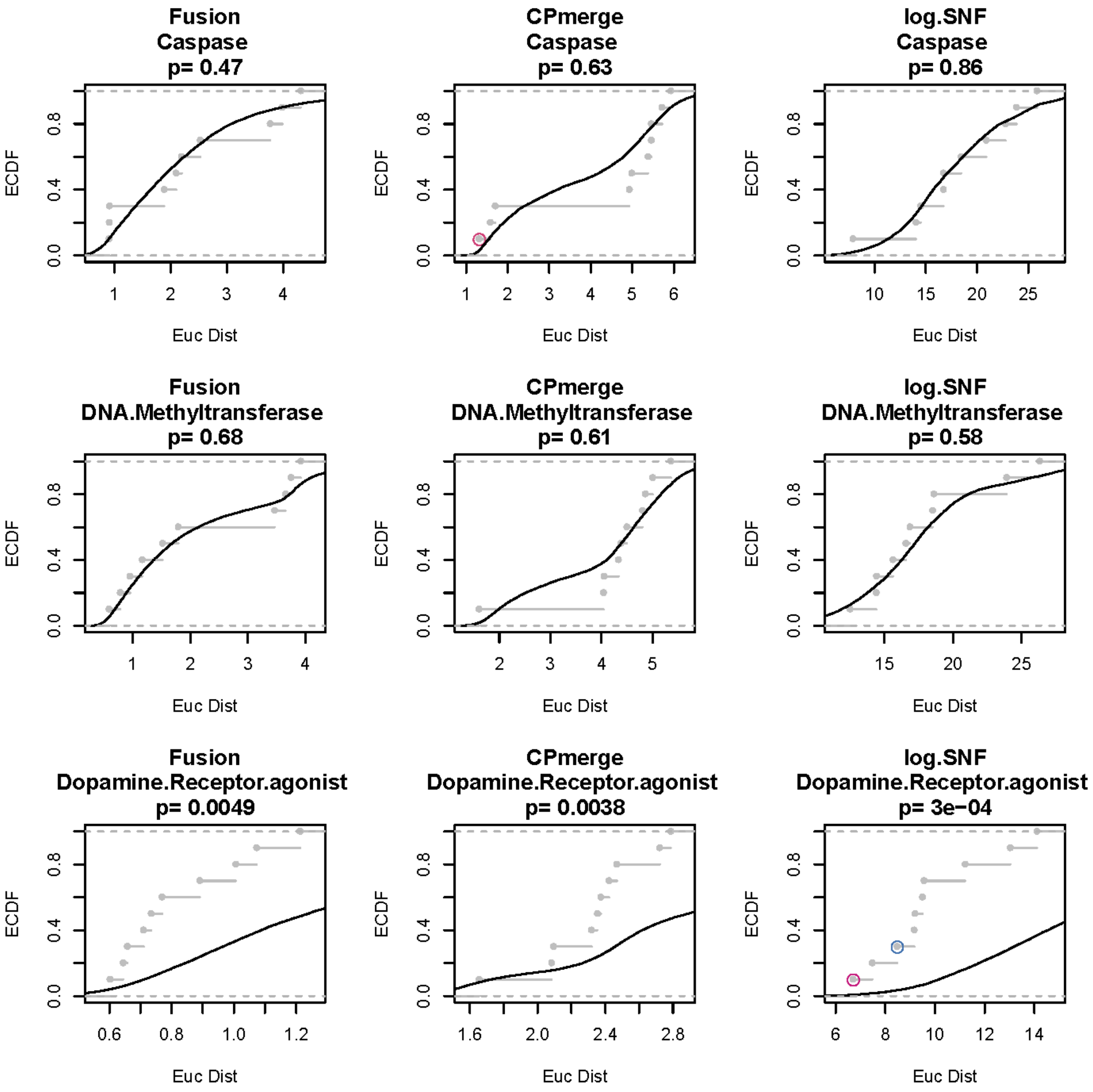
Fusion
Caspase
p= 0.47
CPmerge
Caspase
p= 0.63
log.SNF
Caspase
p= 0.86
Fusion
DNA.Methyltransferase
p= 0.68
CPmerge
DNA.Methyltransferase
p= 0.61
log.SNF
DNA.Methyltransferase
p= 0.58
Fusion
Dopamine.Receptor.agonist
p= 0.0049
CPmerge
Dopamine.Receptor.agonist
p= 0.0038
log.SNF
Dopamine.Receptor.agonist
p= 3e−04
ECDF
Euc Dist

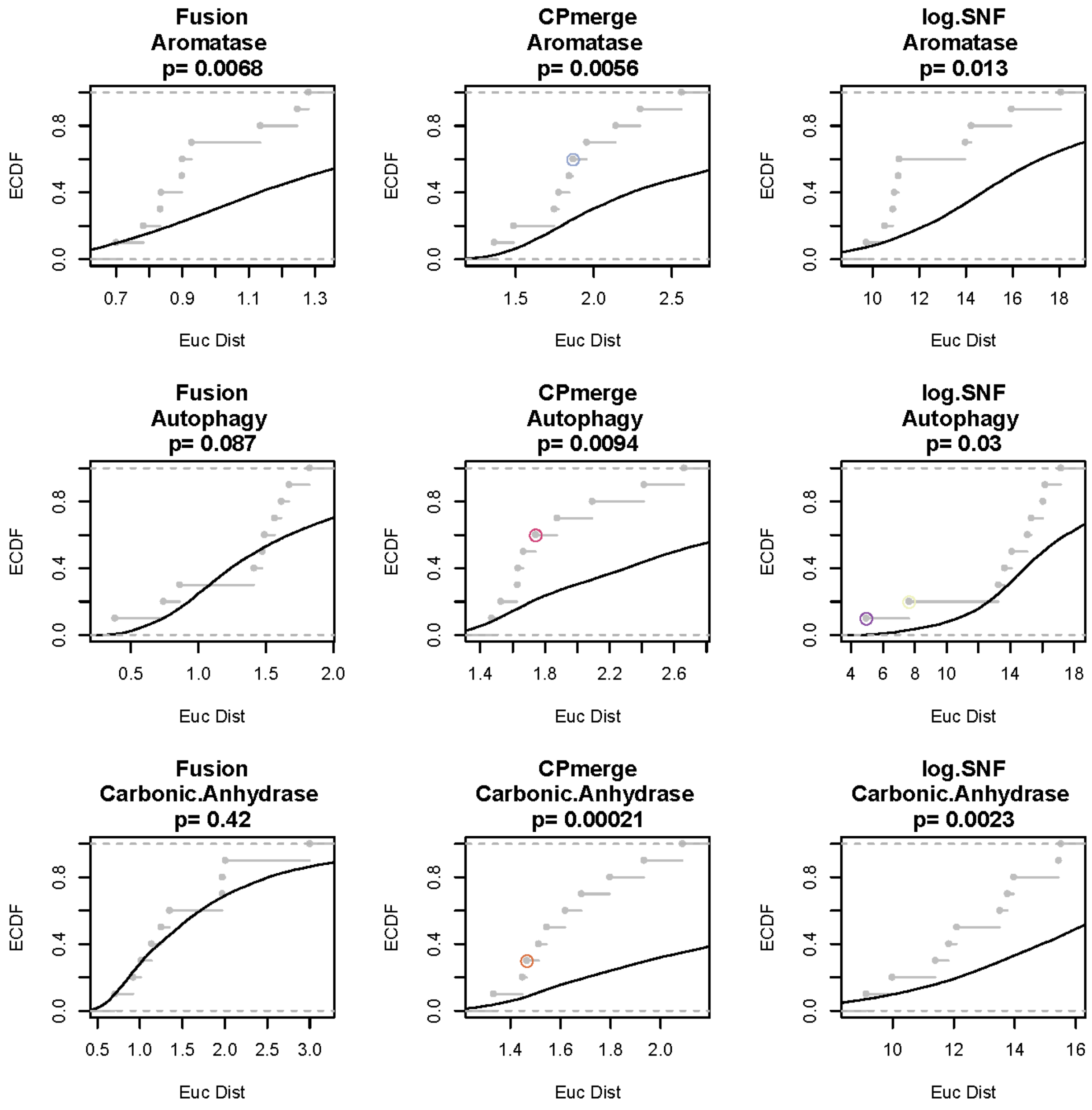
Fusion
Aromatase
p= 0.0068
CPmerge
Aromatase
p= 0.0056
log.SNF
Aromatase
p= 0.013
Fusion
Autophagy
p= 0.087
CPmerge
Autophagy
p= 0.0094
log.SNF
Autophagy
p= 0.03
Fusion
Carbonic.Anhydrase
p= 0.42
CPmerge
Carbonic.Anhydrase
p= 0.00021
log.SNF
Carbonic.Anhydrase
p= 0.0023
ECDF
Euc Dist
0.0
0.4
0.8
0.7
0.9
1.1
1.3
1.5
2.0
2.5
10
12
14
16
18
0.5
1.0
1.4
1.8
2.2
2.6
4
6
8
3.0
1.6

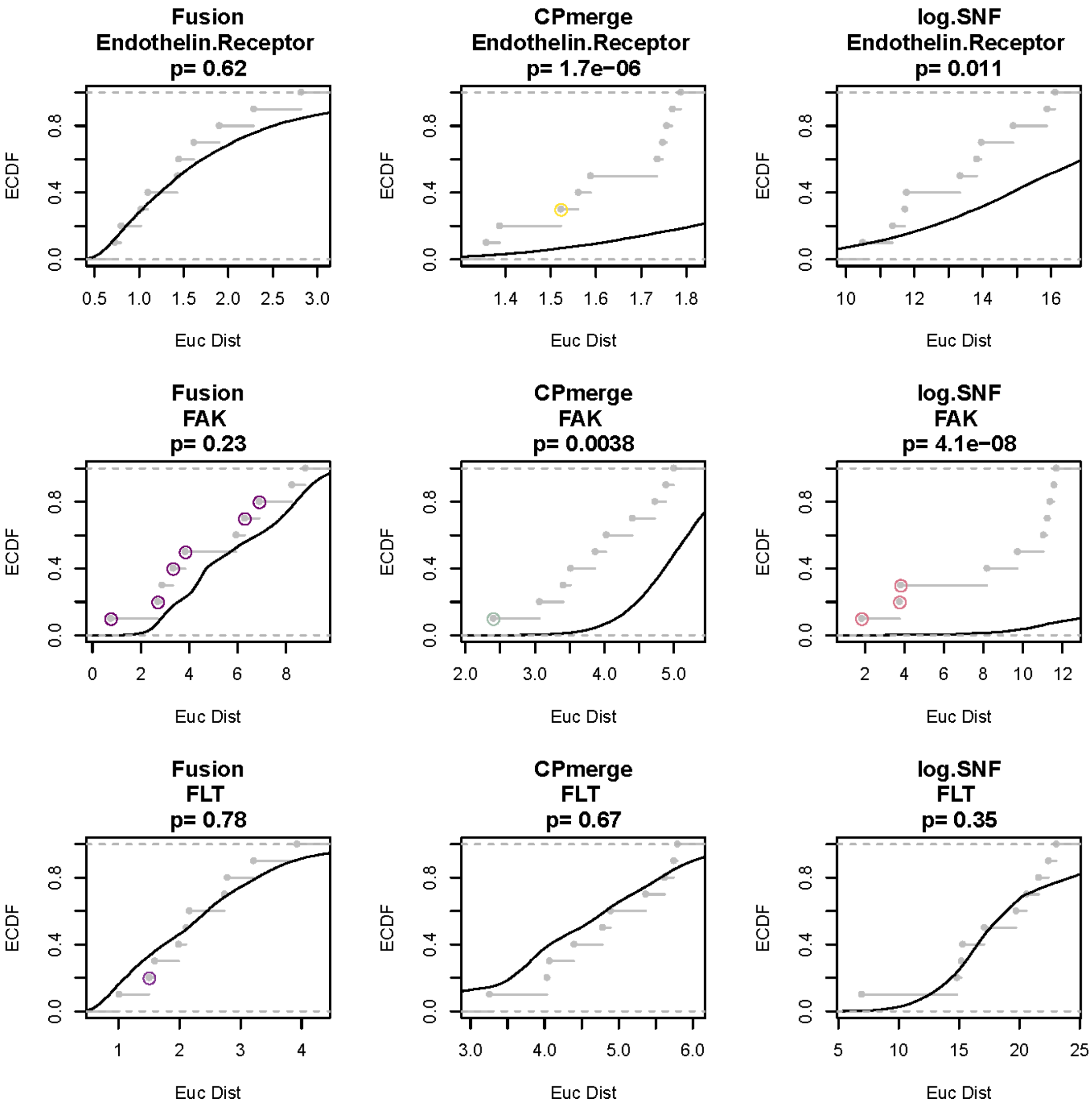
Fusion
Endothelin.Receptor
p= 0.62
CPmerge
Endothelin.Receptor
p= 1.7e−06
log.SNF
Endothelin.Receptor
p= 0.011
Fusion
FAK
p= 0.23
CPmerge
FAK
p= 0.0038
log.SNF
FAK
p= 4.1e−08
Fusion
FLT
p= 0.78
CPmerge
FLT
p= 0.67
log.SNF
FLT
p= 0.35
ECDF
Euc Dist

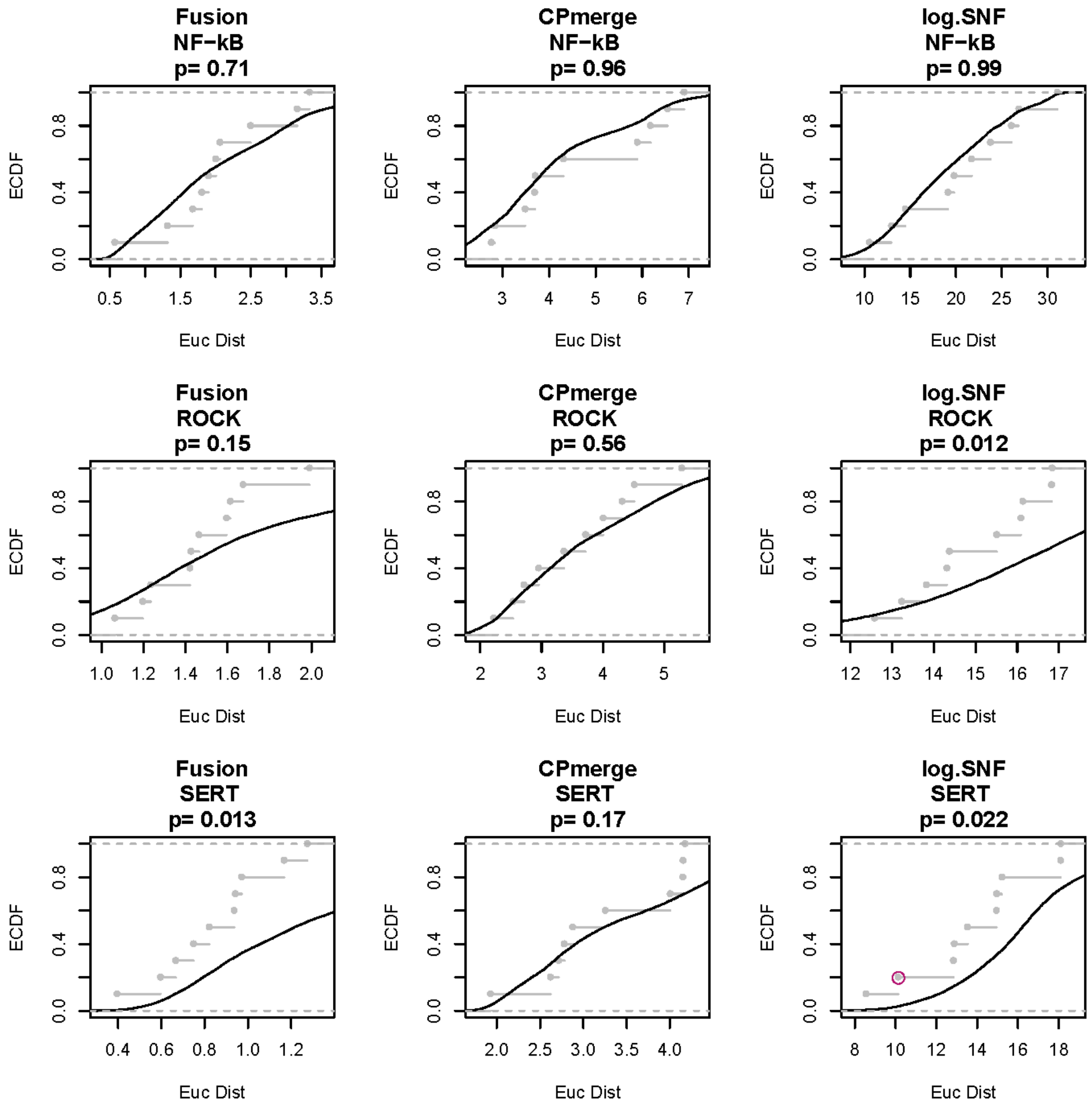
Fusion
NF−kB
p= 0.71
CPmerge
NF−kB
p= 0.96
log.SNF
NF−kB
p= 0.99
Fusion
ROCK
p= 0.15
CPmerge
ROCK
p= 0.56
log.SNF
ROCK
p= 0.012
Fusion
SERT
p= 0.013
CPmerge
SERT
p= 0.17
log.SNF
SERT
p= 0.022
ECDF
Euc Dist

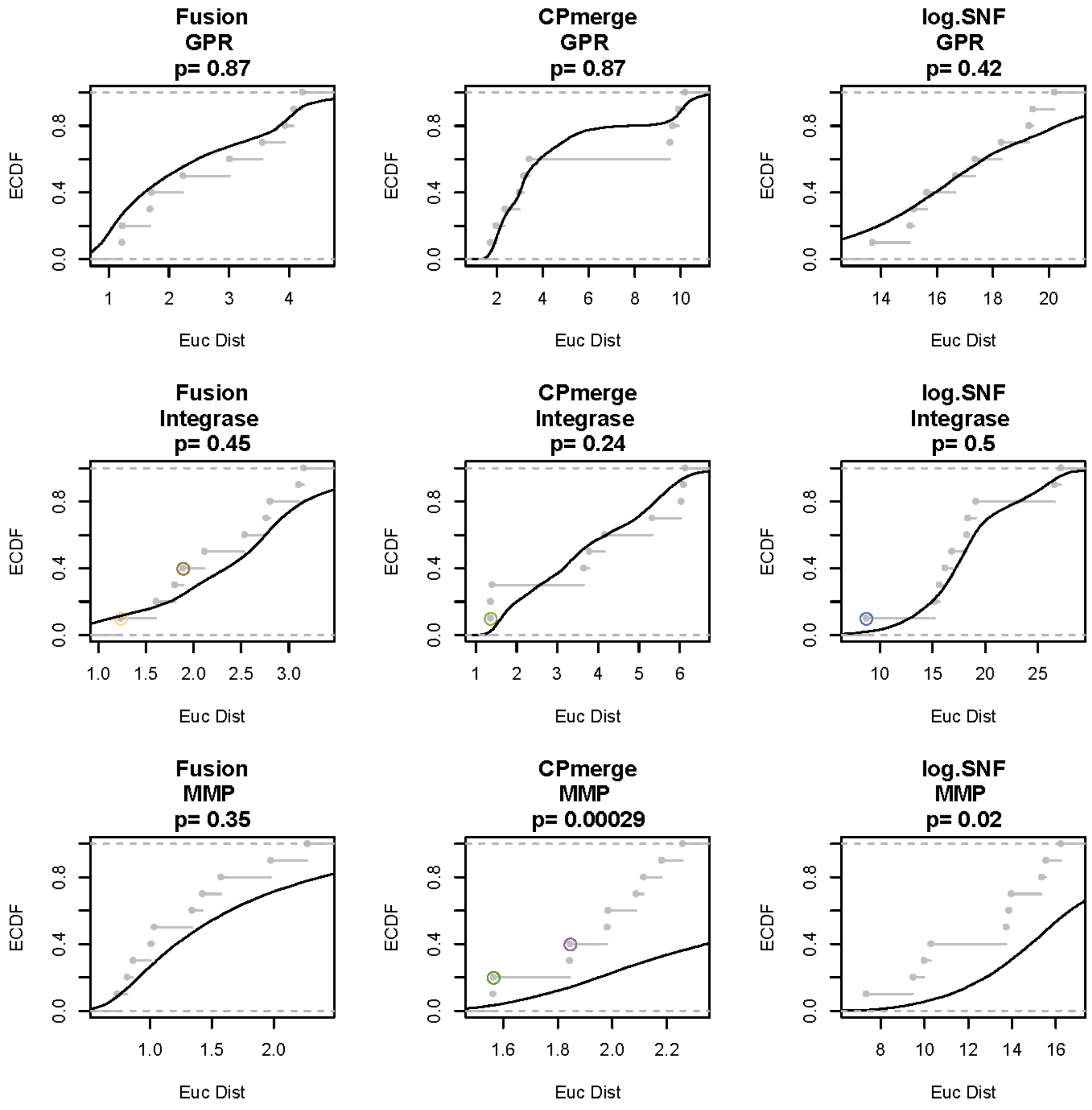
Fusion
GPR
p= 0.87
CPmerge
GPR
p= 0.87
log.SNF
GPR
p= 0.42
Fusion
Integrase
p= 0.45
CPmerge
Integrase
p= 0.24
log.SNF
Integrase
p= 0.5
Fusion
MMP
p= 0.35
CPmerge
MMP
p= 0.00029
log.SNF
MMP
p= 0.02
ECDF
Euc Dist
0.0
0.4
0.8

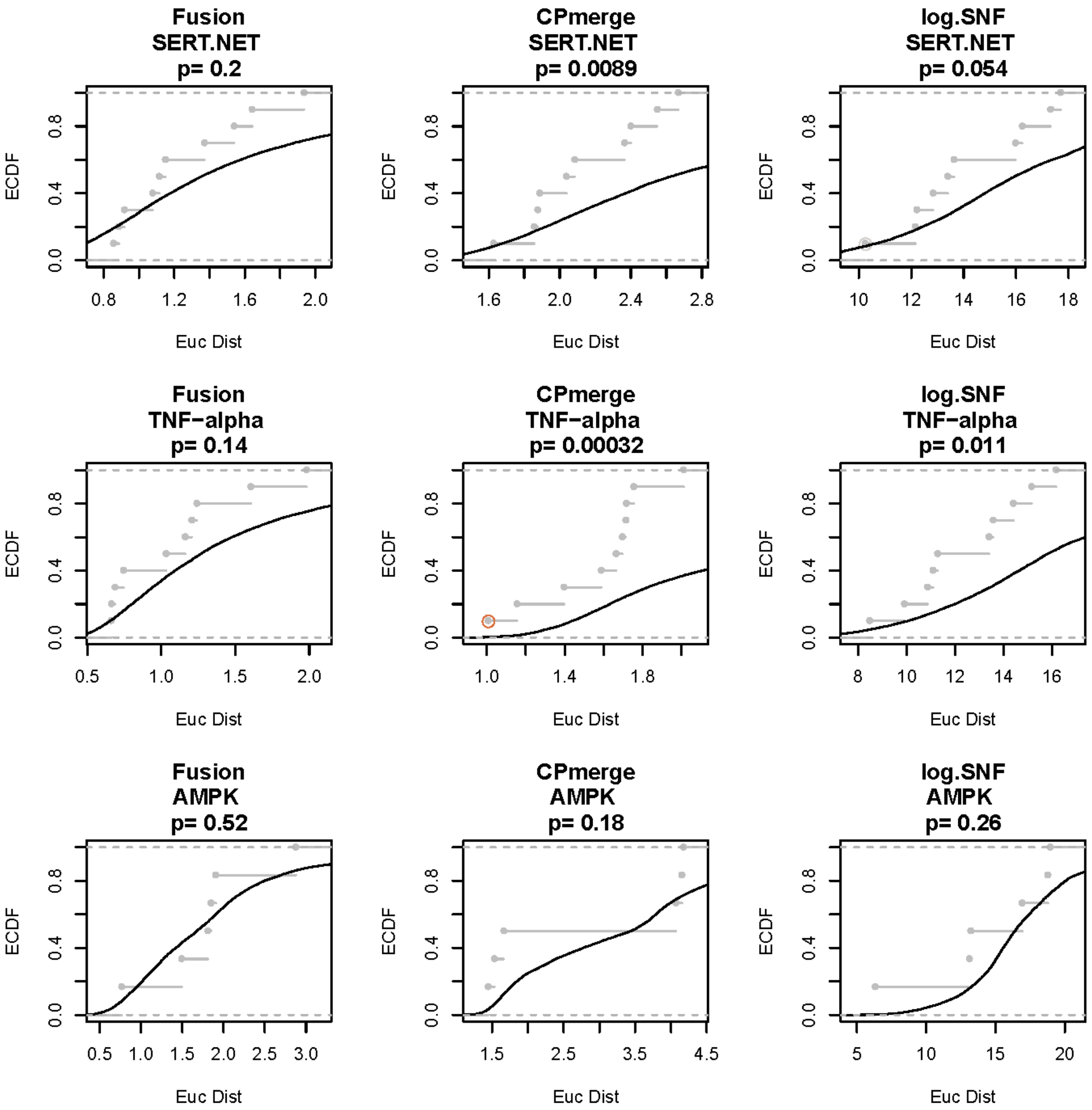
Fusion
SERT.NET
p= 0.2
CPmerge
SERT.NET
p= 0.0089
log.SNF
SERT.NET
p= 0.054
Fusion
TNF−alpha
p= 0.14
CPmerge
TNF−alpha
p= 0.00032
log.SNF
TNF−alpha
p= 0.011
Fusion
AMPK
p= 0.52
CPmerge
AMPK
p= 0.18
log.SNF
AMPK
p= 0.26
ECDF
Euc Dist

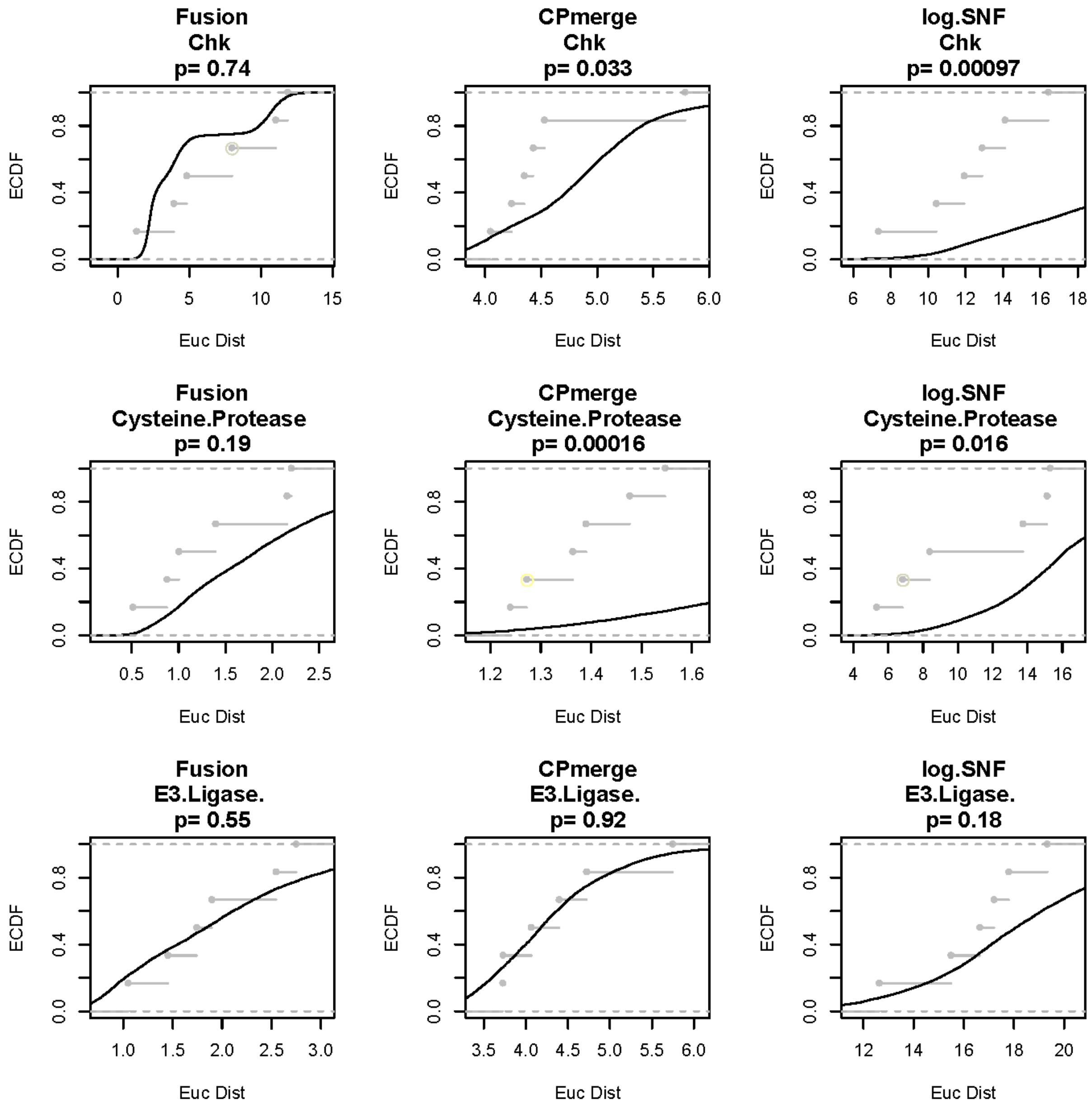
Fusion
Chk
p= 0.74
CPmerge
Chk
p= 0.033
log.SNF
Chk
p= 0.00097
Fusion
Cysteine.Protease
p= 0.19
CPmerge
Cysteine.Protease
p= 0.00016
log.SNF
Cysteine.Protease
p= 0.016
Fusion
E3.Ligase.
p= 0.55
CPmerge
E3.Ligase.
p= 0.92
log.SNF
E3.Ligase.
p= 0.18
ECDF
Euc Dist

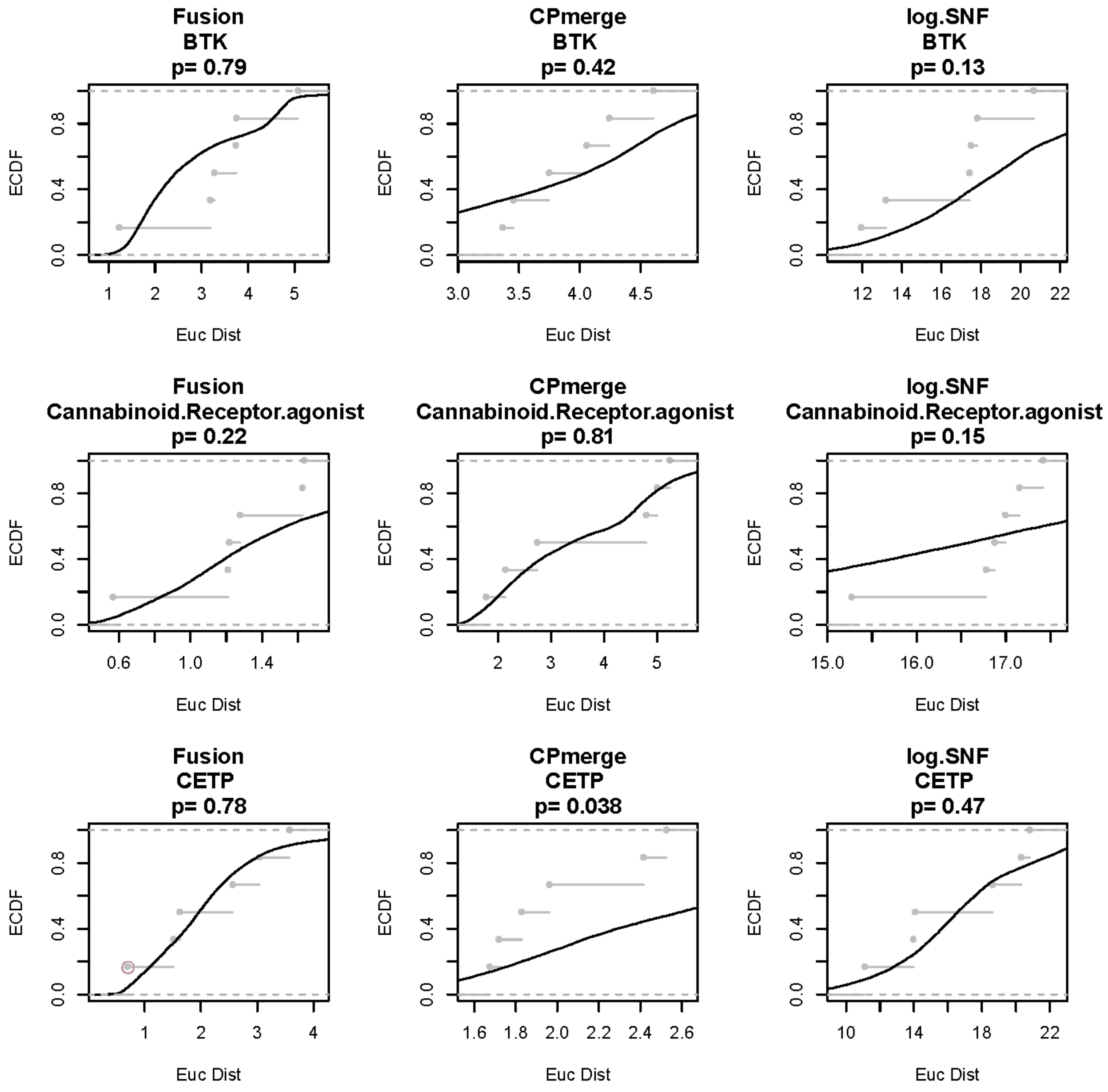
Fusion
BTK
p= 0.79
CPmerge
BTK
p= 0.42
log.SNF
BTK
p= 0.13
Fusion
Cannabinoid.Receptor.agonist
p= 0.22
CPmerge
Cannabinoid.Receptor.agonist
p= 0.81
log.SNF
Cannabinoid.Receptor.agonist
p= 0.15
Fusion
CETP
p= 0.78
CPmerge
CETP
p= 0.038
log.SNF
CETP
p= 0.47
ECDF
0.0
0.4
0.8
Euc Dist
1 2 3 4 5
3.0 3.5 4.0 4.5
12 14 16 18 20 22
0.6 1.0 1.4
2 3 4 5
15.0 16.0 17.0
1 2 3 4
1.6 1.8 2.0 2.2 2.4 2.6
10 14 18 22

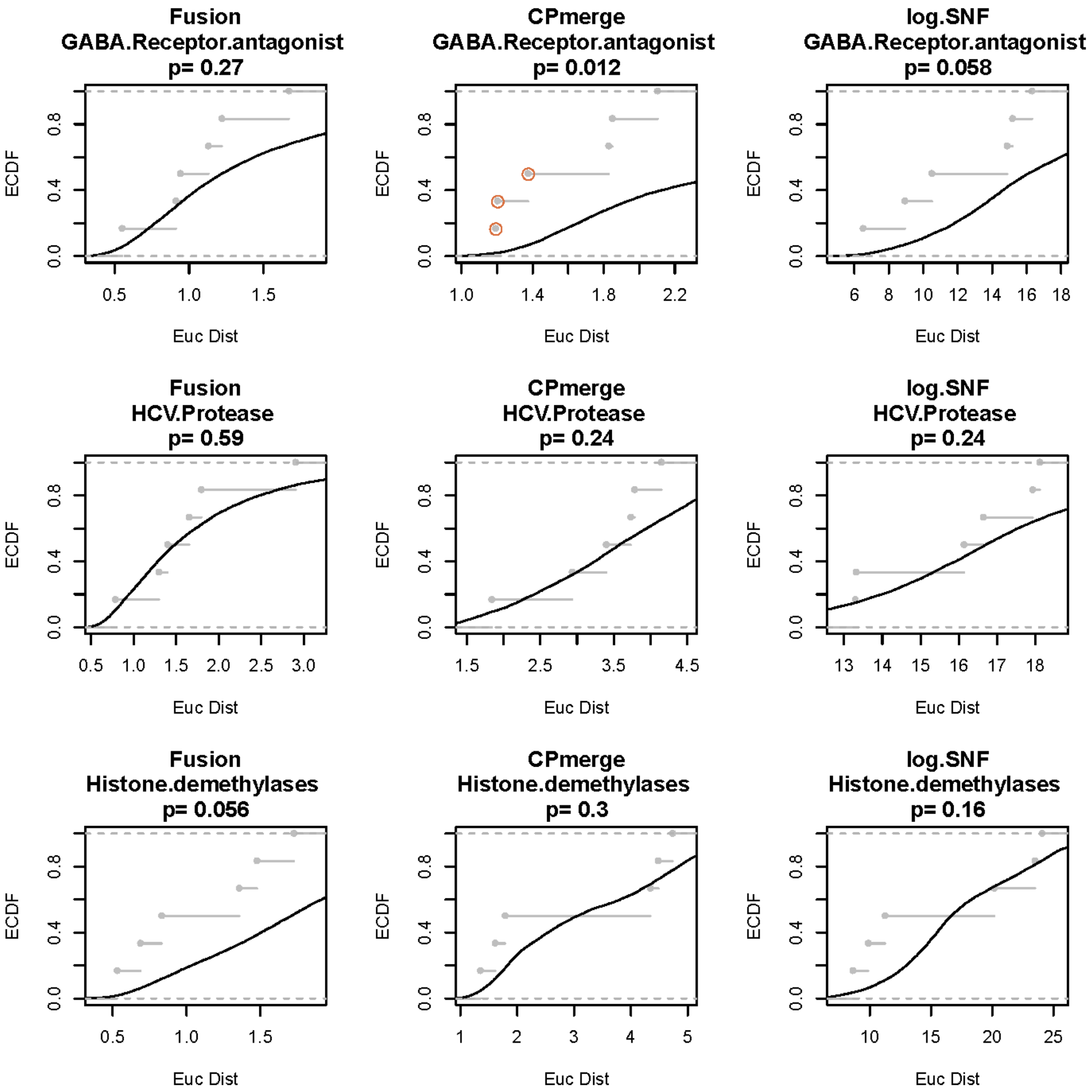
Fusion
GABA.Receptor.antagonist
p= 0.27
CPmerge
GABA.Receptor.antagonist
p= 0.012
log.SNF
GABA.Receptor.antagonist
p= 0.058
Fusion
HCV.Protease
p= 0.59
CPmerge
HCV.Protease
p= 0.24
log.SNF
HCV.Protease
p= 0.24
Fusion
Histone.demethylases
p= 0.056
CPmerge
Histone.demethylases
p= 0.3
log.SNF
Histone.demethylases
p= 0.16
ECDF
Euc Dist

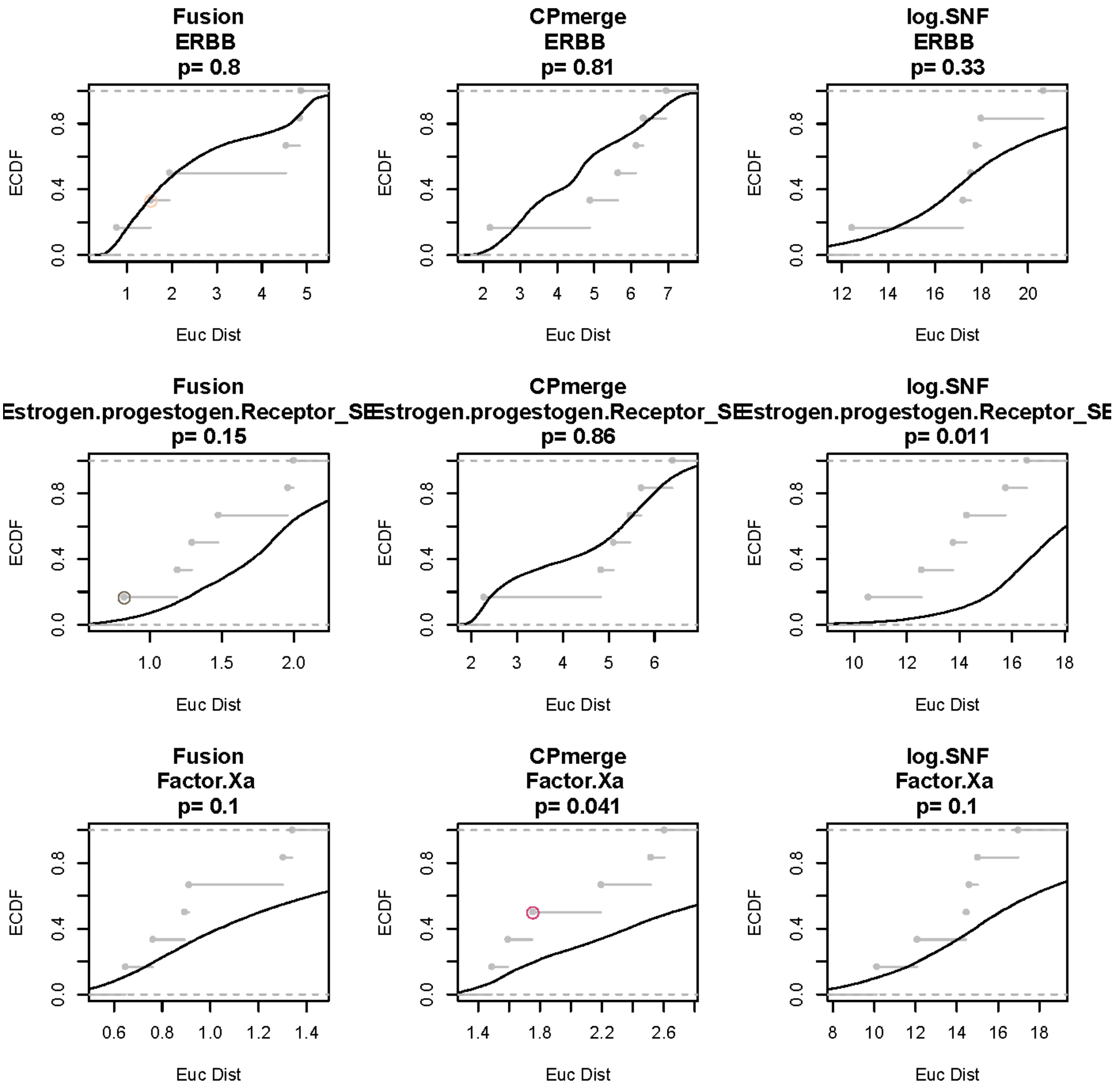
Fusion
ERBB
p= 0.8
CPmerge
ERBB
p= 0.81
log.SNF
ERBB
p= 0.33
Fusion
Estrogen.progestogen.Receptor_SE
p= 0.15
CPmerge
Estrogen.progestogen.Receptor_SE
p= 0.86
log.SNF
Estrogen.progestogen.Receptor_SE
p= 0.011
Fusion
Factor.Xa
p= 0.1
CPmerge
Factor.Xa
p= 0.041
log.SNF
Factor.Xa
p= 0.1
ECDF
Euc Dist

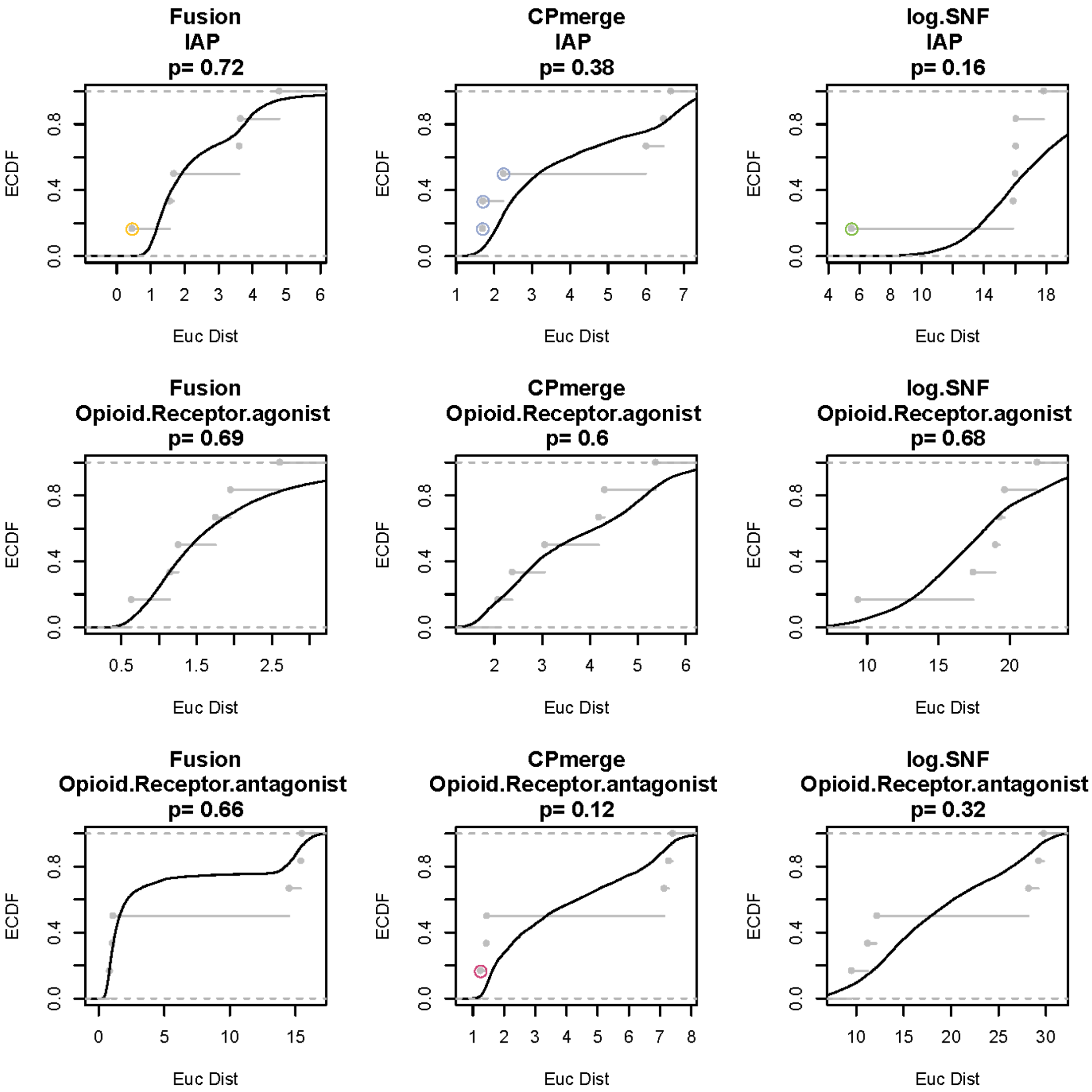

Fusion
IAP
p= 0.72
CPmerge
IAP
p= 0.38
log.SNF
IAP
p= 0.16
Fusion
Opioid.Receptor.agonist
p= 0.69
CPmerge
Opioid.Receptor.agonist
p= 0.6
log.SNF
Opioid.Receptor.agonist
p= 0.68
Fusion
Opioid.Receptor.antagonist
p= 0.66
CPmerge
Opioid.Receptor.antagonist
p= 0.12
log.SNF
Opioid.Receptor.antagonist
p= 0.32
ECDF
Euc Dist

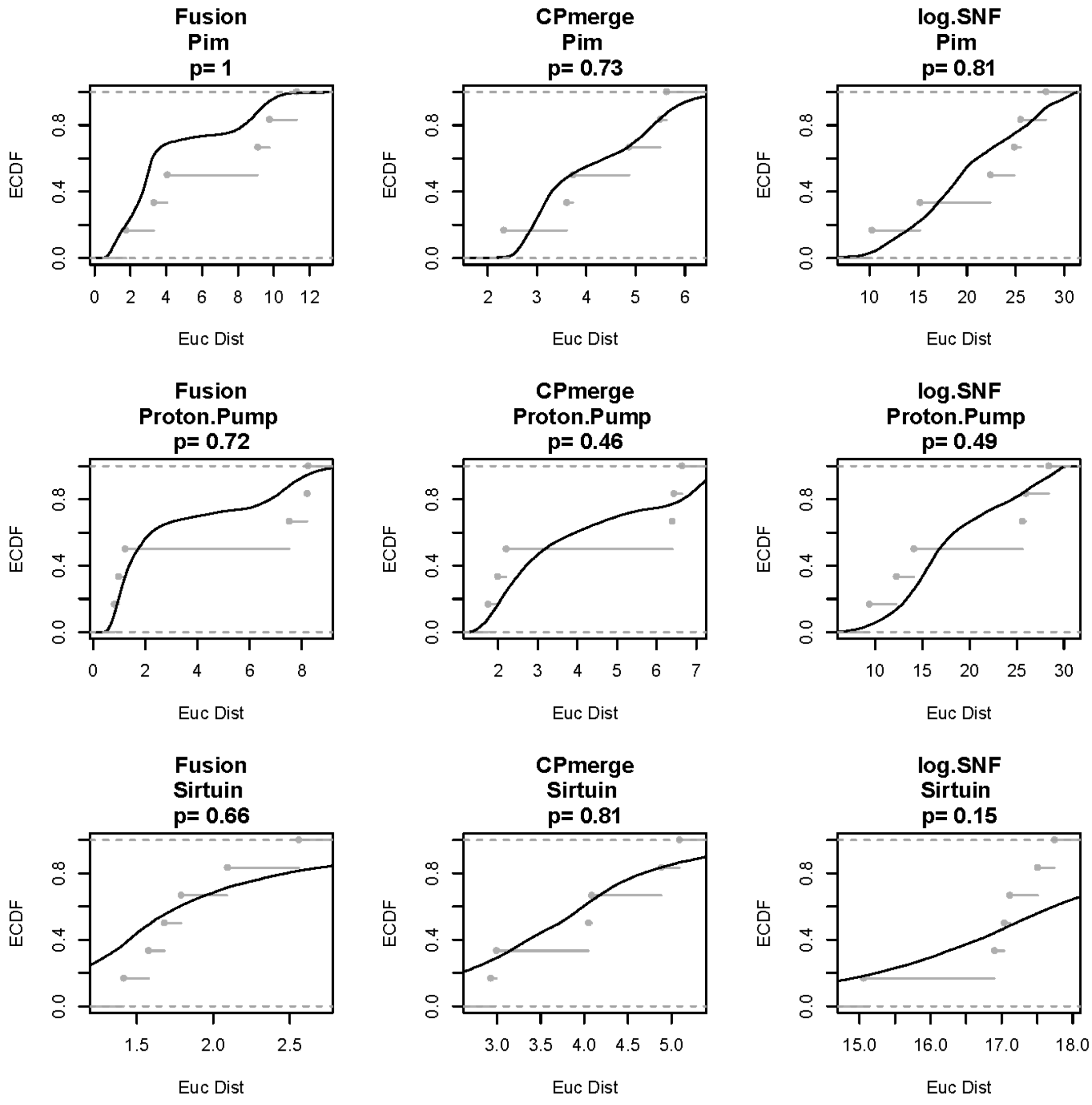
Fusion
Pim
p= 1
CPmerge
Pim
p= 0.73
log.SNF
Pim
p= 0.81
Fusion
Proton.Pump
p= 0.72
CPmerge
Proton.Pump
p= 0.46
log.SNF
Proton.Pump
p= 0.49
Fusion
Sirtuin
p= 0.66
CPmerge
Sirtuin
p= 0.81
log.SNF
Sirtuin
p= 0.15
ECDF
Euc Dist

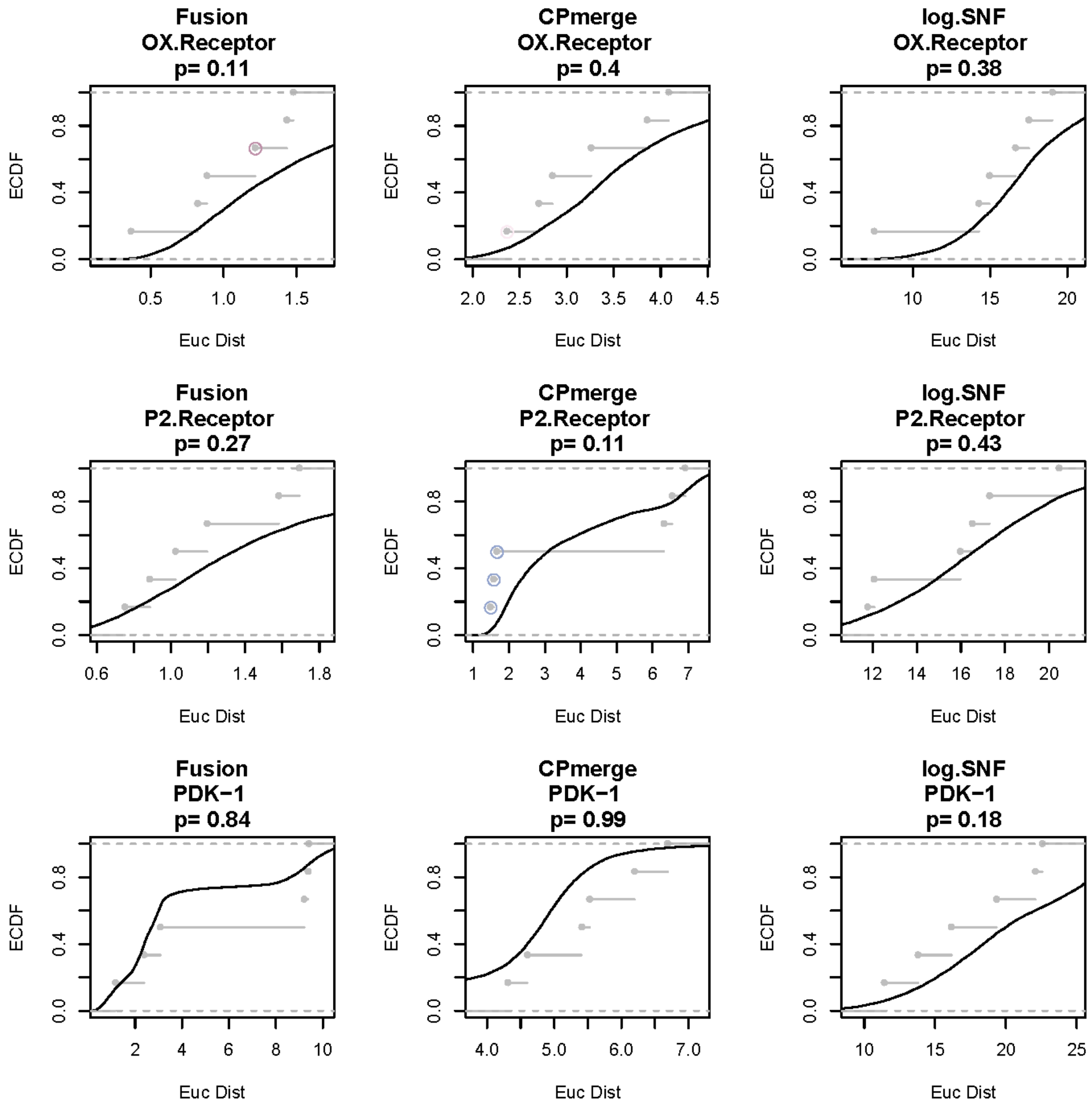
Fusion
OX.Receptor
p= 0.11
CPmerge
OX.Receptor
p= 0.4
log.SNF
OX.Receptor
p= 0.38
Fusion
P2.Receptor
p= 0.27
CPmerge
P2.Receptor
p= 0.11
log.SNF
P2.Receptor
p= 0.43
Fusion
PDK−1
p= 0.84
CPmerge
PDK−1
p= 0.99
log.SNF
PDK−1
p= 0.18
ECDF
Euc Dist

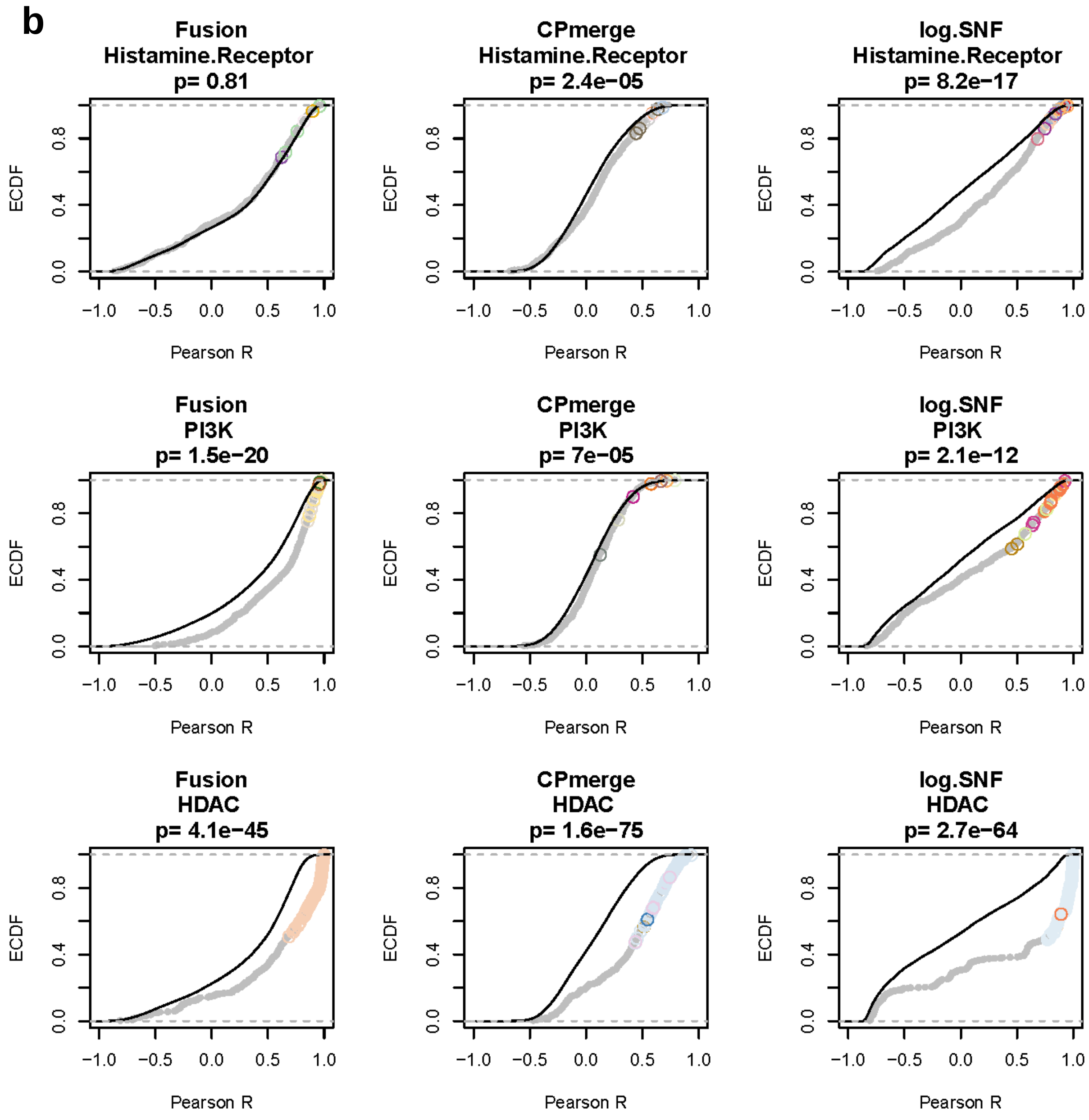
b
Fusion
Histamine.Receptor
p= 0.81
CPmerge
Histamine.Receptor
p= 2.4e−05
log.SNF
Histamine.Receptor
p= 8.2e−17
Fusion
PI3K
p= 1.5e−20
CPmerge
PI3K
p= 7e−05
log.SNF
PI3K
p= 2.1e−12
Fusion
HDAC
p= 4.1e−45
CPmerge
HDAC
p= 1.6e−75
log.SNF
HDAC
p= 2.7e−64
ECDF
0.0
0.4
0.8
−1.0
−0.5
0.0
0.5
1.0
Pearson R

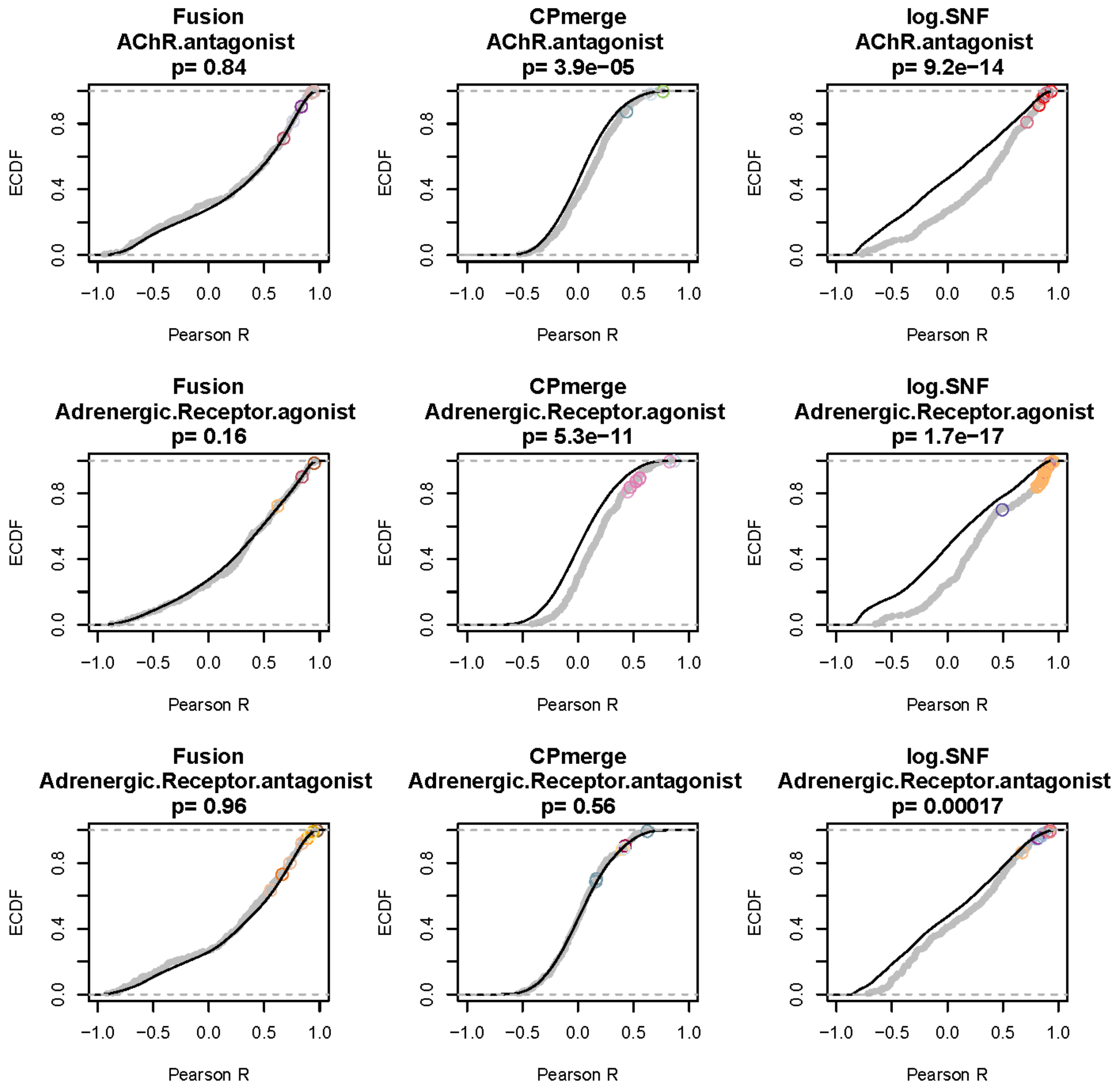
Fusion
AChR.antagonist
p= 0.84
CPmerge
AChR.antagonist
p= 3.9e−05
log.SNF
AChR.antagonist
p= 9.2e−14
Fusion
Adrenergic.Receptor.agonist
p= 0.16
CPmerge
Adrenergic.Receptor.agonist
p= 5.3e−11
log.SNF
Adrenergic.Receptor.agonist
p= 1.7e−17
Fusion
Adrenergic.Receptor.antagonist
p= 0.96
CPmerge
Adrenergic.Receptor.antagonist
p= 0.56
log.SNF
Adrenergic.Receptor.antagonist
p= 0.00017
ECDF
Pearson R
0.0
0.4
0.8
−1.0
−0.5
0.0
0.5
1.0

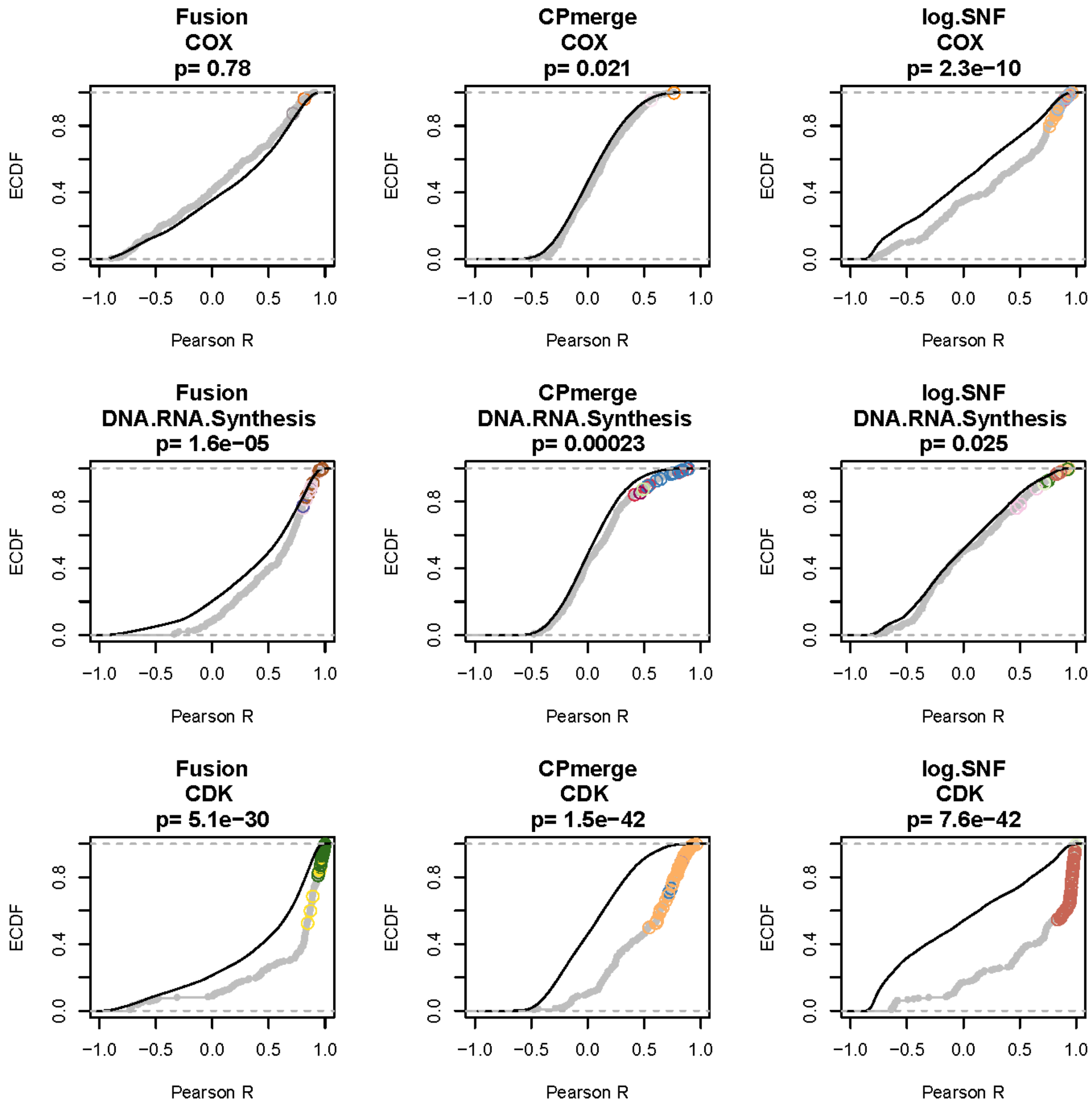
Fusion
COX
p= 0.78
CPmerge
COX
p= 0.021
log.SNF
COX
p= 2.3e−10
Fusion
DNA.RNA.Synthesis
p= 1.6e−05
CPmerge
DNA.RNA.Synthesis
p= 0.00023
log.SNF
DNA.RNA.Synthesis
p= 0.025
Fusion
CDK
p= 5.1e−30
CPmerge
CDK
p= 1.5e−42
log.SNF
CDK
p= 7.6e−42
ECDF
0.0
0.4
0.8
−1.0
−0.5
0.0
0.5
1.0
Pearson R

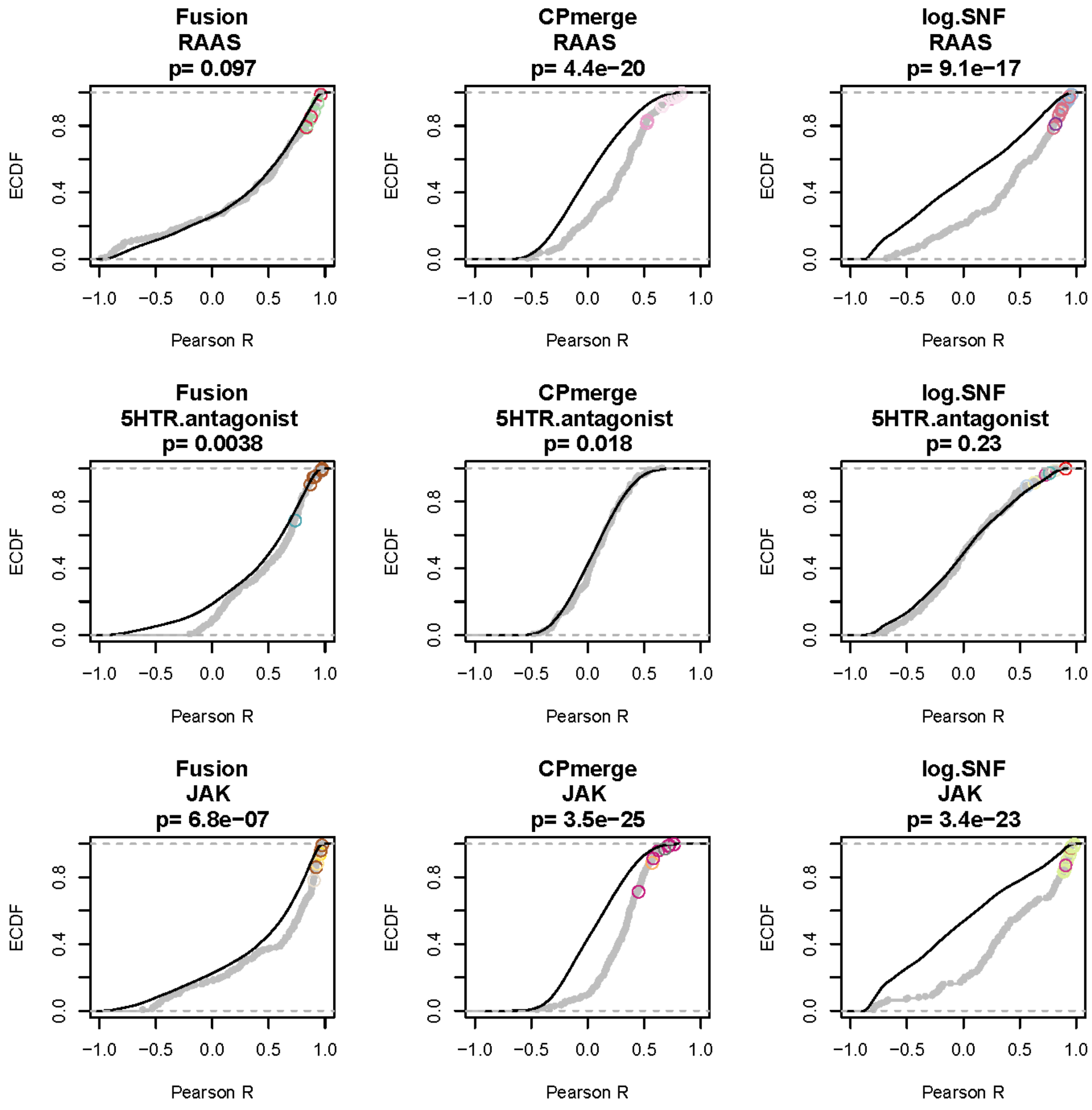

Fusion
RAAS
p= 0.097
CPmerge
RAAS
p= 4.4e−20
log.SNF
RAAS
p= 9.1e−17
Fusion
5HTR.antagonist
p= 0.0038
CPmerge
5HTR.antagonist
p= 0.018
log.SNF
5HTR.antagonist
p= 0.23
Fusion
JAK
p= 6.8e−07
CPmerge
JAK
p= 3.5e−25
log.SNF
JAK
p= 3.4e−23
ECDF
Pearson R
0.0
0.4
0.8
−1.0
−0.5
0.0
0.5
1.0

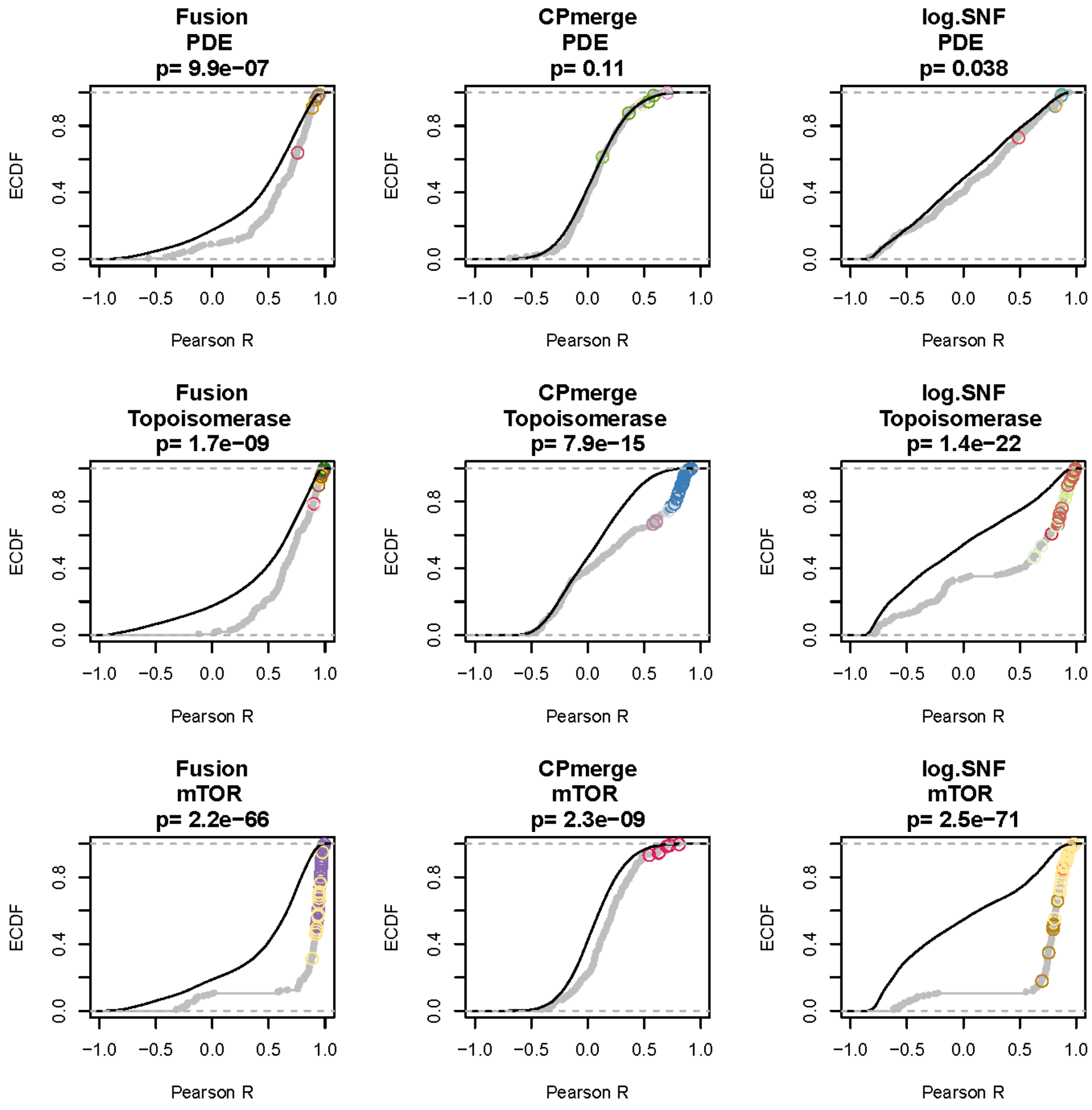

Fusion
PDE
p= 9.9e−07
CPmerge
PDE
p= 0.11
log.SNF
PDE
p= 0.038
Fusion
Topoisomerase
p= 1.7e−09
CPmerge
Topoisomerase
p= 7.9e−15
log.SNF
Topoisomerase
p= 1.4e−22
Fusion
mTOR
p= 2.2e−66
CPmerge
mTOR
p= 2.3e−09
log.SNF
mTOR
p= 2.5e−71
ECDF
0.0
0.4
0.8
Pearson R
−1.0
−0.5
0.0
0.5
1.0

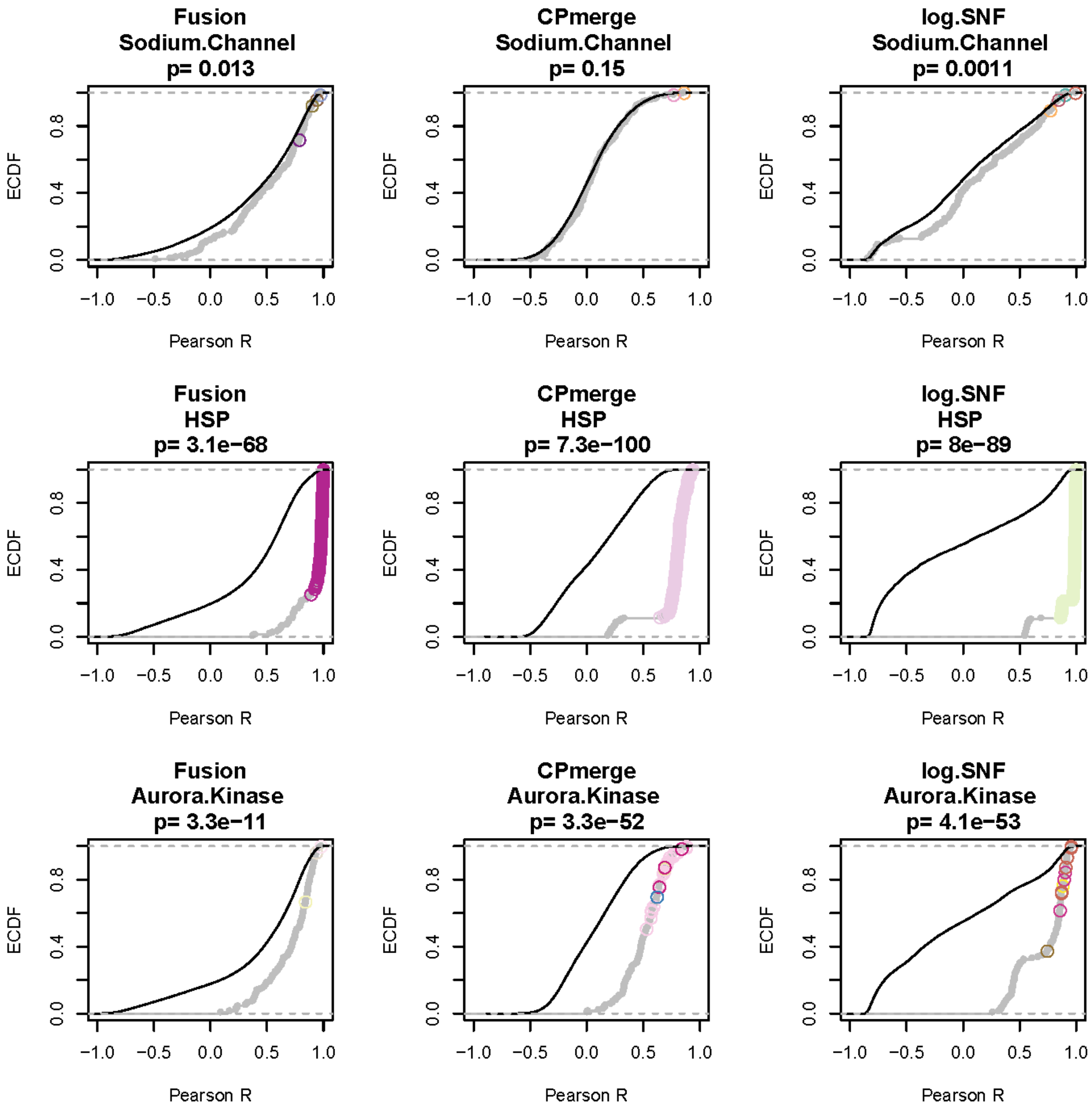
Fusion
Sodium.Channel
p= 0.013
CPmerge
Sodium.Channel
p= 0.15
log.SNF
Sodium.Channel
p= 0.0011
Fusion
HSP
p= 3.1e−68
CPmerge
HSP
p= 7.3e−100
log.SNF
HSP
p= 8e−89
Fusion
Aurora.Kinase
p= 3.3e−11
CPmerge
Aurora.Kinase
p= 3.3e−52
log.SNF
Aurora.Kinase
p= 4.1e−53
ECDF
Pearson R

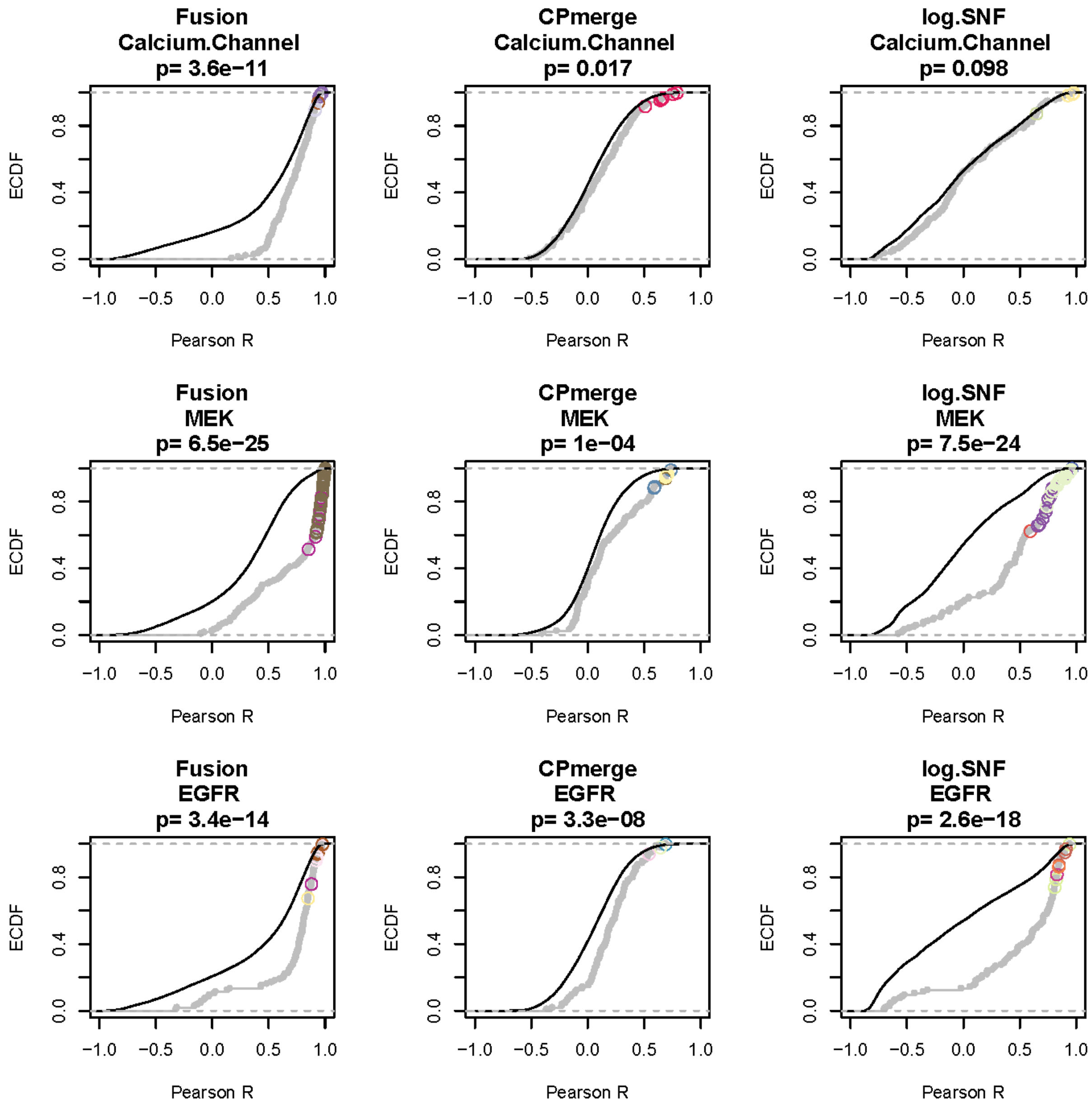
Fusion
Calcium.Channel
p= 3.6e−11
CPmerge
Calcium.Channel
p= 0.017
log.SNF
Calcium.Channel
p= 0.098
Fusion
MEK
p= 6.5e−25
CPmerge
MEK
p= 1e−04
log.SNF
MEK
p= 7.5e−24
Fusion
EGFR
p= 3.4e−14
CPmerge
EGFR
p= 3.3e−08
log.SNF
EGFR
p= 2.6e−18
ECDF
Pearson R
0.0
0.4
0.8
−1.0
−0.5
0.0
0.5
1.0

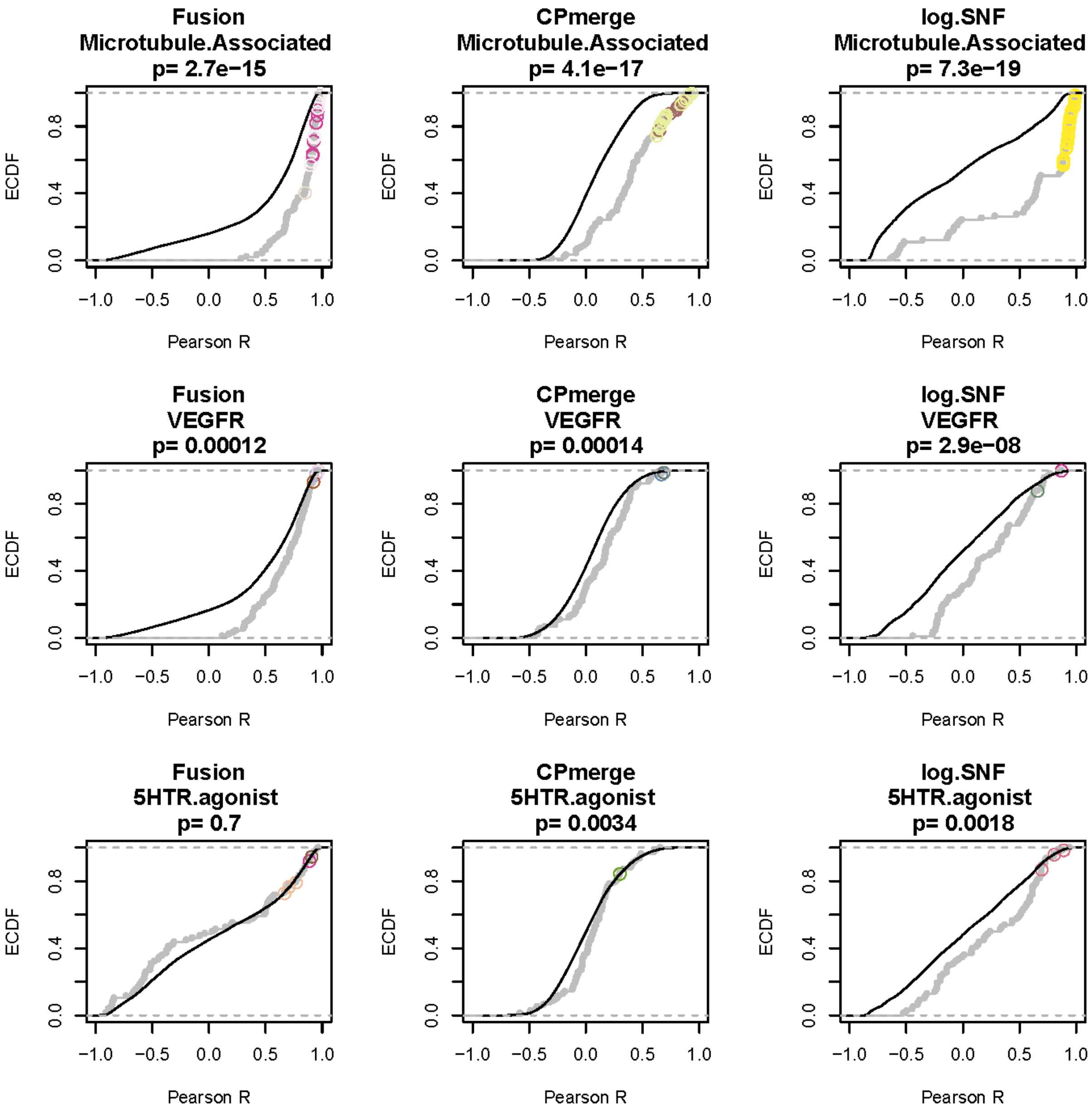

Fusion
Microtubule.Associated
p= 2.7e−15
CPmerge
Microtubule.Associated
p= 4.1e−17
log.SNF
Microtubule.Associated
p= 7.3e−19
Fusion
VEGFR
p= 0.00012
CPmerge
VEGFR
p= 0.00014
log.SNF
VEGFR
p= 2.9e−08
Fusion
5HTR.agonist
p= 0.7
CPmerge
5HTR.agonist
p= 0.0034
log.SNF
5HTR.agonist
p= 0.0018
ECDF
0.0
0.4
0.8
−1.0
−0.5
0.0
0.5
1.0
Pearson R

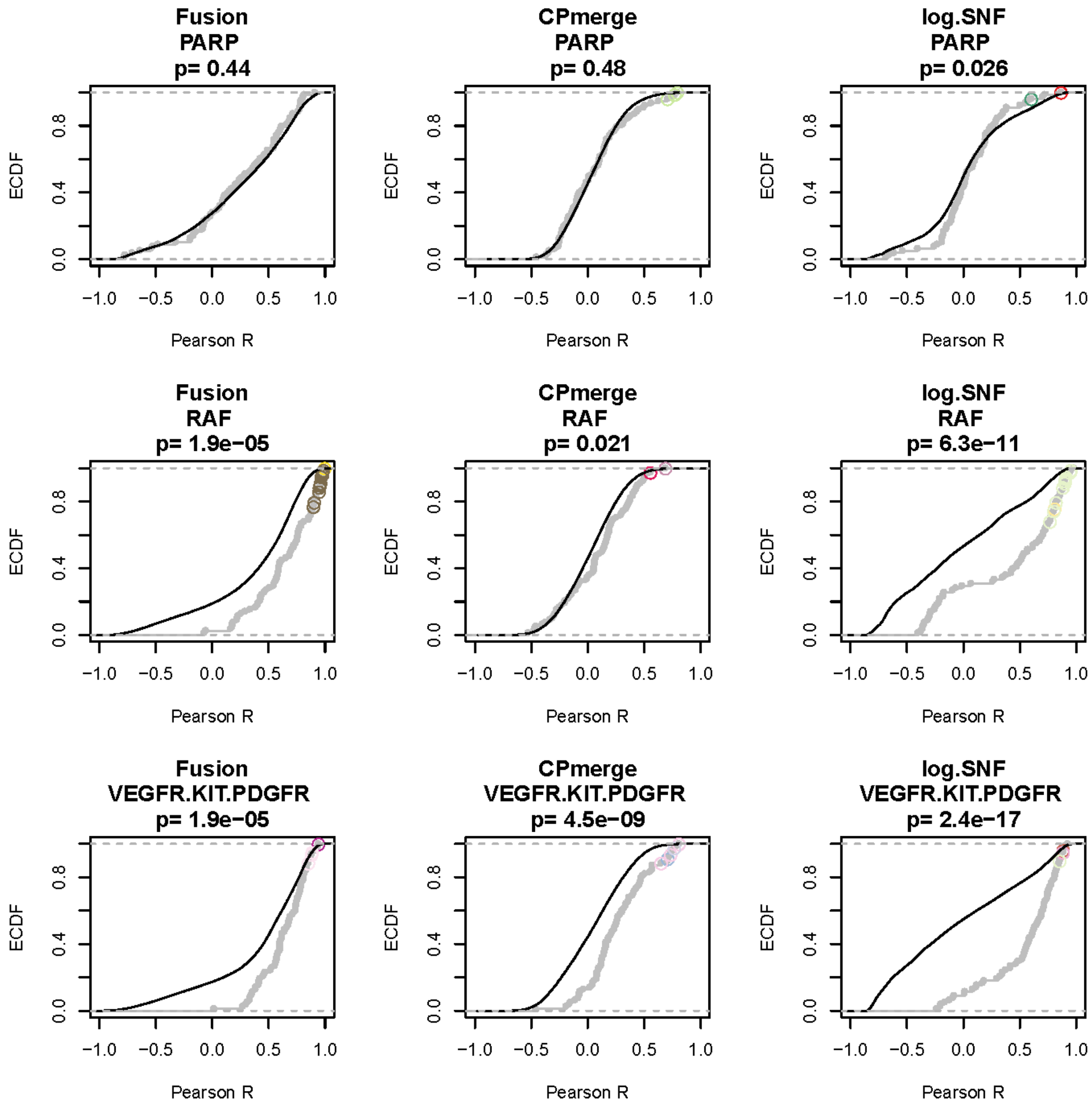
Fusion
PARP
p= 0.44
CPmerge
PARP
p= 0.48
log.SNF
PARP
p= 0.026
Fusion
RAF
p= 1.9e−05
CPmerge
RAF
p= 0.021
log.SNF
RAF
p= 6.3e−11
Fusion
VEGFR.KIT.PDGFR
p= 1.9e−05
CPmerge
VEGFR.KIT.PDGFR
p= 4.5e−09
log.SNF
VEGFR.KIT.PDGFR
p= 2.4e−17
ECDF
Pearson R
0.0
0.4
0.8
−1.0
−0.5
0.0
0.5
1.0

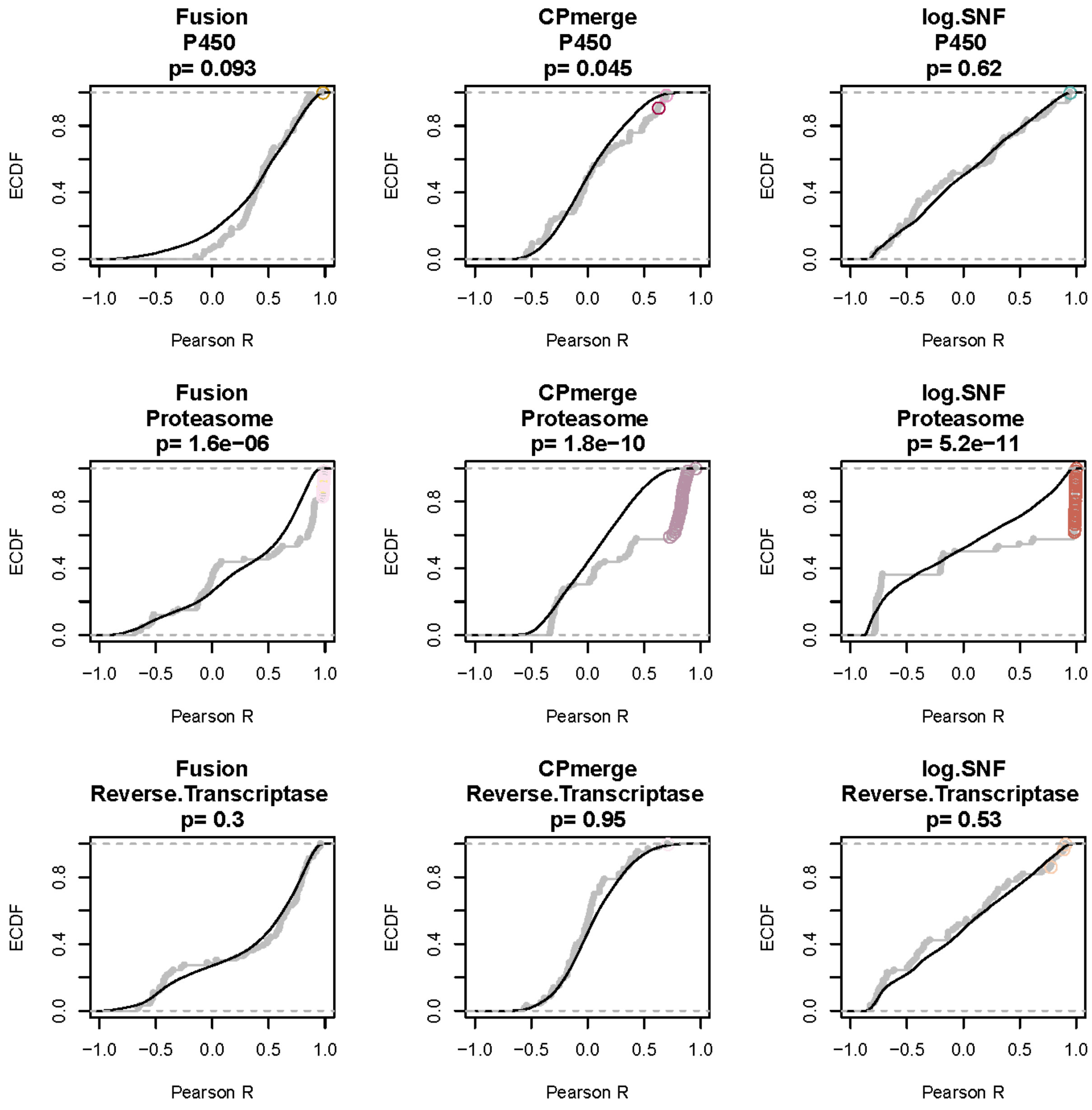
Fusion
P450
p= 0.093
CPmerge
P450
p= 0.045
log.SNF
P450
p= 0.62
Fusion
Proteasome
p= 1.6e−06
CPmerge
Proteasome
p= 1.8e−10
log.SNF
Proteasome
p= 5.2e−11
Fusion
Reverse.Transcriptase
p= 0.3
CPmerge
Reverse.Transcriptase
p= 0.95
log.SNF
Reverse.Transcriptase
p= 0.53
ECDF
0.0
0.4
0.8
−1.0
−0.5
0.0
0.5
1.0
Pearson R

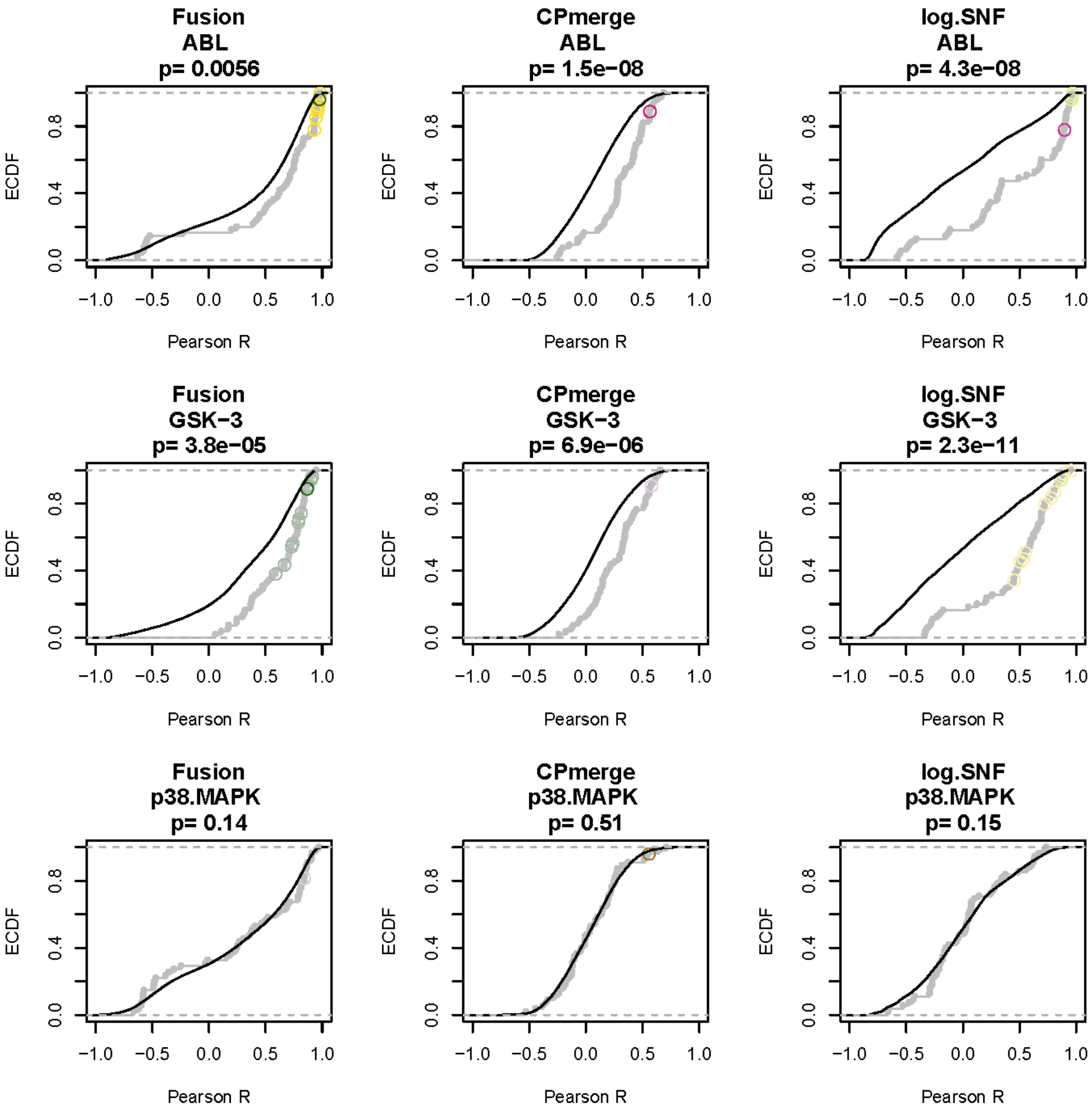
Fusion
ABL
p= 0.0056
CPmerge
ABL
p= 1.5e−08
log.SNF
ABL
p= 4.3e−08
Fusion
GSK−3
p= 3.8e−05
CPmerge
GSK−3
p= 6.9e−06
log.SNF
GSK−3
p= 2.3e−11
Fusion
p38.MAPK
p= 0.14
CPmerge
p38.MAPK
p= 0.51
log.SNF
p38.MAPK
p= 0.15
ECDF
Pearson R
0.0
0.4
0.8
−1.0
−0.5
0.0
0.5
1.0

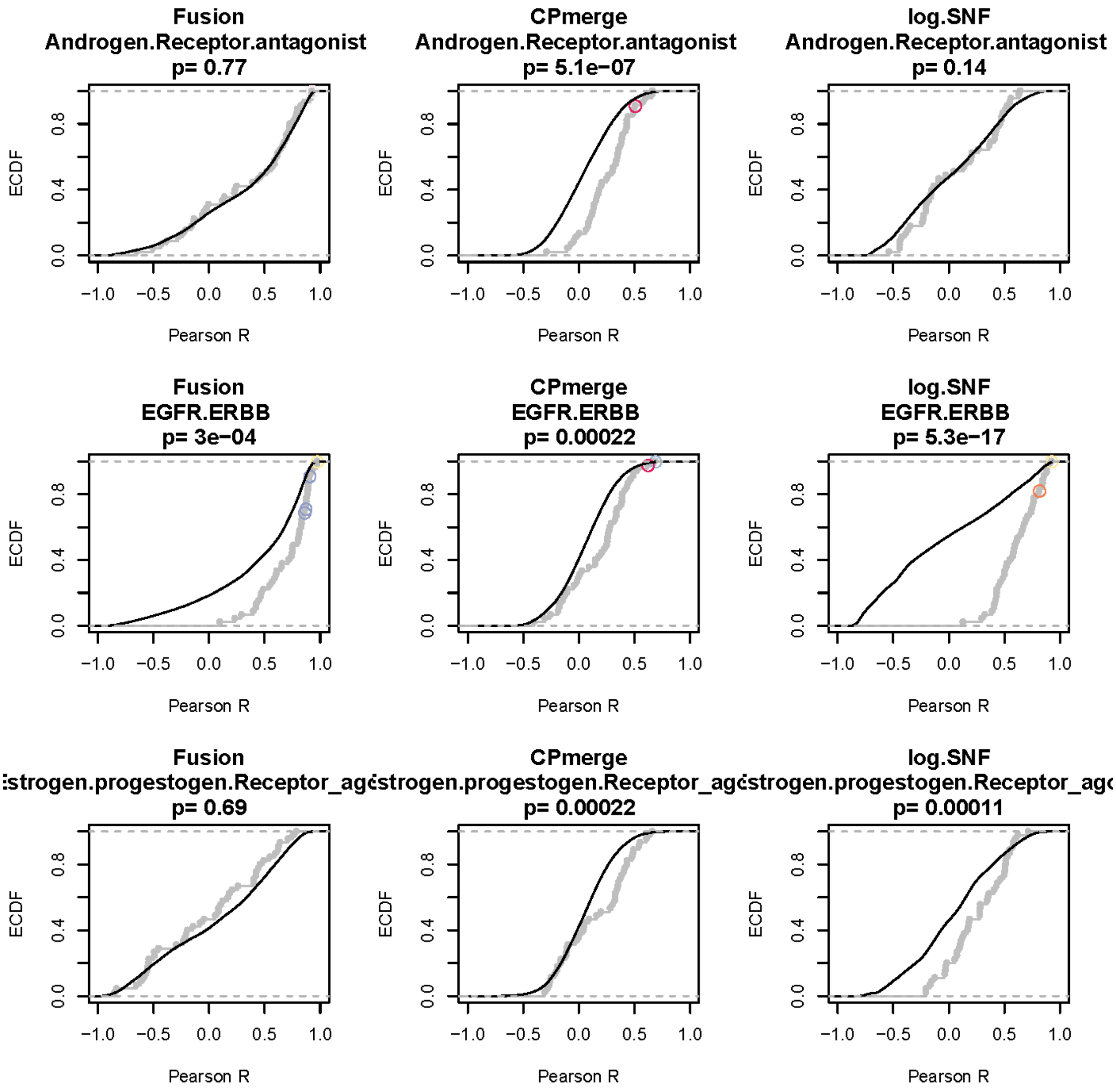
Fusion
Androgen.Receptor.antagonist
p= 0.77
CPmerge
Androgen.Receptor.antagonist
p= 5.1e−07
log.SNF
Androgen.Receptor.antagonist
p= 0.14
Fusion
EGFR.ERBB
p= 3e−04
CPmerge
EGFR.ERBB
p= 0.00022
log.SNF
EGFR.ERBB
p= 5.3e−17
Fusion
:strogen.progestogen.Receptor_agc
p= 0.69
CPmerge
:strogen.progestogen.Receptor_agc
p= 0.00022
log.SNF
:strogen.progestogen.Receptor_agc
p= 0.00011
ECDF
0.0
0.4
0.8
Pearson R
−1.0
−0.5
0.0
0.5
1.0

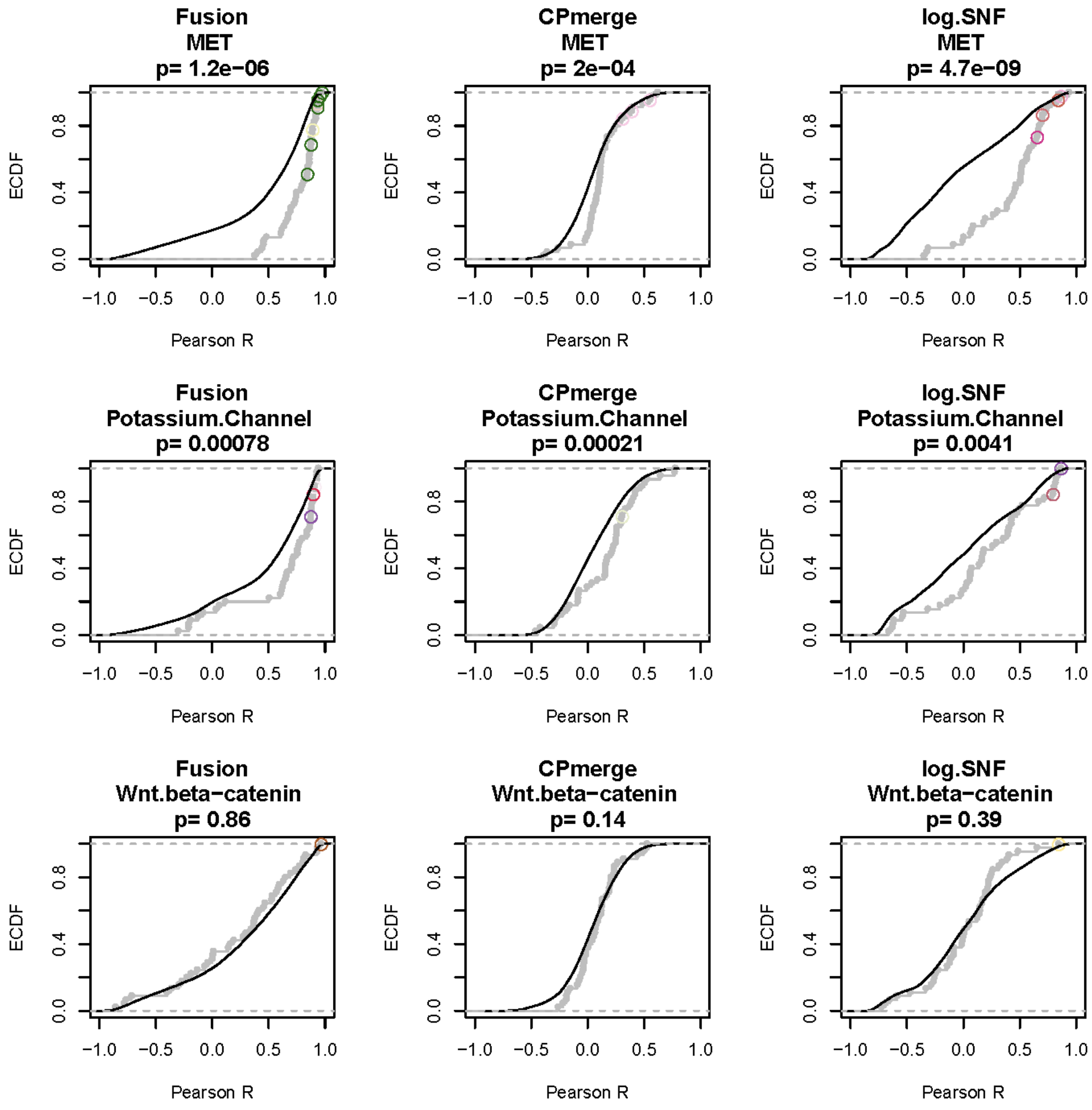

Fusion
MET
p= 1.2e−06
CPmerge
MET
p= 2e−04
log.SNF
MET
p= 4.7e−09
Fusion
Potassium.Channel
p= 0.00078
CPmerge
Potassium.Channel
p= 0.00021
log.SNF
Potassium.Channel
p= 0.0041
Fusion
Wnt.beta−catenin
p= 0.86
CPmerge
Wnt.beta−catenin
p= 0.14
log.SNF
Wnt.beta−catenin
p= 0.39
ECDF
0.0
0.4
0.8
Pearson R
−1.0
−0.5
0.0
0.5
1.0

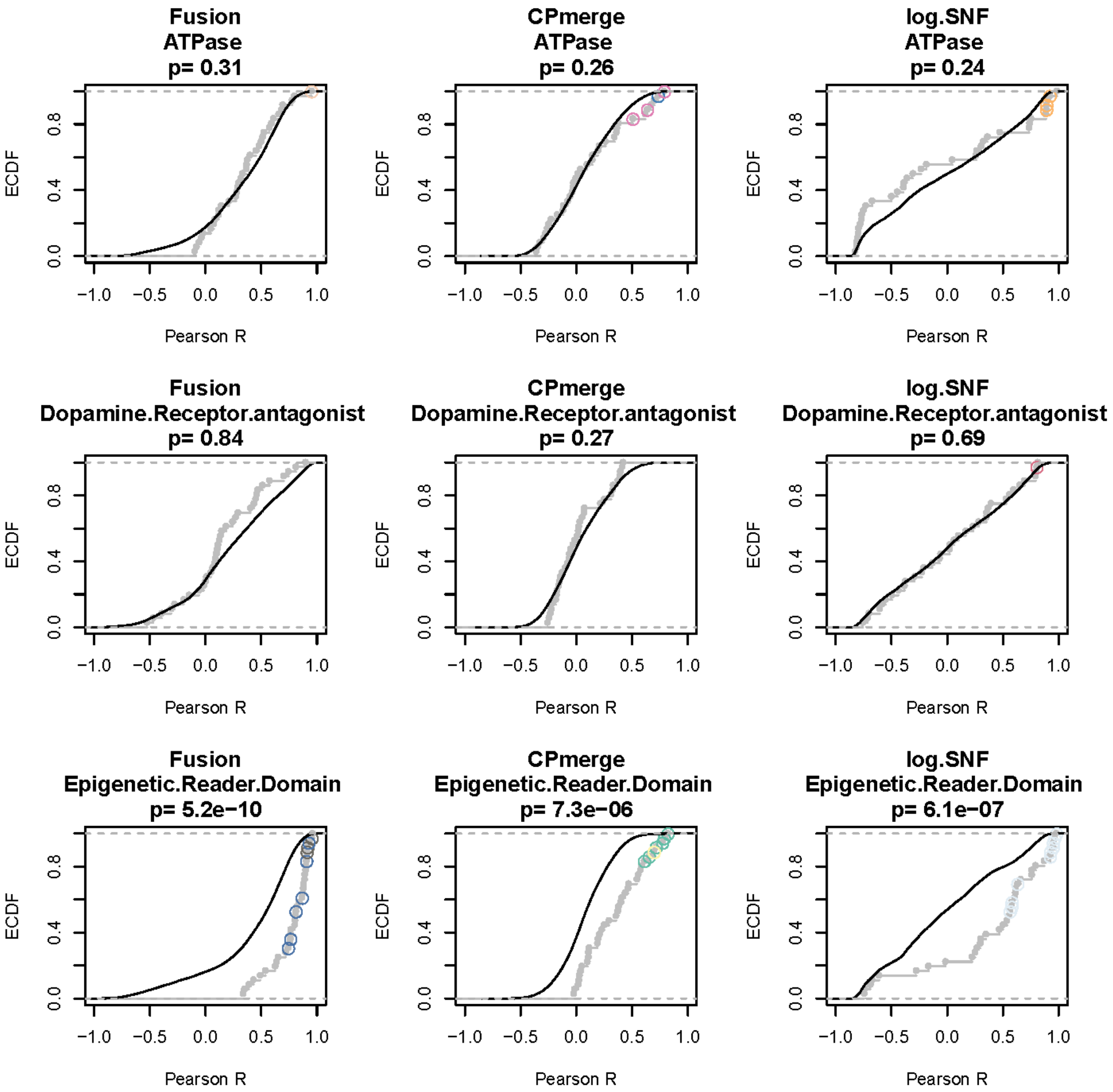
Fusion
ATPase
p= 0.31
CPmerge
ATPase
p= 0.26
log.SNF
ATPase
p= 0.24
Fusion
Dopamine.Receptor.antagonist
p= 0.84
CPmerge
Dopamine.Receptor.antagonist
p= 0.27
log.SNF
Dopamine.Receptor.antagonist
p= 0.69
Fusion
Epigenetic.Reader.Domain
p= 5.2e−10
CPmerge
Epigenetic.Reader.Domain
p= 7.3e−06
log.SNF
Epigenetic.Reader.Domain
p= 6.1e−07
ECDF
0.0
0.4
0.8
−1.0
−0.5
0.0
0.5
1.0
Pearson R

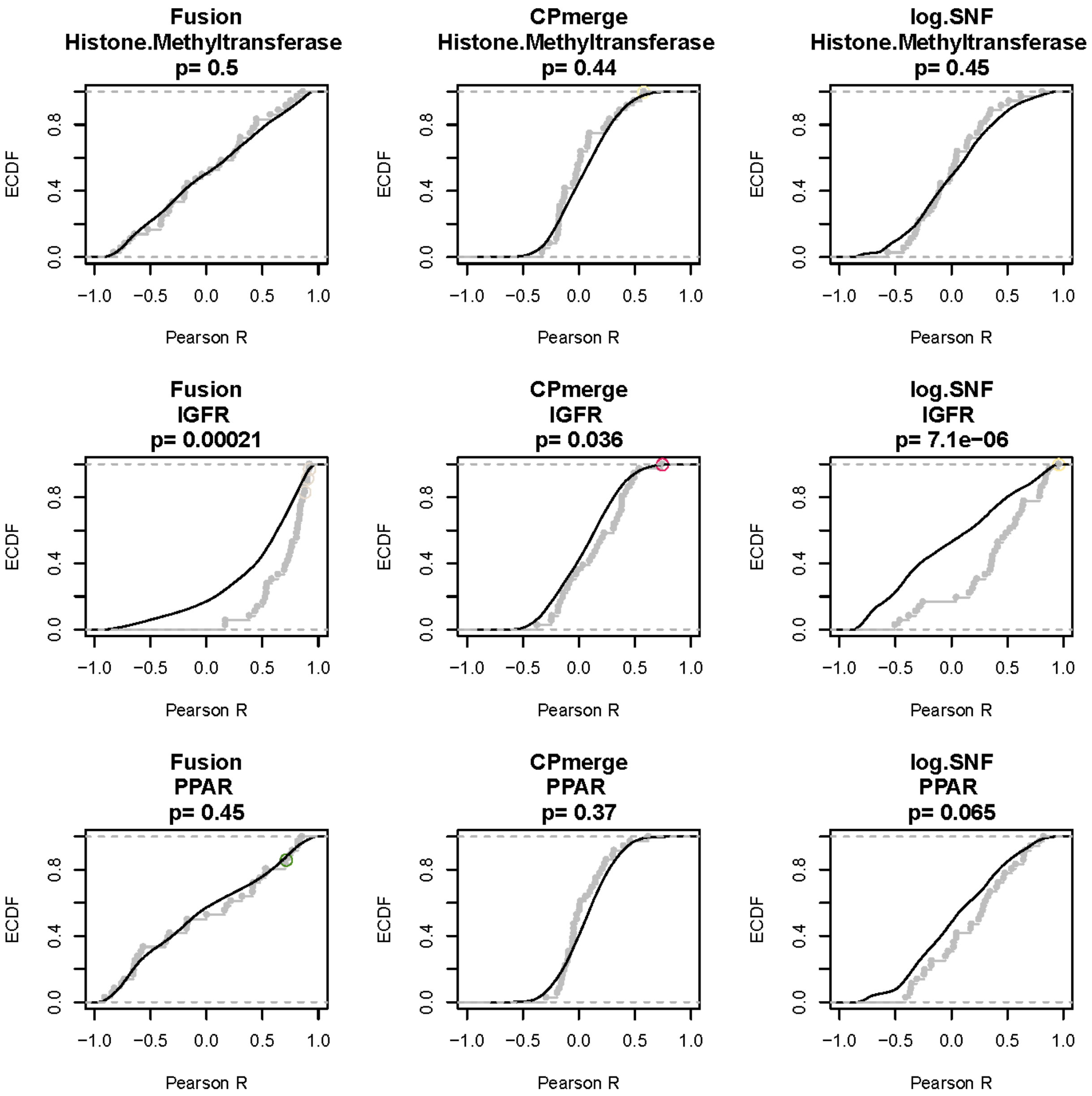
Fusion
Histone.Methyltransferase
p= 0.5
CPmerge
Histone.Methyltransferase
p= 0.44
log.SNF
Histone.Methyltransferase
p= 0.45
Fusion
IGFR
p= 0.00021
CPmerge
IGFR
p= 0.036
log.SNF
IGFR
p= 7.1e−06
Fusion
PPAR
p= 0.45
CPmerge
PPAR
p= 0.37
log.SNF
PPAR
p= 0.065
ECDF
Pearson R

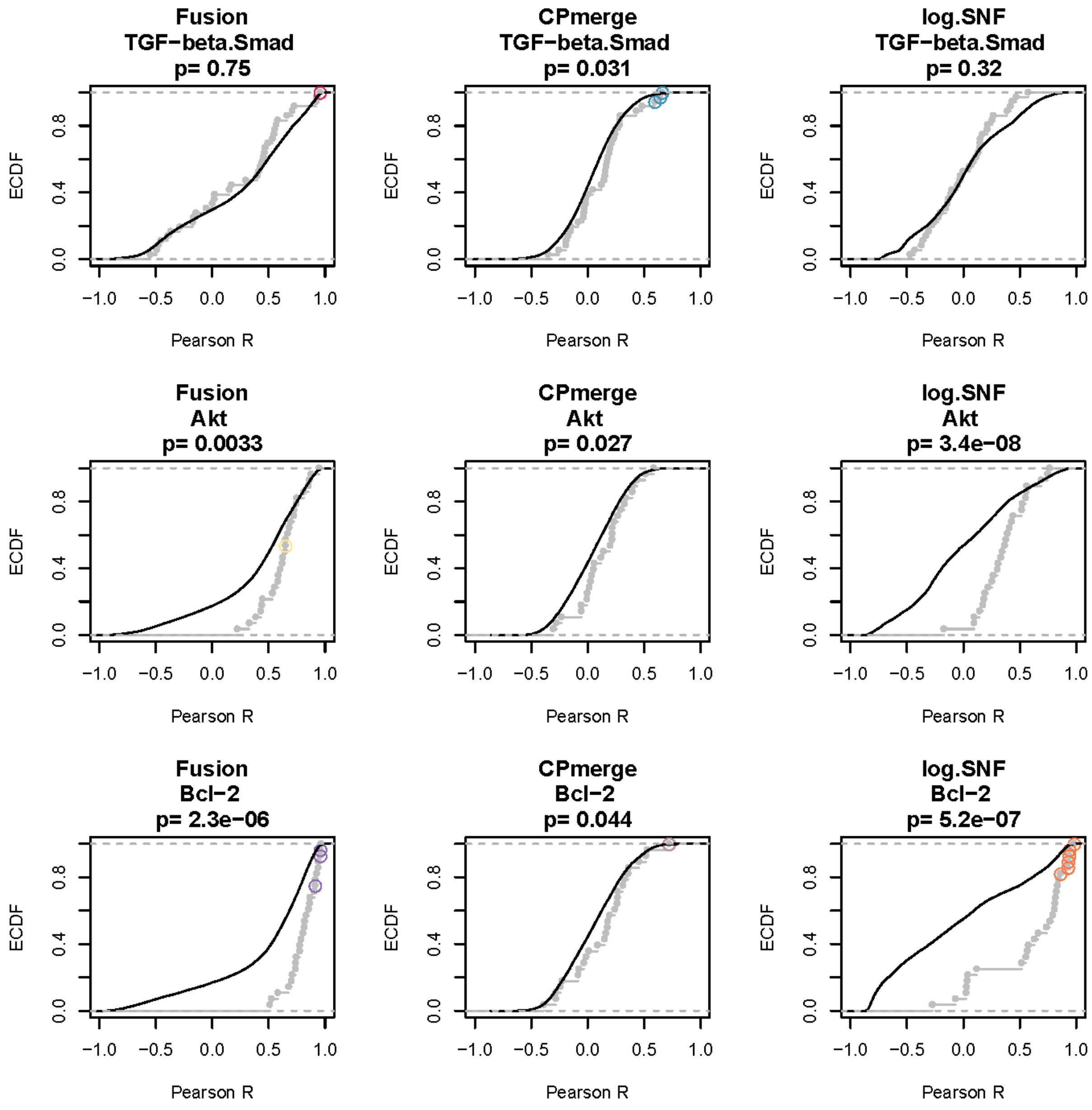
Fusion
TGF−beta.Smad
p= 0.75
CPmerge
TGF−beta.Smad
p= 0.031
log.SNF
TGF−beta.Smad
p= 0.32
Fusion
Akt
p= 0.0033
CPmerge
Akt
p= 0.027
log.SNF
Akt
p= 3.4e−08
Fusion
Bcl−2
p= 2.3e−06
CPmerge
Bcl−2
p= 0.044
log.SNF
Bcl−2
p= 5.2e−07
ECDF
Pearson R
0.0
0.4
0.8
−1.0
−0.5
0.0
0.5
1.0

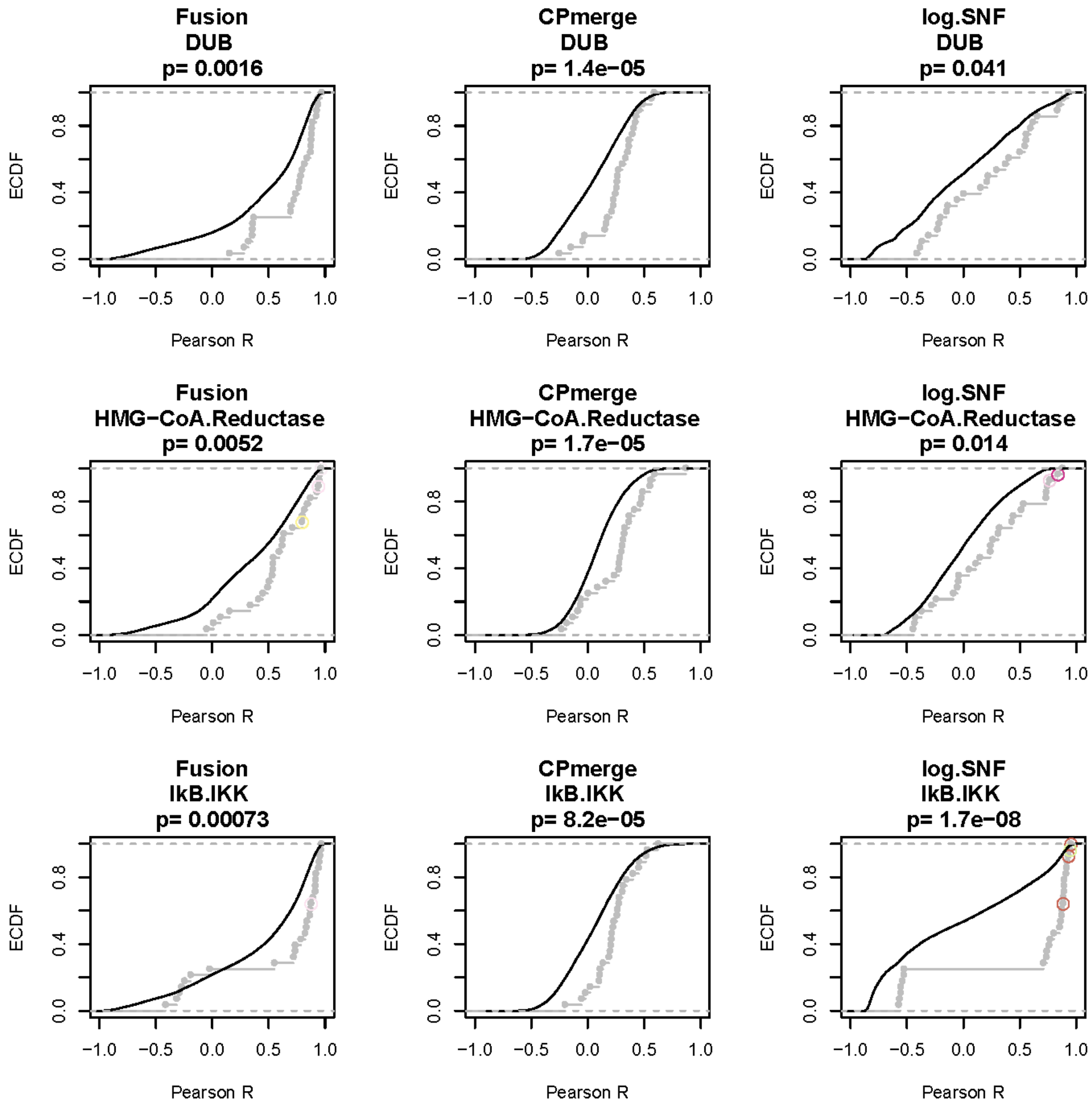

Fusion
DUB
p= 0.0016
CPmerge
DUB
p= 1.4e−05
log.SNF
DUB
p= 0.041
Fusion
HMG−CoA.Reductase
p= 0.0052
CPmerge
HMG−CoA.Reductase
p= 1.7e−05
log.SNF
HMG−CoA.Reductase
p= 0.014
Fusion
IkB.IKK
p= 0.00073
CPmerge
IkB.IKK
p= 8.2e−05
log.SNF
IkB.IKK
p= 1.7e−08
ECDF
0.0
0.4
0.8
−1.0
−0.5
0.0
0.5
1.0
Pearson R

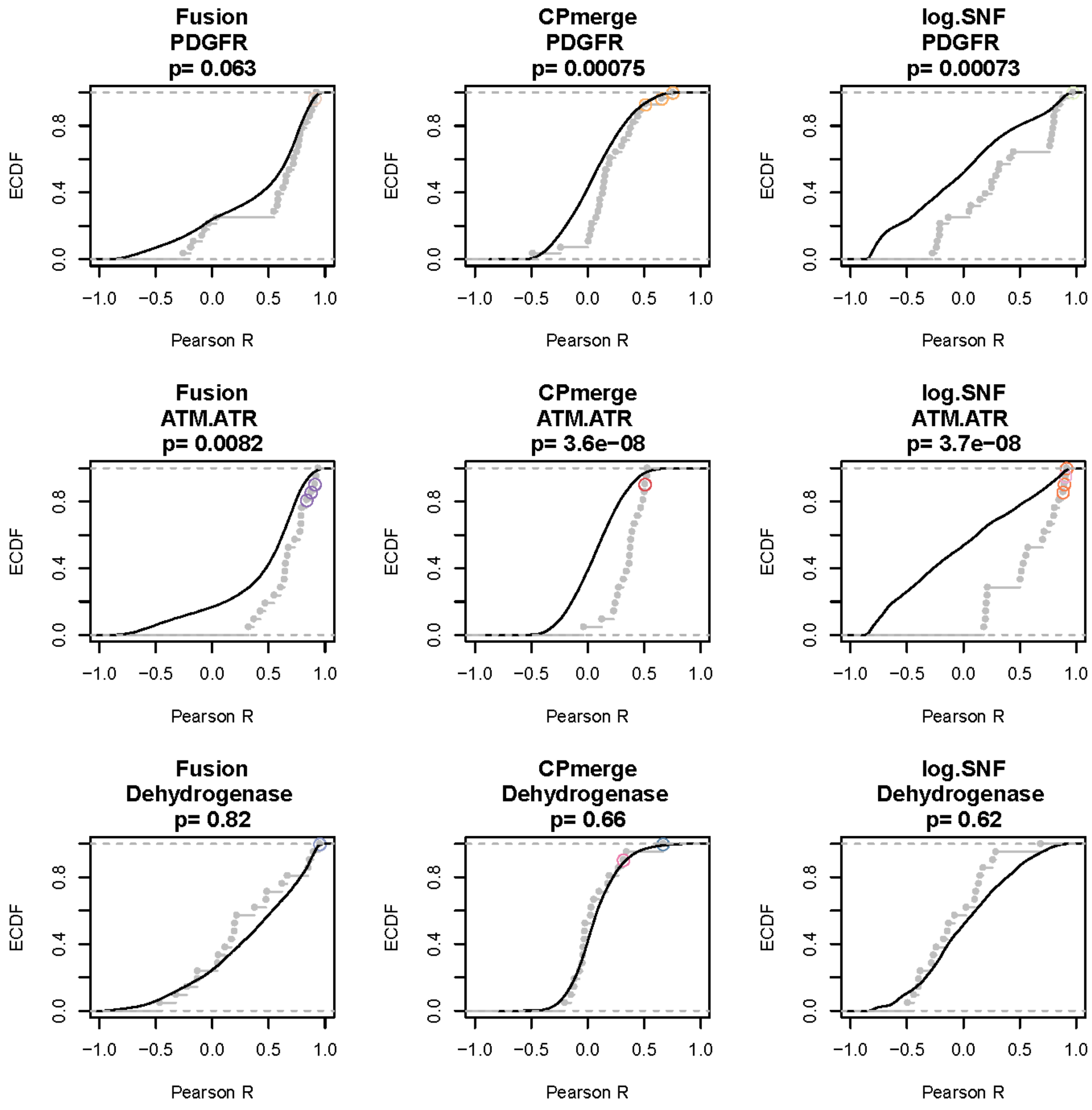

Fusion
PDGFR
p= 0.063
CPmerge
PDGFR
p= 0.00075
log.SNF
PDGFR
p= 0.00073
Fusion
ATM.ATR
p= 0.0082
CPmerge
ATM.ATR
p= 3.6e−08
log.SNF
ATM.ATR
p= 3.7e−08
Fusion
Dehydrogenase
p= 0.82
CPmerge
Dehydrogenase
p= 0.66
log.SNF
Dehydrogenase
p= 0.62
ECDF
Pearson R

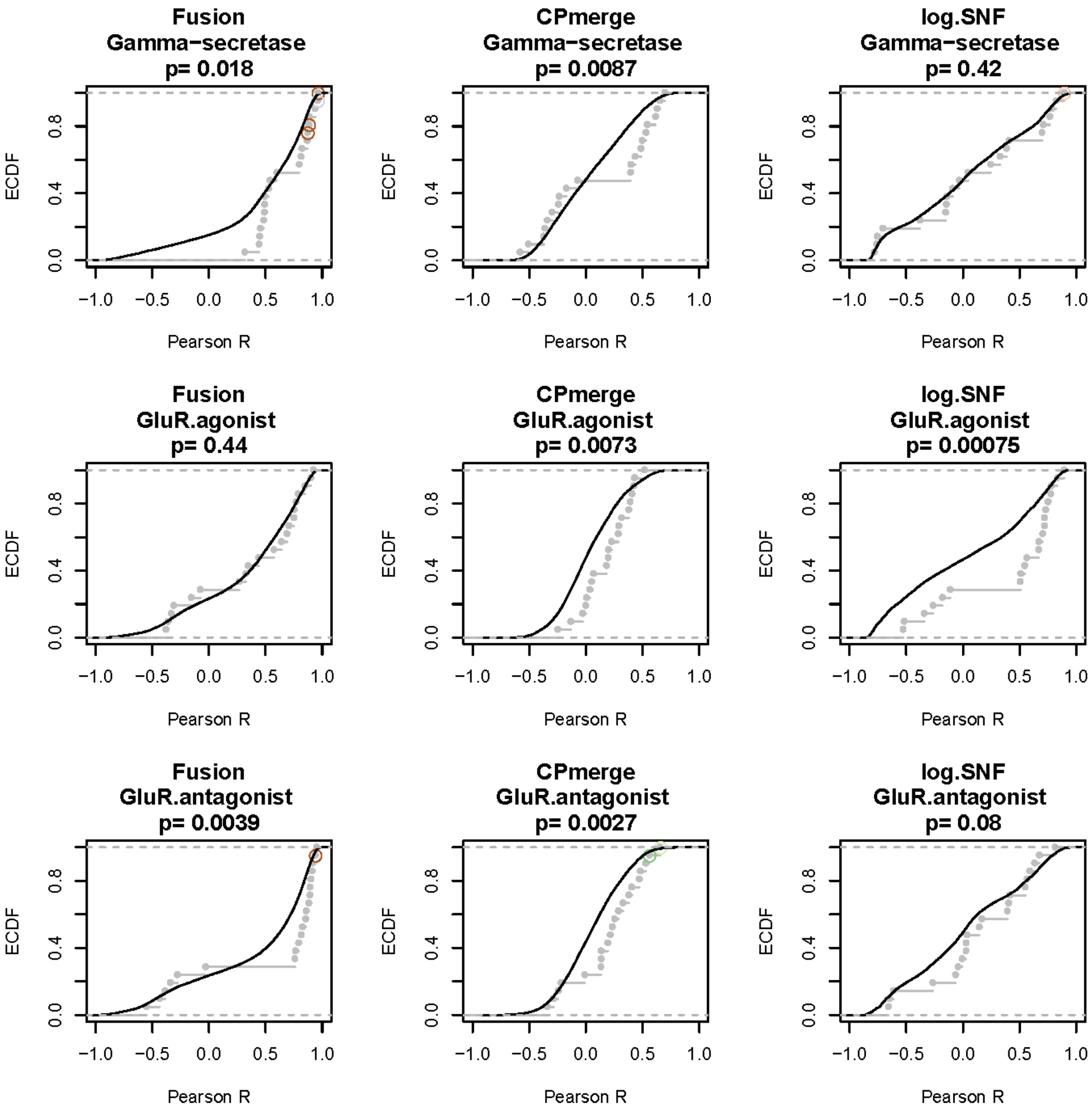
Fusion
Gamma−secretase
p= 0.018
CPmerge
Gamma−secretase
p= 0.0087
log.SNF
Gamma−secretase
p= 0.42
Fusion
GluR.agonist
p= 0.44
CPmerge
GluR.agonist
p= 0.0073
log.SNF
GluR.agonist
p= 0.00075
Fusion
GluR.antagonist
p= 0.0039
CPmerge
GluR.antagonist
p= 0.0027
log.SNF
GluR.antagonist
p= 0.08
ECDF
Pearson R
0.0
0.4
0.8
−1.0
−0.5
0.0
0.5
1.0

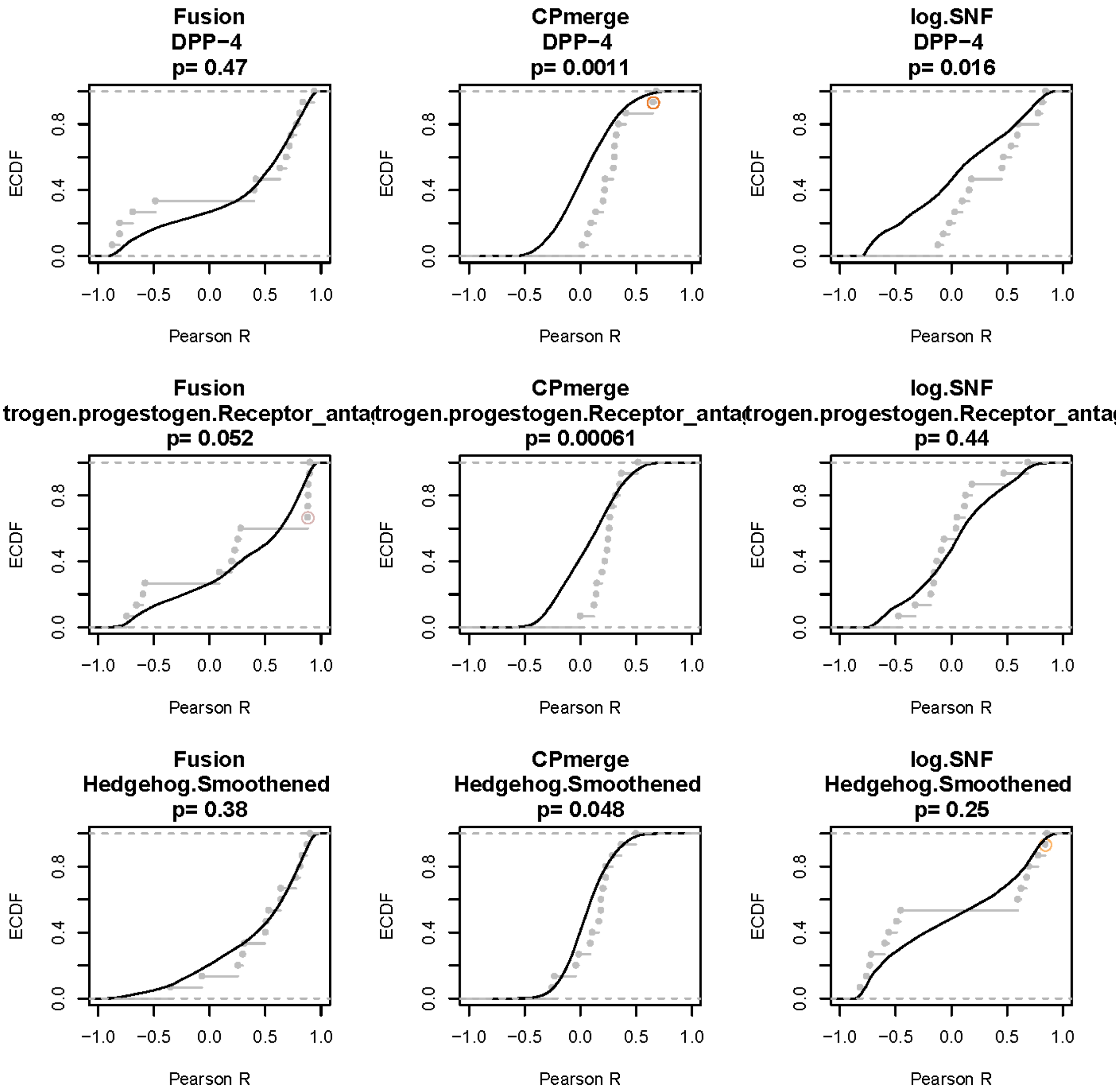
Fusion
DPP−4
p= 0.47
CPmerge
DPP−4
p= 0.0011
log.SNF
DPP−4
p= 0.016
Fusion
trogen.progestogen.Receptor_anta
p= 0.052
CPmerge
trogen.progestogen.Receptor_anta
p= 0.00061
log.SNF
trogen.progestogen.Receptor_anta
p= 0.44
Fusion
Hedgehog.Smoothened
p= 0.38
CPmerge
Hedgehog.Smoothened
p= 0.048
log.SNF
Hedgehog.Smoothened
p= 0.25
ECDF
Pearson R

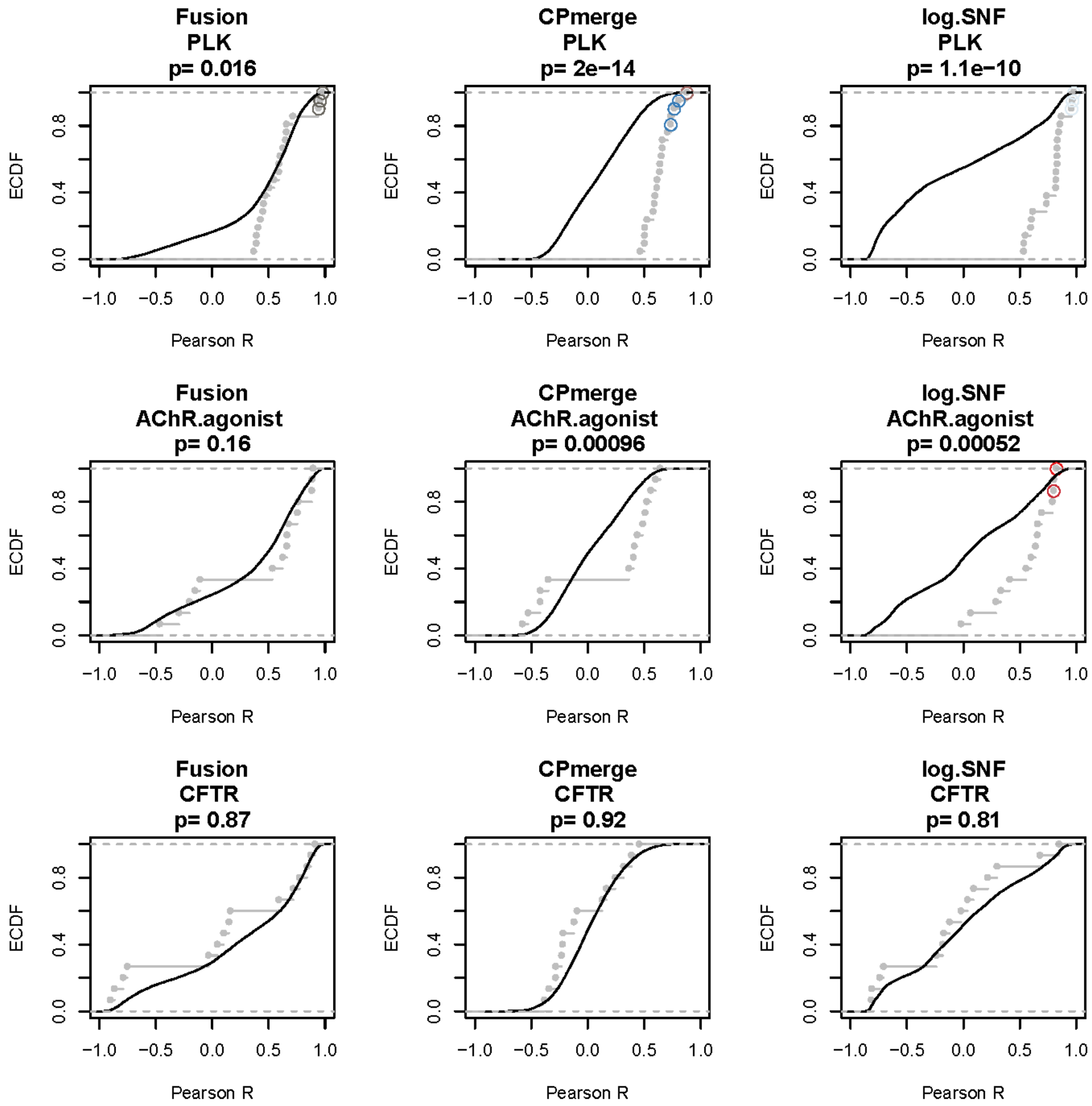
Fusion
PLK
p= 0.016
CPmerge
PLK
p= 2e−14
log.SNF
PLK
p= 1.1e−10
Fusion
AChR.agonist
p= 0.16
CPmerge
AChR.agonist
p= 0.00096
log.SNF
AChR.agonist
p= 0.00052
Fusion
CFTR
p= 0.87
CPmerge
CFTR
p= 0.92
log.SNF
CFTR
p= 0.81
ECDF
Pearson R
0.0
0.4
0.8
−1.0
−0.5
0.0
0.5
1.0

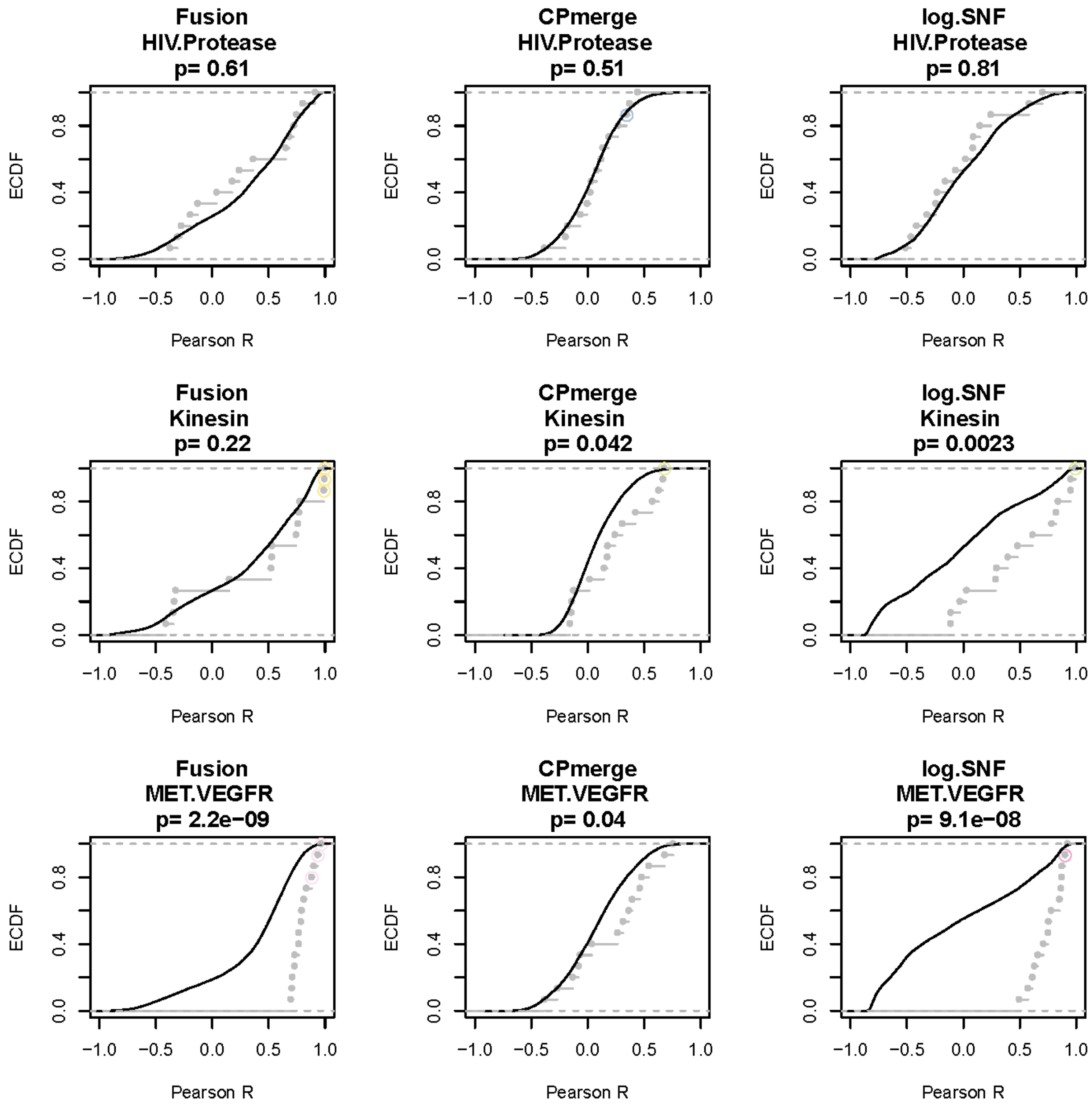
Fusion
HIV.Protease
p= 0.61
CPmerge
HIV.Protease
p= 0.51
log.SNF
HIV.Protease
p= 0.81
Fusion
Kinesin
p= 0.22
CPmerge
Kinesin
p= 0.042
log.SNF
Kinesin
p= 0.0023
Fusion
MET.VEGFR
p= 2.2e−09
CPmerge
MET.VEGFR
p= 0.04
log.SNF
MET.VEGFR
p= 9.1e−08
ECDF
Pearson R
0.0
0.4
0.8
−1.0
−0.5
0.0
0.5
1.0

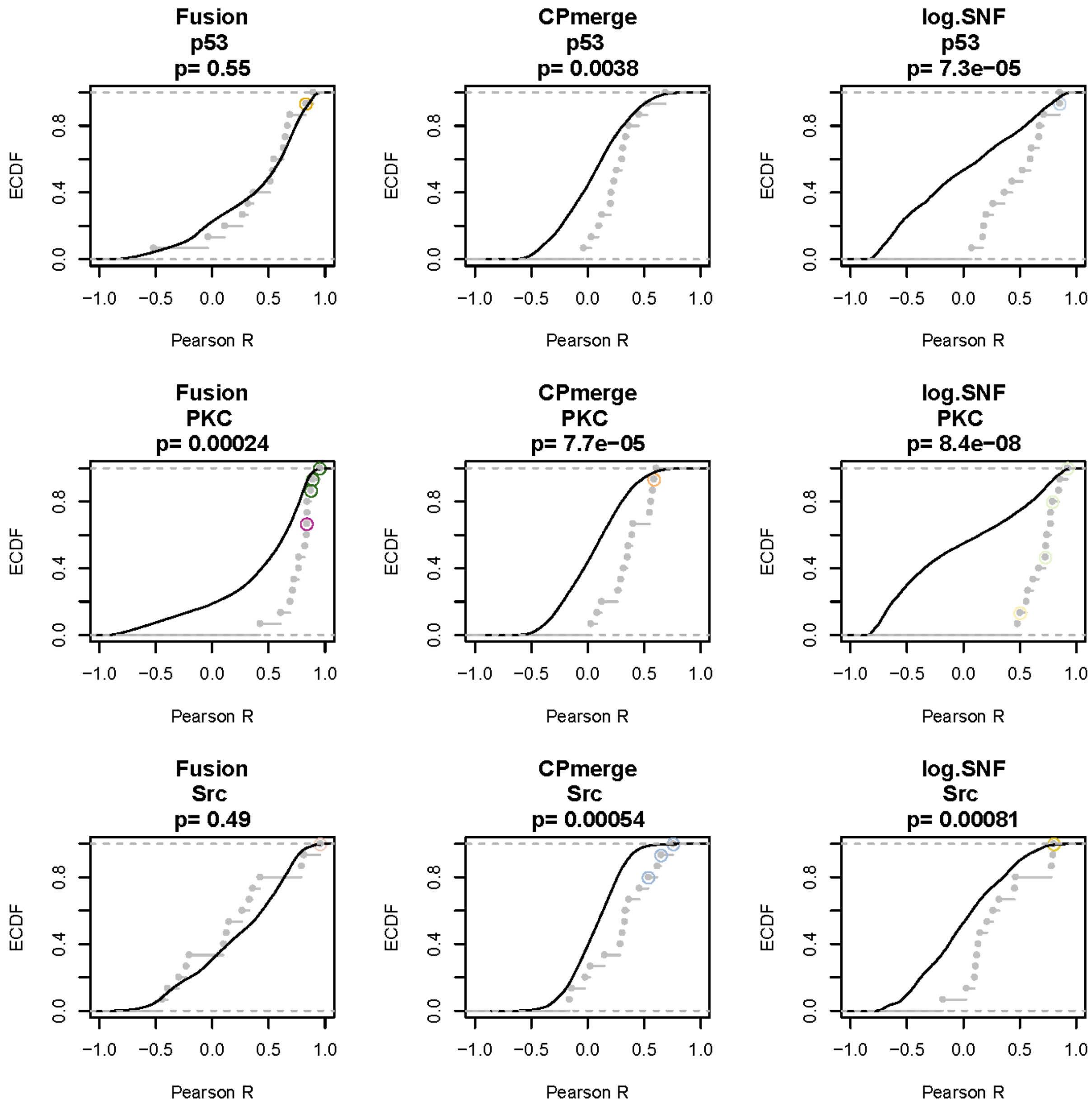

Fusion
p53
p= 0.55
CPmerge
p53
p= 0.0038
log.SNF
p53
p= 7.3e−05
Fusion
PKC
p= 0.00024
CPmerge
PKC
p= 7.7e−05
log.SNF
PKC
p= 8.4e−08
Fusion
Src
p= 0.49
CPmerge
Src
p= 0.00054
log.SNF
Src
p= 0.00081
ECDF
0.0
0.4
0.8
−1.0
−0.5
0.0
0.5
1.0
Pearson R

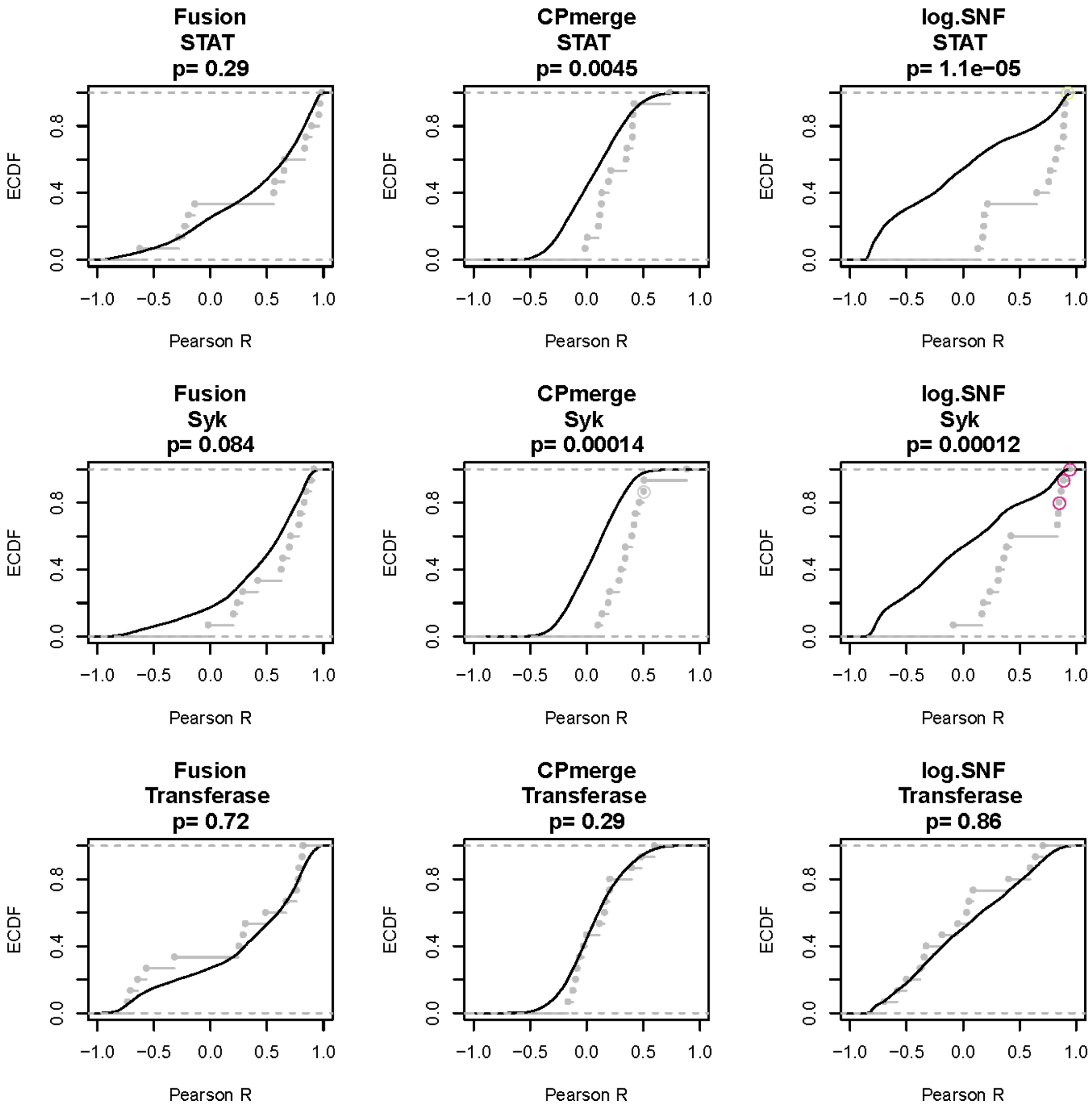
Fusion
STAT
p= 0.29
CPmerge
STAT
p= 0.0045
log.SNF
STAT
p= 1.1e−05
Fusion
Syk
p= 0.084
CPmerge
Syk
p= 0.00014
log.SNF
Syk
p= 0.00012
Fusion
Transferase
p= 0.72
CPmerge
Transferase
p= 0.29
log.SNF
Transferase
p= 0.86
ECDF
Pearson R
0.0
0.4
0.8
−1.0
−0.5
0.0
0.5
1.0

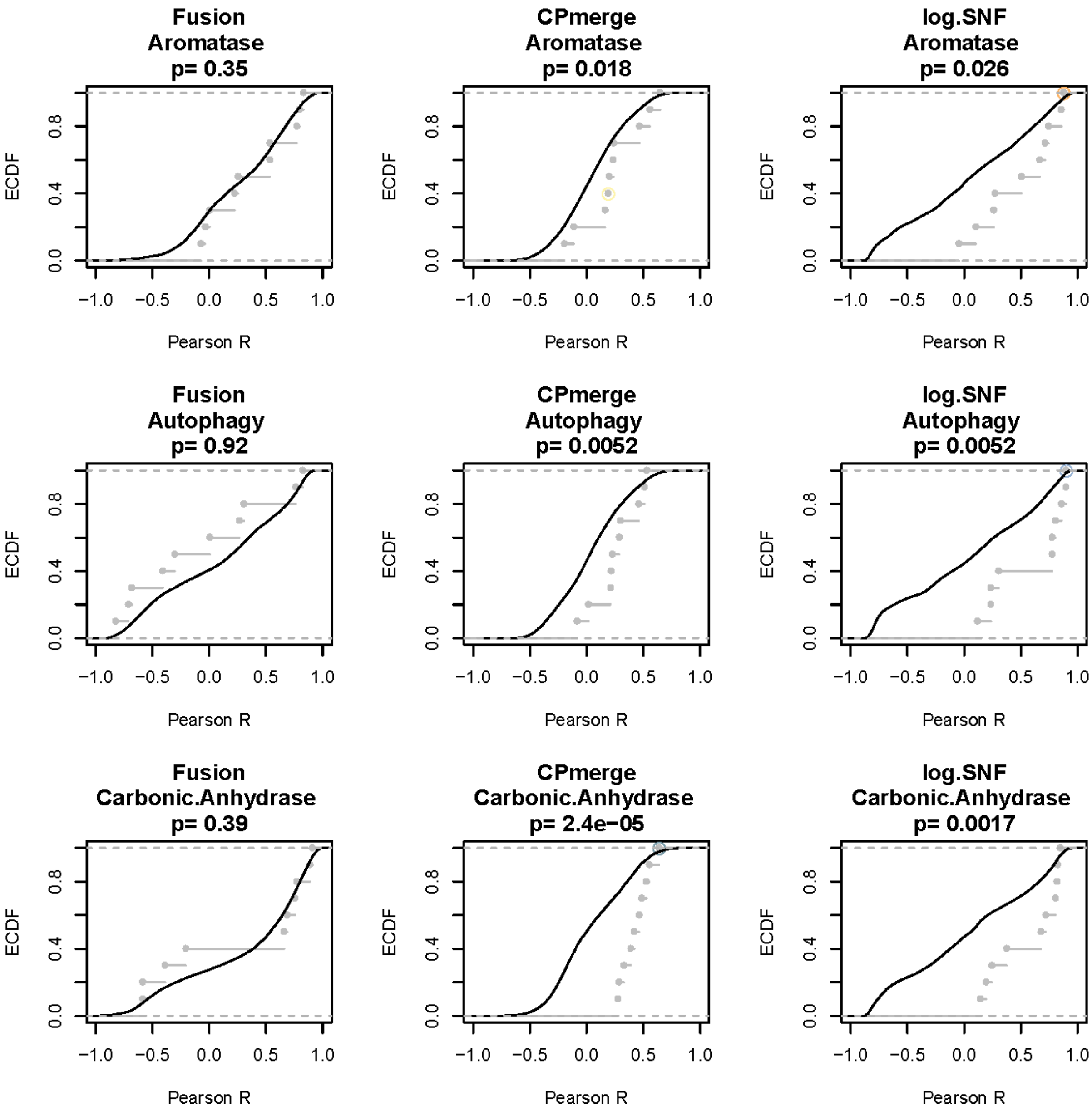
Fusion
Aromatase
p= 0.35
CPmerge
Aromatase
p= 0.018
log.SNF
Aromatase
p= 0.026
Fusion
Autophagy
p= 0.92
CPmerge
Autophagy
p= 0.0052
log.SNF
Autophagy
p= 0.0052
Fusion
Carbonic.Anhydrase
p= 0.39
CPmerge
Carbonic.Anhydrase
p= 2.4e−05
log.SNF
Carbonic.Anhydrase
p= 0.0017
ECDF
0.0
0.4
0.8
−1.0
−0.5
0.0
0.5
1.0
Pearson R

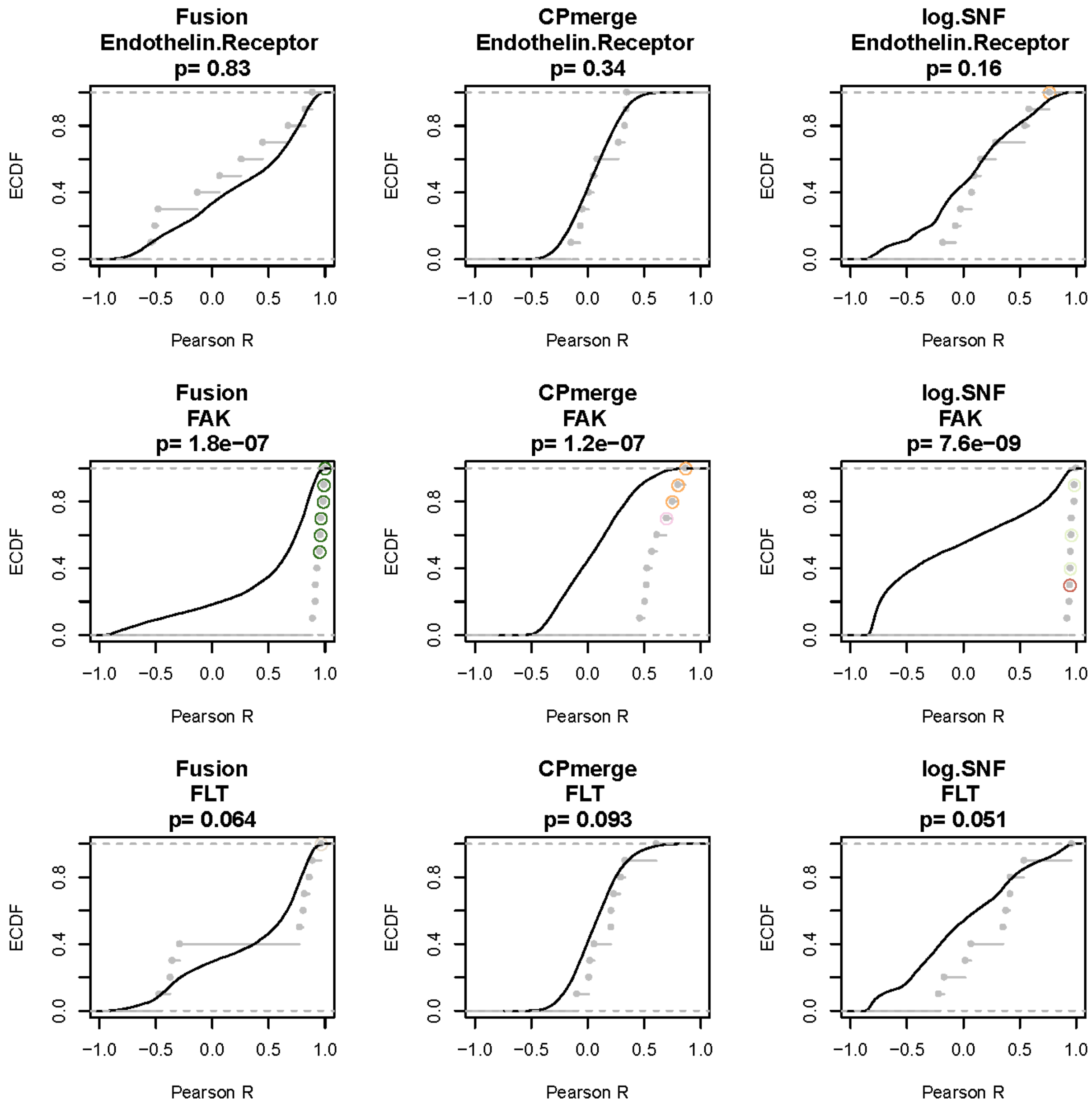
Fusion
Endothelin.Receptor
p= 0.83
CPmerge
Endothelin.Receptor
p= 0.34
log.SNF
Endothelin.Receptor
p= 0.16
Fusion
FAK
p= 1.8e−07
CPmerge
FAK
p= 1.2e−07
log.SNF
FAK
p= 7.6e−09
Fusion
FLT
p= 0.064
CPmerge
FLT
p= 0.093
log.SNF
FLT
p= 0.051
ECDF
Pearson R

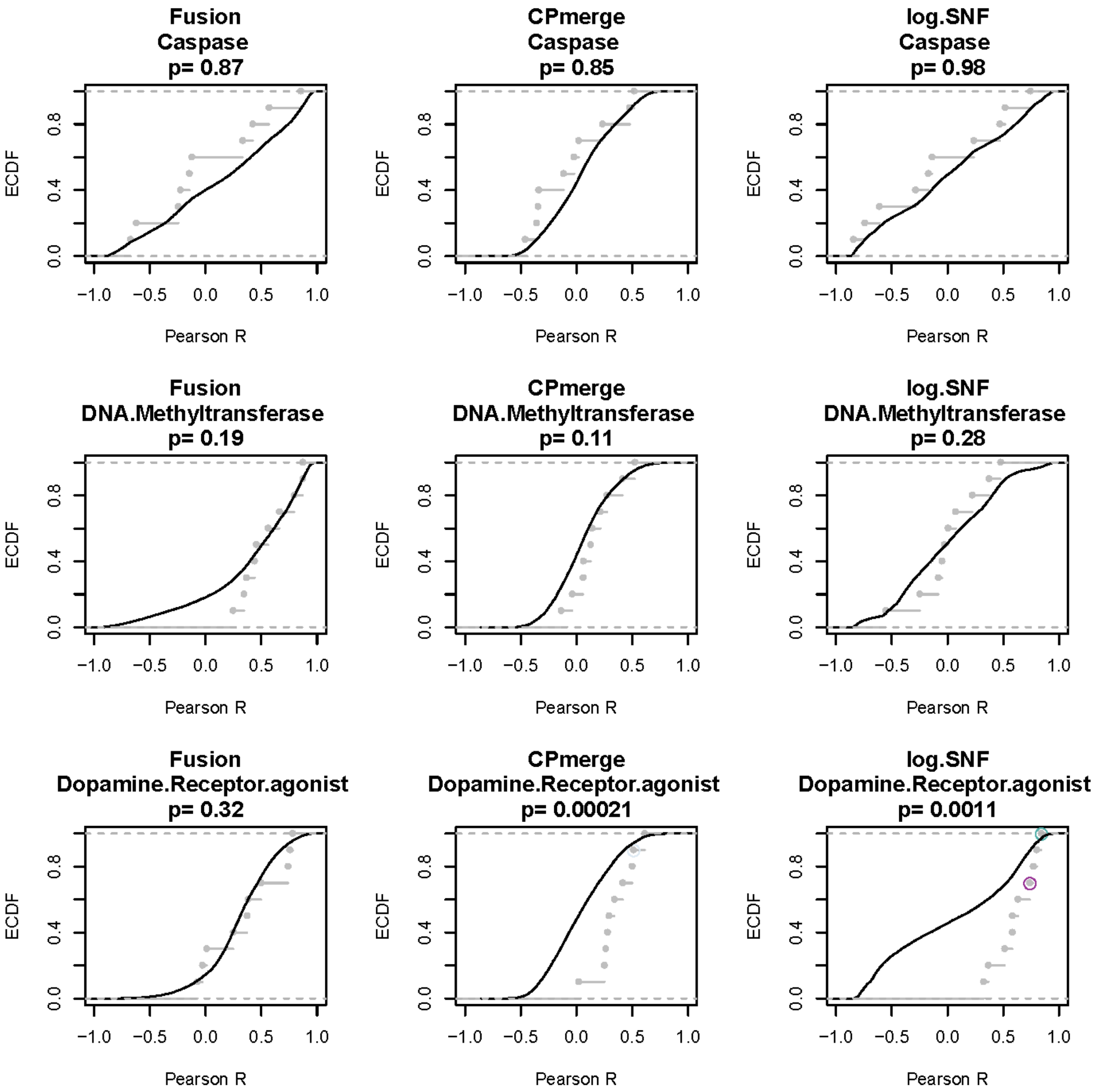
Fusion
Caspase
p= 0.87
CPmerge
Caspase
p= 0.85
log.SNF
Caspase
p= 0.98
Fusion
DNA.Methyltransferase
p= 0.19
CPmerge
DNA.Methyltransferase
p= 0.11
log.SNF
DNA.Methyltransferase
p= 0.28
Fusion
Dopamine.Receptor.agonist
p= 0.32
CPmerge
Dopamine.Receptor.agonist
p= 0.00021
log.SNF
Dopamine.Receptor.agonist
p= 0.0011
ECDF
Pearson R

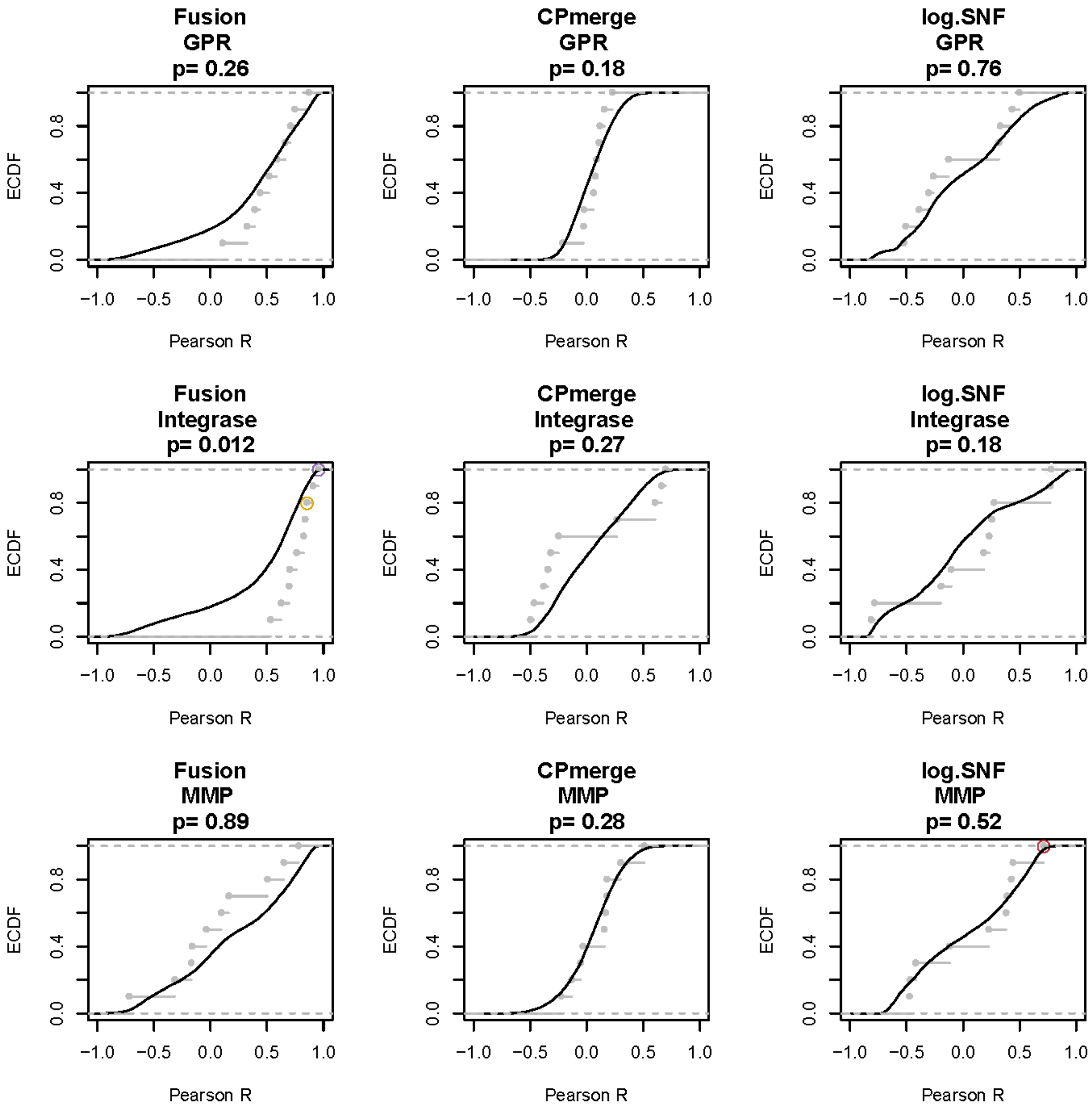
Fusion
GPR
p= 0.26
CPmerge
GPR
p= 0.18
log.SNF
GPR
p= 0.76
Fusion
Integrase
p= 0.012
CPmerge
Integrase
p= 0.27
log.SNF
Integrase
p= 0.18
Fusion
MMP
p= 0.89
CPmerge
MMP
p= 0.28
log.SNF
MMP
p= 0.52
ECDF
Pearson R
0.0
0.4
0.8
−1.0
−0.5
0.0
0.5
1.0

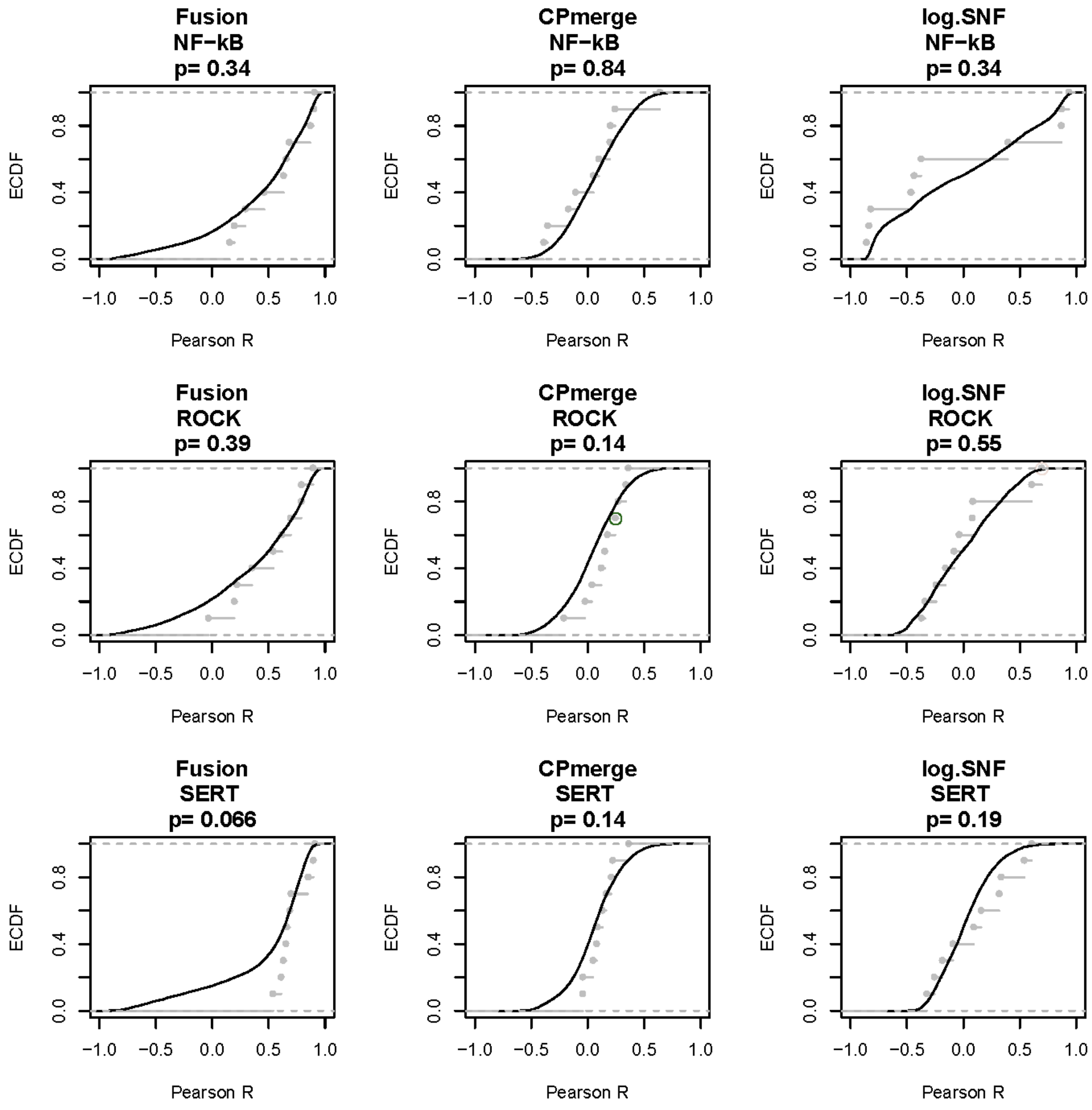
Fusion
NF−kB
p= 0.34
CPmerge
NF−kB
p= 0.84
log.SNF
NF−kB
p= 0.34
Fusion
ROCK
p= 0.39
CPmerge
ROCK
p= 0.14
log.SNF
ROCK
p= 0.55
Fusion
SERT
p= 0.066
CPmerge
SERT
p= 0.14
log.SNF
SERT
p= 0.19
ECDF
Pearson R
0.0
0.4
0.8
−1.0
−0.5
0.0
0.5
1.0

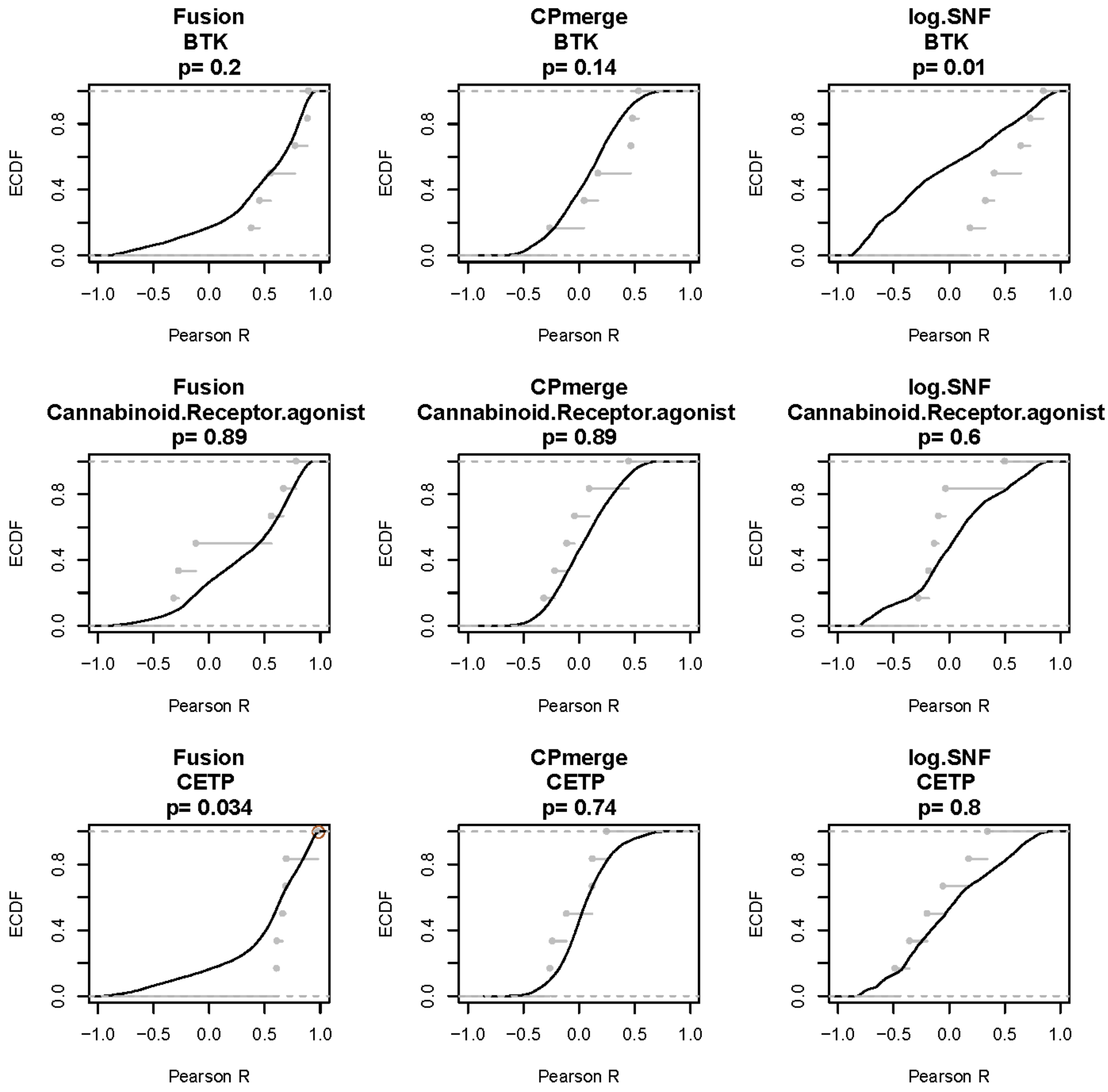
Fusion
BTK
p= 0.2
CPmerge
BTK
p= 0.14
log.SNF
BTK
p= 0.01
Fusion
Cannabinoid.Receptor.agonist
p= 0.89
CPmerge
Cannabinoid.Receptor.agonist
p= 0.89
log.SNF
Cannabinoid.Receptor.agonist
p= 0.6
Fusion
CETP
p= 0.034
CPmerge
CETP
p= 0.74
log.SNF
CETP
p= 0.8
ECDF
0.0
0.4
0.8
−1.0
−0.5
0.0
0.5
1.0
Pearson R

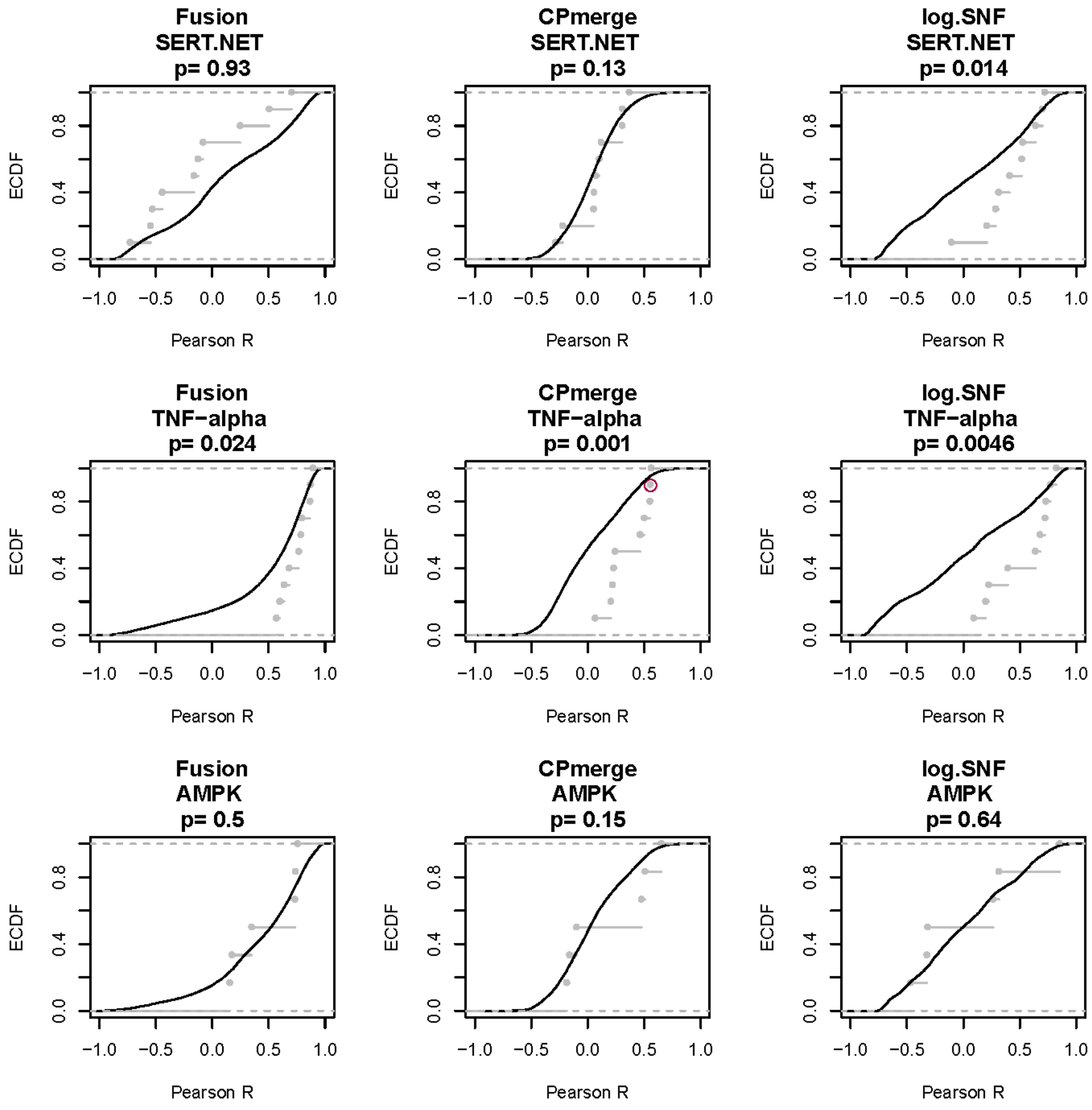

Fusion
SERT.NET
p= 0.93
CPmerge
SERT.NET
p= 0.13
log.SNF
SERT.NET
p= 0.014
Fusion
TNF−alpha
p= 0.024
CPmerge
TNF−alpha
p= 0.001
log.SNF
TNF−alpha
p= 0.0046
Fusion
AMPK
p= 0.5
CPmerge
AMPK
p= 0.15
log.SNF
AMPK
p= 0.64
ECDF
Pearson R
0.0
0.4
0.8
−1.0
−0.5
0.0
0.5
1.0

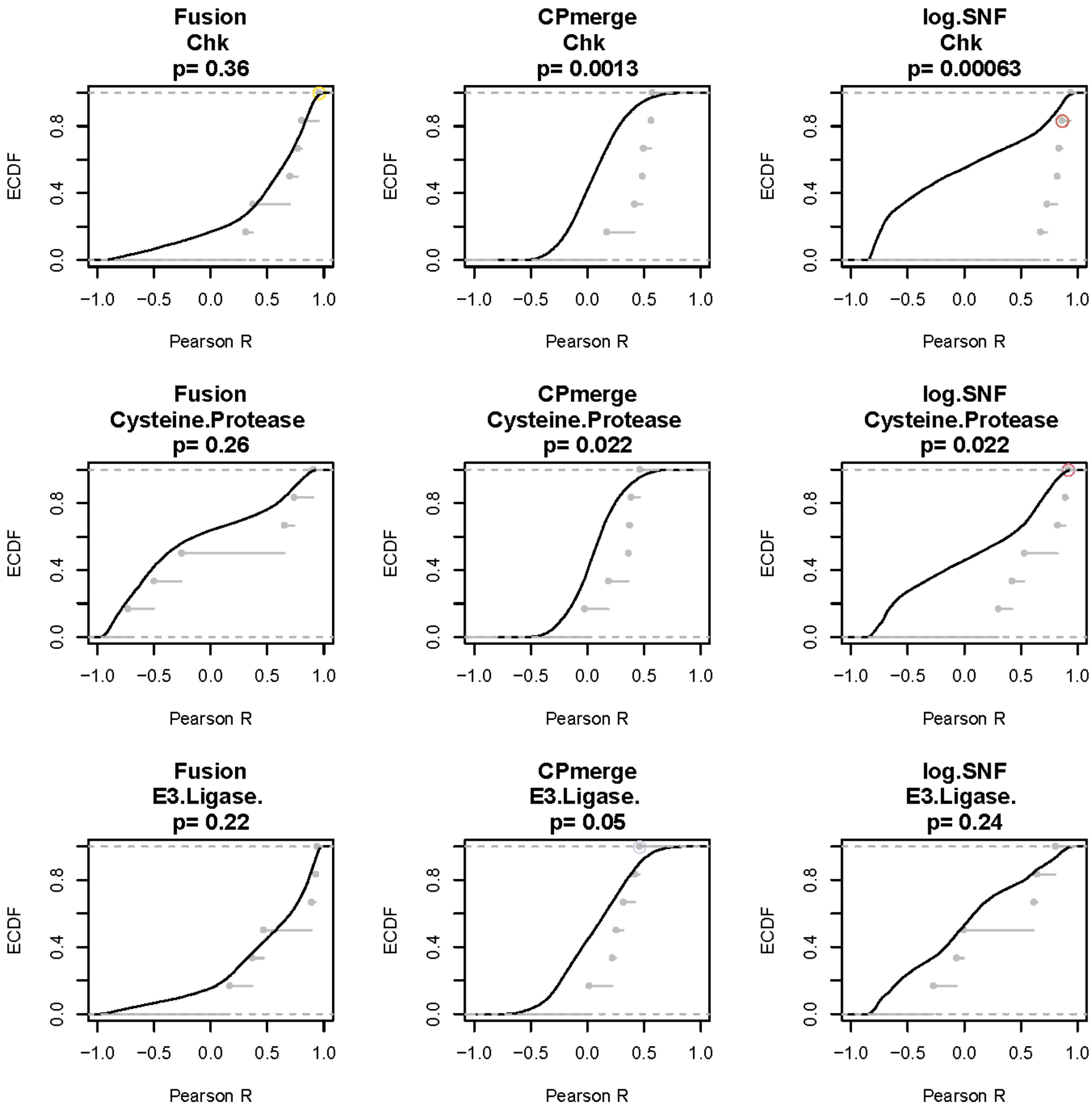
Fusion
Chk
p= 0.36
CPmerge
Chk
p= 0.0013
log.SNF
Chk
p= 0.00063
Fusion
Cysteine.Protease
p= 0.26
CPmerge
Cysteine.Protease
p= 0.022
log.SNF
Cysteine.Protease
p= 0.022
Fusion
E3.Ligase.
p= 0.22
CPmerge
E3.Ligase.
p= 0.05
log.SNF
E3.Ligase.
p= 0.24
ECDF
Pearson R

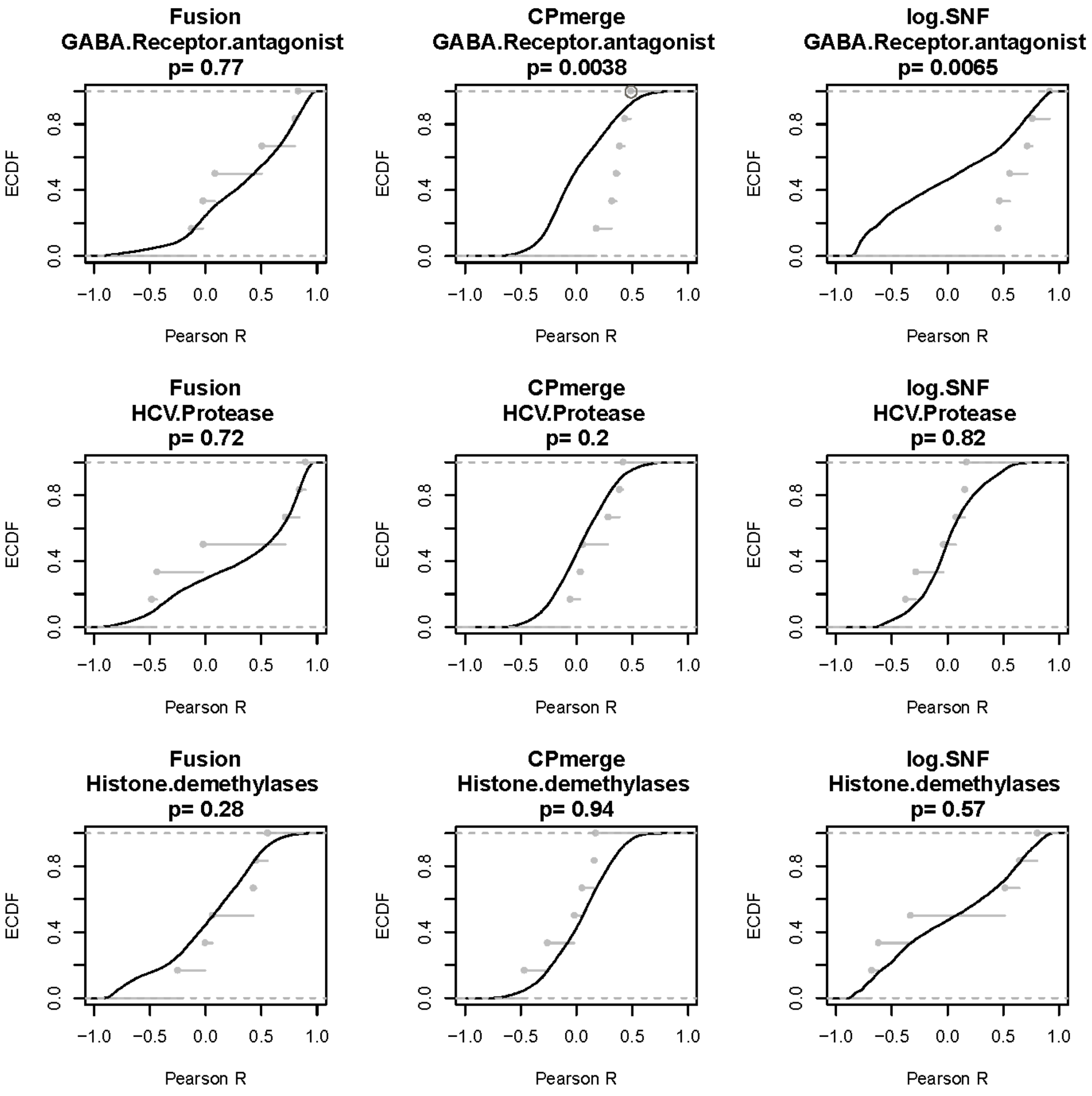
Fusion
GABA.Receptor.antagonist
p= 0.77
CPmerge
GABA.Receptor.antagonist
p= 0.0038
log.SNF
GABA.Receptor.antagonist
p= 0.0065
Fusion
HCV.Protease
p= 0.72
CPmerge
HCV.Protease
p= 0.2
log.SNF
HCV.Protease
p= 0.82
Fusion
Histone.demethylases
p= 0.28
CPmerge
Histone.demethylases
p= 0.94
log.SNF
Histone.demethylases
p= 0.57
ECDF
Pearson R
0.0
0.4
0.8
−1.0
−0.5
0.0
0.5
1.0

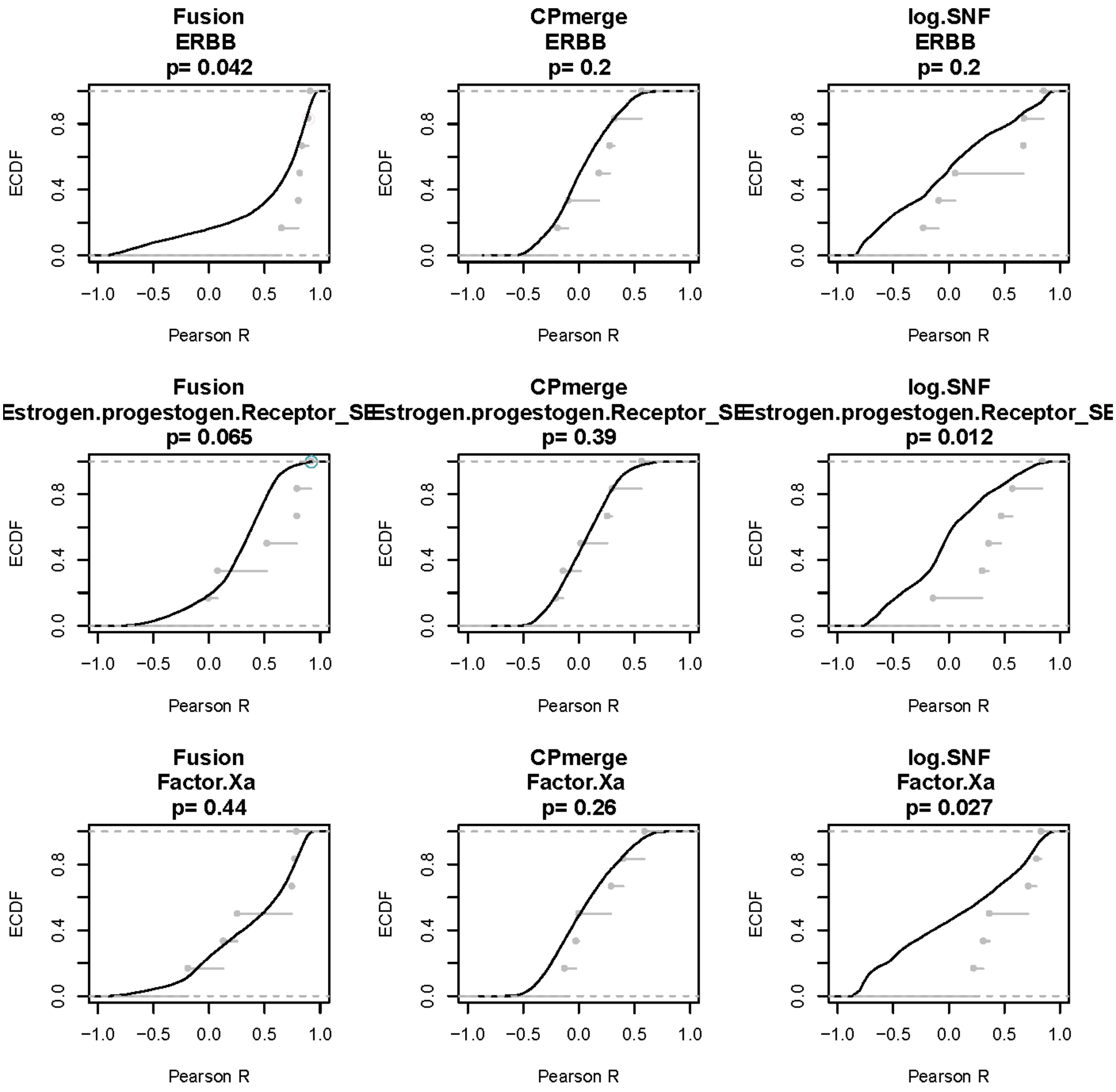
Fusion
ERBB
p= 0.042
CPmerge
ERBB
p= 0.2
log.SNF
ERBB
p= 0.2
Fusion
Estrogen.progestogen.Receptor_SE
p= 0.065
CPmerge
Estrogen.progestogen.Receptor_SE
p= 0.39
log.SNF
Estrogen.progestogen.Receptor_SE
p= 0.012
Fusion
Factor.Xa
p= 0.44
CPmerge
Factor.Xa
p= 0.26
log.SNF
Factor.Xa
p= 0.027
ECDF
0.0
0.4
0.8
−1.0
−0.5
0.0
0.5
1.0
Pearson R

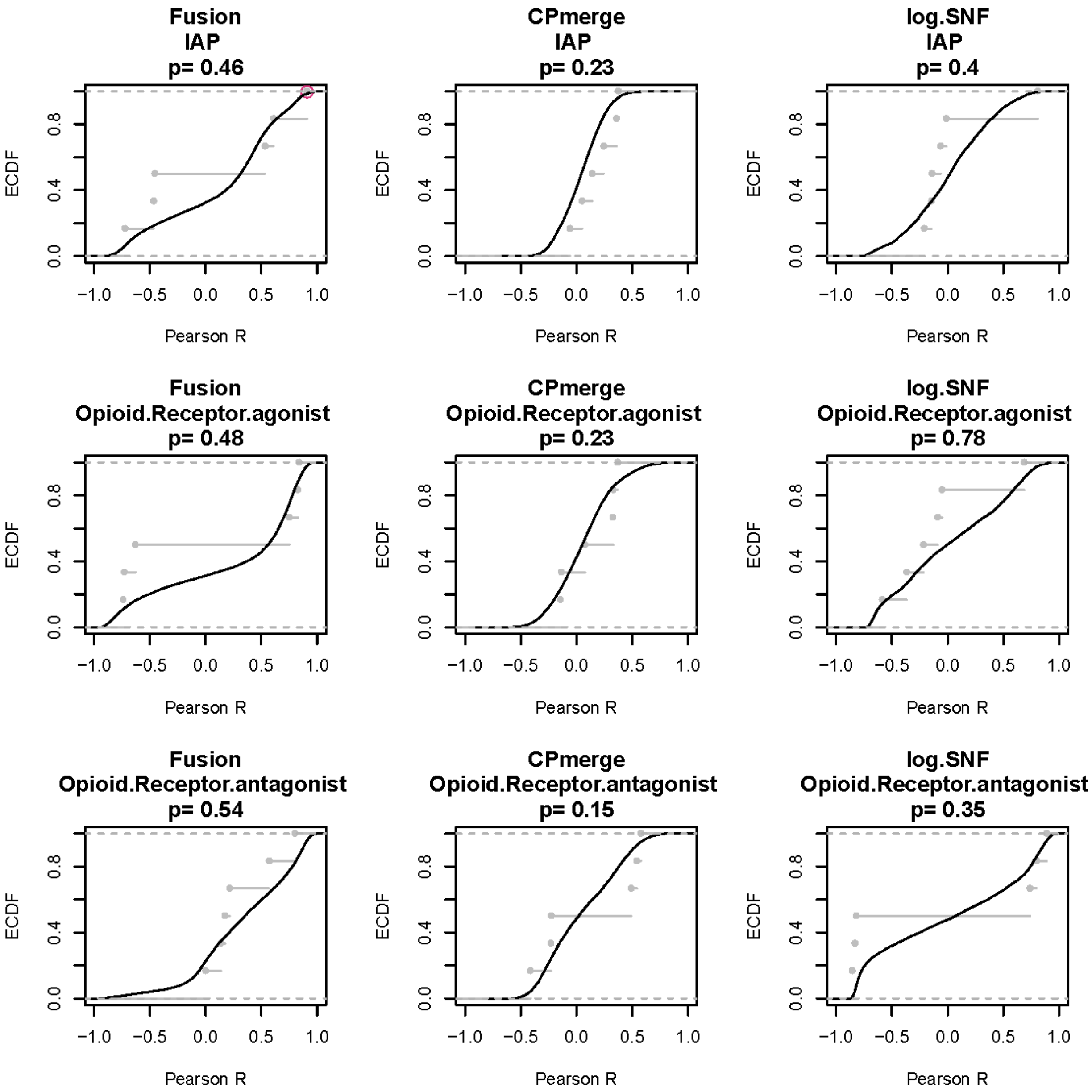

Fusion
IAP
p= 0.46
CPmerge
IAP
p= 0.23
log.SNF
IAP
p= 0.4
Fusion
Opioid.Receptor.agonist
p= 0.48
CPmerge
Opioid.Receptor.agonist
p= 0.23
log.SNF
Opioid.Receptor.agonist
p= 0.78
Fusion
Opioid.Receptor.antagonist
p= 0.54
CPmerge
Opioid.Receptor.antagonist
p= 0.15
log.SNF
Opioid.Receptor.antagonist
p= 0.35
ECDF
Pearson R

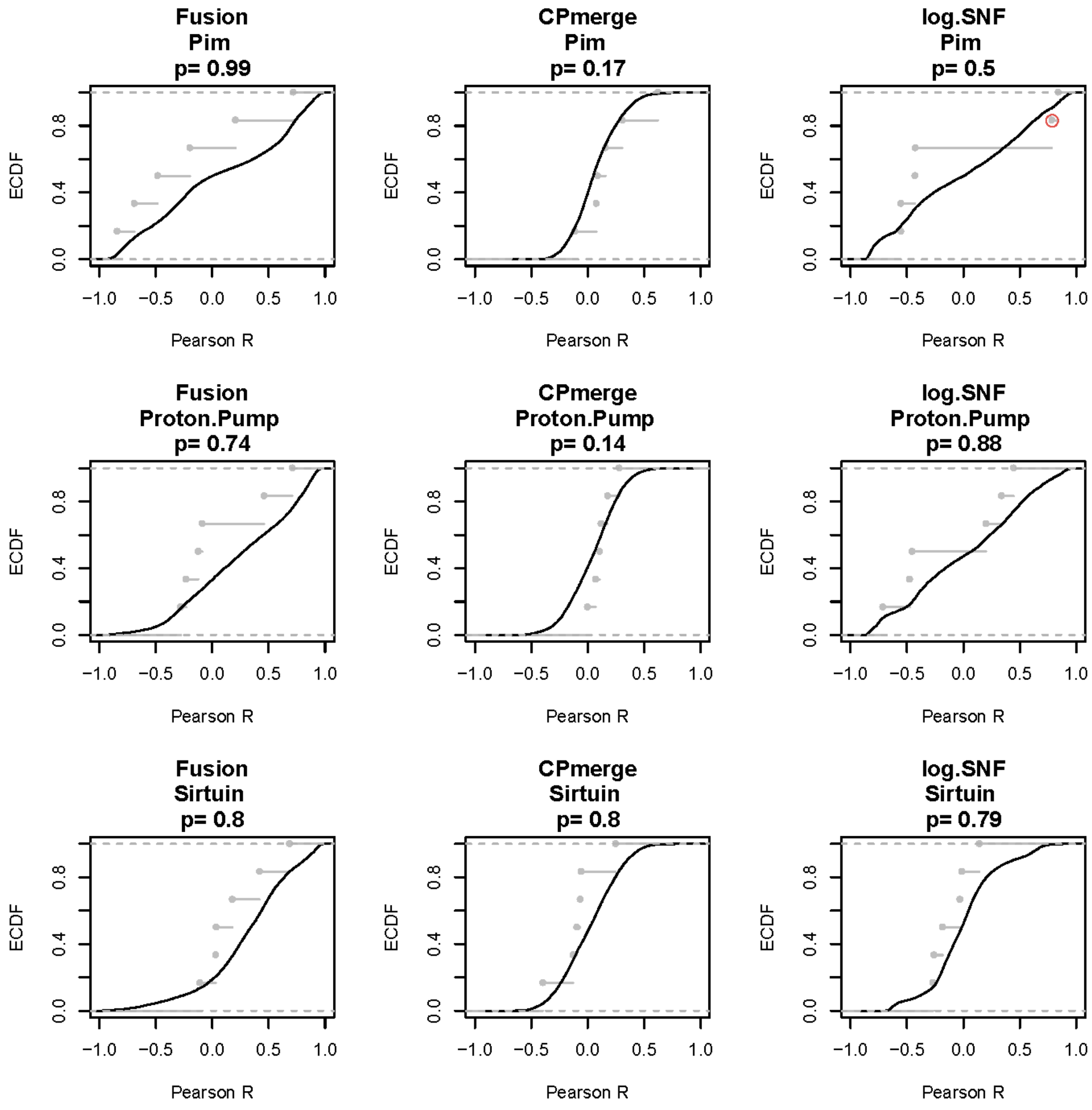
Fusion
Pim
p= 0.99
CPmerge
Pim
p= 0.17
log.SNF
Pim
p= 0.5
Fusion
Proton.Pump
p= 0.74
CPmerge
Proton.Pump
p= 0.14
log.SNF
Proton.Pump
p= 0.88
Fusion
Sirtuin
p= 0.8
CPmerge
Sirtuin
p= 0.8
log.SNF
Sirtuin
p= 0.79
ECDF
Pearson R
0.0
0.4
0.8
−1.0
−0.5
0.0
0.5
1.0

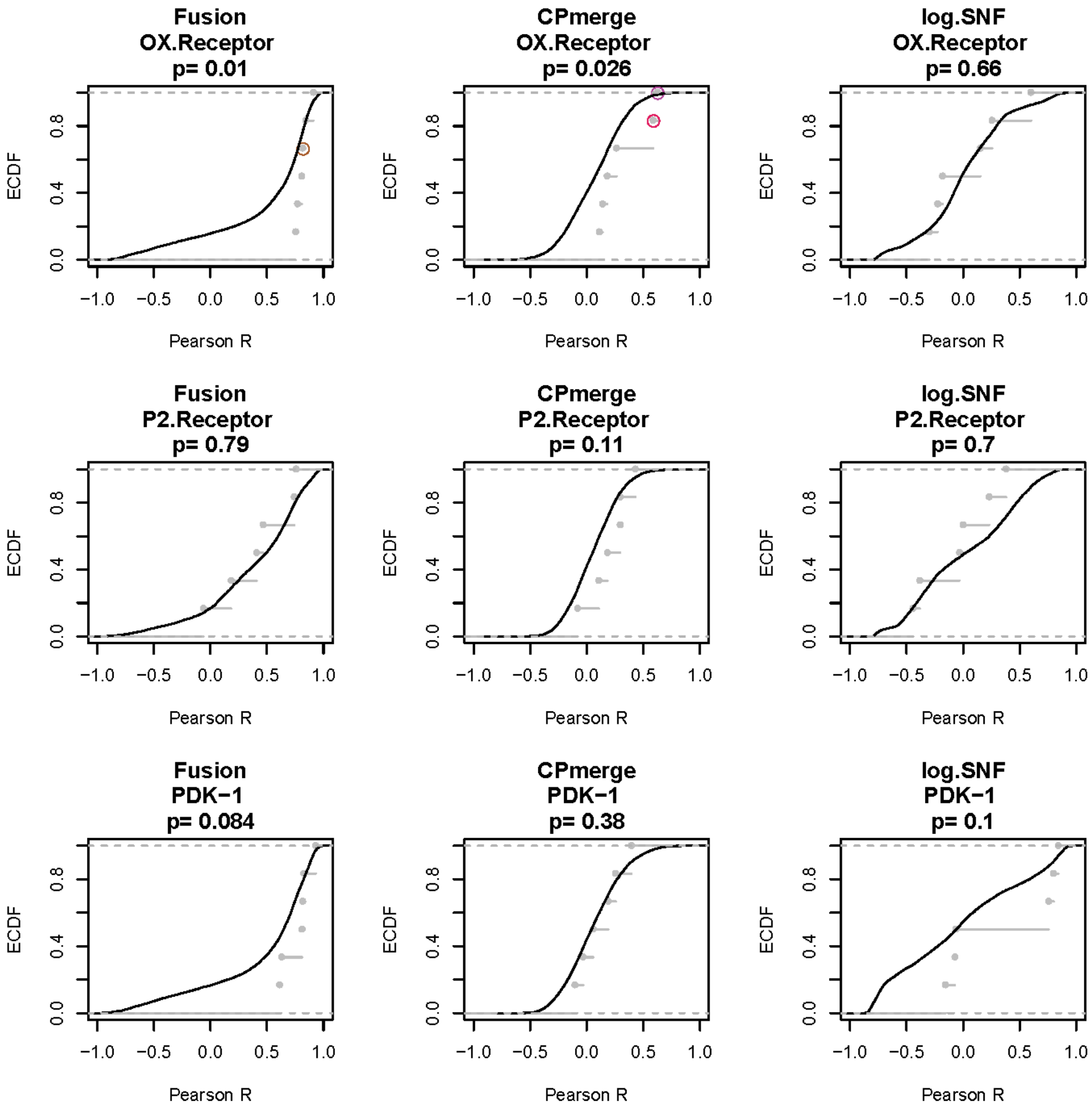
Fusion
OX.Receptor
p= 0.01
CPmerge
OX.Receptor
p= 0.026
log.SNF
OX.Receptor
p= 0.66
Fusion
P2.Receptor
p= 0.79
CPmerge
P2.Receptor
p= 0.11
log.SNF
P2.Receptor
p= 0.7
Fusion
PDK−1
p= 0.084
CPmerge
PDK−1
p= 0.38
log.SNF
PDK−1
p= 0.1
ECDF
0.0
0.4
0.8
−1.0
−0.5
0.0
0.5
1.0
Pearson R

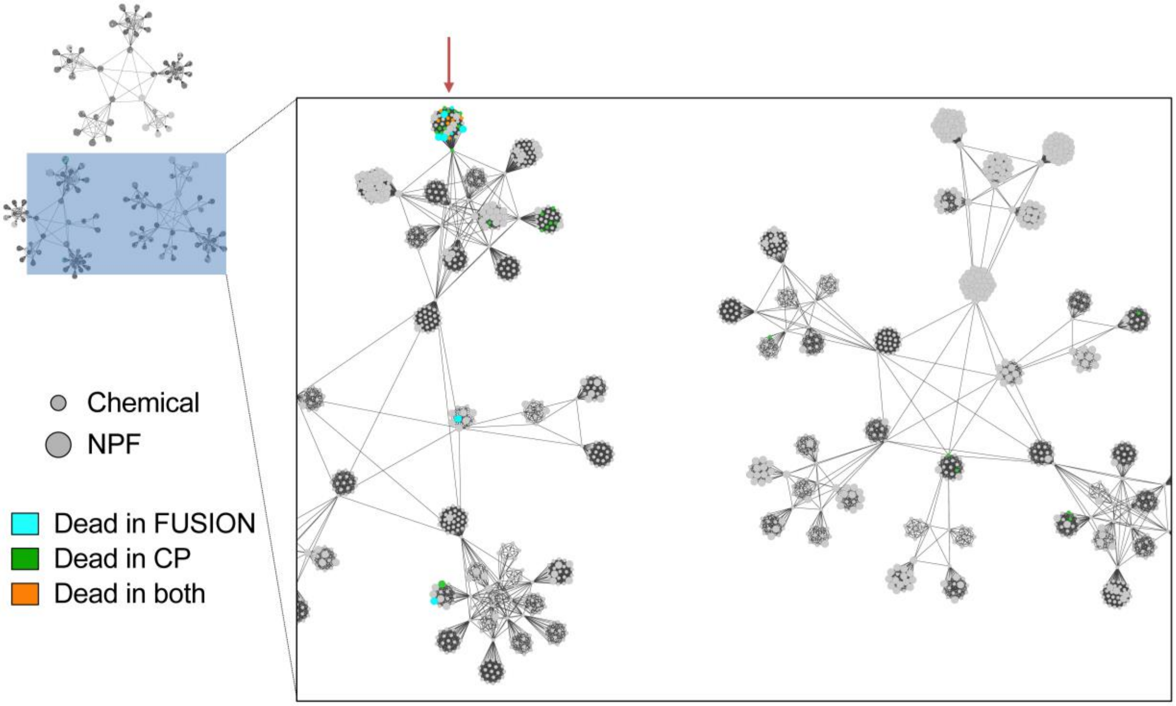

**Supplementary Figure 9: “Dead” Compounds in CP and FUSION cluster together in the SNF-Euclidean APC map.** Red arrow indicates a cluster enriched with compounds that were flagged as dead in either or both datasets.

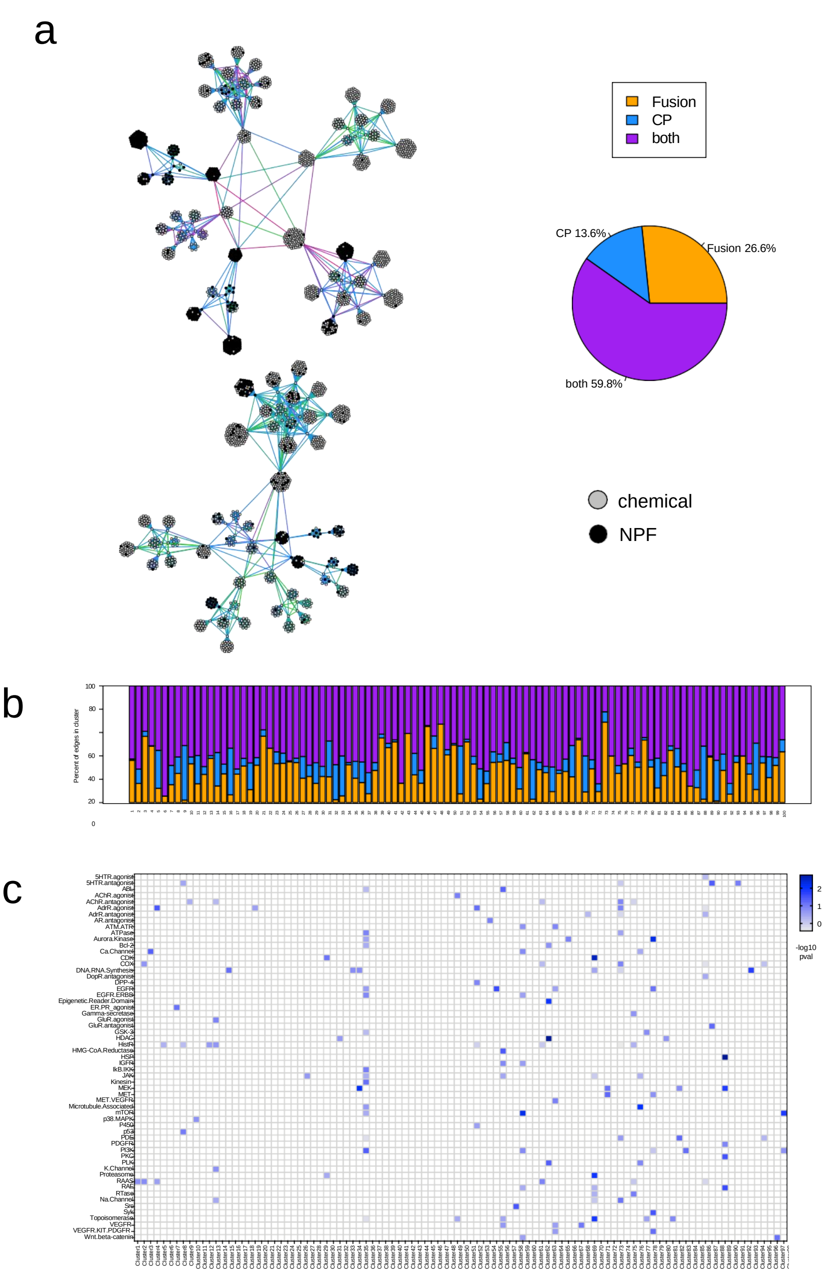

**Supplementary Figure 10. Affinity propagation clustering map of the SNF-pearson network.** A) Hierarchical affinity propagation clustering map of the SNF network using Pearson correlation as the similarity metric. Edges are colored based on contribution from individual datasets: Orange, supported by FUSION; blue, supported by CP; purple, supported by both datasets. Perturbagen type is indicated by node color: black, NPF; gray, pure chemical. B) Bar plot showing the percent of total edges in each APC cluster that are supported by FUSION, CP, or both datasets. Clusters are labeled by cluster number. C) Heatmap showing minus log10 p-values calculated by hypergeometric test for each target annotation class, per APC cluster. Target classes without significant enrichment in any cluster are omitted (Bonferroni-corrected alpha = 0.0016).

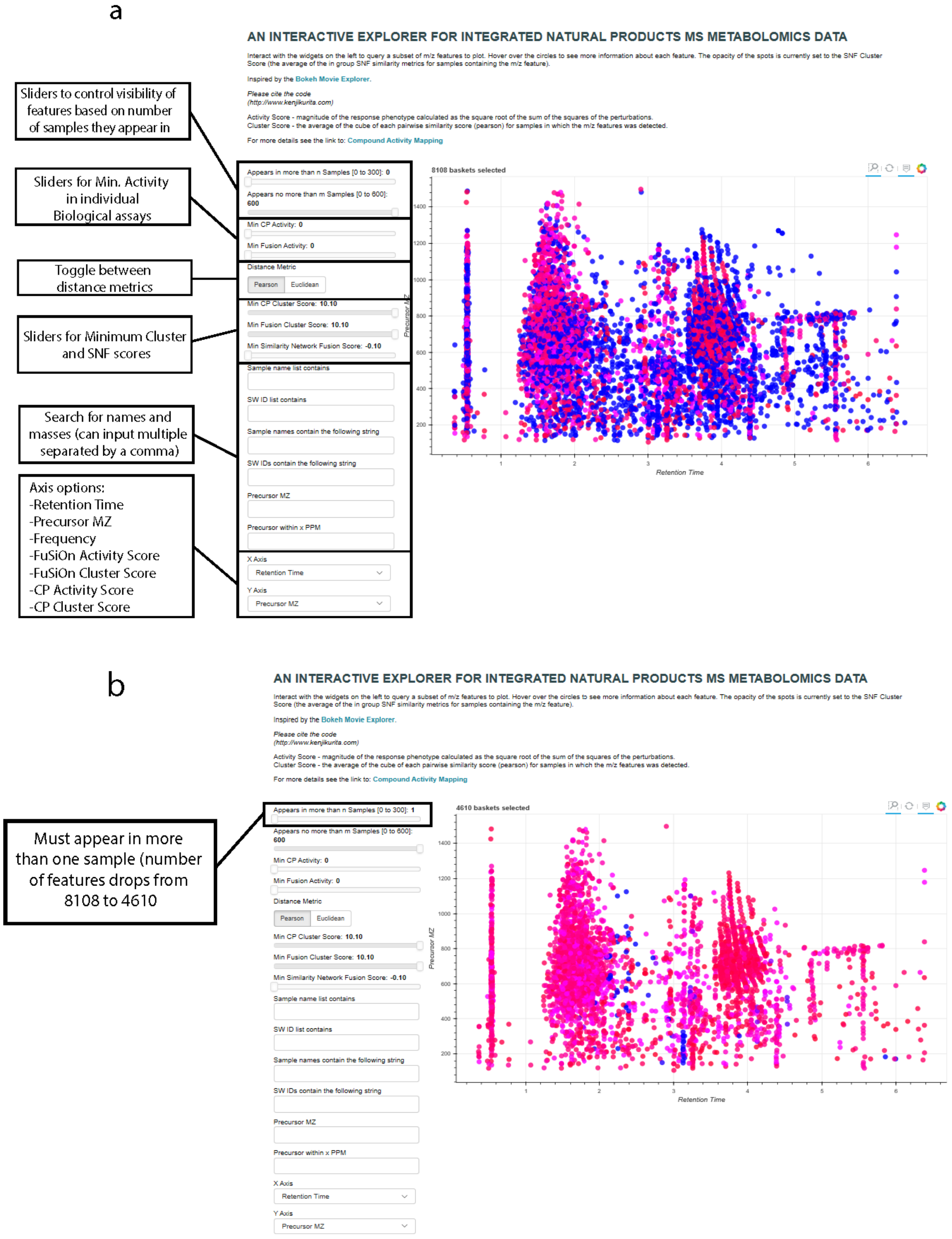
a
Sliders to control visibility of features based on number of samples they appear in
Sliders for Min. Activity in individual Biological assays
Toggle between distance metrics
Sliders for Minimum Cluster and SNF scores
Search for names and masses (can input multiple separated by a comma)
Axis options:
-Retention Time
-Precursor MZ
-Frequency
-FuSiOn Activity Score
-FuSiOn Cluster Score
-CP Activity Score
-CP Cluster Score
AN INTERACTIVE EXPLORER FOR INTEGRATED NATURAL PRODUCTS MS METABOLOMICS DATA
8108 baskets selected
Retention Time
Precursor MZ
b
Must appear in more than one sample (number of features drops from 8108 to 4610
AN INTERACTIVE EXPLORER FOR INTEGRATED NATURAL PRODUCTS MS METABOLOMICS DATA
4610 baskets selected
Retention Time
Precursor MZ

C

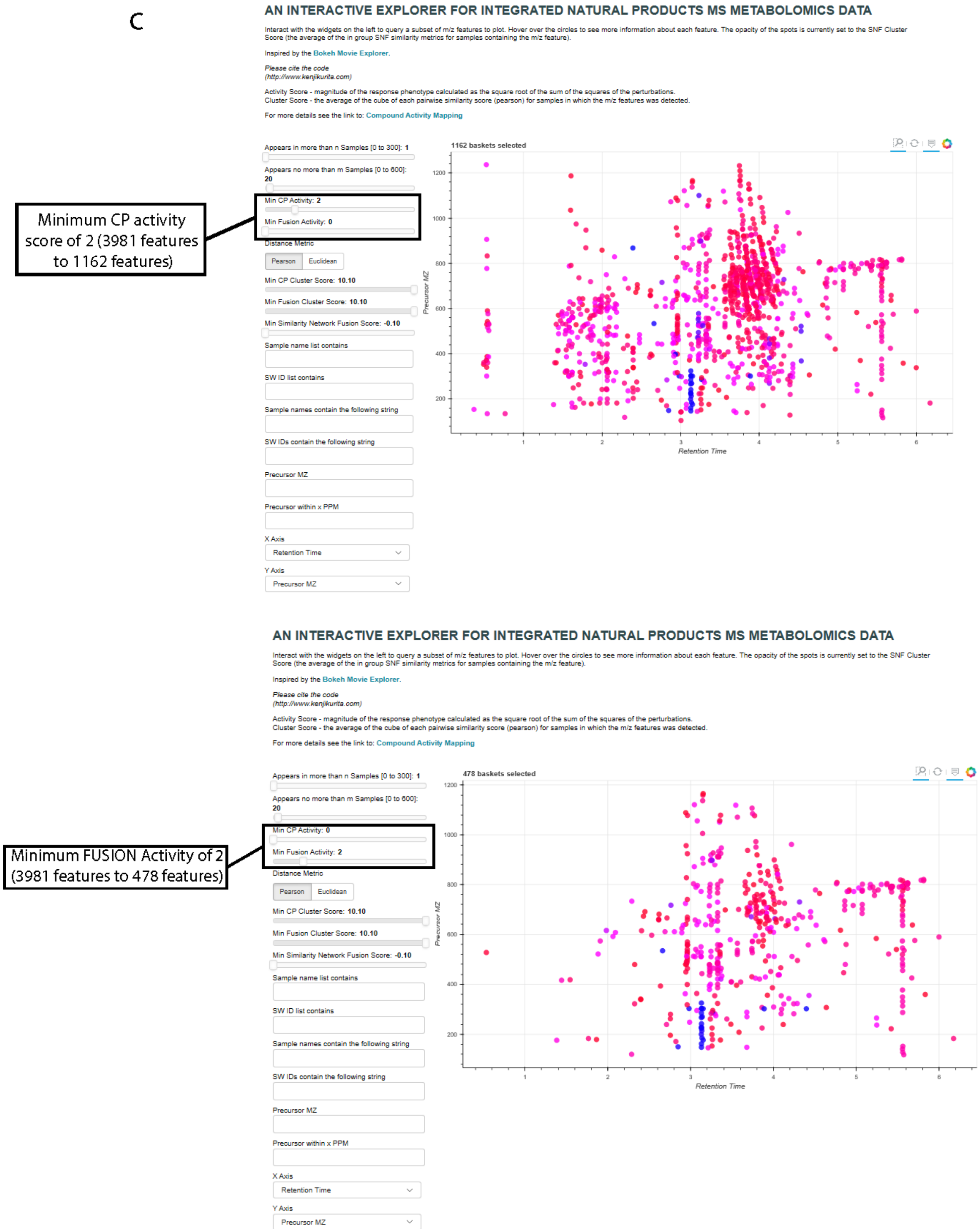

AN INTERACTIVE EXPLORER FOR INTEGRATED NATURAL PRODUCTS MS METABOLOMICS DATA
Interact with the widgets on the left to query a subset of m/z features to plot. Hover over the circles to see more information about each feature. The opacity of the spots is currently set to the SNF Cluster Score (the average of the in group SNF similarity metrics for samples containing the m/z feature).
Inspired by the Bokeh Movie Explorer.
Please cite the code (http://www.kenjikurita.com)
Activity Score - magnitude of the response phenotype calculated as the square root of the sum of the squares of the perturbations.
Cluster Score - the average of the cube of each pairwise similarity score (pearson) for samples in which the m/z features was detected.
For more details see the link to: Compound Activity Mapping
Appears in more than n Samples [0 to 300]: 1
Appears no more than m Samples [0 to 600]: 20
Min CP Activity: 2
Min Fusion Activity: 0
Distance Metric
Pearson
Euclidean
Min CP Cluster Score: 10.10
Min Fusion Cluster Score: 10.10
Min Similarity Network Fusion Score: -0.10
Sample name list contains
SW ID list contains
Sample names contain the following string
SW IDs contain the following string
Precursor MZ
Precursor within x PPM
X Axis
Retention Time
Y Axis
Precursor MZ
1162 baskets selected
Precursor MZ
Retention Time
Minimum CP activity score of 2 (3981 features to 1162 features)
AN INTERACTIVE EXPLORER FOR INTEGRATED NATURAL PRODUCTS MS METABOLOMICS DATA
Min CP Activity: 0
Min Fusion Activity: 2
478 baskets selected
Minimum FUSION Activity of 2 (3981 features to 478 features)

d

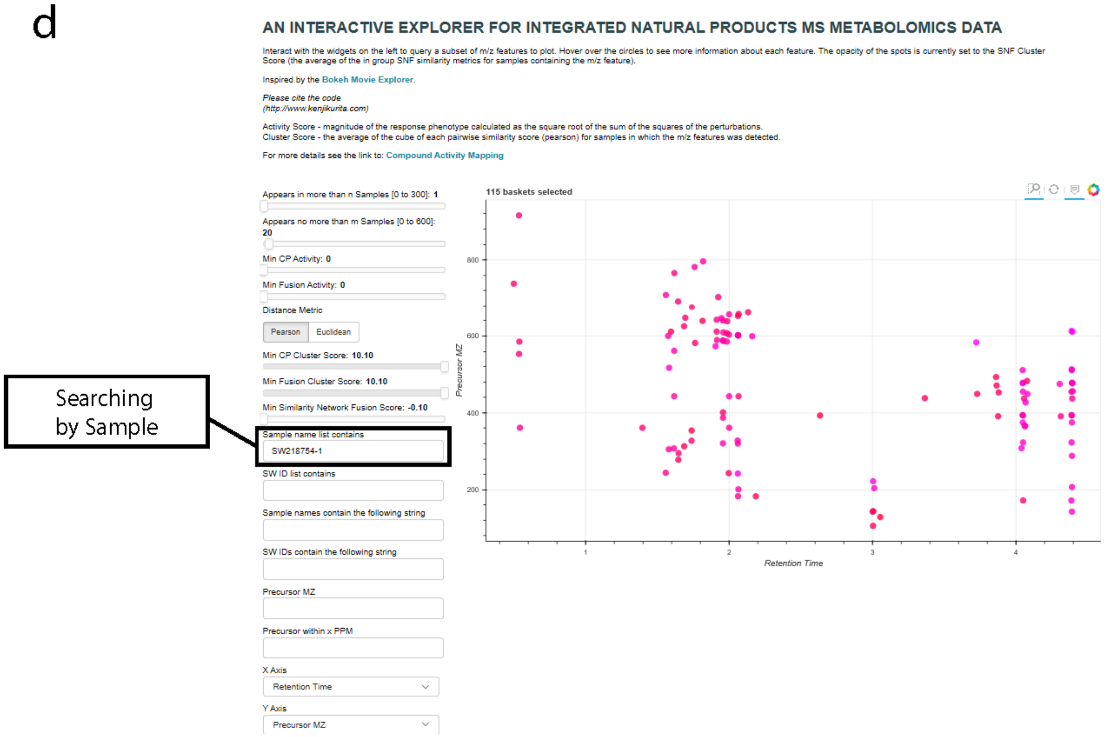

**Supplementary Figure 11: Bokeh Server demonstration of functionality.** (**a**) Labeled webpage for the bokeh server showing functionality and controllable components. (**b**) Minimum Occurrence set to 2 reduces the number of singleton ms features to ~4000. (**c**) Depiction of setting activity minimums in both CP and FUSION. (**d**) Searching bokeh server by sample (SW218754-1) to display all features in that fraction. General search features: Multiple codes may be searched at the same time using “, (space)” to show features in common among all searched codes. Using the text boxes and sliders on the left side of the plot, the feature list can be filtered to include only candidate features of interest. For example, increasing the Minimum Similarity Network Fusion Score slider removes features with low SNF Scores that are poorly correlated with any specific phenotype. Inserting a list of sample names into the 'Sample list name contains' text box filters the results to show only features present in a specific cluster of interest. Features are color-coded by SNF Score, from blue (strongly active) to red (weakly active). In addition, hovering the mouse over each feature in the plot reveals a pop-up window containing information about the feature including mass spectrometric data (*m/z*, rt, CCS) and distribution across the sample set.

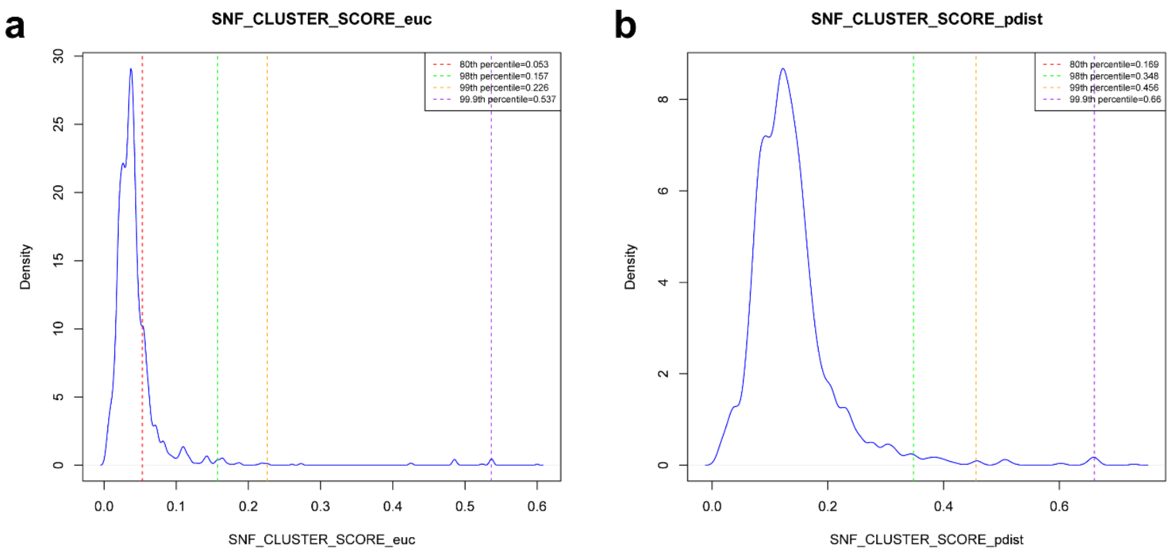

**Supplementary Figure 12. Density distributions of SNF scores for unique metabolites in the natural product fraction library.** SNF scores are shown as density plots using (**a**) Euclidean distance and (**b**) Pearson Correlation as the similarity metric. Percentiles are indicated by dotted lines as labeled.

a

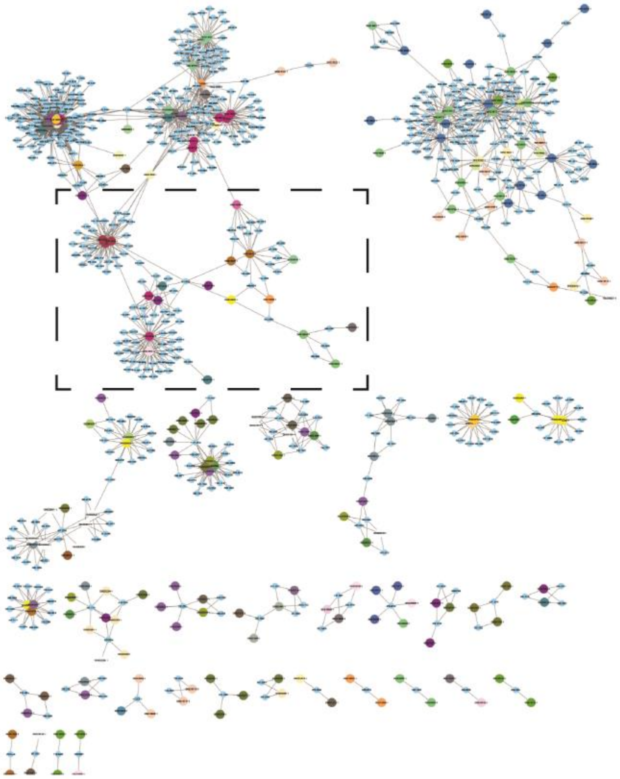

b

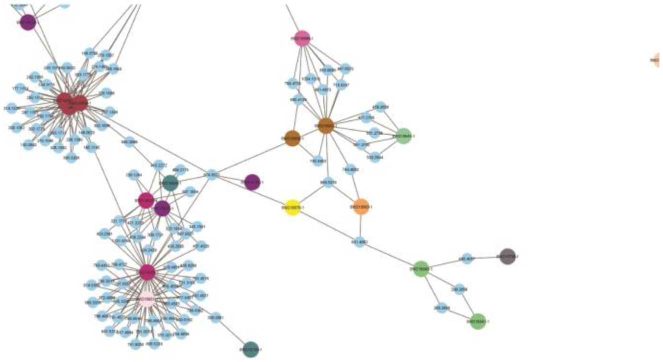

**Supplementary Figure 13: Compound Activity map for combined SNF profiles and untargeted metabolomics features.**

Large nodes represent extracts. Small nodes represent mass spectrometry features. Edges represent presence of mass spectrometric features in connected extracts. Only mass spectrometric features with predicted SNF scores >0.06 are included. A) Full Map. Large nodes color coded by AOC assignment class. B) Expansion of a representative region of the APC map with large nodes coloured by APC cluster.

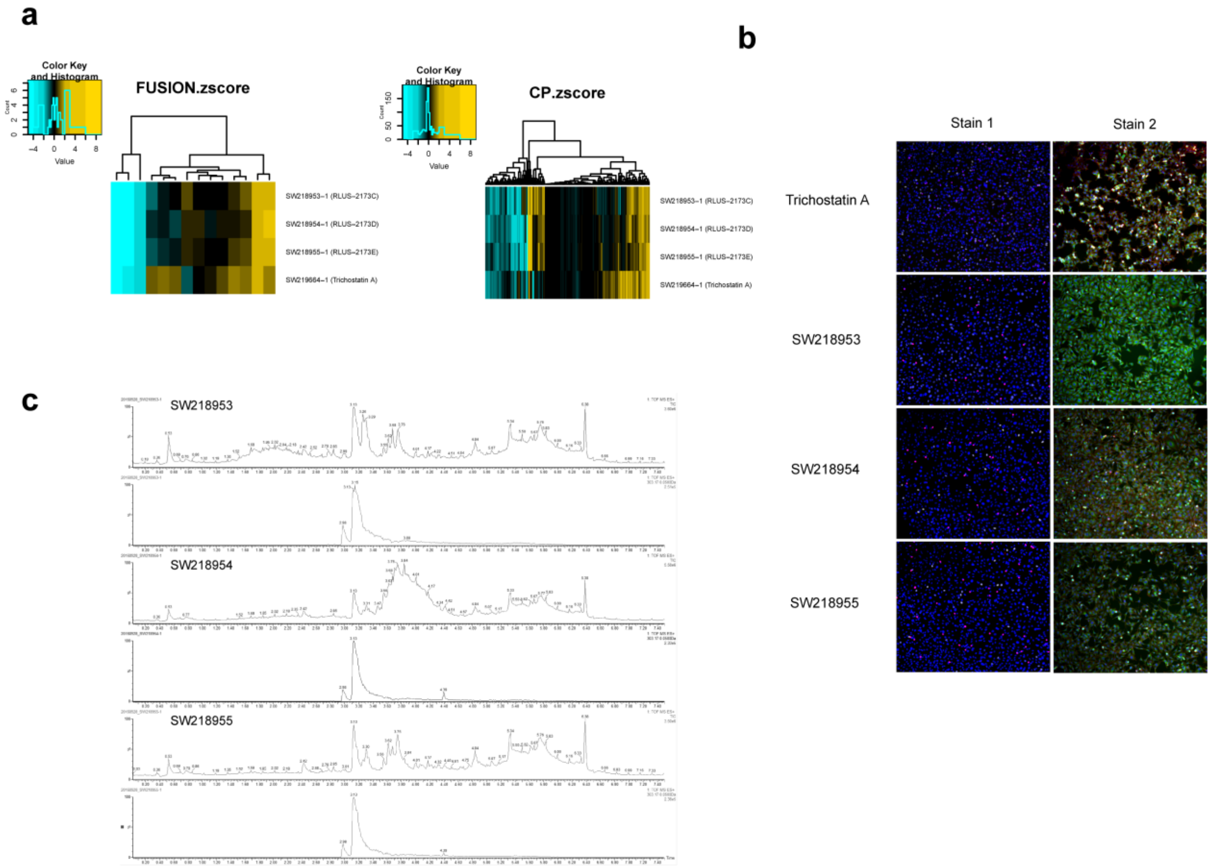

**Supplementary Figure 14: Signatures from natural product fractions containing trichostatin A.** (**a**) Heatmaps of FUSION z-scores and CP fingerprints, and (**b**) CP images from natural products containing trichostatin A and pure trichostatin A. (**c**) TIC and EIC 303.1703 (Trichostatin A) of SW218953, SW218954, and SW218955.

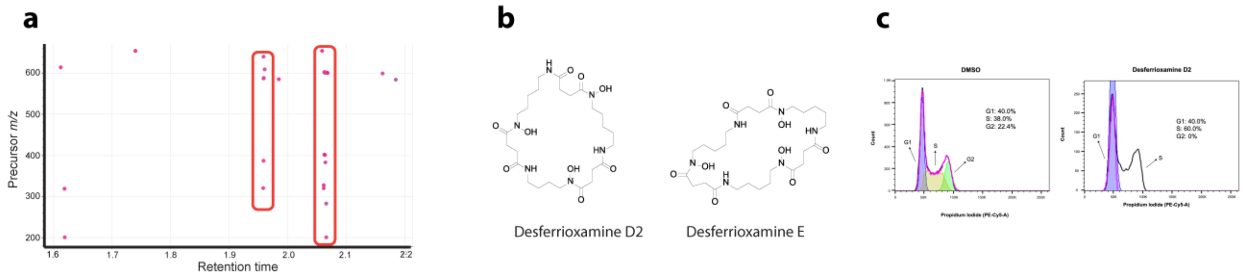

**Supplementary Figure 15: High SNF scores identifying targeted compounds for isolation with predicted modes of action.**

A) Retention plot showing common mass spectrometry features associated with SNF scores above 0.15. B) Chemical structures for desferrioxamines E and D2. C) Flow Cytometric histograms of cell cycle distribution measured with propidium iodide staining. H23 cells are treated with DMSO and desferrioxamine D2 (10mM) for 24 hours.

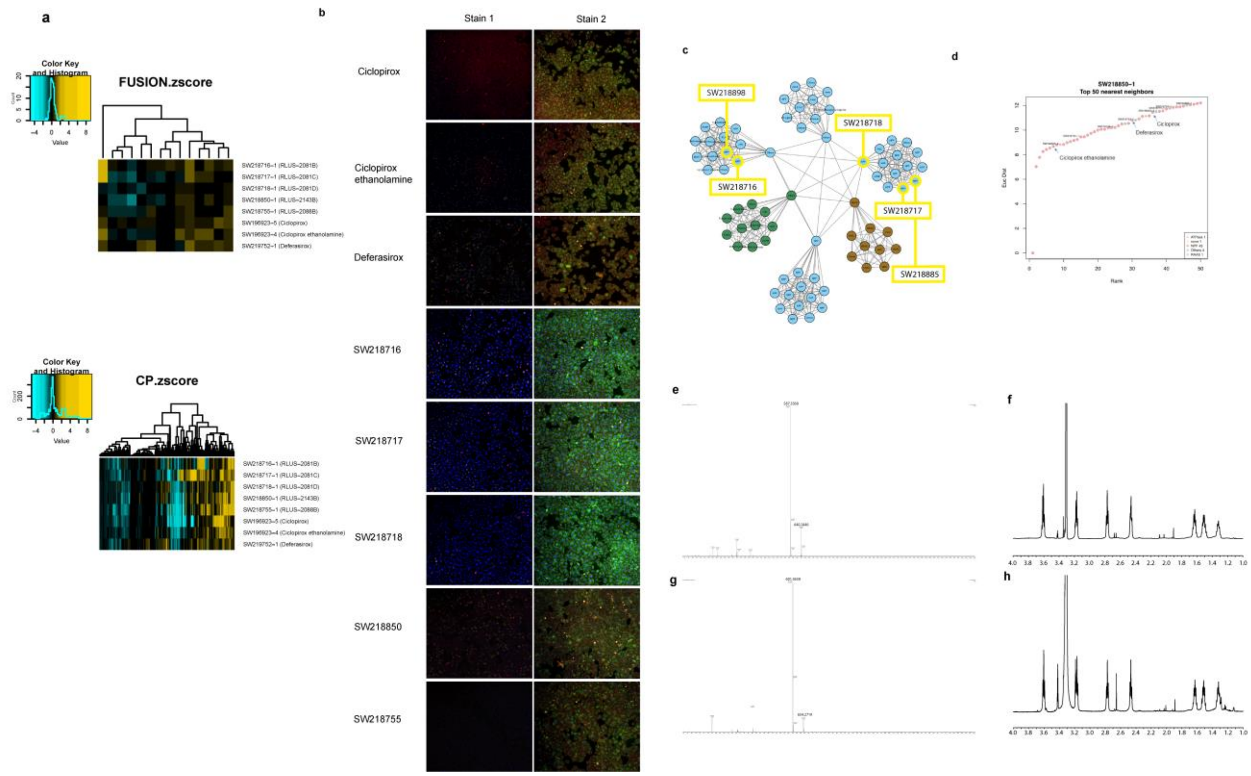

**Supplementary Figure 16. Signatures from natural product fractions containing desferrioxamine.** (**a**) Heatmaps of FUSION z-scores and CP fingerprints, and (**b**) CP images from natural products containing DFO. (**c**) SNF-Euclidean APC cluster showing proximity of DFO-containing natural product fractions. Nodes belonging to chemicals are labeled by target class assignment. Clusters with no significant enrichment for a particular target class are colored light blue. Clusters with significant enrichment for a particular target class are colored by the same scheme as they were in the original map (Figure 3). (**d**) Plot of the top 50 nearest neighbors to SW28850-1 in the SNF-Euclidean network, showing proximity to iron chelators from the Selleck library. (**e**) HRMS spectrum of desferrioxamine D2. (**f**) 600 MHz $^1$H NMR data in MeOD-*d4* of desferrioxamine D2. (**g**) MS spectrum of desferrioxamine E. (**h**) 600 MHz $^1$H NMR data in MeOD-*d4* of desferrioxamine E.

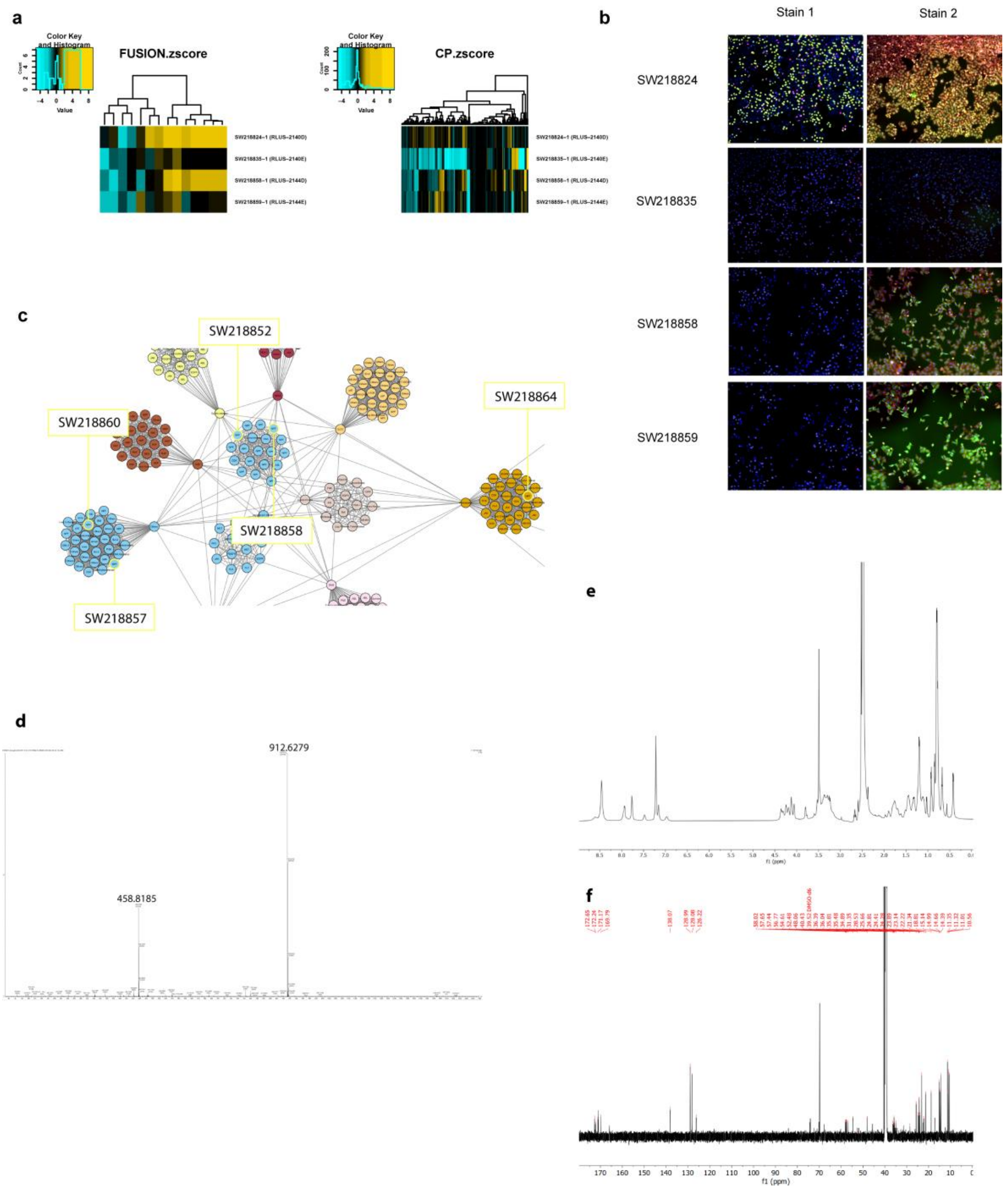

**Supplementary Figure 17. Signatures from natural product fractions containing surugamide.** (**a**) Heatmaps of FUSION z-scores and CP fingerprints, and (**b**) CP images from natural products containing surugamides. (**c**) SNF-Euclidean APC cluster showing proximity of surugamide-containing natural product fractions. Clusters are labeled and colored as described in Supplementary Figure 10. (**d**) HRMS spectrum of surugamide A. (**e**)$^{1}$H NMR spectrum and (**f**) $^{13}$C NMR spectrum for surugamide A in DMSO-*d6* at 600 MHz.

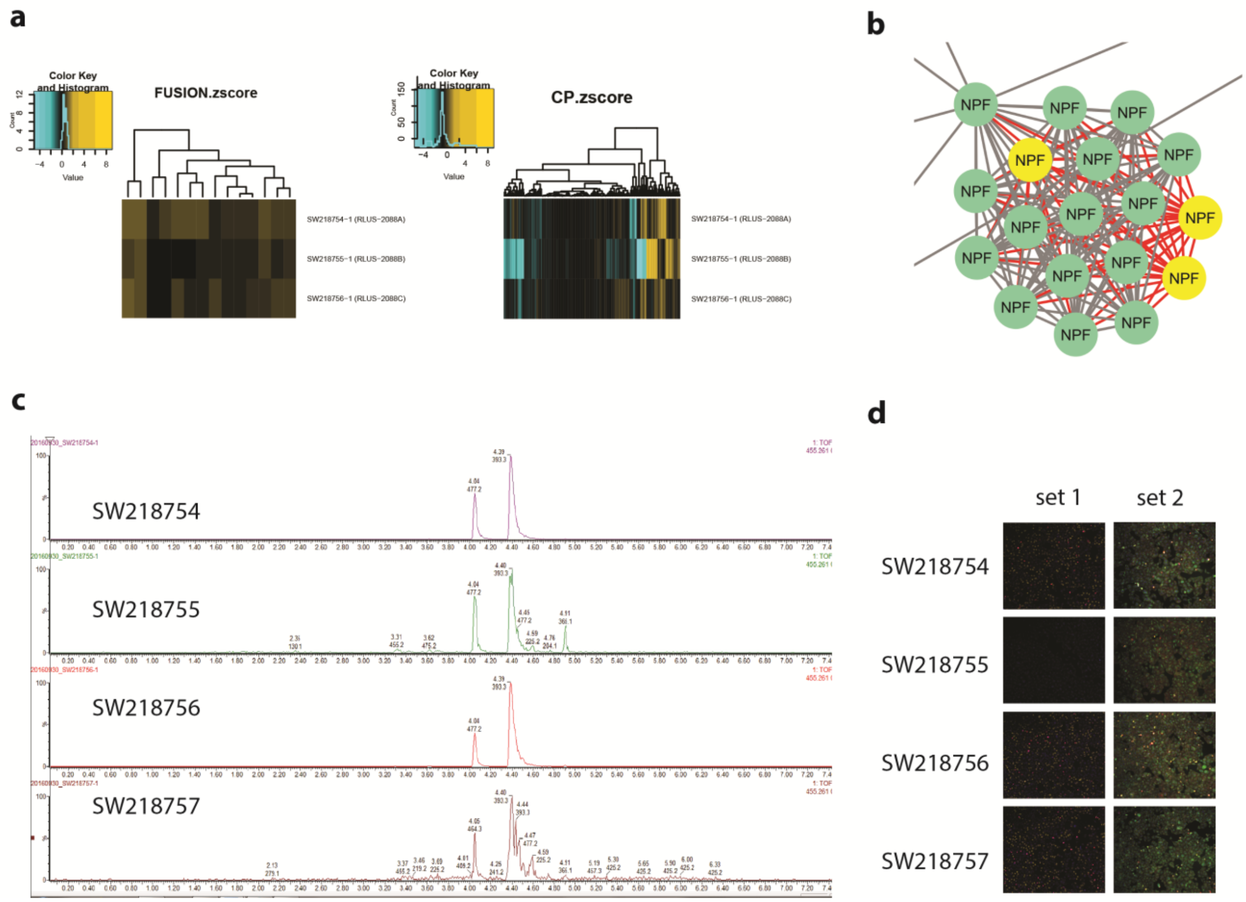

**Supplementary Figure 18. Signatures from natural product fractions containing parkamycin A.** (a) Heatmaps of FUSION z-scores and CP fingerprints, (b) SNF-Euclidean APC cluster showing proximity of parkamycin A containing natural product fractions. Clusters are labeled and colored as described in Supplementary Figure 10., (c) EIC of m/z 455.2560 in adjacent fractions all containing parkamycin A (d) CP images from natural products containing parkamycin A

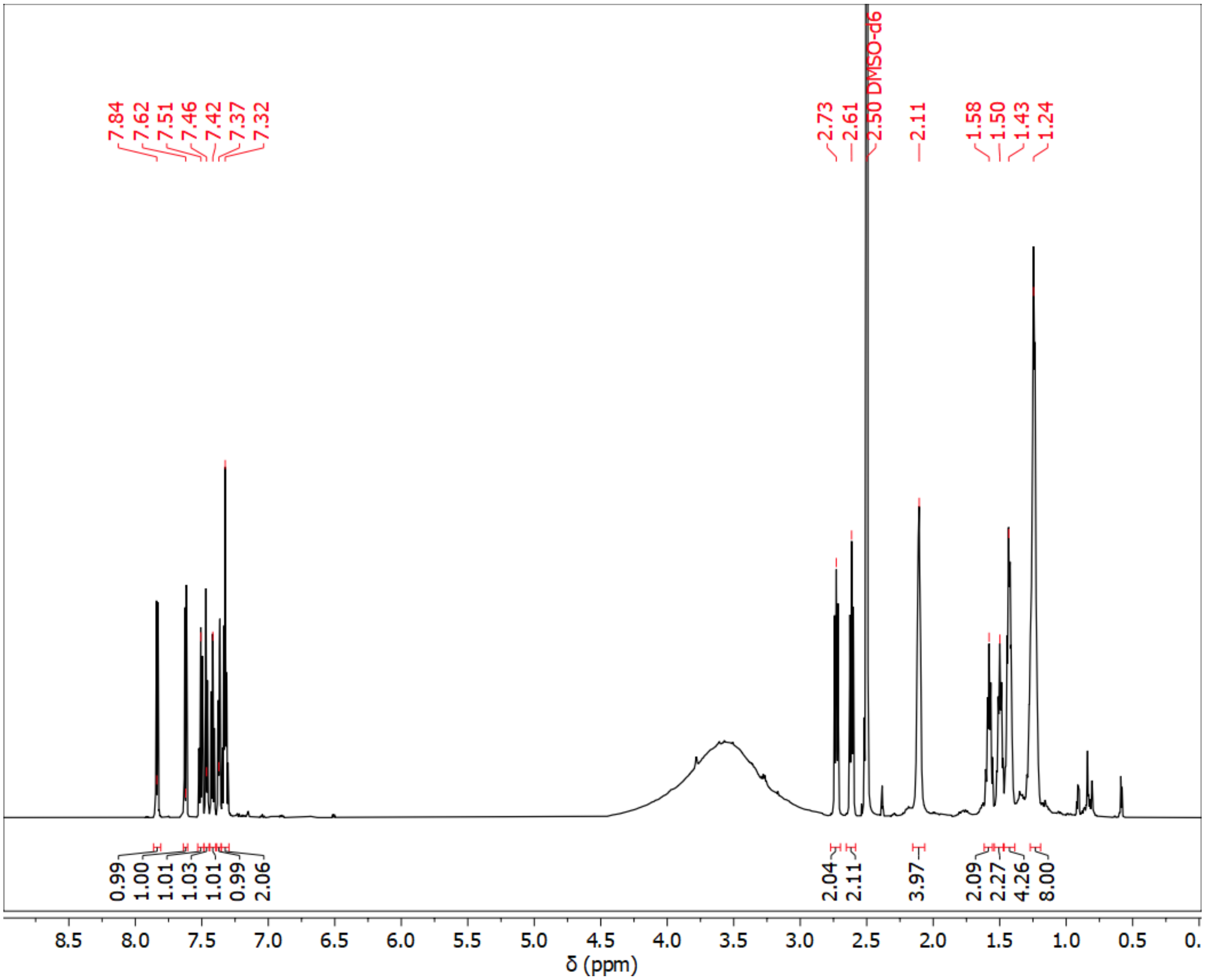

**Supplementary Figure 19. $^{1}$H NMR of parkamycin A.**

$^{1}$H NMR spectrum collected at 600 MHz in DMSO-*d6*

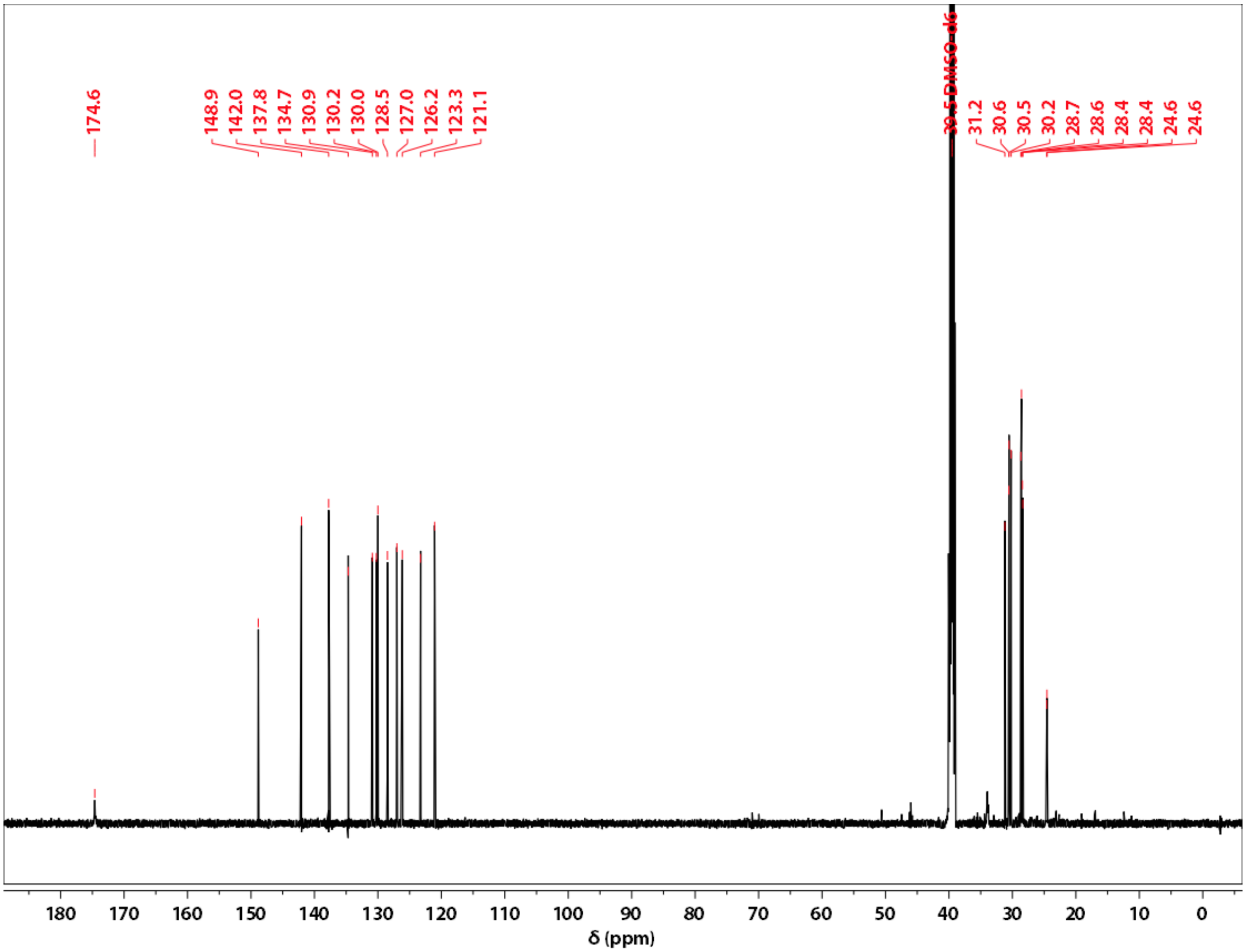

**Supplementary Figure 20. $^{13}$C NMR of parkamycin A.**

$^{13}$C NMR spectrum collected at 150 MHz in DMSO-*d6*

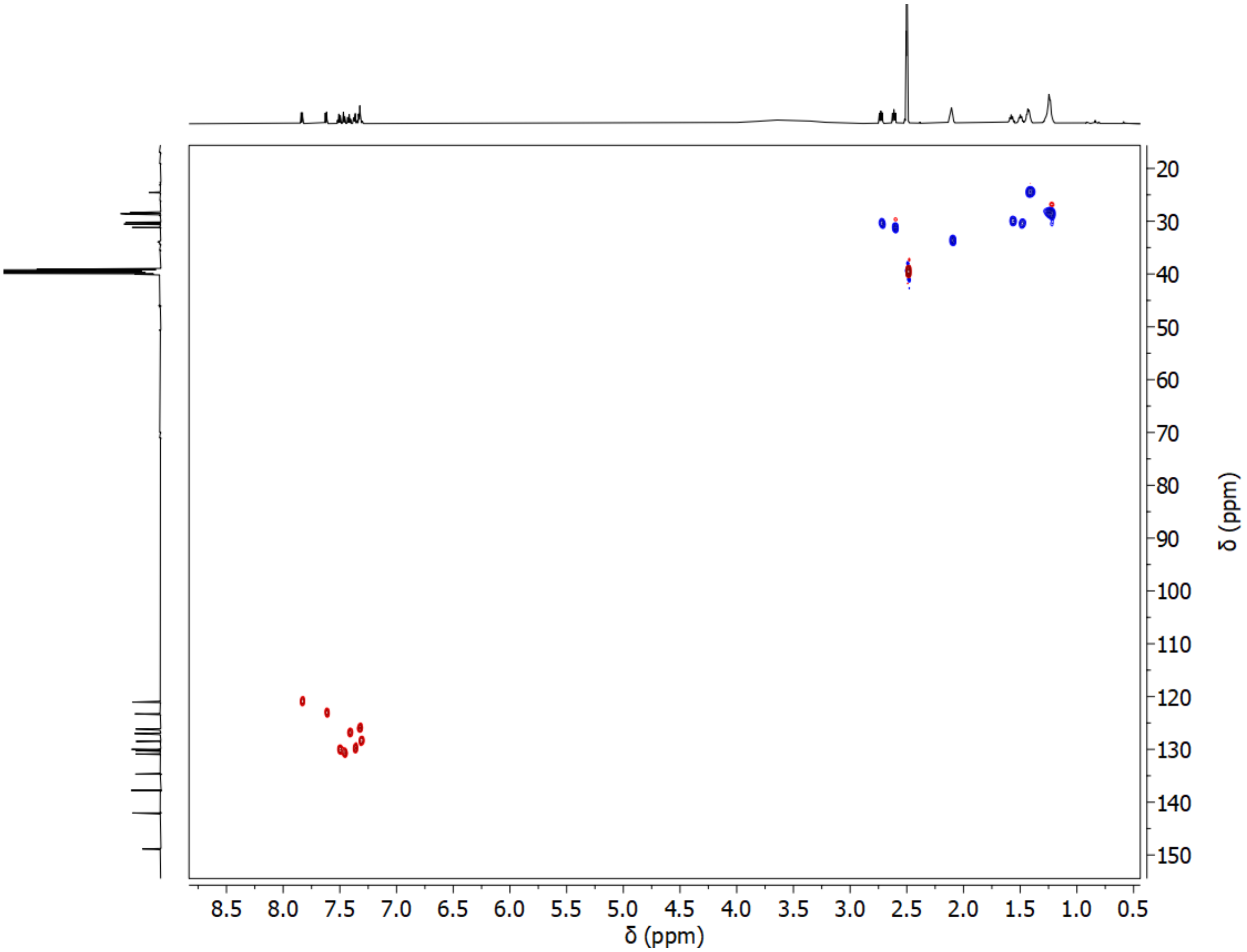

**Supplementary Figure 21. HSQC NMR spectrum of parkamycin A.**
HSQC NMR spectrum collected at 600 MHz in DMSO-*d6*

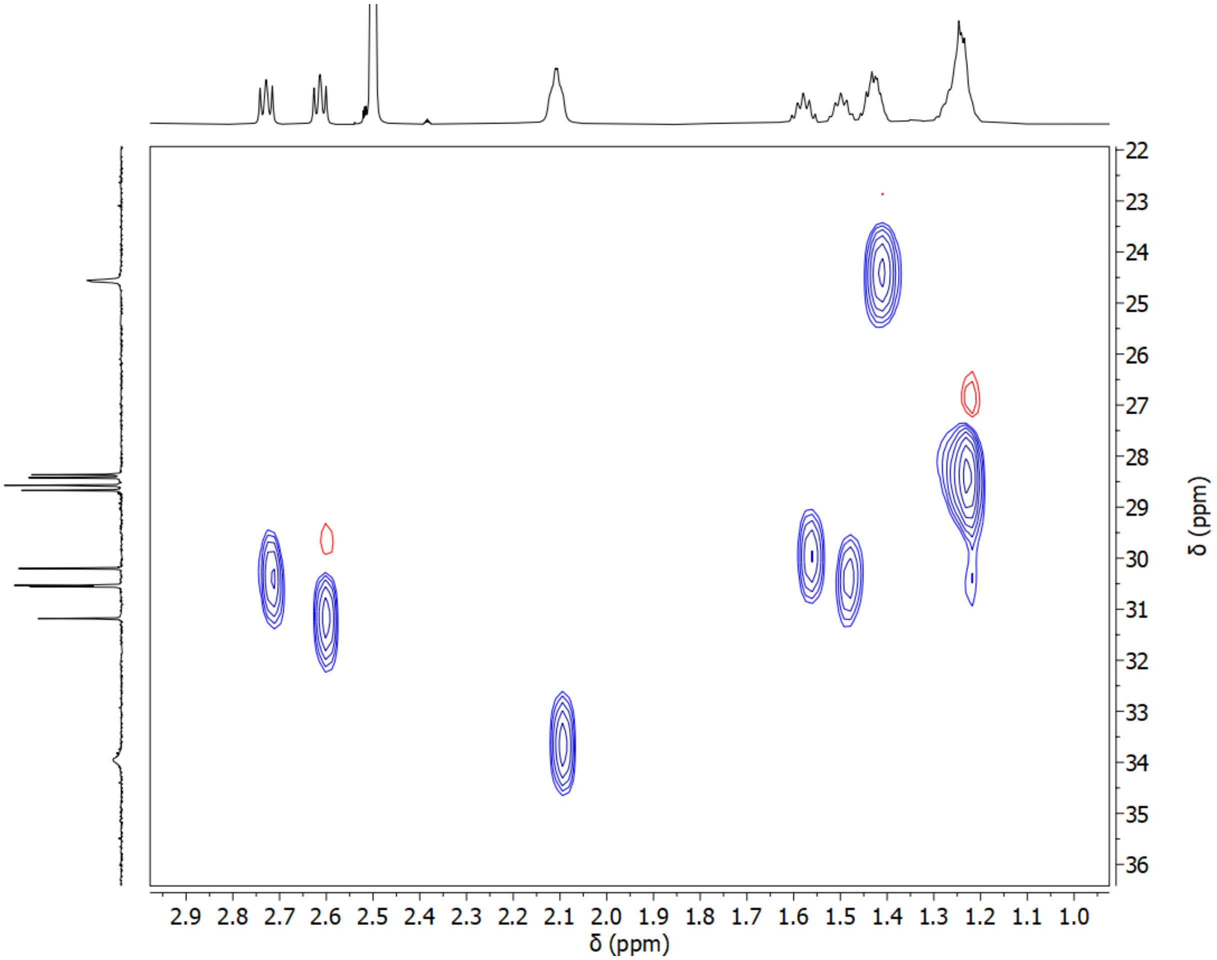

**Supplementary Figure 22. Expanded HSQC NMR spectrum of parkamycin A, region 1.**

Expanded HSQC NMR spectrum collected at 600 MHz in DMSO-*d6*

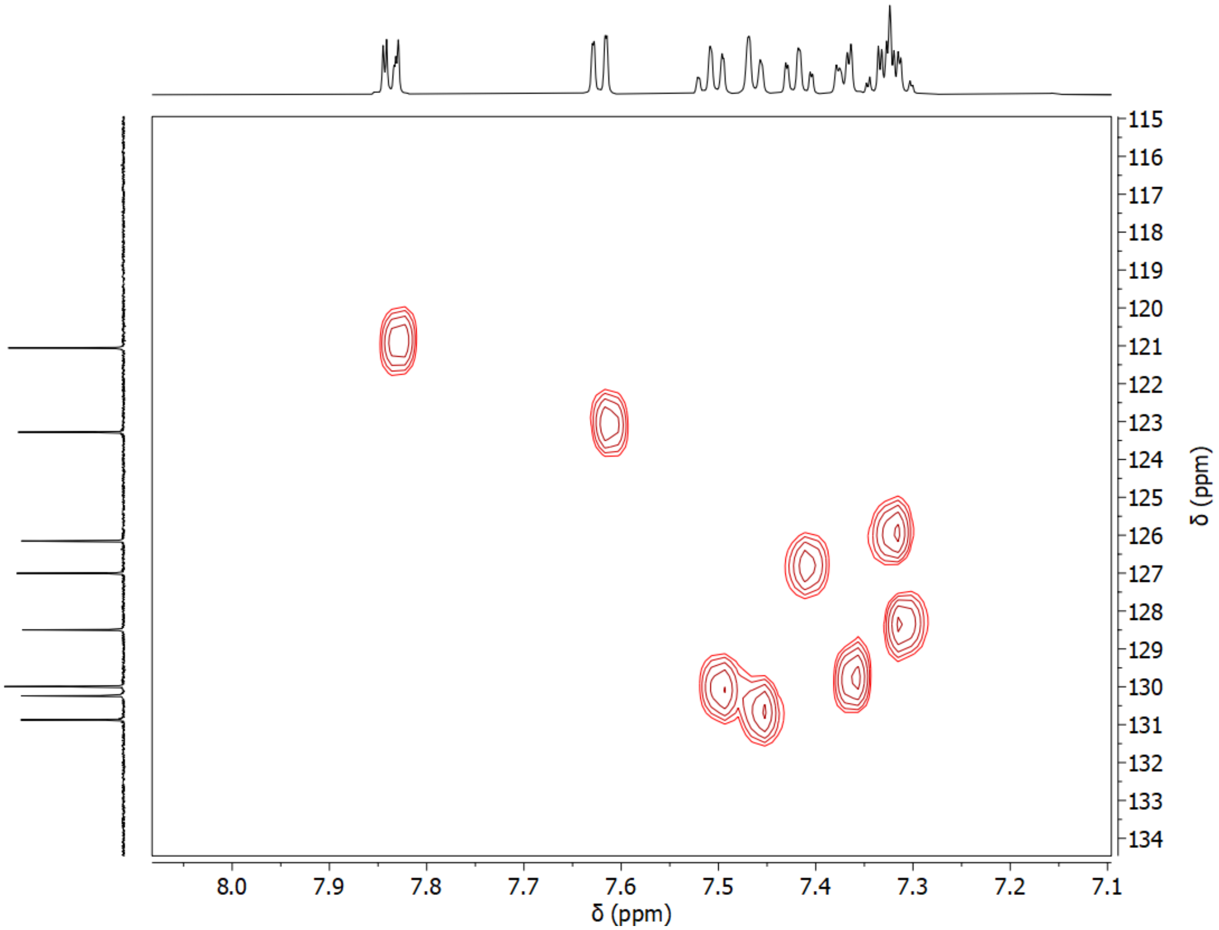

**Supplementary Figure 23. Expanded HSQC NMR spectrum of parkamycin A, region 2.**

Expanded HSQC NMR spectrum collected at 600 MHz in DMSO-*d6*

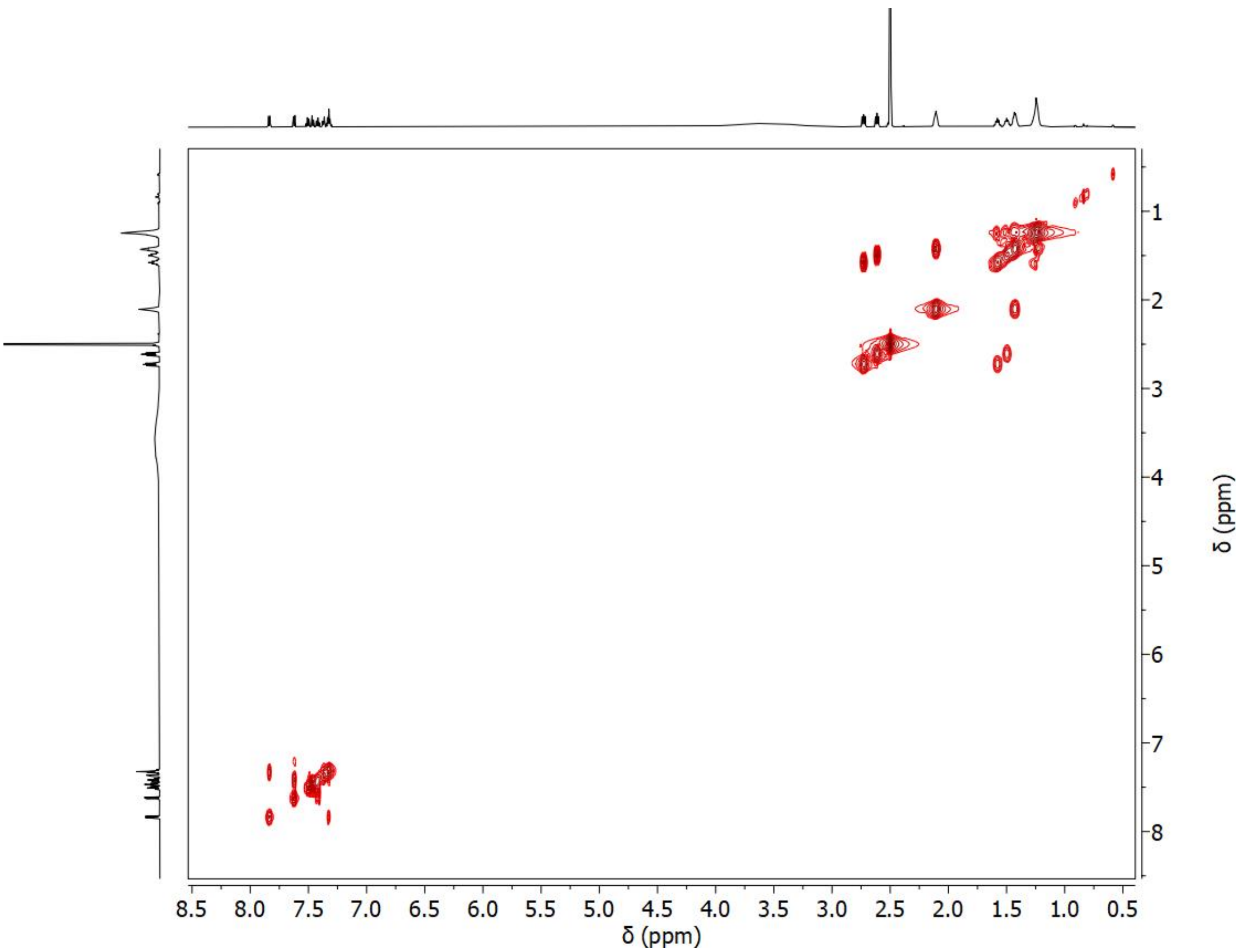

**Supplementary Figure 24. COSY NMR spectrum of parkamycin A.**

COSY NMR spectrum collected at 600 MHz in DMSO-*d6*

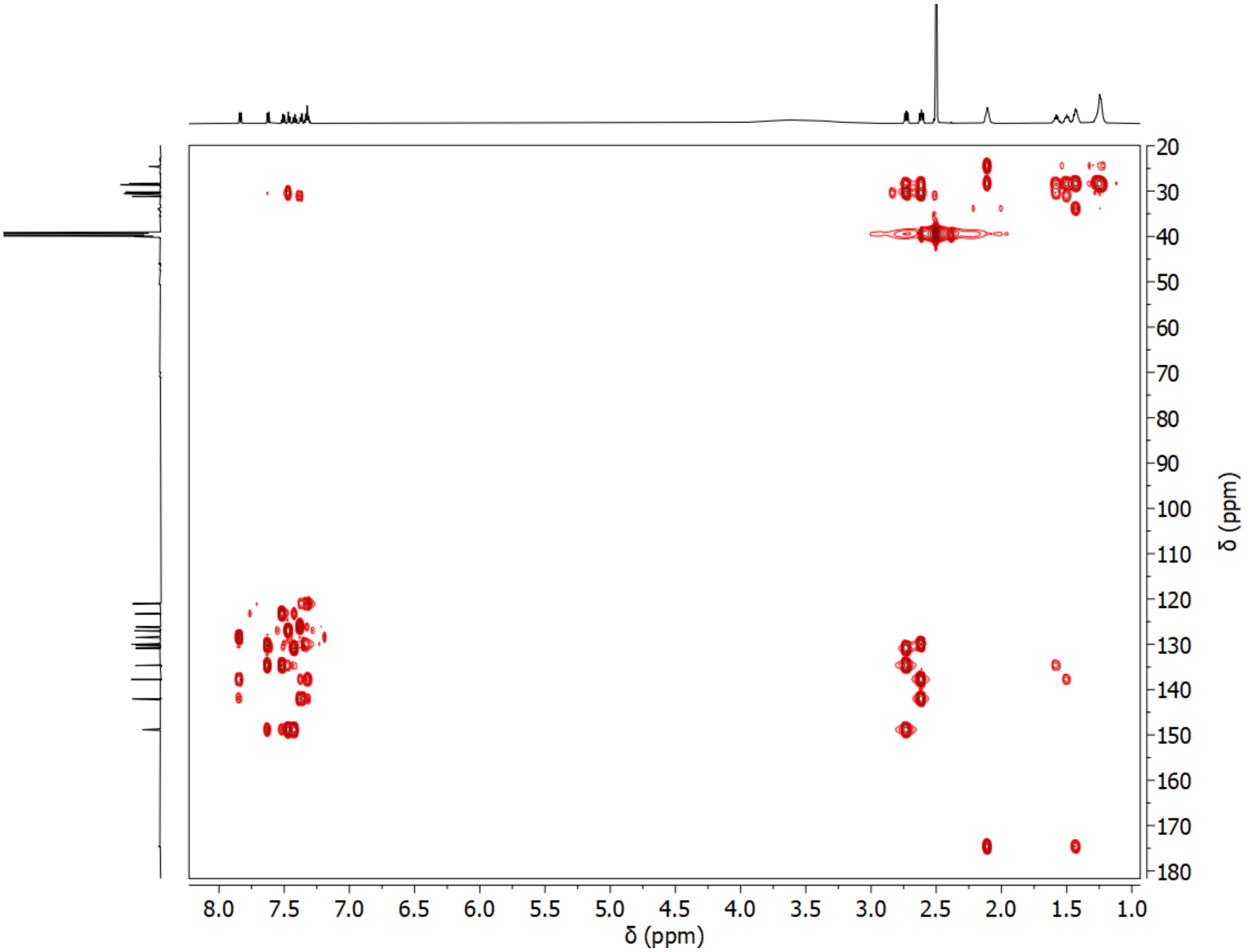

**Supplementary Figure 25. HMBC NMR spectrum of parkamycin A.**

HMBC NMR spectrum collected at 600 MHz in DMSO-*d6*

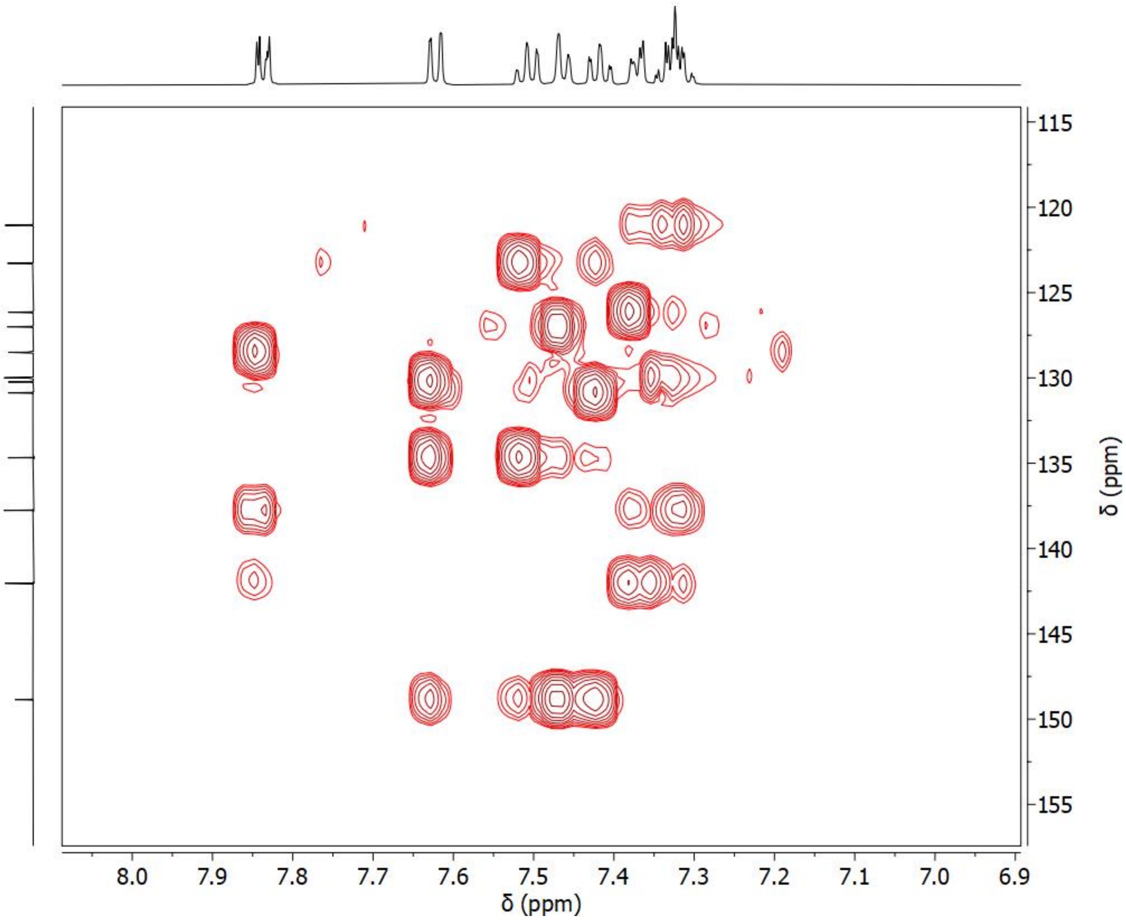

**Supplementary Figure 26. Expanded HMBC NMR spectrum of parkamycin A, region 1.**

Expanded HMBC NMR spectrum collected at 600 MHz in DMSO-*d6*

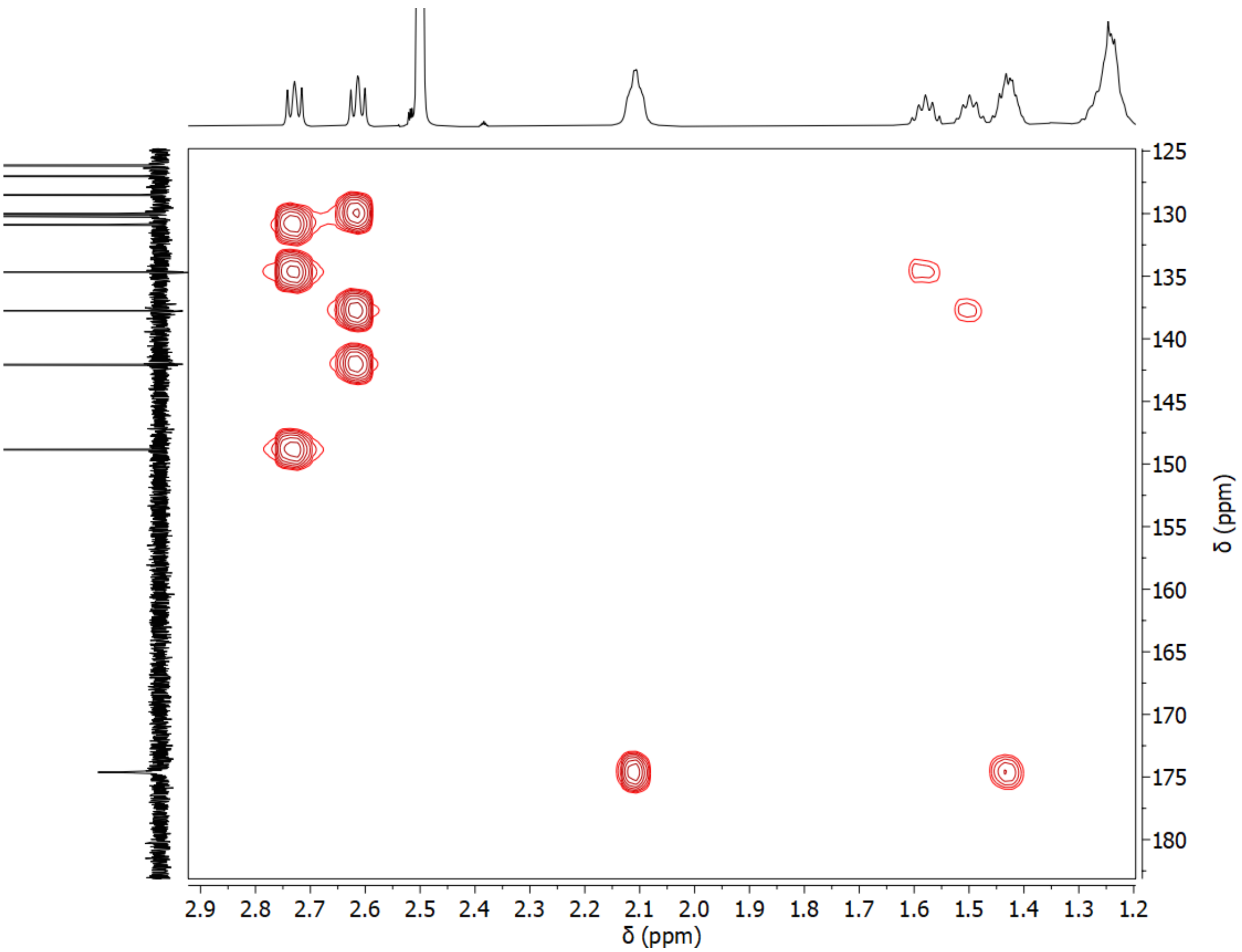

**Supplementary Figure 27. Expanded HMBC NMR spectrum of parkamycin A, region 2.**

Expanded HMBC NMR spectrum collected at 600 MHz in DMSO-*d6*

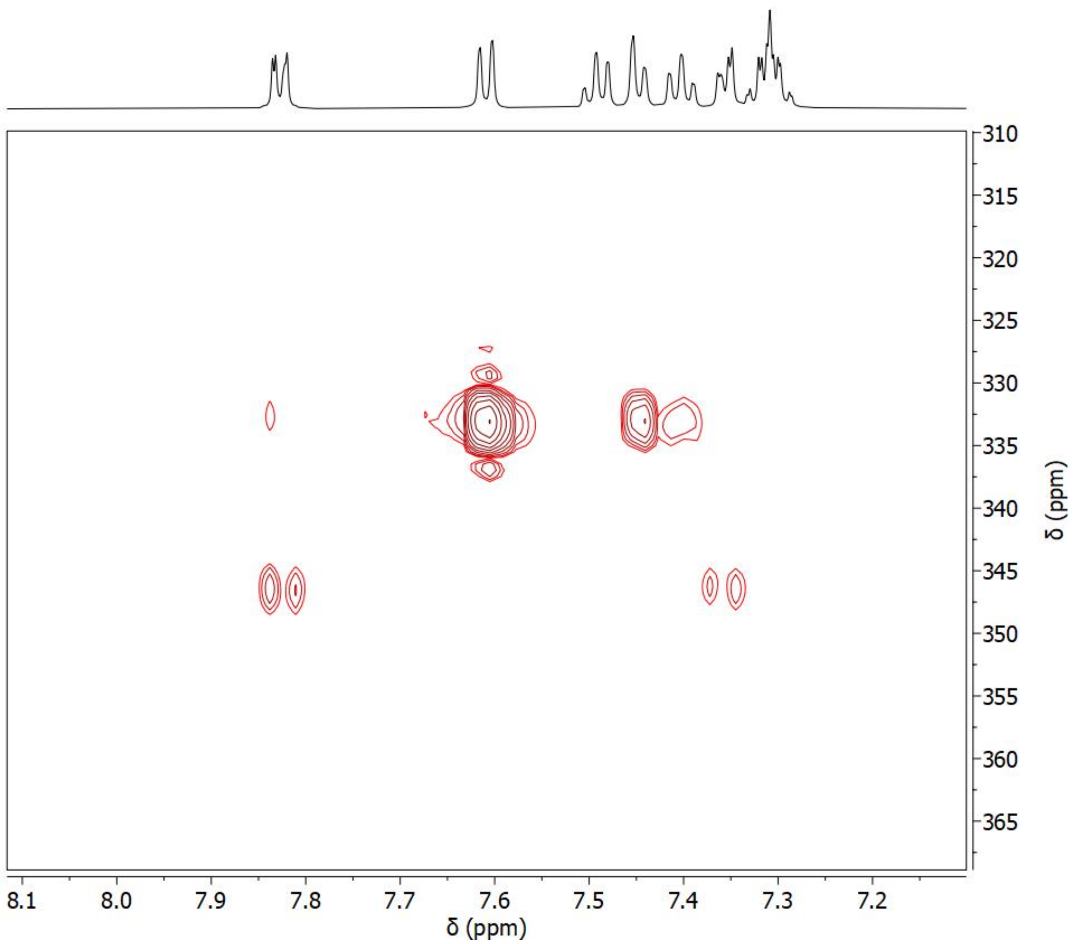

**Supplementary Figure 28. $^{15}$N-HMBC NMR spectrum of parkamycin A.**

$^{15}$N-HMBC NMR spectrum collected at 600 MHz in DMSO-*d6*

**Supplementary Table 1B: Selleck library compounds.** (**a**) List of all Selleck compounds included in the reference library and their target class annotations (For individual compound annotation, see SI Document "Supplemental Table 1A"). (**b**) The number of Selleck compounds in each target class.

| Class | Count |
|---|---|
| Others | 674 |
| none | 115 |
| I3K | 33 |
| Histamine Receptor | 33 |
| HDAC | 32 |
| AChR antagonist | 26 |
| COX | 26 |
| Adrenergic Receptor antagonist | 25 |
| Adrenergic Receptor agonist | 25 |
| DNA/RNA Synthesis | 24 |
| CDK | 22 |
| RAAS | 21 |
| PDE | 20 |
| 5HTR antagonist | 20 |
| VEGFR | 20 |
| JAK | 20 |
| Topoisomerase | 20 |
| Sodium Channel | 19 |
| mTOR | 19 |
| HSP (e.g. HSP90) | 18 |
| Aurora Kinase | 17 |
| EGFR | 17 |
| Calcium Channel | 17 |
| MEK | 16 |
| Microtubule Associated | 14 |
| PARP | 13 |
| RAF | 13 |
| 5HTR agonist | 13 |
| Reverse Transcriptase | 12 |
| P450 (e.g. CYP17) | 12 |
| Proteasome | 12 |
| Estrogen/progestogen Receptor_agonist | 11 |
| p38 MAPK | 11 |
| GSK-3 | 11 |
| EGFR_ERBB | 10 |
| Potassium Channel | 10 |
| Androgen Receptor antagonist | 10 |

| | |
|---|---|
| Wnt/beta-catenin | 10 |
| IGF-1R | 9 |
| ATPase | 9 |
| TGF-beta/Smad | 9 |
| PPAR | 9 |
| ABL | 9 |
| Histone Methyltransferase | 9 |
| Epigenetic Reader Domain | 9 |
| c-Met | 9 |
| Dopamine Receptor antagonist | 9 |
| Bcl-2 | 8 |
| HMG-CoA Reductase | 8 |
| IkB/IKK | 8 |
| DUB | 8 |
| Akt | 8 |
| PDGFR | 7 |
| GluR agonist | 7 |
| Gamma-secretase | 7 |
| GluR antagonist | 7 |
| PLK | 7 |
| Dehydrogenase | 7 |
| FAK | 7 |
| ATM/ATR | 7 |
| HIV Protease | 6 |
| Syk | 6 |
| DPP-4 | 6 |
| STAT | 6 |
| Estrogen/progestogen Receptor_antagonist | 6 |
| Transferase | 6 |
| Src | 6 |
| PKC | 6 |
| AChR agonist | 6 |
| Hedgehog/Smoothened | 6 |
| p53 | 6 |
| Kinesin | 6 |
| CFTR | 6 |
| Autophagy | 5 |
| c-Kit | 5 |
| Endothelin Receptor | 5 |
| SERT/NET | 5 |
| Aromatase | 5 |
| FLT3 | 5 |
| SERT | 5 |
| ROCK | 5 |

| | |
|---|---|
| Caspase | 5 |
| VEGFR_cKIT_PDGFR | 5 |
| Carbonic Anhydrase | 5 |
| MMP | 5 |
| Dopamine Receptor agonist | 5 |
| NF-kB | 5 |
| TNF-alpha | 5 |
| Integrase | 5 |
| DNA Methyltransferase | 5 |
| GPR | 5 |
| Sirtuin | 4 |
| Histone demethylases | 4 |
| HCV Protease | 4 |
| Opioid Receptor agonist | 4 |
| Factor Xa | 4 |
| E3 Ligase | 4 |
| GABA Receptor antagonist | 4 |
| Cysteine Protease | 4 |
| Opioid Receptor antagonist | 4 |
| PDK-1 | 4 |
| BTK | 4 |
| MET_VEGFR | 4 |
| Proton Pump | 4 |
| P2 Receptor | 4 |
| Pim | 4 |
| CETP | 4 |
| Estrogen/progestogen Receptor_SERM | 4 |
| IAP | 4 |
| OX Receptor | 4 |
| Chk | 4 |
| Cannabinoid Receptor agonist | 4 |
| AMPK | 4 |
| Serine Protease | 3 |
| DHFR | 3 |
| BMP | 3 |
| Androgen Receptor agonist | 3 |
| FAAH | 3 |
| FGFR | 3 |
| SGLT | 3 |
| S1P Receptor | 3 |
| LRRK2 | 3 |
| ALK | 3 |
| Rac | 3 |
| Mdm2 | 3 |

| | |
|---|---|
| CRM1 | 3 |
| Hydroxylase | 3 |
| Rho | 3 |
| p97 | 3 |
| 5-alpha Reductase | 3 |
| HIF | 3 |
| MAO | 3 |
| JNK | 3 |
| S6 Kinase | 3 |
| Telomerase | 3 |
| ELF4 | 2 |
| Liver X Receptor | 2 |
| PERK | 2 |
| CaSR | 2 |
| VDA | 2 |
| P-gp | 2 |
| Beta Amyloid | 2 |
| VEGFR_PDGFR | 2 |
| ERK | 2 |
| LPA Receptor | 2 |
| Dynamin | 2 |
| ELF2 | 2 |
| HER2 | 2 |
| PKA | 2 |
| Cannabinoid Receptor antagonist | 2 |
| PAK | 2 |
| cAMP | 2 |
| ERBB | 2 |
| CSF-1R | 2 |
| GABA Receptor agonist | 2 |
| IDO | 2 |
| TRPV | 2 |
| FXR | 2 |
| Phospholipase (e.g. PLA) | 2 |
| Ferroptosis | 2 |
| Cathepsin K | 2 |
| E2 | 1 |
| DDP-4 | 1 |
| MTH | 1 |
| NOD1 | 1 |
| MNK | 1 |
| Survivin | 1 |
| IDH2 | 1 |
| CXCR | 1 |

| | |
|---|---|
| PDHK | 1 |
| BMI | 1 |
| BET | 1 |
| Fo-ATPase | 1 |
| Tie-2 | 1 |
| gp120/CD4 | 1 |
| MBT | 1 |
| Wee1 | 1 |
| Histone Acetyltransferase | 1 |
| Axl | 1 |
| Substance P | 1 |
| AAAD/DOPA decarboxylase inhibitor | 1 |
| MT Receptor | 1 |
| Vasopressin Receptor | 1 |
| ribonucleotide reductase | 1 |
| c-Myc | 1 |
| APE | 1 |
| PAFR | 1 |
| IL Receptor | 1 |
| Ras | 1 |
| Ephrin receptor | 1 |
| GDP/GTP Exchange Factor Inhibitor | 1 |
| Procollagen C Proteinase | 1 |
| CCR | 1 |
| Integrin | 1 |
| Notch | 1 |
| Arp2/3 | 1 |
| E1 Activating | 1 |
| DNA-PK | 1 |
| ATGL | 1 |
| Ftase | 1 |
| Phosphorylase | 1 |

**Supplementary Table 2: K-means cluster membership in FUSION and CP for compounds in the top 30 largest target classes.**

| SWID | Other_ID | Pathway | Cmpd.Name | Fusion_ kmeans | CP_kmeans |
|---|---|---|---|---|---|
| SW220242-1 | S2852 | 5HTR.agonist | BRL-54443 | 1 | 26 |
| SW197244-4 | S2025 | 5HTR.agonist | Urapidil HCl | 6 | 26 |
| SW197596-2 | S1385 | 5HTR.agonist | Mosapride Citrate | 7 | 22 |
| SW197624-3 | S1432 | 5HTR.agonist | Sumatriptan Succinate | 7 | 22 |
| SW219416-1 | S1488 | 5HTR.agonist | Naratriptan | 7 | 8 |
| SW219573-1 | S4109 | 5HTR.agonist | Lorcaserin HCl | 7 | 10 |
| SW219882-1 | S1436 | 5HTR.agonist | Tianeptine sodium | 7 | 12 |
| SW197762-2 | S1649 | 5HTR.agonist | Zolmitriptan | 21 | 3 |
| SW219198-2 | S2875 | 5HTR.agonist | Prucalopride | 21 | 25 |
| SW197669-2 | S1607 | 5HTR.agonist | Rizatriptan Benzoate | 23 | 18 |
| SW197521-3 | S1975 | 5HTR.agonist | Aripiprazole | 25 | 14 |
| SW219418-1 | S2096 | 5HTR.agonist | Almotriptan Malate | 28 | 26 |
| SW220149-1 | S3180 | 5HTR.agonist | Eletriptan HBr | 28 | 3 |
| SW100810-5 | S1390 | 5HTR.antagonist | Ondansetron HCl | 1 | 26 |
| SW219360-1 | S8021 | 5HTR.antagonist | Vortioxetine (Lu AA21004) HBr | 3 | 20 |
| SW219880-1 | S4053 | 5HTR.antagonist | Sertraline HCl | 3 | 20 |
| SW196337-3 | S3183 | 5HTR.antagonist | Amitriptyline HCl | 8 | 24 |
| SW196384-4 | S2541 | 5HTR.antagonist | Clomipramine HCl | 8 | 22 |
| SW219879-1 | S1283 | 5HTR.antagonist | Asenapine | 8 | 5 |
| SW219571-1 | S2663 | 5HTR.antagonist | WAY-100635 Maleate | 19 | 30 |
| SW219811-1 | S2865 | 5HTR.antagonist | VUF 10166 | 19 | 16 |
| SW219375-1 | S2677 | 5HTR.antagonist | BRL-15572 | 20 | 14 |
| SW220247-1 | S2459 | 5HTR.antagonist | Clozapine | 20 | 1 |
| SW220018-1 | S2894 | 5HTR.antagonist | SB742457 | 21 | 12 |
| SW220129-1 | S2849 | 5HTR.antagonist | SB269970 HCl | 21 | 30 |
| SW198927-2 | S1898 | 5HTR.antagonist | Tropisetron | 23 | 18 |
| SW219157-1 | S8010 | 5HTR.antagonist | PRX-08066 Maleic acid | 23 | 27 |
| SW219941-1 | S2856 | 5HTR.antagonist | SB271046 | 25 | 20 |
| SW196882-3 | S2232 | 5HTR.antagonist | Ketanserin | 26 | 11 |
| SW219177-1 | S1243 | 5HTR.antagonist | Agomelatine | 26 | 13 |
| SW219736-1 | S2698 | 5HTR.antagonist | RS-127445 | 26 | 16 |
| SW220248-1 | S2493 | 5HTR.antagonist | Olanzapine | 26 | 11 |
| SW197348-4 | S1615 | 5HTR.antagonist | Risperidone | 28 | 12 |
| SW219131-1 | S2497 | AChR.antagonist | Pancuronium dibromide | 1 | 25 |
| SW219141-1 | S3048 | AChR.antagonist | Solifenacin succinate | 3 | 4 |
| SW102176-4 | S2054 | AChR.antagonist | Orphenadrine Citrate | 6 | 30 |
| SW196544-3 | S2471 | AChR.antagonist | Gallamine Triethiodide | 7 | 1 |
| SW196713-3 | S4034 | AChR.antagonist | Diphemanil Methylsulfate | 7 | 25 |
| SW197005-3 | S4027 | AChR.antagonist | Flavoxate HCl | 7 | 3 |
| SW219039-1 | S4024 | AChR.antagonist | Homatropine Methylbromide | 7 | 25 |

| | | | | | |
|---|---|---|---|---|---|
| SW219041-1 | S2547 | AChR.antagonist | Tiotropium Bromide hydrate | 7 | 11 |
| SW220294-1 | S2130 | AChR.antagonist | Atropine | 7 | 8 |
| SW219082-1 | S4025 | AChR.antagonist | Homatropine Bromide | 12 | 25 |
| SW219426-1 | S3144 | AChR.antagonist | Darifenacin HBr | 12 | 24 |
| SW219846-1 | S2659 | AChR.antagonist | 5-hydroxymethyl Tolterodine (PNU 200577; 5-HMT; 5-HM) | 12 | 17 |
| SW196970-2 | S1285 | AChR.antagonist | Biperiden HCl | 15 | 14 |
| SW197495-2 | S2550 | AChR.antagonist | Tolterodine tartrate | 15 | 14 |
| SW196787-4 | S3117 | AChR.antagonist | Oxybutynin chloride | 16 | 4 |
| SW196691-3 | S1913 | AChR.antagonist | Tropicamide | 23 | 18 |
| SW199029-2 | S1608 | AChR.antagonist | Pyridostigmine Bromide | 23 | 26 |
| SW199663-2 | S2087 | AChR.antagonist | Rivastigmine Tartrate | 23 | 26 |
| SW219290-1 | S2240 | AChR.antagonist | Fesoterodine Fumarate | 24 | 8 |
| SW199343-2 | S2508 | AChR.antagonist | Scopolamine HBr | 26 | 30 |
| SW219038-1 | S1978 | AChR.antagonist | Methscopolamine | 26 | 22 |
| SW219548-1 | S4014 | AChR.antagonist | Hyoscyamine | 26 | 1 |
| SW219130-1 | S1397 | AChR.antagonist | Rocuronium Bromide | 28 | 20 |
| SW219146-1 | S2549 | AChR.antagonist | Trospium chloride | 28 | 18 |
| SW219176-1 | S4031 | AChR.antagonist | Aclidinium Bromide | 28 | 26 |
| SW219209-1 | S3047 | AChR.antagonist | Otilonium Bromide | 29 | 9 |
| SW220145-1 | S3083 | Adrenergic.Receptor. agonist | Indacaterol Maleate | 3 | 4 |
| SW199659-2 | S2020 | Adrenergic.Receptor. agonist | Formoterol Hemifumarate | 6 | 16 |
| SW196632-3 | S2495 | Adrenergic.Receptor. agonist | Oxymetazoline HCl | 7 | 25 |
| SW197098-3 | S4043 | Adrenergic.Receptor. agonist | Tetrahydrozoline HCl | 7 | 11 |
| SW197214-4 | S2438 | Adrenergic.Receptor. agonist | Synephrine HCl | 7 | 8 |
| SW219274-1 | S2522 | Adrenergic.Receptor. agonist | L-Adrenaline | 7 | 25 |
| SW219274-2 | S2521 | Adrenergic.Receptor. agonist | Epinephrine Bitartrate | 7 | 11 |
| SW196701-3 | S2519 | Adrenergic.Receptor. agonist | Naphazoline HCl | 12 | 23 |
| SW197212-2 | S2566 | Adrenergic.Receptor. agonist | Isoprenaline HCl | 12 | 11 |
| SW197214-5 | S2362 | Adrenergic.Receptor. agonist | Synephrine | 13 | 25 |
| SW219274-3 | S3061 | Adrenergic.Receptor. agonist | Epinephrine HCl | 13 | 25 |
| SW219607-1 | S3075 | Adrenergic.Receptor. agonist | Dexmedetomidine | 13 | 1 |
| SW196915-3 | S4277 | Adrenergic.Receptor. agonist | Bambuterol HCl | 16 | 20 |

| | | | | | |
|---|---|---|---|---|---|
| SW197048-3 | S2516 | Adrenergic.Receptor. agonist | Xylazine HCl | 20 | 30 |
| SW197681-3 | S2458 | Adrenergic.Receptor. agonist | Clonidine HCl | 20 | 11 |
| SW219096-1 | S2545 | Adrenergic.Receptor. agonist | Scopine | 20 | 1 |
| SW220261-1 | S3185 | Adrenergic.Receptor. agonist | Adrenalone HCl | 20 | 25 |
| SW220301-1 | S4009 | Adrenergic.Receptor. agonist | Mirabegron | 20 | 26 |
| SW199068-2 | S1437 | Adrenergic.Receptor. agonist | Tizanidine HCl | 21 | 22 |
| SW219239-1 | S3060 | Adrenergic.Receptor. agonist | Medetomidine HCl | 24 | 3 |
| SW197044-3 | S4065 | Adrenergic.Receptor. agonist | Guanabenz Acetate | 26 | 30 |
| SW219275-1 | S2533 | Adrenergic.Receptor. agonist | Ritodrine HCl | 26 | 1 |
| SW219446-1 | S2569 | Adrenergic.Receptor. agonist | Phenylephrine HCl | 28 | 3 |
| SW219456-1 | S2092 | Adrenergic.Receptor. agonist | Detomidine HCl | 28 | 8 |
| SW219607-2 | S2090 | Adrenergic.Receptor. agonist | Dexmedetomidine HCl (Precedex) | 28 | 26 |
| SW197547-3 | S1831 | Adrenergic.Receptor. antagonist | Carvedilol | 3 | 2 |
| SW219269-1 | S1549 | Adrenergic.Receptor. antagonist | Nebivolol | 3 | 16 |
| SW199199-2 | S1856 | Adrenergic.Receptor. antagonist | Metoprolol Tartrate | 6 | 26 |
| SW196705-3 | S4010 | Adrenergic.Receptor. antagonist | Acebutolol HCl | 7 | 26 |
| SW196913-4 | S1409 | Adrenergic.Receptor. antagonist | Alfuzosin HCl | 7 | 18 |
| SW219414-1 | S4076 | Adrenergic.Receptor. antagonist | Propranolol HCl | 7 | 24 |
| SW196892-3 | S2517 | Adrenergic.Receptor. antagonist | Maprotiline HCl | 8 | 27 |
| SW197099-3 | S1324 | Adrenergic.Receptor. antagonist | Doxazosin Mesylate | 8 | 5 |
| SW219365-1 | S2691 | Adrenergic.Receptor. antagonist | BMY 7378 | 12 | 24 |
| SW196595-3 | S4291 | Adrenergic.Receptor. antagonist | Labetalol HCl | 15 | 22 |
| SW197155-3 | S4278 | Adrenergic.Receptor. antagonist | Carteolol HCl | 15 | 10 |
| SW196909-3 | S4124 | Adrenergic.Receptor. antagonist | Tolazoline HCl | 16 | 3 |
| SW197151-3 | S1206 | Adrenergic.Receptor. antagonist | Bisoprolol fumarate | 20 | 26 |
| SW197352-4 | S2509 | Adrenergic.Receptor. antagonist | Sotalol | 20 | 1 |
| SW196570-4 | S2126 | Adrenergic.Receptor. antagonist | Naftopidil | 21 | 11 |
| SW196637-3 | S2038 | Adrenergic.Receptor. antagonist | Phentolamine Mesylate | 23 | 11 |

| | | | | | |
|---|---|---|---|---|---|
| SW196917-4 | S1827 | Adrenergic.Receptor. antagonist | Betaxolol hydrochloride (Betoptic) | 23 | 27 |
| SW196917-5 | S2091 | Adrenergic.Receptor. antagonist | Betaxolol | 23 | 30 |
| SW197327-3 | S2499 | Adrenergic.Receptor. antagonist | Phenoxybenzamin e HCl | 24 | 29 |
| SW222230-1 | S2113 | Adrenergic.Receptor. antagonist | Cisatracurium Besylate | 24 | 8 |
| SW196457-4 | S2059 | Adrenergic.Receptor. antagonist | Terazosin HCl | 28 | 26 |
| SW196570-5 | S1387 | Adrenergic.Receptor. antagonist | Naftopidil DiHCl | 28 | 23 |
| SW197240-3 | S4123 | Adrenergic.Receptor. antagonist | Timolol Maleate | 28 | 12 |
| SW219300-1 | S2086 | Adrenergic.Receptor. antagonist | Ivabradine HCl | 28 | 30 |
| SW219765-1 | S1613 | Adrenergic.Receptor. antagonist | Silodosin | 28 | 26 |
| SW219886-1 | S1154 | Aurora.Kinase | SNS-314 Mesylate | 2 | 21 |
| SW219458-1 | S7065 | Aurora.Kinase | MK-8745 | 6 | 2 |
| SW220051-1 | S2718 | Aurora.Kinase | TAK-901 | 10 | 2 |
| SW219462-1 | S1107 | Aurora.Kinase | Danusertib (PHA-739358) | 13 | 15 |
| SW219643-1 | S1100 | Aurora.Kinase | MLN8054 | 15 | 21 |
| SW219491-1 | S1171 | Aurora.Kinase | CYC116 | 17 | 28 |
| SW220282-1 | S1103 | Aurora.Kinase | ZM 447439 | 18 | 6 |
| SW219480-1 | S1519 | Aurora.Kinase | CCT129202 | 19 | 6 |
| SW219431-1 | S1147 | Aurora.Kinase | Barasertib (AZD1152-HQPA) | 21 | 28 |
| SW219731-1 | S2719 | Aurora.Kinase | AMG-900 | 22 | 28 |
| SW220174-1 | S2770 | Aurora.Kinase | MK-5108 (VX-689) | 23 | 6 |
| SW212828-2 | S1048 | Aurora.Kinase | VX-680 (Tozasertib; MK-0457) | 25 | 6 |
| SW219481-1 | S2744 | Aurora.Kinase | CCT137690 | 25 | 21 |
| SW219771-1 | S1133 | Aurora.Kinase | Alisertib (MLN8237) | 27 | 6 |
| SW219373-1 | S1454 | Aurora.Kinase | PHA-680632 | 29 | 15 |
| SW220199-1 | S1451 | Aurora.Kinase | Aurora A Inhibitor I | 29 | 27 |
| SW219406-1 | S1529 | Aurora.Kinase | Hesperadin | 30 | 15 |
| SW219347-1 | S2481 | Calcium.Channel | Manidipine | 2 | 29 |
| SW219784-1 | S1293 | Calcium.Channel | Cilnidipine | 2 | 29 |
| SW219238-1 | S1747 | Calcium.Channel | Nimodipine | 4 | 29 |
| SW219299-1 | S1885 | Calcium.Channel | Felodipine | 6 | 27 |
| SW197582-2 | S2017 | Calcium.Channel | Benidipine HCl | 14 | 4 |
| SW197620-3 | S1425 | Calcium.Channel | Ranolazine 2HCl | 15 | 14 |
| SW219737-1 | S2491 | Calcium.Channel | Nitrendipine | 15 | 5 |
| SW219720-1 | S2403 | Calcium.Channel | Tetrandrine | 18 | 12 |
| SW196411-3 | S2573 | Calcium.Channel | Tetracaine HCl | 20 | 26 |
| SW219572-1 | S2721 | Calcium.Channel | Nilvadipine | 21 | 20 |
| SW220228-1 | S1905 | Calcium.Channel | Amlodipine | 23 | 3 |
| SW219840-1 | S1994 | Calcium.Channel | Lacidipine | 27 | 5 |
| SW196530-3 | S2030 | Calcium.Channel | Flunarizine 2HCl | 28 | 4 |
| SW220017-1 | S1662 | Calcium.Channel | Isradipine | 28 | 23 |

| | | | | | |
|---|---|---|---|---|---|
| SW220087-1 | S2080 | Calcium.Channel | Clevidipine Butyrate | 28 | 16 |
| SW219236-1 | S3053 | Calcium.Channel | Azelnidipine | 29 | 12 |
| SW219348-1 | S2482 | Calcium.Channel | Manidipine 2HCl | 29 | 16 |
| SW219166-1 | S2014 | CDK | BMS-265246 | 5 | 19 |
| SW219356-1 | S1572 | CDK | BS-181 HCl | 6 | 20 |
| SW220101-1 | S7440 | CDK | LEE011 | 8 | 5 |
| SW219405-1 | S7158 | CDK | LY2835219 | 10 | 19 |
| SW219631-1 | S7461 | CDK | LDC000067 | 10 | 19 |
| SW219878-1 | S2742 | CDK | PHA-767491 | 10 | 28 |
| SW219949-1 | S2751 | CDK | Milciclib (PHA-848125) | 10 | 28 |
| SW219954-1 | S2735 | CDK | MK-8776 (SCH 900776) | 10 | 21 |
| SW220252-1 | S1487 | CDK | PHA-793887 | 10 | 28 |
| SW219231-1 | S2679 | CDK | Flavopiridol HCl | 11 | 28 |
| SW219310-1 | S8058 | CDK | P276-00 | 11 | 28 |
| SW219478-1 | S1145 | CDK | SNS-032 (BMS-387032) | 11 | 28 |
| SW219680-1 | S2621 | CDK | AZD5438 | 11 | 28 |
| SW220016-1 | S2768 | CDK | Dinaciclib (SCH727965) | 11 | 28 |
| SW220191-1 | S2688 | CDK | R547 | 11 | 28 |
| SW219950-1 | S7320 | CDK | TG003 | 12 | 6 |
| SW220195-1 | S1153 | CDK | Roscovitine (Seliciclib;CYC202) | 13 | 28 |
| SW220039-1 | S7114 | CDK | NU6027 | 14 | 26 |
| SW220083-1 | S7509 | CDK | ML167 | 14 | 8 |
| SW219396-1 | S1249 | CDK | JNJ-7706621 | 19 | 19 |
| SW220131-1 | S1116 | CDK | Palbociclib (PD-0332991) HCl | 27 | 5 |
| SW219609-1 | S1524 | CDK | AT7519 | 30 | 21 |
| SW219668-1 | S3043 | COX | Rofecoxib | 5 | 14 |
| SW196700-3 | S4078 | COX | Mefenamic Acid | 7 | 14 |
| SW196989-3 | S2577 | COX | Phenacetin | 7 | 26 |
| SW197564-2 | S4049 | COX | Valdecoxib | 7 | 10 |
| SW219542-1 | S4011 | COX | Ampiroxicam | 7 | 26 |
| SW196831-3 | S3023 | COX | Bufexamac | 14 | 13 |
| SW219543-1 | S2121 | COX | Licofelone | 14 | 16 |
| SW196404-3 | S1903 | COX | Diclofenac Sodium | 16 | 20 |
| SW219723-1 | S4295 | COX | Meclofenamate Sodium | 19 | 4 |
| SW196982-3 | S3200 | COX | Triflusal | 20 | 26 |
| SW219769-1 | S2903 | COX | Lumiracoxib | 20 | 26 |
| SW196753-3 | S1959 | COX | Tolfenamic Acid | 23 | 26 |
| SW197105-3 | S1626 | COX | Naproxen | 23 | 26 |
| SW219862-1 | S1713 | COX | Piroxicam | 23 | 20 |
| SW196824-3 | S2602 | COX | Acemetacin | 26 | 26 |
| SW197312-3 | S4051 | COX | Nabumetone | 26 | 30 |
| SW219241-1 | S2531 | COX | Asaraldehyde | 26 | 22 |
| SW199611-3 | S1261 | COX | Celecoxib | 27 | 20 |
| SW196448-3 | S2108 | COX | Flunixin Meglumin | 28 | 30 |
| SW196784-3 | S1645 | COX | Ketoprofen | 28 | 2 |
| SW196785-3 | S2040 | COX | Nimesulide | 28 | 11 |
| SW197293-4 | S1646 | COX | Ketorolac | 28 | 30 |
| SW203738-2 | S1638 | COX | Ibuprofen | 28 | 3 |

| | | | | | |
|---|---|---|---|---|---|
| SW219801-1 | S3008 | COX | Zaltoprofen | 28 | 3 |
| SW220125-1 | S2047 | COX | Lornoxicam | 28 | 14 |
| SW197496-2 | S1289 | DNA.RNA.Synthesis | Carmofur | 1 | 7 |
| SW197705-2 | S1192 | DNA.RNA.Synthesis | Raltitrexed | 1 | 7 |
| SW199617-3 | S1209 | DNA.RNA.Synthesis | Fluorouracil (5-Fluoracil; 5-FU) | 1 | 7 |
| SW199090-2 | S1305 | DNA.RNA.Synthesis | Mercaptopurine (6-MP) | 2 | 19 |
| SW197258-4 | S4288 | DNA.RNA.Synthesis | Chloroambucil | 4 | 28 |
| SW219115-1 | S4297 | DNA.RNA.Synthesis | Mupirocin | 5 | 11 |
| SW220050-1 | S2029 | DNA.RNA.Synthesis | Uridine | 6 | 25 |
| SW220241-1 | S1300 | DNA.RNA.Synthesis | FT-207 (NSC 148958) | 7 | 20 |
| SW222225-1 | S1166 | DNA.RNA.Synthesis | Cisplatin | 11 | 10 |
| SW000346-2 | S2554 | DNA.RNA.Synthesis | Daphnetin | 12 | 22 |
| SW218080-2 | S1218 | DNA.RNA.Synthesis | Clofarabine | 12 | 28 |
| SW218086-2 | S1213 | DNA.RNA.Synthesis | Nelarabine | 12 | 7 |
| SW218146-2 | S1229 | DNA.RNA.Synthesis | Fludarabine Phosphate | 12 | 7 |
| SW222226-1 | S1214 | DNA.RNA.Synthesis | Bleomycin Sulfate | 13 | 19 |
| SW197746-4 | S1199 | DNA.RNA.Synthesis | Cladribine | 16 | 28 |
| SW199649-2 | S1714 | DNA.RNA.Synthesis | Gemcitabine | 16 | 28 |
| SW219867-1 | S1221 | DNA.RNA.Synthesis | Dacarbazine | 19 | 30 |
| SW220273-1 | S1299 | DNA.RNA.Synthesis | Floxuridine | 19 | 7 |
| SW219881-1 | S1983 | DNA.RNA.Synthesis | Adenine HCl | 20 | 24 |
| SW197177-4 | S1302 | DNA.RNA.Synthesis | Ifosfamide | 21 | 14 |
| SW219116-1 | S2794 | DNA.RNA.Synthesis | Sofosbuvir (PSI-7977; GS-7977) | 26 | 11 |
| SW219388-1 | S1334 | DNA.RNA.Synthesis | Flupirtine maleate | 26 | 8 |
| SW219151-1 | S1224 | DNA.RNA.Synthesis | Oxaliplatin | 27 | 28 |
| SW220171-1 | S1156 | DNA.RNA.Synthesis | Capecitabine | 30 | 14 |
| SW219272-1 | S1143 | EGFR | AG-490 (Tyrphostin B42) | 1 | 13 |
| SW219447-1 | S1023 | EGFR | Erlotinib HCl (OSI-744) | 2 | 18 |
| SW219475-1 | S7284 | EGFR | CO-1686 (AVL-301) | 2 | 6 |
| SW219315-1 | S1173 | EGFR | WZ4002 | 17 | 3 |
| SW219395-1 | S1179 | EGFR | WZ8040 | 17 | 2 |
| SW219714-1 | S2728 | EGFR | AG-1478 (Tyrphostin AG-1478) | 17 | 27 |
| SW199108-4 | S1025 | EGFR | Gefitinib (ZD1839) | 18 | 4 |
| SW219863-1 | S7297 | EGFR | AZD9291 | 18 | 9 |
| SW219293-1 | S2205 | EGFR | OSI-420 | 19 | 17 |
| SW219394-1 | S1170 | EGFR | WZ3146 | 25 | 15 |
| SW219476-1 | S7206 | EGFR | CNX-2006 | 25 | 9 |
| SW218184-2 | S7039 | EGFR | PD168393 | 27 | 5 |
| SW219523-1 | S8009 | EGFR | AG-18 | 28 | 10 |
| SW219267-1 | S1392 | EGFR | Pelitinib (EKB-569) | 30 | 2 |
| SW219698-1 | S2922 | EGFR | Icotinib | 30 | 17 |
| SW219372-1 | S2818 | HDAC | CI994 (Tacedinaline) | 5 | 28 |
| SW219374-1 | S7292 | HDAC | RG2833 (RGFP109) | 5 | 28 |

| | | | | | |
|---|---|---|---|---|---|
| SW219836-1 | S8001 | HDAC | Rocilinostat (ACY-1215) | 5 | 15 |
| SW219287-2 | S8049 | HDAC | Tubastatin A | 7 | 30 |
| SW219287-1 | S2627 | HDAC | Tubastatin A HCl | 8 | 28 |
| SW219401-1 | S7229 | HDAC | RGFP966 | 8 | 5 |
| SW219449-1 | S7324 | HDAC | TMP269 | 12 | 9 |
| SW219695-1 | S7595 | HDAC | Santacruzamate A (CAY10683) | 12 | 20 |
| SW219169-2 | S1168 | HDAC | Valproic acid sodium salt (Sodium valproate) | 15 | 22 |
| SW220150-1 | S2012 | HDAC | PCI-34051 | 16 | 30 |
| SW219627-1 | S1422 | HDAC | Droxinostat | 18 | 29 |
| SW219738-1 | S1484 | HDAC | MC1568 | 20 | 20 |
| SW199536-4 | S1047 | HDAC | Vorinostat (SAHA; MK0683) | 22 | 15 |
| SW218130-2 | S1122 | HDAC | Mocetinostat (MGCD0103) | 22 | 28 |
| SW218266-2 | S1090 | HDAC | PCI-24781 (Abexinostat) | 22 | 15 |
| SW219369-1 | S1030 | HDAC | Panobinostat (LBH589) | 22 | 15 |
| SW219379-1 | S2779 | HDAC | M344 | 22 | 15 |
| SW219385-1 | S1095 | HDAC | LAQ824 (Dacinostat) | 22 | 15 |
| SW219429-1 | S1515 | HDAC | Pracinostat (SB939) | 22 | 15 |
| SW219445-1 | S1085 | HDAC | Belinostat (PXD101) | 22 | 15 |
| SW219469-1 | S2170 | HDAC | Givinostat (ITF2357) | 22 | 15 |
| SW219664-1 | S1045 | HDAC | Trichostatin A (TSA) | 22 | 15 |
| SW219667-1 | S1053 | HDAC | Entinostat (MS-275) | 22 | 9 |
| SW219675-1 | S2693 | HDAC | Resminostat | 22 | 15 |
| SW219772-1 | S2244 | HDAC | AR-42 | 22 | 15 |
| SW219796-1 | S1096 | HDAC | Quisinostat (JNJ-26481585) | 22 | 15 |
| SW219824-1 | S8043 | HDAC | Scriptaid | 22 | 15 |
| SW219934-1 | S1194 | HDAC | CUDC-101 | 22 | 15 |
| SW220090-1 | S7473 | HDAC | Nexturastat A | 22 | 9 |
| SW220304-1 | S3020 | HDAC | Romidepsin (FK228; Depsipeptide) | 22 | 5 |
| SW219199-1 | S4125 | HDAC | Sodium Phenylbutyrate | 26 | 25 |
| SW219084-1 | S2239 | HDAC | Tubacin | 29 | 29 |
| SW196835-3 | S1847 | Histamine.Receptor | Clemastine Fumarate | 3 | 14 |
| SW197416-3 | S1358 | Histamine.Receptor | Loratadine | 3 | 5 |
| SW196450-4 | S2044 | Histamine.Receptor | Cyproheptadine HCl | 5 | 4 |
| SW196380-2 | S1845 | Histamine.Receptor | Cimetidine | 6 | 26 |
| SW196927-3 | S4139 | Histamine.Receptor | Cyclizine 2HCl | 7 | 20 |
| SW196969-3 | S3176 | Histamine.Receptor | Betahistine 2HCl | 7 | 11 |

| | | | | | |
|---|---|---|---|---|---|
| SW197446-3 | S3146 | Histamine.Receptor | Tripelennamine HCl | 7 | 1 |
| SW199568-2 | S3208 | Histamine.Receptor | Fexofenadine HCl | 7 | 3 |
| SW219552-1 | S4118 | Histamine.Receptor | Histamine 2HCl | 7 | 30 |
| SW197471-3 | S2552 | Histamine.Receptor | Azelastine HCl | 8 | 24 |
| SW219888-1 | S3186 | Histamine.Receptor | Azatadine dimaleate | 12 | 26 |
| SW196707-3 | S1382 | Histamine.Receptor | Mianserin HCl | 14 | 27 |
| SW219889-1 | S3052 | Histamine.Receptor | Rupatadine Fumarate | 14 | 29 |
| SW196598-4 | S1357 | Histamine.Receptor | Lidocaine | 15 | 25 |
| SW196972-3 | S1291 | Histamine.Receptor | Cetirizine DiHCl | 15 | 22 |
| SW219424-1 | S2905 | Histamine.Receptor | JNJ-7777120 | 15 | 22 |
| SW220161-1 | S3037 | Histamine.Receptor | Bepotastine Besilate | 15 | 26 |
| SW197397-2 | S2078 | Histamine.Receptor | Famotidine | 16 | 26 |
| SW197031-3 | S2585 | Histamine.Receptor | Brompheniramine hydrogen maleate | 20 | 26 |
| SW197771-3 | S2494 | Histamine.Receptor | Olopatadine HCl | 20 | 25 |
| SW196372-4 | S1816 | Histamine.Receptor | Chlorpheniramine Maleate | 23 | 1 |
| SW196887-4 | S2024 | Histamine.Receptor | Ketotifen Fumarate | 23 | 11 |
| SW197026-2 | S2308 | Histamine.Receptor | Hesperetin | 23 | 11 |
| SW197646-3 | S1880 | Histamine.Receptor | Roxatidine Acetate HCl | 23 | 12 |
| SW219167-1 | S2813 | Histamine.Receptor | Ciproxifan | 23 | 12 |
| SW196800-3 | S1801 | Histamine.Receptor | Ranitidine | 26 | 13 |
| SW219837-1 | S4131 | Histamine.Receptor | Levodropropizine | 26 | 25 |
| SW196508-3 | S1890 | Histamine.Receptor | Nizatidine | 28 | 18 |
| SW196577-3 | S4026 | Histamine.Receptor | Hydroxyzine 2HCl | 28 | 16 |
| SW197234-3 | S4293 | Histamine.Receptor | Promethazine HCl | 28 | 5 |
| SW197792-3 | S4012 | Histamine.Receptor | Desloratadine | 28 | 3 |
| SW219521-1 | S3163 | Histamine.Receptor | Benztropine mesylate | 28 | 12 |
| SW219706-1 | S2065 | Histamine.Receptor | Lafutidine | 28 | 26 |
| SW219489-1 | S7122 | HSP | XL888 | 9 | 15 |
| SW220214-1 | S8039 | HSP | PU-H71 | 9 | 15 |
| SW219742-1 | S2639 | HSP | SNX-2112 (PF-04928473) | 13 | 15 |
| SW219775-1 | S1052 | HSP | Elesclomol (STA-4783) | 19 | 16 |
| SW218138-2 | S1163 | HSP | AT13387 | 30 | 15 |
| SW219302-1 | S1141 | HSP | 17-AAG (Tanespimycin) | 30 | 15 |
| SW219303-1 | S1142 | HSP | 17-DMAG (Alvespimycin) HCl | 30 | 19 |
| SW219319-1 | S1069 | HSP | AUY922 (NVP-AUY922) | 30 | 15 |
| SW219510-1 | S2713 | HSP | Geldanamycin | 30 | 15 |
| SW219606-1 | S2685 | HSP | KW-2478 | 30 | 15 |
| SW219719-1 | S7340 | HSP | CH5138303 | 30 | 15 |
| SW220092-1 | S7458 | HSP | VER-49009 | 30 | 19 |
| SW220107-1 | S7282 | HSP | NMS-E973 | 30 | 15 |
| SW220153-1 | S2656 | HSP | PF-04929113 (SNX-5422) | 30 | 15 |
| SW220170-1 | S7459 | HSP | VER-50589 | 30 | 15 |

| | | | | | |
|---|---|---|---|---|---|
| SW220175-1 | S7097 | HSP | HSP990 (NVP-HSP990) | 30 | 15 |
| SW220186-1 | S1175 | HSP | BIIB021 | 30 | 15 |
| SW220253-1 | S1159 | HSP | Ganetespib (STA-9090) | 30 | 15 |
| SW219203-1 | S8004 | JAK | ZM 39923 HCl | 7 | 19 |
| SW219757-1 | S7036 | JAK | XL019 | 8 | 6 |
| SW219153-1 | S2796 | JAK | WP1066 | 10 | 27 |
| SW219679-1 | S2219 | JAK | CYT387 | 10 | 19 |
| SW219864-1 | S8057 | JAK | Pacritinib (SB1518) | 10 | 21 |
| SW220119-1 | S1134 | JAK | AT9283 | 11 | 21 |
| SW220020-1 | S7605 | JAK | Filgotinib (GLPG0634) | 15 | 28 |
| SW220096-1 | S2851 | JAK | Baricitinib (LY3009104; INCB028050) | 15 | 28 |
| SW220207-1 | S2902 | JAK | S-Ruxolitinib (INCB018424) | 15 | 8 |
| SW220133-2 | S2789 | JAK | Tofacitinib (CP-690550;Tasocitinib) | 16 | 1 |
| SW218187-2 | S2736 | JAK | TG101348 (SAR302503) | 17 | 6 |
| SW219632-1 | S2692 | JAK | TG101209 | 17 | 2 |
| SW219960-1 | S2179 | JAK | LY2784544 | 17 | 2 |
| SW219437-1 | S2686 | JAK | NVP-BSK805 2HCl | 18 | 2 |
| SW219486-1 | S2162 | JAK | AZD1480 | 19 | 2 |
| SW219623-1 | S2214 | JAK | AZ 960 | 19 | 27 |
| SW220243-1 | S2867 | JAK | WHI-P154 | 19 | 8 |
| SW219454-1 | S2806 | JAK | CEP-33779 | 25 | 5 |
| SW222338-1 | S1378 | JAK | Ruxolitinib (INCB018424) | 27 | 23 |
| SW220133-1 | S5001 | JAK | Tofacitinib (CP-690550) Citrate | 28 | 8 |
| SW202561-3 | S1008 | MEK | Selumetinib (AZD6244) | 9 | 23 |
| SW218101-2 | S1036 | MEK | PD0325901 | 9 | 10 |
| SW219366-1 | S1102 | MEK | U0126-EtOH | 9 | 29 |
| SW219605-1 | S1568 | MEK | PD318088 | 9 | 2 |
| SW219691-1 | S1475 | MEK | Pimasertib (AS-703026) | 9 | 10 |
| SW219692-1 | S2134 | MEK | AZD8330 | 9 | 13 |
| SW219839-1 | S1066 | MEK | SL-327 | 9 | 14 |
| SW219910-1 | S7007 | MEK | MEK162 (ARRY-162; ARRY-438162) | 9 | 29 |
| SW220152-1 | S2617 | MEK | TAK-733 | 9 | 13 |
| SW218089-2 | S2673 | MEK | Trametinib (GSK1120212) | 13 | 10 |
| SW218254-2 | S1177 | MEK | PD98059 | 13 | 10 |
| SW219634-1 | S1531 | MEK | BIX 02189 | 16 | 23 |
| SW197494-3 | S2310 | MEK | Honokiol | 24 | 20 |
| SW219635-1 | S1530 | MEK | BIX 02188 | 28 | 12 |
| SW218136-2 | S1089 | MEK | Refametinib (RDEA119; Bay 86-9766) | 30 | 5 |

| | | | | | |
|---|---|---|---|---|---|
| SW219604-1 | S1020 | MEK | PD184352 (CI-1040) | 30 | 23 |
| SW219216-1 | S7493 | Microtubule.Associated | INH1 | 1 | 24 |
| SW219847-1 | S1148 | Microtubule.Associated | Docetaxel | 10 | 2 |
| SW220268-1 | S1364 | Microtubule.Associated | Epothilone B (EPO906; Patupilone) | 10 | 2 |
| SW219685-1 | S1165 | Microtubule.Associated | ABT-751 (E7010) | 11 | 27 |
| SW220274-1 | S1297 | Microtubule.Associated | Epothilone A | 11 | 2 |
| SW219359-1 | S7336 | Microtubule.Associated | CW069 | 15 | 30 |
| SW219257-1 | S4269 | Microtubule.Associated | Vinorelbine Tartrate | 18 | 2 |
| SW219306-1 | S7494 | Microtubule.Associated | INH6 | 19 | 24 |
| SW102861-5 | S2775 | Microtubule.Associated | Nocodazole | 21 | 27 |
| SW219468-1 | S1241 | Microtubule.Associated | Vincristine | 21 | 2 |
| SW219940-1 | S1248 | Microtubule.Associated | Vinblastine | 21 | 5 |
| SW220198-1 | S2195 | Microtubule.Associated | CYT997 (Lexibulin) | 21 | 5 |
| SW219615-1 | S4071 | Microtubule.Associated | Griseofulvin | 24 | 8 |
| SW219830-1 | S1150 | Microtubule.Associated | Paclitaxel | 29 | 2 |
| SW219073-1 | S1039 | mTOR | Rapamycin (Sirolimus) | 2 | 29 |
| SW219138-1 | S1044 | mTOR | Temsirolimus (CCI-779; NSC 683864) | 2 | 29 |
| SW219218-1 | S1120 | mTOR | Everolimus (RAD001) | 2 | 23 |
| SW219472-1 | S2696 | mTOR | GDC-0980 (RG7422) | 2 | 9 |
| SW220210-1 | S2811 | mTOR | INK 128 (MLN0128) | 2 | 17 |
| SW220211-1 | S2218 | mTOR | PP242 | 2 | 17 |
| SW222224-1 | S1022 | mTOR | Ridaforolimus (Deforolimus; MK-8669) | 2 | 16 |
| SW219676-1 | S2238 | mTOR | Palomid 529 (P529) | 10 | 2 |
| SW219762-1 | S8050 | mTOR | ETP-46464 | 27 | 20 |
| SW220190-1 | S2699 | mTOR | CH5132799 | 27 | 17 |
| SW220246-1 | S2624 | mTOR | OSI-027 | 27 | 23 |
| SW219732-1 | S2406 | mTOR | Chrysophanic Acid | 28 | 12 |
| SW218287-2 | S1555 | mTOR | AZD8055 | 29 | 20 |
| SW219487-1 | S2661 | mTOR | WYE-125132 (WYE-132) | 29 | 12 |
| SW219671-1 | S1266 | mTOR | WYE-354 | 29 | 16 |
| SW219704-1 | S2783 | mTOR | AZD2014 | 29 | 27 |

| | | | | | |
|---|---|---|---|---|---|
| SW219922-1 | S2689 | mTOR | WAY-600 | 29 | 27 |
| SW220188-1 | S1226 | mTOR | KU-0063794 | 29 | 3 |
| SW222245-1 | S7091 | mTOR | Zotarolimus(ABT-578) | 29 | 16 |
| SW219628-1 | S2187 | P450 | Avasimibe | 2 | 4 |
| SW196888-4 | S1353 | P450 | Ketoconazole | 3 | 29 |
| SW197561-4 | S2046 | P450 | Pioglitazone HCl | 4 | 11 |
| SW196866-2 | S2262 | P450 | Apigenin | 6 | 21 |
| SW219553-1 | S2900 | P450 | Cobicistat (GS-9350) | 8 | 29 |
| SW219642-1 | S1195 | P450 | TAK-700 (Orteronel) | 15 | 30 |
| SW101224-2 | S2526 | P450 | Alizarin | 20 | 22 |
| SW197571-2 | S1442 | P450 | Voriconazole | 20 | 22 |
| SW219043-1 | S2555 | P450 | Clarithromycin | 20 | 11 |
| SW219329-1 | S2394 | P450 | Naringenin | 23 | 11 |
| SW219546-1 | S2921 | P450 | PF-4981517 | 24 | 22 |
| SW219229-1 | S2268 | P450 | Baicalein | 27 | 10 |
| SW219192-1 | S4273 | PARP | 3-Aminobenzamide | 7 | 25 |
| SW219820-1 | S1004 | PARP | Veliparib (ABT-888) | 7 | 14 |
| SW218142-2 | S1060 | PARP | Olaparib (AZD2281; Ku-0059436) | 8 | 7 |
| SW219891-2 | S2886 | PARP | PJ34 | 8 | 20 |
| SW219192-2 | S1132 | PARP | INO-1001 | 12 | 12 |
| SW219655-1 | S7048 | PARP | BMN 673 | 12 | 28 |
| SW218112-2 | S1087 | PARP | Iniparib (BSI-201) | 13 | 10 |
| SW219891-1 | S7300 | PARP | PJ34 HCl | 15 | 2 |
| SW219733-1 | S8038 | PARP | UPF 1069 | 20 | 9 |
| SW219936-1 | S2178 | PARP | AG-14361 | 21 | 28 |
| SW219802-1 | S7029 | PARP | AZD2461 | 23 | 7 |
| SW219544-1 | S1098 | PARP | Rucaparib (AG-014699;PF-01367338) | 26 | 7 |
| SW220067-1 | S7438 | PARP | ME0328 | 28 | 20 |
| SW197603-2 | S1512 | PDE | Tadalafil | 1 | 22 |
| SW219717-1 | S1550 | PDE | Pimobendan | 5 | 2 |
| SW219816-1 | S7224 | PDE | Deltarasin | 10 | 20 |
| SW196433-3 | S2320 | PDE | Luteolin | 11 | 8 |
| SW199053-2 | S1294 | PDE | Cilostazol | 15 | 14 |
| SW222234-1 | S2687 | PDE | PF-2545920 | 18 | 28 |
| SW197648-2 | S3172 | PDE | Anagrelide HCl | 20 | 9 |
| SW197737-2 | S2515 | PDE | Vardenafil HCl Trihydrate | 20 | 14 |
| SW199664-3 | S1431 | PDE | Sildenafil Citrate | 20 | 14 |
| SW219217-1 | S4019 | PDE | Avanafil | 20 | 16 |
| SW219244-1 | S2127 | PDE | S- (+)-Rolipram | 20 | 17 |
| SW219280-1 | S1455 | PDE | Cilomilast | 20 | 18 |
| SW197542-3 | S1929 | PDE | Irsogladine | 23 | 3 |
| SW219125-1 | S2312 | PDE | Icariin | 24 | 11 |
| SW196583-4 | S1430 | PDE | Rolipram | 26 | 22 |
| SW196679-3 | S1504 | PDE | Dyphylline | 26 | 8 |
| SW219741-1 | S2620 | PDE | GSK256066 | 26 | 24 |
| SW219856-1 | S8034 | PDE | Apremilast (CC-10004) | 26 | 22 |

| | | | | | |
|---|---|---|---|---|---|
| SW220196-1 | S2131 | PDE | Roflumilast | 26 | 11 |
| SW199286-2 | S1673 | PDE | Aminophylline | 28 | 23 |
| SW219311-1 | S2671 | PI3K | AS-252424 | 1 | 20 |
| SW202556-4 | S1065 | PI3K | GDC-0941 | 2 | 12 |
| SW218117-2 | S1038 | PI3K | PI-103 | 2 | 12 |
| SW219415-1 | S1360 | PI3K | GSK1059615 | 2 | 27 |
| SW219812-1 | S1118 | PI3K | XL147 | 2 | 19 |
| SW220128-1 | S2814 | PI3K | BYL719 | 2 | 6 |
| SW220216-1 | S2767 | PI3K | 3-Methyladenine | 5 | 26 |
| SW219482-1 | S7018 | PI3K | CZC24832 | 7 | 18 |
| SW219650-1 | S8002 | PI3K | GSK2636771 | 8 | 30 |
| SW219158-1 | S2749 | PI3K | BGT226 (NVP-BGT226) | 10 | 27 |
| SW219506-1 | S1205 | PI3K | PIK-75 | 11 | 28 |
| SW113275-2 | S2682 | PI3K | CAY10505 | 12 | 13 |
| SW219297-1 | S1352 | PI3K | TG100-115 | 12 | 1 |
| SW219823-1 | S2226 | PI3K | CAL-101 (Idelalisib; GS-1101) | 12 | 18 |
| SW219919-1 | S2681 | PI3K | AS-604850 | 12 | 10 |
| SW220182-1 | S1072 | PI3K | ZSTK474 | 13 | 4 |
| SW220212-1 | S2227 | PI3K | PIK-294 | 13 | 4 |
| SW217688-2 | S1105 | PI3K | LY294002 | 14 | 30 |
| SW218196-2 | S2636 | PI3K | A66 | 14 | 3 |
| SW218249-2 | S1169 | PI3K | TGX-221 | 14 | 16 |
| SW219525-1 | S2870 | PI3K | TG100713 | 14 | 9 |
| SW219822-1 | S7028 | PI3K | IPI-145 (INK1197) | 14 | 17 |
| SW219502-1 | S2658 | PI3K | GSK2126458 (GSK458) | 17 | 21 |
| SW219871-1 | S2759 | PI3K | CUDC-907 | 17 | 20 |
| SW218129-2 | S1219 | PI3K | YM201636 | 18 | 27 |
| SW219556-1 | S2207 | PI3K | PIK-293 | 24 | 22 |
| SW219873-1 | S7356 | PI3K | HS-173 | 25 | 27 |
| SW219187-1 | S1462 | PI3K | AZD6482 | 27 | 3 |
| SW219245-1 | S1489 | PI3K | PIK-93 | 27 | 5 |
| SW220201-1 | S7016 | PI3K | VS-5584 (SB2343) | 27 | 7 |
| SW218114-2 | S1523 | PI3K | SAR245409 (XL765) | 28 | 20 |
| SW218149-2 | S2247 | PI3K | BKM120 (NVP-BKM120; Buparlisib) | 29 | 27 |
| SW219545-1 | S2758 | PI3K | Wortmannin | 29 | 4 |
| SW219392-1 | S1386 | Proteasome | Nafamostat Mesylate | 12 | 12 |
| SW219683-1 | S7462 | Proteasome | PI-1840 | 12 | 9 |
| SW199665-2 | S3017 | Proteasome | Aspirin | 15 | 26 |
| SW208077-3 | S1013 | Proteasome | Bortezomib (PS-341) | 17 | 19 |
| SW218090-2 | S2853 | Proteasome | Carfilzomib (PR-171) | 17 | 19 |
| SW219161-1 | S1157 | Proteasome | CEP-18770 (Delanzomib) | 17 | 19 |
| SW219743-1 | S2180 | Proteasome | MLN2238 | 17 | 19 |
| SW219744-1 | S2181 | Proteasome | MLN9708 | 17 | 19 |
| SW219780-1 | S2619 | Proteasome | MG-132 | 17 | 19 |
| SW220115-1 | S7172 | Proteasome | ONX-0914 (PR-957) | 17 | 19 |

| | | | | | |
|---|---|---|---|---|---|
| SW220116-1 | S7049 | Proteasome | Oprozomib (ONX 0912) | 17 | 19 |
| SW197284-3 | S2101 | Proteasome | Gabexate Mesylate | 28 | 25 |
| SW219493-1 | S3046 | RAAS | Azilsartan | 1 | 12 |
| SW219848-1 | S1793 | RAAS | Ramipril | 1 | 1 |
| SW220041-1 | S2037 | RAAS | Candesartan Cilexetil | 4 | 4 |
| SW197658-2 | S1894 | RAAS | Valsartan | 6 | 28 |
| SW197676-3 | S1738 | RAAS | Telmisartan | 6 | 26 |
| SW199393-2 | S2581 | RAAS | Quinapril HCl | 7 | 12 |
| SW220093-1 | S2109 | RAAS | Imidapril HCl | 7 | 26 |
| SW222231-1 | S2199 | RAAS | Aliskiren Hemifumarate | 7 | 22 |
| SW197672-2 | S1657 | RAAS | Enalaprilat Dihydrate | 13 | 22 |
| SW220029-1 | S2664 | RAAS | Clinofibrate | 19 | 11 |
| SW199641-2 | S1359 | RAAS | Losartan Potassium (DuP 753) | 20 | 26 |
| SW199650-2 | S1604 | RAAS | Olmesartan Medoxomil | 23 | 11 |
| SW219263-1 | S2079 | RAAS | Moexipril HCl | 23 | 26 |
| SW220127-1 | S7098 | RAAS | PD123319 | 23 | 1 |
| SW197591-3 | S1284 | RAAS | Benazepril HCl | 24 | 22 |
| SW198780-2 | S1941 | RAAS | Enalapril Maleate | 28 | 26 |
| SW199612-2 | S1578 | RAAS | Candesartan | 28 | 11 |
| SW219164-1 | S2051 | RAAS | Captopril | 28 | 11 |
| SW219383-1 | S2081 | RAAS | Cilazapril Monohydrate | 28 | 26 |
| SW219451-1 | S2099 | RAAS | Temocapril HCl | 28 | 18 |
| SW219494-1 | S3057 | RAAS | Azilsartan Medoxomil | 28 | 3 |
| SW212797-2 | S2872 | RAF | GW5074 | 6 | 3 |
| SW218185-2 | S2720 | RAF | ZM 336372 | 6 | 12 |
| SW219204-1 | S1104 | RAF | GDC-0879 | 9 | 18 |
| SW219448-1 | S2746 | RAF | AZ 628 | 9 | 21 |
| SW219503-1 | S2807 | RAF | Dabrafenib (GSK2118436) | 9 | 21 |
| SW219895-1 | S7291 | RAF | TAK-632 | 9 | 7 |
| SW220064-1 | S7108 | RAF | LGX818 | 9 | 4 |
| SW202562-3 | S1040 | RAF | Sorafenib Tosylate | 10 | 7 |
| SW202562-4 | S7397 | RAF | Sorafenib | 10 | 29 |
| SW219923-1 | S2161 | RAF | RAF265 (CHIR-265) | 11 | 16 |
| SW218095-2 | S1267 | RAF | Vemurafenib (PLX4032; RG7204) | 27 | 4 |
| SW218119-2 | S1152 | RAF | PLX-4720 | 29 | 24 |
| SW220229-1 | S2220 | RAF | SB590885 | 30 | 9 |
| SW220279-1 | S1398 | Reverse.Transcriptase | Stavudine (d4T) | 1 | 12 |
| SW220193-1 | S2914 | Reverse.Transcriptase | Dapivirine (TMC120) | 2 | 2 |
| SW198799-2 | S2579 | Reverse.Transcriptase | Zidovudine | 7 | 13 |

| | | | | | |
|---|---|---|---|---|---|
| SW220232-1 | S7303 | Reverse.Transcriptase | Rilpivirine | 10 | 9 |
| SW219933-1 | S1718 | Reverse.Transcriptase | Adefovir Dipivoxil | 13 | 19 |
| SW220151-1 | S1400 | Reverse.Transcriptase | Tenofovir Disoproxil Fumarate | 16 | 7 |
| SW197364-4 | S1719 | Reverse.Transcriptase | Zalcitabine | 23 | 26 |
| SW197614-3 | S1706 | Reverse.Transcriptase | Lamivudine | 23 | 26 |
| SW220172-1 | S1704 | Reverse.Transcriptase | Emtricitabine | 23 | 14 |
| SW219570-1 | S3080 | Reverse.Transcriptase | Etravirine (TMC125) | 27 | 7 |
| SW197569-2 | S1742 | Reverse.Transcriptase | Nevirapine | 28 | 26 |
| SW198619-2 | S1702 | Reverse.Transcriptase | Didanosine | 28 | 8 |
| SW219101-1 | S4016 | Sodium.Channel | Ouabain | 2 | 21 |
| SW219113-1 | S4290 | Sodium.Channel | Digoxin | 2 | 21 |
| SW196688-3 | S4080 | Sodium.Channel | Triamterene | 7 | 20 |
| SW220143-1 | S2524 | Sodium.Channel | Phenytoin sodium | 7 | 11 |
| SW196719-3 | S4023 | Sodium.Channel | Procaine HCl | 12 | 30 |
| SW219770-1 | S1256 | Sodium.Channel | Rufinamide | 15 | 11 |
| SW220114-1 | S2118 | Sodium.Channel | Ibutilide Fumarate | 15 | 14 |
| SW196333-5 | S2560 | Sodium.Channel | Amiloride hydrochloride dihydrate | 20 | 11 |
| SW197468-3 | S1391 | Sodium.Channel | Oxcarbazepine | 20 | 14 |
| SW203757-2 | S2525 | Sodium.Channel | Phenytoin | 20 | 1 |
| SW197486-3 | S3024 | Sodium.Channel | Lamotrigine | 21 | 30 |
| SW196805-4 | S1614 | Sodium.Channel | Riluzole | 23 | 9 |
| SW196878-3 | S4294 | Sodium.Channel | Procainamide HCl | 23 | 23 |
| SW197338-3 | S1828 | Sodium.Channel | Proparacaine HCl | 23 | 22 |
| SW220141-1 | S1693 | Sodium.Channel | Carbamazepine | 23 | 11 |
| SW196964-3 | S2500 | Sodium.Channel | Propafenone HCl | 24 | 24 |
| SW198832-2 | S3064 | Sodium.Channel | Ambroxol HCl | 24 | 14 |
| SW196475-3 | S4038 | Sodium.Channel | Dibucaine HCl | 28 | 12 |
| SW220277-1 | S2785 | Sodium.Channel | A-803467 | 28 | 4 |
| SW219924-1 | S7518 | Topoisomerase | Voreloxin (SNS-595) | 4 | 19 |
| SW203763-2 | S1342 | Topoisomerase | Genistein | 6 | 28 |
| SW196414-3 | S1288 | Topoisomerase | Camptothecin | 10 | 19 |
| SW196745-6 | S1889 | Topoisomerase | Mitoxantrone | 10 | 9 |
| SW219079-1 | S1393 | Topoisomerase | Pirarubicin | 10 | 19 |
| SW219806-1 | S7261 | Topoisomerase | Beta-Lapachone | 10 | 27 |
| SW219421-1 | S1228 | Topoisomerase | Idarubicin HCl | 11 | 4 |
| SW219441-1 | S1208 | Topoisomerase | Doxorubicin (Adriamycin) | 11 | 20 |
| SW219442-1 | S1223 | Topoisomerase | Epirubicin HCl | 11 | 9 |
| SW197554-3 | S1465 | Topoisomerase | Moxifloxacin HCl | 13 | 26 |
| SW219048-1 | S1225 | Topoisomerase | Etoposide | 13 | 19 |
| SW220163-1 | S1198 | Topoisomerase | Irinotecan | 13 | 19 |

| | | | | | |
|---|---|---|---|---|---|
| SW196774-3 | S3181 | Topoisomerase | Flumequine | 15 | 11 |
| SW197608-4 | S4119 | Topoisomerase | Pefloxacin Mesylate Dihydrate | 16 | 26 |
| SW197790-4 | S2217 | Topoisomerase | Irinotecan HCl Trihydrate | 16 | 28 |
| SW219056-1 | S3603 | Topoisomerase | Betulinic acid | 18 | 30 |
| SW197557-5 | S1231 | Topoisomerase | Topotecan HCl | 19 | 19 |
| SW219948-1 | S4908 | Topoisomerase | SN-38 | 19 | 28 |
| SW220315-1 | S2423 | Topoisomerase | 10-Hydroxycamptothecin | 19 | 19 |
| SW219946-1 | S1367 | Topoisomerase | Amonafide | 27 | 19 |
| SW219896-1 | S1084 | VEGFR | Brivanib (BMS-540215) | 1 | 17 |
| SW218092-2 | S1046 | VEGFR | Vandetanib (ZD6474) | 3 | 2 |
| SW218116-2 | S1003 | VEGFR | Linifanib (ABT-869) | 10 | 19 |
| SW220084-1 | S7258 | VEGFR | SKLB1002 | 12 | 20 |
| SW219259-1 | S1164 | VEGFR | Lenvatinib (E7080) | 13 | 23 |
| SW219897-1 | S1138 | VEGFR | Brivanib Alaninate (BMS-582664) | 13 | 1 |
| SW198937-2 | S1101 | VEGFR | Vatalanib (PTK787) 2HCl | 15 | 29 |
| SW218301-2 | S1010 | VEGFR | Nintedanib (BIBF 1120) | 18 | 8 |
| SW219500-1 | S2896 | VEGFR | ZM 323881 HCl | 19 | 4 |
| SW219791-1 | S2845 | VEGFR | Semaxanib (SU5416) | 19 | 8 |
| SW218300-2 | S1032 | VEGFR | Motesanib Diphosphate (AMG-706) | 22 | 8 |
| SW220296-1 | S2221 | VEGFR | Apatinib | 24 | 4 |
| SW219944-1 | S2842 | VEGFR | SAR131675 | 26 | 17 |
| SW219943-1 | S2897 | VEGFR | ZM 306416 | 27 | 17 |
| SW219464-1 | S1005 | VEGFR.KIT.PDGFR | Axitinib | 1 | 21 |
| SW218156-2 | S1363 | VEGFR.KIT.PDGFR | Ki8751 | 5 | 21 |
| SW218097-2 | S1178 | VEGFR.KIT.PDGFR | Regorafenib (BAY 73-4506) | 11 | 29 |
| SW218082-2 | S1035 | VEGFR.KIT.PDGFR | Pazopanib HCl (GW786034 HCl) | 13 | 28 |
| SW219261-1 | S1017 | VEGFR.KIT.PDGFR | Cediranib (AZD2171) | 18 | 18 |
| SW219787-1 | S1018 | VEGFR.KIT.PDGFR | Dovitinib (TKI-258; CHIR-258) | 18 | 6 |
| SW218082-3 | S3012 | VEGFR.KIT.PDGFR | Pazopanib | 19 | 28 |
| SW219794-1 | S1220 | VEGFR.KIT.PDGFR | OSI-930 | 19 | 4 |
| SW219407-1 | S1042 | VEGFR.KIT.PDGFR | Sunitinib Malate | 25 | 2 |
| SW219787-2 | S2769 | VEGFR.KIT.PDGFR | Dovitinib (TKI-258) Dilactic Acid | 25 | 21 |
| SW219262-1 | S1557 | VEGFR.KIT.PDGFR | KRN 633 | 28 | 8 |
| SW220176-1 | S2231 | VEGFR.KIT.PDGFR | Telatinib | 29 | 5 |
| SW219364-1 | S1207 | VEGFR.KIT.PDGFR | Tivozanib (AV-951) | 30 | 17 |

**Supplementary Table 3: SNF-Euclidean APC hypergeometric test p-values.**
Hypergeometric test p-values for Selleck target classes in SNF-Euclidean APC clusters. Bonferroni-corrected alpha = 0.0006

| Class | Cluster Number | -log10 p-value | Significant? |
|---|---|---|---|
| P450 | 1 | 4.25605576 | TRUE |
| ATM.ATR | 1 | 5.04517835 | TRUE |
| Hist.Receptor | 4 | 2.2897748 | FALSE |
| COX | 6 | 1.91982167 | FALSE |
| RAAS | 6 | 2.13377808 | FALSE |
| Hist.Receptor | 6 | 3.43010914 | TRUE |
| AdrR.agonist | 6 | 4.02855491 | TRUE |
| Hist.Receptor | 9 | 1.95941317 | FALSE |
| Hist.Receptor | 10 | 1.75917031 | FALSE |
| RAAS | 10 | 3.45658465 | TRUE |
| Reverse.Transcriptase | 10 | 4.49051096 | TRUE |
| Hist.Receptor | 13 | 2.69927303 | FALSE |
| AChR.antagonist | 13 | 3.0056789 | FALSE |
| PDE | 14 | 2.68768419 | FALSE |
| K.Channel | 14 | 3.62607173 | TRUE |
| Adr.Receptor.antagonist | 15 | 2.5148078 | FALSE |
| PI3K | 17 | 3.14300227 | FALSE |
| COX | 19 | 3.3097259 | TRUE |
| HDAC | 20 | 1.79545626 | FALSE |
| K.Channel | 20 | 3.29865781 | TRUE |
| DNA.RNA.Synthesis | 25 | 6.20147175 | TRUE |
| AChR.antagonist | 26 | 2.84683694 | FALSE |
| RAAS | 26 | 3.21292655 | FALSE |
| COX | 26 | 4.02855491 | TRUE |
| PI3K | 27 | 3.05651353 | FALSE |
| JAK | 39 | 3.71566364 | TRUE |
| CDK | 42 | 8.86025664 | TRUE |
| PDE | 43 | 3.48380466 | TRUE |
| DNA.RNA.Synthesis | 45 | 3.1743779 | FALSE |
| Aurora.Kinase | 45 | 3.63343507 | TRUE |
| Reverse.Transcriptase | 45 | 4.11305913 | TRUE |
| VEGFR.KIT.PDGFR | 45 | 5.79439169 | TRUE |
| MEK | 46 | 4.4173404 | TRUE |
| RAF | 48 | 5.61784122 | TRUE |
| HSP | 48 | 15.6575773 | TRUE |

| | | | |
|---|---|---|---|
| PLK | 49 | 4.77132139 | TRUE |
| HDAC | 49 | 15.6575773 | TRUE |
| Hist.Receptor | 64 | 2.05864879 | FALSE |
| AChR.antagonist | 64 | 2.35315313 | FALSE |
| Adr.Receptor.agonist | 64 | 2.40247918 | FALSE |
| COX | 64 | 2.40247918 | FALSE |
| Dopamine.Receptor.antagonist | 65 | 4.92012054 | TRUE |
| Topoisomerase | 66 | 3.41531571 | TRUE |
| PI3K | 67 | 2.13035048 | FALSE |
| Topoisomerase | 67 | 2.7625374 | FALSE |
| Bcl-2 | 67 | 4.0269807 | TRUE |
| Hist.Receptor | 68 | 1.95941317 | FALSE |
| COX | 68 | 2.30034797 | FALSE |
| VEGFR | 71 | 4.97183704 | TRUE |
| HMG-CoA.Reductase | 73 | 5.67048662 | TRUE |
| Src | 74 | 6.64992842 | TRUE |
| Epigenetic.Reader.Domain | 76 | 14.3111371 | TRUE |
| Hist.Receptor | 77 | 2.13035048 | FALSE |
| Adr.Receptor.antagonist | 77 | 2.4761153 | FALSE |
| AChR.antagonist | 80 | 2.88696492 | FALSE |
| Hist.Receptor | 81 | 2.42654657 | FALSE |
| ER.PR_antagonist | 81 | 4.78604926 | TRUE |
| RAF | 82 | 5.49803799 | TRUE |
| MEK | 82 | 7.51565297 | TRUE |
| HDAC | 83 | 9.41983968 | TRUE |
| AChR.antagonist | 86 | 2.21944195 | FALSE |
| Adr.Receptor.antagonist | 86 | 2.26828361 | FALSE |
| 5HTR.agonist | 86 | 4.58709819 | TRUE |
| p38.MAPK | 87 | 4.82954492 | TRUE |
| STAT | 88 | 4.17883559 | TRUE |
| ABL | 88 | 4.8251888 | TRUE |
| Topoisomerase | 88 | 5.04851238 | TRUE |
| RAF | 90 | 4.14395889 | TRUE |
| ATPase | 90 | 6.73816904 | TRUE |
| ABL | 91 | 4.37914804 | TRUE |
| Adr.Receptor.antagonist | 93 | 3.50631673 | TRUE |
| ER.PR_agonist | 97 | 4.67738131 | TRUE |
| EGFR.ERBB | 98 | 4.44183346 | TRUE |
| DNA.RNA.Synthesis | 99 | 3.4727626 | TRUE |
| PI3K | 100 | 3.20179911 | FALSE |
| Wnt.beta-catenin | 100 | 3.70337698 | TRUE |

| | | | |
|---|---|---|---|
| IGFR | 100 | 3.85458471 | TRUE |
| Ca.Channel | 100 | 4.37616146 | TRUE |
| mTOR | 100 | 14.7784985 | TRUE |
| Microtubule.Associated | 101 | 7.23326999 | TRUE |
| MEK | 102 | 9.30487308 | TRUE |
| Topoisomerase | 103 | 2.97361122 | FALSE |
| Sodium.Channel | 103 | 3.04111994 | FALSE |
| IkB.IKK | 103 | 4.24542125 | TRUE |
| JAK | 103 | 4.3690592 | TRUE |
| CDK | 103 | 5.68016436 | TRUE |
| Proteasome | 103 | 15.9545898 | TRUE |
| MET | 105 | 4.9666167 | TRUE |
| PDGFR | 105 | 5.29514851 | TRUE |
| PARP | 106 | 4.70583723 | TRUE |
| PDE | 107 | 2.84249965 | FALSE |
| Histone.Methyltransferase | 108 | 4.59476586 | TRUE |
| Aurora.Kinase | 109 | 5.95513653 | TRUE |
| p38.MAPK | 110 | 5.06645713 | TRUE |
| JAK | 111 | 3.48380466 | TRUE |
| EGFR | 111 | 3.87285091 | TRUE |
| MET.VEGFR | 111 | 5.21210644 | TRUE |
| ABL | 111 | 6.22196478 | TRUE |
| DNA.RNA.Synthesis | 115 | 4.31493643 | TRUE |
| RAAS | 117 | 2.86452526 | FALSE |
| Sodium.Channel | 117 | 2.99566445 | FALSE |
| DNA.RNA.Synthesis | 117 | 3.9787859 | TRUE |
| Kinesin | 118 | 6.76986956 | TRUE |
| Topoisomerase | 119 | 5.95198865 | TRUE |
| PLK | 122 | 4.83449691 | TRUE |
| STAT | 122 | 5.0753188 | TRUE |
| Microtubule.Associated | 122 | 9.48944986 | TRUE |
| EGFR | 123 | 4.39394247 | TRUE |
| PI3K | 123 | 4.88317226 | TRUE |
| mTOR | 123 | 10.0294242 | TRUE |
| RAAS | 124 | 2.77899116 | FALSE |
| 5HTR.antagonist | 124 | 2.84249965 | FALSE |
| 5HTR.antagonist | 127 | 2.48838075 | FALSE |
| Adr.Receptor.agonist | 127 | 3.3097259 | TRUE |
| GluR.antagonist | 127 | 5.7801431 | TRUE |

**Supplementary Table 4: Target class enrichment in APC clusters.** APC clusters that are enriched with a single class from the Selleck library and the NPFs that are associated with those clusters

| Class | Cluster number | Number of NPFs | NPF IDs |
|---|---|---|---|
| HSP | 48 | 0 | |
| HDAC | 49 | 3 | SW218953 through SW218955 |
| Epigenetic.Reader.Domain | 76 | 0 | |
| mTOR | 100 | 1 | SW218859 |
| Proteasome | 103 | 1 | SW218864 |
| mTOR | 123 | 0 | |

**Supplementary Table 5: Cluster #49 members.** NPFs and Selleck compounds that are present in the APC cluster #49, which is enriched in HDAC inhibitors.

| Name | Class | Compound | MW |
|---|---|---|---|
| SW218140-2 | PLK | Volasertib (BI 6727) | 618.81 |
| SW218130-2 | HDAC | Mocetinostat (MGCD0103) | 396.44 |
| SW199536-4 | HDAC | Vorinostat (SAHA, MK0683) | 264.3 |
| SW218954-1 | NPF | RLUS-2173D | NA |
| SW218953-1 | NPF | RLUS-2173C | NA |
| SW218266-2 | HDAC | PCI-24781 (Abexinostat) | 397.42 |
| SW218187-2 | JAK | TG101348 (SAR302503) | 524.68 |
| SW219379-1 | HDAC | M344 | 307.39 |
| SW219369-1 | HDAC | Panobinostat (LBH589) | 349.43 |
| SW219316-1 | PLK | Ro3280 | 543.61 |
| SW218955-1 | NPF | RLUS-2173E | NA |
| SW219469-1 | HDAC | Givinostat (ITF2357) | 475.97 |
| SW219445-1 | HDAC | Belinostat (PXD101) | 318.35 |
| SW219429-1 | HDAC | Pracinostat (SB939) | 358.48 |
| SW219385-1 | HDAC | LAQ824 (Dacinostat) | 379.46 |
| SW219796-1 | HDAC | Quisinostat (JNJ-26481585) | 394.48 |
| SW219772-1 | HDAC | AR-42 | 312.36 |
| SW219675-1 | HDAC | Resminostat | 349.4 |
| SW219664-1 | HDAC | Trichostatin A (TSA) | 302.4 |
| SW220090-1 | HDAC | Nexturastat A | 341.4 |
| SW219934-1 | HDAC | CUDC-101 | 434.49 |
| SW219824-1 | HDAC | Scriptaid | 326.35 |
